# Mapping long-distance jump dispersal risk by leveraging transport infrastructure data and sampling bias correction: a case study with the invasive spotted lanternfly (*Lycorma delicatula*)

**DOI:** 10.64898/2026.10.29.762583

**Authors:** Joseph Keller, Nadège Belouard, Sebastiano De Bona, Jocelyn E. Behm, Matthew R. Helmus

## Abstract

Long-distance jump dispersal accelerates invasive species spread. Analysis of past jumps can identify invasion pathways and point to locations at risk of future colonization, but spatial bias in survey records obscures patterns. To inform surveillance for the spreading invasive grape pest *Lycorma delicatula* (Hemiptera: Fulgoridae), we gathered 82,890 agency records of established invasive populations from the United States. We used a clustering algorithm to identify 207 jump dispersal sites, and we developed a bias-aware comparison dataset by filtering records to identify proactive search effort at locations not yet known to hold *L. delicatula*. We measured points’ proximity to a co-produced list of 22 property types encompassing categories with high vehicle traffic, railroads, and related contexts that may transport *L. delicatula*. Bootstrapping analysis showed that proactive surveys were concentrated near hypothesized influential property types, confirming substantial sampling bias. With this bias accounted for, historical jump dispersal sites were closer than expected to 15 property types, particularly distribution centers, truck rentals, flea markets, and importers. We also compared the lengths of road and rail around these points, finding that jump sites were strongly associated with elevated rail density, whereas models showed limited support for roads as jump sites. Finally, we used the high-risk features to project risk of future jumps across the eastern U.S. using ensemble random forests. Predicted high-risk areas were northeast Illinois, southeast Michigan, and central Ohio. This framework is generalizable, allowing various pathways to be assessed and jump dispersal risk to be projected based on spatially biased surveillance data.

## 1. Introduction

Managers working to eradicate or slow the spread of harmful invasive species are challenged by jump dispersal events, extra-range dispersal events distinct from diffusive spread (See Table 1 for definitions of terminology used, Wilson et al., 2009, Haubrock et al., 2025). These events are marked by exceptional long movement distances, relative rarity, and non-standard mechanisms of dispersal, including transport by humans (Eyer et al., 2021; Suarez et al., 2001). Jump dispersal generates small, difficult-to-detect satellite populations (Wilson et al., 2009) that create spatial discontinuities in the invaded range, fragment management into isolated foci, and compound surveillance and control effort (Clark et al., 2003; Melbourne & Hastings, 2009). Early detection and rapid response to manage jump events can often provide numerous benefits (Burgos-Rodríguez & Burgiel, 2020; Greenlees et al., 2018; Simberloff, 2001) including local eradication (Brockerhoff et al., 2010). Thus, prediction of jump dispersal events is key, necessitating approaches that identify areas of elevated risk (Saffer et al., 2025).

**Table 1:** Terminology used.

| Term | Definition |
| --- | --- |
| Jump dispersal | Dispersal beyond the contiguous range margin that establishes new, spatially disjunct (satellite) populations, as opposed to the incremental advance of the invasion front. |
| Invasion pathway | The process by which a non-native species is transported and introduced beyond its current range. |
| Invasion hub | A location with elevated risk of receiving jump-dispersing individuals. |
| Transport infrastructure | A physical network over which people and goods move (i.e. roads, railways). |

Examining locations of satellite populations that resulted from previous jump dispersal events can reveal potential jump dispersal pathways. Invasion pathways are the processes that transport propagules to new areas, and the consequences of those pathways are concentrated at invasion hubs: locations with elevated risk, relative to the surrounding landscape, of receiving jump-dispersing individuals (Table 1). For instance, the emerald ash borer’s (*Agrilus planipennis*) colonization in Michigan implicated firewood transport as a jump dispersal pathway and campgrounds as hubs (Muirhead et al., 2006). Once invasion hubs have been identified, they can become focal points for surveillance and management to reduce or halt jump events. For example, when analyses indicated a link between emerald ash borer jump dispersal and camp sites, additional quarantine measures to reduce the movement of firewood were recommended to reduce colonization at these hubs (Muirhead et al., 2006).

To identify associations between potential pathways and jump dispersal events, two types of spatial data are needed: species occurrence data and landscape feature data. Species occurrence datasets are increasingly available (Bois et al., 2011; De Bona et al., 2023; Saul et al., 2017; Van Rees et al., 2022; Wallace & Bargeron, 2014), and methods are available to identify jump dispersal events based on these data (Belouard et al., 2025; Goldstein et al., 2019; Shigesada et al., 1995). Linear data describing road and rail locations can characterize the influence of transport infrastructure, (Benedetti & Morelli, 2017), while point-of-interest (POI) datasets map the locations of businesses, infrastructure, and other activity centers that may facilitate human-mediated transport (Psyllidis et al., 2022; Sun et al., 2023). These data underpin modern location-based services and digital mapping platforms and have been widely applied in fields such as epidemiology, urban analytics, and conservation planning (Du & Mao, 2024; Ferro-Diez et al., 2020; Huang et al., 2022; Jiao et al., 2023). However, point of interest data have not been explicitly integrated into methodological frameworks for estimating jump dispersal risk.

Identifying links between transport infrastructure, property types, and jump dispersal sites can be difficult because pest surveillance effort is often non-randomly distributed with respect to human-built features (Dimson et al., 2023). Species occurrence data are often collected opportunistically, resulting in spatial sampling bias that obscures true associations between pest occurrence and landscape features (De Groot et al., 2022). In practice, both public reporting and agency-led surveys are often concentrated in accessible or high-priority locations, including vulnerable areas (Dennis & Thomas, 2000), roadsides (Daru et al., 2018), and near other transport infrastructure (Penhacek et al., 2025). Consequently, analyses that do not account for this bias may confound elevated surveillance effort near certain property types with true ecological processes or important invasion pathways (Phillips et al., 2009). While methods incorporating both species presences and absences can reduce bias provided that survey bias equally affects both (Zadrozny, 2004), explicitly characterizing sampling bias remains necessary to define the scope of inference and avoid spurious associations between landscape features and jump dispersal risk that could misdirect management actions (Baumbusch et al., 2025).

The spotted lanternfly (*Lycorma delicatula*, Hemiptera: Fulgoridae) provides a well-documented system for studying long-distance jump dispersal. Since its introduction to the eastern United States, this grape pest’s rapid spread has been characterized by repeated establishment of outlying satellite populations (Belouard et al., 2025; Elsensohn et al., 2024; Keller et al., 2025; Urban et al., 2021). Eggs, nymphs, and adults can be moved on a wide range of materials and vehicles (Elsensohn et al., 2024; Liu, 2019). Consistent with these mechanisms, prior studies have linked *L. delicatula* spread to transportation networks and human activity, including associations with rail infrastructure, population density, and interstate trade (Cook et al., 2021; Huron et al., 2022; Jones et al., 2021; Ladin et al., 2023). Despite extensive survey efforts and detailed occurrence data compiled by state and federal agencies (De Bona et al., 2023), the role of specific features of the human-built landscape in facilitating jump dispersal remains untested. This system therefore provides an opportunity to evaluate which features contribute to the spatial patterning of jump dispersal events by invasive species.

We developed a framework to identify invasion hubs and map jump dispersal risk by integrating POI data with bias-corrected occurrence data. We quantified associations between *L. delicatula* jump dispersal events and features of the human-built landscape while explicitly accounting for spatial sampling bias in survey effort. We addressed three questions: (1) Do proactive state and federal survey locations exhibit spatial sampling bias toward property types and transport infrastructure hypothesized to facilitate *L. delicatula* dispersal? (2) Which property types and transport infrastructure types are associated with jump dispersal events after accounting for any sampling bias? (3) Can bias-corrected pathway information accurately predict the most recent dispersal jumps? While our results focus on *L. delicatula*, the framework is broadly applicable to other invasive species with sufficient occurrence data that spread by jump dispersal.

## 2. Methods

### 2.1 State and federal survey data and POI/transport infrastructure mapping datasets

We compiled spatial occurrence data and defined a study region to encompass all established *Lycorma delicatula* populations confirmed by federal and state agency surveys in the United States by December 2024. The study region included the District of Columbia and 18 northeastern states: Massachusetts, Rhode Island, Connecticut, New York, New Jersey, Pennsylvania, Delaware, Maryland, Virginia, North Carolina, South Carolina, Tennessee, Kentucky, West Virginia, Ohio, Michigan, Illinois, and Indiana. All species occurrence, POI, and transport infrastructure datasets were clipped to this extent. Occurrence data were derived from the *lydemapr* database, which compiles *L. delicatula* records from its first detection in 2014 through 2025 (De Bona et al., 2023). While the publicly available *lydemapr* data are aggregated to 1 km resolution, we conducted analyses using the original survey records with unrounded coordinates, accessed under data-sharing agreements. To reduce potential false positives, we excluded records from public reporting and community science initiatives and restricted analyses to confirmed reports of established populations recorded by state and federal agency surveyors (n = 82,890). Records were assigned to biological years based on the species’ univoltine life cycle, with turnover at the end of April. Analyses of POI and transport infrastructure associations with jump dispersal were conducted using *L. delicatula* occurrence data from 2014–2023, with 2024-2025 data withheld to evaluate the accuracy of derived risk metrics.

The Dataplor POI database contains more than 18 million georeferenced records from across the U.S. and is regularly updated to reflect changes in business status (Dataplor, 2022). POIs are assigned to categorical labels using a proprietary machine-learning classification. In consultation with U.S. federal and state agency operations managers, we identified 21 POI categories where accidental human-mediated transport of *L. delicatula* individuals or egg masses is likely. These categories were amusement parks, auction centers, campgrounds, casinos, colleges, fairgrounds, farmers’ markets, flea markets, importers, junkyards, landfills, landscaping businesses, racetracks, railroad stations, shipping and distribution centers, stadiums, summer camps, timber facilities, truck rentals, truck stops, and wineries, including vineyards. Dataplor records were filtered to retain only POIs assigned to these property categories, and additional manual cleaning was conducted to remove mislabeled, inactive, or otherwise irrelevant locations (Figures S1.1–S1.22).

Ports were included as an additional class of potential influential points due to their role in concentrating trade and movement. Ports were not consistently represented within the Dataplor classifications. We therefore compiled port locations from the Gulf Coast Prairie Landscape Conservation Cooperative (LCC Network Data Steward & Gulf Coast Prairie LCC Data Manager, 2014) and World Port Index (National Geospatial-Intelligence Agency, 2020), capturing ports of varying sizes along coastlines and inland waterways.

In addition to the 22 total property types, we evaluated linear transport infrastructure features hypothesized to facilitate *L. delicatula* jump dispersal across the landscape. Rail lines, primary roads, and secondary roads were included as potential continuous dispersal pathways. Spatial data were obtained from TIGER/Line 2023 datasets provided by the U.S. Census Bureau (2023).

### 2.2 Identifying jump-dispersal events from surveillance records

We used the DBSCAN clustering algorithm to identify locations where *L. delicatula* jump dispersal events have occurred in the invaded range in the eastern United States (i.e., Density-Based Spatial Clustering of Applications with Noise, Ester et al., 1996). This approach provides an alternative to prior methods while achieving similar results (Belouard et al., 2025). The DBSCAN algorithm groups points based on density using a search radius and minimum number of points, allowing identification of irregularly shaped clusters without specifying the number of clusters *a priori*, as would be needed for k-means clustering (Kanagala & Krishnaiah, 2016). Instead, groups of points are clustered based on density, and irregular shaped point clusters including rings and u-shapes are readily picked out (Ester et al., 1996). Although DBSCAN can filter low-density observations as noise (Santosuosso & Papini, 2018), we set the minimum cluster size to one point to retain isolated established populations because these single records represent candidate jump-dispersal events.

To distinguish long-distance jump dispersal from local diffusion and to resolve distinct jump events occurring in close proximity, we applied a two-step clustering approach. Using the R package *dbscan* (Hahsler et al., 2019), we carried out two sequential iterations of DBSCAN to spatially cluster *L. delicatula* establishment records (See Figure S1.23 for a schematic). For each year, we first clustered records of established *L. delicatula* from that year and all prior years using a 20 km search radius. Including prior years’ records accounted for lack of evidence of local eradication even when intense management has occurred (Keller et al., 2023; Urban & Leach, 2023). This coarse-scale clustering separated candidate jump points from the existing invasion front, with the distance parameter based on observed annual spread distances and rates (Belouard et al., 2025; Cook et al., 2021; Keller et al., 2025). Jump dispersal was defined by the presence of spatial gaps such that detections >20 km from prior years’ records were not classified as jump events if contemporaneous records formed a continuous chain connecting them to existing populations, leaving no gap greater than 20 km. We then applied a second clustering step to the isolated jump points using a 5 km search radius to distinguish nearby but independent jump events. This finer-scale step reflects the limited spatial extent of newly *L. delicatula* satellite populations and prevents merging nearby but independent events. Finally, we conducted a sensitivity analysis to evaluate how the number of clusters varied with the biologically informed distance parameters (20 km and 5 km), testing values ranging from 10–50 km for the first clustering step and 2.5– 30 km for the second.

### 2.3 Bias-corrected single feature analysis

The agency-derived *L. delicatula* survey data we used contained records gathered in pursuit of multiple objectives including proactively searching for new colonization events, delimiting newly detected populations, and monitoring known infestations. Observations focused on existing populations and/or made in response to the discovery of an outlying population in part reflect *L. delicatula*’s jump dispersal and colonization processes and are therefore unsuitable for characterizing sampling bias uninfluenced by actual jump patterns. Only surveys predating any nearby detection capture proactive search effort, where location choice is independent of local establishment. Proactive search points were therefore defined as absence records collected by government surveyors at least 20 km from any previously verified established population. Prior presence records were defined at daily temporal resolution to exclude follow-up survey effort after detection.

To quantify associations between jump-dispersal sites and property types while accounting for search bias, we compared spatial proximity to 22 POI types among three groups: (1) 130 jump dispersal sites from prior to 2024, (2) 121,904 proactive search locations from the same period, and (3) 5,000 randomly sampled background points across the study region. For each location, we calculated the distance to the nearest POI point of each type using the sf package in R (Pebesma, 2018). For jump events represented by multiple records clustered together, distances were averaged to obtain a single value per event. Distances were resampled with replacement over 50,000 bootstrap iterations to estimate group means. Differences between proactive search (group 2) and regional background (group 3) means quantified sampling bias, while differences between jump dispersal (group 1) and proactive search (group 2) means measured property-type associations independent of bias. We assessed whether jump locations were closer than expected to each POI type by calculating a one-sided empirical p-value, derived by evaluating the proportion of iterations where the jump site mean distance was greater than the proactive search site mean distance (first comparison) or where the proactive search site mean distance was greater than the regional background mean distance (second comparison). One-sided tests were used because we prioritized identifying positive associations between POI types and jump dispersal sites. We controlled the false discovery rate (FDR) across the 22 variables using the Benjamini-Hochberg procedure (Benjamini & Hochberg, 1995), with variables considered significant at FDR-adjusted q < 0.05. Finally, we quantified the effect size of each property type using Cliff’s Δ, a non-parametric metric ranging from −1 to 1 (Cliff, 1993).

We used the same approach to assess associations between jump dispersal sites and transport infrastructure (rail, primary roads, secondary roads). We measured infrastructure density as total length within a 500 m buffer around each location. Sensitivity analyses across buffer sizes (50–1000 m; Figure S1.24) indicated that 500 m maximized effect size and was used in subsequent analyses. As with the POI analysis, one-sided empirical p-values corrected with the Benjamini-Hochberg procedure were used to assess significance.

### 2.4. Ensemble random forest models to map jump dispersal risk

We used ensemble random forests (ERFs, Siders et al., 2020) to project *L. delicatula* jump risk across the study region (Fig. 1). ERFs address highly imbalanced classification problems by fitting multiple down-sampled random forests and averaging predictions across models. Models were fit on a binary response indicating whether a jump-dispersal event was observed during a survey event, with jump events identified 2014-2023 treated as presences (N = 130, see Section 3.1) and proactive search locations without detections from the same period treated as absences (N = 121,904). Model predictors included the distances to 22 POI types; densities of rail, primary road, and primary + secondary roads; and distance to the nearest established *L. delicatula* population from the prior biological year.

**Figure 1:**
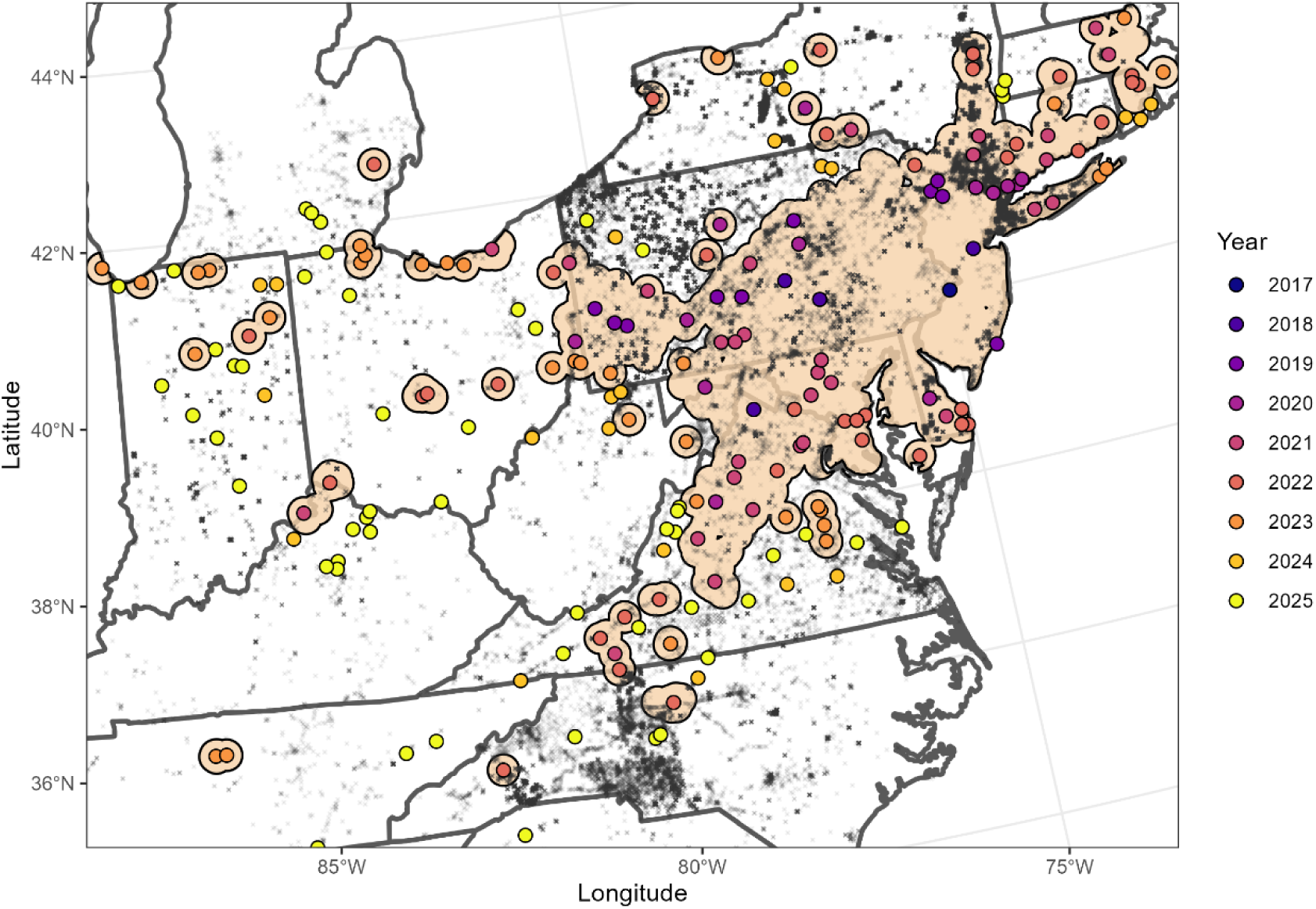
The spread of *L. delicatula* within its invaded range in the eastern U.S. has been marked by repeated long distance jump dispersal events, with jump dispersal sites identified algorithmically shown here colored by the biological year in which populations were first discovered. The tan area shows the 20 km buffer around all 2023 established *L. delicatula* populations, marking the extent of the invasion used to assess associations between jumps sites, properties, and transport infrastructure. Gray points show the locations of proactively searched recorded absences.

To identify a robust model, we compared six options: (1) a full model using all predictors, (2) a reduced full model using all POI distances and transport infrastructure values but excluding distance to existing *L. delicatula* populations, (3) a significant-pathways model using all predictors that were found to have significant associations with jump sites in the bootstrapping analysis (along with distance to *L. delicatula*), (4) a reduced significant-pathways model excluding distance from *L. delicatula*, (5) a forward-selection-based model obtained by iteratively assessing single predictors and adding whichever best improved model fit until additions stopped making improvements, and (6) a backward-selection-based model with the least important variable removed iteratively until model performance was reduced. For iterative model selection, we used 5-fold cross validation to assess model AUC with variance and selected the simplest model with AUC within 1 standard error of the highest value among all submodels. After exploring model settings, we used an ensemble size of 9 forests, 500 trees per forest, and sampled 3 covariates at each node. For each model, we calculated Accumulated Local Effects (ALEs) using the calc_ALE function (Siders et al., 2020).

Finally, projected 2024 risk was mapped for each of the six ERFs by generating predictions for the centroid of 2.5 × 2.5 km grid cells spanning the study region. Areas within 20 km of previously established populations were excluded from projections as they could not be colonized by jump dispersal under our definition. To evaluate out-of-sample predictive performance of each model, all 2024 and 2025 jump sites were withheld from model fitting and used for validation. We calculated the final AUC values for each model using 2024-2025 jump sites as positive cases and centroids of the uninvaded 2.5 × 2.5 km grid cells as negative cases using the roc.curve function in the ROCR package in R (Sing et al., 2005). We estimated confidence intervals for each model’s AUC based on 10,000 bootstrapping iterations and selected the simplest model that did not significantly differ from the highest observed mean AUC, using the Holm-Bonferroni method to correct for multiple comparisons among models.

### 2.5 State and municipality risk prioritization for jump dispersal surveillance

Using the selected ERF model (significant-pathways), we translated projected jump-dispersal risk into state- and municipality-level surveillance priorities. County subdivision boundaries were obtained from U.S. Census Bureau TIGER/Line shapefiles accessed with the *tigris* package in R (Walker, 2023). Each 2.5 × 2.5 km grid cell was assigned to the county subdivision containing its centroid, and subdivisions were scored using the maximum projected risk value among contained grid cells. For each state in the study region, we produced a focused jump-dispersal risk map and ranked county subdivisions by projected risk (Supplements 2–17). We additionally quantified the proportion of each state exceeding the 90^th^, 75^th^ and 50^th^ percentile projected risk value for confirmed 2024-2025 jump-dispersal events to rank states according to the extent of elevated jump-dispersal risk therein.

## 3. Results

### 3.1 Do survey locations exhibit proactive bias toward hypothesized property types and transport infrastructure?

Our workflow identified 207 *L. delicatula* jump-dispersal events from 2014–2025, with 130 events of those events from 2014-2023 which we used in the pathway analyses (Figure 1). Annual jump-dispersal counts were 0, 0, 0, 1, 4, 10, 13, 30, 38, 34, 22, and 55 from 2014–2025, respectively, indicating an overall increase in jump-event frequency through time (Mann–Kendall tau = 0.78, z = 2.81, p = 0.002). Inferred jump-event counts were moderately sensitive to the clustering distance parameters used to distinguish jumps from diffusive spread and to separate nearby satellite populations (Figure S1.25). Of the jump-dispersal events identified using our selected parameters, 73 were represented by a single established presence record in the year of discovery, whereas 135 included multiple established presence records.

State and federal surveillance showed strong proactive spatial sampling bias toward all hypothesized property types and transport infrastructure. We identified 121,904 proactive absence records located at least 20 km from previously confirmed *L. delicatula* populations from 2014–2023 used in pathway analyses (Section 3.2). Proactive search locations were closer to all 22 property types and occurred in areas with greater rail and road corridor density than randomly sampled regional background points (Table S1.1; Figure 2). For example, the mean distance from proactive search locations to the nearest shipping and distribution center was 55% of the mean distance from random background points (95% CI [54 %, 57 %] Figure 2A). Sampling bias was strongest for wineries. The average distance to the nearest winery for proactive sampling points was just 41% of that of random background points (95% CI [40%, 43%]). These results confirm that agency surveys were concentrated near all hypothesized influential property types, making regional background point comparisons alone insufficient for statistically identifying jump dispersal pathways. Instead, proactive search points must be used.

**Figure 2:**
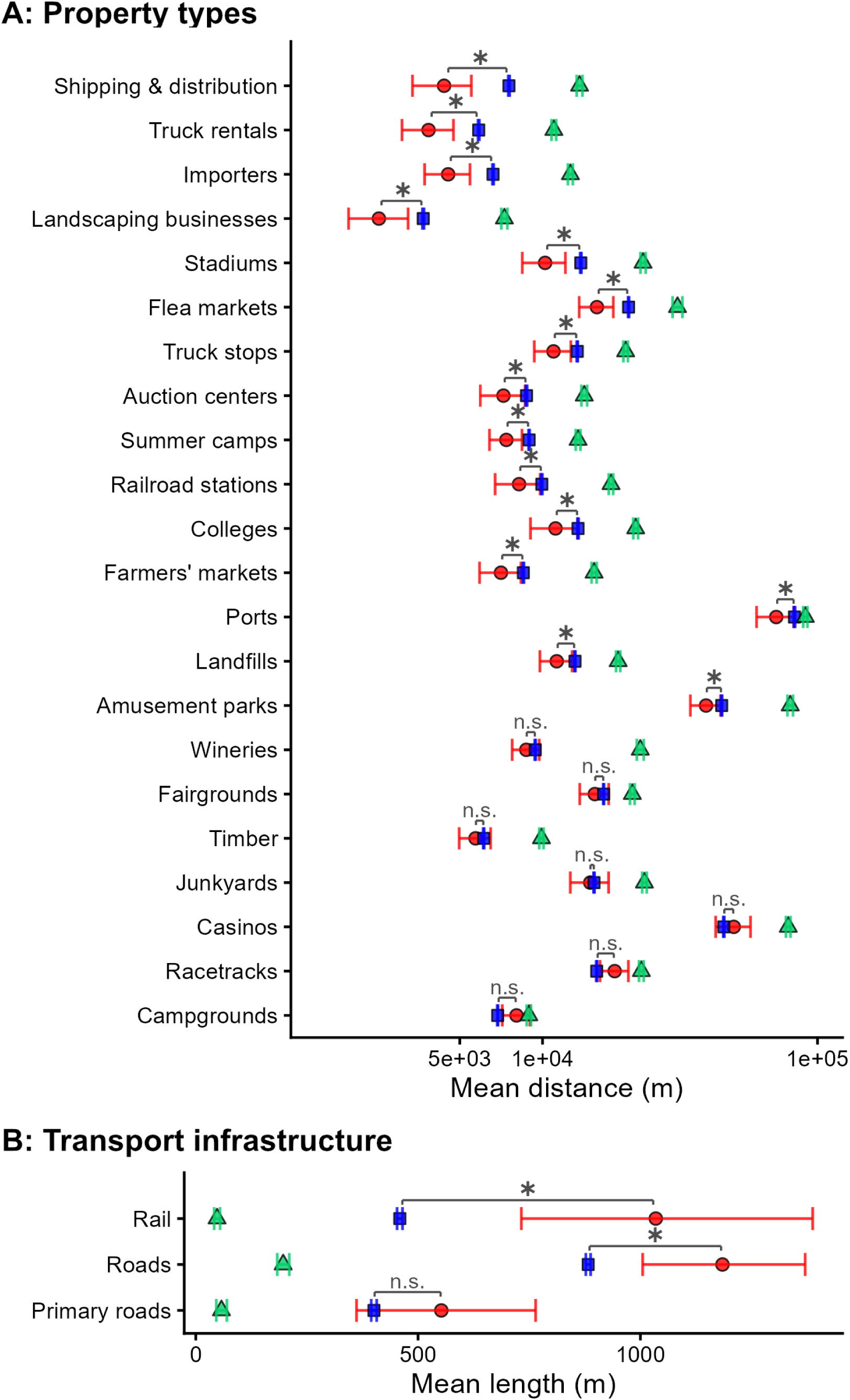
*Lycorma delicatula* jump dispersal sites were significantly associated with several property and transport infrastructure types. In panel A, jump dispersal sites (red circles) were located closer to some property types than expected based on two null models: regional background points (green triangles) and proactive sampling sites (blue squares). For each property type, we measured the mean distance between 130 *L. delicatula* jump dispersal sites from 2014-2023 and the nearest property of that type. For comparison, we bootstrapped estimated means for 5000 random points spanning the study region and for the 121,904 points where proactive search by government surveyors occurred over the same time period. Points indicate the mean, while error bars show the 95% confidence interval. Annotated brackets above points indicate the significance of the difference between jump dispersal sites’ distances and proactive search’s, corrected for multiple comparisons using the Benjamini-Hochberg procedure. Panel B shows that greater lengths of transport infrastructure were found around jump dispersal sites than around either of the other types of points. Jump dispersal sites had significantly more rail within a 500 m radius than random regional background points or proactively surveyed locations. Similarly, jump dispersal sites’ buffers held greater lengths of roads, on average, than proactive search sites or regional background sites. When only primary roads (interstate highways) were considered, though, no difference between jump sites and proactive search sites was found.

### 3.2 Which features are associated with jump dispersal after bias correction?

After accounting for survey bias, jump-dispersal events were significantly associated with 15 of 22 property types and 2 of 3 transport infrastructure types. Jump sites were closer than proactive search locations to the following property types: shipping and distribution centers, truck rentals, importers, landscaping businesses, stadiums, flea markets, truck stops, auction centers, summer camps, railroad stations, colleges, farmers’ markets, ports, landfills, and amusement parks (Figure 2; Table S1.1). The strongest association was with shipping and distribution centers (Cliff’s Δ = 0.27, 95% CI [0.18, 0.36]). More than 78% of jump points occurred within 5 km of a shipping and distribution center, compared with 54% of proactive search locations. Other notable associations included truck rentals (Δ = 0.24, 95% CI [0.15, 0.33]), importers (Δ = 0.20, 95% CI [0.11, 0.29]), and flea markets (Δ = 0.18, 95% CI [0.09, 0.27]). Jump-dispersal sites had greater rail density within 500 m than proactive search locations (Figure 2B; Cliff’s Δ = −0.21, 95% CI [−0.30, −0.12]). Jump sites were also found more in areas of higher combined primary and secondary road density, but the effect was weaker (Figure 2B; Cliff’s Δ = −0.17, 95% CI [−0.27, −0.07]). Some notable operationally relevant hypothesized property types including wineries, junkyards, and timber facilities, were not associated with jump-dispersal events after accounting for proactive survey bias. Similarly, primary roads alone, such as interstate highways, were also not associated with jump-dispersal events.

### 3.3 Can POI and transport infrastructure data predict risk and prioritize surveillance?

Ensemble random forest models successfully discriminated the withheld 2024-2025 jump-dispersal sites from uninvaded background locations, with AUC values ranging from 0.76 to 0.93 across model formulations (Table 2, selected predictors for each model listed in Table S1.2). Models that included distance to previously established *L. delicatula* populations outperformed corresponding models without this predictor. The significant-pathways model, which included only features identified as significant in the bias-corrected bootstrapping analyses plus distance to prior-year established populations, performed best (2024 AUC = 0.93, 95% CI [0.90, 0.96]), slightly exceeding the full model with all predictors (AUC = 0.91, 95% CI [0.87, 0.95]) and the model selected based on forward selection (AUC = 0.92, 95% CI [0.89, 0.95]). Because the significant-pathways model performed comparably to the more complex full model while using fewer predictors, it was selected for further analyses.

**Table 2:** Area under the receiver operating characteristic curve for six ensemble random forest.

| Model | 2024 AUC*† | 2024-2025 AUC*† |
| --- | --- | --- |
| Full | 0.91 [0.86, 0.95]<sup>ab</sup> | 0.90 [0.88, 0.92]<sup>a</sup> |
| Full (no distance to *L. delicatula*) | 0.78 [0.65, 0.88]<sup>c</sup> | 0.76 [0.70, 0.82]<sup>b</sup> |
| Significant pathways | 0.93 [0.90, 0.96]<sup>a</sup> | 0.90 [0.88, 0.92]<sup>a</sup> |
| Significant pathways (no distance to *L. delicatula*) | 0.83 [0.72, 0.91]<sup>c</sup> | 0.79 [0.73, 0.84]<sup>b</sup> |
| Forward selection | 0.92 [0.89, 0.95]<sup>a</sup> | 0.89 [0.86, 0.92]<sup>a</sup> |
| Backward selection | 0.87 [0.82, 0.92]<sup>ab</sup> | 0.85 [0.82, 0.88]<sup>c</sup> |

\* Values are AUC [95% bootstrap CI]. Models sharing a superscript letter within a column did not differ significantly (bootstrap comparison, $\alpha = 0.05$ ).
† Evaluated against jump dispersal sites detected in 2024 only, or 2024 and 2025 combined, versus background points from a $2.5 \times 2.5$ km grid spanning the region available for colonization in 2024. Models were fit using *L. delicatula* distribution through 2023.

Projected risk maps identified elevated jump-dispersal risk beyond the existing invaded range, particularly in developed and transportation-dense regions. High-risk areas included portions of northeast Illinois and Ohio, southeast Michigan around the Detroit outbreak, and central North Carolina near the Kernersville-centered population (Figure 3; Appendices 2–17). Plots of ALEs indicated that risk generally increased near significant property types and with greater rail density, whereas road-density effects were weaker and less consistent (Figure S1.26). Distance to previously established populations also contributed strongly to projected risk, consistent with risk being highest in areas beyond but still near the expanding invasion.

**Figure 3:**
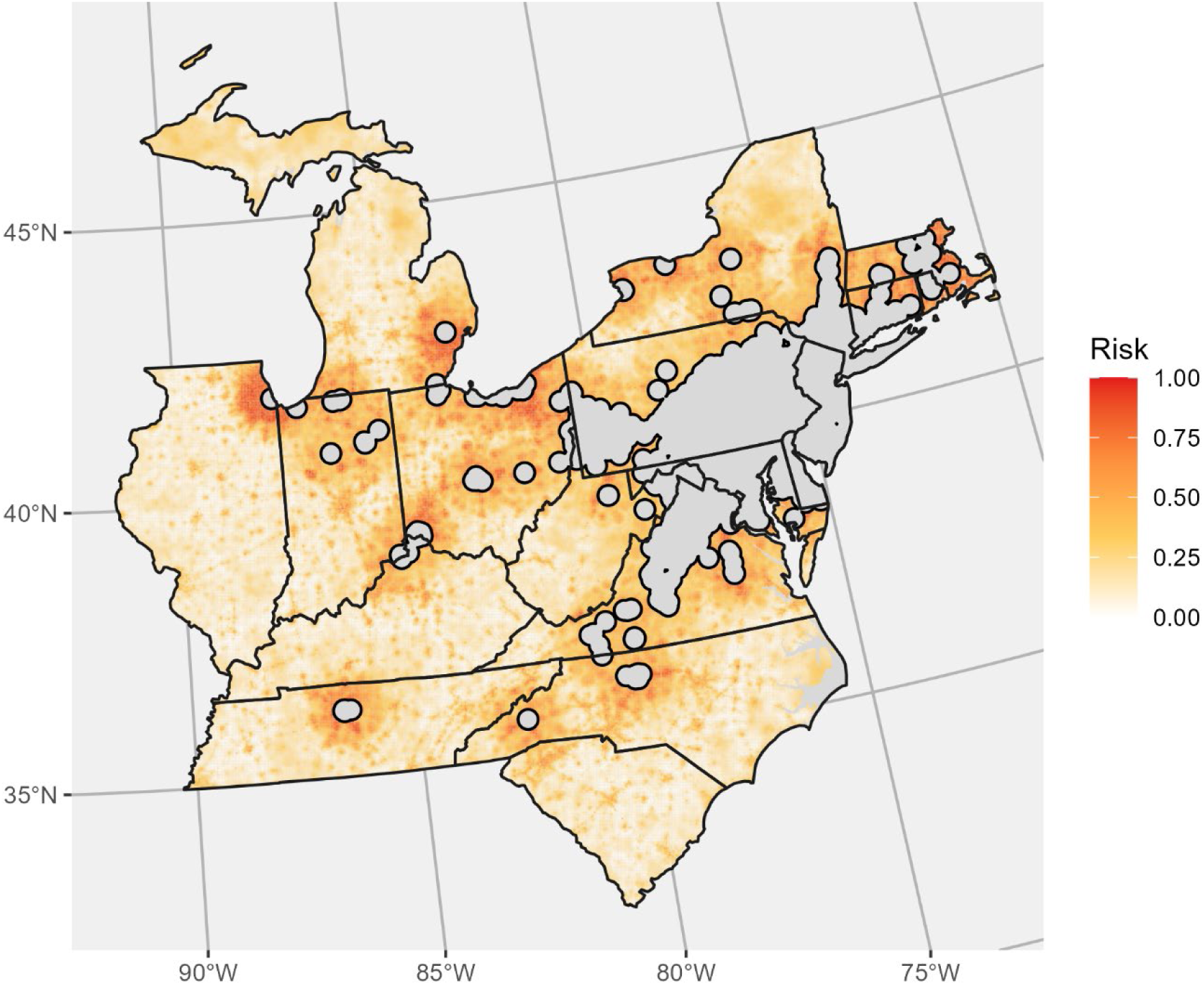
Projected risk of colonization by jump dispersal varied across the study region in the northeast United States, with developed areas near existing *L. delicatula* populations generally identified as having the greatest risk. The map is colored based on the scaled risk projected by a fitted ensemble random forest model fit using spatial predictors identified in bias-corrected pathway analysis, along with distance from existing *L. delicatula* populations. Solid grey areas fall within 20 km of existing established populations of *L. delicatula* and therefore can no longer be newly colonized by jump dispersal, though diffusive spread and infilling may still occur in these locations. Values are scaled to range from 0 (least risk) to 1 (greatest risk).

Management-relevant areas (2.5 × 2.5 km grid cells) that were projected to be at elevated risk of colonization by jump dispersal were unevenly distributed across states within the study region (Figure 4). Illinois held the greatest proportion of area at the highest risk level, with 37% of that area falling within the state. Ohio, Michigan, Massachusetts, and Indiana also held substantial proportions of the high-risk area across levels. Of the area with projected risk greater than 0.77, 25% was in Illinois, 20% was in Ohio, and 17% was in Michigan. As expected, states with longer history of invasion (e.g. Pennsylvania and New Jersey) held less high-risk area than more recently colonized states in part because much of the area there is already occupied and therefore can no longer be colonized by jump dispersal. The risk described here is an assessment of the hazard of future colonization by jump dispersal, not of persistence of existing populations or impact.

**Figure 4:**
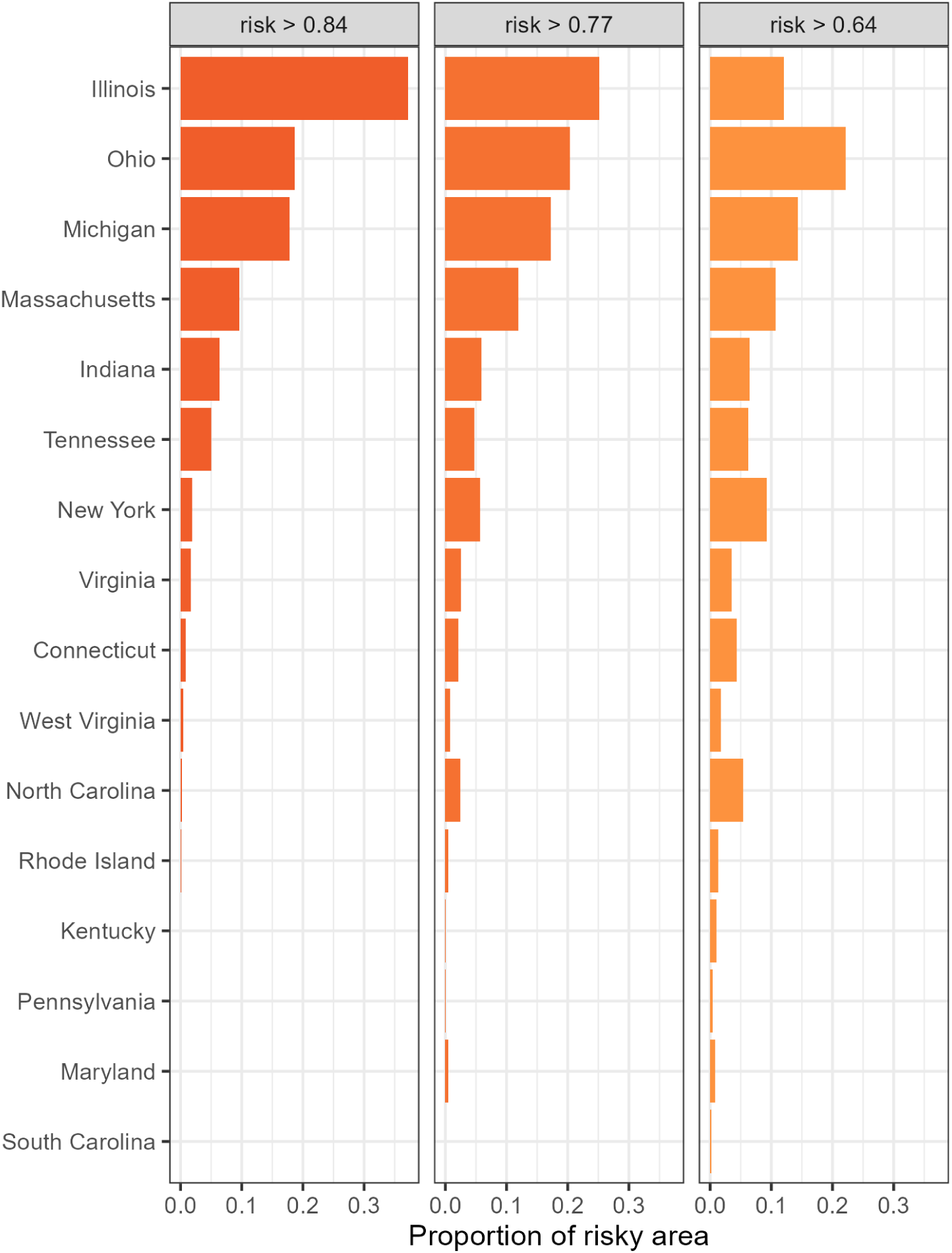
Management-relevant areas (2.5 × 2.5 km grid cells) at elevated risk of colonization by jump dispersal were unevenly distributed across states within the study region. To compare the amount of at-risk area among states, we identified risky areas based on three risk levels: areas with risk greater than the 90^th^ percentile among 2024 jump sites (risk score = 0.83), the 75^th^ percentile (0.77), and the 50^th^ percentile (0.63). Then for each level of risk, we identified the proportion of that total area falling into each of the states within the study region. Illinois held the greatest proportion of area at the highest risk level, with 37% of that area falling within the state. Ohio and Michigan also held substantial proportions of the high-risk area across levels.

## 4 Discussion

### 4.1 Key challenge: Operational surveillance records require bias-aware comparison sets

Here, we developed a workflow to estimate invasive species jump-dispersal risk by integrating agency surveillance records with point-of-interest and transport infrastructure data while accounting for spatial sampling bias in search effort. With this workflow, we found that state and federal agency survey effort for invasive *L. delicatula* was strongly concentrated near hypothesized pathways (Figure 2). Proactive surveys were closer to all evaluated property types and occurred in areas with greater rail and road density than expected under random spatial sampling. The results are consistent with prior studies showing that opportunistic, preferential, or unstructured biodiversity records are often biased toward accessible or human-influenced locations (Kadmon et al., 2004; Phillips et al., 2009). As a consequence of such bias, invasion pathway analyses will overstate the importance of many pathways if invasion processes are confounded with patterns in where surveyors were directed to look (Hughes et al., 2021; Pennino et al., 2019; Sorbe et al., 2023; Szymura et al., 2023).

A key challenge in accounting for operational surveillance bias is that the purpose of individual surveys is often not fully recoverable from the data alone. If the purpose were known, the bias could theoretically be modeled. For the *L. delicatula* invasion, records were generated by multiple agencies and operational programs conducting surveys for different purposes, including delimiting known populations, monitoring occupied areas, evaluating management actions, and searching ahead of the invasion front for new satellite populations. No metadata exists on these purposes. We isolated the surveys ahead of the invasion front for use as our comparison dataset. This approach parallels bias-correction strategies in species distribution modeling, where background or comparison data are selected to match the bias structure of occurrence data rather than represent the landscape at random (Phillips et al., 2009). When survey metadata on intent or effort can be directly characterized, alternative approaches such as explicit effort modeling, simulated bias surfaces, or spatial filtering may be appropriate (Barber et al., 2022; Stolar & Nielsen, 2015). In our workflow, we produced bias-aware comparison sets with proactive absences, which provided an empirical comparison set for estimating the spatial footprint of search effort. Then, nested clustering reduced spatial over-representation by treating clustered detections from the same satellite population as a single jump-dispersal event (Inman et al., 2021; Vollering et al., 2019). Thus, operational surveillance records are biased, but usable if the relevant search process is characterized to produce a bias-aware comparison set.

### 4.2 Bias-corrected associations support pathway-diverse surveillance

For *L. delicatula*, most of the 25 proposed landscape features were found to be significantly associated with jump dispersal sites. After comparing jump-dispersal sites to the bias-aware proactive search locations, *L. delicatula* jump events were associated with 15 property types and two transport infrastructure types (Figure 2). These included shipping and distribution centers, truck stops and rentals, importers, landscaping businesses, and rail yards and stations. In contrast, seven property types, including wineries and timber facilities, and one transport infrastructure type (primary roads alone) were not associated with jump-dispersal events after accounting for proactive survey bias. The significant associations are consistent with prior evidence that *L. delicatula* spreads along transportation networks at broad spatial scales (Cook et al., 2021; Huron et al., 2022; C. M. Jones et al., 2021; Ladin et al., 2023; Vetrovec & Payne, 2023). The identified features associated with *L. delicatula* jump dispersal share characteristics that plausibly facilitate movement of egg masses or mobile life stages: high vehicle turnover, freight handling, outdoor storage, and repeated movement of goods or materials. These mechanisms are consistent with the species’ biology, including oviposition on transported substrates (Liu, 2019; Urban & Leach, 2023) and experimental evidence that nymphs and adults can remain attached to vehicle surfaces under high wind speeds (Elsensohn et al., 2024).

*L. delicatula* jump dispersal is associated with a diverse set of human-mediated transport contexts (Gippet et al., 2019). There is not a single dominant pathway. This distinction is important for management because it indicates that the relevant target is not a single property type, but multiple property and transport infrastructure types statistically associated with newly detected satellite populations after accounting for survey bias. While we have demonstrated predictability in *L. delicatula* jump dispersal sites, long distance dispersal events are rare and random, and even sites that are not identified as likely jump dispersal sites may end up being epicenters of new outbreaks given the right confluence of events. Operationally, these results support pathway-diverse surveillance combined with public outreach, including continued educational campaigns that encourage “look before you leave” practices to reduce human-mediated dispersal of *L. delicatula* (Roe et al., 2024; Urban et al., 2021). Concentrating surveillance only around the strongest associated property types, such as shipping and distribution centers or rail yards, could reduce operational costs and detect some jump events, but would likely miss others arising through less frequent transport contexts, weakening early detection and increasing the risk of costly secondary outbreaks (Foxcroft et al., 2019; Hulme, 2009; Turbelin et al., 2022). However, future work is needed to understand if there are interactions among pathways and proximity to established populations that may cause some pathways to be dominant the further an uninvaded area is from the invasion front.

### 4.3 Pathway-informed models prioritize surveillance beyond the invasion front

Ensemble random forest models integrated the effects of multiple landscape features on jump dispersal risk to successfully discern withheld jump site locations from the regional background. The best-performing model had strong predictive power (AUC 0.93). This model included the 15 property types and 2 transport infrastructure types identified under the bias-aware comparison set and also distance to the nearest *L. delicatula* population present in the prior year as predictors. Indeed, distance to the invasion front was identified as impactful and included in all of the top best fit models (Table 2). Projecting forward one year to withheld data, models including distance from prior *L. delicatula* populations had AUC ranging from 0.87 to 0.93, while those without had AUC ranging from 0.78 to 0.83. Other predictors that were included in all models regardless of selection procedure were: distance to shipping centers, distance away from summer camps, and distance to truck stops. The repeated identification of these variables as valuable predictors points toward pathway relationships with *L. delicatula* jump dispersal that may be distinct from the broader association with overall human traffic.

### 4.4 Jump dispersal risk is unevenly spread over the Eastern U.S

The best performing model identified several areas in the Eastern U.S. with high risk of jump dispersal (Fig. 3). At the state level, states relatively recently colonized by *L. delicatula* with large as-yet uninvaded areas including cities emerged as having the largest proportions of highly at-risk area. Illinois, Ohio, and Michigan all had areas identified as high-risk for future colonization by jump dispersal (Figure 4). States with longer histories of invasion by *L. delicatula* such as Pennsylvania, New Jersey, and Maryland, were not as high-risk because the potential for colonization there has already largely been realized. Within states, risk was highest in developed areas near previously invaded areas. For example, human population centers in Ohio including, Akron and Dayton were similarly identified as very high-risk. The densely populated area around the *L. delicatula* population in Chicago, Illinois, has the highest risk of subsequent jump dispersal events of the states within our study region. Colonization of additional shipping centers in this area poses considerable risk due to long-distance transport emanating from Illinois to the uninvaded western states. Prioritizing survey effort in the highest risk locations is likely to aid early detection within affected states.

### 4.5 Future risk evaluations can be refined

Projections of long-distance jump dispersal risk informed by invasion pathways can provide useful information for mechanistic spread projections. Mechanistic spatial spread models, for example, have previously included information on railways, simulating hitchhiking travel along rail to capture patterns of spread (Jones et al., 2021). With evidence that *L. delicatula* makes use of multiple dispersal modes to travel long distances, an alternative long distance dispersal component may provide a useful addition to such models. Integrating refined long-distance dispersal risk with mechanistic spread models can guide managers preparing for potential spread into as-yet unaffected grape growing regions that may suffer substantial economic harm if *L. delicatula* disperses there (Huron et al., 2022; C. Jones et al., 2022).

As additional data on *L. delicatula* biology and spread patterns are gathered, refined estimates of *L. delicatula* jump dispersal risk may be derived. For an outlying population of *L. delicatula* to become established, individuals must disperse there, and conditions there must be suitable for survival and reproduction (Renault et al., 2018). Because populations only result when both factors (dispersal and habitat suitability) are present (Gallien et al., 2015), we were unable to parse the factors influencing one of these processes from those influencing the other. We focused on characterizing relationships between human infrastructure and *L. delicatula* jump dispersal sites, but adding additional predictors including land cover variables, climate variables, and/or host plant distribution data may result in stronger performance (With, 2002). In particular, we expect detailed distribution data describing where *L. delicatula*’s preferred host plant *Ailanthus altissima* occurs may be highly informative. Efforts to map this plant have the potential to improve surveillance.

This analysis focused on 18 northeastern U.S. states (and the District of Columbia) where *L. delicatula* presence has been reported, and the associations between jump dispersal sites and human infrastructure that we observed in this region may have limited applicability to other regions with differing land use patterns, history of development, or host plant availability. In a hypothetical region where suitable hostplants for *L. delicatula* are absent, for example, proximity to impactful property types is irrelevant to whether or not *L. delicatula* will colonize. Without suitable hosts, no new population can arise regardless of dispersal frequency. In the study region, *L. delicatula*’s preferred host plant *Ailanthus altissima* is frequently found along rails, due in part to the timing of rail construction disturbance co-occurring with range expansion by this invasive plant (Kasson et al., 2013). Associations with rail may therefore be weaker in areas where *A. altissima* is absent or native. As the invasion continues to grow, updated and expanded risk mapping will be needed to incorporate new information.

### 4.6 Conclusions

Our workflow identified associations between property types, transport infrastructure, and long-distance jump dispersal by invasive pests. Significantly associations informed a strongly discriminative ensemble random forest model to project future jump dispersal risk. We presented model output in three formats: the projected risk maps themselves (Fig 4, Supplemental materials 2-17), state-by-state lists of subcounty jurisdictions ranked based on the maximum projected risk therein (Supplemental materials 2-17), and a summary of which states held the greatest proportion of risky locations under three data-derived threshold levels of risk (Figure 4). These options allow stakeholders to make use of relevant projections to meet their particular goals. When prioritizing surveillance effort across states, the region-wide map and state rankings can guide decisions. For state agencies allocating early detection efforts within a state, the state maps and subcounty lists may guide targeted early detection surveys. Early detection efforts can be strengthened with targeted surveillance informed by bias-aware modeling.

## Declaration of generative AI and AI-assisted technologies in the manuscript preparation process

During the preparation of this work, the authors used Claude by Anthropic to parallelize code for faster processing. After using this tool/service, the authors reviewed and edited the content as needed and take full responsibility for the content of the published article.

## Data statement

Anonymized *L. delicatula* occurrence data are available in the lydemapr data base online at https://github.com/ieco-lab/lydemapr. The full dataset used in these analyses cannot be shared due to data privacy requirements.

## CRediT author statement

**Joseph Keller**: Conceptualization, Methodology, Software, Formal Analysis, Data Curation, Writing – Original Draft, Funding Acquisition. **Nadège Belouard:** Conceptualization, Methodology, Writing – Review & Editing. **Sebastiano De Bona:** Conceptualization, Data Curation, Writing – Review & Editing. **Jocelyn E. Behm:** Conceptualization, Methodology, Supervision, Writing – Review & Editing. **Matthew R. Helmus:** Conceptualization, Methodology, Data Curation, Writing – Review & Editing, Supervision, Project administration, Funding acquisition.

## Supporting information

S2 Connecticut risk

S3 Illinois risk

S4 Indiana risk

S5 Kentucky risk

S6 Maryland risk

S7 Massachusetts risk

S8 Michigan risk

S9 New York risk

S10 North Carolina risk

S11 Ohio risk

S12 Pennsylvania risk

S13 Rhode Island risk

S14 South Carolina risk

S15 Tennessee risk

S16 Virginia risk

S17 West Virginia risk

