## Supplementary material for "Mapping long-distance jump dispersal risk by leveraging transport infrastructure data and sampling bias correction: a case study with the invasive spotted lanternfly (*Lycorma delicatula*)": S1 supporting information

**Table S1.1:** Bootstrapping results comparing proximity between potential invasion hubs and locations where jump dispersal by *L. delicatula* has been observed (jump sites), proactive search sites that were surveyed before nearby *L. delicatula* populations were known to exist (proactive search) and randomly located points spanning the study region (regional background). Pathway names in bold were identified as showing a significant association with jump dispersal sites based on the proactive search/jump site distance comparisons.

| **Pathway** | **Regional background distance (km)** | **Proactive search distance (km)** | **Jump site distance (km)** | **Regional background - proactive search distance (km)** | **Proactive search/regional background Cliff's delta** | **Jump site - proactive search distance (km)** | **Jump site/Proactive search Cliff's delta** |
|---|---|---|---|---|---|---|---|
| **Shipping and distribution center** | 13.6 [13.3, 14.0] | 7.5 [7.5, 7.6] | 4.4 [3.4, 5.5] | 6.1 [5.8, 6.4] | -0.41 | -3.2 [-4.2, -2.0] | 0.27 |
| **Truck rental** | 11.0 [10.8, 11.2] | 5.9 [5.8, 5.9] | 3.8 [3.1, 4.7] | 5.1 [4.9, 5.4] | -0.47 | -2.0 [-2.8, -1.1] | 0.24 |
| **Importer** | 12.6 [12.4, 12.9] | 6.6 [6.6, 6.6] | 4.5 [3.7, 5.4] | 6.0 [5.8, 6.3] | -0.47 | -2.1 [-2.9, -1.2] | 0.20 |
| **Flea market** | 31.0 [29.7, 32.3] | 20.6 [20.5, 20.6] | 15.8 [13.6, 18.1] | 10.4 [9.2, 11.7] | -0.18 | -4.8 [-7.0, -2.5] | 0.18 |
| **Landscaping business** | 7.3 [7.1, 7.5] | 3.7 [3.7, 3.7] | 2.5 [2.0, 3.2] | 3.6 [3.4, 3.8] | -0.45 | -1.1 [-1.7, -0.4] | 0.18 |
| **Stadium** | 23.2 [22.7, 23.7] | 13.8 [13.7, 13.9] | 10.2 [8.4, 12.1] | 9.4 [8.9, 9.9] | -0.37 | -3.6 [-5.3, -1.7] | 0.18 |
| **Railroad station** | 17.7 [17.4, 18.1] | 9.9 [9.9, 10.0] | 8.2 [6.7, 9.8] | 7.8 [7.5, 8.1] | -0.44 | -1.7 [-3.2, -0.1] | 0.16 |
| **College** | 21.8 [21.4, 22.3] | 13.5 [13.4, 13.5] | 11.2 [9.0, 13.4] | 8.4 [7.9, 8.8] | -0.37 | -2.3 [-4.4, -0.1] | 0.16 |
| **Auction center** | 14.2 [13.8, 14.5] | 8.7 [8.7, 8.8] | 7.2 [5.9, 8.6] | 5.4 [5.1, 5.8] | -0.36 | -1.5 [-2.8, -0.1] | 0.14 |
| **Truck stop** | 20.1 [19.7, 20.4] | 13.4 [13.3, 13.4] | 11.0 [9.3, 12.7] | 6.7 [6.3, 7.1] | -0.33 | -2.4 [-4.0, -0.7] | 0.14 |
| **Amusement park** | 79.6 [77.7, 81.5] | 44.8 [44.7, 45.0] | 39.3 [34.5, 44.4] | 34.8 [32.9, 36.6] | -0.38 | -5.5 [-10.4, -0.5] | 0.12 |
| **Landfill** | 18.8 [18.5, 19.2] | 13.1 [13.1, 13.2] | 11.3 [9.8, 12.8] | 5.7 [5.4, 6.1] | -0.30 | -1.9 [-3.3, -0.3] | 0.11 |
| **Farmers' market** | 15.4 [15.1, 15.7] | 8.5 [8.5, 8.6] | 7.1 [5.9, 8.3] | 6.9 [6.6, 7.2] | -0.43 | -1.5 [-2.6, -0.2] | 0.10 |

| | | | | | | | |
|---|---|---|---|---|---|---|---|
| Junkyard | 23.5 [23.0, 23.9] | 15.4 [15.3, 15.5] | 14.9 [12.6, 17.4] | 8.1 [7.6, 8.5] | -0.32 | -0.5 [-2.8, 2.0] | 0.09 |
| **Summer camp** | 13.5 [13.2, 13.7] | 8.9 [8.9, 9.0] | 7.4 [6.4, 8.4] | 4.5 [4.3, 4.8] | -0.32 | -1.5 [-2.5, -0.5] | 0.08 |
| Fairgrounds | 21.2 [20.8, 21.6] | 16.7 [16.6, 16.7] | 15.5 [13.7, 17.4] | 4.5 [4.1, 5.0] | -0.16 | -1.2 [-3.0, 0.7] | 0.08 |
| **Port** | 90.3 [88.7, 91.9] | 82.5 [82.0, 82.9] | 70.8 [60.0, 81.9] | 7.9 [6.2, 9.5] | -0.14 | -11.7 [-22.4, -0.5] | 0.07 |
| Timber | 9.9 [9.7, 10.1] | 6.1 [6.1, 6.1] | 5.7 [5.0, 6.5] | 3.8 [3.6, 4.0] | -0.39 | -0.4 [-1.1, 0.4] | 0.06 |
| Winery | 22.7 [22.0, 23.3] | 9.4 [9.4, 9.4] | 8.7 [7.8, 9.7] | 13.3 [12.6, 13.9] | -0.49 | -0.7 [-1.7, 0.3] | 0.01 |
| Casino | 78.3 [76.8, 79.8] | 45.6 [45.4, 45.8] | 49.6 [42.6, 57.1] | 32.7 [31.2, 34.1] | -0.40 | 4.0 [-3.0, 11.4] | -0.02 |
| Racetrack | 22.9 [22.4, 23.3] | 15.7 [15.7, 15.8] | 18.3 [16.2, 20.5] | 7.1 [6.7, 7.6] | -0.29 | 2.6 [0.4, 4.8] | -0.10 |
| Campground | 8.9 [8.8, 9.1] | 6.9 [6.8, 6.9] | 8.0 [7.1, 9.0] | 2.0 [1.9, 2.2] | -0.20 | 1.2 [0.3, 2.1] | -0.10 |
| **Primary and secondary roads** | 196.6 [183.3, 210.3] | 882.6 [877.3, 888.0] | 1184.6 [1005.2, 1370.6] | 686.1 [671.4, 700.3] | 0.46 | 301.9 [122.3, 488.1] | -0.17 |
| **Rail** | 47.9 [41.3, 54.7] | 458.8 [452.7, 464.9] | 1034.6 [732.2, 1387.5] | 410.9 [401.8, 420.1] | 0.25 | 575.8 [273.7, 928.2] | -0.21 |
| Primary roads | 57.7 [46.3, 70.0] | 400.7 [394.6, 406.8] | 552.2 [361.5, 764.4] | 343.0 [329.2, 356.0] | 0.13 | 151.5 [-39.1, 363.9] | -0.10 |

**Table S1.2:** Predictors included in the ensemble random forest models

| Predictor | Included in model: Full | Full w/o invasion context | Significant pathways | Significant pathways w/o invasion context | Forward selection | Backward selection |
|---|---|---|---|---|---|---|
| Distance to amusement park | X | X | X | X | | X |
| Distance to casino | X | X | | | X | X |
| Distance to auction center | X | X | X | X | | |
| Distance to campground | X | X | | | X | |
| Distance to college | X | X | X | X | | |
| Distance to fairgrounds | X | X | | | | X |
| Distance to farmers' market | X | X | X | X | | |
| Distance to flea market | X | X | X | X | | |
| Distance to importer | X | X | X | X | | |
| Distance to junkyard | X | X | | | | |
| Distance to landfill | X | X | X | X | | |
| Distance to landscaping business | X | X | X | X | X | |
| Distance to race track | X | X | | | | |
| Distance to railroad station | X | X | X | X | X | |
| Distance to shipping center | X | X | X | X | X | X |
| Distance to stadium | X | X | X | X | | |
| Distance to summer camp | X | X | X | X | X | X |
| Distance to timber facility | X | X | | | | |
| Distance to truck rental | X | X | X | X | | |
| Distance to truck stop | X | X | X | X | X | X |
| Distance to winery | X | X | | | | |
| Distance to port | X | X | X | X | | |
| Rail length within 500 m | X | X | X | X | X | |
| Primary road length within 500 m | X | X | | | X | |

| | | | | | | |
|---|---|---|---|---|---|---|
| Primary + secondary road length within 500 m | X | X | X | X | X | |
| Distance to existing spotted lanternfly pop. | X | | X | | X | X |

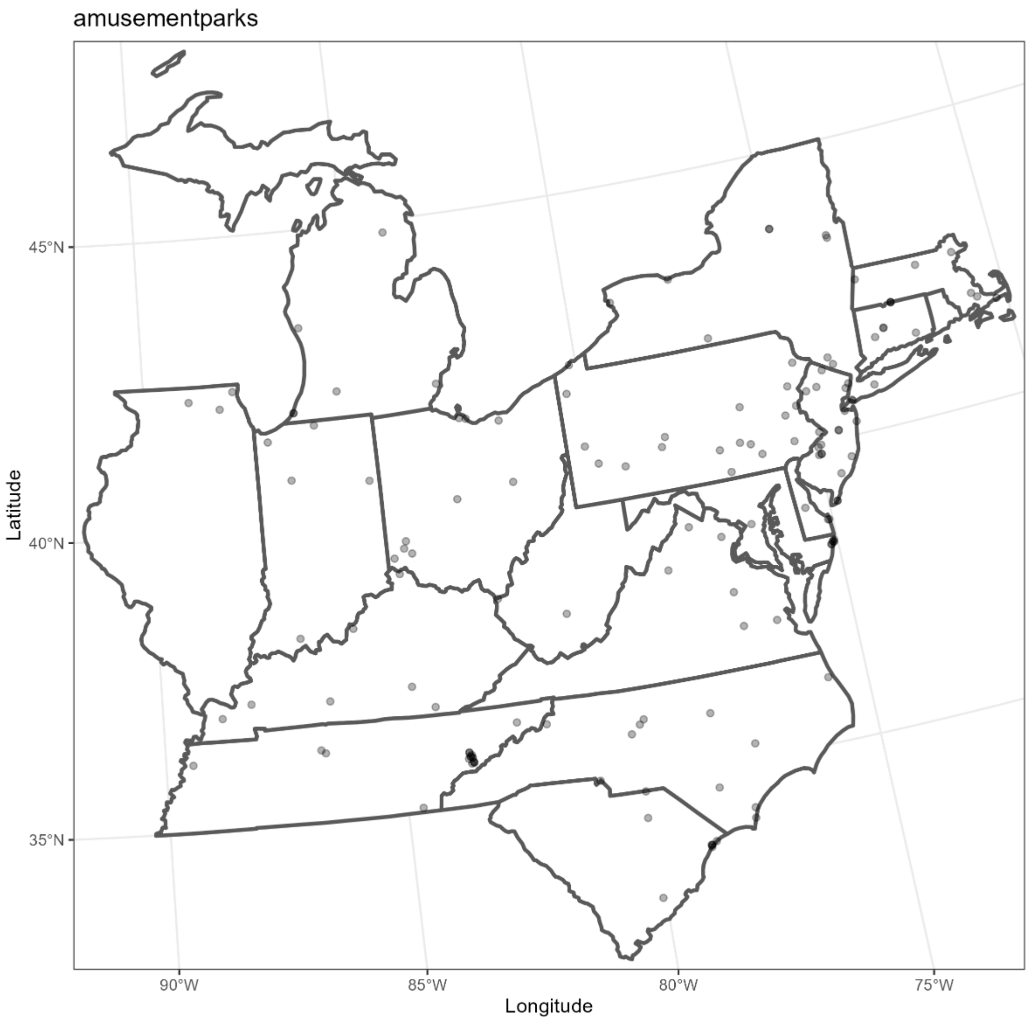


**Figure S1.1:** Amusement park locations within the study region in the Dataplor point of interest database used in the pathway analyses (Dataplor, 2022).

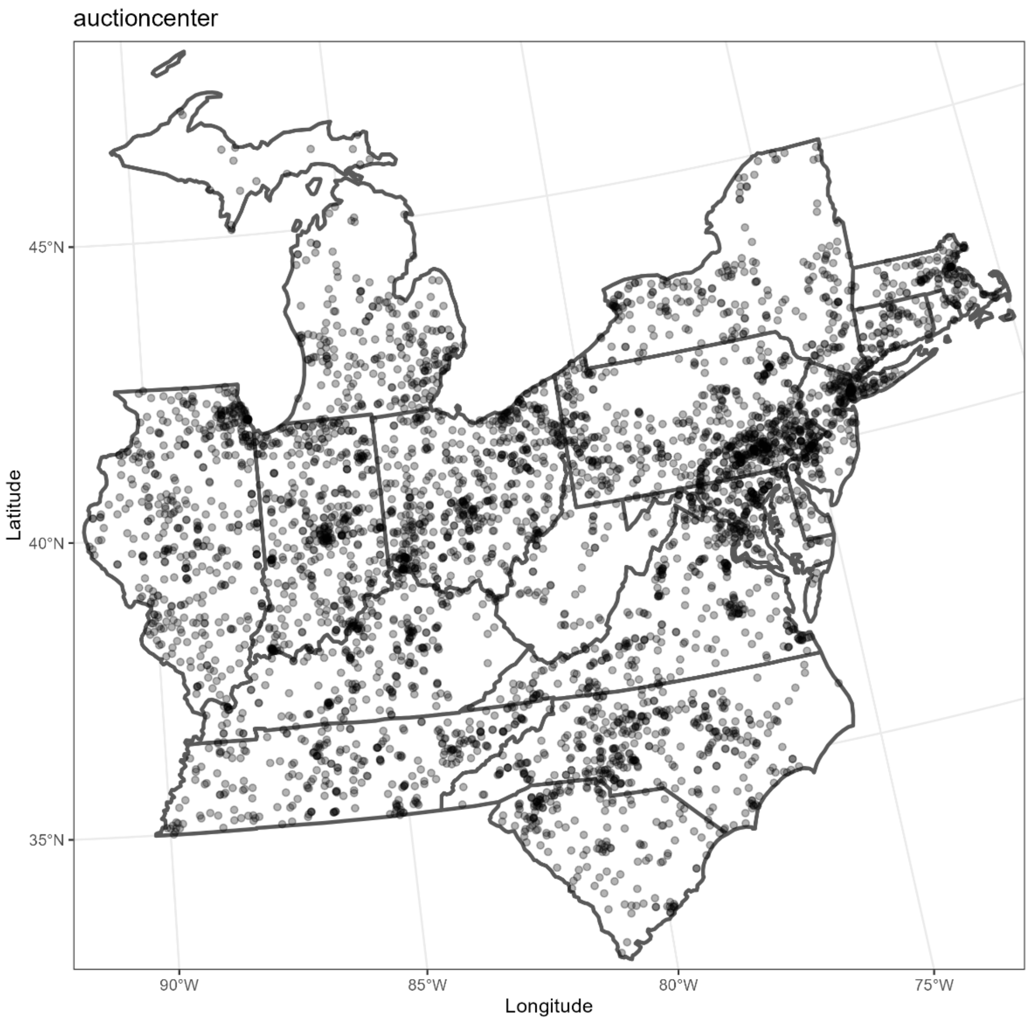


**Figure S1.2:** Auction center locations within the study region in the Dataplor point of interest database used in the pathway analyses (Dataplor, 2022).

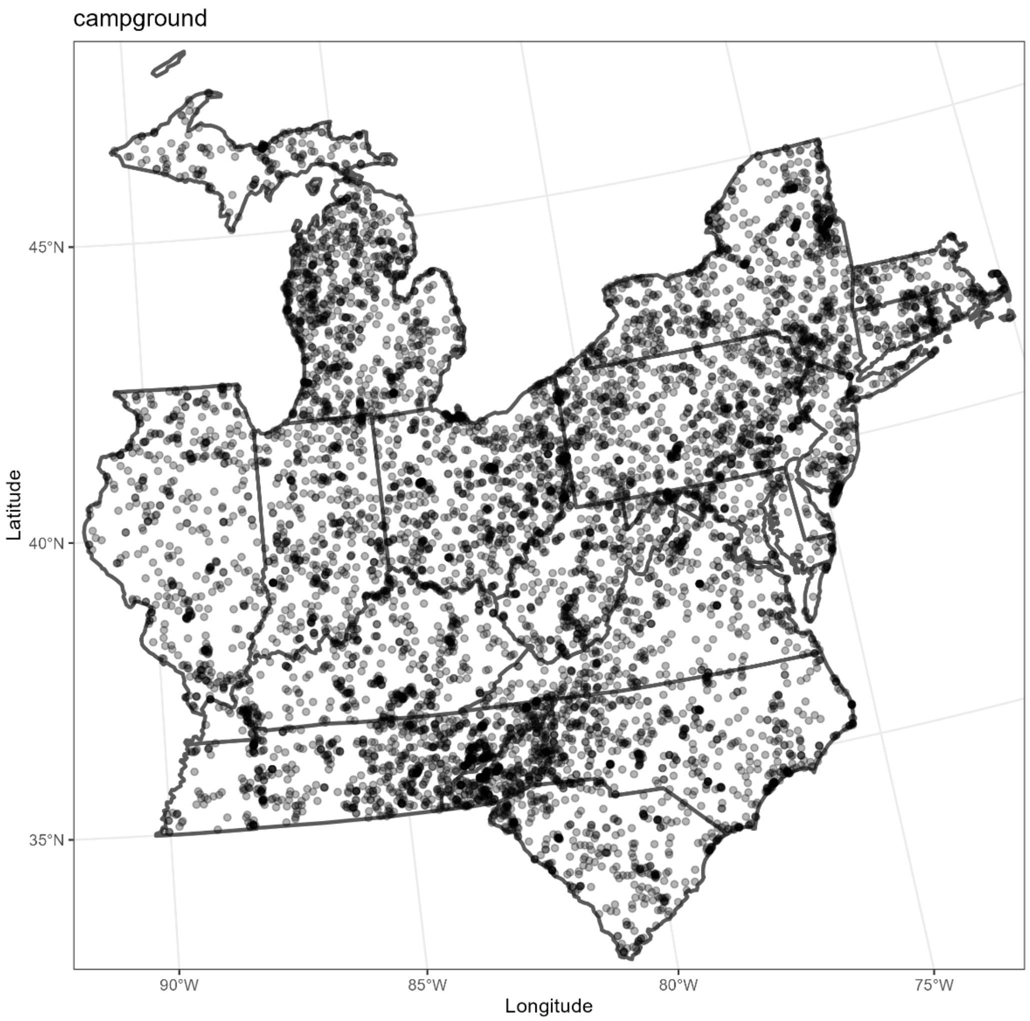


**Figure S1.3:** Campground locations within the study region in the Dataplor point of interest database used in the pathway analyses (Dataplor, 2022).

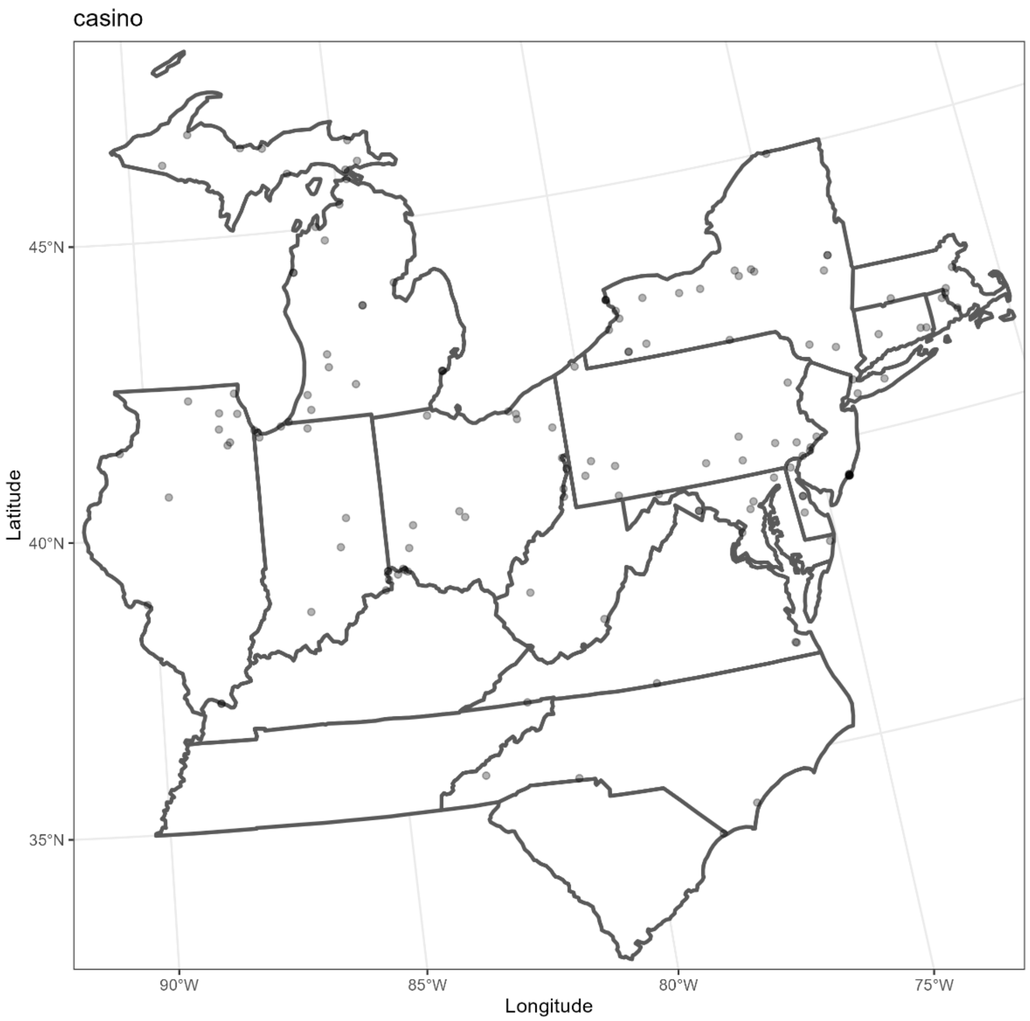


**Figure S1.4:** Casino locations within the study region in the Dataplor point of interest database used in the pathway analyses (Dataplor, 2022).

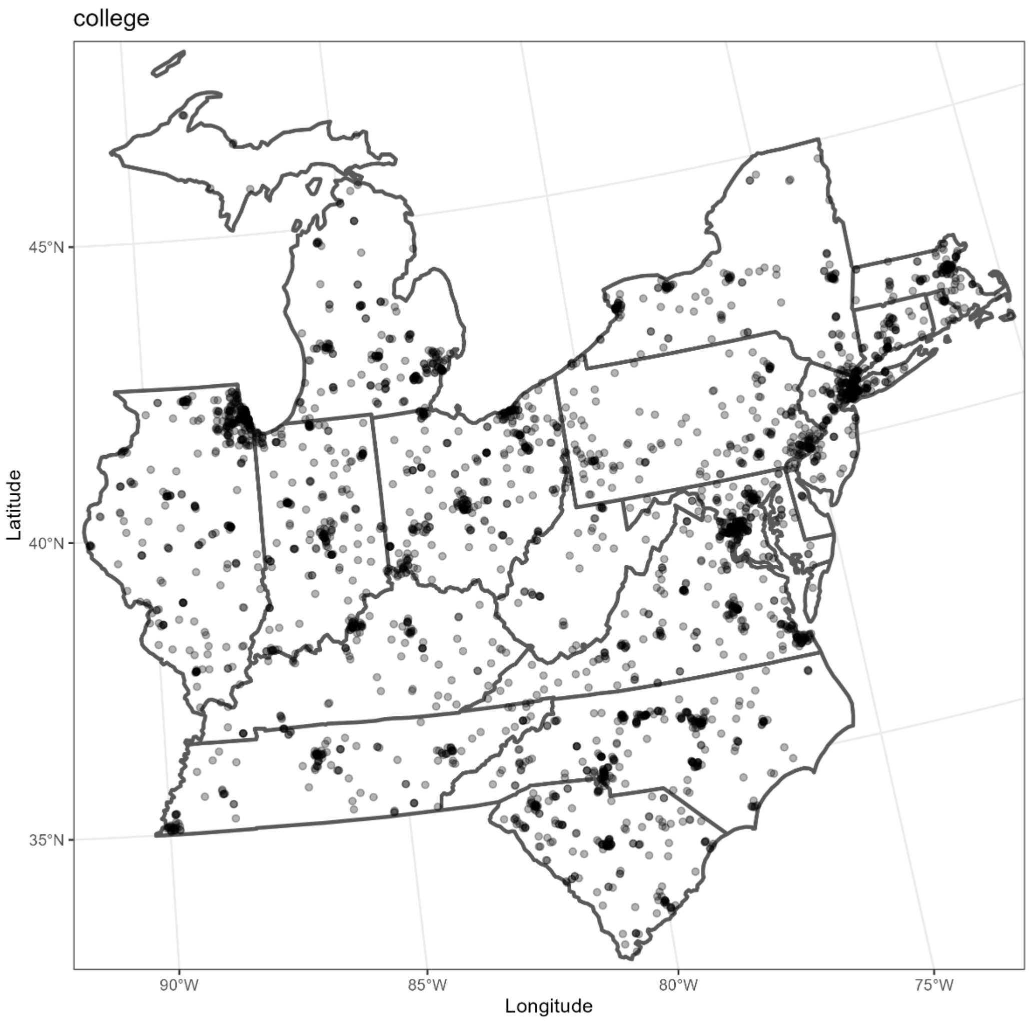


**Figure S1.5:** College locations within the study region in the Dataplor point of interest database used in the pathway analyses (Dataplor, 2022).

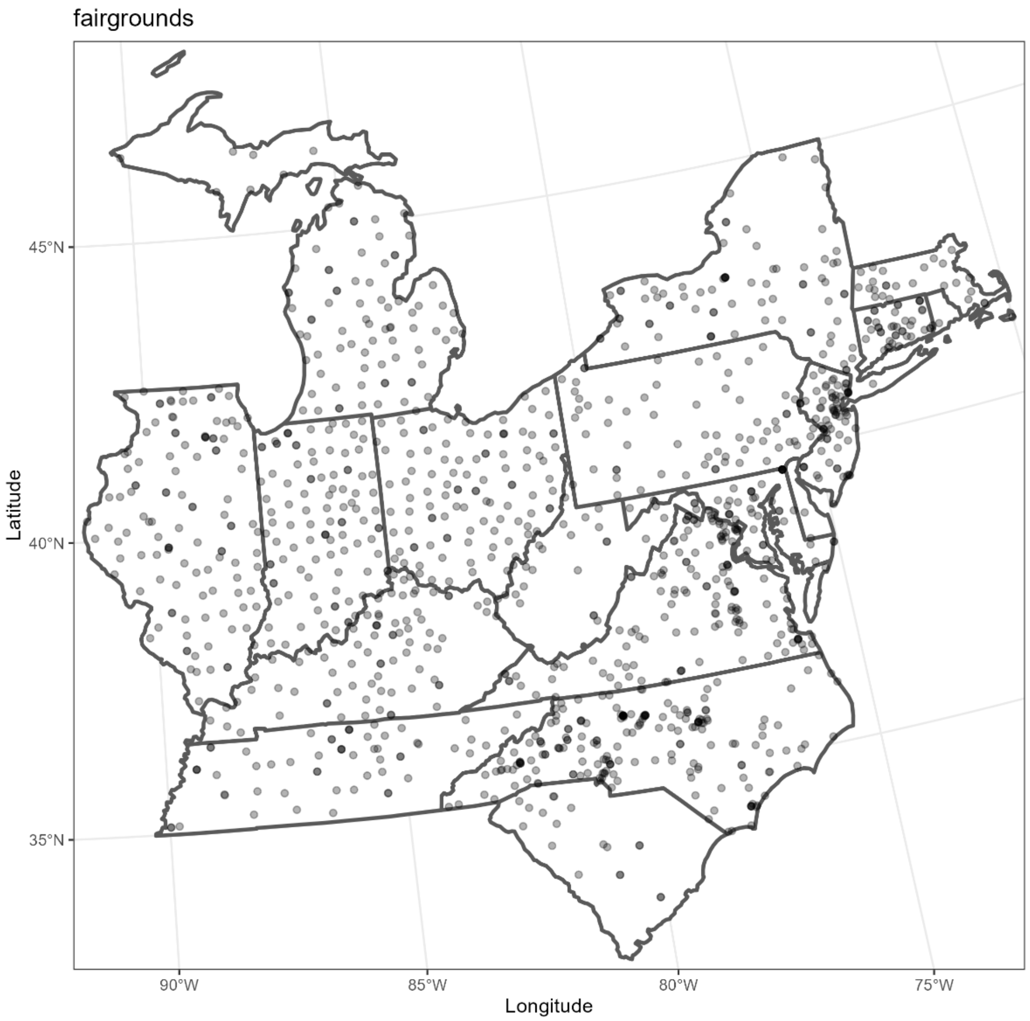


**Figure S1.6:** Fairgrounds locations within the study region in the Dataplor point of interest database used in the pathway analyses (Dataplor, 2022).

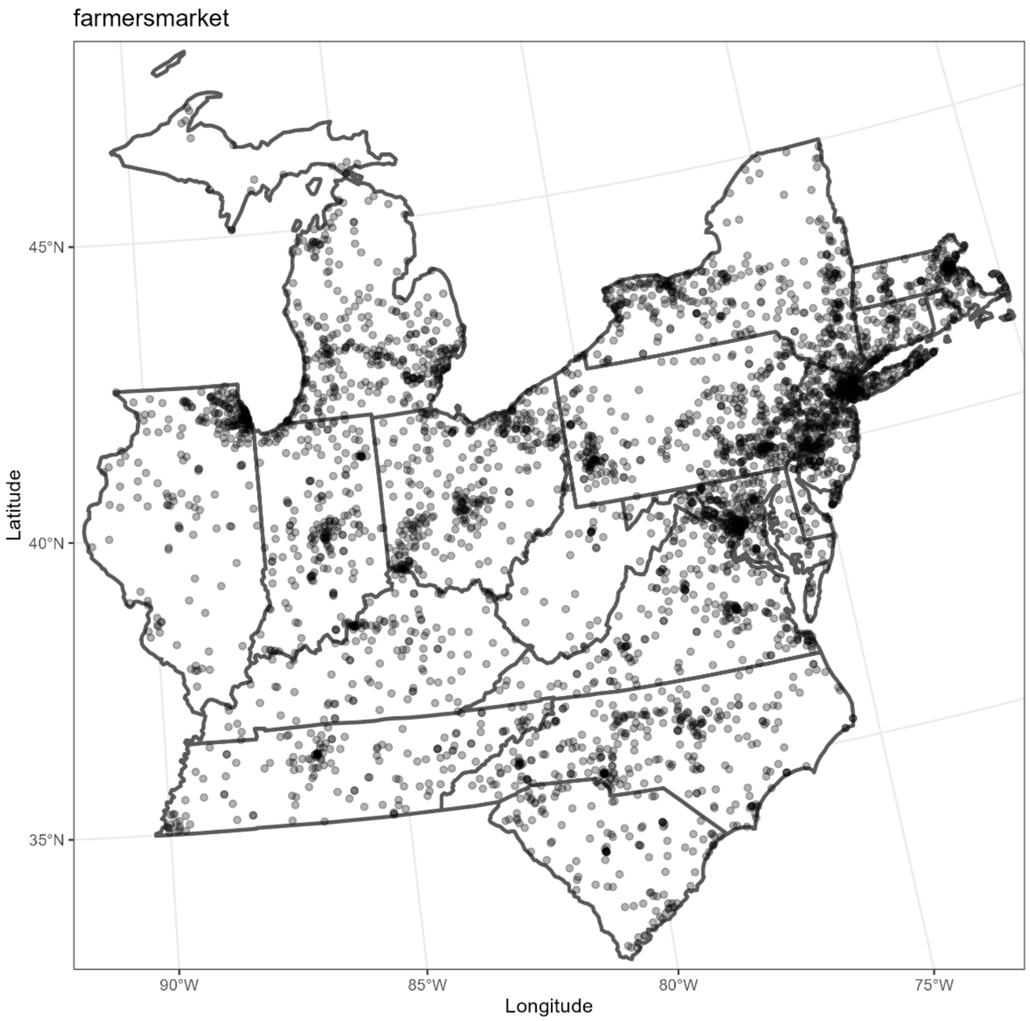


**Figure S1.7:** Farmers' market locations within the study region in the Dataplor point of interest database used in the pathway analyses (Dataplor, 2022).

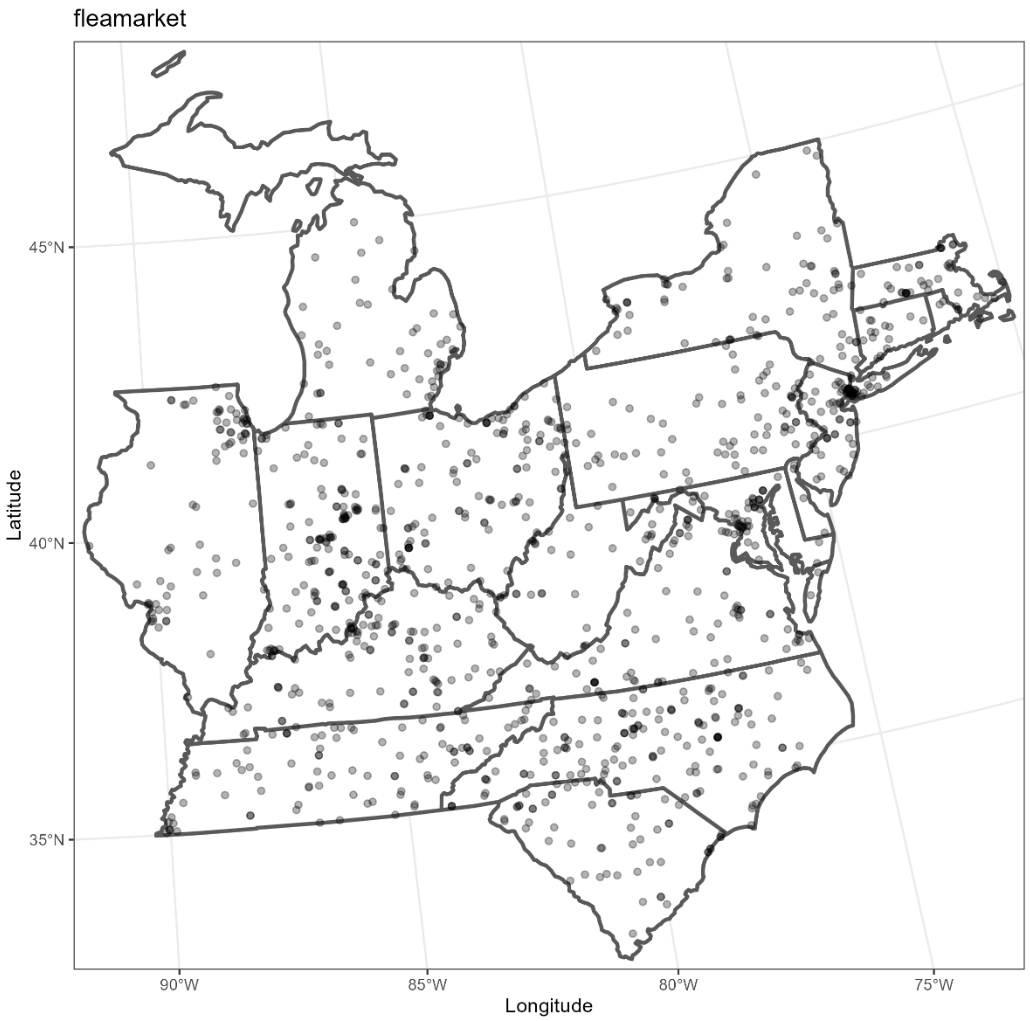


**Figure S1.8:** Flea market locations within the study region in the Dataplor point of interest database used in the pathway analyses (Dataplor, 2022).

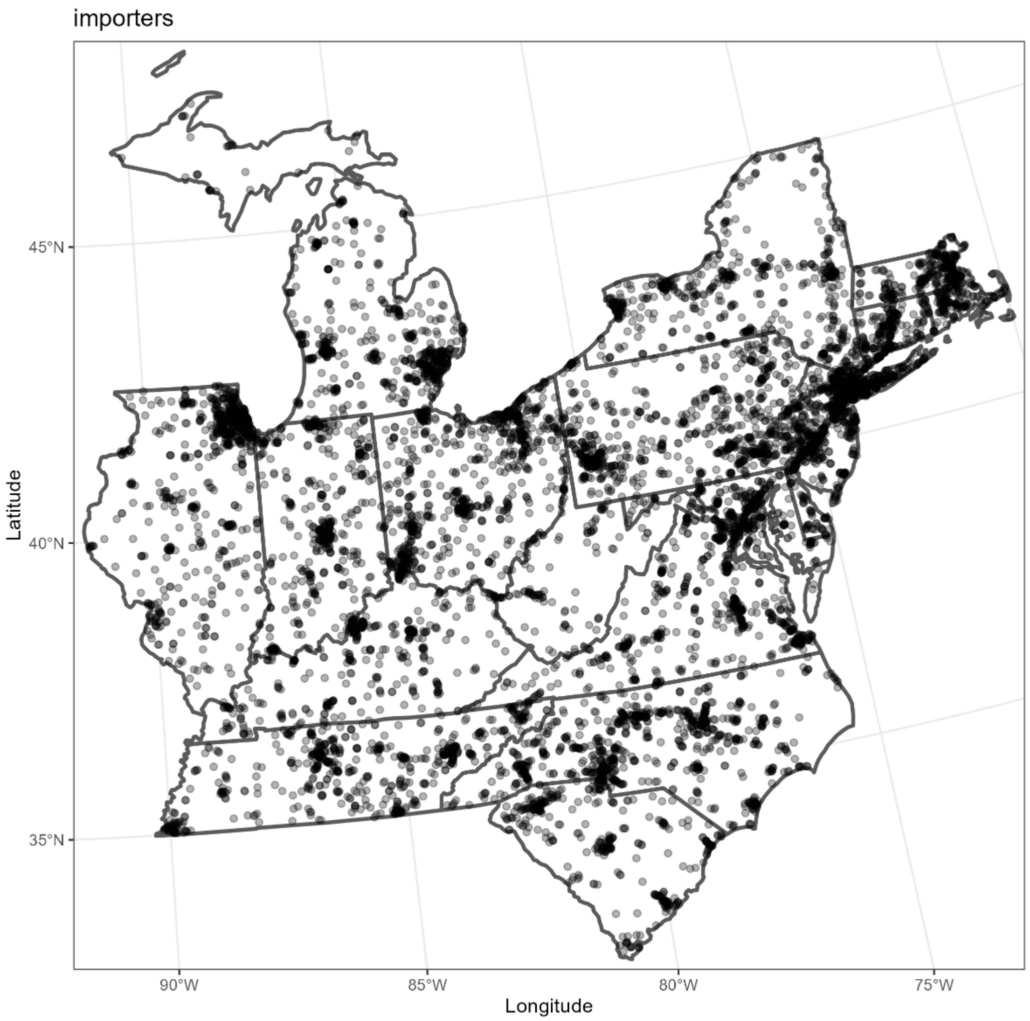


**Figure S1.9:** Importer locations within the study region in the Dataplor point of interest database used in the pathway analyses (Dataplor, 2022).

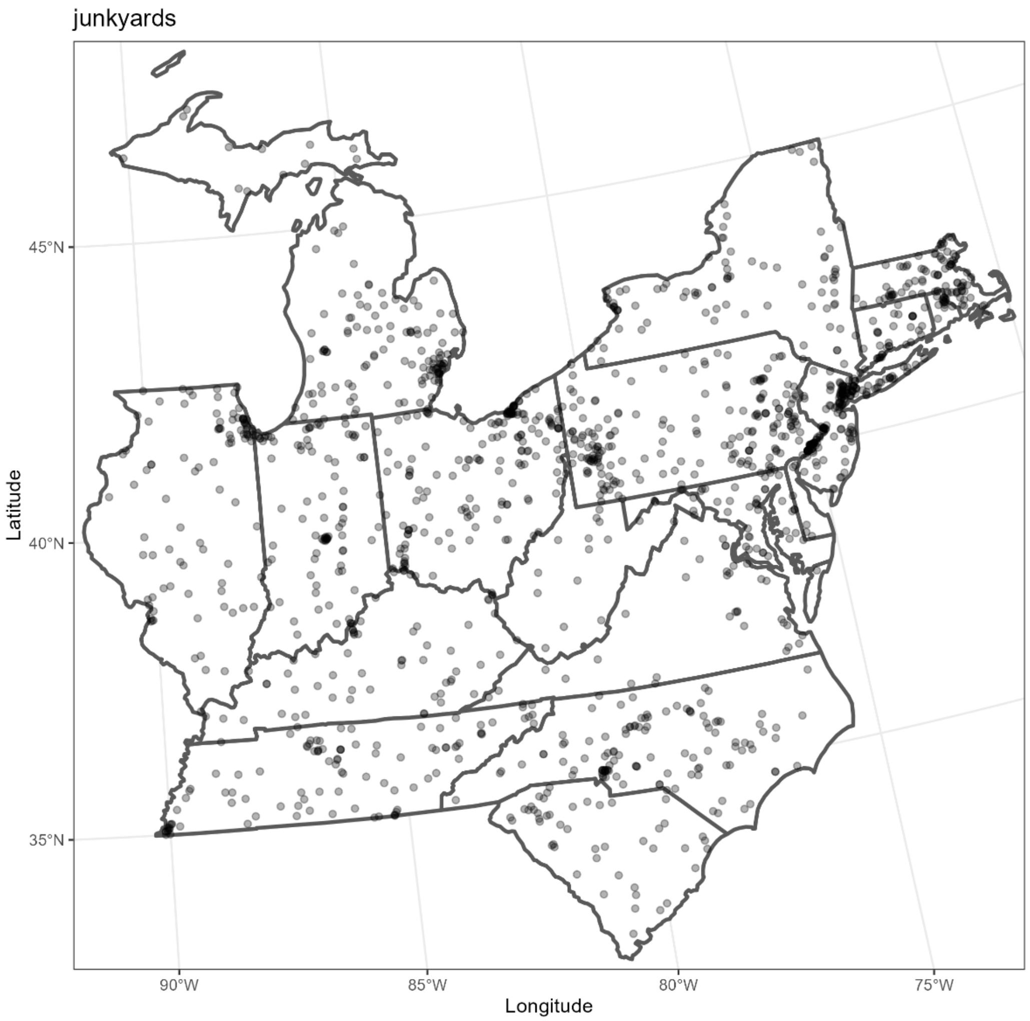


**Figure S1.10:** Junkyard locations within the study region in the Dataplor point of interest database used in the pathway analyses (Dataplor, 2022).

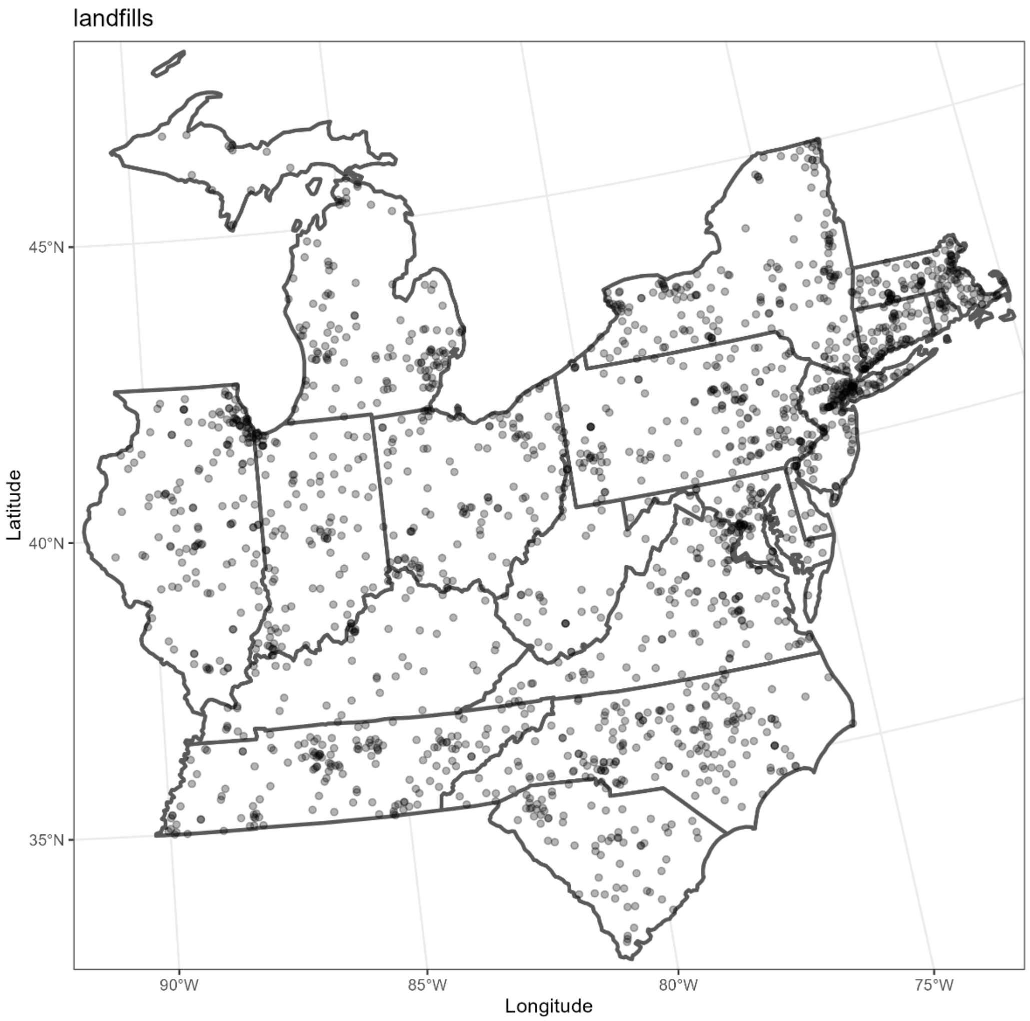


**Figure S1.11:** Landfill locations within the study region in the Dataplor point of interest database used in the pathway analyses (Dataplor, 2022).

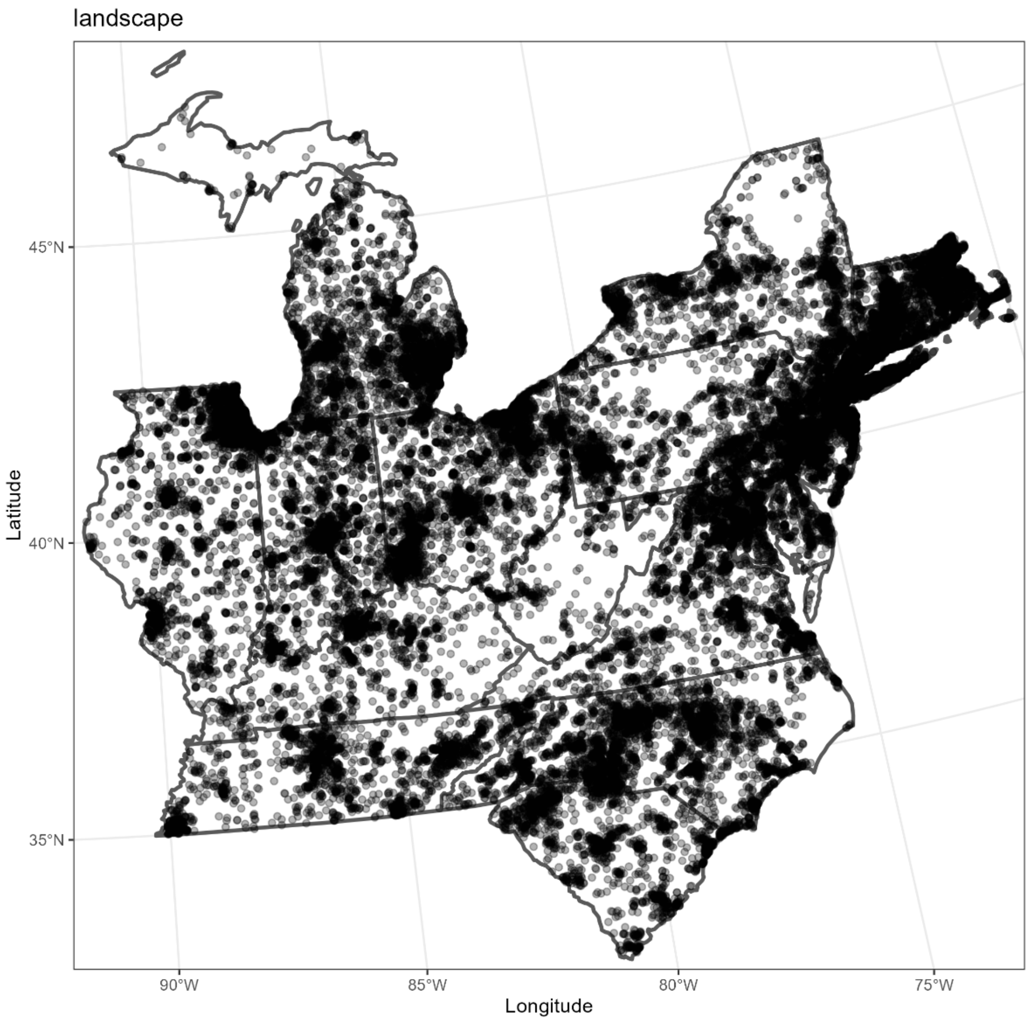


**Figure S1.12:** Landscaping business locations within the study region in the Dataplor point of interest database used in the pathway analyses (Dataplor, 2022).

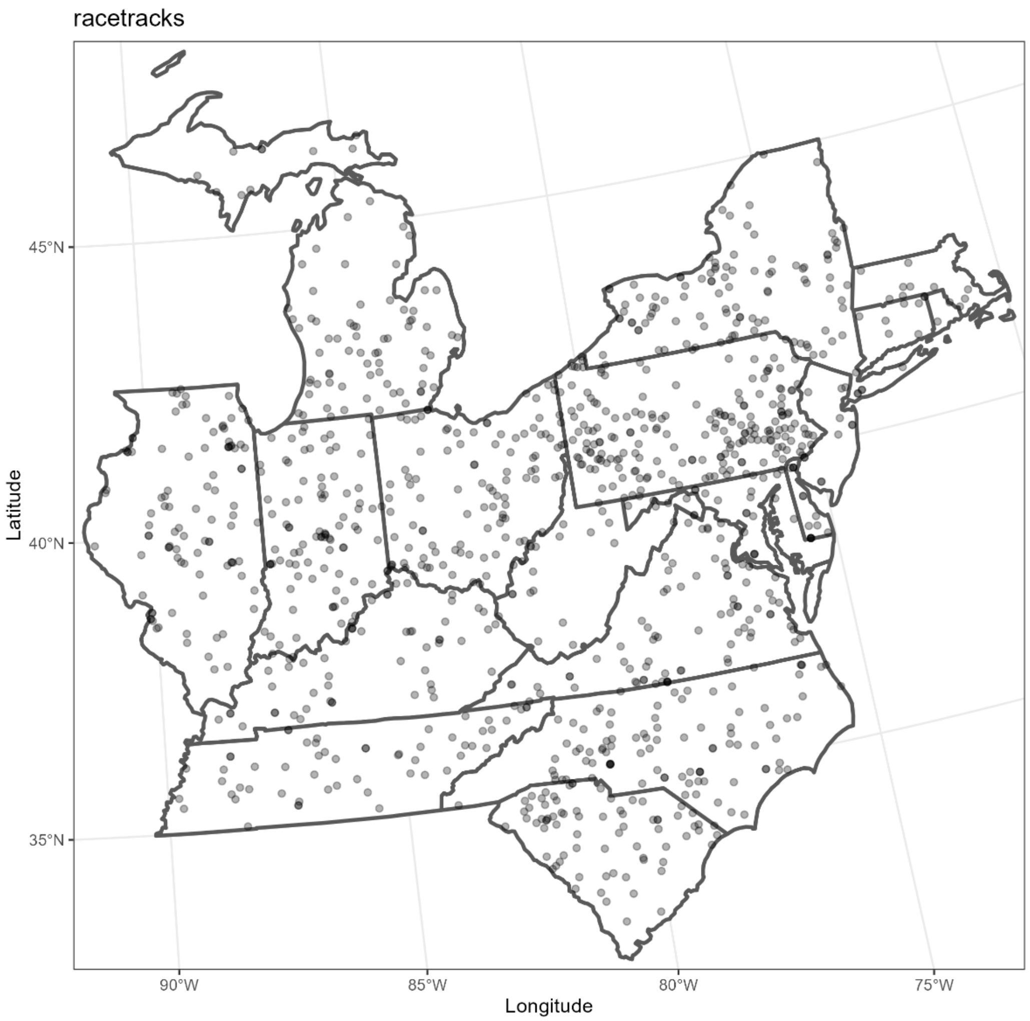


**Figure S1.13:** Race track locations within the study region in the Dataplor point of interest database used in the pathway analyses (Dataplor, 2022).

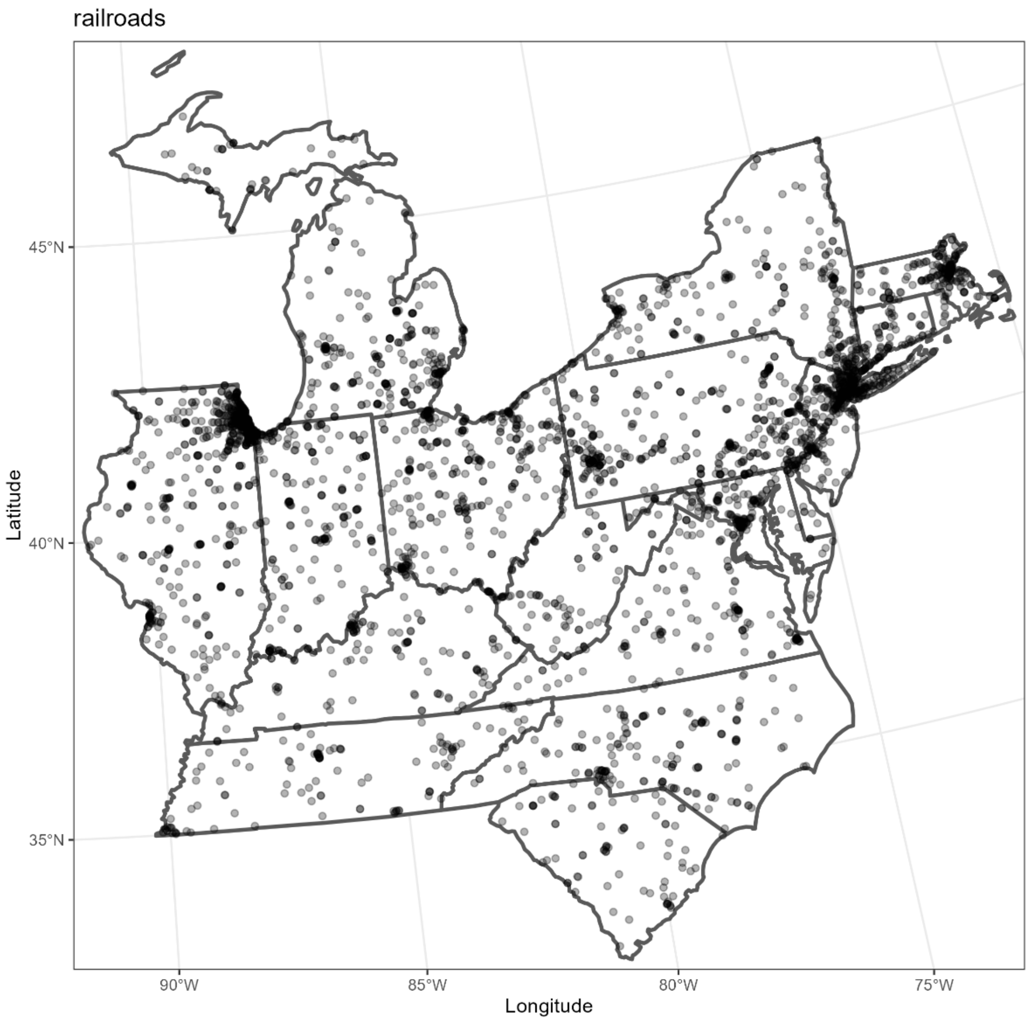


**Figure S1.14:** Railroad station locations within the study region in the Dataplor point of interest database used in the pathway analyses (Dataplor, 2022).

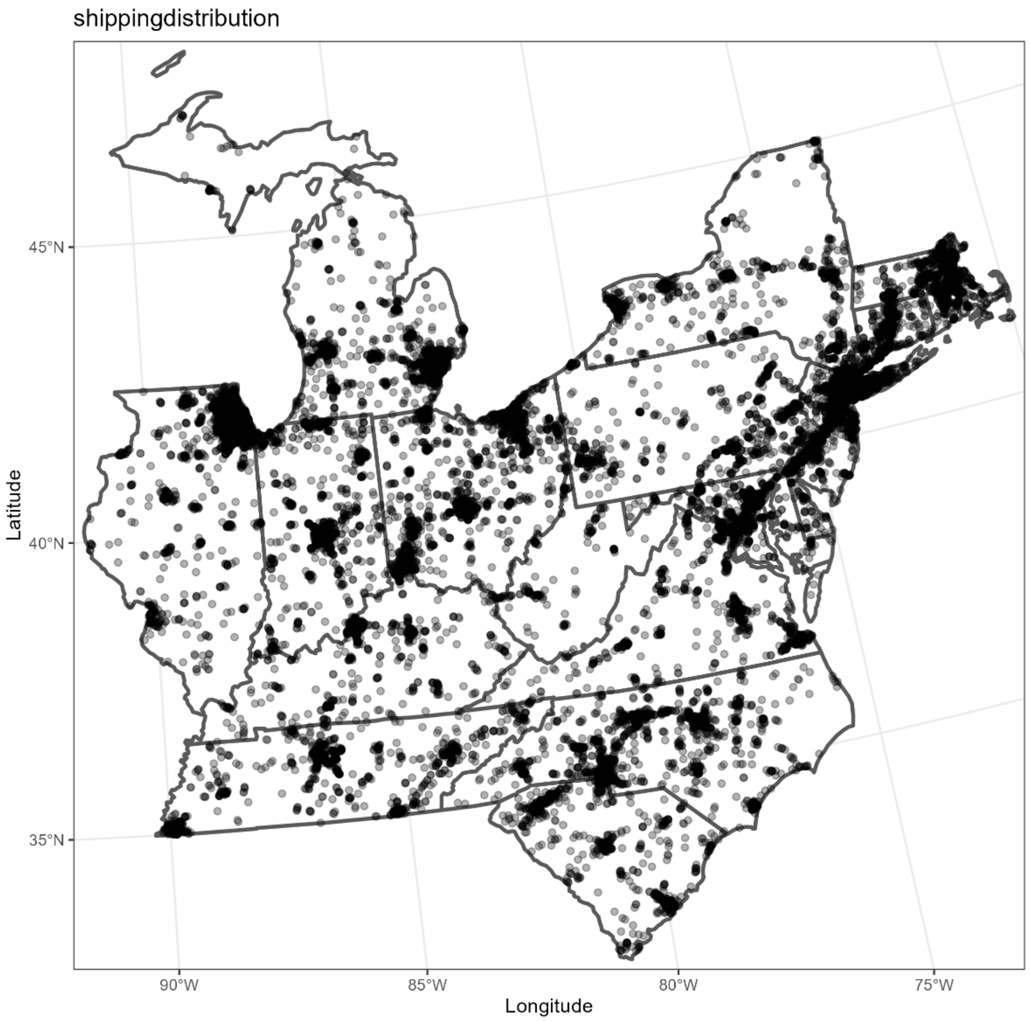


**Figure S1.15:** Shipping and distribution center locations within the study region in the Dataplor point of interest database used in the pathway analyses (Dataplor, 2022).

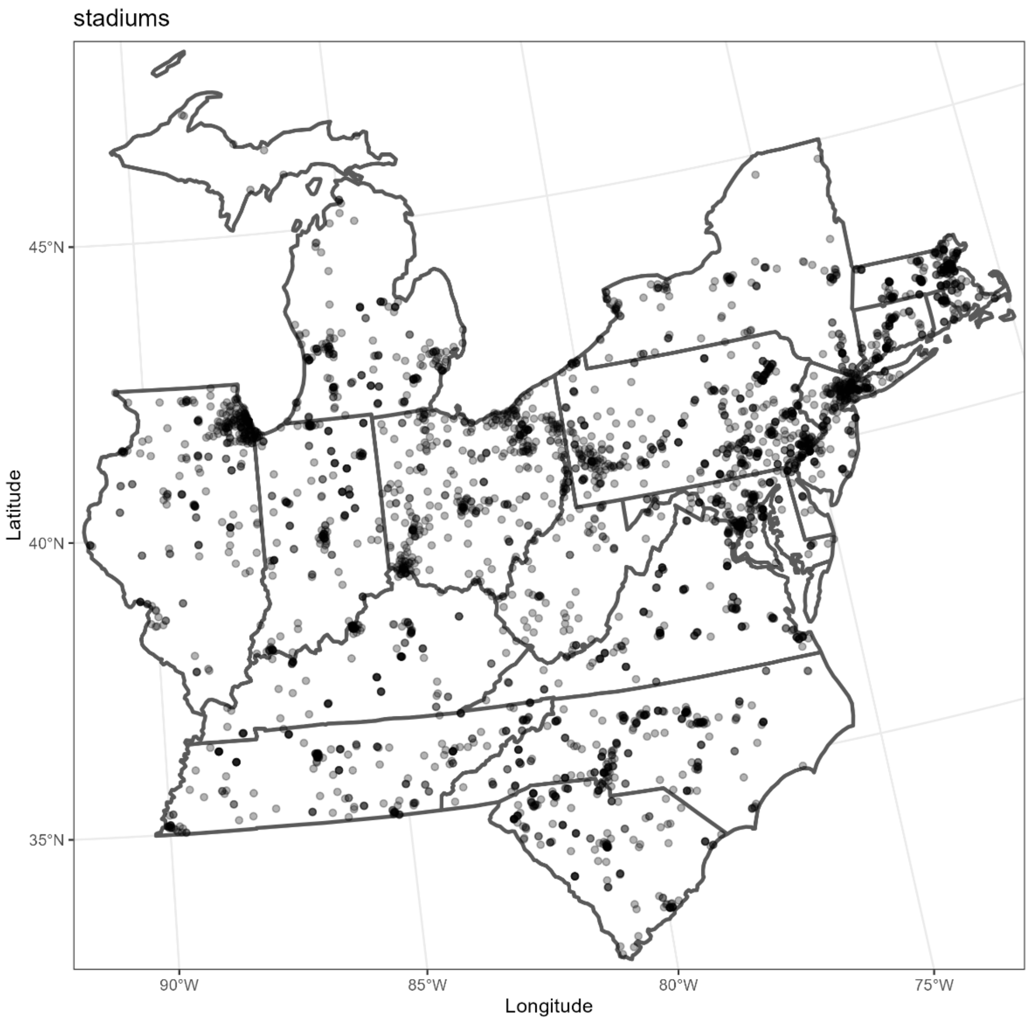


**Figure S1.16:** Stadium locations within the study region in the Dataplor point of interest database used in the pathway analyses (Dataplor, 2022).

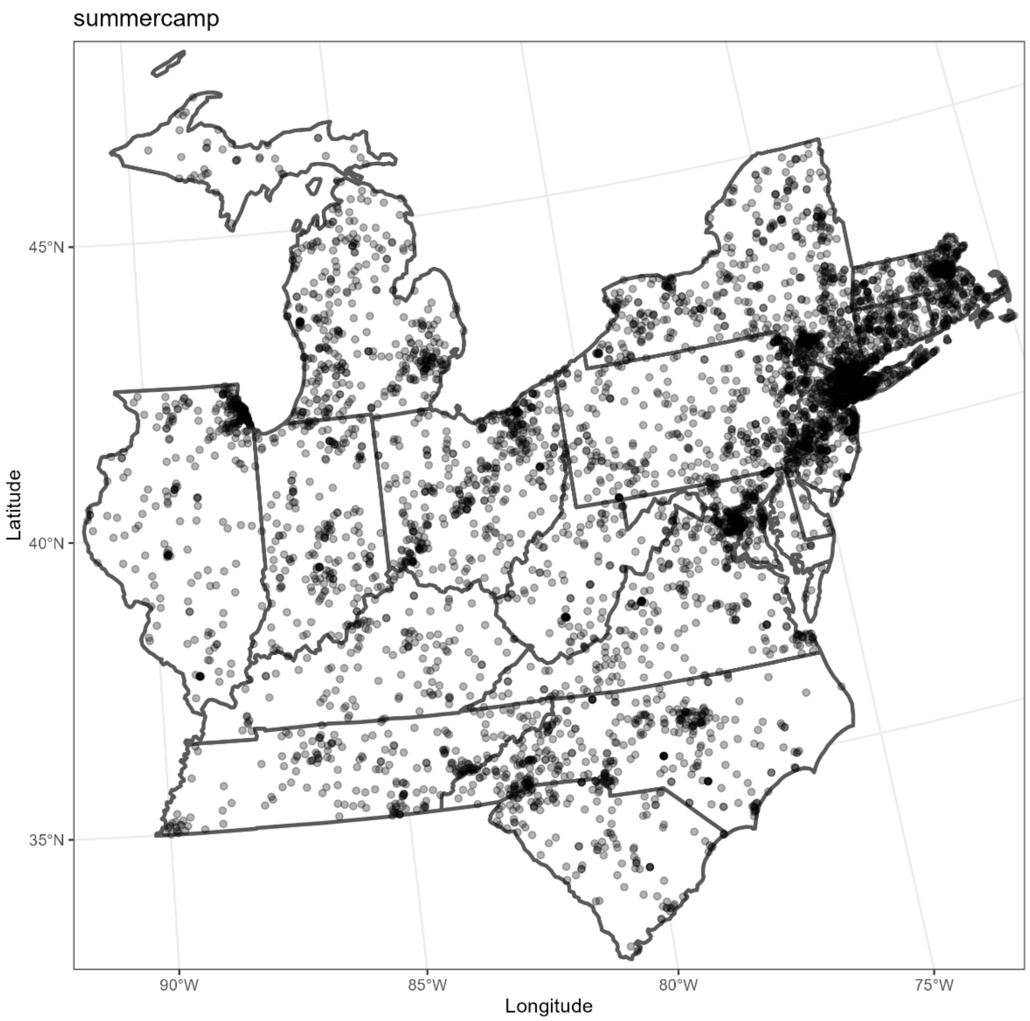


**Figure S1.17:** Summer camp locations within the study region in the Dataplor point of interest database used in the pathway analyses (Dataplor, 2022).

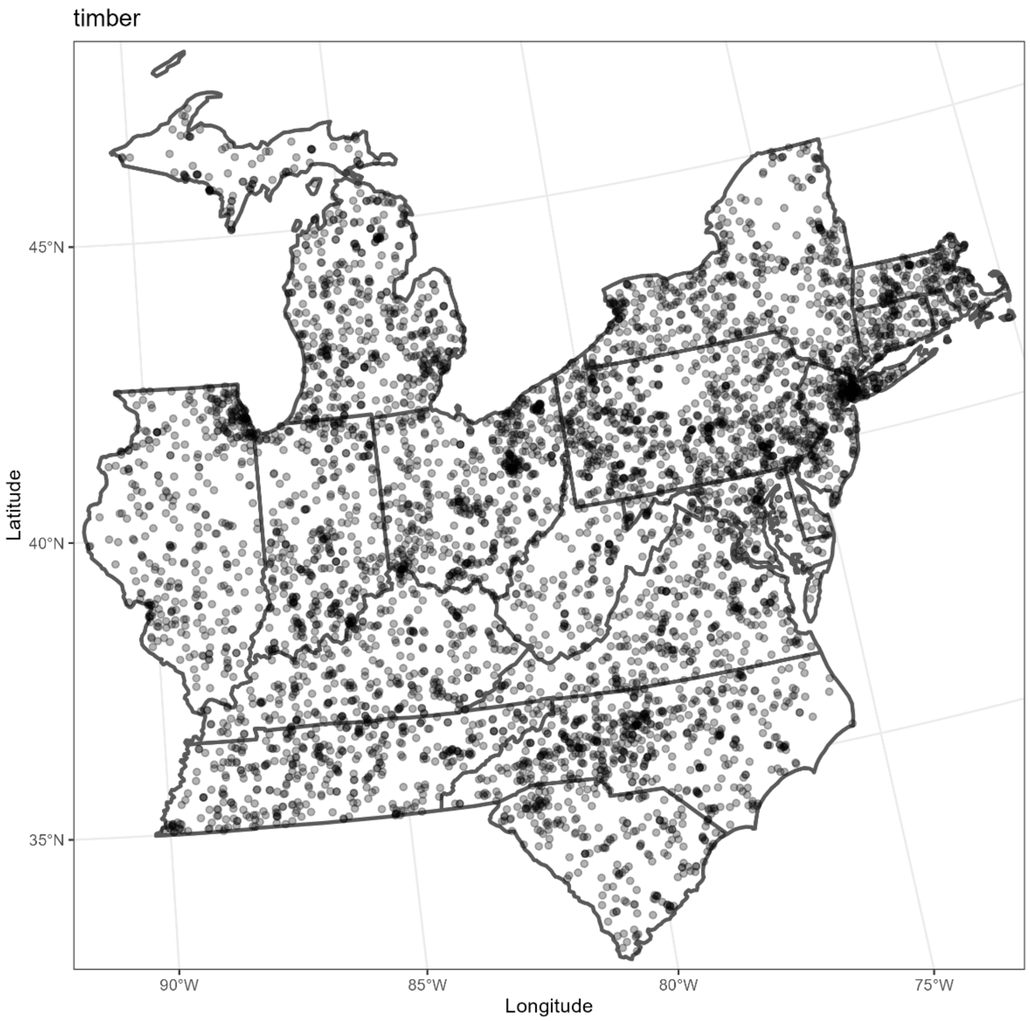


**Figure S1.18:** Timber processing facility locations within the study region in the Dataplor point of interest database used in the pathway analyses (Dataplor, 2022).

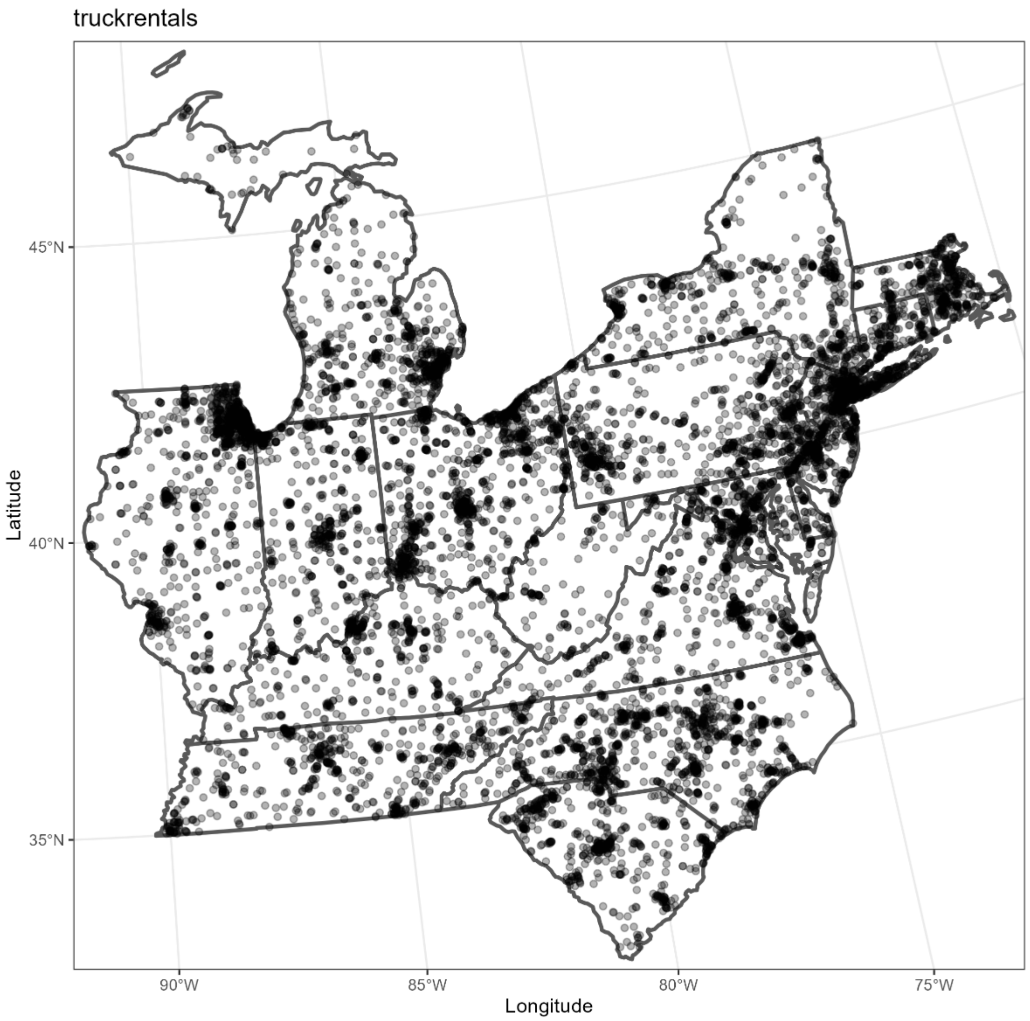


**Figure S1.19:** Truck rental locations within the study region in the Dataplor point of interest database used in the pathway analyses (Dataplor, 2022).

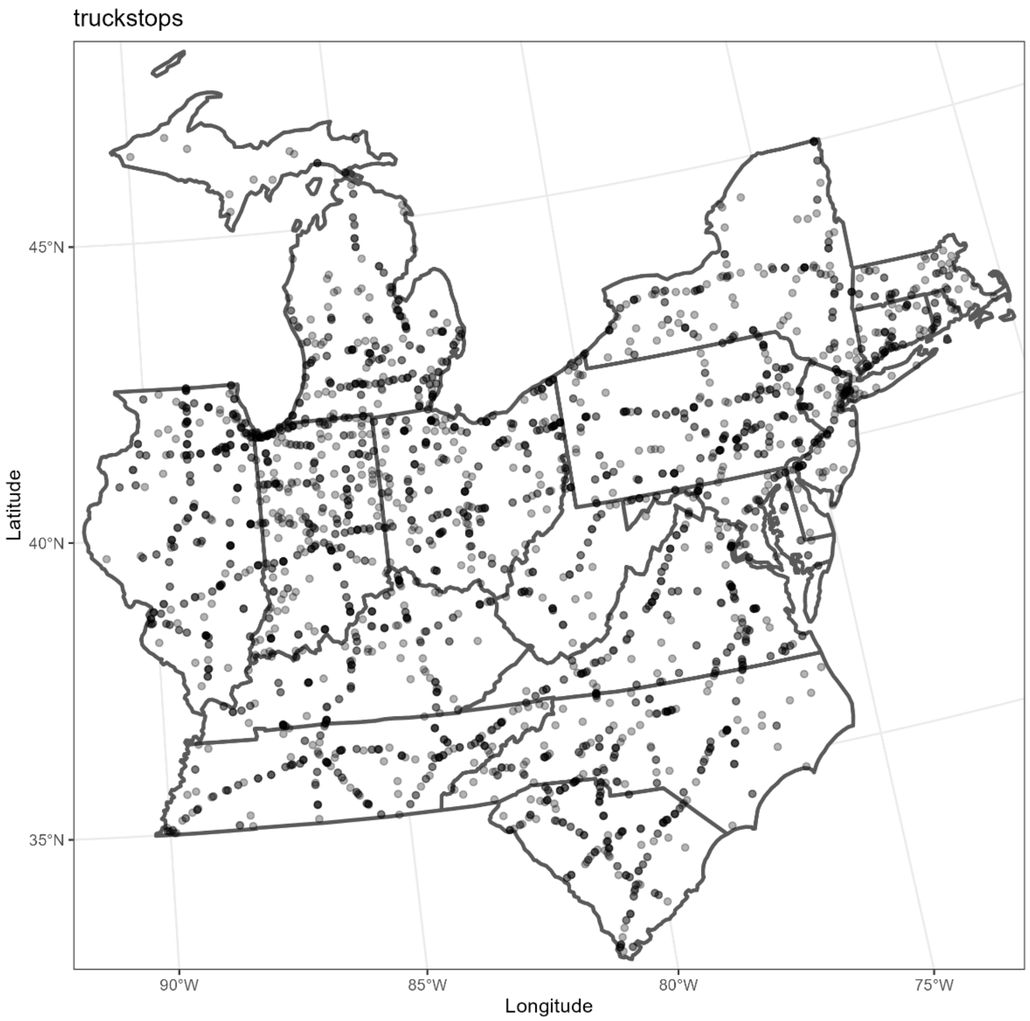


**Figure S1.20:** Truck stop locations within the study region in the Dataplor point of interest database used in the pathway analyses (Dataplor, 2022).

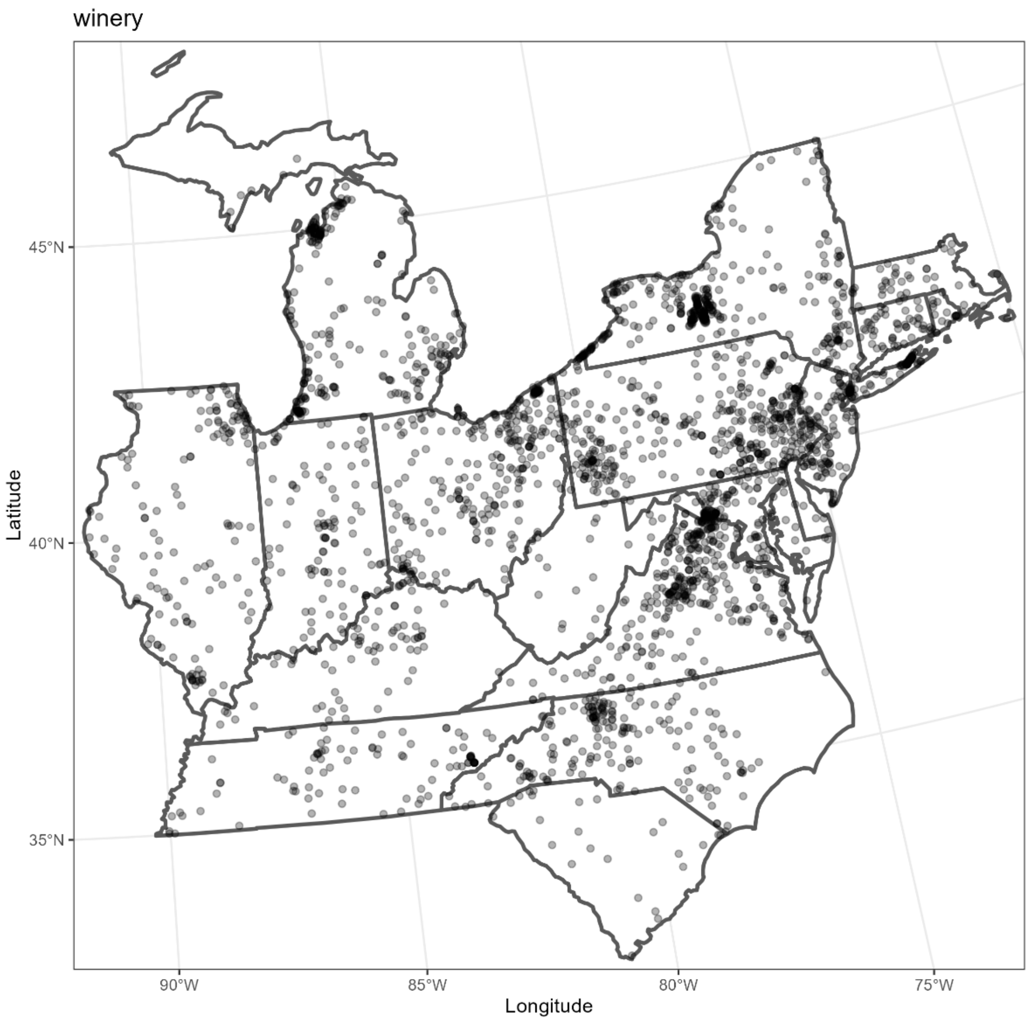


**Figure S1.21:** Winery locations within the study region in the Dataplor point of interest database used in the pathway analyses (Dataplor, 2022).

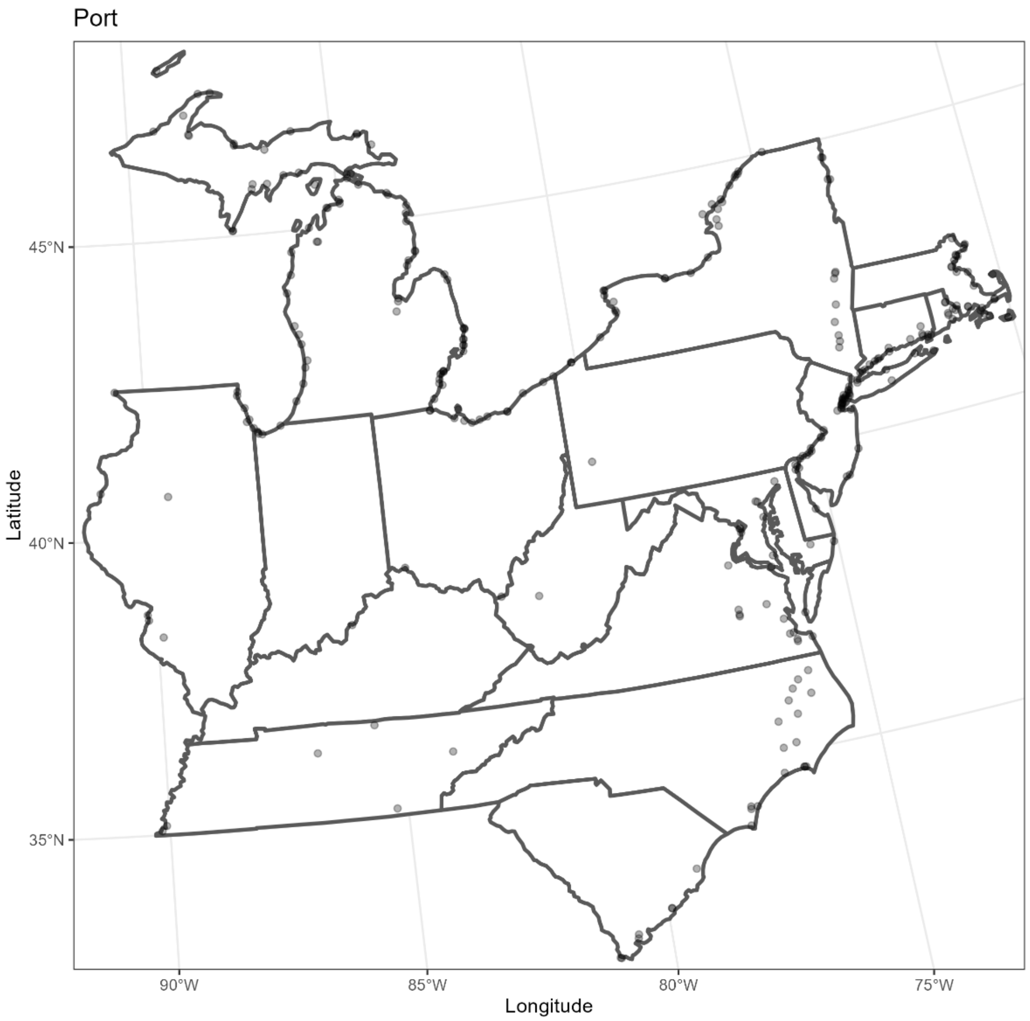


**Figure S1.22:** Port locations within the study region. Locations are from the Gulf Coast Prairie Landscape Conservation Cooperative (LCC Network Data Steward & Gulf Coast Prairie LCC Data Manager, 2014) and World Port Index (National Geospatial-Intelligence Agency, 2020)

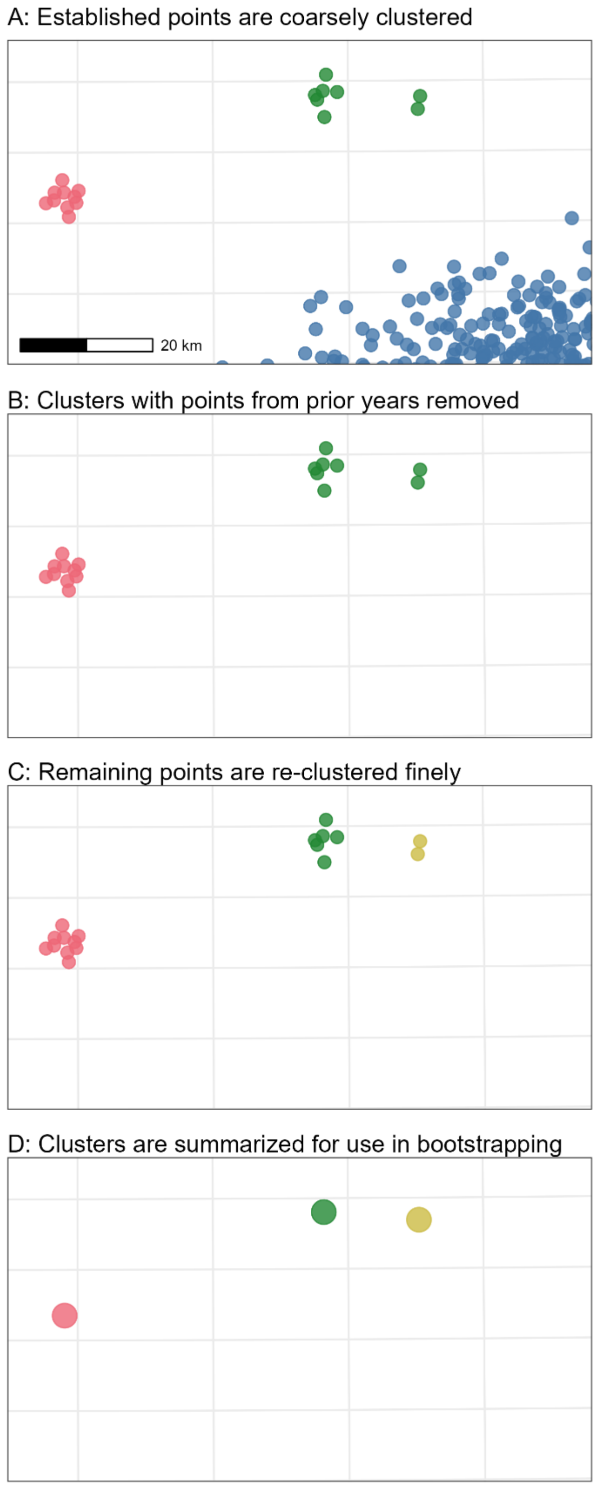


**Figure S1.23**: The DBSCAN clustering algorithm was used to identify outlying nascent *L. delicatula* populations resulting from jump dispersal. For each year of the invasion (2014-2024, simulated data shown for clarity), records of established *L. delicatula* populations in that year or any prior year made by government agencies were first coarsely clustered (shown by point colors) with search radius 20 km (A). Clusters holding previous year's records (blue points) were removed, because these did not represent newly discovered nascent populations (B). Then, retained points were re-clustered with a finer search radius of 5 km to discern spatially near but distinct nascent populations (C). Finally, for use in bootstrapping analysis, spatial variables (including proximity to points of interest and length of nearby transportation infrastructure) were calculated for each point within a cluster and then averaged (D).

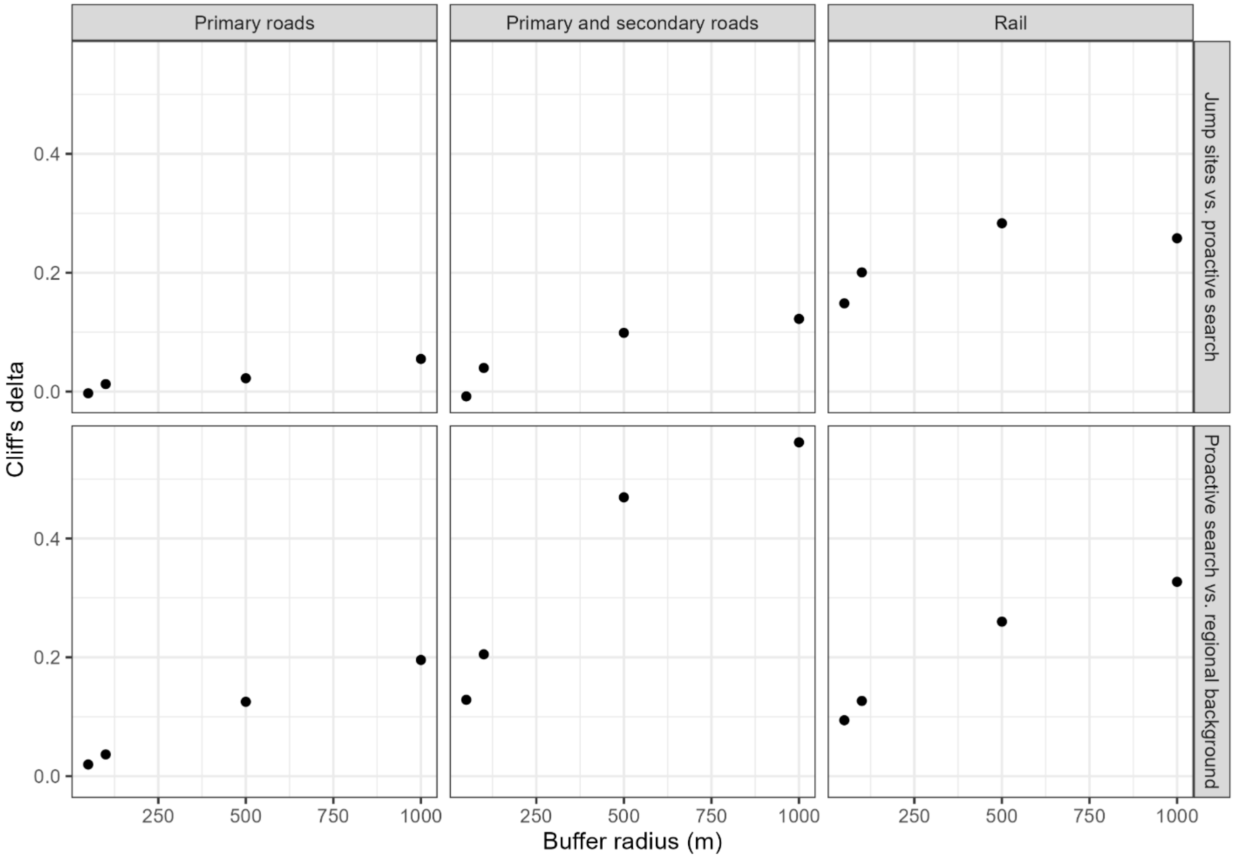


**Figure S1.24:** A 500 m buffer around study sites resulted in the strongest apparent difference between jump locations and proactive search sites in the amount of nearby transport infrastructure. Sensitivity analysis for the effect of buffer radius around points on the estimated effect size (Cliff's delta) describing the difference in road density (left and center) or rail density (right) around proactive search points and *L. delicatula* jump dispersal sites (top row) and between proactive search points and regional background (bottom row).

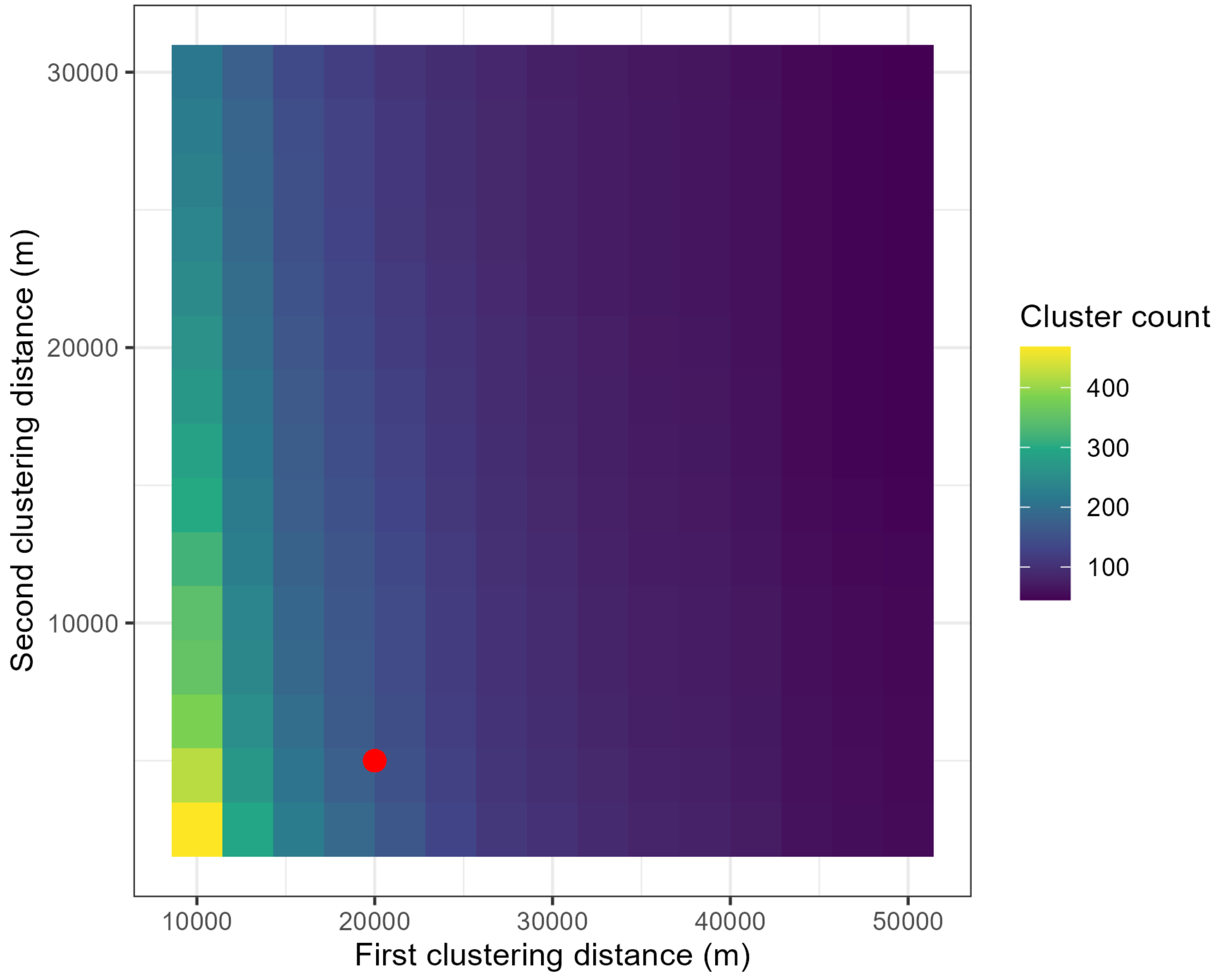


**Figure S1.25:** The selected clustering parameters (20,000 m first clustering distance, 5000 m second clustering distance, shown with a red point) resulted in an intermediate number of clusters compared to the high numbers possible with fine-scale clustering or the minimum number with coarse clustering. The selected parameterization was based on prior knowledge of *L. delicatula* dispersal capacity and nascent population sizes (Belouard et al., 2025; Keller et al., 2026). Evaluated values for the first search radius used in initial clustering ranged from 10,000 m to 50,000 m (along the x axis), while evaluated values for the second search radius used to finely distinguish individual clusters ranged from 2,500 m to 30,000 m along the y axis.

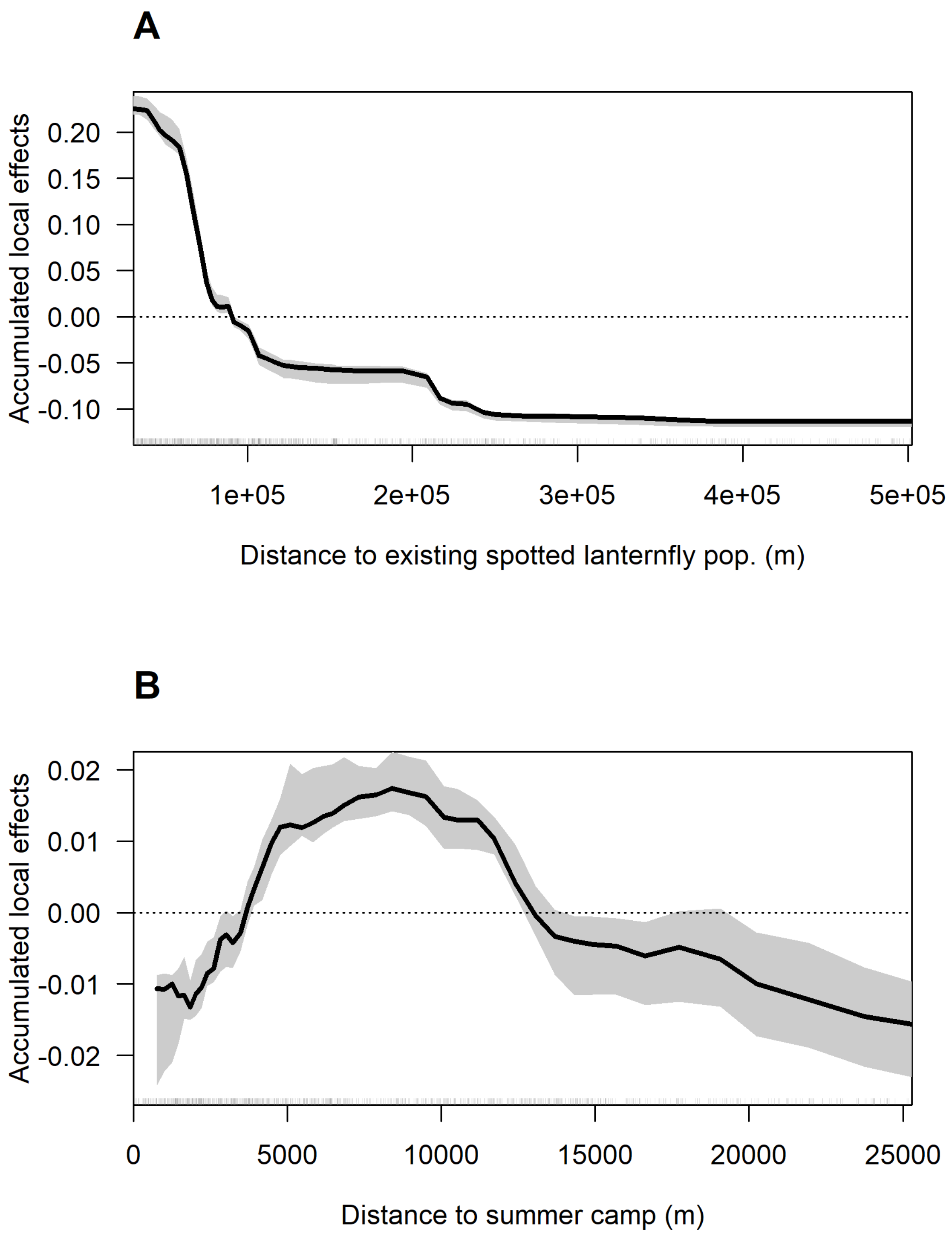
A
Accumulated local effects
0.20
0.15
0.10
0.05
0.00
-0.05
-0.10
1e+05
2e+05
3e+05
4e+05
5e+05
Distance to existing spotted lanternfly pop. (m)
B
Accumulated local effects
0.02
0.01
0.00
-0.01
-0.02
0
5000
10000
15000
20000
25000
Distance to summer camp (m)

**C**

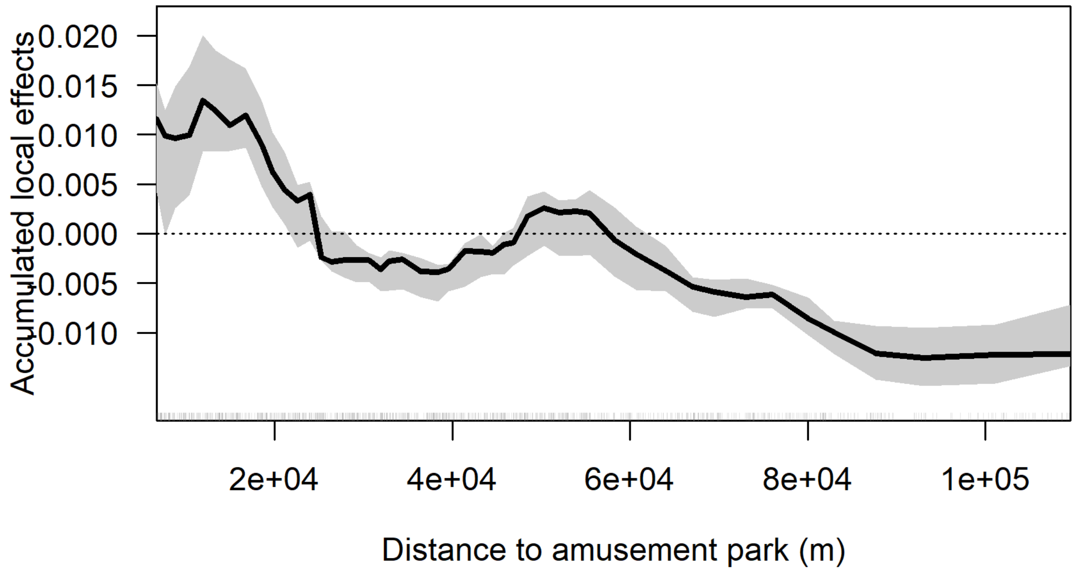

Accumulated local effects
0.020
0.015
0.010
0.005
0.000
-0.005
-0.010
2e+04
4e+04
6e+04
8e+04
1e+05
Distance to amusement park (m)


**D**

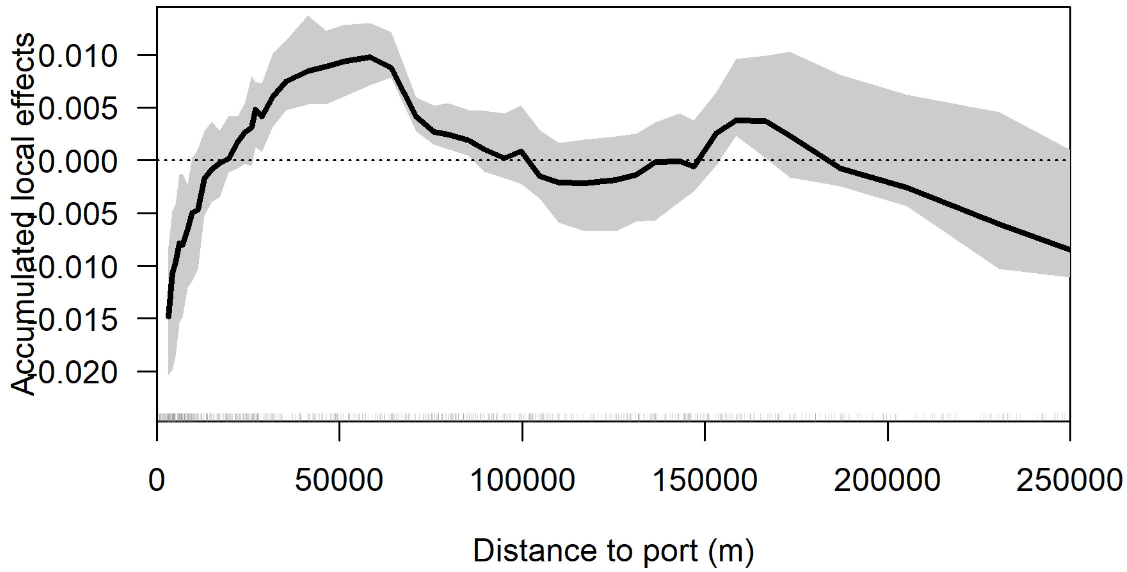

Accumulated local effects
0.010
0.005
0.000
-0.005
-0.010
-0.015
-0.020
0
50000
100000
150000
200000
250000
Distance to port (m)

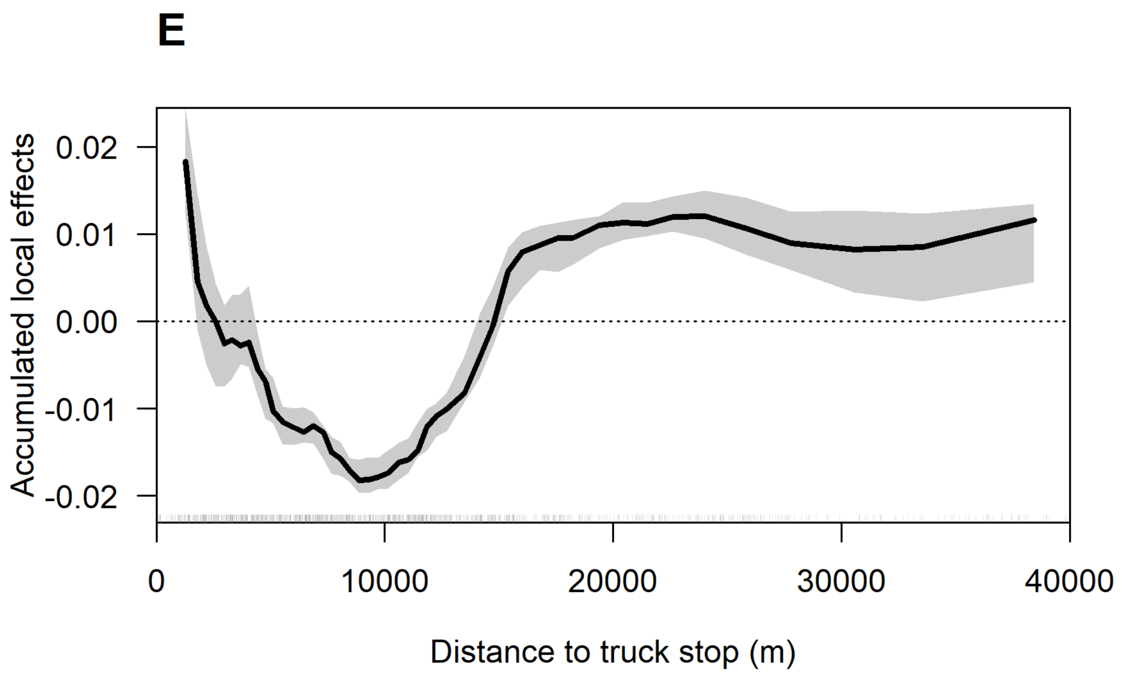
E
Accumulated local effects
0.02
0.01
0.00
-0.01
-0.02
0
10000
20000
30000
40000
Distance to truck stop (m)

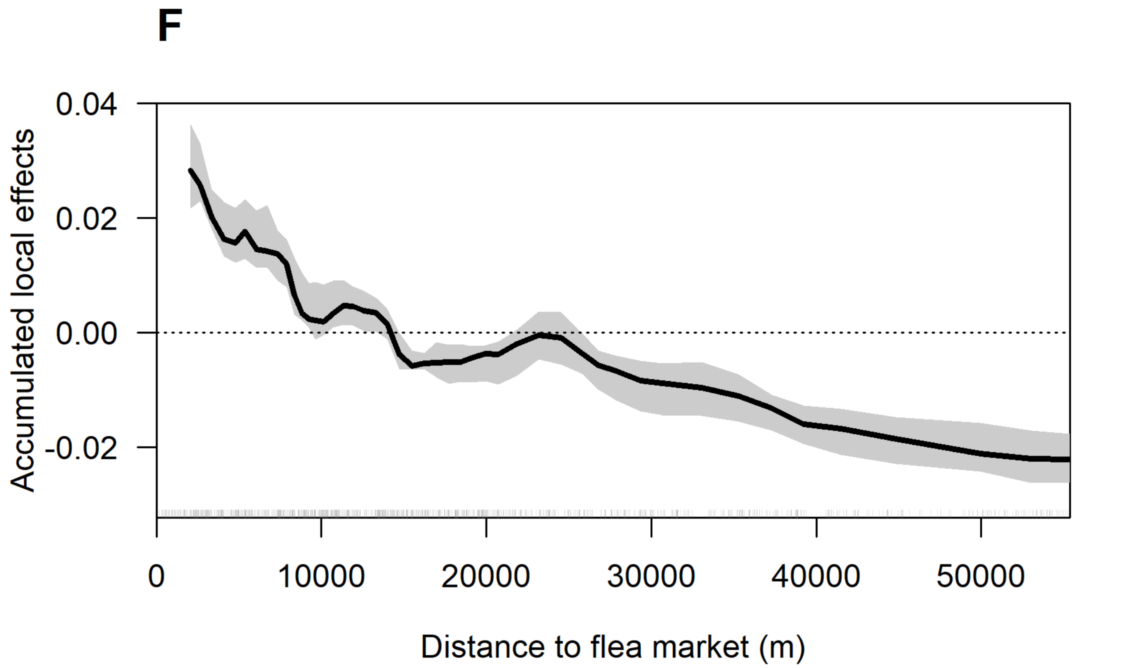
F
Accumulated local effects
0.04
0.02
0.00
-0.02
0
10000
20000
30000
40000
50000
Distance to flea market (m)

G

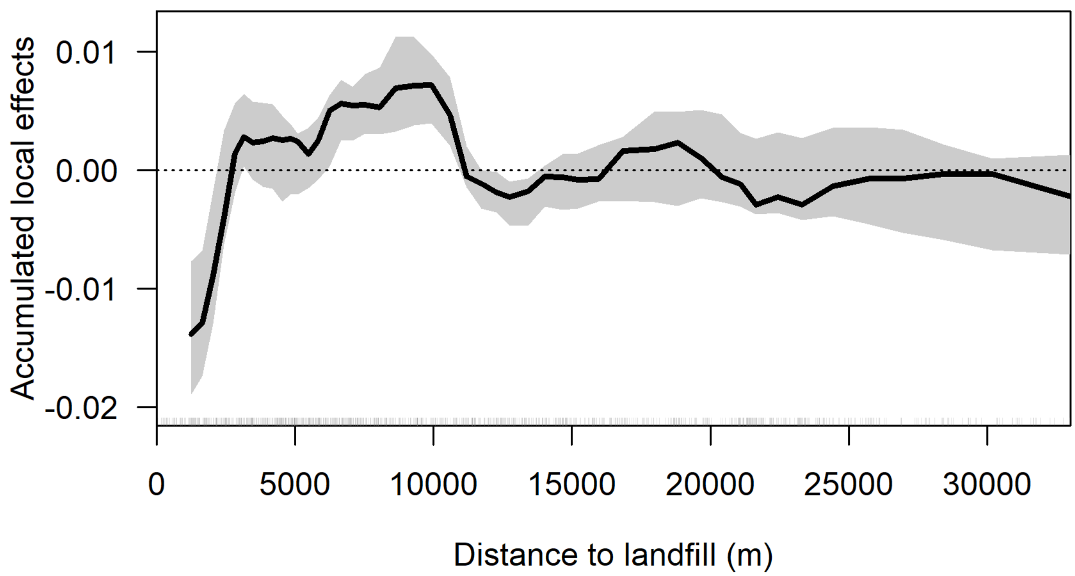

Accumulated local effects
0.01
0.00
-0.01
-0.02
0
5000
10000
15000
20000
25000
30000
Distance to landfill (m)

H

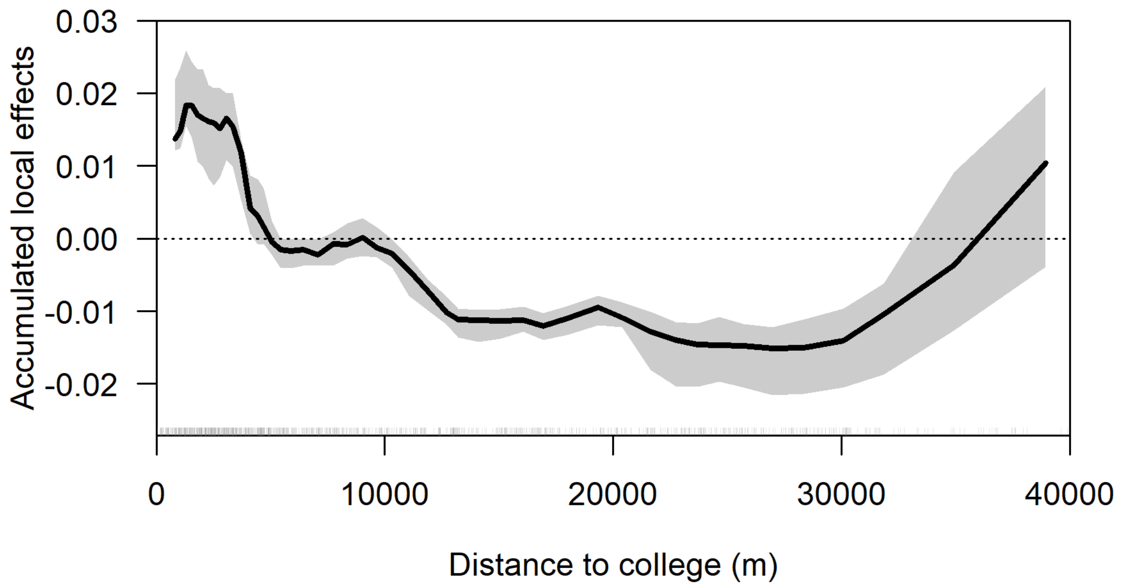
Accumulated local effects
0.03
0.02
0.01
0.00
-0.01
-0.02
0
10000
20000
30000
40000
Distance to college (m)

I

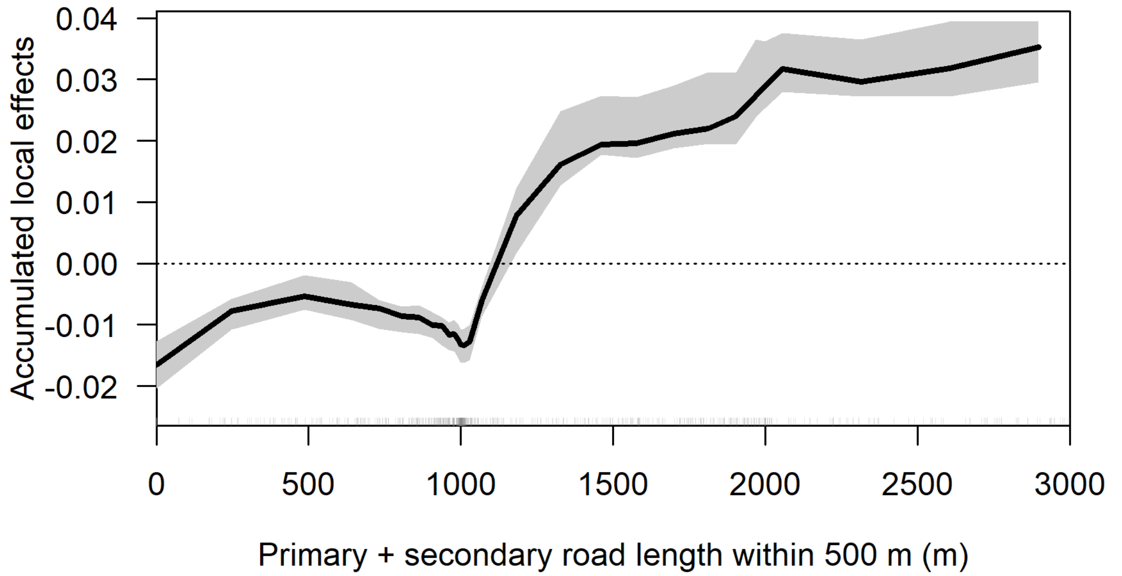
Accumulated local effects
0.04
0.03
0.02
0.01
0.00
-0.01
-0.02
0
500
1000
1500
2000
2500
3000
Primary + secondary road length within 500 m (m)

J

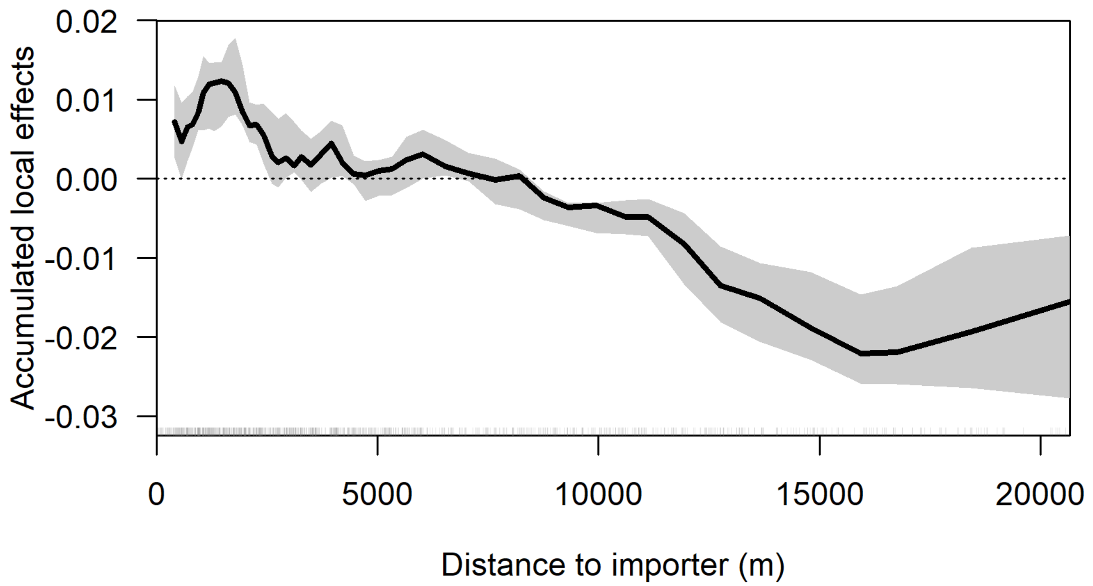
0.02
0.01
0.00
-0.01
-0.02
-0.03
Accumulated local effects
0
5000
10000
15000
20000
Distance to importer (m)

K

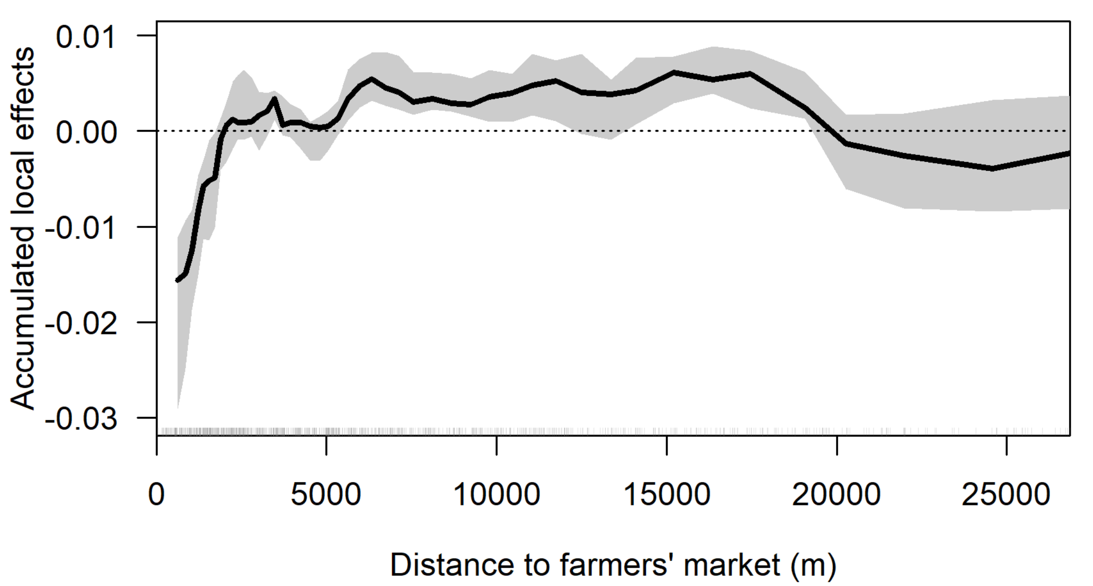
0.01
0.00
-0.01
-0.02
-0.03
Accumulated local effects
0
5000
10000
15000
20000
25000
Distance to farmers' market (m)

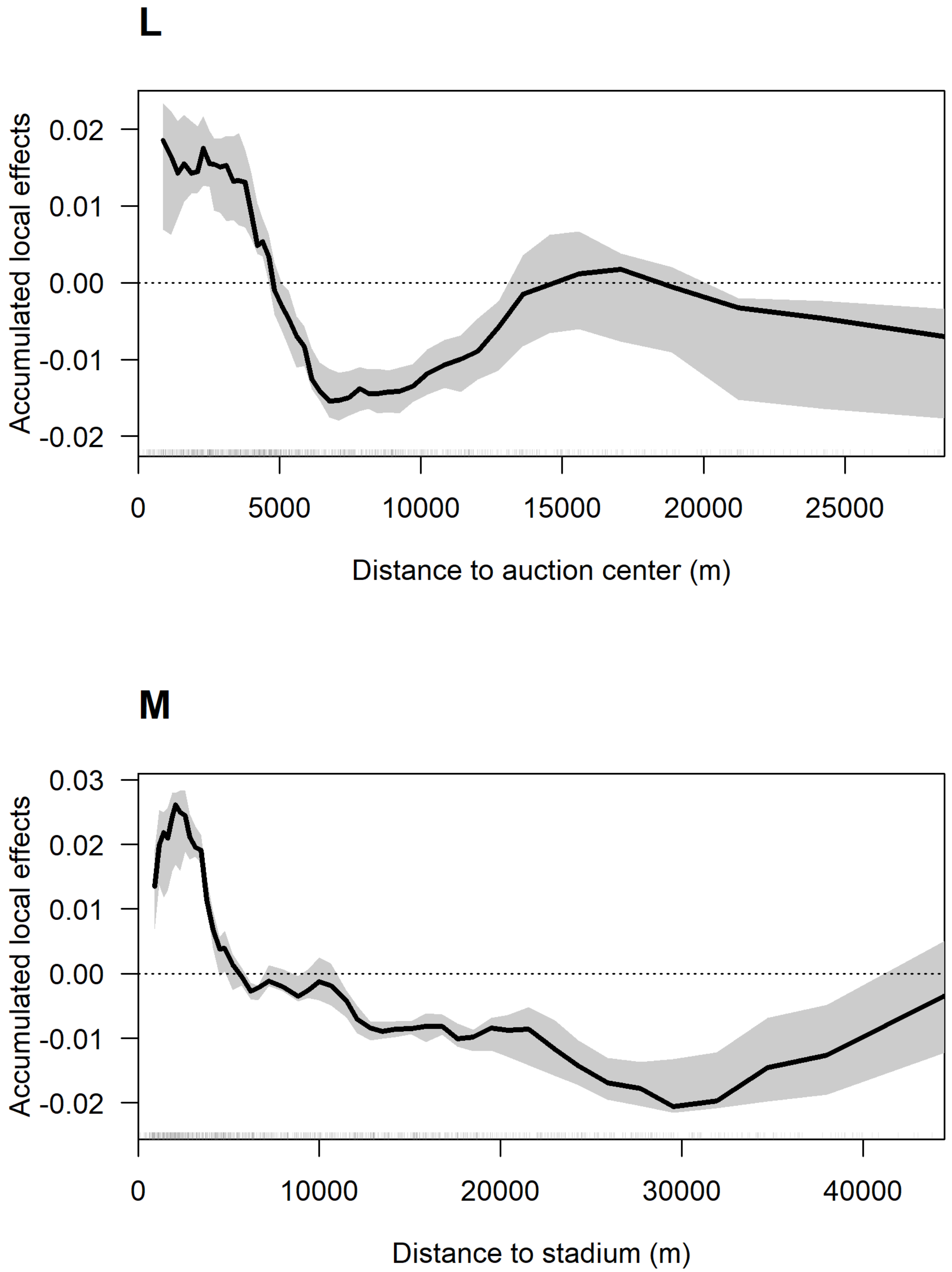
L
Accumulated local effects
0.02
0.01
0.00
-0.01
-0.02
0
5000
10000
15000
20000
25000
Distance to auction center (m)
M
Accumulated local effects
0.03
0.02
0.01
0.00
-0.01
-0.02
0
10000
20000
30000
40000
Distance to stadium (m)

**N**

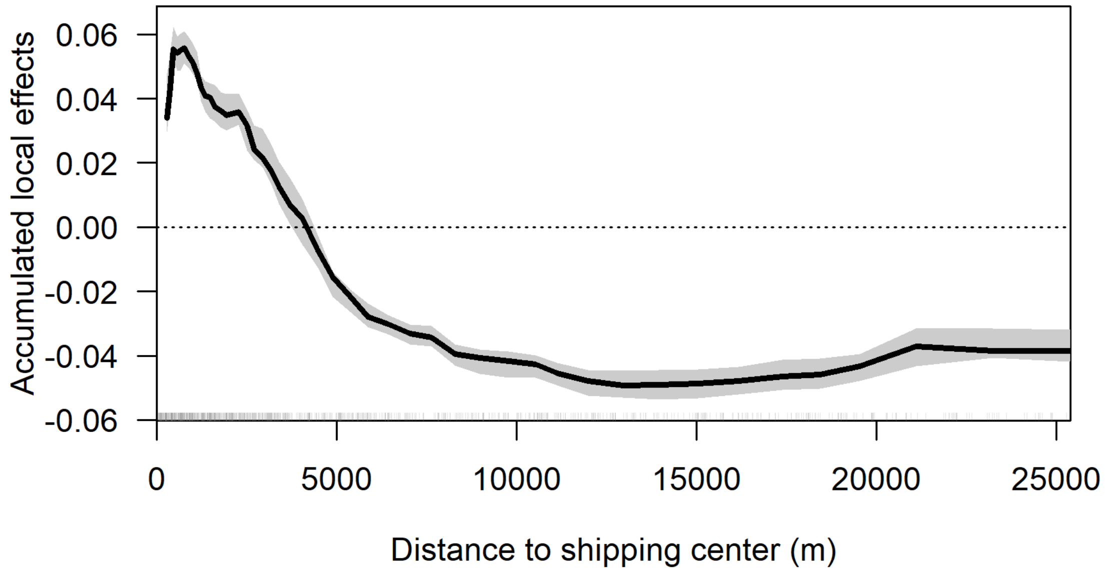

Accumulated local effects
0.06
0.04
0.02
0.00
-0.02
-0.04
-0.06
0
5000
10000
15000
20000
25000
Distance to shipping center (m)


**O**

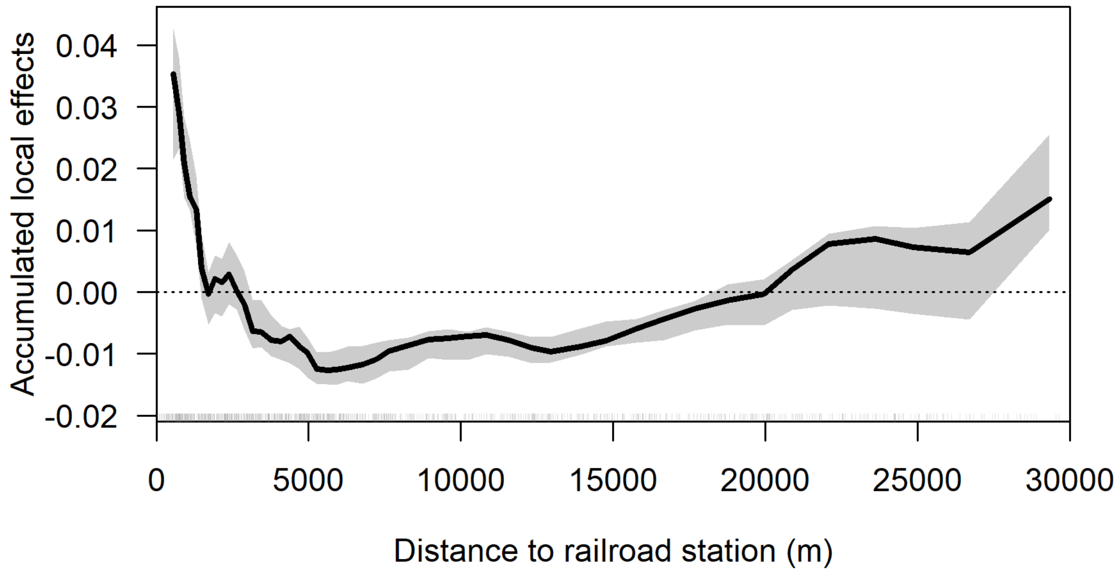

Accumulated local effects
0.04
0.03
0.02
0.01
0.00
-0.01
-0.02
0
5000
10000
15000
20000
25000
30000
Distance to railroad station (m)

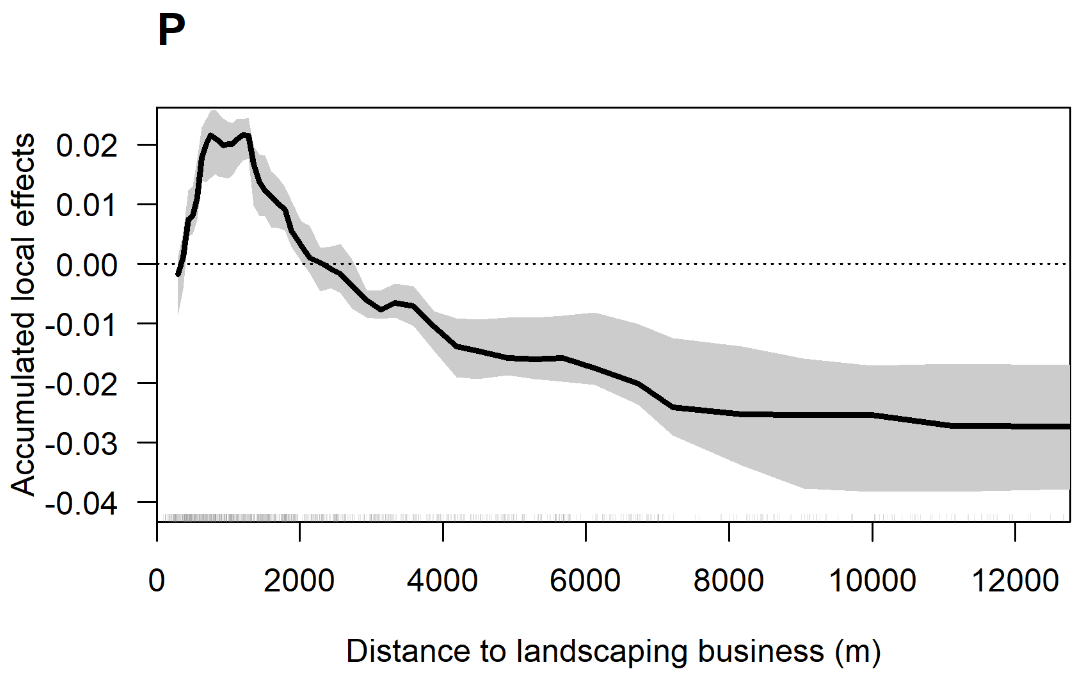

P
Accumulated local effects
0.02
0.01
0.00
-0.01
-0.02
-0.03
-0.04
0
2000
4000
6000
8000
10000
12000
Distance to landscaping business (m)


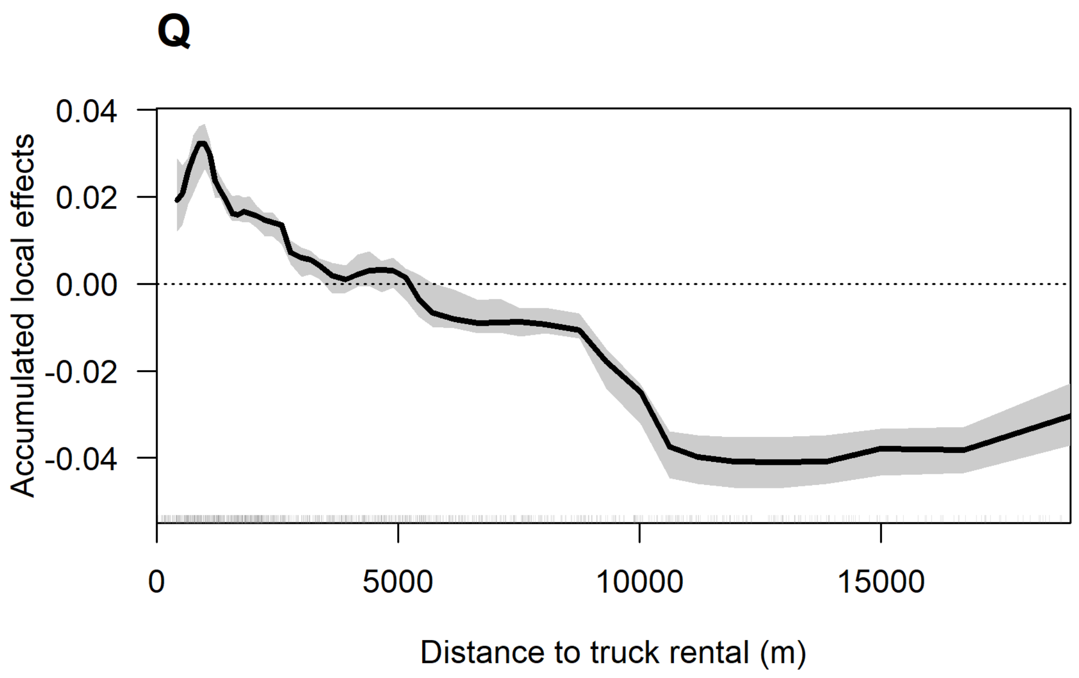

Q
Accumulated local effects
0.04
0.02
0.00
-0.02
-0.04
0
5000
10000
15000
Distance to truck rental (m)

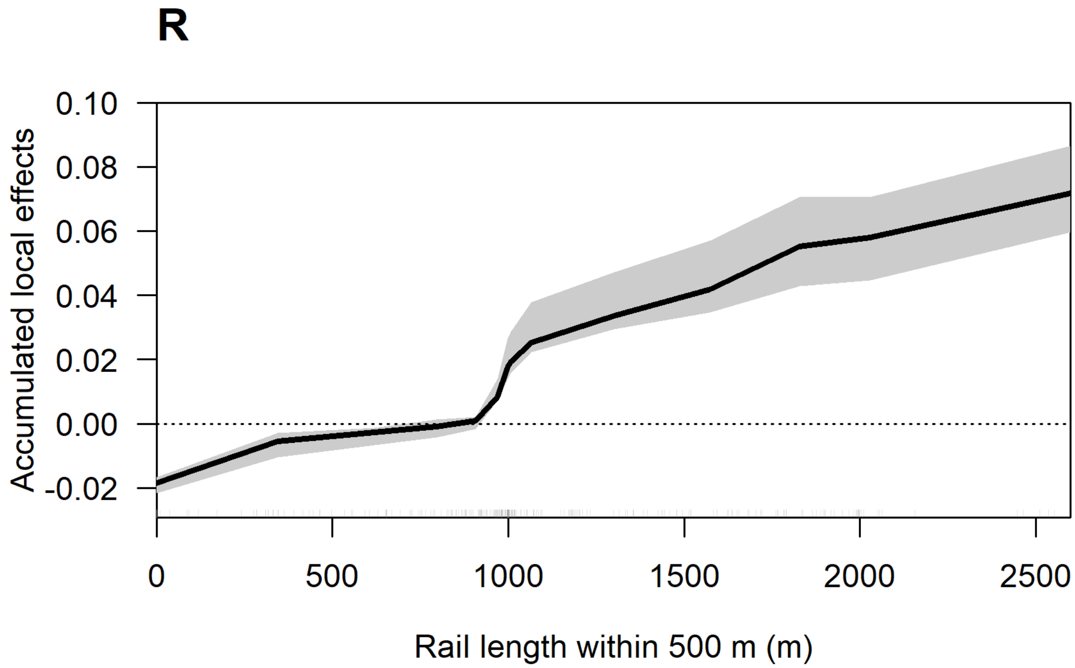


**Figure S1.26:** Accumulated local effects of the best fit ensemble random forest (ERF) model, which was identified by forward selection. Plots show the relationship between various predictors and the estimated risk of colonization by jump dispersal in the ERF with reduced pathway predictors and distance from prior *L. delicatula* established populations. In these plots, the range of values for a given predictor are displayed on the x-axis, while the y-axis shows the estimated impact on risk for that value. Positive y values indicate increased risk of jump dispersal, while negative values indicate reduced risk. Note that the y-axis range varies from plot to plot.

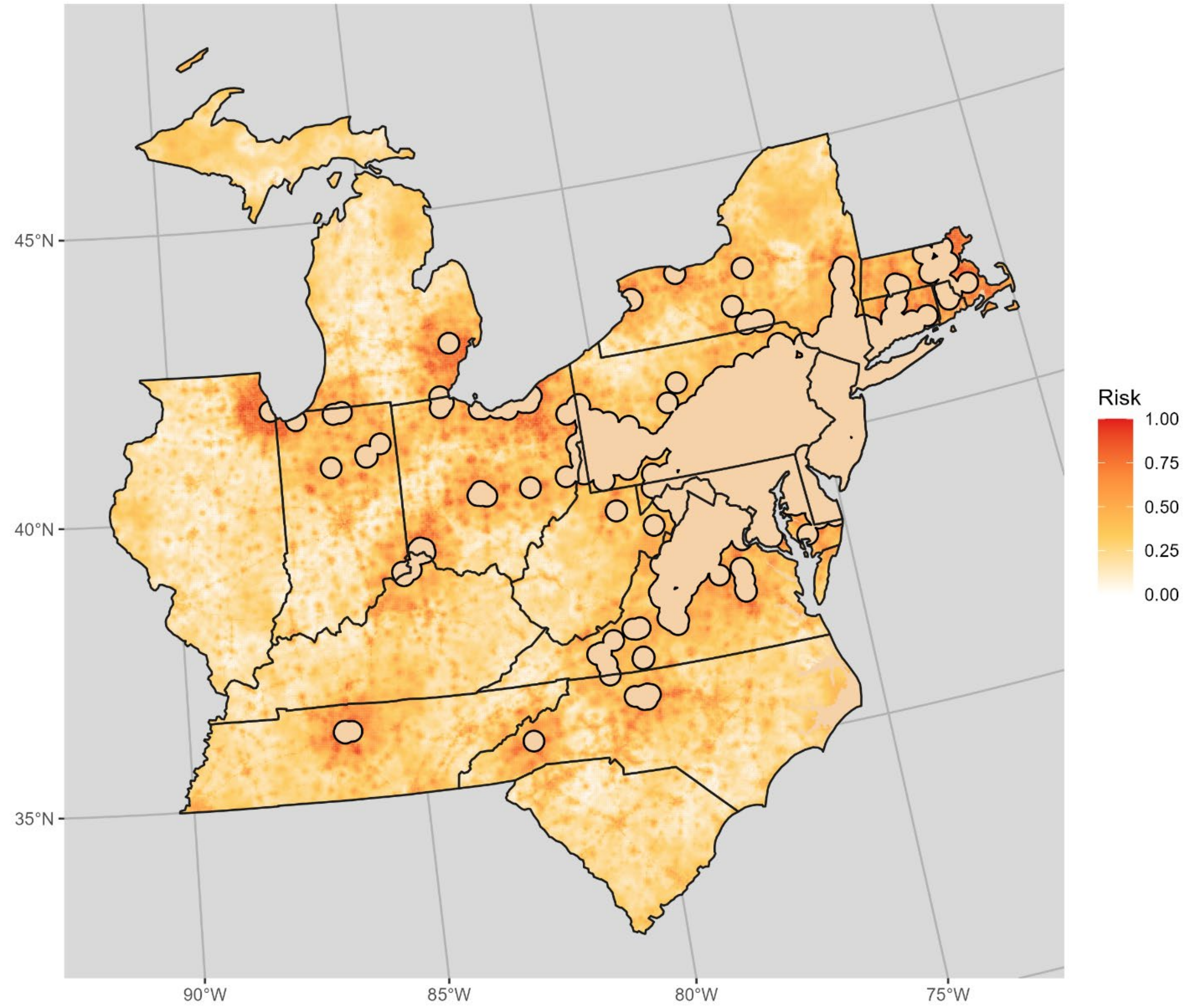


**Figure S1.26:** Projected risk of jump dispersal colonization based on the full ensemble random forest model with all predictors for 2024 including distance from previously established *L. delicatula* populations. Note that solid tan areas were already close to existing *L. delicatula* populations and therefore not able to be colonized by jump dispersal. Darker, redder colors indicate greater projected risk of *L. delicatula* colonization by jump dispersal.

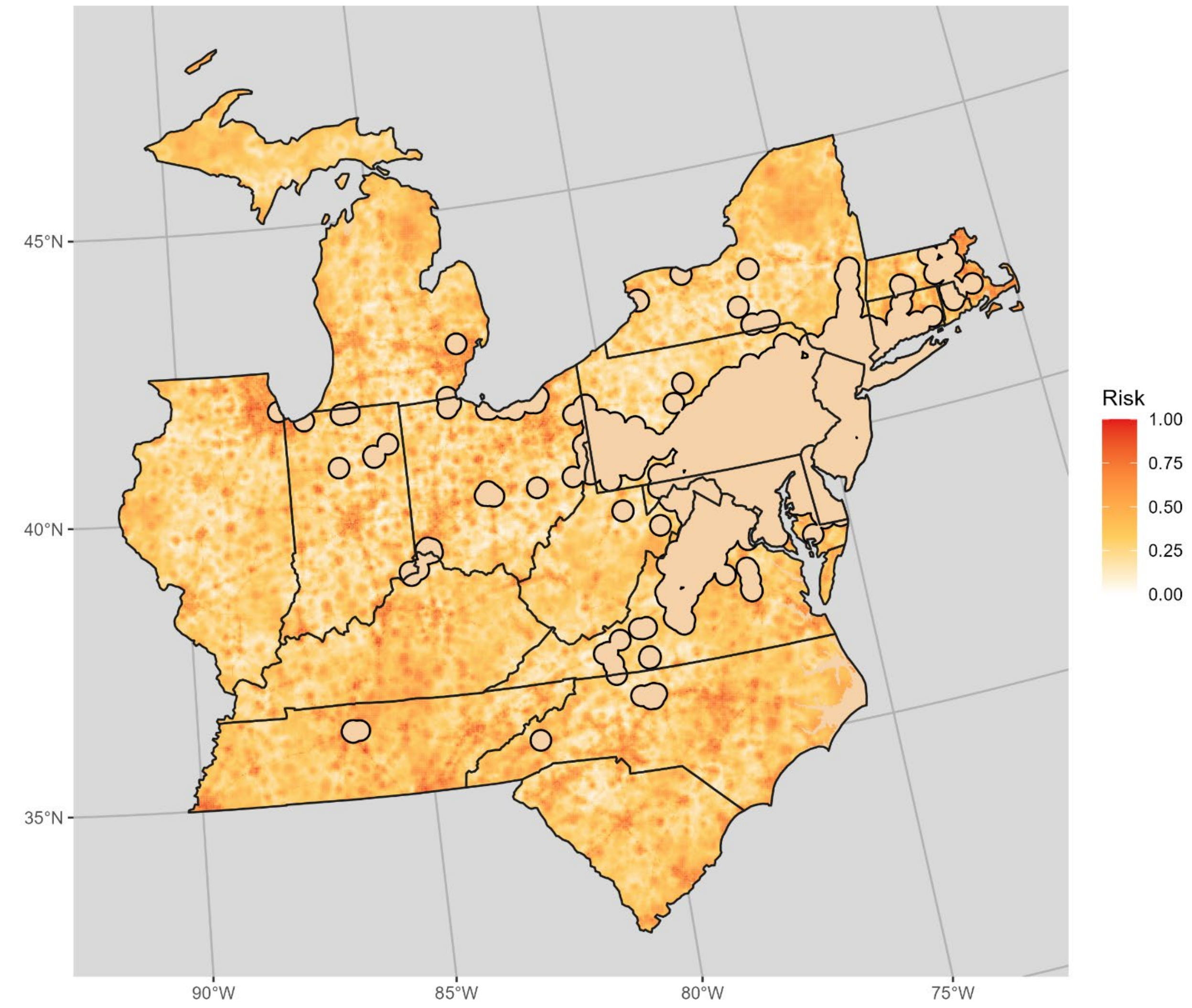


**Figure S1.27:** Projected risk of jump dispersal colonization based on the full ensemble random forest model including all predictors for 2024 except distance from previously established *L. delicatula* populations. Note that solid tan areas were already close to existing *L. delicatula* populations and therefore not able to be colonized by jump dispersal. Darker, redder colors indicate greater projected risk of *L. delicatula* colonization by jump dispersal.

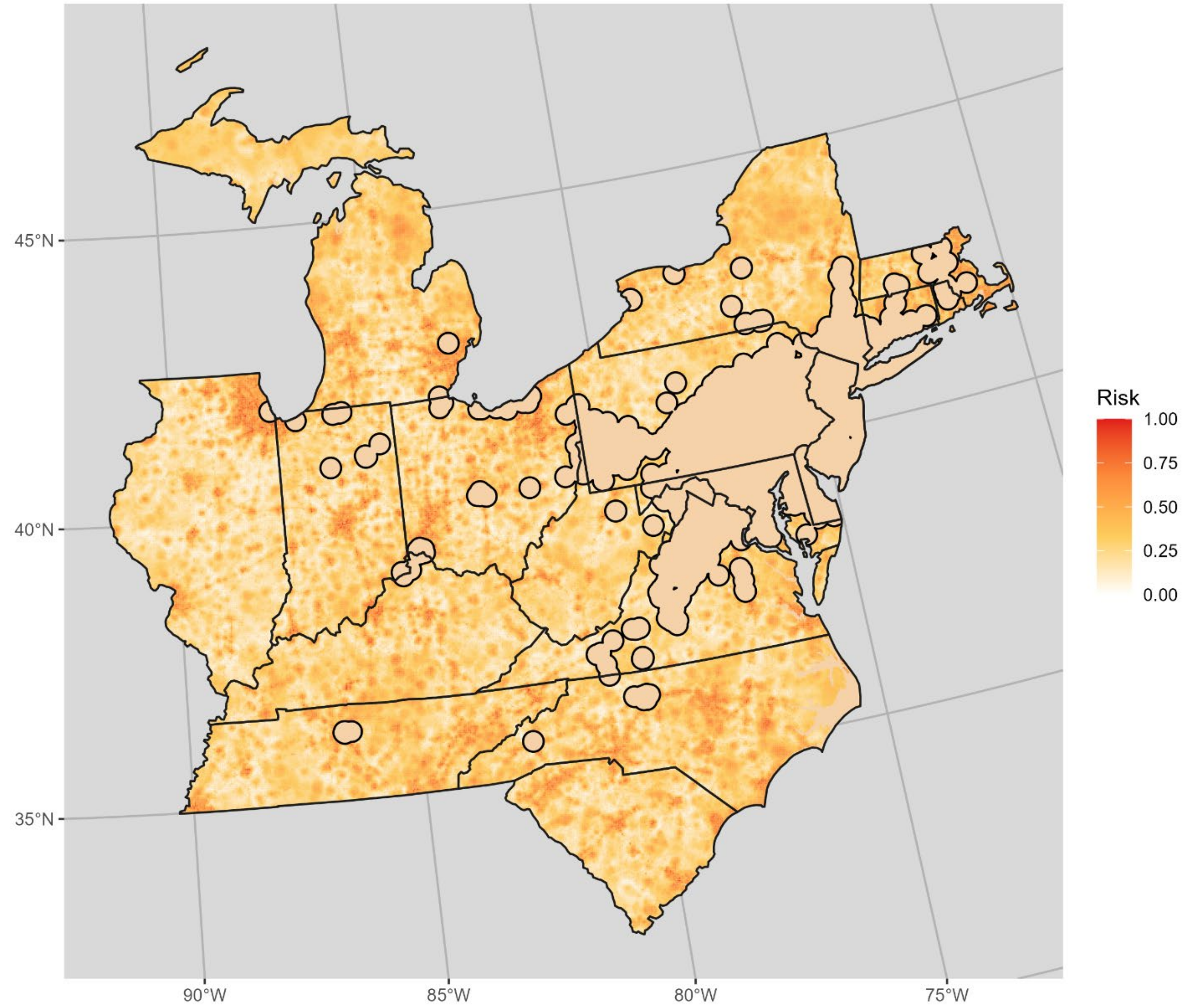


**Figure S1.28:** Projected risk of jump dispersal colonization based on an ensemble random forest model including all significant pathway predictors for 2024 while omitting and distance from existing *L. delicatula* populations. Note that solid tan areas were already close to existing *L. delicatula* populations and therefore not able to be colonized by jump dispersal. Darker, redder colors indicate greater projected risk of *L. delicatula* colonization by jump dispersal.

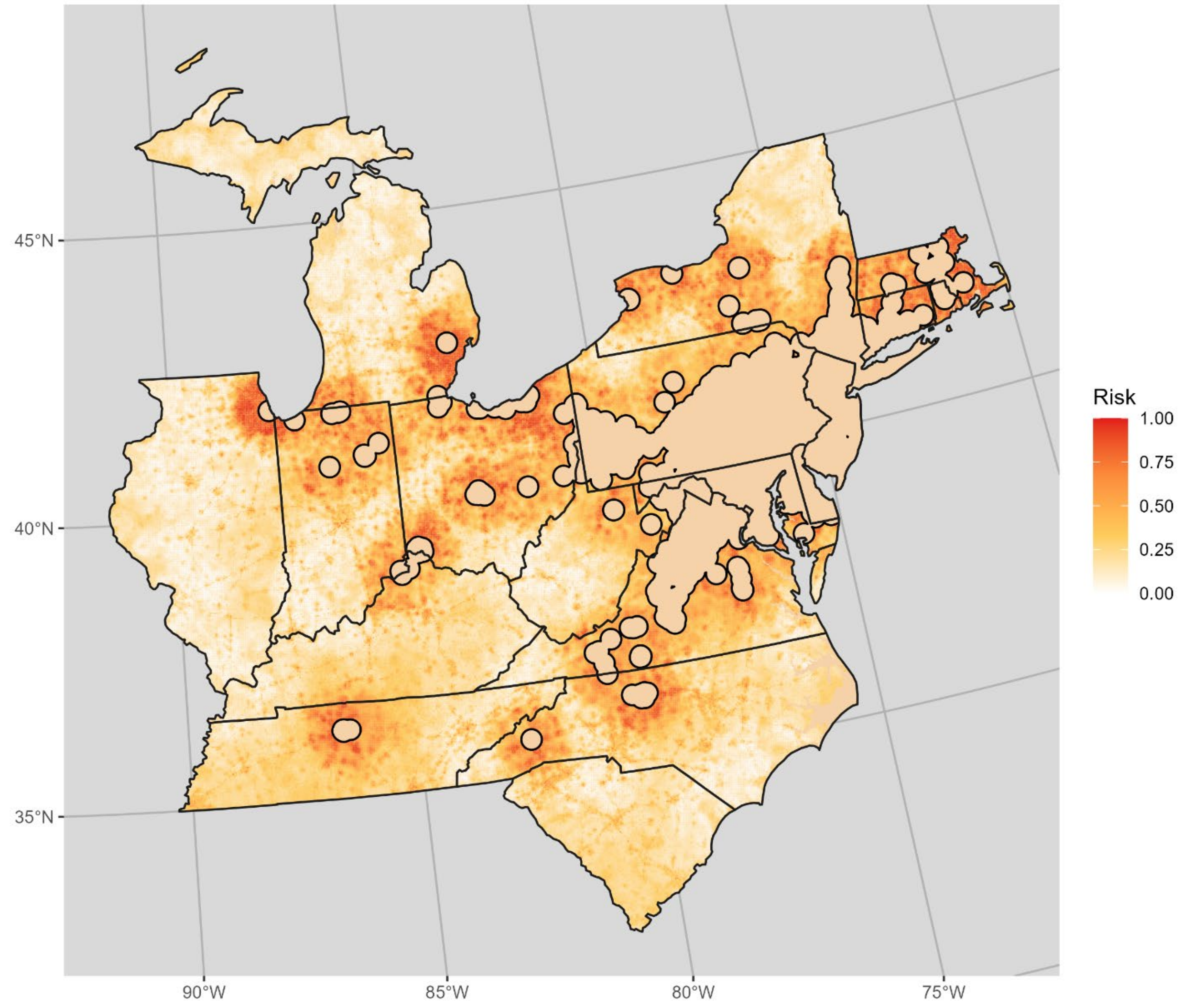


**Figure S1.29:** Projected risk of jump dispersal colonization based on the ensemble random forest model with predictors selected by forward selection. Note that solid tan areas were already close to existing *L. delicatula* populations and therefore not able to be colonized by jump dispersal. Darker, redder colors indicate greater projected risk of *L. delicatula* colonization by jump dispersal.

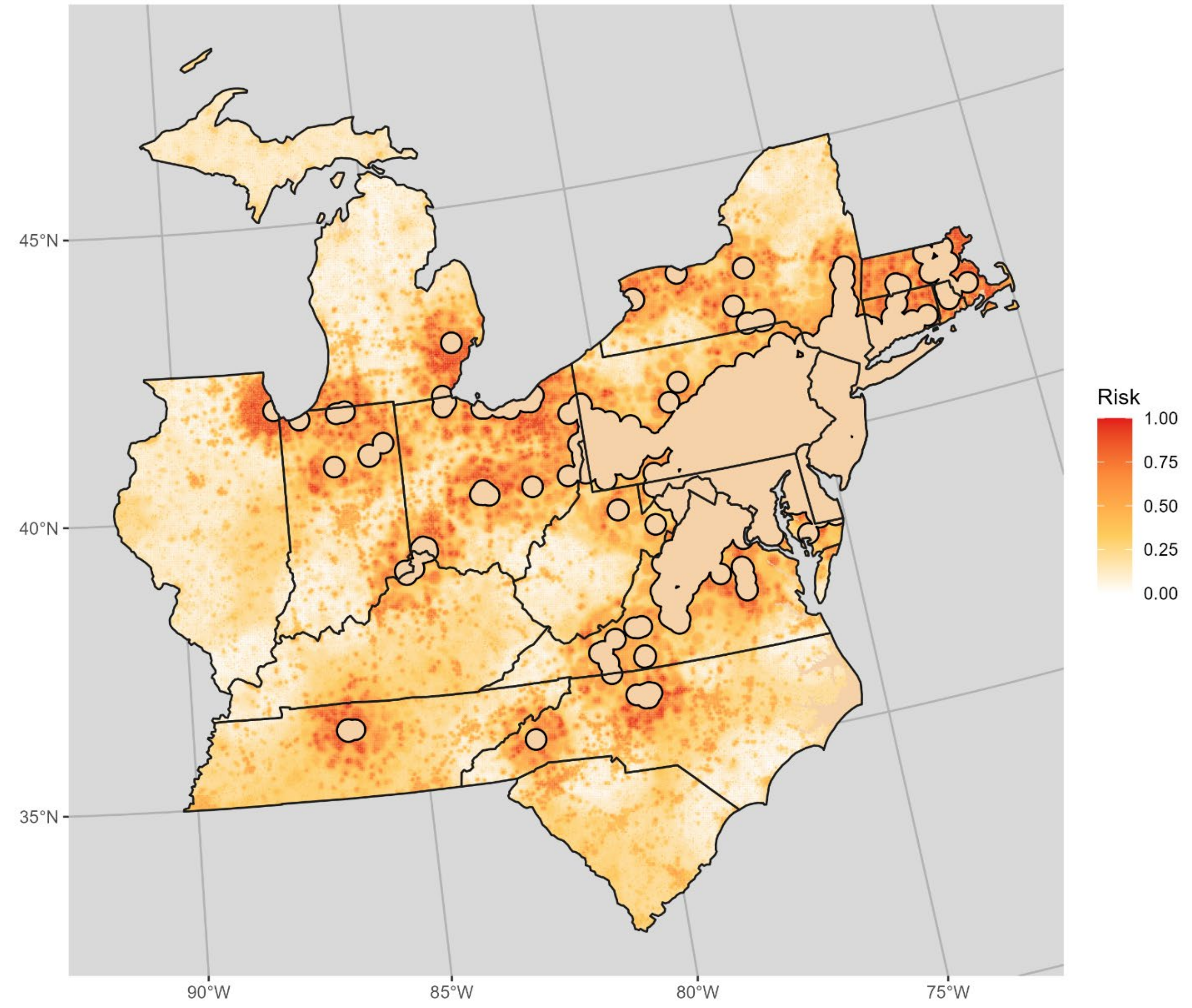


**Figure S1.30:** Projected risk of jump dispersal colonization based on the ensemble random forest model with predictors selected by backward selection. Note that solid tan areas were already close to existing *L. delicatula* populations and therefore not able to be colonized by jump dispersal. Darker, redder colors indicate greater projected risk of *L. delicatula* colonization by jump dispersal.
