## Supplementary material for "Mapping long-distance jump dispersal risk by leveraging transport infrastructure data and sampling bias correction: a case study with the invasive spotted lanternfly (*Lycorma delicatula*)": S2 Connecticut risk

*Supplemental material S2.*  Mapping long-distance jump dispersal risk by leveraging transport infrastructure data and sampling bias correction: a case study with the invasive spotted lanternfly (*Lycorma delicatula*)
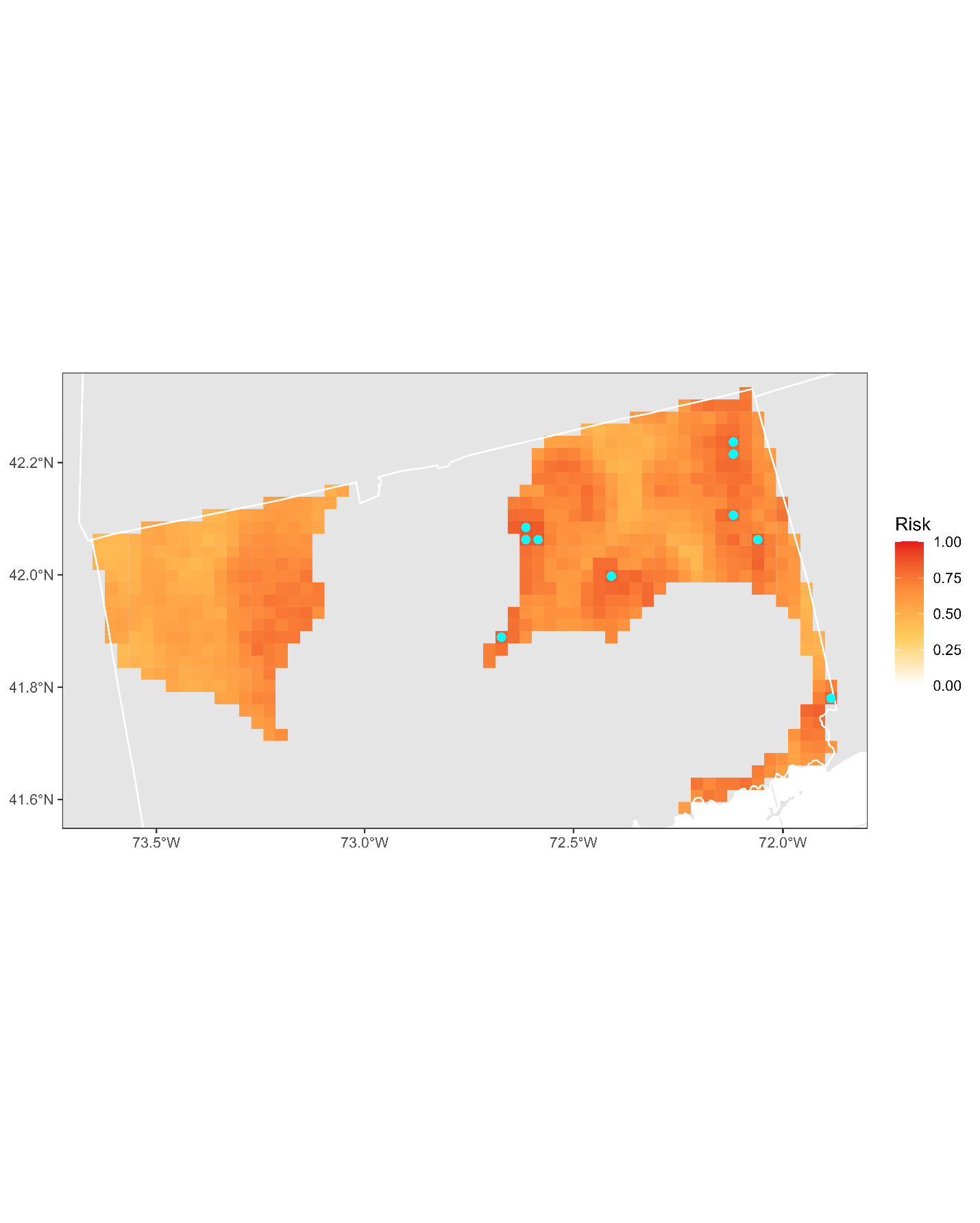

**Figure S2.1:** Projected risk of colonization by *L. delicatula* jump dispersal for Connecticut. Risk projections are based on an ensemble random forest model including distances from existing *L. delicatula* populations. Note that areas falling less than 20 km from existing *L. delicatula* populations are not included in the map because any colonization there would be categorized as diffusive spread based on our methodology (regardless of the actual source of the resulting population). Darker, redder colors indicate increased risk of colonization by jump dispersal. Cyan points show the top ten highest risk locations in the state.

**Table S2.1:** Ranked list of Connecticut county subdivisions sorted by the maximum projected risk of colonization by jump dispersal occurring there. For each subdivision, we identified the 2.5 x 2.5 km grid cell centroid included there with the highest projected risk of colonization by jump dispersal and report that location and risk value.

| **County, subdivision** | **Median risk value** | **Maximum risk value** | **Latitude of maximum risk** | **Longitude of maximum risk** |
| --- | --- | --- | --- | --- |
| Capitol Planning Region, Ellington town | 0.732921 | 0.87816 | 41.88090589 | -72.43140804 |
| Capitol Planning Region, Tolland town | 0.680473 | 0.869554 | 41.85375711 | -72.40945423 |
| Capitol Planning Region, Vernon town | 0.770576 | 0.864443 | 41.85924382 | -72.43885265 |
| Capitol Planning Region, Mansfield town | 0.621436 | 0.852878 | 41.75574976 | -72.25564517 |
| Capitol Planning Region, Glastonbury town | 0.722162 | 0.842926 | 41.6968748 | -72.55691874 |
| Northeastern Connecticut Planning Region, Plainfield town | 0.724045 | 0.839968 | 41.75385797 | -71.8810752 |
| Southeastern Connecticut Planning Region, North Stonington town | 0.724314 | 0.831092 | 41.43907798 | -71.80438778 |
| Northeastern Connecticut Planning Region, Thompson town | 0.737224 | 0.828402 | 41.93811187 | -71.87886792 |
| Northeastern Connecticut Planning Region, Brooklyn town | 0.723776 | 0.820065 | 41.8083257 | -71.92452721 |
| Northeastern Connecticut Planning Region, Putnam town | 0.773669 | 0.80581 | 41.91648064 | -71.88648957 |
| Capitol Planning Region, Coventry town | 0.635557 | 0.802851 | 41.73410086 | -72.26313075 |
| Southeastern Connecticut Planning Region, Windham town | 0.745024 | 0.798817 | 41.69585963 | -72.18265445 |
| Capitol Planning Region, Columbia town | 0.719204 | 0.788327 | 41.71796953 | -72.29994005 |
| Capitol Planning Region, Willington town | 0.700377 | 0.781065 | 41.91835576 | -72.26193035 |
| Northeastern Connecticut Planning Region, Killingly town | 0.686122 | 0.780796 | 41.85158745 | -71.90932625 |
| Capitol Planning Region, Stafford town | 0.69177 | 0.779989 | 41.9727046 | -72.305781 |
| Southeastern Connecticut Planning Region, Stonington town | 0.643222 | 0.777031 | 41.40146557 | -71.84872507 |
| Southeastern Connecticut Planning Region, Groton town | 0.696611 | 0.775148 | 41.32689611 | -72.06135549 |
| Northwest Hills Planning Region, New Hartford town | 0.732652 | 0.773265 | 41.82669707 | -73.01245639 |
| Northwest Hills Planning Region, Litchfield town | 0.599919 | 0.770576 | 41.78295024 | -73.15187909 |
| Capitol Planning Region, Bolton town | 0.711942 | 0.767348 | 41.79425819 | -72.46115885 |
| Northwest Hills Planning Region, Torrington town | 0.720414 | 0.761162 | 41.79932564 | -73.11523321 |
| Northeastern Connecticut Planning Region, Woodstock town | 0.613098 | 0.758203 | 41.94932142 | -71.93765627 |
| Southeastern Connecticut Planning Region, Waterford town | 0.738838 | 0.756052 | 41.34910859 | -72.17798926 |
| Northeastern Connecticut Planning Region, Pomfret town | 0.657612 | 0.752286 | 41.87882092 | -71.93108513 |
| Northeastern Connecticut Planning Region, Canterbury town | 0.655729 | 0.751748 | 41.73783564 | -71.91798247 |
| Northwest Hills Planning Region, Barkhamsted town | 0.675901 | 0.748252 | 41.8917794 | -72.99065213 |
| Northwest Hills Planning Region, Harwinton town | 0.7188 | 0.747176 | 41.79399359 | -73.08580768 |
| Southeastern Connecticut Planning Region, Lebanon town | 0.685853 | 0.746638 | 41.70692787 | -72.24128801 |
| Southeastern Connecticut Planning Region, New London town | 0.739645 | 0.739645 | 41.33245914 | -72.09050704 |
| Northeastern Connecticut Planning Region, Eastford town | 0.628564 | 0.719742 | 41.88509142 | -72.08554421 |
| Northeastern Connecticut Planning Region, Scotland town | 0.661377 | 0.711673 | 41.72804416 | -72.10898831 |
| Northwest Hills Planning Region, Winchester town | 0.67886 | 0.711404 | 41.91315667 | -73.10851668 |
| Northeastern Connecticut Planning Region, Chaplin town | 0.617267 | 0.704142 | 41.75524019 | -72.13078634 |
| Northwest Hills Planning Region, Morris town | 0.626009 | 0.698763 | 41.69614823 | -73.18069797 |
| Northeastern Connecticut Planning Region, Sterling town | 0.63085 | 0.680473 | 41.69937882 | -71.83769065 |
| Naugatuck Valley Planning Region, Watertown town | 0.669984 | 0.678322 | 41.66913325 | -73.15851285 |
| Northwest Hills Planning Region, Norfolk town | 0.480097 | 0.677246 | 41.92380511 | -73.16747593 |
| Northwest Hills Planning Region, Colebrook town | 0.604626 | 0.66326 | 41.97292197 | -73.05727623 |
| Capitol Planning Region, Manchester town | 0.657343 | 0.657343 | 41.75093487 | -72.47600668 |
| Capitol Planning Region, Hebron town | 0.613233 | 0.655729 | 41.71831515 | -72.42473343 |
| Northeastern Connecticut Planning Region, Voluntown town | 0.490855 | 0.651425 | 41.63450966 | -71.86050183 |
| Capitol Planning Region, Andover town | 0.615116 | 0.64766 | 41.71282608 | -72.39539516 |
| Northwest Hills Planning Region, Goshen town | 0.550296 | 0.644164 | 41.90210017 | -73.17469702 |
| Northwest Hills Planning Region, Hartland town | 0.579344 | 0.636902 | 41.96757563 | -73.02778206 |
| Southeastern Connecticut Planning Region, East Lyme town | 0.612157 | 0.633943 | 41.3274731 | -72.18545552 |
| Naugatuck Valley Planning Region, Bethlehem town | 0.597902 | 0.621033 | 41.64743486 | -73.16571053 |
| Northeastern Connecticut Planning Region, Ashford town | 0.494352 | 0.618343 | 41.88564481 | -72.21063439 |
| Northwest Hills Planning Region, Cornwall town | 0.550834 | 0.594406 | 41.83646267 | -73.32133697 |
| Northeastern Connecticut Planning Region, Hampton town | 0.526493 | 0.576385 | 41.77132377 | -72.09391861 |
| Northwest Hills Planning Region, Sharon town | 0.529317 | 0.568585 | 41.88450849 | -73.46156148 |
| Northwest Hills Planning Region, Canaan town | 0.505917 | 0.556482 | 41.93387397 | -73.35163052 |
| Northwest Hills Planning Region, Warren town | 0.511027 | 0.555944 | 41.77660209 | -73.37225234 |
| Northwest Hills Planning Region, Salisbury town | 0.481173 | 0.554061 | 41.95492234 | -73.46967525 |
| Northwest Hills Planning Region, North Canaan town | 0.512372 | 0.54922 | 42.04245349 | -73.31573874 |
| Capitol Planning Region, Granby town | 0.540882 | 0.540882 | 42.02750159 | -72.85108614 |
| Northeastern Connecticut Planning Region, Union town | 0.498521 | 0.5234 | 41.99389321 | -72.17299759 |
| Northwest Hills Planning Region, Kent town | 0.503228 | 0.503228 | 41.78712844 | -73.43113651 |
