## Supplementary material for "Mapping long-distance jump dispersal risk by leveraging transport infrastructure data and sampling bias correction: a case study with the invasive spotted lanternfly (*Lycorma delicatula*)": S3 Illinois risk

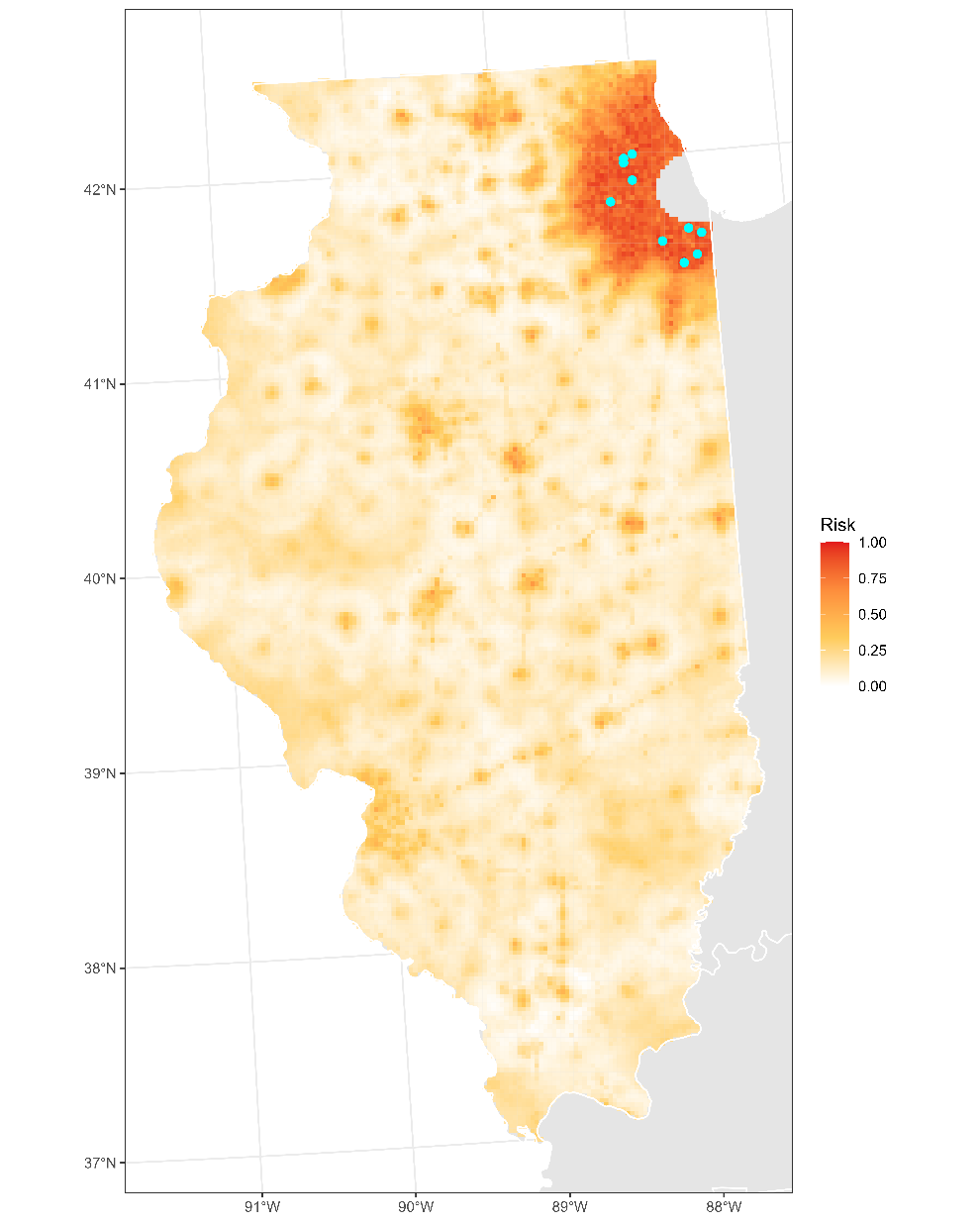

**Figure S3.1:** Projected risk of colonization by *L. delicatula* jump dispersal for Illinois. Risk projections are based on an ensemble random forest model including distances from existing *L. delicatula* populations. Note that areas falling less than 20 km from existing *L. delicatula* populations are not included in the map because any colonization there would be categorized as diffusive spread based on our methodology (regardless of the actual source of the resulting population). Darker, redder colors indicate increased risk of colonization by jump dispersal. Cyan points show the top ten highest risk locations in the state.

**Table S3.1:** Ranked list of Illinois county subdivisions sorted by the maximum projected risk of colonization by jump dispersal occurring there. For each subdivision, we identified the 2.5 x 2.5 km grid cell centroid included there with the highest projected risk of colonization by jump dispersal and report that location and risk value.

| **County, subdivision** | **Median risk value** | **Maximum risk value** | **Latitude of maximum risk** | **Longitude of maximum risk** |
| --- | --- | --- | --- | --- |
| Will County, Frankfort township | 0.86040882 | 1 | 41.5531482 | -87.86910003 |
| Cook County, Schaumburg township | 0.84749866 | 0.99220011 | 41.99310725 | -88.09223931 |
| DuPage County, Bloomingdale township | 0.8922808 | 0.97660032 | 41.97083615 | -88.09476403 |
| Will County, Monee township | 0.85529855 | 0.97417967 | 41.43225643 | -87.73181012 |
| Cook County, Elk Grove township | 0.82571275 | 0.97095212 | 42.01164841 | -88.02913503 |
| DuPage County, Naperville township | 0.84104357 | 0.96180742 | 41.77594113 | -88.20797799 |
| Cook County, Thornton township | 0.83996772 | 0.957773 | 41.58018477 | -87.59306262 |
| Cook County, Bremen township | 0.82786444 | 0.95346961 | 41.60832722 | -87.6806975 |
| DuPage County, Milton township | 0.86498117 | 0.94620764 | 41.87803422 | -88.04438374 |
| Cook County, Bloom township | 0.86686391 | 0.94217321 | 41.47090094 | -87.6364286 |
| Will County, Lockport township | 0.85032275 | 0.93921463 | 41.63312196 | -88.07218905 |
| DuPage County, Wayne township | 0.85287789 | 0.93706294 | 41.91321219 | -88.2535621 |
| Cook County, Maine township | 0.83862292 | 0.93006993 | 42.04863943 | -87.9028261 |
| Cook County, Orland township | 0.83499193 | 0.93006993 | 41.57731067 | -87.89661468 |
| DuPage County, York township | 0.8091716 | 0.92872512 | 41.8984293 | -88.01160763 |
| Will County, Joliet township | 0.82329209 | 0.92011834 | 41.49955761 | -88.08727401 |
| DuPage County, Downers Grove township | 0.81414739 | 0.91931146 | 41.740695 | -87.99923203 |
| Cook County, Palatine township | 0.80621302 | 0.91823561 | 42.12488023 | -88.04671932 |
| Cook County, Rich township | 0.83351264 | 0.91527703 | 41.49901076 | -87.72394467 |
| Kane County, Geneva township | 0.86605702 | 0.91447015 | 41.8704831 | -88.28873388 |
| Cook County, Hanover township | 0.78160301 | 0.91258741 | 42.04319545 | -88.17809279 |
| Lake County, Libertyville township | 0.75672405 | 0.9112426 | 42.2955385 | -87.90466531 |
| Kane County, Aurora township | 0.80500269 | 0.91070468 | 41.77958683 | -88.26836162 |
| Will County, Crete township | 0.81145777 | 0.90989779 | 41.46893423 | -87.60638543 |
| DuPage County, Addison township | 0.85032275 | 0.90909091 | 41.96146324 | -87.94342714 |
| DuPage County, Winfield township | 0.8006993 | 0.90855299 | 41.89094002 | -88.25603261 |
| Kane County, St. Charles township | 0.79182356 | 0.90586337 | 41.91502817 | -88.28381355 |
| Will County, Troy township | 0.76465842 | 0.90505648 | 41.50325887 | -88.14741416 |
| Will County, Wheatland township | 0.84615385 | 0.90398063 | 41.64234107 | -88.2228353 |
| Lake County, Vernon township | 0.78133405 | 0.9020979 | 42.18416615 | -87.9176272 |
| Cook County, Wheeling township | 0.85045724 | 0.90182894 | 42.09885669 | -87.98861186 |
| Cook County, Lemont township | 0.81414739 | 0.8986014 | 41.65164601 | -88.00940298 |
| DuPage County, Lisle township | 0.83216783 | 0.89779451 | 41.79268603 | -88.11490407 |
| Will County, New Lenox township | 0.79034427 | 0.8964497 | 41.51621061 | -87.99453719 |
| Kane County, Batavia township | 0.80486821 | 0.89537386 | 41.82412659 | -88.26343472 |
| Will County, Du Page township | 0.83835395 | 0.89080151 | 41.68135844 | -88.12743969 |
| Cook County, Proviso township | 0.84669177 | 0.89080151 | 41.84633941 | -87.89583777 |
| Will County, Green Garden township | 0.73695535 | 0.88434642 | 41.46221697 | -87.8493615 |
| Cook County, Chicago city | 0.78886498 | 0.87331899 | 41.97610258 | -87.81977621 |
| Will County, Homer township | 0.82477138 | 0.87170522 | 41.60712434 | -88.01447879 |
| Cook County, Palos township | 0.81065089 | 0.87116729 | 41.68477762 | -87.82344673 |
| Lake County, Moraine township | 0.81253362 | 0.86982249 | 42.17651385 | -87.79620334 |
| Lake County, Ela township | 0.79800968 | 0.86928456 | 42.17129551 | -88.07198344 |
| Will County, Plainfield township | 0.78738569 | 0.86578806 | 41.59414754 | -88.16754988 |
| Cook County, Niles township | 0.80607854 | 0.86309844 | 42.03904069 | -87.75134695 |
| Kendall County, Oswego township | 0.77541689 | 0.86229155 | 41.692323 | -88.30835275 |
| Cook County, Leyden township | 0.81226466 | 0.85556751 | 41.91503453 | -87.91832406 |
| Lake County, Shields township | 0.72969338 | 0.85529855 | 42.27135782 | -87.87686399 |
| Cook County, Barrington township | 0.80204411 | 0.85449166 | 42.08958045 | -88.20341466 |
| Kane County, Elgin township | 0.73964497 | 0.8507262 | 42.00412457 | -88.27395407 |
| Cook County, Northfield township | 0.81253362 | 0.84991931 | 42.06900256 | -87.8699286 |
| Lake County, West Deerfield township | 0.80554061 | 0.84938139 | 42.22489879 | -87.85169362 |
| Cook County, Lyons township | 0.81280258 | 0.82974718 | 41.75538431 | -87.87598201 |
| Cook County, Evanston city | 0.78644432 | 0.82920925 | 42.05936502 | -87.71840985 |
| Kane County, Sugar Grove township | 0.67455621 | 0.82598171 | 41.78679339 | -88.3891476 |
| Lake County, Waukegan township | 0.63367402 | 0.82167832 | 42.33627027 | -87.83861861 |
| Kane County, Dundee township | 0.71543841 | 0.81253362 | 42.11550716 | -88.26159431 |
| Will County, Jackson township | 0.72794513 | 0.81038193 | 41.43092216 | -88.06475565 |
| Kankakee County, Manteno township | 0.66379774 | 0.80930608 | 41.23778932 | -87.84516156 |
| Cook County, Worth township | 0.79948897 | 0.80876815 | 41.6605968 | -87.79591516 |
| Cook County, New Trier township | 0.77245831 | 0.80607854 | 42.15231356 | -87.76848707 |
| Cook County, Norwood Park township | 0.78994083 | 0.78994083 | 41.95383734 | -87.82238707 |
| Will County, Channahon township | 0.68786982 | 0.7880581 | 41.46057733 | -88.18244555 |
| Kankakee County, Kankakee township | 0.54962345 | 0.78214094 | 41.12654895 | -87.85802327 |
| Lake County, Fremont township | 0.65868747 | 0.76089295 | 42.325369 | -88.02374985 |
| Lake County, Warren township | 0.68477676 | 0.75901022 | 42.3400915 | -87.89946893 |
| Kane County, Blackberry township | 0.6105433 | 0.7506724 | 41.83133808 | -88.384297 |
| Lake County, Wauconda township | 0.64873588 | 0.74905863 | 42.26411927 | -88.12261271 |
| Kane County, Campton township | 0.60476062 | 0.74341044 | 41.944497 | -88.40241074 |
| Kendall County, Bristol township | 0.61820871 | 0.73614847 | 41.71998007 | -88.39641194 |
| Lake County, Cuba township | 0.66729424 | 0.73184508 | 42.19728389 | -88.13018237 |
| McHenry County, Algonquin township | 0.61915008 | 0.71570737 | 42.18596906 | -88.31489759 |
| Kendall County, Seward township | 0.49260355 | 0.71113502 | 41.46605159 | -88.27261558 |
| Kendall County, Na-Au-Say township | 0.55433029 | 0.70064551 | 41.57736825 | -88.26034831 |
| Kane County, Plato township | 0.38353954 | 0.697149 | 41.98905016 | -88.39755765 |
| Kankakee County, Bourbonnais township | 0.52850995 | 0.69176977 | 41.17104395 | -87.85288345 |
| Kane County, Rutland township | 0.50430339 | 0.68370091 | 42.10044248 | -88.38539783 |
| Lake County, Avon township | 0.58270576 | 0.68343195 | 42.35138411 | -88.08205968 |
| Will County, Wilmington township | 0.44566971 | 0.67697687 | 41.3492809 | -88.19481549 |
| Kendall County, Kendall township | 0.39725659 | 0.67670791 | 41.63090244 | -88.40607624 |
| Will County, Manhattan township | 0.52514793 | 0.67079075 | 41.44943884 | -88.00214795 |
| Will County, Will township | 0.52501345 | 0.67052179 | 41.38386268 | -87.6770261 |
| McLean County, Normal township | 0.38058096 | 0.66594944 | 40.54515881 | -88.99967699 |
| McHenry County, Nunda township | 0.48843464 | 0.66110812 | 42.25281415 | -88.30750641 |
| Champaign County, Champaign City township | 0.43719742 | 0.65922539 | 40.12199445 | -88.26935719 |
| Boone County, Belvidere township | 0.43033889 | 0.6570737 | 42.2602212 | -88.82682317 |
| Will County, Peotone township | 0.52837547 | 0.65250134 | 41.30261891 | -87.80744719 |
| Vermilion County, Danville township | 0.36713287 | 0.64819796 | 40.10462501 | -87.64769616 |
| Lake County, Lake Villa township | 0.42011834 | 0.64443249 | 42.39220908 | -88.01606392 |
| Will County, Florence township | 0.27272727 | 0.6422808 | 41.36785962 | -88.13231451 |
| Winnebago County, Rockford township | 0.40182894 | 0.63340506 | 42.25280771 | -89.1028369 |
| Will County, Washington township | 0.54034427 | 0.6309844 | 41.38190564 | -87.64701891 |
| McLean County, Bloomington City township | 0.42011834 | 0.62721893 | 40.47838433 | -89.00624341 |
| Grundy County, Saratoga township | 0.41568047 | 0.62506724 | 41.38597707 | -88.43252707 |
| Boone County, Flora township | 0.2232383 | 0.62210866 | 42.23792918 | -88.82912262 |
| Lake County, Grant township | 0.43961807 | 0.61807423 | 42.37737389 | -88.1404131 |
| Macon County, Decatur township | 0.40478752 | 0.60731576 | 39.84870425 | -88.94933268 |
| Lake County, Benton township | 0.44419042 | 0.60435718 | 42.42729492 | -87.85859147 |
| Grundy County, Aux Sable township | 0.47081764 | 0.59870898 | 41.44378959 | -88.27506438 |
| Stephenson County, Freeport township | 0.40613233 | 0.59332975 | 42.3016412 | -89.61811462 |
| Sangamon County, Capital township | 0.29263045 | 0.5809575 | 39.84039787 | -89.62982294 |
| Winnebago County, Cherry Valley township | 0.33109198 | 0.58041958 | 42.22561747 | -89.01382577 |
| Peoria County, Peoria City township | 0.4232114 | 0.57504034 | 40.66719938 | -89.61542628 |
| Tazewell County, Fondulac township | 0.36242604 | 0.5739645 | 40.66569744 | -89.5856741 |
| Champaign County, Urbana township | 0.24569661 | 0.56455083 | 40.11652401 | -88.18087785 |
| Kankakee County, Sumner township | 0.43410436 | 0.56293706 | 41.27844381 | -87.78007019 |
| LaSalle County, Bruce township | 0.26654115 | 0.55271651 | 41.11730828 | -88.82253935 |
| Lake County, Newport township | 0.39940828 | 0.55110274 | 42.43110722 | -87.91952259 |
| Kankakee County, Rockville township | 0.33082302 | 0.54599247 | 41.26385362 | -87.90250743 |
| Grundy County, Morris township | 0.49045186 | 0.54007531 | 41.36371377 | -88.43492258 |
| McLean County, Bloomington township | 0.34884346 | 0.53711673 | 40.45448283 | -88.97877874 |
| Boone County, Bonus township | 0.16837009 | 0.52770307 | 42.25853179 | -88.79640807 |
| Kane County, Kaneville township | 0.25712749 | 0.52205487 | 41.88298791 | -88.50040366 |
| McHenry County, Grafton township | 0.39940828 | 0.51936525 | 42.17085202 | -88.43881463 |
| Effingham County, Douglas township | 0.15868747 | 0.51748252 | 39.10973281 | -88.55293721 |
| Coles County, Charleston township | 0.30823023 | 0.51748252 | 39.48844673 | -88.16067648 |
| Macon County, Hickory Point township | 0.21893491 | 0.51694459 | 39.8964989 | -89.00379779 |
| Boone County, Spring township | 0.13246369 | 0.51398601 | 42.23624036 | -88.79871719 |
| Champaign County, Cunningham township | 0.48251748 | 0.51398601 | 40.11835439 | -88.21036946 |
| Kendall County, Little Rock township | 0.43060785 | 0.51344809 | 41.68076184 | -88.49171 |
| DeKalb County, Cortland township | 0.32813341 | 0.51048951 | 41.91742626 | -88.70982261 |
| DeKalb County, DeKalb township | 0.4338354 | 0.50726197 | 41.89685396 | -88.74240321 |
| McHenry County, Dorr township | 0.38864981 | 0.50726197 | 42.30455893 | -88.4242593 |
| Kankakee County, Limestone township | 0.19607316 | 0.5056482 | 41.10810816 | -87.92038285 |
| Peoria County, Limestone township | 0.34185046 | 0.50430339 | 40.64642339 | -89.6471717 |
| Madison County, Nameoki township | 0.28644432 | 0.49919311 | 38.74608188 | -90.10368069 |
| Kankakee County, Yellowhead township | 0.39671867 | 0.49542765 | 41.24646504 | -87.63290508 |
| Champaign County, Somer township | 0.14470145 | 0.49462076 | 40.14241712 | -88.23744808 |
| DeKalb County, Sandwich township | 0.31118881 | 0.4943518 | 41.64321209 | -88.61704434 |
| DeKalb County, Sycamore township | 0.29209252 | 0.49381388 | 41.98427168 | -88.70283741 |
| Kankakee County, Ganeer township | 0.37452932 | 0.48547606 | 41.15944607 | -87.67337802 |
| Lake County, Antioch township | 0.33647122 | 0.48547606 | 42.48694224 | -88.09727524 |
| Kane County, Hampshire township | 0.32786444 | 0.4795589 | 42.15210738 | -88.50195336 |
| Lee County, South Dixon township | 0.22686928 | 0.47794513 | 41.82426269 | -89.47936343 |
| Vermilion County, Newell township | 0.11941904 | 0.47660032 | 40.14711654 | -87.61300989 |
| Ogle County, Dement township | 0.16568047 | 0.47471759 | 41.93583544 | -89.04281495 |
| McLean County, Dry Grove township | 0.18047337 | 0.47310382 | 40.50391133 | -89.06339837 |
| Stephenson County, Silver Creek township | 0.16568047 | 0.47068316 | 42.27783145 | -89.58972035 |
| Kane County, Virgil township | 0.19338354 | 0.46853147 | 41.90526448 | -88.49801122 |
| McHenry County, McHenry township | 0.36941904 | 0.46772458 | 42.3419491 | -88.2976294 |
| Whiteside County, Coloma township | 0.32436794 | 0.46637977 | 41.76792857 | -89.69711435 |
| Tazewell County, Pekin township | 0.30742335 | 0.46584185 | 40.55734267 | -89.65512719 |
| Kankakee County, Momence township | 0.37022593 | 0.46422808 | 41.15748859 | -87.6434662 |
| Lake County, Zion township | 0.42791824 | 0.46180742 | 42.44765652 | -87.82550166 |
| St. Clair County, Centreville township | 0.34991931 | 0.46073158 | 38.56799041 | -90.11804844 |
| Kane County, Big Rock township | 0.35153308 | 0.45804196 | 41.77161273 | -88.51234308 |
| Williamson County, West Marion precinct | 0.23224852 | 0.45589026 | 37.74615788 | -88.94895668 |
| Perry County, Du Quoin No. 4 precinct | 0.17374933 | 0.45131791 | 38.00868294 | -89.2408454 |
| Clark County, Martinsville township | 0.19903174 | 0.45051103 | 39.35859602 | -87.88092297 |
| McHenry County, Greenwood township | 0.30177515 | 0.44809037 | 42.32862288 | -88.45226362 |
| Edgar County, Paris township | 0.24488973 | 0.44513179 | 39.61566918 | -87.70444125 |
| Winnebago County, Rockton township | 0.08472297 | 0.44486283 | 42.47417674 | -89.05013432 |
| Kankakee County, Otto township | 0.13501883 | 0.44405594 | 41.05980927 | -87.86572089 |
| Whiteside County, Sterling township | 0.2027972 | 0.44244217 | 41.79022271 | -89.69510635 |
| DeKalb County, Squaw Grove township | 0.30150619 | 0.44244217 | 41.77859049 | -88.63313981 |
| Peoria County, Richwoods township | 0.43195266 | 0.44136633 | 40.73250928 | -89.57963496 |
| Peoria County, Kickapoo township | 0.33001614 | 0.44109736 | 40.78154444 | -89.66500217 |
| Rock Island County, Rock Island township | 0.43249059 | 0.43786982 | 41.49435997 | -90.56804512 |
| Madison County, Wood River township | 0.31952663 | 0.4373319 | 38.90052317 | -90.0620104 |
| Sangamon County, Williams township | 0.12426036 | 0.4373319 | 39.88342031 | -89.59644296 |
| Rock Island County, South Moline township | 0.40263583 | 0.43598709 | 41.48919931 | -90.44756784 |
| Clark County, Marshall township | 0.22700377 | 0.4351802 | 39.41371834 | -87.69824152 |
| LaSalle County, LaSalle township | 0.34064013 | 0.4351802 | 41.33425844 | -89.10226244 |
| McHenry County, Richmond township | 0.36915008 | 0.4338354 | 42.45338127 | -88.2852475 |
| LaSalle County, Ottawa township | 0.26788596 | 0.43275955 | 41.36561755 | -88.85760316 |
| Effingham County, Summit township | 0.11296396 | 0.43114578 | 39.11147553 | -88.58202926 |
| LaSalle County, Northville township | 0.23749328 | 0.43060785 | 41.62093969 | -88.61939012 |
| Bond County, Central township | 0.18652501 | 0.42738031 | 38.86669114 | -89.39438752 |
| Logan County, East Lincoln township | 0.12022593 | 0.42684239 | 40.16209256 | -89.36372687 |
| Rock Island County, Hampton township | 0.21167294 | 0.42657343 | 41.48789148 | -90.4174513 |
| McLean County, Mount Hope township | 0.09319527 | 0.42630446 | 40.30842805 | -89.17164648 |
| LaSalle County, Peru township | 0.29451318 | 0.4260355 | 41.35814965 | -89.13011192 |
| Vermilion County, Grant township | 0.24865519 | 0.4260355 | 40.46421196 | -87.66517684 |
| LaSalle County, Dayton township | 0.12937063 | 0.42280796 | 41.38620752 | -88.82530178 |
| Champaign County, Champaign township | 0.26533082 | 0.42227004 | 40.0757 | -88.24468709 |
| Grundy County, Goose Lake township | 0.24448628 | 0.42200108 | 41.3992669 | -88.27995734 |
| Madison County, Alton township | 0.39133943 | 0.42092523 | 38.90603981 | -90.17818052 |
| LaSalle County, Mendota township | 0.11915008 | 0.41796665 | 41.55864587 | -89.11048184 |
| Adams County, Quincy township | 0.35610543 | 0.41527703 | 39.96113306 | -91.3641406 |
| Coles County, Mattoon township | 0.23036579 | 0.41500807 | 39.47891002 | -88.36770491 |
| St. Clair County, Belleville city | 0.30325444 | 0.41366326 | 38.49422967 | -89.9790162 |
| Morgan County, Jacksonville No. 2 precinct | 0.41366326 | 0.41366326 | 39.73529337 | -90.22902134 |
| LaSalle County, Eagle township | 0.12130178 | 0.41285637 | 41.11898957 | -88.85246444 |
| Ford County, Drummer township | 0.13313609 | 0.41151157 | 40.46463945 | -88.38129939 |
| McHenry County, Coral township | 0.24448628 | 0.41151157 | 42.15386563 | -88.53231479 |
| Livingston County, Pontiac township | 0.25 | 0.4112426 | 40.88274952 | -88.63632999 |
| Sangamon County, Springfield township | 0.32544379 | 0.41070468 | 39.84188881 | -89.65923473 |
| Kankakee County, St. Anne township | 0.25389995 | 0.40909091 | 41.02793533 | -87.7189715 |
| Jackson County, Carbondale township | 0.2506724 | 0.4077461 | 37.71627016 | -89.21017777 |
| Franklin County, Benton township | 0.20266272 | 0.40693921 | 37.99084099 | -88.92552344 |
| Macon County, Whitmore township | 0.17025282 | 0.40693921 | 39.89154483 | -88.91554243 |
| Rock Island County, Blackhawk township | 0.30957504 | 0.40693921 | 41.44848188 | -90.54138813 |
| Madison County, Godfrey village | 0.22498655 | 0.40613233 | 38.92693335 | -90.14734542 |
| St. Clair County, Stookey township | 0.24072082 | 0.40613233 | 38.56660742 | -90.08913956 |
| Coles County, Lafayette township | 0.24045186 | 0.40586337 | 39.45488194 | -88.34083144 |
| Christian County, Pana township | 0.21664874 | 0.40586337 | 39.38640118 | -89.0825649 |
| Boone County, Poplar Grove township | 0.2175901 | 0.40559441 | 42.35107986 | -88.848066 |
| Fayette County, Vandalia township | 0.18840775 | 0.40505648 | 38.96209025 | -89.09376018 |
| Rock Island County, South Rock Island township | 0.37399139 | 0.40505648 | 41.47333211 | -90.5998807 |
| McDonough County, Macomb City township | 0.40102205 | 0.4042496 | 40.47024061 | -90.67588089 |
| Winnebago County, Winnebago township | 0.2049489 | 0.40263583 | 42.27993772 | -89.19191746 |
| Fayette County, Otego township | 0.20521786 | 0.40182894 | 38.97780185 | -88.97544074 |
| Bureau County, Princeton township | 0.16742873 | 0.40102205 | 41.39774911 | -89.45849444 |
| St. Clair County, East St. Louis township | 0.31603012 | 0.39967725 | 38.5916285 | -90.14517503 |
| Kendall County, Fox township | 0.30043034 | 0.39752555 | 41.63622036 | -88.49648208 |
| Morgan County, South Jacksonville No. 4 precinct | 0.3964497 | 0.3964497 | 39.69075683 | -90.23259179 |
| Winnebago County, Harlem township | 0.33082302 | 0.39591178 | 42.33873916 | -89.03306614 |
| Peoria County, Medina township | 0.17993545 | 0.39564282 | 40.80232691 | -89.63320156 |
| Grundy County, Erienna township | 0.19499731 | 0.39537386 | 41.36548625 | -88.46494284 |
| Champaign County, Rantoul township | 0.13609467 | 0.3951049 | 40.31667468 | -88.15892915 |
| Winnebago County, Roscoe township | 0.22525551 | 0.3929532 | 42.44859361 | -88.99136178 |
| Will County, Wilton township | 0.31065089 | 0.39187735 | 41.37890569 | -87.94971437 |
| Madison County, Chouteau township | 0.31253362 | 0.39133943 | 38.76556294 | -90.04390502 |
| Boone County, Caledonia township | 0.22162453 | 0.39133943 | 42.35275735 | -88.87852281 |
| McLean County, Money Creek township | 0.12735342 | 0.39107047 | 40.64645849 | -88.81032372 |
| McHenry County, Hartland township | 0.190156 | 0.38999462 | 42.33039316 | -88.48270082 |
| DeKalb County, Mayfield township | 0.14080151 | 0.3894567 | 41.98598248 | -88.73313093 |
| Madison County, Edwardsville township | 0.28886498 | 0.38891877 | 38.78360075 | -89.95510517 |
| Ogle County, Flagg township | 0.14254976 | 0.38596019 | 41.91517878 | -89.07530599 |
| Effingham County, Watson township | 0.0837816 | 0.38515331 | 39.08749581 | -88.55522087 |
| Tazewell County, Morton township | 0.24663798 | 0.38434642 | 40.61508435 | -89.47077193 |
| Montgomery County, South Litchfield township | 0.18047337 | 0.38407746 | 39.17116701 | -89.65936944 |
| Madison County, Fort Russell township | 0.21920387 | 0.38192577 | 38.899127 | -90.03297063 |
| Champaign County, Hensley township | 0.15895643 | 0.38058096 | 40.14604455 | -88.296457 |
| Jefferson County, Shiloh township | 0.20548682 | 0.38058096 | 38.30555491 | -88.95303507 |
| Henry County, Kewanee township | 0.20239376 | 0.37896719 | 41.24169946 | -89.92491238 |
| Rock Island County, Moline township | 0.34709521 | 0.37789134 | 41.51279916 | -90.47593282 |
| St. Clair County, Canteen township | 0.34521248 | 0.37789134 | 38.63339047 | -90.08373355 |
| Madison County, Jarvis township | 0.16917698 | 0.37681549 | 38.73623471 | -89.90085014 |
| Grundy County, Wauponsee township | 0.19378698 | 0.37654653 | 41.3414509 | -88.43731658 |
| St. Clair County, Mascoutah township | 0.18612157 | 0.37547068 | 38.55240514 | -89.80011188 |
| DeKalb County, Genoa township | 0.16944594 | 0.37358795 | 42.09569153 | -88.69116568 |
| Kankakee County, Aroma township | 0.16594944 | 0.37278107 | 41.12078537 | -87.76831925 |
| Morgan County, South Jacksonville No. 2 precinct | 0.3725121 | 0.3725121 | 39.71302502 | -90.23080712 |
| Tazewell County, Cincinnati township | 0.19419042 | 0.3725121 | 40.53655832 | -89.68681209 |
| LaSalle County, Rutland township | 0.25026896 | 0.37089833 | 41.36224955 | -88.79754469 |
| Sangamon County, Clear Lake township | 0.24865519 | 0.37009145 | 39.83588374 | -89.54159494 |
| McLean County, Dale township | 0.17280796 | 0.3690156 | 40.48165239 | -89.06556739 |
| McHenry County, Burton township | 0.3091716 | 0.36793975 | 42.42745398 | -88.22677612 |
| Knox County, Galesburg City township | 0.25766541 | 0.36713287 | 40.94893197 | -90.36997458 |
| St. Clair County, Caseyville township | 0.27541689 | 0.36686391 | 38.60834147 | -90.02768713 |
| LaSalle County, Otter Creek township | 0.1520979 | 0.36686391 | 41.13788479 | -88.79034122 |
| Madison County, Granite City township | 0.28778913 | 0.36632598 | 38.70155901 | -90.10727919 |
| Marion County, Centralia township | 0.12829478 | 0.3655191 | 38.51717226 | -89.1357926 |
| Iroquois County, Lovejoy township | 0.16863905 | 0.3655191 | 40.48839629 | -87.69220237 |
| Will County, Reed township | 0.24731038 | 0.36498117 | 41.28433496 | -88.2321985 |
| St. Clair County, Lebanon township | 0.17832168 | 0.36444325 | 38.59691852 | -89.79633378 |
| Williamson County, East Marion precinct | 0.11363636 | 0.36390533 | 37.74450105 | -88.9203929 |
| Henry County, Wethersfield township | 0.21785906 | 0.36256052 | 41.21941399 | -89.92683226 |
| Peoria County, Radnor township | 0.2239107 | 0.36229155 | 40.80381786 | -89.66301199 |
| LaSalle County, South Ottawa township | 0.2027972 | 0.36229155 | 41.34166705 | -88.8298432 |
| Tazewell County, Washington township | 0.23426573 | 0.36040882 | 40.70107432 | -89.40305156 |
| LaSalle County, Wallace township | 0.112695 | 0.35906401 | 41.38788857 | -88.85534117 |
| Franklin County, Tyrone township | 0.16581495 | 0.35879505 | 37.97517767 | -89.04227062 |
| Lee County, Dixon township | 0.1344809 | 0.35718128 | 41.84808368 | -89.50753632 |
| McHenry County, Chemung township | 0.16191501 | 0.35664336 | 42.42830689 | -88.62544063 |
| Madison County, Collinsville township | 0.2732652 | 0.35476062 | 38.67230425 | -89.96432413 |
| St. Clair County, Shiloh Valley township | 0.26748252 | 0.35476062 | 38.53447887 | -89.88866946 |
| McDonough County, Macomb township | 0.15169446 | 0.35476062 | 40.46898745 | -90.64619033 |
| Grundy County, Goodfarm township | 0.17832168 | 0.35287789 | 41.11529174 | -88.4013442 |
| Richland County, Olney township | 0.2584723 | 0.35233997 | 38.72342855 | -88.09735492 |
| Rock Island County, Coal Valley township | 0.24636902 | 0.35126412 | 41.4459033 | -90.48118752 |
| St. Clair County, O'Fallon township | 0.22417967 | 0.35099516 | 38.5983701 | -89.82524898 |
| Macon County, Long Creek township | 0.16191501 | 0.3507262 | 39.82313491 | -88.89273866 |
| St. Clair County, St. Clair township | 0.30742335 | 0.35018827 | 38.51507307 | -89.94829774 |
| McLean County, Old Town township | 0.18665949 | 0.34938139 | 40.45117419 | -88.91948137 |
| DeKalb County, Afton township | 0.13313609 | 0.34938139 | 41.87627391 | -88.77496465 |
| Livingston County, Reading township | 0.14497041 | 0.34911243 | 41.09504328 | -88.824803 |
| Logan County, West Lincoln township | 0.17684239 | 0.34830554 | 40.16519261 | -89.42280469 |
| Adams County, Riverside township | 0.232383 | 0.34749866 | 39.94100938 | -91.42454709 |
| Champaign County, Tolono township | 0.14349112 | 0.34588488 | 39.98674668 | -88.25431817 |
| Massac County, Jackson precinct | 0.10260893 | 0.345078 | 37.09789034 | -88.58314456 |
| Woodford County, Spring Bay township | 0.20790748 | 0.34427111 | 40.75326965 | -89.54783135 |
| McLean County, Empire township | 0.19472835 | 0.34319527 | 40.32982225 | -88.75294569 |
| McDonough County, Emmet township | 0.13743948 | 0.34211942 | 40.47148682 | -90.70557249 |
| Woodford County, Cruger township | 0.19069392 | 0.33862292 | 40.71866456 | -89.31167229 |
| McHenry County, Seneca township | 0.15868747 | 0.33808499 | 42.30810539 | -88.48511278 |
| Sangamon County, Curran township | 0.18612157 | 0.33808499 | 39.73501092 | -89.75715322 |
| Livingston County, Indian Grove township | 0.16729424 | 0.33700914 | 40.74225576 | -88.5311464 |
| Fayette County, Sefton township | 0.15814954 | 0.3359333 | 39.00004592 | -88.97328899 |
| Knox County, Galesburg township | 0.22592792 | 0.33566434 | 40.92796353 | -90.40162394 |
| McDonough County, Scotland township | 0.13851533 | 0.33512641 | 40.44670499 | -90.64786421 |
| Montgomery County, North Litchfield township | 0.19096288 | 0.33485745 | 39.19342317 | -89.65742169 |
| Morgan County, Literberry precinct | 0.09709521 | 0.33405056 | 39.7575619 | -90.22723446 |
| Tazewell County, Groveland township | 0.25053792 | 0.33351264 | 40.64040368 | -89.52820259 |
| Stephenson County, Harlem township | 0.10502959 | 0.33324368 | 42.32544547 | -89.64652831 |
| Jefferson County, Spring Garden township | 0.10193653 | 0.33324368 | 38.14654101 | -88.91053026 |
| Effingham County, Teutopolis township | 0.31374395 | 0.33297472 | 39.12846289 | -88.49245454 |
| Bureau County, Hall township | 0.23924153 | 0.33297472 | 41.31520826 | -89.16447467 |
| Massac County, Metropolis No. 4 precinct | 0.33270576 | 0.33270576 | 37.15099495 | -88.72035505 |
| Warren County, Monmouth township | 0.20252824 | 0.33243679 | 40.91596482 | -90.64236851 |
| Henry County, Geneseo township | 0.22229693 | 0.33109198 | 41.43121634 | -90.15016469 |
| Madison County, Foster township | 0.23050027 | 0.33055406 | 38.94919588 | -90.1455534 |
| Stephenson County, Lancaster township | 0.08203335 | 0.33001614 | 42.32394827 | -89.6160659 |
| Sangamon County, Divernon township | 0.13058096 | 0.32974718 | 39.59554337 | -89.65146788 |
| Fulton County, Canton township | 0.13905325 | 0.32894029 | 40.55383629 | -90.04329642 |
| Madison County, Venice township | 0.22915546 | 0.32894029 | 38.68204744 | -90.16698557 |
| Piatt County, Monticello township | 0.14201183 | 0.32813341 | 40.0508005 | -88.57362767 |
| Morgan County, Jacksonville No. 15 precinct | 0.3267886 | 0.3267886 | 39.73664387 | -90.258402 |
| Adams County, Ellington township | 0.19096288 | 0.32598171 | 39.96004163 | -91.33464924 |
| LaSalle County, Troy Grove township | 0.15761162 | 0.32598171 | 41.5363667 | -89.11266849 |
| LaSalle County, Fall River township | 0.15626681 | 0.32571275 | 41.3399801 | -88.79982411 |
| Clark County, Douglas township | 0.18652501 | 0.3246369 | 39.43594044 | -87.69568414 |
| Jefferson County, Mount Vernon township | 0.16917698 | 0.3246369 | 38.30223663 | -88.89547352 |
| Peoria County, West Peoria township | 0.32436794 | 0.32436794 | 40.64492891 | -89.61742754 |
| Franklin County, Denning township | 0.16931146 | 0.32383002 | 37.90186713 | -88.93406258 |
| Grundy County, Braceville township | 0.30016138 | 0.32383002 | 41.26389679 | -88.26462568 |
| Bond County, Pleasant Mound township | 0.14604626 | 0.32329209 | 38.88738894 | -89.3633454 |
| Sangamon County, Rochester township | 0.1366326 | 0.32275417 | 39.77061773 | -89.57697512 |
| LaSalle County, Earl township | 0.13259817 | 0.32248521 | 41.59336389 | -88.92530121 |
| Sangamon County, Woodside township | 0.28671329 | 0.3211404 | 39.7305865 | -89.66904634 |
| Jefferson County, Dodds township | 0.17267348 | 0.3211404 | 38.27999462 | -88.89762839 |
| Fayette County, La Clede township | 0.22054868 | 0.32006455 | 38.83266265 | -88.78538694 |
| LaSalle County, Waltham township | 0.08445401 | 0.32006455 | 41.37391441 | -89.00777423 |
| Macon County, Harristown township | 0.13582571 | 0.31979559 | 39.83138378 | -89.03969519 |
| Wayne County, Grover township | 0.24905863 | 0.31952663 | 38.38232221 | -88.36608543 |
| Lawrence County, Allison township | 0.12116729 | 0.31845078 | 38.6870083 | -87.54802728 |
| Macoupin County, Carlinville township | 0.25255514 | 0.31683701 | 39.27188541 | -89.88501914 |
| Clark County, Wabash township | 0.11699839 | 0.31629909 | 39.41176874 | -87.66905052 |
| Douglas County, Tuscola township | 0.17011834 | 0.3154922 | 39.78661247 | -88.27590108 |
| Alexander County, Cairo precinct | 0.24018289 | 0.31441635 | 37.02168964 | -89.18807027 |
| Madison County, Saline township | 0.18585261 | 0.31387843 | 38.79282102 | -89.69229946 |
| Grundy County, Felix township | 0.22807961 | 0.3119957 | 41.33067204 | -88.2572835 |
| Macon County, South Wheatland township | 0.17455621 | 0.31118881 | 39.80749775 | -89.01246631 |
| Franklin County, Ewing township | 0.13434642 | 0.31091985 | 38.12429845 | -88.91267603 |
| DeKalb County, Malta township | 0.14362561 | 0.31011296 | 41.92421984 | -88.83089041 |
| Ogle County, Oregon-Nashua township | 0.11538462 | 0.309844 | 42.01870676 | -89.33930394 |
| Hancock County, Montebello township | 0.26398601 | 0.30930608 | 40.38459503 | -91.33662222 |
| Moultrie County, Sullivan township | 0.21651425 | 0.30876815 | 39.58370313 | -88.62178031 |
| Fayette County, Wheatland township | 0.16245293 | 0.30849919 | 38.99673688 | -88.91517703 |
| Morgan County, Jacksonville No. 6 precinct | 0.30796127 | 0.30796127 | 39.71166807 | -90.20143661 |
| Edgar County, Symmes township | 0.17590102 | 0.30715438 | 39.57122217 | -87.70956179 |
| Champaign County, Mahomet township | 0.20575578 | 0.30715438 | 40.19770269 | -88.40976844 |
| Morgan County, Jacksonville No. 11 precinct | 0.30688542 | 0.30688542 | 39.71437508 | -90.26017873 |
| Winnebago County, Owen township | 0.15384615 | 0.30688542 | 42.36267226 | -89.06129425 |
| Vermilion County, Catlin township | 0.15021517 | 0.3063475 | 40.0843524 | -87.67975587 |
| Jo Daviess County, Dunleith township | 0.28483055 | 0.30607854 | 42.48246702 | -90.64387973 |
| DeKalb County, Kingston township | 0.16594944 | 0.30607854 | 42.09740529 | -88.72150727 |
| Jo Daviess County, Warren township | 0.22068316 | 0.30580958 | 42.49906742 | -89.99868685 |
| Wayne County, Berry township | 0.28214094 | 0.30554061 | 38.45964556 | -88.532096 |
| De Witt County, Clintonia township | 0.23574502 | 0.30554061 | 40.16352094 | -88.97766295 |
| Perry County, Du Quoin No. 9 precinct | 0.19392146 | 0.30500269 | 38.03093144 | -89.23880203 |
| Champaign County, Ludlow township | 0.13206025 | 0.30500269 | 40.31483205 | -88.12935702 |
| Iroquois County, Douglas township | 0.10543303 | 0.30500269 | 40.75406769 | -87.99069656 |
| Williamson County, Herrin precinct | 0.18181818 | 0.30446477 | 37.79558024 | -89.03045787 |
| Adams County, Melrose township | 0.19728349 | 0.30365788 | 39.89536189 | -91.39792086 |
| Washington County, Okawville township | 0.20306616 | 0.30338892 | 38.45003365 | -89.54794185 |
| St. Clair County, Freeburg township | 0.18558365 | 0.30311996 | 38.44687468 | -89.92495989 |
| McHenry County, Dunham township | 0.13609467 | 0.30311996 | 42.40427721 | -88.59733624 |
| Morgan County, Pisgah precinct | 0.15438408 | 0.30258203 | 39.733936 | -90.19964179 |
| Ogle County, Monroe township | 0.16151157 | 0.30231307 | 42.13644364 | -89.02277117 |
| Greene County, Kane township | 0.25201721 | 0.30123722 | 39.25074615 | -90.38479313 |
| Livingston County, Dwight township | 0.11995697 | 0.3006993 | 41.09481384 | -88.43364496 |
| Randolph County, Sparta precinct | 0.17105971 | 0.30043034 | 38.12368346 | -89.72085778 |
| Perry County, Du Quoin No. 6 precinct | 0.16890802 | 0.30016138 | 38.0070945 | -89.2121735 |
| Wayne County, Big Mound township | 0.17791824 | 0.29854761 | 38.38410858 | -88.39488455 |
| Saline County, Harrisburg township | 0.22377622 | 0.29747176 | 37.7206035 | -88.52063687 |
| Kane County, Burlington township | 0.17912856 | 0.29747176 | 42.0629782 | -88.51154277 |
| Wayne County, Indian Prairie township | 0.24919311 | 0.29693384 | 38.48362892 | -88.55866709 |
| McLean County, Randolph township | 0.15640129 | 0.29666487 | 40.32094615 | -88.9919217 |
| DeKalb County, Somonauk township | 0.15465304 | 0.29532006 | 41.64666559 | -88.67733443 |
| Cumberland County, Greenup township | 0.14739107 | 0.2950511 | 39.24206804 | -88.15803855 |
| Effingham County, Mound township | 0.18921463 | 0.29424422 | 39.05506688 | -88.76327166 |
| Lee County, May township | 0.14120495 | 0.29370629 | 41.6589355 | -89.31285476 |
| Woodford County, Metamora township | 0.16231845 | 0.29343733 | 40.78859418 | -89.36496483 |
| Effingham County, Jackson township | 0.08297472 | 0.29316837 | 39.089238 | -88.58430407 |
| Vermilion County, Blount township | 0.0979021 | 0.29316837 | 40.15103618 | -87.67198991 |
| Wayne County, Lamard township | 0.27945132 | 0.29263045 | 38.43212876 | -88.44789957 |
| Shelby County, Richland township | 0.19553523 | 0.29101668 | 39.35956966 | -88.61529783 |
| Schuyler County, Rushville township | 0.22175901 | 0.29020979 | 40.10744508 | -90.55468441 |
| LaSalle County, Dimmick township | 0.13232921 | 0.28994083 | 41.37880808 | -89.09789372 |
| Coles County, Seven Hickory township | 0.09830554 | 0.28886498 | 39.53291047 | -88.15583947 |
| Crawford County, Robinson township | 0.1732114 | 0.28859602 | 38.98957381 | -87.71751208 |
| Marion County, Salem township | 0.1810113 | 0.28832706 | 38.61860765 | -88.95187758 |
| Monroe County, Precinct 11 | 0.22915546 | 0.28752017 | 38.3912709 | -90.16118648 |
| Wayne County, Mount Erie township | 0.22754169 | 0.28698225 | 38.50483533 | -88.17907249 |
| Livingston County, Avoca township | 0.14066703 | 0.28698225 | 40.7645092 | -88.52880199 |
| Iroquois County, Onarga township | 0.10462614 | 0.2859064 | 40.71145639 | -88.02545288 |
| Peoria County, Chillicothe township | 0.16594944 | 0.2859064 | 40.92992616 | -89.50172534 |
| Champaign County, South Homer township | 0.11350188 | 0.2859064 | 40.034934 | -87.95250656 |
| Iroquois County, Middleport township | 0.12385691 | 0.28536848 | 40.78328921 | -87.74756517 |
| Perry County, Du Quoin No. 10 precinct | 0.17616998 | 0.28536848 | 37.98484637 | -89.21422424 |
| Pulaski County, Mound City precinct | 0.24623453 | 0.28429263 | 37.08845098 | -89.18197035 |
| Pike County, Newburg township | 0.14308768 | 0.28375471 | 39.60384578 | -90.79871143 |
| Schuyler County, Buena Vista township | 0.21086606 | 0.28267886 | 40.13099671 | -90.58253783 |
| Wabash County, Mount Carmel precinct | 0.16272189 | 0.2824099 | 38.41184334 | -87.78292372 |
| St. Clair County, Sugar Loaf township | 0.17482517 | 0.28133405 | 38.52757331 | -90.20831007 |
| De Witt County, Wapella township | 0.17791824 | 0.28079613 | 40.23028133 | -88.97109876 |
| Warren County, Hale township | 0.15344271 | 0.2802582 | 40.91721926 | -90.67224622 |
| Greene County, Carrollton township | 0.23883808 | 0.27945132 | 39.27432765 | -90.41226215 |
| Vermilion County, Georgetown township | 0.1105433 | 0.27918236 | 40.03598421 | -87.62603581 |
| Woodford County, Worth township | 0.19741797 | 0.27918236 | 40.79014796 | -89.39476376 |
| Monroe County, Precinct 30 | 0.27891339 | 0.27891339 | 38.36900878 | -90.16296013 |
| McHenry County, Hebron township | 0.20481442 | 0.27891339 | 42.48286815 | -88.40476498 |
| Shelby County, Prairie township | 0.14254976 | 0.27891339 | 39.33733025 | -88.61757022 |
| Madison County, Hamel township | 0.14349112 | 0.27864443 | 38.8891634 | -89.8297238 |
| Williamson County, Southern precinct | 0.07127488 | 0.27837547 | 37.61433427 | -88.99018717 |
| Pike County, Cincinnati township | 0.22834857 | 0.27837547 | 39.64365818 | -91.23739902 |
| Wayne County, Orchard township | 0.22781065 | 0.27810651 | 38.48537023 | -88.58751066 |
| Grundy County, Nettle Creek township | 0.13878429 | 0.27810651 | 41.41178111 | -88.49020979 |
| Winnebago County, Pecatonica township | 0.16917698 | 0.27810651 | 42.2894106 | -89.3745408 |
| Cass County, Panther Creek township | 0.21611081 | 0.2767617 | 40.04433604 | -90.14489156 |
| Shelby County, Tower Hill township | 0.13851533 | 0.27622378 | 39.38314294 | -89.02413772 |
| Morgan County, Arcadia precinct | 0.09117805 | 0.27622378 | 39.75891283 | -90.25662417 |
| Franklin County, Browning township | 0.1520979 | 0.27568585 | 37.99249653 | -88.95418273 |
| Clay County, Harter township | 0.24488973 | 0.27541689 | 38.61529183 | -88.51616925 |
| Winnebago County, Seward township | 0.09951587 | 0.27434104 | 42.28784964 | -89.34410014 |
| LaSalle County, Eden township | 0.0648198 | 0.27380312 | 41.26581573 | -89.07881111 |
| Clinton County, Breese township | 0.18827327 | 0.27353416 | 38.60580227 | -89.53417979 |
| McLean County, Towanda township | 0.16164605 | 0.27299623 | 40.4940206 | -88.88538761 |
| Macoupin County, Staunton township | 0.214766 | 0.27272727 | 38.99899952 | -89.79119654 |
| Wayne County, Bedford township | 0.20683163 | 0.27245831 | 38.47659619 | -88.44330659 |
| Wayne County, Keith township | 0.20077999 | 0.27218935 | 38.59305674 | -88.51844865 |
| Tazewell County, Elm Grove township | 0.10623991 | 0.27218935 | 40.55435068 | -89.59571416 |
| Alexander County, Cache precinct | 0.19445939 | 0.27218935 | 37.04553872 | -89.2143399 |
| Hancock County, Wythe township | 0.21490048 | 0.27192039 | 40.36120741 | -91.30842207 |
| Sangamon County, Chatham township | 0.12950511 | 0.27165143 | 39.68754749 | -89.70231219 |
| Iroquois County, Papineau township | 0.13528779 | 0.27165143 | 41.00569306 | -87.72157909 |
| Mason County, Lynchburg township | 0.22001076 | 0.2711135 | 40.11793727 | -90.28710257 |
| St. Clair County, Stites township | 0.27084454 | 0.27084454 | 38.65978517 | -90.16876452 |
| Peoria County, Hollis township | 0.2049489 | 0.27084454 | 40.58109844 | -89.68285781 |
| Woodford County, Clayton township | 0.07436794 | 0.27084454 | 40.90625834 | -89.05396318 |
| Fulton County, Astoria township | 0.22821409 | 0.27057558 | 40.21234555 | -90.39829018 |
| Hancock County, Appanoose township | 0.22404519 | 0.27003765 | 40.58631119 | -91.35322309 |
| Kankakee County, Pilot township | 0.14833244 | 0.27003765 | 41.05074928 | -88.07738785 |
| Hardin County, McFarlan precinct | 0.23561054 | 0.26976869 | 37.45998247 | -88.28931176 |
| Wayne County, Jasper township | 0.24327595 | 0.26949973 | 38.40276078 | -88.33496 |
| Pulaski County, Mounds precinct | 0.21893491 | 0.26923077 | 37.13295732 | -89.17789768 |
| LaSalle County, Manlius township | 0.24825175 | 0.26923077 | 41.33316201 | -88.67976222 |
| Wayne County, Hickory Hill township | 0.18477676 | 0.26896181 | 38.46313388 | -88.58976437 |
| McLean County, Lexington township | 0.11135019 | 0.26815492 | 40.64476797 | -88.78059832 |
| LaSalle County, Brookfield township | 0.08821947 | 0.26788596 | 41.30917375 | -88.65207548 |
| Crawford County, Lamotte township | 0.18760086 | 0.26681011 | 39.00593732 | -87.62788967 |
| Ogle County, Scott township | 0.12197418 | 0.26519634 | 42.0951229 | -89.08793265 |
| Clay County, Stanford township | 0.20306616 | 0.26465842 | 38.67847094 | -88.4514855 |
| Fayette County, Bear Grove township | 0.08647122 | 0.26438946 | 38.96531567 | -89.15185246 |
| Shelby County, Shelbyville township | 0.20817644 | 0.26438946 | 39.38868322 | -88.72984384 |
| Calhoun County, Hamburg precinct | 0.23130715 | 0.26412049 | 39.28956464 | -90.76269621 |
| Hancock County, Warsaw township | 0.25094137 | 0.26385153 | 40.36555685 | -91.42705398 |
| Christian County, Taylorville township | 0.14416353 | 0.26277569 | 39.55492646 | -89.30188286 |
| Rock Island County, Canoe Creek township | 0.16393222 | 0.2619688 | 41.61231618 | -90.1956201 |
| Woodford County, Olio township | 0.1366326 | 0.26169984 | 40.71709139 | -89.28190515 |
| Clinton County, Germantown township | 0.13232921 | 0.26143088 | 38.56280912 | -89.56701369 |
| Madison County, Moro township | 0.14214632 | 0.26116192 | 38.9436486 | -90.02931692 |
| Mercer County, Eliza township | 0.21732114 | 0.26116192 | 41.24737833 | -91.09893281 |
| Schuyler County, Hickory township | 0.19445939 | 0.26089295 | 40.14154808 | -90.31487199 |
| Schuyler County, Bainbridge township | 0.19405594 | 0.26035503 | 40.08516847 | -90.55637828 |
| Henry County, Annawan township | 0.16083916 | 0.26035503 | 41.39627623 | -89.88137223 |
| St. Clair County, Smithton township | 0.1958042 | 0.25981711 | 38.47197027 | -89.98084768 |
| Will County, Custer township | 0.12896719 | 0.25927918 | 41.28067422 | -88.1722405 |
| Calhoun County, Belleview precinct | 0.24045186 | 0.2584723 | 39.35882858 | -90.81633127 |
| Iroquois County, Prairie Green township | 0.17267348 | 0.25820334 | 40.50671719 | -87.63032601 |
| Alexander County, McClure precinct | 0.21247983 | 0.25820334 | 37.30595794 | -89.47600803 |
| Bond County, Old Ripley township | 0.09548144 | 0.25793437 | 38.82984558 | -89.54341894 |
| Grundy County, Mazon township | 0.20683163 | 0.25793437 | 41.25062705 | -88.41690574 |
| Madison County, Marine township | 0.19956966 | 0.25685853 | 38.77350988 | -89.75219098 |
| Greene County, Bluffdale township | 0.2204142 | 0.25658956 | 39.27822927 | -90.49985678 |
| Clark County, Casey township | 0.20750403 | 0.25658956 | 39.33827009 | -87.91258708 |
| Hancock County, Rocky Run-Wilcox township | 0.1633943 | 0.25658956 | 40.36447992 | -91.39739465 |
| Lee County, Alto township | 0.07342657 | 0.25658956 | 41.88962485 | -89.01699347 |
| Fulton County, Farmington township | 0.1451049 | 0.2563206 | 40.68609006 | -90.00231658 |
| Pike County, Ross township | 0.21342119 | 0.25605164 | 39.40818874 | -90.9301672 |
| Mason County, Bath township | 0.13555675 | 0.25605164 | 40.06660645 | -90.14307084 |
| Greene County, Woodville township | 0.22552448 | 0.25578268 | 39.25466708 | -90.4723578 |
| Moultrie County, East Nelson township | 0.12305003 | 0.25551372 | 39.60246903 | -88.56091821 |
| Piatt County, Sangamon township | 0.09077461 | 0.25497579 | 40.07129936 | -88.54183704 |
| Sangamon County, Gardner township | 0.10919849 | 0.25470683 | 39.82258825 | -89.72000857 |
| Lawrence County, Lawrence township | 0.13394298 | 0.25443787 | 38.74126736 | -87.68746423 |
| Edwards County, Shelby No. 1 precinct | 0.19284562 | 0.25389995 | 38.52522655 | -88.14784285 |
| Sangamon County, Island Grove township | 0.1274879 | 0.25363098 | 39.74224736 | -89.90402183 |
| Edwards County, Albion No. 1 precinct | 0.23023131 | 0.25309306 | 38.38445368 | -88.04697461 |
| Washington County, Nashville township | 0.17092523 | 0.2528241 | 38.35183305 | -89.3829003 |
| Perry County, Du Quoin No. 5 precinct | 0.13986014 | 0.25228618 | 38.01026467 | -89.26951853 |
| Coles County, Hutton township | 0.09924691 | 0.25228618 | 39.46437411 | -88.13386834 |
| LaSalle County, Miller township | 0.07356105 | 0.25228618 | 41.37769675 | -88.6751265 |
| Greene County, Linder township | 0.21947283 | 0.25201721 | 39.29395931 | -90.35213392 |
| LaSalle County, Utica township | 0.15909091 | 0.25174825 | 41.35327874 | -89.0400166 |
| Randolph County, Red Bud precinct | 0.14604626 | 0.25094137 | 38.20485405 | -90.00273905 |
| Brown County, Mount Sterling township | 0.13582571 | 0.25094137 | 39.98254707 | -90.77124416 |
| Schuyler County, Woodstock township | 0.20441097 | 0.25094137 | 40.08770948 | -90.6154394 |
| Vermilion County, Oakwood township | 0.1422808 | 0.25094137 | 40.11046967 | -87.73611896 |
| Cass County, Sangamon Valley township | 0.20266272 | 0.25040344 | 40.04570643 | -90.17439722 |
| Cass County, Bluff Springs township | 0.20884884 | 0.25013448 | 40.01055841 | -90.38446006 |
| McLean County, Chenoa township | 0.16917698 | 0.25013448 | 40.66871545 | -88.80806981 |
| Madison County, Pin Oak township | 0.20467994 | 0.24986552 | 38.80301006 | -89.89525919 |
| Schuyler County, Browning township | 0.22458311 | 0.24878967 | 40.16515307 | -90.34265973 |
| Calhoun County, Hardin precinct | 0.22108661 | 0.24852071 | 39.14971408 | -90.62658234 |
| Will County, Wesley township | 0.17267348 | 0.24852071 | 41.27883333 | -88.14226383 |
| Carroll County, Salem township | 0.13286713 | 0.24852071 | 42.07231165 | -89.97444261 |
| Stephenson County, Florence township | 0.12883271 | 0.24852071 | 42.28231991 | -89.68104931 |
| Cass County, Beardstown township | 0.19674556 | 0.24852071 | 40.01187262 | -90.41395631 |
| Fayette County, Lone Grove township | 0.19419042 | 0.24798279 | 38.85659175 | -88.81217725 |
| Edwards County, Albion No. 2 precinct | 0.22781065 | 0.24771382 | 38.3863147 | -88.07576665 |
| Fayette County, Avena township | 0.17186659 | 0.24744486 | 39.03620543 | -88.82362322 |
| Logan County, Broadwell township | 0.06105433 | 0.24744486 | 40.11912629 | -89.39737632 |
| McDonough County, Chalmers township | 0.14967725 | 0.2471759 | 40.42567518 | -90.67920911 |
| Hardin County, Rock precinct | 0.1851802 | 0.24690694 | 37.51939311 | -88.16852145 |
| Pike County, Pleasant Hill township | 0.21987628 | 0.24690694 | 39.40458946 | -90.84239275 |
| Jo Daviess County, Elizabeth township | 0.18195266 | 0.24690694 | 42.33157023 | -90.25788741 |
| Kankakee County, Pembroke township | 0.16702528 | 0.24663798 | 41.11103948 | -87.61884531 |
| Randolph County, Chester precinct | 0.16702528 | 0.24636902 | 37.88178447 | -89.79896526 |
| Carroll County, Mount Carroll township | 0.09548144 | 0.24636902 | 42.09461781 | -89.97251448 |
| Clark County, Parker township | 0.1253362 | 0.24556213 | 39.36049605 | -87.91009816 |
| Kendall County, Lisbon township | 0.18370091 | 0.24529317 | 41.47325206 | -88.39286396 |
| Macoupin County, Shaws Point township | 0.17038731 | 0.24529317 | 39.28981047 | -89.79555944 |
| Macoupin County, Virden township | 0.13690156 | 0.24529317 | 39.49014429 | -89.7783352 |
| Edwards County, Salem No. 1 precinct | 0.197149 | 0.24502421 | 38.51593825 | -88.00361503 |
| Madison County, St. Jacob township | 0.1422808 | 0.24502421 | 38.72899744 | -89.75600513 |
| Sangamon County, New Berlin township | 0.13474987 | 0.24502421 | 39.7199837 | -89.90590822 |
| Rock Island County, Drury township | 0.21100054 | 0.24502421 | 41.33543243 | -91.06266293 |
| Wayne County, Massilon township | 0.23359333 | 0.24475524 | 38.41774822 | -88.19094934 |
| Macoupin County, Bird township | 0.19445939 | 0.24475524 | 39.27474038 | -89.94339278 |
| Grundy County, Maine township | 0.17912856 | 0.24448628 | 41.2675142 | -88.32457418 |
| Mercer County, New Boston township | 0.21328671 | 0.24394836 | 41.22276118 | -91.04045251 |
| De Witt County, Texas township | 0.14416353 | 0.2436794 | 40.13961766 | -88.95032852 |
| Henry County, Oxford township | 0.19284562 | 0.24341044 | 41.17313955 | -90.3821732 |
| Wayne County, Barnhill township | 0.19889726 | 0.24314147 | 38.26936966 | -88.34890513 |
| Calhoun County, Gilead precinct | 0.22996235 | 0.24260355 | 39.15097121 | -90.65573317 |
| Vermilion County, Ross township | 0.15613233 | 0.24260355 | 40.37723424 | -87.70516727 |
| Montgomery County, Hillsboro township | 0.13542227 | 0.24206563 | 39.1398681 | -89.48654729 |
| Hancock County, Sonora township | 0.19553523 | 0.24179666 | 40.47484985 | -91.360491 |
| Massac County, Adkins precinct | 0.2415277 | 0.2415277 | 37.17494413 | -88.74652039 |
| Macon County, Oakley township | 0.1486014 | 0.2415277 | 39.86595978 | -88.85892672 |
| Iroquois County, Loda township | 0.14201183 | 0.2415277 | 40.5353833 | -88.075071 |
| Knox County, Knox township | 0.12157074 | 0.2415277 | 40.94627423 | -90.31020458 |
| Clark County, Auburn township | 0.20467994 | 0.24125874 | 39.39730238 | -87.78835306 |
| Macoupin County, Cahokia township | 0.17132867 | 0.24125874 | 39.17412764 | -89.71765307 |
| Livingston County, Owego township | 0.12116729 | 0.24098978 | 40.87927938 | -88.57669099 |
| Edgar County, Stratton township | 0.14147391 | 0.24072082 | 39.58954342 | -87.64847601 |
| Tazewell County, Hopedale township | 0.13636364 | 0.24045186 | 40.41315752 | -89.45948703 |
| Grundy County, Norman township | 0.13690156 | 0.24045186 | 41.30396704 | -88.56207708 |
| Edwards County, Albion No. 3 precinct | 0.20548682 | 0.24018289 | 38.36408798 | -88.07817235 |
| DeKalb County, Pierce township | 0.11067778 | 0.24018289 | 41.87286528 | -88.71447199 |
| Moultrie County, Marrowbone township | 0.16473911 | 0.24018289 | 39.63334877 | -88.70513356 |
| Effingham County, St. Francis township | 0.14887036 | 0.23964497 | 39.1266992 | -88.46335785 |
| Union County, Anna District 2 precinct | 0.20293168 | 0.23937601 | 37.44926688 | -89.23461646 |
| Jackson County, Bradley township | 0.18652501 | 0.23937601 | 37.89067699 | -89.53934989 |
| Jersey County, Jersey township | 0.18652501 | 0.23937601 | 39.13676714 | -90.3351307 |
| Marion County, Kinmundy township | 0.12789134 | 0.23910705 | 38.81211011 | -88.81656923 |
| Mason County, Mason City township | 0.17576654 | 0.23910705 | 40.20254331 | -89.71631235 |
| Knox County, Truro township | 0.18074233 | 0.23883808 | 40.88940521 | -90.04504481 |
| Vermilion County, McKendree township | 0.09440559 | 0.23856912 | 40.05624266 | -87.59398254 |
| Henry County, Alba township | 0.16608392 | 0.23856912 | 41.41999864 | -89.9095096 |
| Cass County, Chandlerville township | 0.18639053 | 0.23856912 | 40.06522872 | -90.11355718 |
| Coles County, Ashmore township | 0.09144701 | 0.23830016 | 39.50698756 | -88.09977567 |
| Jasper County, South Muddy township | 0.13488435 | 0.2380312 | 38.93631177 | -88.27866854 |
| Macoupin County, Dorchester township | 0.17697687 | 0.2380312 | 39.00045233 | -89.82027145 |
| Marion County, Meacham township | 0.16554599 | 0.23776224 | 38.81042236 | -88.78759105 |
| Pike County, Spring Creek township | 0.20817644 | 0.23776224 | 39.40215566 | -90.78388133 |
| Henderson County, Oquawka township | 0.20710059 | 0.23722431 | 40.92819388 | -90.94119231 |
| Hardin County, Cave-In-Rock precinct | 0.21490048 | 0.23695535 | 37.49716135 | -88.1708713 |
| Douglas County, Camargo township | 0.16379774 | 0.23641743 | 39.86824976 | -88.14875868 |
| Edwards County, Salem No. 2 precinct | 0.20871436 | 0.23614847 | 38.5178094 | -88.03245766 |
| Jo Daviess County, East Galena township | 0.18706294 | 0.23614847 | 42.40519273 | -90.4049028 |
| Wayne County, Elm River township | 0.21678322 | 0.23587951 | 38.46945552 | -88.3279684 |
| Macon County, South Macon township | 0.14349112 | 0.23507262 | 39.71685903 | -88.99176589 |
| Massac County, Jefferson precinct | 0.17374933 | 0.23480366 | 37.17323964 | -88.71817993 |
| Whiteside County, Hopkins township | 0.10516407 | 0.23480366 | 41.79317108 | -89.75557196 |
| Hamilton County, McLeansboro township | 0.17912856 | 0.2345347 | 38.07989035 | -88.54180994 |
| Edwards County, Browns precinct | 0.20723507 | 0.23426573 | 38.38258592 | -88.01818403 |
| Clinton County, Sugar Creek township | 0.18746638 | 0.23399677 | 38.61032086 | -89.62092359 |
| Richland County, Noble township | 0.18948359 | 0.23399677 | 38.68814691 | -88.24673594 |
| Brown County, Missouri township | 0.20172136 | 0.23372781 | 40.02710391 | -90.76799365 |
| Calhoun County, Richwood precinct | 0.17065627 | 0.23345885 | 39.063139 | -90.69140565 |
| Calhoun County, Crater precinct | 0.22243141 | 0.23345885 | 39.26106515 | -90.61831972 |
| Macoupin County, Honey Point township | 0.1422808 | 0.23318989 | 39.19638472 | -89.71572312 |
| Mason County, Havana township | 0.1422808 | 0.23292093 | 40.28655211 | -90.06559197 |
| Bureau County, Walnut township | 0.17778375 | 0.232383 | 41.56134241 | -89.59459484 |
| Clay County, Louisville township | 0.1668908 | 0.23211404 | 38.75045643 | -88.53141067 |
| Ford County, Patton township | 0.11592254 | 0.23184508 | 40.44827301 | -88.11460119 |
| Pike County, Pittsfield township | 0.15438408 | 0.23130715 | 39.6050627 | -90.82804868 |
| Clinton County, Brookside township | 0.19688004 | 0.23103819 | 38.54421814 | -89.22034293 |
| Hardin County, Stone Church precinct | 0.21893491 | 0.23076923 | 37.46358515 | -88.34619257 |
| Fulton County, Vermont township | 0.14523938 | 0.23050027 | 40.27785491 | -90.36344249 |
| Iroquois County, Sheldon township | 0.09359871 | 0.22996235 | 40.76956655 | -87.53931131 |
| Greene County, White Hall township | 0.1958042 | 0.22969338 | 39.43020434 | -90.40020548 |
| Perry County, Du Quoin No. 12 precinct | 0.1810113 | 0.22915546 | 37.96259812 | -89.21627375 |
| Hancock County, Carthage township | 0.16662184 | 0.2288865 | 40.4213373 | -91.1259356 |
| Pike County, Atlas township | 0.18639053 | 0.22861754 | 39.52193601 | -90.98095413 |
| Hardin County, Rosiclare precinct | 0.22767617 | 0.22834857 | 37.41911389 | -88.35078106 |
| Richland County, Decker township | 0.19338354 | 0.22834857 | 38.6232692 | -88.28271309 |
| Macoupin County, Gillespie township | 0.13125336 | 0.22807961 | 39.08948436 | -89.81268872 |
| Fulton County, Buckheart township | 0.09696073 | 0.22781065 | 40.53156103 | -90.04516077 |
| Ogle County, Byron township | 0.08754707 | 0.22727273 | 42.12703625 | -89.26790425 |
| Saline County, Cottage township | 0.17186659 | 0.22700377 | 37.71708265 | -88.46355018 |
| Jackson County, Somerset township | 0.09467456 | 0.22700377 | 37.79090252 | -89.34697707 |
| Alexander County, Sandusky precinct | 0.20979021 | 0.2267348 | 37.20920511 | -89.34196091 |
| Monroe County, Precinct 21 | 0.1570737 | 0.2267348 | 38.41489951 | -90.1882616 |
| Montgomery County, Nokomis township | 0.16500807 | 0.22646584 | 39.28633396 | -89.29742383 |
| Champaign County, Sidney township | 0.10368478 | 0.22646584 | 40.02019275 | -88.07278742 |
| Sangamon County, Mechanicsburg township | 0.07073696 | 0.22592792 | 39.85509799 | -89.48076212 |
| Macoupin County, Mount Olive township | 0.19029048 | 0.22565896 | 39.06431112 | -89.75638429 |
| Alexander County, Thebes precinct | 0.20199032 | 0.22538999 | 37.21076353 | -89.37032866 |
| Calhoun County, Carlin precinct | 0.21086606 | 0.22512103 | 39.33039065 | -90.67179556 |
| Macon County, Illini township | 0.08821947 | 0.22512103 | 39.89813648 | -89.03321894 |
| Hancock County, Nauvoo township | 0.22512103 | 0.22512103 | 40.54281244 | -91.38586547 |
| Schuyler County, Frederick township | 0.20118343 | 0.22485207 | 40.0354542 | -90.4417264 |
| Madison County, Olive township | 0.14389457 | 0.22458311 | 38.93222738 | -89.79689796 |
| Menard County, Petersburg North No. 14 precinct | 0.21785906 | 0.22458311 | 40.03025206 | -89.84989804 |
| Marshall County, Bennington township | 0.12358795 | 0.22458311 | 41.0001885 | -89.13485267 |
| Madison County, Omphghent township | 0.1492738 | 0.22431415 | 38.97819446 | -89.82216423 |
| Montgomery County, Raymond township | 0.14725659 | 0.22431415 | 39.32245992 | -89.55810638 |
| Moultrie County, Jonathan Creek township | 0.15357719 | 0.22431415 | 39.62470924 | -88.55862001 |
| Grundy County, Garfield township | 0.14093599 | 0.22431415 | 41.19893558 | -88.30189931 |
| Pike County, Kinderhook township | 0.16352878 | 0.22404519 | 39.68596979 | -91.17569748 |
| Vermilion County, Middlefork township | 0.13232921 | 0.22404519 | 40.35887831 | -87.76691224 |
| Wayne County, Leech township | 0.19849381 | 0.22296934 | 38.37510933 | -88.25090305 |
| Peoria County, Princeville township | 0.16164605 | 0.22296934 | 40.92109383 | -89.772483 |
| Washington County, Covington township | 0.14981173 | 0.22270038 | 38.44083331 | -89.37485357 |
| Cumberland County, Neoga township | 0.10543303 | 0.22270038 | 39.30458475 | -88.44481389 |
| Union County, Anna District 4 precinct | 0.14712211 | 0.22216245 | 37.47151813 | -89.23258662 |
| Clay County, Clay City township | 0.18921463 | 0.22216245 | 38.69356949 | -88.33349046 |
| Pulaski County, Villa Ridge precinct | 0.2056213 | 0.22189349 | 37.17906053 | -89.20217227 |
| Washington County, Beaucoup township | 0.13421194 | 0.22162453 | 38.3502779 | -89.3540913 |
| Cass County, Virginia township | 0.17146315 | 0.22162453 | 39.93571669 | -90.21289951 |
| Macoupin County, Brushy Mound township | 0.16299086 | 0.22135557 | 39.25105581 | -89.91607645 |
| Rock Island County, Bowling township | 0.11672942 | 0.22135557 | 41.40516305 | -90.5749307 |
| Greene County, Rockbridge township | 0.16648736 | 0.22108661 | 39.31087036 | -90.23355479 |
| McLean County, Downs township | 0.10166756 | 0.22108661 | 40.35546113 | -88.80989112 |
| Macoupin County, Polk township | 0.16931146 | 0.22081764 | 39.25247944 | -89.94525492 |
| Henry County, Clover township | 0.119688 | 0.22054868 | 41.17048699 | -90.32221307 |
| Montgomery County, Zanesville township | 0.14792899 | 0.22054868 | 39.32396741 | -89.58730638 |
| Pulaski County, America precinct | 0.17832168 | 0.22027972 | 37.13135379 | -89.14956359 |
| Greene County, Roodhouse township | 0.1640667 | 0.22027972 | 39.47342365 | -90.3674744 |
| Hardin County, Monroe precinct | 0.19943518 | 0.22001076 | 37.52849097 | -88.31083305 |
| Monroe County, Precinct 6 | 0.13421194 | 0.2197418 | 38.22711404 | -90.00092085 |
| Montgomery County, Pitman township | 0.11161915 | 0.21947283 | 39.34921865 | -89.64375388 |
| Richland County, Madison township | 0.18827327 | 0.21893491 | 38.58448663 | -88.02517934 |
| Jersey County, Richwood township | 0.20172136 | 0.21893491 | 39.19043949 | -90.53578047 |
| Jersey County, English township | 0.1732114 | 0.21866595 | 39.16167799 | -90.39169088 |
| White County, Carmi township | 0.15626681 | 0.21839699 | 38.07881964 | -88.16666126 |
| McDonough County, Bushnell township | 0.13636364 | 0.21839699 | 40.55303132 | -90.52058457 |
| Whiteside County, Montmorency township | 0.13757396 | 0.21839699 | 41.74563495 | -89.69912106 |
| Brown County, Lee township | 0.11995697 | 0.21839699 | 40.00604931 | -90.79912009 |
| Wayne County, Arrington township | 0.19203873 | 0.21812803 | 38.37070389 | -88.54116666 |
| Marion County, Stevenson township | 0.09601937 | 0.21812803 | 38.63919396 | -88.92081588 |
| Alexander County, Olive Branch precinct | 0.18854223 | 0.21785906 | 37.18695001 | -89.34395098 |
| Effingham County, Mason township | 0.11457773 | 0.21785906 | 38.91651219 | -88.68949864 |
| Douglas County, Bourbon township | 0.15478752 | 0.21785906 | 39.72886621 | -88.42974257 |
| Monroe County, Precinct 25 | 0.2175901 | 0.2175901 | 38.43852215 | -90.21535417 |
| Henry County, Colona township | 0.12305003 | 0.2175901 | 41.46427929 | -90.38911597 |
| Monroe County, Precinct 36 | 0.21732114 | 0.21732114 | 38.43716196 | -90.18649452 |
| McHenry County, Marengo township | 0.12493276 | 0.21732114 | 42.24824562 | -88.61394874 |
| Pope County, Eddyville No. 6 precinct | 0.1584185 | 0.21705218 | 37.59584766 | -88.67660478 |
| Monroe County, Precinct 28 | 0.1422808 | 0.21705218 | 38.38989802 | -90.13234643 |
| Greene County, Wrights township | 0.18934911 | 0.21705218 | 39.42888723 | -90.37094668 |
| De Witt County, Santa Anna township | 0.15653577 | 0.21705218 | 40.28019202 | -88.66874722 |
| Marshall County, Henry township | 0.15532544 | 0.21705218 | 41.12422407 | -89.3634985 |
| Mercer County, Rivoli township | 0.1401291 | 0.21705218 | 41.1534724 | -90.44389278 |
| De Witt County, Harp township | 0.1514255 | 0.21678322 | 40.18246456 | -88.91640156 |
| Henderson County, Bald Bluff township | 0.18585261 | 0.21678322 | 40.99147804 | -90.84666803 |
| Vermilion County, Butler township | 0.09117805 | 0.21651425 | 40.38111123 | -87.76434238 |
| Union County, Union precinct | 0.0718128 | 0.21624529 | 37.34739457 | -89.41525424 |
| Calhoun County, Point precinct | 0.17348037 | 0.21624529 | 38.90222204 | -90.58656802 |
| Jo Daviess County, West Galena township | 0.18154922 | 0.21597633 | 42.40650484 | -90.43541672 |
| Wayne County, Zif township | 0.18585261 | 0.21570737 | 38.57699621 | -88.25853976 |
| Piatt County, Willow Branch township | 0.10812265 | 0.21570737 | 40.03202479 | -88.63486681 |
| Jefferson County, Pendleton township | 0.12641205 | 0.21516945 | 38.27496819 | -88.81132186 |
| Shelby County, Moweaqua township | 0.18154922 | 0.21490048 | 39.6501134 | -88.99826423 |
| Sangamon County, Fancy Creek township | 0.10274341 | 0.21490048 | 39.88491909 | -89.62587166 |
| Henderson County, Rozetta township | 0.17186659 | 0.21490048 | 40.94809726 | -90.87980762 |
| Pope County, Golconda No. 2 precinct | 0.19069392 | 0.21463152 | 37.42446486 | -88.43606185 |
| Marion County, Tonti township | 0.09319527 | 0.21436256 | 38.66474368 | -88.97650298 |
| Cumberland County, Union township | 0.09494352 | 0.21436256 | 39.28097876 | -88.06576573 |
| Peoria County, Rosefield township | 0.12506724 | 0.21436256 | 40.74142514 | -89.75833384 |
| McHenry County, Alden township | 0.17267348 | 0.21436256 | 42.42482568 | -88.56447478 |
| Pike County, Barry township | 0.17442173 | 0.2140936 | 39.68483433 | -91.14632161 |
| Iroquois County, Danforth township | 0.08391608 | 0.2140936 | 40.79855461 | -87.9856668 |
| Tazewell County, Deer Creek township | 0.10018827 | 0.21382464 | 40.65498873 | -89.37744443 |
| Edgar County, Grandview township | 0.15034965 | 0.21355568 | 39.55287122 | -87.77061654 |
| Pulaski County, Pulaski precinct | 0.19338354 | 0.21328671 | 37.20290499 | -89.22850185 |
| Sangamon County, Auburn township | 0.11847768 | 0.21328671 | 39.59997998 | -89.73940881 |
| Randolph County, Kaskaskia precinct | 0.1753631 | 0.21301775 | 37.8875485 | -89.91351319 |
| Jersey County, Elsah township | 0.1851802 | 0.21301775 | 38.93504752 | -90.3216798 |
| Greene County, Rubicon township | 0.15922539 | 0.21301775 | 39.35268066 | -90.17155391 |
| Crawford County, Oblong township | 0.12627757 | 0.21274879 | 38.99728553 | -87.83361299 |
| Saline County, Mountain township | 0.17912856 | 0.21247983 | 37.60057665 | -88.38938208 |
| Jefferson County, McClellan township | 0.112695 | 0.2119419 | 38.28331191 | -88.95517261 |
| Shelby County, Rose township | 0.12560516 | 0.2119419 | 39.41601034 | -88.81526046 |
| Carroll County, Savanna township | 0.12022593 | 0.21140398 | 42.07930358 | -90.12625528 |
| Fulton County, Woodland township | 0.17146315 | 0.21113502 | 40.20837066 | -90.30955773 |
| Pike County, Pleasant Vale township | 0.1640667 | 0.21086606 | 39.66255437 | -91.14782276 |
| Alexander County, Tamms precinct | 0.19150081 | 0.2105971 | 37.2283223 | -89.28322087 |
| Clinton County, Looking Glass township | 0.14362561 | 0.2105971 | 38.50202482 | -89.68837611 |
| Clay County, Songer township | 0.16675632 | 0.21032813 | 38.73170844 | -88.59157022 |
| Shelby County, Clarksburg township | 0.15303927 | 0.21032813 | 39.34590036 | -88.76350924 |
| Peoria County, Trivoli township | 0.08431953 | 0.21005917 | 40.70695531 | -89.97065744 |
| Whiteside County, Prophetstown township | 0.13179129 | 0.20979021 | 41.66806495 | -89.94856883 |
| Jackson County, Murphysboro township | 0.09655729 | 0.20952125 | 37.76708808 | -89.3204001 |
| White County, Hawthorne township | 0.08647122 | 0.20952125 | 38.09920825 | -88.13560884 |
| Edwards County, Ellery precinct | 0.18908015 | 0.20952125 | 38.39001654 | -88.1333551 |
| Stark County, Elmira township | 0.13569123 | 0.20952125 | 41.21654773 | -89.86685072 |
| Schuyler County, Birmingham township | 0.15075309 | 0.20925229 | 40.21140852 | -90.90287659 |
| Tazewell County, Tremont township | 0.12022593 | 0.20925229 | 40.52601674 | -89.47894439 |
| Jo Daviess County, Menominee township | 0.15586337 | 0.20925229 | 42.50225968 | -90.58103463 |
| White County, Gray township | 0.14470145 | 0.2084454 | 38.24736017 | -88.00396293 |
| Edgar County, Buck township | 0.16594944 | 0.2084454 | 39.59924758 | -87.79480356 |
| Hancock County, Augusta township | 0.19123185 | 0.2084454 | 40.25835425 | -90.95891161 |
| Jackson County, Makanda township | 0.09911243 | 0.20817644 | 37.67335935 | -89.24280528 |
| Clay County, Larkinsburg township | 0.15653577 | 0.20817644 | 38.84807995 | -88.66721196 |
| Moultrie County, Lowe township | 0.12869822 | 0.20790748 | 39.73240304 | -88.48842226 |
| De Witt County, Wilson township | 0.13206025 | 0.20790748 | 40.22696957 | -88.91198753 |
| Macon County, Blue Mound township | 0.10785368 | 0.20736955 | 39.80913324 | -89.04185127 |
| Menard County, Petersburg South No. 15 precinct | 0.16541151 | 0.20736955 | 40.00942543 | -89.88129379 |
| Marshall County, Hopewell township | 0.12641205 | 0.20736955 | 41.10038869 | -89.33566236 |
| Bureau County, Dover township | 0.10328133 | 0.20736955 | 41.42003097 | -89.4564215 |
| Union County, Jonesboro District 1 precinct | 0.09628833 | 0.20710059 | 37.4508503 | -89.26307359 |
| Menard County, Petersburg West No. 16 precinct | 0.20710059 | 0.20710059 | 40.0079864 | -89.8518088 |
| LaSalle County, Meriden township | 0.08364712 | 0.20710059 | 41.55539575 | -89.05024649 |
| Macoupin County, Barr township | 0.18437332 | 0.20683163 | 39.36942009 | -90.05283676 |
| Pike County, Pearl township | 0.19163529 | 0.20656267 | 39.41947772 | -90.66522666 |
| Mercer County, Keithsburg township | 0.17832168 | 0.20656267 | 41.1535201 | -90.98518235 |
| Randolph County, Prairie du Rocher precinct | 0.13125336 | 0.20629371 | 38.09911722 | -90.1267071 |
| Greene County, Walkerville township | 0.17670791 | 0.20629371 | 39.37241867 | -90.61003166 |
| Brown County, Ripley township | 0.18746638 | 0.20629371 | 40.045667 | -90.67781612 |
| Knox County, Elba township | 0.16541151 | 0.20629371 | 40.86712454 | -90.04691743 |
| Piatt County, Blue Ridge township | 0.13219473 | 0.20602474 | 40.23050715 | -88.58466561 |
| Hancock County, Prairie township | 0.16016676 | 0.20602474 | 40.40356874 | -91.2461423 |
| Coles County, Morgan township | 0.08687466 | 0.20575578 | 39.53106782 | -88.12658826 |
| Livingston County, Nevada township | 0.09830554 | 0.2049489 | 41.03157668 | -88.500554 |
| Randolph County, Walsh precinct | 0.11108123 | 0.2049489 | 38.10289633 | -89.75147728 |
| St. Clair County, Millstadt township | 0.16473911 | 0.2049489 | 38.43025961 | -90.04221248 |
| Cumberland County, Woodbury township | 0.12291555 | 0.2049489 | 39.20490299 | -88.27933439 |
| St. Clair County, Marissa township | 0.12910167 | 0.20467994 | 38.23643367 | -89.7401102 |
| Wayne County, Four Mile township | 0.15115654 | 0.20467994 | 38.35368278 | -88.62980527 |
| Marshall County, Evans township | 0.11135019 | 0.20467994 | 41.06375526 | -89.06854151 |
| McLean County, Funks Grove township | 0.09400215 | 0.20441097 | 40.3926194 | -89.07422994 |
| Edwards County, Bone Gap precinct | 0.18235611 | 0.20441097 | 38.42703692 | -88.01333359 |
| Madison County, Alhambra township | 0.11659494 | 0.20441097 | 38.88771283 | -89.80069311 |
| Henry County, Burns township | 0.1048951 | 0.20414201 | 41.24453854 | -89.98491763 |
| Union County, Anna District 7 precinct | 0.18020441 | 0.20414201 | 37.46992758 | -89.20412223 |
| Livingston County, Newtown township | 0.12452932 | 0.20414201 | 41.09335554 | -88.7948887 |
| Jo Daviess County, Apple River township | 0.1605702 | 0.20414201 | 42.48094942 | -90.09223389 |
| Henry County, Galva township | 0.16715976 | 0.20387305 | 41.15820253 | -90.05244916 |
| Grundy County, Greenfield township | 0.12668101 | 0.20387305 | 41.19712519 | -88.27195398 |
| Randolph County, Tilden precinct | 0.15357719 | 0.20360409 | 38.16671826 | -89.68830975 |
| Coles County, Paradise township | 0.12385691 | 0.20360409 | 39.41220357 | -88.37475744 |
| Schuyler County, Oakland township | 0.13690156 | 0.20360409 | 40.21496082 | -90.45745059 |
| Jo Daviess County, Rice township | 0.17401829 | 0.20360409 | 42.31327001 | -90.35114144 |
| Henderson County, Carman township | 0.18558365 | 0.20333513 | 40.73339937 | -91.07470549 |
| Ogle County, Rockvale township | 0.09628833 | 0.20306616 | 42.04100153 | -89.33717452 |
| Gallatin County, Eagle Creek township | 0.14725659 | 0.2027972 | 37.62102355 | -88.35859131 |
| Knox County, Haw Creek township | 0.09830554 | 0.2027972 | 40.8699056 | -90.10661071 |
| Moultrie County, Dora township | 0.11081226 | 0.20252824 | 39.79076458 | -88.71869128 |
| Vermilion County, South Ross township | 0.11457773 | 0.20252824 | 40.35694391 | -87.73733302 |
| McLean County, Hudson township | 0.13811189 | 0.20252824 | 40.58967657 | -88.99529254 |
| Peoria County, Akron township | 0.12910167 | 0.20252824 | 40.9196284 | -89.74262097 |
| Sangamon County, Ball township | 0.13555675 | 0.20225928 | 39.61780209 | -89.64950623 |
| Hancock County, Chili township | 0.16648736 | 0.20225928 | 40.21730317 | -91.05082677 |
| Massac County, East Brooklyn precinct | 0.0866057 | 0.20172136 | 37.12186968 | -88.60924724 |
| Richland County, Preston township | 0.15034965 | 0.20145239 | 38.76788304 | -88.09253004 |
| Greene County, Patterson township | 0.14550834 | 0.20145239 | 39.45508738 | -90.45701761 |
| White County, Phillips township | 0.10462614 | 0.20118343 | 38.15281084 | -87.92754606 |
| Moultrie County, Lovington township | 0.11915008 | 0.20118343 | 39.71888236 | -88.6374248 |
| Henderson County, Lomax township | 0.17132867 | 0.20118343 | 40.66652361 | -91.10911959 |
| Champaign County, St. Joseph township | 0.11659494 | 0.20091447 | 40.13136919 | -88.06045749 |
| Fulton County, Pleasant township | 0.15438408 | 0.20091447 | 40.27652866 | -90.33383746 |
| Union County, Anna District 1 precinct | 0.20064551 | 0.20064551 | 37.473102 | -89.26105222 |
| Saline County, Stonefort township | 0.17105971 | 0.20064551 | 37.61808825 | -88.67440349 |
| Marion County, Romine township | 0.12950511 | 0.20037655 | 38.51450638 | -88.70067833 |
| Schuyler County, Huntsville township | 0.17805272 | 0.20037655 | 40.16564506 | -90.8764828 |
| Ogle County, Pine Rock township | 0.12721893 | 0.20037655 | 41.99009728 | -89.22019086 |
| Madison County, Helvetia township | 0.12910167 | 0.20010758 | 38.72605504 | -89.69807522 |
| Fayette County, Wilberton township | 0.16487359 | 0.20010758 | 38.86658197 | -88.98617978 |
| Cass County, Newmansville township | 0.19351802 | 0.20010758 | 40.01511249 | -89.9992455 |
| LaSalle County, Osage township | 0.08176439 | 0.20010758 | 41.0175896 | -89.04302115 |
| Henry County, Cornwall township | 0.11807423 | 0.19983862 | 41.40196781 | -90.00164743 |
| Champaign County, Pesotum township | 0.08620226 | 0.19956966 | 39.96450883 | -88.25672221 |
| Jo Daviess County, Rush township | 0.1246638 | 0.19956966 | 42.4586305 | -90.09413429 |
| Mason County, Allens Grove township | 0.14308768 | 0.1993007 | 40.31090758 | -89.64729675 |
| Wayne County, Orel township | 0.1492738 | 0.19903174 | 38.35195136 | -88.60101244 |
| Livingston County, Pike township | 0.12506724 | 0.19876278 | 40.77490246 | -88.70744039 |
| Washington County, Hoyleton township | 0.11484669 | 0.19876278 | 38.39321435 | -89.32122783 |
| Marion County, Sandoval township | 0.11672942 | 0.19876278 | 38.56166418 | -89.13161235 |
| Union County, Dongola District 2 precinct | 0.07961269 | 0.19849381 | 37.37772598 | -89.15541087 |
| Peoria County, Logan township | 0.0922539 | 0.19849381 | 40.69687772 | -89.76224975 |
| Clark County, Johnson township | 0.16352878 | 0.19849381 | 39.25314074 | -87.98079364 |
| Union County, Balcom precinct | 0.08445401 | 0.19822485 | 37.42542584 | -89.20819766 |
| Perry County, Du Quoin No. 7 precinct | 0.12883271 | 0.19795589 | 38.02934252 | -89.21012152 |
| Monroe County, Precinct 20 | 0.18464228 | 0.19795589 | 38.30897703 | -90.31231671 |
| Washington County, Irvington township | 0.10516407 | 0.19795589 | 38.49653201 | -89.16674204 |
| Cumberland County, Spring Point township | 0.08351264 | 0.19741797 | 39.18627629 | -88.33993567 |
| Richland County, Bonpas township | 0.08741259 | 0.19741797 | 38.58261377 | -87.99631061 |
| Macoupin County, Shipman township | 0.11995697 | 0.19741797 | 39.12313751 | -90.0437857 |
| Clay County, Oskaloosa township | 0.15935987 | 0.197149 | 38.75394504 | -88.58930877 |
| Menard County, Sandridge No. 8 precinct | 0.15572889 | 0.197149 | 40.05251794 | -89.8479861 |
| McDonough County, Blandinsville township | 0.12694997 | 0.197149 | 40.61373806 | -90.90387754 |
| Bureau County, Wyanet township | 0.15021517 | 0.197149 | 41.35927086 | -89.58280343 |
| Knox County, Rio township | 0.10112964 | 0.19688004 | 41.14952652 | -90.35397582 |
| Hancock County, Pontoosuc township | 0.17428725 | 0.19688004 | 40.60418346 | -91.23273453 |
| Montgomery County, Audubon township | 0.13232921 | 0.19661108 | 39.2386768 | -89.24322368 |
| Bureau County, Mineral township | 0.09548144 | 0.19607316 | 41.39483571 | -89.85130648 |
| Brown County, Pea Ridge township | 0.15357719 | 0.19607316 | 40.07410494 | -90.8237963 |
| Peoria County, Jubilee township | 0.12668101 | 0.1958042 | 40.8126168 | -89.8419006 |
| Iroquois County, Belmont township | 0.10314685 | 0.1958042 | 40.76105063 | -87.75015483 |
| Fayette County, Loudon township | 0.15572889 | 0.19553523 | 39.17133848 | -88.83952554 |
| Wabash County, Compton precinct | 0.1492738 | 0.19526627 | 38.37883021 | -87.96060729 |
| Woodford County, El Paso township | 0.14981173 | 0.19526627 | 40.72488056 | -89.01186916 |
| Putnam County, Magnolia township | 0.09185046 | 0.19526627 | 41.12266252 | -89.33355944 |
| Macoupin County, Scottville township | 0.12802582 | 0.19499731 | 39.43760303 | -90.07659293 |
| Peoria County, Brimfield township | 0.15384615 | 0.19499731 | 40.8823278 | -89.89578562 |
| Mercer County, Abington township | 0.14940828 | 0.19499731 | 41.07944061 | -90.81021556 |
| Franklin County, Goode township | 0.08606778 | 0.19472835 | 38.08803349 | -89.06044506 |
| Henderson County, Biggsville township | 0.16097364 | 0.19472835 | 40.88121902 | -90.88463976 |
| Menard County, Greenview No. 6 precinct | 0.17482517 | 0.19445939 | 40.11495135 | -89.75365846 |
| Shelby County, Oconee township | 0.15922539 | 0.19419042 | 39.34190565 | -89.08682289 |
| Mason County, Salt Creek township | 0.15976331 | 0.19419042 | 40.18027771 | -89.71826925 |
| McHenry County, Riley township | 0.10476062 | 0.19392146 | 42.18138379 | -88.62104562 |
| White County, Mill Shoals township | 0.10718128 | 0.19392146 | 38.24713755 | -88.3512244 |
| McDonough County, Prairie City township | 0.1274879 | 0.19392146 | 40.61729033 | -90.45593365 |
| Menard County, Petersburg East No. 13 precinct | 0.13582571 | 0.19338354 | 40.02880566 | -89.82040512 |
| Menard County, Indian Creek No. 7 precinct | 0.17105971 | 0.19338354 | 40.09268621 | -89.75560137 |
| Livingston County, Esmen township | 0.15814954 | 0.19338354 | 40.97350946 | -88.65691601 |
| Pike County, Martinsburg township | 0.14658419 | 0.19311458 | 39.49609671 | -90.8946147 |
| Macon County, Friends Creek township | 0.08162991 | 0.19311458 | 39.99607995 | -88.78670443 |
| Edwards County, Shelby No. 2 precinct | 0.17711135 | 0.19311458 | 38.43076795 | -88.07095087 |
| Marion County, Odin township | 0.14281872 | 0.19311458 | 38.62353521 | -89.0385992 |
| Clark County, Dolson township | 0.15236686 | 0.19311458 | 39.39922408 | -87.81754141 |
| Menard County, Atterberry No. 10 precinct | 0.18370091 | 0.19311458 | 40.03312415 | -89.90888743 |
| McDonough County, Lamoine township | 0.1577461 | 0.19311458 | 40.34510822 | -90.893328 |
| Mercer County, Millersburg township | 0.16958042 | 0.19311458 | 41.19213711 | -90.83201365 |
| Macoupin County, Girard township | 0.14900484 | 0.19257665 | 39.46788451 | -89.78025372 |
| McLean County, Gridley township | 0.13340506 | 0.19257665 | 40.67039957 | -88.83780587 |
| Union County, Mill Creek precinct | 0.14107047 | 0.19230769 | 37.34116332 | -89.30157829 |
| Jasper County, Wade township | 0.13084992 | 0.19230769 | 38.99570359 | -88.15541849 |
| Champaign County, Philo township | 0.10018827 | 0.19230769 | 39.98492671 | -88.2248795 |
| Lee County, Willow Creek township | 0.12210866 | 0.19230769 | 41.73199729 | -89.00235929 |
| Saline County, Independence township | 0.16016676 | 0.19176977 | 37.67437288 | -88.49660956 |
| Macoupin County, Western Mound township | 0.17495966 | 0.19176977 | 39.32766864 | -90.11492048 |
| Monroe County, Precinct 14 | 0.13421194 | 0.19150081 | 38.39399639 | -90.2188698 |
| Warren County, Point Pleasant township | 0.14080151 | 0.19150081 | 40.71662311 | -90.68729476 |
| Wabash County, Bellmont precinct | 0.15895643 | 0.19150081 | 38.40105461 | -87.95816548 |
| DeKalb County, Clinton township | 0.11377084 | 0.19150081 | 41.76487463 | -88.78646671 |
| Richland County, Claremont township | 0.13421194 | 0.19123185 | 38.73818893 | -87.97921508 |
| Whiteside County, Albany township | 0.16447015 | 0.19123185 | 41.76977523 | -90.21291106 |
| Randolph County, Brewerville precinct | 0.13528779 | 0.19096288 | 38.02677127 | -90.01724516 |
| Clinton County, Meridian township | 0.10018827 | 0.19069392 | 38.5632714 | -89.16049976 |
| Woodford County, Montgomery township | 0.13259817 | 0.19069392 | 40.62486675 | -89.23086948 |
| Shelby County, Windsor township | 0.12210866 | 0.19042496 | 39.44852826 | -88.60619431 |
| Schuyler County, Camden township | 0.1662184 | 0.19042496 | 40.11744619 | -90.79102691 |
| Clay County, Hoosier township | 0.13905325 | 0.190156 | 38.74517271 | -88.44457395 |
| Washington County, Ashley township | 0.15734266 | 0.18988704 | 38.31695667 | -89.15454101 |
| Kendall County, Big Grove township | 0.12076385 | 0.18988704 | 41.47857849 | -88.48306629 |
| Putnam County, Granville township | 0.08633674 | 0.18934911 | 41.29612821 | -89.22665327 |
| Effingham County, West township | 0.1753631 | 0.18934911 | 38.98834527 | -88.76992044 |
| Henry County, Cambridge township | 0.10906401 | 0.18908015 | 41.29883437 | -90.19130008 |
| Williamson County, Blairsville precinct | 0.14712211 | 0.18908015 | 37.77820595 | -89.11829335 |
| Jefferson County, Webber township | 0.1098709 | 0.18908015 | 38.3194491 | -88.80695882 |
| Stark County, Essex township | 0.14322216 | 0.18908015 | 41.05622566 | -89.79062201 |
| Livingston County, Odell township | 0.1070468 | 0.18881119 | 41.01107514 | -88.53278786 |
| Christian County, Rosamond township | 0.10866057 | 0.18881119 | 39.38963206 | -89.14099731 |
| Livingston County, Eppards Point township | 0.14712211 | 0.18881119 | 40.83823508 | -88.64095773 |
| McLean County, Martin township | 0.11444325 | 0.18854223 | 40.56772141 | -88.60922295 |
| Franklin County, Frankfort township | 0.10301237 | 0.18827327 | 37.90020685 | -88.90543892 |
| St. Clair County, New Athens township | 0.12775686 | 0.18827327 | 38.33272236 | -89.87655572 |
| LaSalle County, Deer Park township | 0.05782679 | 0.1880043 | 41.30709554 | -89.01441016 |
| Macoupin County, Chesterfield township | 0.15021517 | 0.1880043 | 39.23723198 | -90.09298068 |
| Clark County, Anderson township | 0.17267348 | 0.18773534 | 39.35092768 | -87.76423762 |
| Christian County, Prairieton township | 0.08848843 | 0.18773534 | 39.65175213 | -89.02758473 |
| Johnson County, Vienna No. 3 precinct | 0.12977407 | 0.18746638 | 37.40750053 | -88.89535745 |
| Jersey County, Rosedale township | 0.13959118 | 0.18746638 | 39.08037203 | -90.57328259 |
| Winnebago County, Burritt township | 0.11350188 | 0.18719742 | 42.30223721 | -89.18973283 |
| Fulton County, Waterford township | 0.14174287 | 0.18665949 | 40.31021387 | -90.09335367 |
| Hancock County, La Harpe township | 0.15344271 | 0.18665949 | 40.6149309 | -90.93363289 |
| Hamilton County, South Crouch township | 0.14523938 | 0.18639053 | 38.12610866 | -88.56599796 |
| Hancock County, Bear Creek township | 0.1662184 | 0.18639053 | 40.35899096 | -91.24911158 |
| Mercer County, Mercer township | 0.16325982 | 0.18639053 | 41.18846422 | -90.74202637 |
| Edgar County, Young America township | 0.13743948 | 0.18612157 | 39.80120674 | -87.80129028 |
| Warren County, Lenox township | 0.11256052 | 0.18612157 | 40.87138623 | -90.6457387 |
| Stark County, Penn township | 0.15613233 | 0.18612157 | 41.09489516 | -89.66695873 |
| Union County, Lick Creek precinct | 0.04289941 | 0.18585261 | 37.48736514 | -89.11667141 |
| Washington County, Plum Hill township | 0.11350188 | 0.18585261 | 38.38023817 | -89.49617333 |
| LaSalle County, Ophir township | 0.12573964 | 0.18585261 | 41.50920554 | -89.02455975 |
| Fulton County, Orion township | 0.10610543 | 0.18558365 | 40.55102625 | -89.98387044 |
| Warren County, Sumner township | 0.12049489 | 0.18558365 | 41.05592413 | -90.78191297 |
| Perry County, Pinckneyville No. 8 precinct | 0.16702528 | 0.18531469 | 38.10862049 | -89.43364342 |
| De Witt County, Rutledge township | 0.14120495 | 0.18531469 | 40.28190835 | -88.69831842 |
| Jo Daviess County, Nora township | 0.11081226 | 0.18531469 | 42.49623053 | -89.9375961 |
| Adams County, Lima township | 0.15599785 | 0.18504572 | 40.1806908 | -91.26095171 |
| Massac County, Benton precinct | 0.08391608 | 0.18477676 | 37.26392254 | -88.73784057 |
| Madison County, Leef township | 0.13300161 | 0.18477676 | 38.83733174 | -89.68844308 |
| Christian County, Assumption township | 0.09655729 | 0.18477676 | 39.5627579 | -89.03619455 |
| Union County, Stokes precinct | 0.06589564 | 0.18423884 | 37.44447666 | -89.1492524 |
| Jackson County, Degognia township | 0.16312534 | 0.18423884 | 37.76314809 | -89.66536161 |
| Fulton County, Isabel township | 0.11740183 | 0.18423884 | 40.28794125 | -90.09519607 |
| Lee County, Palmyra township | 0.06562668 | 0.18423884 | 41.84960551 | -89.53778946 |
| Williamson County, Carterville precinct | 0.09655729 | 0.18396988 | 37.75434175 | -89.09179461 |
| Sangamon County, Pawnee township | 0.08284024 | 0.18396988 | 39.59255131 | -89.59284664 |
| Randolph County, Palestine precinct | 0.13824637 | 0.18370091 | 37.99597252 | -89.84695338 |
| Edwards County, Dixon precinct | 0.17213556 | 0.18370091 | 38.31963433 | -88.08297939 |
| Wabash County, Lick Prairie precinct | 0.16191501 | 0.18370091 | 38.48995159 | -87.94838342 |
| Macoupin County, South Palmyra township | 0.14550834 | 0.18370091 | 39.41254729 | -90.0199259 |
| Washington County, Pilot Knob township | 0.1542496 | 0.18343195 | 38.26283192 | -89.39092753 |
| Warren County, Swan township | 0.07880581 | 0.18343195 | 40.7153724 | -90.65750109 |
| Jo Daviess County, Guilford township | 0.16837009 | 0.18343195 | 42.37755337 | -90.28469575 |
| Rock Island County, Andalusia township | 0.1394567 | 0.18316299 | 41.45480476 | -90.69190875 |
| Ogle County, Pine Creek township | 0.09548144 | 0.18289403 | 42.02182282 | -89.39995193 |
| Shelby County, Ash Grove township | 0.12130178 | 0.18289403 | 39.40231262 | -88.58153848 |
| Peoria County, Hallock township | 0.11807423 | 0.18289403 | 40.93144806 | -89.53158665 |
| Washington County, Bolo township | 0.13528779 | 0.18262507 | 38.26127865 | -89.36215326 |
| McLean County, West township | 0.09440559 | 0.18262507 | 40.30244122 | -88.66646078 |
| Peoria County, Millbrook township | 0.13367402 | 0.18262507 | 40.90887533 | -89.98344508 |
| Henry County, Munson township | 0.11403981 | 0.18262507 | 41.40892362 | -90.15201845 |
| McLean County, Yates township | 0.1281603 | 0.18235611 | 40.75093138 | -88.67996178 |
| Clinton County, Carlyle township | 0.15451856 | 0.18235611 | 38.64108184 | -89.35667681 |
| Fayette County, Sharon township | 0.09655729 | 0.18208714 | 39.0065824 | -89.08952866 |
| Pike County, Hadley township | 0.11296396 | 0.18208714 | 39.70250226 | -91.02728919 |
| Edgar County, Edgar township | 0.11726735 | 0.18208714 | 39.68428391 | -87.72604995 |
| Rock Island County, Buffalo Prairie township | 0.12641205 | 0.18208714 | 41.35064774 | -90.88068728 |
| Coles County, Humboldt township | 0.09494352 | 0.18181818 | 39.63096151 | -88.29260499 |
| Marshall County, Saratoga township | 0.11646046 | 0.18181818 | 41.11568333 | -89.63501774 |
| Randolph County, Rockwood precinct | 0.13797741 | 0.18181818 | 37.83287748 | -89.7168576 |
| Fulton County, Lewistown township | 0.14039806 | 0.18181818 | 40.40206896 | -90.14527621 |
| Hamilton County, Dahlgren township | 0.15303927 | 0.18128026 | 38.1775037 | -88.67638724 |
| Clay County, Xenia township | 0.15451856 | 0.18128026 | 38.64276196 | -88.60060228 |
| Shelby County, Big Spring township | 0.12600861 | 0.18128026 | 39.33385441 | -88.55920434 |
| Logan County, Atlanta township | 0.13851533 | 0.1810113 | 40.26710028 | -89.23505765 |
| DeKalb County, Paw Paw township | 0.10570199 | 0.1810113 | 41.63289925 | -88.83036463 |
| Cass County, Philadelphia township | 0.14174287 | 0.18074233 | 39.93298722 | -90.15397819 |
| Monroe County, Precinct 10 | 0.09897795 | 0.18047337 | 38.43987558 | -90.24421489 |
| Whiteside County, Lyndon township | 0.09494352 | 0.18047337 | 41.69036053 | -89.94664358 |
| Kankakee County, Salina township | 0.14349112 | 0.18020441 | 41.13975239 | -88.06738894 |
| Brown County, Cooperstown township | 0.10973642 | 0.17966649 | 40.00111211 | -90.68112345 |
| Morgan County, Chapin precinct | 0.12547068 | 0.17939753 | 39.76817646 | -90.46238263 |
| Ogle County, Marion township | 0.08418505 | 0.17939753 | 42.10315521 | -89.23970065 |
| Christian County, Buckhart township | 0.1070468 | 0.17939753 | 39.67243079 | -89.40887028 |
| Hancock County, Walker township | 0.14819796 | 0.17939753 | 40.20408658 | -91.28906523 |
| Warren County, Ellison township | 0.15330823 | 0.17912856 | 40.74015431 | -90.71543092 |
| Henry County, Andover township | 0.12399139 | 0.17912856 | 41.30019503 | -90.22133165 |
| Hamilton County, Crouch township | 0.14322216 | 0.1788596 | 38.24892742 | -88.37997019 |
| Bureau County, Selby township | 0.0746369 | 0.1788596 | 41.38840535 | -89.27816984 |
| Rock Island County, Cordova township | 0.13891877 | 0.1788596 | 41.68192713 | -90.25047057 |
| Edgar County, Embarrass township | 0.15277031 | 0.17859064 | 39.71612532 | -87.87006662 |
| Menard County, Sugar Grove No. 5 precinct | 0.10247445 | 0.17859064 | 40.06895322 | -89.72803551 |
| Champaign County, Scott township | 0.10556751 | 0.17859064 | 40.10873132 | -88.419211 |
| Jefferson County, Casner township | 0.12049489 | 0.17832168 | 38.31048181 | -89.03938709 |
| Ogle County, White Rock township | 0.05204411 | 0.17832168 | 42.00433256 | -89.06645593 |
| Edgar County, Shiloh township | 0.1401291 | 0.17805272 | 39.74025658 | -87.8968802 |
| Lee County, Hamilton township | 0.11377084 | 0.17805272 | 41.58513153 | -89.62270136 |
| Pulaski County, Ullin precinct | 0.12910167 | 0.17805272 | 37.24741086 | -89.22445072 |
| Jefferson County, Grand Prairie township | 0.10005379 | 0.17805272 | 38.47268022 | -89.13996776 |
| Iroquois County, Pigeon Grove township | 0.06145777 | 0.17778375 | 40.56858012 | -87.89205821 |
| Mercer County, North Henderson township | 0.11605702 | 0.17778375 | 41.13118122 | -90.44564666 |
| Clay County, Pixley township | 0.13703604 | 0.17751479 | 38.73982805 | -88.29987425 |
| Edwards County, French Creek precinct | 0.15034965 | 0.17751479 | 38.317775 | -88.05421331 |
| Shelby County, Herrick township | 0.16137708 | 0.17751479 | 39.24473421 | -88.94953261 |
| Williamson County, Stonefort precinct | 0.08996772 | 0.17697687 | 37.62151051 | -88.73142378 |
| Rock Island County, Rural township | 0.09709521 | 0.17697687 | 41.40388521 | -90.54484774 |
| Pulaski County, Olmsted precinct | 0.14174287 | 0.17697687 | 37.17585826 | -89.14546924 |
| Shelby County, Okaw township | 0.11108123 | 0.17670791 | 39.43656498 | -88.78381739 |
| Knox County, Persifer township | 0.06495428 | 0.17670791 | 40.89218719 | -90.10475677 |
| Whiteside County, Garden Plain township | 0.14550834 | 0.17643895 | 41.79071336 | -90.18082493 |
| Jasper County, North Muddy township | 0.10247445 | 0.17590102 | 39.02523698 | -88.26920633 |
| Jo Daviess County, Stockton township | 0.09709521 | 0.17590102 | 42.34285419 | -90.01217753 |
| Coles County, North Okaw township | 0.09682625 | 0.17590102 | 39.52338153 | -88.36299599 |
| Hancock County, Pilot Grove township | 0.11000538 | 0.17590102 | 40.46591387 | -91.12288674 |
| Johnson County, Vienna No. 2 precinct | 0.09386767 | 0.17563206 | 37.40583101 | -88.86692498 |
| Jersey County, Mississippi township | 0.14093599 | 0.17563206 | 39.04503582 | -90.28387189 |
| Henderson County, Media township | 0.15115654 | 0.17563206 | 40.76613382 | -90.80321761 |
| Macoupin County, North Palmyra township | 0.12856374 | 0.1753631 | 39.4348103 | -90.01808262 |
| Shelby County, Todds Point township | 0.10892953 | 0.1753631 | 39.52384 | -88.7456483 |
| Montgomery County, Butler Grove township | 0.13488435 | 0.1753631 | 39.18892965 | -89.56997852 |
| McLean County, Danvers township | 0.09709521 | 0.1753631 | 40.53262752 | -89.1799659 |
| Jefferson County, Elk Prairie township | 0.0718128 | 0.17509414 | 38.14820003 | -88.93924984 |
| Stephenson County, Rock Run township | 0.11013986 | 0.17509414 | 42.33557171 | -89.40074805 |
| Adams County, Northeast township | 0.15640129 | 0.17482517 | 40.12465612 | -90.96832627 |
| Warren County, Tompkins township | 0.15034965 | 0.17482517 | 40.87263984 | -90.67559769 |
| Jo Daviess County, Woodbine township | 0.10341581 | 0.17482517 | 42.3511723 | -90.19506941 |
| De Witt County, Creek township | 0.09776762 | 0.17455621 | 40.13629636 | -88.89129306 |
| Henderson County, Stronghurst township | 0.1309844 | 0.17455621 | 40.72991978 | -91.01509285 |
| Saline County, Raleigh township | 0.09037117 | 0.17428725 | 37.78555554 | -88.48530759 |
| Washington County, Du Bois township | 0.13582571 | 0.17428725 | 38.29631161 | -89.18540082 |
| Warren County, Roseville township | 0.11888112 | 0.17428725 | 40.73765905 | -90.65582391 |
| Stephenson County, Ridott township | 0.07826789 | 0.17374933 | 42.31326776 | -89.40286612 |
| Macon County, Mount Zion township | 0.10018827 | 0.17348037 | 39.77863924 | -88.89713983 |
| Christian County, Mosquito township | 0.11161915 | 0.17348037 | 39.82038905 | -89.24758297 |
| McDonough County, Mound township | 0.1042227 | 0.1732114 | 40.5320316 | -90.55201379 |
| Mason County, Kilbourne township | 0.12318451 | 0.17294244 | 40.12645448 | -89.98990799 |
| Mason County, Manito township | 0.13555675 | 0.17267348 | 40.40734982 | -89.78758406 |
| Woodford County, Roanoke township | 0.13878429 | 0.17267348 | 40.80298666 | -89.21385891 |
| Champaign County, Ogden township | 0.09628833 | 0.17240452 | 40.12763135 | -88.00147284 |
| Randolph County, Central precinct | 0.11296396 | 0.17240452 | 38.05691524 | -89.72655522 |
| Cumberland County, Crooked Creek township | 0.13528779 | 0.17213556 | 39.26060636 | -88.09734404 |
| Henderson County, Gladstone township | 0.14456697 | 0.17213556 | 40.88241661 | -90.91450701 |
| Marshall County, Steuben township | 0.0929263 | 0.17213556 | 41.0382269 | -89.43167182 |
| Perry County, Willisville precinct | 0.14658419 | 0.17186659 | 37.95895124 | -89.56215526 |
| Woodford County, Cazenovia township | 0.09736417 | 0.17186659 | 40.91907698 | -89.29273249 |
| Massac County, West Brooklyn precinct | 0.11457773 | 0.17159763 | 37.12359971 | -88.63756569 |
| Menard County, Tallula No. 11 precinct | 0.1274879 | 0.17159763 | 39.9914338 | -89.97162896 |
| Franklin County, Barren township | 0.11242604 | 0.17132867 | 38.08640283 | -89.03174622 |
| Douglas County, Arcola township | 0.11888112 | 0.17105971 | 39.70127251 | -88.34410218 |
| Clark County, Orange township | 0.16447015 | 0.17105971 | 39.24746999 | -87.89339669 |
| Jasper County, Willow Hill township | 0.12291555 | 0.17105971 | 39.00864947 | -88.00781008 |
| Winnebago County, Durand township | 0.13125336 | 0.17079075 | 42.42167229 | -89.33126284 |
| Tazewell County, Boynton township | 0.09588488 | 0.17079075 | 40.39089327 | -89.46152979 |
| Tazewell County, Mackinaw township | 0.11108123 | 0.17079075 | 40.54210438 | -89.35811264 |
| Kankakee County, Essex township | 0.09386767 | 0.17079075 | 41.19530779 | -88.24201019 |
| Pope County, Golconda No. 1 precinct | 0.1316568 | 0.17079075 | 37.36128272 | -88.49966832 |
| Stark County, Valley township | 0.10664336 | 0.17052179 | 41.05329597 | -89.73078438 |
| Massac County, Grant precinct | 0.12157074 | 0.17052179 | 37.17664199 | -88.77486214 |
| Hamilton County, Beaver Creek township | 0.13851533 | 0.17052179 | 38.20624396 | -88.41331717 |
| Wabash County, Lancaster precinct | 0.10274341 | 0.17052179 | 38.51217571 | -87.94593419 |
| Logan County, Mount Pulaski township | 0.11350188 | 0.17052179 | 40.00158939 | -89.28974183 |
| Hancock County, St. Mary township | 0.13932222 | 0.17052179 | 40.28299137 | -91.01657531 |
| Mercer County, Suez township | 0.13501883 | 0.17025282 | 41.09302948 | -90.59889352 |
| Saline County, East Eldorado township | 0.10274341 | 0.16998386 | 37.80602577 | -88.45446634 |
| Marion County, Alma township | 0.10462614 | 0.16998386 | 38.66143632 | -88.91865748 |
| Clark County, Melrose township | 0.13044648 | 0.16998386 | 39.26203629 | -87.77436168 |
| Hancock County, Harmony township | 0.15075309 | 0.16998386 | 40.28531681 | -91.07581542 |
| Stark County, Toulon township | 0.14066703 | 0.16998386 | 41.0814085 | -89.84852499 |
| Monroe County, Precinct 16 | 0.1697149 | 0.1697149 | 38.34674662 | -90.16473271 |
| Iroquois County, Ridgeland township | 0.10745024 | 0.16944594 | 40.7133238 | -88.05518731 |
| Stephenson County, Rock Grove township | 0.1338085 | 0.16944594 | 42.44865114 | -89.4206488 |
| Adams County, Clayton township | 0.14658419 | 0.16917698 | 40.10355099 | -90.99943614 |
| Pike County, Levee township | 0.10919849 | 0.16890802 | 39.69044279 | -91.29321011 |
| Mason County, Quiver township | 0.10166756 | 0.16890802 | 40.35056761 | -90.00077383 |
| Marshall County, Lacon township | 0.13717052 | 0.16890802 | 41.01440811 | -89.40384655 |
| Lee County, East Grove township | 0.14618074 | 0.16863905 | 41.62365692 | -89.49801442 |
| McDonough County, Colchester township | 0.15250134 | 0.16863905 | 40.45167427 | -90.76659567 |
| Livingston County, Sunbury township | 0.10866057 | 0.16863905 | 41.01975032 | -88.68216853 |
| Logan County, Chester township | 0.0718128 | 0.16863905 | 40.1160144 | -89.33833759 |
| Gallatin County, Bowlesville township | 0.14846692 | 0.16837009 | 37.67501023 | -88.15203302 |
| Menard County, Oakford No. 9 precinct | 0.16083916 | 0.16837009 | 40.08191701 | -89.99364648 |
| Randolph County, Ellis Grove precinct | 0.12372243 | 0.16810113 | 37.99741232 | -89.87563396 |
| White County, Burnt Prairie township | 0.09252286 | 0.16810113 | 38.23625711 | -88.17877929 |
| Jo Daviess County, Thompson township | 0.12775686 | 0.16810113 | 42.3734895 | -90.19320434 |
| Woodford County, Partridge township | 0.12103281 | 0.16783217 | 40.90612475 | -89.47391904 |
| Jo Daviess County, Scales Mound township | 0.1457773 | 0.16783217 | 42.48780614 | -90.24495172 |
| Pulaski County, Wetaug precinct | 0.12937063 | 0.16783217 | 37.31097348 | -89.1615583 |
| Kankakee County, Norton township | 0.10005379 | 0.16756321 | 41.05445374 | -88.13714616 |
| DeKalb County, Franklin township | 0.07261969 | 0.16756321 | 42.13149273 | -88.93167798 |
| Ford County, Dix township | 0.09198494 | 0.16729424 | 40.46105101 | -88.32202258 |
| Lee County, Bradford township | 0.05123722 | 0.16729424 | 41.83565373 | -89.26555917 |
| Jo Daviess County, Hanover township | 0.12264658 | 0.16729424 | 42.24230627 | -90.26525318 |
| Marshall County, Bell Plain township | 0.10193653 | 0.16702528 | 41.00179647 | -89.16473552 |
| Henry County, Atkinson township | 0.11484669 | 0.16702528 | 41.42425818 | -89.99974626 |
| Franklin County, Northern township | 0.12654653 | 0.16675632 | 38.06972687 | -88.74482876 |
| Adams County, Keene township | 0.13206025 | 0.16675632 | 40.15728906 | -91.23285656 |
| Mercer County, Duncan township | 0.14470145 | 0.16675632 | 41.32834683 | -90.88230793 |
| Richland County, German township | 0.12076385 | 0.16648736 | 38.76229133 | -88.00570547 |
| Menard County, Athens South No. 1 precinct | 0.08418505 | 0.16648736 | 39.95616161 | -89.70831069 |
| Ogle County, Mount Morris township | 0.09669177 | 0.16648736 | 42.04411867 | -89.39784177 |
| Hamilton County, Crook township | 0.13421194 | 0.1662184 | 38.11908291 | -88.45119691 |
| Champaign County, Harwood township | 0.10408822 | 0.1662184 | 40.3129825 | -88.09978642 |
| De Witt County, Nixon township | 0.08862292 | 0.16594944 | 40.1278722 | -88.74372855 |
| Monroe County, Precinct 34 | 0.16594944 | 0.16594944 | 38.46078498 | -90.21359475 |
| Bond County, Burgess township | 0.10381926 | 0.16594944 | 38.80910286 | -89.5743854 |
| Richland County, Denver township | 0.13125336 | 0.16568047 | 38.77157701 | -88.15042045 |
| Scott County, Naples-Bluffs precinct | 0.08270576 | 0.16568047 | 39.70902554 | -90.64376556 |
| Monroe County, Precinct 15 | 0.12668101 | 0.16514255 | 38.28671268 | -90.31404257 |
| Jersey County, Ruyle township | 0.11215707 | 0.16514255 | 39.22180943 | -90.24064397 |
| Bureau County, Macon township | 0.11081226 | 0.16514255 | 41.25085694 | -89.65294352 |
| Perry County, Pinckneyville No. 6 precinct | 0.12977407 | 0.16487359 | 38.08327665 | -89.37821873 |
| Bureau County, Milo township | 0.14416353 | 0.16433566 | 41.22708208 | -89.62495985 |
| Perry County, Beaucoup precinct | 0.13421194 | 0.1640667 | 38.1945285 | -89.36818692 |
| Marion County, Raccoon township | 0.12022593 | 0.1640667 | 38.50739292 | -88.96259973 |
| Cass County, Hagener township | 0.10745024 | 0.1640667 | 39.94635839 | -90.44862889 |
| Rock Island County, Coe township | 0.10005379 | 0.1640667 | 41.65827343 | -90.22211314 |
| Morgan County, Woodson precinct | 0.12910167 | 0.16379774 | 39.64486527 | -90.20681443 |
| Clinton County, Wade township | 0.14281872 | 0.16379774 | 38.64418412 | -89.41452533 |
| Hancock County, St. Albans township | 0.14846692 | 0.16379774 | 40.20186129 | -91.22988553 |
| Saline County, Carrier Mills township | 0.11941904 | 0.16352878 | 37.64032867 | -88.67220087 |
| Iroquois County, Stockland township | 0.10032275 | 0.16352878 | 40.61195075 | -87.52817736 |
| Jackson County, Kinkaid township | 0.10543303 | 0.16352878 | 37.84768 | -89.57186306 |
| Gallatin County, Gold Hill township | 0.10597095 | 0.16352878 | 37.69907776 | -88.17819928 |
| Effingham County, Liberty township | 0.14147391 | 0.16352878 | 39.21244964 | -88.77684745 |
| Warren County, Greenbush township | 0.08808499 | 0.16325982 | 40.63957272 | -90.45419566 |
| Mason County, Pennsylvania township | 0.12694997 | 0.16299086 | 40.22628054 | -89.74392579 |
| Fulton County, Putman township | 0.09965035 | 0.16299086 | 40.49390787 | -90.19733253 |
| Hancock County, Dallas City township | 0.12990855 | 0.16218397 | 40.60193111 | -91.173224 |
| Lee County, Nelson township | 0.07840237 | 0.16218397 | 41.76345437 | -89.60645425 |
| Grundy County, Vienna township | 0.12345347 | 0.16191501 | 41.25417308 | -88.47685067 |
| Jersey County, Piasa township | 0.09951587 | 0.16164605 | 39.039618 | -90.16747167 |
| Mercer County, Ohio Grove township | 0.14712211 | 0.16164605 | 41.07821836 | -90.78026692 |
| Perry County, Pinckneyville No. 5 precinct | 0.1422808 | 0.16164605 | 38.11015705 | -89.46235954 |
| Coles County, East Oakland township | 0.09642281 | 0.16137708 | 39.63289573 | -87.96796758 |
| Schuyler County, Littleton township | 0.09749866 | 0.16137708 | 40.19782993 | -90.57747424 |
| Bureau County, Neponset township | 0.10381926 | 0.16137708 | 41.23738808 | -89.83491362 |
| Fayette County, Pope township | 0.13367402 | 0.16137708 | 38.74284666 | -89.17275662 |
| Scott County, Alsey precinct | 0.14201183 | 0.16110812 | 39.54154845 | -90.39156171 |
| Clinton County, Wheatfield township | 0.09924691 | 0.16110812 | 38.66488658 | -89.38358343 |
| Woodford County, Panola township | 0.0894298 | 0.16083916 | 40.77104188 | -89.03726412 |
| Henry County, Phenix township | 0.08996772 | 0.16083916 | 41.49946913 | -90.17471209 |
| Whiteside County, Fenton township | 0.1344809 | 0.16083916 | 41.66948303 | -89.9787536 |
| Jefferson County, Rome township | 0.12856374 | 0.16083916 | 38.43901167 | -88.94018258 |
| Jo Daviess County, Rawlins township | 0.15693921 | 0.1605702 | 42.44983686 | -90.40130338 |
| Adams County, Mendon township | 0.10731576 | 0.1605702 | 40.15951292 | -91.29199953 |
| Henderson County, Terre Haute township | 0.1457773 | 0.1605702 | 40.66304746 | -91.019783 |
| Woodford County, Minonk township | 0.11726735 | 0.16030124 | 40.86009563 | -89.02850896 |
| Stark County, Osceola township | 0.12708445 | 0.16030124 | 41.22857546 | -89.65494948 |
| Macoupin County, Brighton township | 0.10919849 | 0.16030124 | 39.0382465 | -90.13837434 |
| Whiteside County, Portland township | 0.11377084 | 0.15976331 | 41.58865212 | -90.16730924 |
| Fayette County, Bowling Green township | 0.10193653 | 0.15976331 | 39.19692655 | -88.89559245 |
| Clinton County, Lake township | 0.11659494 | 0.15949435 | 38.54738427 | -89.27810909 |
| Shelby County, Dry Point township | 0.13932222 | 0.15949435 | 39.24141384 | -88.89122564 |
| Cumberland County, Sumpter township | 0.10785368 | 0.15949435 | 39.26613401 | -88.1847725 |
| Christian County, South Fork township | 0.07154384 | 0.15922539 | 39.5818629 | -89.38771102 |
| Livingston County, Chatsworth township | 0.09736417 | 0.15922539 | 40.75026131 | -88.29069994 |
| Macon County, Maroa township | 0.09064013 | 0.15895643 | 40.02835858 | -88.96128044 |
| Jefferson County, Farrington township | 0.10852609 | 0.15895643 | 38.47002953 | -88.70511749 |
| Perry County, Tamaroa No. 1 precinct | 0.07544379 | 0.15868747 | 38.20572378 | -89.16491268 |
| Jersey County, Quarry township | 0.13905325 | 0.15868747 | 38.98615631 | -90.46359748 |
| Ogle County, Buffalo township | 0.10919849 | 0.15868747 | 41.98640394 | -89.58602876 |
| Jackson County, Ora township | 0.10274341 | 0.1584185 | 37.9321388 | -89.47815047 |
| Washington County, Richview township | 0.11161915 | 0.1584185 | 38.38689553 | -89.20593955 |
| Pike County, Montezuma township | 0.12856374 | 0.1584185 | 39.50856678 | -90.65864157 |
| Fulton County, Young Hickory township | 0.11753631 | 0.1584185 | 40.69847759 | -90.27028116 |
| Bureau County, Concord township | 0.0831092 | 0.1584185 | 41.38900316 | -89.73105588 |
| Tazewell County, Dillon township | 0.07342657 | 0.1584185 | 40.39546474 | -89.55043872 |
| Livingston County, Amity township | 0.12331899 | 0.1584185 | 40.99920314 | -88.71434436 |
| DeKalb County, Shabbona township | 0.0957504 | 0.15814954 | 41.76824912 | -88.84686966 |
| Knox County, Chestnut township | 0.1225121 | 0.15814954 | 40.72075867 | -90.26848299 |
| Iroquois County, Iroquois township | 0.06750941 | 0.15814954 | 40.78522176 | -87.77732226 |
| Peoria County, Timber township | 0.11403981 | 0.15788058 | 40.56470085 | -89.80368139 |
| Morgan County, Lynnville precinct | 0.0957504 | 0.15788058 | 39.69344922 | -90.29131803 |
| Clay County, Blair township | 0.13273265 | 0.15761162 | 38.9096014 | -88.57343993 |
| Bond County, Mulberry Grove township | 0.07127488 | 0.15761162 | 38.927187 | -89.27217278 |
| Fulton County, Union township | 0.09776762 | 0.15761162 | 40.63826743 | -90.42443845 |
| Fayette County, Ramsey township | 0.11753631 | 0.1570737 | 39.18778846 | -89.10168476 |
| Perry County, Tamaroa No. 2 precinct | 0.12600861 | 0.1570737 | 38.1866682 | -89.22446383 |
| Hancock County, Durham township | 0.11525013 | 0.15680473 | 40.62079047 | -91.08242443 |
| Clinton County, East Fork township | 0.10032275 | 0.15680473 | 38.72059983 | -89.17484003 |
| Crawford County, Honey Creek township | 0.11713287 | 0.15680473 | 38.94513317 | -87.7225806 |
| Knox County, Henderson township | 0.10879505 | 0.15680473 | 40.99350583 | -90.36642713 |
| Crawford County, Licking township | 0.13004303 | 0.15680473 | 39.1585704 | -87.90334817 |
| Fulton County, Bernadotte township | 0.12560516 | 0.15653577 | 40.40615975 | -90.23424024 |
| Jefferson County, Blissville township | 0.11820871 | 0.15653577 | 38.28986565 | -89.07027653 |
| Winnebago County, Laona township | 0.13528779 | 0.15653577 | 42.49171663 | -89.38588349 |
| Hamilton County, Knight Prairie township | 0.1451049 | 0.15626681 | 38.08510617 | -88.62786928 |
| St. Clair County, Fayetteville township | 0.11753631 | 0.15626681 | 38.37289664 | -89.78635174 |
| Macoupin County, Nilwood township | 0.12197418 | 0.15626681 | 39.35658743 | -89.78982862 |
| McLean County, Blue Mound township | 0.11242604 | 0.15626681 | 40.55063874 | -88.70057988 |
| Pope County, Jefferson No. 4 precinct | 0.0866057 | 0.15599785 | 37.25008482 | -88.5108845 |
| Pike County, Hardin township | 0.12668101 | 0.15599785 | 39.55807292 | -90.77260573 |
| Iroquois County, Concord township | 0.10395374 | 0.15599785 | 40.79180093 | -87.53665532 |
| Macoupin County, Bunker Hill township | 0.13300161 | 0.15572889 | 39.05071495 | -89.9328645 |
| Scott County, Winchester No. 3 precinct | 0.14671867 | 0.15572889 | 39.60967014 | -90.41569451 |
| Iroquois County, Chebanse township | 0.06603012 | 0.15572889 | 40.99497626 | -87.90325264 |
| Putnam County, Hennepin township | 0.09198494 | 0.15572889 | 41.2563139 | -89.32091402 |
| Pike County, Detroit township | 0.11928456 | 0.15572889 | 39.59639986 | -90.622709 |
| McDonough County, Bethel township | 0.11686391 | 0.15545992 | 40.29569373 | -90.77802534 |
| Cass County, Arenzville township | 0.12963959 | 0.15545992 | 39.94109278 | -90.33075547 |
| Shelby County, Cold Spring township | 0.12291555 | 0.15545992 | 39.3370111 | -88.99923766 |
| Menard County, Irish Grove No. 4 precinct | 0.10839161 | 0.15545992 | 40.13278748 | -89.66311314 |
| Johnson County, Ozark precinct | 0.11081226 | 0.15519096 | 37.53593465 | -88.79711069 |
| Edgar County, Elbridge township | 0.08969876 | 0.15519096 | 39.56536047 | -87.62180154 |
| Champaign County, Sadorus township | 0.07759548 | 0.15519096 | 39.96992568 | -88.34501996 |
| Pike County, Derry township | 0.11915008 | 0.154922 | 39.58994219 | -91.00563621 |
| Adams County, Ursa township | 0.11807423 | 0.154922 | 40.03015383 | -91.41885743 |
| Mason County, Crane Creek township | 0.13542227 | 0.154922 | 40.14158369 | -89.84032643 |
| Logan County, Hurlbut township | 0.07235073 | 0.154922 | 40.01243932 | -89.49606291 |
| Jo Daviess County, Council Hill township | 0.14604626 | 0.15465304 | 42.47083921 | -90.36895985 |
| Pope County, Webster No. 5 precinct | 0.09534696 | 0.15438408 | 37.46068272 | -88.66133608 |
| Warren County, Berwick township | 0.11430877 | 0.15438408 | 40.75229004 | -90.47529418 |
| Jackson County, De Soto township | 0.10018827 | 0.15411512 | 37.78301736 | -89.20404053 |
| Clark County, Darwin township | 0.11296396 | 0.15411512 | 39.30066262 | -87.68185832 |
| Christian County, Johnson township | 0.12426036 | 0.15411512 | 39.51042131 | -89.3060157 |
| Morgan County, Meredosia No. 2 precinct | 0.09467456 | 0.15384615 | 39.79174478 | -90.49007456 |
| Massac County, Washington precinct | 0.10785368 | 0.15384615 | 37.19377247 | -88.68765596 |
| Livingston County, Nebraska township | 0.11632598 | 0.15384615 | 40.91861304 | -88.8727021 |
| Stark County, Goshen township | 0.1161915 | 0.15384615 | 41.13027624 | -89.93449968 |
| Lee County, Ashton township | 0.09924691 | 0.15384615 | 41.88023194 | -89.26126314 |
| Fulton County, Fairview township | 0.11955352 | 0.15357719 | 40.62757395 | -90.18642062 |
| Adams County, Concord township | 0.12062937 | 0.15330823 | 39.94522396 | -90.95134541 |
| Christian County, May township | 0.07423346 | 0.15303927 | 39.52951907 | -89.24539455 |
| Knox County, Salem township | 0.11807423 | 0.15303927 | 40.73204962 | -90.02833954 |
| Sangamon County, Cooper township | 0.10341581 | 0.15303927 | 39.72002214 | -89.46350626 |
| Knox County, Victoria township | 0.1316568 | 0.15277031 | 41.00081404 | -90.03566403 |
| Jefferson County, Moores Prairie township | 0.1048951 | 0.15250134 | 38.1792175 | -88.70511355 |
| De Witt County, De Witt township | 0.12426036 | 0.15250134 | 40.17236978 | -88.73921058 |
| Adams County, Houston township | 0.13071544 | 0.15223238 | 40.12816755 | -91.05698918 |
| Whiteside County, Newton township | 0.13986014 | 0.15223238 | 41.7433641 | -90.12408825 |
| Stephenson County, West Point township | 0.08741259 | 0.15223238 | 42.37890174 | -89.82536489 |
| Gallatin County, Shawnee township | 0.10126412 | 0.15223238 | 37.69724055 | -88.14967188 |
| Washington County, Lively Grove township | 0.12170522 | 0.15196342 | 38.25126248 | -89.5943242 |
| Montgomery County, East Fork township | 0.0866057 | 0.15196342 | 39.16058988 | -89.45541464 |
| Woodford County, Greene township | 0.11323292 | 0.15196342 | 40.77752222 | -89.15640923 |
| Vermilion County, Sidell township | 0.11672942 | 0.15169446 | 39.91426666 | -87.81797555 |
| Piatt County, Goose Creek township | 0.10570199 | 0.1514255 | 40.09876229 | -88.62799545 |
| Lee County, Nachusa township | 0.06428187 | 0.1514255 | 41.81963496 | -89.38864468 |
| Macoupin County, South Otter township | 0.12345347 | 0.1514255 | 39.42481993 | -89.81333257 |
| Boone County, Manchester township | 0.07719204 | 0.1514255 | 42.4229848 | -88.93263251 |
| Iroquois County, Artesia township | 0.06885422 | 0.15115654 | 40.64472834 | -88.03294861 |
| Jasper County, Smallwood township | 0.12587413 | 0.15088757 | 38.8900368 | -88.25439392 |
| Fulton County, Cass township | 0.1077192 | 0.15088757 | 40.47434024 | -90.25851532 |
| Lee County, Viola township | 0.08203335 | 0.15088757 | 41.71784294 | -89.15551053 |
| Shelby County, Ridge township | 0.11403981 | 0.15061861 | 39.43993577 | -88.84227676 |
| Jo Daviess County, Vinegar Hill township | 0.14497041 | 0.15061861 | 42.47347338 | -90.43004517 |
| Champaign County, Compromise township | 0.07288865 | 0.15034965 | 40.30926265 | -88.04064982 |
| Mercer County, Perryton township | 0.08552985 | 0.15034965 | 41.2553543 | -90.73703802 |
| LaSalle County, Serena township | 0.08418505 | 0.15034965 | 41.51474378 | -88.7213633 |
| Tazewell County, Delavan township | 0.09844002 | 0.15008069 | 40.39242403 | -89.49116484 |
| Tazewell County, Little Mackinaw township | 0.11847768 | 0.15008069 | 40.47531195 | -89.36434313 |
| Piatt County, Bement township | 0.09386767 | 0.15008069 | 39.91733637 | -88.58744945 |
| Marshall County, Richland township | 0.13017751 | 0.15008069 | 40.96831165 | -89.37812565 |
| Mercer County, Preemption township | 0.10086068 | 0.15008069 | 41.3159734 | -90.58179887 |
| Hamilton County, Twigg township | 0.11296396 | 0.14981173 | 38.03190932 | -88.48898707 |
| Shelby County, Holland township | 0.12426036 | 0.14981173 | 39.23469123 | -88.77462782 |
| Douglas County, Garrett township | 0.09642281 | 0.14981173 | 39.77511701 | -88.45441276 |
| Perry County, Sunfield precinct | 0.11592254 | 0.14981173 | 38.05317983 | -89.23675743 |
| Fulton County, Banner township | 0.08149543 | 0.14954276 | 40.5287519 | -89.98575329 |
| Vermilion County, Vance township | 0.09588488 | 0.14954276 | 40.07179082 | -87.82963791 |
| De Witt County, Barnett township | 0.08445401 | 0.14954276 | 40.21131064 | -89.03238658 |
| Fayette County, Hurricane township | 0.12869822 | 0.1492738 | 39.21324524 | -89.18700219 |
| Clark County, York township | 0.12856374 | 0.1492738 | 39.20982812 | -87.66296673 |
| Scott County, Manchester precinct | 0.12210866 | 0.1492738 | 39.54022926 | -90.36225795 |
| Edgar County, Kansas township | 0.11888112 | 0.14900484 | 39.56243247 | -87.91688591 |
| Henderson County, Raritan township | 0.13690156 | 0.14873588 | 40.70048312 | -90.83789894 |
| Wayne County, Garden Hill township | 0.1429532 | 0.14846692 | 38.57950499 | -88.66512559 |
| Adams County, Camp Point township | 0.12694997 | 0.14846692 | 40.10472109 | -91.02898129 |
| Livingston County, Forrest township | 0.09776762 | 0.14846692 | 40.75744099 | -88.40973902 |
| Montgomery County, Witt township | 0.07261969 | 0.14846692 | 39.240258 | -89.27238326 |
| Stephenson County, Buckeye township | 0.06253362 | 0.14792899 | 42.41468197 | -89.63835921 |
| Douglas County, Murdock township | 0.10852609 | 0.14792899 | 39.79785393 | -88.097333 |
| Iroquois County, Crescent township | 0.07947821 | 0.14792899 | 40.76298253 | -87.77990268 |
| Hancock County, Hancock township | 0.12937063 | 0.14792899 | 40.45891938 | -90.94472342 |
| Mercer County, Greene township | 0.12950511 | 0.14792899 | 41.20576313 | -90.62035854 |
| Carroll County, Freedom township | 0.08203335 | 0.14792899 | 42.11692455 | -89.97058511 |
| Hancock County, Rock Creek township | 0.11457773 | 0.14766003 | 40.46705525 | -91.15258399 |
| Fulton County, Lee township | 0.07732652 | 0.14766003 | 40.61598546 | -90.42618572 |
| Iroquois County, Milford township | 0.10005379 | 0.14766003 | 40.64598399 | -87.70368029 |
| Clinton County, Santa Fe township | 0.11861216 | 0.14739107 | 38.55672035 | -89.45143726 |
| Wabash County, Friendsville precinct | 0.12049489 | 0.14739107 | 38.50265134 | -87.80175823 |
| Knox County, Lynn township | 0.10785368 | 0.14739107 | 41.13591681 | -90.05432657 |
| Monroe County, Precinct 22 | 0.10086068 | 0.14712211 | 38.32448442 | -90.16650421 |
| McDonough County, Walnut Grove township | 0.07342657 | 0.14712211 | 40.55937505 | -90.66921196 |
| Lee County, Marion township | 0.08351264 | 0.14712211 | 41.75585567 | -89.45538008 |
| Monroe County, Precinct 26 | 0.14685315 | 0.14685315 | 38.34537458 | -90.13591009 |
| Shelby County, Sigel township | 0.07450242 | 0.14685315 | 39.23964358 | -88.48091956 |
| LaSalle County, Adams township | 0.09050565 | 0.14685315 | 41.6089365 | -88.80250833 |
| Monroe County, Precinct 7 | 0.09454008 | 0.14658419 | 38.32311278 | -90.13769029 |
| Monroe County, Precinct 24 | 0.11941904 | 0.14658419 | 38.46213882 | -90.2424642 |
| Iroquois County, Martinton township | 0.08351264 | 0.14658419 | 40.9167275 | -87.73199309 |
| Champaign County, Newcomb township | 0.112695 | 0.14658419 | 40.26443357 | -88.4026711 |
| Pike County, Griggsville township | 0.1190156 | 0.14631522 | 39.66944672 | -90.76452187 |
| Bureau County, Bureau township | 0.10919849 | 0.14631522 | 41.44991114 | -89.60474855 |
| Marion County, Carrigan township | 0.08929532 | 0.14604626 | 38.6490318 | -89.09431548 |
| Lawrence County, Bond township | 0.08768155 | 0.14604626 | 38.78570551 | -87.68238848 |
| Effingham County, Lucas township | 0.12318451 | 0.14604626 | 38.96394489 | -88.36338415 |
| Scott County, Winchester No. 2 precinct | 0.11995697 | 0.14604626 | 39.63454757 | -90.47265334 |
| Fulton County, Kerton township | 0.12789134 | 0.14604626 | 40.20297376 | -90.19126327 |
| Douglas County, Bowdre township | 0.10341581 | 0.1457773 | 39.75891306 | -88.19025114 |
| Bureau County, Leepertown township | 0.1035503 | 0.1457773 | 41.25787859 | -89.3509095 |
| Lee County, Brooklyn township | 0.07315761 | 0.14550834 | 41.69555932 | -89.15768785 |
| Cass County, Ashland township | 0.13206025 | 0.14523938 | 39.94971303 | -90.03429763 |
| Putnam County, Senachwine township | 0.12721893 | 0.14523938 | 41.1926056 | -89.38717852 |
| Jo Daviess County, Wards Grove township | 0.10731576 | 0.14523938 | 42.36375659 | -89.97976656 |
| Monroe County, Precinct 13 | 0.1098709 | 0.14497041 | 38.12548303 | -90.21113638 |
| Montgomery County, Harvel township | 0.10274341 | 0.14497041 | 39.36697075 | -89.55413926 |
| Schuyler County, Brooklyn township | 0.11471221 | 0.14497041 | 40.20656794 | -90.78453422 |
| Christian County, Ricks township | 0.08445401 | 0.14470145 | 39.42916715 | -89.46045261 |
| Marion County, Iuka township | 0.11000538 | 0.14470145 | 38.64964736 | -88.71623987 |
| Adams County, Honey Creek township | 0.08902636 | 0.14470145 | 40.02245804 | -91.21223237 |
| LaSalle County, Groveland township | 0.10260893 | 0.14470145 | 40.94915489 | -89.0197318 |
| Washington County, Oakdale township | 0.10018827 | 0.14443249 | 38.27351614 | -89.59238113 |
| Marion County, Omega township | 0.11753631 | 0.14443249 | 38.6718861 | -88.7140189 |
| Bureau County, Gold township | 0.10126412 | 0.14443249 | 41.41712368 | -89.84935792 |
| Scott County, Exeter-Bluffs precinct | 0.09400215 | 0.14443249 | 39.76947228 | -90.49178098 |
| Whiteside County, Mount Pleasant township | 0.10610543 | 0.14416353 | 41.80041791 | -89.90675801 |
| Perry County, Pinckneyville No. 4 precinct | 0.10597095 | 0.14416353 | 38.06411745 | -89.43761557 |
| Tazewell County, Malone township | 0.1344809 | 0.14389457 | 40.38067709 | -89.70061322 |
| Montgomery County, Walshville township | 0.11632598 | 0.14389457 | 39.03464917 | -89.61285862 |
| Knox County, Ontario township | 0.08714363 | 0.14362561 | 41.14686079 | -90.29403678 |
| McLean County, White Oak township | 0.10381926 | 0.14362561 | 40.59294997 | -89.05470874 |
| Edgar County, Ross township | 0.12668101 | 0.14308768 | 39.7973483 | -87.74258992 |
| Jackson County, Pomona township | 0.07530931 | 0.14281872 | 37.67494074 | -89.27134866 |
| Crawford County, Prairie township | 0.11861216 | 0.14281872 | 39.15284547 | -87.81607117 |
| Sangamon County, Illiopolis township | 0.1218397 | 0.14281872 | 39.84422413 | -89.27489472 |
| Stark County, West Jersey township | 0.11350188 | 0.14281872 | 40.9979994 | -89.97586323 |
| Perry County, Pinckneyville No. 7 precinct | 0.13259817 | 0.14254976 | 38.0801574 | -89.32081221 |
| Livingston County, Saunemin township | 0.10032275 | 0.14254976 | 40.89095449 | -88.39541887 |
| Bureau County, Wheatland township | 0.12116729 | 0.1422808 | 41.19875976 | -89.50706589 |
| LaSalle County, Mission township | 0.09642281 | 0.1422808 | 41.53012034 | -88.59866299 |
| Randolph County, Bremen precinct | 0.11484669 | 0.14201183 | 38.01240262 | -89.73034778 |
| Livingston County, Waldo township | 0.1013986 | 0.14201183 | 40.76278452 | -88.88836359 |
| Livingston County, Rooks Creek township | 0.10543303 | 0.14201183 | 40.88790238 | -88.72579938 |
| Edgar County, Hunter township | 0.09467456 | 0.14174287 | 39.65424756 | -87.61144681 |
| Adams County, Fall Creek township | 0.09494352 | 0.14174287 | 39.82850806 | -91.4022009 |
| Shelby County, Lakewood township | 0.11444325 | 0.14147391 | 39.30479457 | -88.82631036 |
| McDonough County, Tennessee township | 0.10193653 | 0.14147391 | 40.38967695 | -90.89013719 |
| Coles County, Pleasant Grove township | 0.10193653 | 0.14093599 | 39.4104122 | -88.34554953 |
| Monroe County, Precinct 12 | 0.10220549 | 0.14093599 | 38.30629533 | -90.2546967 |
| Piatt County, Unity township | 0.08055406 | 0.14093599 | 39.79912339 | -88.48144372 |
| Randolph County, Steeleville precinct | 0.10866057 | 0.14039806 | 38.05544053 | -89.69785444 |
| Fayette County, South Hurricane township | 0.08781603 | 0.14039806 | 39.14489993 | -89.1641558 |
| Knox County, Indian Point township | 0.11081226 | 0.14039806 | 40.74968213 | -90.41568565 |
| Fulton County, Joshua township | 0.10381926 | 0.1401291 | 40.62483085 | -90.12692746 |
| Johnson County, Bloomfield precinct | 0.07880581 | 0.1401291 | 37.42807625 | -88.86478626 |
| Morgan County, Markham precinct | 0.11094675 | 0.1401291 | 39.73932421 | -90.3171666 |
| Clay County, Bible Grove township | 0.12896719 | 0.13986014 | 38.902582 | -88.45740341 |
| Crawford County, Hutsonville township | 0.11027434 | 0.13986014 | 39.07845542 | -87.70735644 |
| Macoupin County, Hillyard township | 0.10462614 | 0.13959118 | 39.12173701 | -90.01465728 |
| Jefferson County, Field township | 0.11861216 | 0.13932222 | 38.41511073 | -88.91350308 |
| Jackson County, Elk township | 0.09642281 | 0.13932222 | 37.94193679 | -89.24696812 |
| Effingham County, Moccasin township | 0.09386767 | 0.13932222 | 39.10124273 | -88.78792519 |
| Wabash County, Coffee precinct | 0.10933297 | 0.13905325 | 38.35282426 | -87.9054941 |
| Hancock County, Fountain Green township | 0.12291555 | 0.13905325 | 40.54806697 | -90.93839192 |
| Clinton County, Irishtown township | 0.11188811 | 0.13878429 | 38.66333158 | -89.35465102 |
| Christian County, Locust township | 0.07665412 | 0.13851533 | 39.48501616 | -89.24956072 |
| Knox County, Copley township | 0.09010221 | 0.13851533 | 41.04955238 | -90.12166978 |
| Jersey County, Otter Creek township | 0.11780527 | 0.13824637 | 39.00449551 | -90.3746331 |
| Tazewell County, Sand Prairie township | 0.10260893 | 0.13824637 | 40.40146283 | -89.66900152 |
| Shelby County, Rural township | 0.1007262 | 0.13797741 | 39.44988394 | -89.01768726 |
| Pike County, Flint township | 0.11861216 | 0.13797741 | 39.71027882 | -90.67314298 |
| Vermilion County, Elwood township | 0.1013986 | 0.13797741 | 39.92485663 | -87.63902143 |
| Scott County, Bloomfield precinct | 0.11202259 | 0.13770845 | 39.61485567 | -90.53303257 |
| Montgomery County, Irving township | 0.09494352 | 0.13743948 | 39.18284274 | -89.45340462 |
| Adams County, Richfield township | 0.10314685 | 0.13743948 | 39.77166137 | -91.08148626 |
| Lee County, Reynolds township | 0.04612695 | 0.13743948 | 41.87060508 | -89.07972257 |
| Ford County, Pella township | 0.09171598 | 0.13743948 | 40.81337424 | -88.22386933 |
| Jersey County, Fidelity township | 0.09736417 | 0.13717052 | 39.17319832 | -90.15672341 |
| Bond County, Shoal Creek township | 0.08136095 | 0.13717052 | 39.01089346 | -89.58574025 |
| Scott County, Winchester No. 1 precinct | 0.09279182 | 0.13717052 | 39.63324734 | -90.44331032 |
| Whiteside County, Hume township | 0.08041958 | 0.13717052 | 41.75004385 | -89.78976395 |
| Jasper County, Grandville township | 0.10623991 | 0.13690156 | 39.09567536 | -87.96893506 |
| Menard County, Athens North No. 2 precinct | 0.11108123 | 0.13690156 | 40.00216119 | -89.73388073 |
| Champaign County, Crittenden township | 0.08848843 | 0.1366326 | 39.96268944 | -88.2272926 |
| White County, Enfield township | 0.08216783 | 0.13636364 | 38.0638805 | -88.28370599 |
| Marion County, Foster township | 0.08418505 | 0.13636364 | 38.82044713 | -88.9614801 |
| McDonough County, Eldorado township | 0.10341581 | 0.13636364 | 40.30536855 | -90.48015173 |
| Lee County, Wyoming township | 0.06401291 | 0.13636364 | 41.70971594 | -89.00458568 |
| Christian County, King township | 0.09696073 | 0.13609467 | 39.36545548 | -89.5249226 |
| Livingston County, Long Point township | 0.10946746 | 0.13609467 | 40.96146121 | -88.83835508 |
| Bond County, Lagrange township | 0.0831092 | 0.13582571 | 39.02244733 | -89.38021773 |
| Scott County, Merritt precinct | 0.09427111 | 0.13582571 | 39.74590455 | -90.46409706 |
| Sangamon County, Cartwright township | 0.08848843 | 0.13582571 | 39.87726235 | -89.92211053 |
| Bureau County, Ohio township | 0.08001614 | 0.13582571 | 41.57755353 | -89.4720087 |
| Randolph County, Evansville precinct | 0.12049489 | 0.13555675 | 38.08930911 | -89.92565494 |
| Crawford County, Martin township | 0.11457773 | 0.13555675 | 38.95284013 | -87.83861117 |
| Clinton County, Clement township | 0.11013986 | 0.13528779 | 38.56963264 | -89.27606201 |
| Morgan County, Meredosia No. 1 precinct | 0.10274341 | 0.13528779 | 39.83499325 | -90.45723296 |
| Lee County, Amboy township | 0.08512641 | 0.13528779 | 41.68122086 | -89.31072785 |
| DeKalb County, Milan township | 0.05110274 | 0.13501883 | 41.87965419 | -88.83546318 |
| Madison County, New Douglas township | 0.1042227 | 0.13501883 | 38.99460023 | -89.70397881 |
| Shelby County, Flat Branch township | 0.09117805 | 0.13501883 | 39.60561705 | -89.00258977 |
| Jasper County, Ste. Marie township | 0.12103281 | 0.13474987 | 38.87153862 | -87.96450368 |
| Ford County, Lyman township | 0.07665412 | 0.13474987 | 40.60949672 | -88.18639498 |
| Henry County, Edford township | 0.112695 | 0.1344809 | 41.45624432 | -90.20850438 |
| Johnson County, Simpson precinct | 0.08902636 | 0.13421194 | 37.51030076 | -88.74233217 |
| Washington County, Johannisburg township | 0.09238838 | 0.13421194 | 38.36403135 | -89.61341566 |
| Morgan County, Waverly No. 1 precinct | 0.09817106 | 0.13421194 | 39.52665982 | -90.0692803 |
| Livingston County, Charlotte township | 0.10502959 | 0.13421194 | 40.79294623 | -88.2560839 |
| Carroll County, Washington township | 0.07665412 | 0.13421194 | 42.12392133 | -90.12249447 |
| Livingston County, Sullivan township | 0.11108123 | 0.13394298 | 40.88376057 | -88.27615716 |
| Shelby County, Penn township | 0.1070468 | 0.13394298 | 39.64013695 | -88.82236945 |
| Champaign County, Stanton township | 0.07006455 | 0.13394298 | 40.15173902 | -88.02848472 |
| Adams County, Beverly township | 0.12009145 | 0.13367402 | 39.78926824 | -90.96229441 |
| Scott County, Glasgow precinct | 0.11672942 | 0.13367402 | 39.54546482 | -90.47947937 |
| Randolph County, Baldwin precinct | 0.11457773 | 0.13340506 | 38.1711288 | -89.77454519 |
| Jackson County, Levan township | 0.07504034 | 0.13340506 | 37.86538012 | -89.48405245 |
| Livingston County, Germanville township | 0.11161915 | 0.13340506 | 40.65945825 | -88.27064578 |
| Adams County, McKee township | 0.10664336 | 0.13340506 | 39.85728501 | -90.9870506 |
| Perry County, Cutler precinct | 0.0915815 | 0.13313609 | 38.02267676 | -89.49894734 |
| Macoupin County, North Otter township | 0.10381926 | 0.13313609 | 39.44996948 | -89.86993754 |
| Henry County, Hanna township | 0.08633674 | 0.13286713 | 41.47853889 | -90.20666727 |
| Crawford County, Montgomery township | 0.09655729 | 0.13286713 | 38.93928174 | -87.63557389 |
| Macon County, Niantic township | 0.077461 | 0.13286713 | 39.84105535 | -89.21608699 |
| McLean County, Lawndale township | 0.08230231 | 0.13286713 | 40.58997354 | -88.60690868 |
| Henry County, Yorktown township | 0.11161915 | 0.13259817 | 41.55517166 | -89.92804089 |
| Lee County, Lee Center township | 0.0929263 | 0.13259817 | 41.71944711 | -89.18569968 |
| St. Clair County, Prairie Du Long township | 0.08041958 | 0.13232921 | 38.3829323 | -89.98816252 |
| Peoria County, Elmwood township | 0.09951587 | 0.13232921 | 40.77236678 | -89.93517294 |
| Carroll County, Woodland township | 0.07719204 | 0.13232921 | 42.11833816 | -90.00096454 |
| Johnson County, Burnside precinct | 0.12627757 | 0.13206025 | 37.57702714 | -88.73579106 |
| Vermilion County, Carroll township | 0.10718128 | 0.13206025 | 39.93457277 | -87.78603078 |
| Lee County, Sublette township | 0.10395374 | 0.13206025 | 41.65736125 | -89.28268875 |
| Pike County, Fairmount township | 0.10287789 | 0.13179129 | 39.80676196 | -90.84304441 |
| Adams County, Gilmer township | 0.09306079 | 0.13179129 | 39.95672587 | -91.24618057 |
| Whiteside County, Fulton township | 0.09467456 | 0.13179129 | 41.85624849 | -90.1449875 |
| Clinton County, St. Rose township | 0.10946746 | 0.13179129 | 38.6755819 | -89.5861444 |
| Lawrence County, Christy township | 0.05863367 | 0.13179129 | 38.73061382 | -87.86351079 |
| Massac County, Georges Creek precinct | 0.09171598 | 0.13152232 | 37.2878669 | -88.7640512 |
| Perry County, Swanwick precinct | 0.08136095 | 0.13152232 | 38.17691232 | -89.45641824 |
| St. Clair County, Engelmann township | 0.09817106 | 0.13152232 | 38.46337812 | -89.80765432 |
| Jasper County, Hunt City township | 0.10785368 | 0.13152232 | 39.03087572 | -88.00536155 |
| Fulton County, Ellisville township | 0.10368478 | 0.13152232 | 40.69981909 | -90.30006071 |
| DeKalb County, Victor township | 0.06939215 | 0.13152232 | 41.63121192 | -88.80022103 |
| Henry County, Lynn township | 0.10946746 | 0.13125336 | 41.24133125 | -90.40685505 |
| Union County, Dongola District 1 precinct | 0.06320602 | 0.1309844 | 37.35547535 | -89.15746124 |
| Randolph County, Coulterville precinct | 0.11081226 | 0.13071544 | 38.16374424 | -89.63082524 |
| Boone County, Boone township | 0.09642281 | 0.13071544 | 42.39059282 | -88.75205047 |
| Mason County, Forest City township | 0.10543303 | 0.13044648 | 40.36281142 | -89.79146342 |
| McDonough County, New Salem township | 0.09480904 | 0.13044648 | 40.44290497 | -90.55882656 |
| LaSalle County, Freedom township | 0.07974718 | 0.13044648 | 41.45804861 | -88.90868641 |
| Henry County, Osco township | 0.10220549 | 0.13017751 | 41.34882074 | -90.30783089 |
| Logan County, Prairie Creek township | 0.07261969 | 0.12990855 | 40.28564752 | -89.59008859 |
| Bureau County, Greenville township | 0.10301237 | 0.12990855 | 41.56433726 | -89.65485935 |
| St. Clair County, Lenzburg township | 0.09844002 | 0.12963959 | 38.26306504 | -89.82455799 |
| Macon County, Pleasant View township | 0.11726735 | 0.12963959 | 39.67887757 | -89.11342543 |
| Carroll County, Rock Creek-Lima township | 0.06374395 | 0.12937063 | 42.08889406 | -89.85104783 |
| Mason County, Sherman township | 0.10193653 | 0.12937063 | 40.25292113 | -89.83072503 |
| Randolph County, Wine Hill precinct | 0.11148467 | 0.12910167 | 37.92190406 | -89.70927008 |
| Marshall County, Whitefield township | 0.09386767 | 0.12910167 | 41.06050112 | -89.42960097 |
| Livingston County, Round Grove township | 0.09279182 | 0.12910167 | 41.11171081 | -88.34152082 |
| Mercer County, Richland Grove township | 0.09494352 | 0.12856374 | 41.24264127 | -90.43686621 |
| Hamilton County, Flannigan township | 0.11699839 | 0.12856374 | 38.0183928 | -88.63455312 |
| Marshall County, Roberts township | 0.10919849 | 0.12856374 | 41.0240656 | -89.16258175 |
| Edgar County, Prairie township | 0.09386767 | 0.12829478 | 39.81373336 | -87.65196694 |
| Knox County, Maquon township | 0.08754707 | 0.12829478 | 40.71805385 | -90.20890867 |
| Iroquois County, Beaver township | 0.08674018 | 0.12829478 | 40.86443668 | -87.61803015 |
| Bureau County, Arispie township | 0.09306079 | 0.12829478 | 41.24025216 | -89.44297708 |
| Menard County, Rock Creek No. 12 precinct | 0.09817106 | 0.12802582 | 39.94119065 | -89.85753396 |
| Fulton County, Liverpool township | 0.08068854 | 0.12775686 | 40.37703332 | -90.08781956 |
| Piatt County, Cerro Gordo township | 0.08741259 | 0.1274879 | 39.81300927 | -88.71643584 |
| Whiteside County, Erie township | 0.12331899 | 0.1274879 | 41.60957487 | -90.13529225 |
| Sangamon County, Cotton Hill township | 0.09359871 | 0.12721893 | 39.67703795 | -89.49688989 |
| Knox County, Cedar township | 0.09440559 | 0.12694997 | 40.88339103 | -90.405146 |
| Gallatin County, North Fork township | 0.08122647 | 0.12694997 | 37.79529027 | -88.28303454 |
| Iroquois County, Beaverville township | 0.08122647 | 0.12694997 | 41.00179421 | -87.66188432 |
| Livingston County, Pleasant Ridge township | 0.08579882 | 0.12668101 | 40.78324163 | -88.46690312 |
| Iroquois County, Ashkum township | 0.0809575 | 0.12668101 | 40.86528741 | -87.97811035 |
| Bureau County, Clarion township | 0.08405056 | 0.12668101 | 41.56186776 | -89.17072273 |
| Stephenson County, Kent township | 0.07073696 | 0.12668101 | 42.35658983 | -89.82734811 |
| Pulaski County, Perks precinct | 0.09077461 | 0.12641205 | 37.26325149 | -89.10888249 |
| McDonough County, Industry township | 0.11188811 | 0.12641205 | 40.33403935 | -90.62658377 |
| Randolph County, Ruma precinct | 0.11659494 | 0.12587413 | 38.16033385 | -90.00637216 |
| Woodford County, Kansas township | 0.08795051 | 0.12560516 | 40.63747108 | -89.05035577 |
| Franklin County, Eastern township | 0.09400215 | 0.12560516 | 38.00639291 | -88.80872684 |
| McLean County, Cropsey township | 0.09696073 | 0.12560516 | 40.60522962 | -88.48577712 |
| Jackson County, Vergennes township | 0.09722969 | 0.1253362 | 37.9435171 | -89.27561545 |
| Sangamon County, Lanesville township | 0.08539537 | 0.12506724 | 39.82354306 | -89.30637519 |
| Effingham County, Union township | 0.09346423 | 0.12479828 | 38.93183809 | -88.57116742 |
| Warren County, Spring Grove township | 0.09064013 | 0.12452932 | 40.98409119 | -90.66721109 |
| Washington County, Venedy township | 0.10395374 | 0.12452932 | 38.43079269 | -89.60759133 |
| Pike County, Perry township | 0.10193653 | 0.12452932 | 39.75731528 | -90.7286382 |
| Morgan County, Concord precinct | 0.08768155 | 0.12452932 | 39.85596151 | -90.42608069 |
| Rock Island County, Edgington township | 0.08176439 | 0.12452932 | 41.41020381 | -90.69527336 |
| Williamson County, Crab Orchard precinct | 0.07275417 | 0.12399139 | 37.73271601 | -88.72048262 |
| McDonough County, Sciota township | 0.07813341 | 0.12372243 | 40.62995539 | -90.75347184 |
| Whiteside County, Tampico township | 0.10005379 | 0.12345347 | 41.61918148 | -89.86192596 |
| White County, Indian Creek township | 0.0739645 | 0.12345347 | 37.976758 | -88.32166574 |
| Ogle County, Brookville township | 0.0894298 | 0.12345347 | 42.10090442 | -89.63649419 |
| Bureau County, Indiantown township | 0.09131253 | 0.12291555 | 41.24936306 | -89.62294446 |
| Tazewell County, Hittle township | 0.09776762 | 0.12264658 | 40.36243883 | -89.34508067 |
| Livingston County, Union township | 0.09359871 | 0.12264658 | 40.93901913 | -88.45031059 |
| Champaign County, Condit township | 0.06737493 | 0.12264658 | 40.30534172 | -88.33878366 |
| White County, Heralds Prairie township | 0.06186122 | 0.12237762 | 38.0380187 | -88.22871456 |
| Knox County, Walnut Grove township | 0.10583647 | 0.12237762 | 41.07321459 | -90.14974971 |
| Hamilton County, Mayberry township | 0.0739645 | 0.12210866 | 38.03014492 | -88.46032327 |
| Bureau County, Fairfield township | 0.10637439 | 0.12157074 | 41.54941004 | -89.80750963 |
| Fayette County, Shafter township | 0.04626143 | 0.12130178 | 39.00980982 | -89.14765629 |
| Jasper County, Crooked Creek township | 0.09965035 | 0.12103281 | 39.15130808 | -88.13856668 |
| Jasper County, Fox township | 0.11067778 | 0.12076385 | 38.94570109 | -88.07317161 |
| Iroquois County, Fountain Creek township | 0.08431953 | 0.12076385 | 40.56667473 | -87.86238852 |
| Lee County, Franklin Grove township | 0.06132329 | 0.12076385 | 41.88181159 | -89.29152473 |
| Bureau County, Manlius township | 0.09763314 | 0.12049489 | 41.45140955 | -89.63483269 |
| Bureau County, Westfield township | 0.0591716 | 0.12049489 | 41.40910159 | -89.24598126 |
| Vermilion County, Love township | 0.10220549 | 0.12049489 | 39.94314596 | -87.57761692 |
| Jefferson County, Bald Hill township | 0.07221625 | 0.12022593 | 38.20411759 | -89.13616562 |
| Ford County, Button township | 0.07988166 | 0.12022593 | 40.42045376 | -88.02821987 |
| Christian County, Stonington township | 0.05661646 | 0.12022593 | 39.68048935 | -89.14276022 |
| Edgar County, Brouilletts Creek township | 0.07315761 | 0.12022593 | 39.78954908 | -87.6252082 |
| Montgomery County, Fillmore Consolidated township | 0.05755783 | 0.119688 | 39.04314203 | -89.34910509 |
| Adams County, Burton township | 0.07894029 | 0.11941904 | 39.93444219 | -91.24765642 |
| Jo Daviess County, Derinda township | 0.07988166 | 0.11941904 | 42.26190968 | -90.20251777 |
| Douglas County, Sargent township | 0.10247445 | 0.11915008 | 39.72743912 | -88.04600785 |
| Pulaski County, Grand Chain precinct | 0.10086068 | 0.11915008 | 37.23118624 | -88.94071993 |
| Moultrie County, Whitley township | 0.09763314 | 0.11915008 | 39.51698036 | -88.62860844 |
| Tazewell County, Spring Lake township | 0.0887574 | 0.11915008 | 40.45188933 | -89.78369986 |
| Gallatin County, Ridgway township | 0.06710597 | 0.11888112 | 37.79347762 | -88.25446743 |
| Perry County, Pinckneyville No. 2 precinct | 0.10650888 | 0.11888112 | 38.03565789 | -89.32485094 |
| Stephenson County, Oneco township | 0.07799892 | 0.11861216 | 42.50242712 | -89.59962873 |
| Ogle County, Lafayette township | 0.06818182 | 0.11861216 | 41.90252184 | -89.25911308 |
| Monroe County, Precinct 9 | 0.09306079 | 0.1183432 | 38.11999788 | -90.09619533 |
| Marion County, Haines township | 0.09077461 | 0.1183432 | 38.48183707 | -88.90703708 |
| De Witt County, Waynesville township | 0.10274341 | 0.1183432 | 40.27970835 | -89.05544777 |
| Bond County, Tamalco township | 0.09037117 | 0.11753631 | 38.76986739 | -89.25758209 |
| Gallatin County, Equality township | 0.09359871 | 0.11726735 | 37.70454933 | -88.26378994 |
| Logan County, Eminence township | 0.06159225 | 0.11726735 | 40.22416591 | -89.26883372 |
| Ogle County, Lincoln township | 0.08526089 | 0.11699839 | 42.07710463 | -89.60817528 |
| Grundy County, Highland township | 0.07235073 | 0.11699839 | 41.18739007 | -88.48397735 |
| McLean County, Anchor township | 0.08848843 | 0.11646046 | 40.56423942 | -88.549849 |
| Carroll County, Cherry Grove-Shannon township | 0.07369554 | 0.1161915 | 42.11119881 | -89.8490798 |
| Ford County, Mona township | 0.08203335 | 0.11592254 | 40.90419509 | -88.24390712 |
| Ogle County, Forreston township | 0.07288865 | 0.11592254 | 42.18861369 | -89.59793566 |
| Christian County, Mount Auburn township | 0.08916084 | 0.11592254 | 39.7758813 | -89.25176462 |
| Morgan County, Franklin precinct | 0.08579882 | 0.11565358 | 39.52804765 | -90.09857308 |
| Henry County, Western township | 0.06952663 | 0.11565358 | 41.32786044 | -90.33967826 |
| Champaign County, Raymond township | 0.08485745 | 0.11565358 | 39.8886394 | -88.11692797 |
| Warren County, Coldbrook township | 0.08848843 | 0.11538462 | 40.95286603 | -90.45963792 |
| Whiteside County, Union Grove township | 0.09413663 | 0.11511565 | 41.76008324 | -90.00130773 |
| Bond County, Mills township | 0.07611619 | 0.11484669 | 38.82527239 | -89.45641887 |
| Ford County, Sullivant township | 0.0809575 | 0.11484669 | 40.53316505 | -88.40381702 |
| Massac County, Logan precinct | 0.09467456 | 0.11484669 | 37.27402185 | -88.90811225 |
| Montgomery County, Grisham township | 0.09696073 | 0.11484669 | 39.03012455 | -89.52560915 |
| Monroe County, Precinct 8 | 0.08821947 | 0.11457773 | 38.29669679 | -90.05306026 |
| Bureau County, Berlin township | 0.059844 | 0.11457773 | 41.43453918 | -89.30397877 |
| Johnson County, Tunnel Hill precinct | 0.06105433 | 0.11430877 | 37.52039254 | -88.91316952 |
| Brown County, Versailles township | 0.07544379 | 0.11430877 | 39.8414096 | -90.60437134 |
| Union County, Anna District 6 precinct | 0.11403981 | 0.11403981 | 37.49535352 | -89.25902964 |
| Christian County, Greenwood township | 0.07920925 | 0.11403981 | 39.35308703 | -89.29123393 |
| McLean County, Cheney's Grove township | 0.09534696 | 0.11403981 | 40.44948467 | -88.50221461 |
| Carroll County, York township | 0.06952663 | 0.11403981 | 41.9663874 | -90.10531962 |
| Effingham County, Bishop township | 0.06885422 | 0.11403981 | 39.08045903 | -88.43890207 |
| Logan County, Laenna township | 0.09548144 | 0.11377084 | 39.97617185 | -89.23290636 |
| Adams County, Columbus township | 0.07705756 | 0.11377084 | 40.01905603 | -91.12369253 |
| Ford County, Peach Orchard township | 0.08230231 | 0.11377084 | 40.61496471 | -88.2754876 |
| Woodford County, Linn township | 0.08929532 | 0.11350188 | 40.91749913 | -89.26288163 |
| Whiteside County, Hahnaman township | 0.07530931 | 0.11350188 | 41.58959354 | -89.71313243 |
| Perry County, Pinckneyville No. 3 precinct | 0.08337816 | 0.11323292 | 37.99889642 | -89.4722378 |
| Crawford County, Southwest township | 0.10301237 | 0.11323292 | 38.86775763 | -87.90654344 |
| Montgomery County, Rountree township | 0.08633674 | 0.11296396 | 39.34015037 | -89.46850395 |
| Woodford County, Palestine township | 0.09252286 | 0.11296396 | 40.73299307 | -89.16070434 |
| Ford County, Wall township | 0.07557827 | 0.112695 | 40.49460283 | -88.13931572 |
| Douglas County, Newman township | 0.10368478 | 0.112695 | 39.8793105 | -87.96996832 |
| Hamilton County, South Flannigan township | 0.09077461 | 0.11242604 | 37.92944005 | -88.6434461 |
| Livingston County, Belle Prairie township | 0.09252286 | 0.11242604 | 40.65149145 | -88.5107877 |
| Jackson County, Fountain Bluff township | 0.0689887 | 0.11188811 | 37.75866072 | -89.57960828 |
| Pike County, Chambersburg township | 0.09117805 | 0.11081226 | 39.79937745 | -90.66654107 |
| Bureau County, La Moille township | 0.08041958 | 0.11081226 | 41.57136397 | -89.3514782 |
| Champaign County, East Bend township | 0.08216783 | 0.11027434 | 40.32216106 | -88.24765464 |
| Stephenson County, Winslow township | 0.09991931 | 0.11027434 | 42.49480133 | -89.9070526 |
| Morgan County, Prentice-Sinclair precinct | 0.06428187 | 0.11000538 | 39.85924258 | -90.01226945 |
| McLean County, Allin township | 0.08203335 | 0.11000538 | 40.44517866 | -89.21814284 |
| Johnson County, Lake No. 1 precinct | 0.08189887 | 0.11000538 | 37.59044132 | -88.96378823 |
| Adams County, Liberty township | 0.08203335 | 0.11000538 | 39.85962386 | -91.04593909 |
| Franklin County, Six Mile township | 0.07786444 | 0.10973642 | 37.93068682 | -89.0464715 |
| Randolph County, Percy precinct | 0.10032275 | 0.10973642 | 37.98421234 | -89.61754959 |
| Marion County, Patoka township | 0.0626681 | 0.10973642 | 38.76187698 | -89.11273818 |
| Iroquois County, Milks Grove township | 0.07759548 | 0.10946746 | 40.98585468 | -88.11472096 |
| Fayette County, Seminary township | 0.06508876 | 0.10919849 | 38.90335744 | -89.2452057 |
| Morgan County, Murrayville precinct | 0.08284024 | 0.10919849 | 39.5362305 | -90.27435314 |
| Iroquois County, Ash Grove township | 0.07369554 | 0.10919849 | 40.58891253 | -87.85984098 |
| Pike County, New Salem township | 0.08512641 | 0.10892953 | 39.67188976 | -90.82324959 |
| Ford County, Rogers township | 0.08539537 | 0.10866057 | 40.98770157 | -88.14457293 |
| Monroe County, Precinct 29 | 0.08512641 | 0.10785368 | 38.27443615 | -90.05486439 |
| Fulton County, Farmers township | 0.08136095 | 0.10785368 | 40.3682828 | -90.38604461 |
| Franklin County, Cave township | 0.07665412 | 0.10758472 | 37.86615758 | -88.70730975 |
| Logan County, Sheridan township | 0.06670253 | 0.10677784 | 40.15206328 | -89.60206077 |
| Champaign County, Ayers township | 0.09413663 | 0.10677784 | 39.90342202 | -87.99687631 |
| Williamson County, Creal Springs precinct | 0.03227542 | 0.10623991 | 37.67443703 | -88.86970997 |
| Henry County, Weller township | 0.09265734 | 0.10623991 | 41.23196405 | -90.19680847 |
| Morgan County, Nortonville precinct | 0.07961269 | 0.10623991 | 39.53080272 | -90.12786698 |
| Vermilion County, Jamaica township | 0.09655729 | 0.10623991 | 39.96064058 | -87.8423149 |
| Ford County, Brenton township | 0.0922539 | 0.10623991 | 40.6762321 | -88.17904694 |
| McDonough County, Hire township | 0.08203335 | 0.10543303 | 40.47518367 | -90.79465347 |
| Logan County, Aetna township | 0.05836471 | 0.10516407 | 40.06200214 | -89.16550439 |
| Greene County, Athensville township | 0.08512641 | 0.1048951 | 39.48763784 | -90.19004478 |
| Fulton County, Deerfield township | 0.0831092 | 0.1048951 | 40.60665615 | -90.21798114 |
| Massac County, Hillerman precinct | 0.09413663 | 0.10462614 | 37.24675636 | -88.82511727 |
| Wabash County, Wabash precinct | 0.06401291 | 0.10462614 | 38.4987944 | -87.74409839 |
| Adams County, Payson township | 0.06925767 | 0.10462614 | 39.77622281 | -91.19913293 |
| Lawrence County, Bridgeport township | 0.06347499 | 0.10435718 | 38.70264058 | -87.77925625 |
| Macon County, Austin township | 0.07961269 | 0.10435718 | 40.01427393 | -89.110806 |
| Morgan County, Waverly No. 3 precinct | 0.0802851 | 0.10408822 | 39.59065493 | -90.00514782 |
| Johnson County, Goreville No. 1 precinct | 0.06239914 | 0.10381926 | 37.59208807 | -88.99229364 |
| Williamson County, Lake Creek precinct | 0.03899946 | 0.10381926 | 37.78898912 | -88.91612676 |
| Sangamon County, Buffalo Hart township | 0.06535772 | 0.10381926 | 39.9203392 | -89.44525297 |
| De Witt County, Tunbridge township | 0.07409898 | 0.1035503 | 40.12229394 | -89.04105767 |
| Henry County, Loraine township | 0.08579882 | 0.1035503 | 41.53571692 | -89.99022229 |
| Johnson County, Vienna No. 1 precinct | 0.09359871 | 0.10328133 | 37.4108196 | -88.95222621 |
| Saline County, Brushy township | 0.07557827 | 0.10328133 | 37.75324582 | -88.68972853 |
| Monroe County, Precinct 23 | 0.09628833 | 0.10328133 | 38.23815477 | -90.23114098 |
| Knox County, Orange township | 0.05177515 | 0.10328133 | 40.87808085 | -90.28571828 |
| Livingston County, Broughton township | 0.08162991 | 0.10220549 | 40.95232548 | -88.29870628 |
| Logan County, Elkhart township | 0.07611619 | 0.10112964 | 39.9840228 | -89.38021391 |
| Lawrence County, Petty township | 0.06562668 | 0.10086068 | 38.84363377 | -87.88004753 |
| Johnson County, Belknap precinct | 0.09252286 | 0.10059172 | 37.36467036 | -88.92803097 |
| Fayette County, Carson township | 0.09319527 | 0.10059172 | 39.13515873 | -88.98946188 |
| Montgomery County, Bois D'Arc township | 0.0648198 | 0.10059172 | 39.44121859 | -89.69441519 |
| Johnson County, Grantsburg No. 2 precinct | 0.06858526 | 0.10005379 | 37.35629436 | -88.78595955 |
| Union County, Jonesboro District 2 precinct | 0.07772996 | 0.09978483 | 37.38409404 | -89.26913043 |
| Logan County, Orvil township | 0.06159225 | 0.09978483 | 40.23197549 | -89.41665576 |
| Carroll County, Fairhaven township | 0.03684777 | 0.09978483 | 42.00397827 | -89.94988972 |
| Johnson County, Elvira precinct | 0.0541958 | 0.09978483 | 37.4330663 | -88.95011289 |
| Saline County, Rector township | 0.07894029 | 0.09924691 | 37.87449898 | -88.47624153 |
| Logan County, Lake Fork township | 0.06912318 | 0.09924691 | 39.95391659 | -89.23500692 |
| Marshall County, La Prairie township | 0.06387843 | 0.09924691 | 41.04735246 | -89.6111241 |
| Christian County, Bear Creek township | 0.07772996 | 0.09924691 | 39.49129428 | -89.36660479 |
| Pulaski County, Karnak precinct | 0.09144701 | 0.09897795 | 37.29792927 | -88.93438114 |
| Clark County, Westfield township | 0.07880581 | 0.09897795 | 39.47352335 | -87.92685186 |
| DeKalb County, South Grove township | 0.04747176 | 0.09897795 | 42.03732683 | -88.84975159 |
| Jo Daviess County, Pleasant Valley township | 0.05540613 | 0.09870898 | 42.27450511 | -89.98749221 |
| Ogle County, Lynnville township | 0.05486821 | 0.09736417 | 41.9804112 | -89.03837072 |
| Williamson County, Corinth precinct | 0.05096826 | 0.09682625 | 37.77719711 | -88.71609694 |
| Lawrence County, Russell township | 0.06239914 | 0.09655729 | 38.84252075 | -87.52995087 |
| Warren County, Floyd township | 0.08405056 | 0.09655729 | 40.82044025 | -90.49991913 |
| Stephenson County, Dakota township | 0.06670253 | 0.09655729 | 42.3863339 | -89.51844496 |
| Gallatin County, Asbury township | 0.07611619 | 0.09655729 | 37.90098389 | -88.18556732 |
| Vermilion County, Pilot township | 0.07530931 | 0.09601937 | 40.17909412 | -87.75790736 |
| Warren County, Kelly township | 0.07625067 | 0.09601937 | 40.9974425 | -90.45614671 |
| McLean County, Bellflower township | 0.06724045 | 0.0957504 | 40.36399305 | -88.57077801 |
| Stephenson County, Waddams township | 0.02998924 | 0.0957504 | 42.37744642 | -89.79487526 |
| Sangamon County, Loami township | 0.0535234 | 0.09548144 | 39.69628759 | -89.87843545 |
| Hamilton County, South Twigg township | 0.06374395 | 0.09548144 | 37.94821412 | -88.58395371 |
| Johnson County, Lake No. 2 precinct | 0.05029586 | 0.09494352 | 37.58210749 | -88.82128032 |
| Gallatin County, New Haven township | 0.06804734 | 0.09494352 | 37.89730637 | -88.12835759 |
| LaSalle County, Allen township | 0.05029586 | 0.09494352 | 41.13277612 | -88.70055071 |
| Union County, Alto Pass precinct | 0.06051641 | 0.09467456 | 37.59218867 | -89.39348577 |
| Brown County, Buckhorn township | 0.07315761 | 0.09413663 | 39.91815123 | -90.83504177 |
| Brown County, Elkhorn township | 0.06643357 | 0.09413663 | 39.91571342 | -90.77611237 |
| McLean County, Dawson township | 0.07423346 | 0.09386767 | 40.46672741 | -88.79865278 |
| LaSalle County, Farm Ridge township | 0.06643357 | 0.09359871 | 41.27485951 | -88.83664456 |
| LaSalle County, Grand Rapids township | 0.05110274 | 0.09332975 | 41.20637216 | -88.81347044 |
| Ogle County, Maryland township | 0.06589564 | 0.09306079 | 42.13793642 | -89.48051442 |
| Union County, Cobden District 1 precinct | 0.05083378 | 0.0922539 | 37.51760486 | -89.25700585 |
| Saline County, Tate township | 0.07840237 | 0.09198494 | 37.88839735 | -88.70510965 |
| Effingham County, Banner township | 0.07154384 | 0.09198494 | 39.17992601 | -88.6043165 |
| Sangamon County, Maxwell township | 0.06885422 | 0.09144701 | 39.67688216 | -89.93902765 |
| Carroll County, Wysox township | 0.05513717 | 0.09144701 | 41.95071882 | -89.77191641 |
| Jo Daviess County, Berreman township | 0.06253362 | 0.09144701 | 42.27308805 | -89.95704492 |
| Johnson County, Cache precinct | 0.07988166 | 0.09144701 | 37.39022178 | -88.9827661 |
| Stephenson County, Erin township | 0.04908553 | 0.09117805 | 42.35513502 | -89.79686821 |
| Union County, Cobden District 2 precinct | 0.03980635 | 0.09090909 | 37.58277159 | -89.22241911 |
| McLean County, Arrowsmith township | 0.077461 | 0.09090909 | 40.45473034 | -88.59113098 |
| Boone County, LeRoy township | 0.077461 | 0.09037117 | 42.4334751 | -88.7169008 |
| Morgan County, Alexander precinct | 0.06387843 | 0.09010221 | 39.72982256 | -90.11150981 |
| Jackson County, Grand Tower township | 0.05957504 | 0.08983324 | 37.59527427 | -89.45051757 |
| Jasper County, Grove township | 0.06858526 | 0.08983324 | 39.14180962 | -88.34463685 |
| Menard County, Fancy Prairie No. 3 precinct | 0.07127488 | 0.0887574 | 39.99472433 | -89.58649787 |
| Champaign County, Colfax township | 0.05970952 | 0.0887574 | 40.01619803 | -88.36971841 |
| Shelby County, Pickaway township | 0.06912318 | 0.08821947 | 39.57340209 | -88.82901749 |
| Champaign County, Kerr township | 0.07127488 | 0.08821947 | 40.39634231 | -88.0011064 |
| Ogle County, Leaf River township | 0.07409898 | 0.08821947 | 42.13175049 | -89.35901475 |
| Logan County, Corwin township | 0.06320602 | 0.08795051 | 40.05999964 | -89.55101579 |
| Lee County, Harmon township | 0.05876815 | 0.08795051 | 41.6727855 | -89.58440889 |
| Lawrence County, Lukin township | 0.05123722 | 0.08768155 | 38.57123485 | -87.82312949 |
| Lawrence County, Denison township | 0.06213018 | 0.08741259 | 38.65626954 | -87.75537576 |
| Champaign County, Brown township | 0.05567509 | 0.08687466 | 40.39611285 | -88.35882499 |
| Whiteside County, Clyde township | 0.04545455 | 0.0866057 | 41.84501472 | -89.90287237 |
| LaSalle County, Hope township | 0.06213018 | 0.08552985 | 41.13380578 | -89.12185354 |
| Livingston County, Fayette township | 0.06939215 | 0.08526089 | 40.62215804 | -88.3942989 |
| Rock Island County, Zuma township | 0.06239914 | 0.08472297 | 41.5271828 | -90.29337576 |
| White County, Emma township | 0.04410974 | 0.08445401 | 38.03067733 | -88.1140838 |
| Cumberland County, Cottonwood township | 0.06750941 | 0.08445401 | 39.33465753 | -88.20673166 |
| Ogle County, Woosung township | 0.05715438 | 0.08445401 | 41.91648801 | -89.53160229 |
| Johnson County, Grantsburg No. 1 precinct | 0.07383002 | 0.08418505 | 37.41962633 | -88.7226009 |
| Whiteside County, Ustick township | 0.03926842 | 0.0831092 | 41.85348265 | -90.08445146 |
| Ogle County, Taylor township | 0.06562668 | 0.08284024 | 41.926393 | -89.28724241 |
| Stephenson County, Jefferson township | 0.0654922 | 0.08284024 | 42.27023249 | -89.89615408 |
| Johnson County, Goreville No. 2 precinct | 0.04410974 | 0.08230231 | 37.56819544 | -88.96590194 |
| Monroe County, Precinct 27 | 0.08203335 | 0.08203335 | 38.30222216 | -90.16827464 |
| Winnebago County, Harrison township | 0.04518558 | 0.08203335 | 42.39144123 | -89.18098029 |
| Sangamon County, Talkington township | 0.06239914 | 0.08176439 | 39.56127584 | -89.86047375 |
| Fulton County, Harris township | 0.06293706 | 0.08176439 | 40.52686075 | -90.43316388 |
| Saline County, Galatia township | 0.05406132 | 0.08149543 | 37.81478813 | -88.59736382 |
| Knox County, Sparta township | 0.06253362 | 0.0809575 | 41.00908974 | -90.21509463 |
| Fayette County, Kaskaskia township | 0.05244755 | 0.08068854 | 38.89210267 | -89.04206358 |
| Williamson County, Grassy precinct | 0.03819258 | 0.07880581 | 37.66696697 | -89.12864402 |
| Ogle County, Eagle Point township | 0.0626681 | 0.07799892 | 41.96710573 | -89.64868863 |
| LaSalle County, Vermillion township | 0.05271651 | 0.07719204 | 41.19573913 | -89.02544214 |
| LaSalle County, Richland township | 0.0535234 | 0.07584723 | 41.17346907 | -89.02764437 |
| Ogle County, Grand Detour township | 0.07288865 | 0.07557827 | 41.91496461 | -89.44076049 |
| Winnebago County, Shirland township | 0.04868209 | 0.07450242 | 42.48065513 | -89.17220527 |
| Whiteside County, Jordan township | 0.04922001 | 0.07450242 | 41.85710821 | -89.6890747 |
| Union County, Jonesboro District 3 precinct | 0.06172673 | 0.07423346 | 37.47467922 | -89.28951904 |
| Saline County, Long Branch township | 0.0591716 | 0.0739645 | 37.87801374 | -88.53345046 |
| Stephenson County, Loran township | 0.06186122 | 0.07261969 | 42.24358723 | -89.80681382 |
| Logan County, Oran township | 0.04948897 | 0.07073696 | 40.22099362 | -89.20971399 |
| Morgan County, Waverly No. 2 precinct | 0.06885422 | 0.06885422 | 39.61009174 | -89.94464581 |
| Whiteside County, Genesee township | 0.03604088 | 0.06374395 | 41.92841948 | -89.77390452 |
| Gallatin County, Omaha township | 0.04868209 | 0.06186122 | 37.90644998 | -88.27139254 |
| Carroll County, Elkhorn Grove township | 0.05271651 | 0.0591716 | 42.01615153 | -89.73561366 |
| Jackson County, Sand Ridge township | 0.02635826 | 0.05190963 | 37.70807695 | -89.46922084 |
