## Supplementary material for "Mapping long-distance jump dispersal risk by leveraging transport infrastructure data and sampling bias correction: a case study with the invasive spotted lanternfly (*Lycorma delicatula*)": S4 Indiana risk

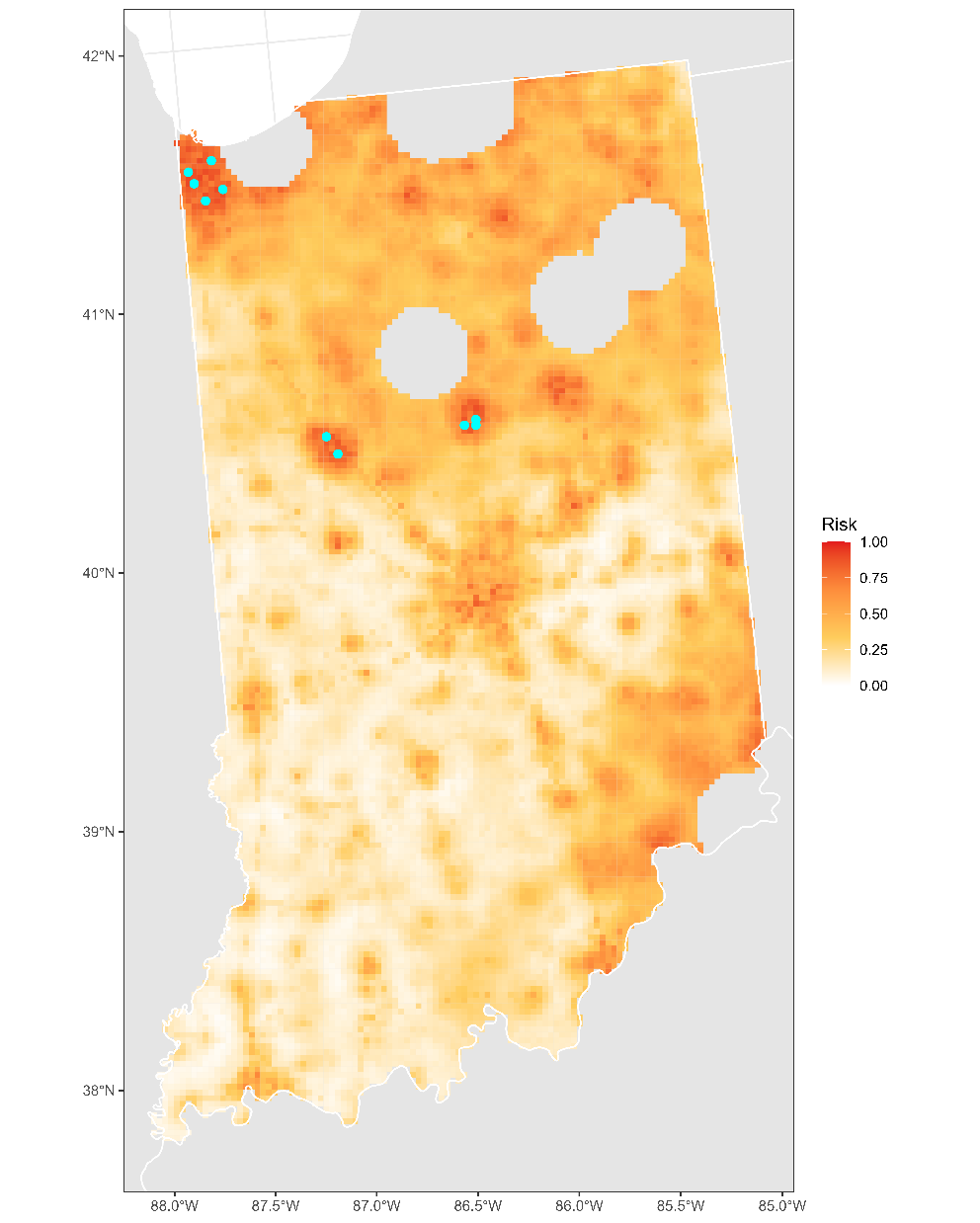

**Figure S4.1:** Projected risk of colonization by *L. delicatula* jump dispersal for Indiana. Risk projections are based on an ensemble random forest model including distances from existing *L. delicatula* populations. Note that areas falling less than 20 km from existing *L. delicatula* populations are not included in the map because any colonization there would be categorized as diffusive spread based on our methodology (regardless of the actual source of the resulting population). Darker, redder colors indicate increased risk of colonization by jump dispersal. Cyan points show the top ten highest risk locations in the state.

**Table S4.1:** Ranked list of Indiana county subdivisions sorted by the maximum projected risk of colonization by jump dispersal occurring there. For each subdivision, we identified the 2.5 x 2.5 km grid cell centroid included there with the highest projected risk of colonization by jump dispersal and report that location and risk value.

| **County, subdivision** | **Median risk value** | **Maximum risk value** | **Latitude of maximum risk** | **Longitude of maximum risk** |
| --- | --- | --- | --- | --- |
| Lake County, St. John township | 0.81118881 | 0.91662184 | 41.48124111 | -87.45348662 |
| Lake County, Ross township | 0.79532006 | 0.90398063 | 41.4488806 | -87.30604719 |
| Lake County, Calumet township | 0.79827864 | 0.9020979 | 41.56417099 | -87.35240106 |
| Howard County, Center township | 0.83781603 | 0.89564282 | 40.46419096 | -86.11179686 |
| Lake County, Center township | 0.8302851 | 0.89429801 | 41.41048272 | -87.40157974 |
| Tippecanoe County, Wea township | 0.74341044 | 0.89214632 | 40.384549 | -86.83930732 |
| Tippecanoe County, Fairfield township | 0.80043034 | 0.88596019 | 40.45547592 | -86.88990984 |
| Howard County, Taylor township | 0.58741259 | 0.88569123 | 40.44199384 | -86.11488427 |
| Lake County, North township | 0.80231307 | 0.8781603 | 41.52773804 | -87.47812786 |
| Howard County, Harrison township | 0.70575578 | 0.87036041 | 40.44661452 | -86.17401869 |
| Marshall County, West township | 0.57208176 | 0.8633674 | 41.35777386 | -86.34921684 |
| Kosciusko County, Wayne township | 0.66460463 | 0.84749866 | 41.22945448 | -85.85305645 |
| Grant County, Center township | 0.68544917 | 0.84346423 | 40.53957602 | -85.65225164 |
| Howard County, Howard township | 0.6457773 | 0.83969876 | 40.50626296 | -86.07602463 |
| Grant County, Franklin township | 0.61417429 | 0.83485745 | 40.51981047 | -85.68506769 |
| Marion County, Wayne township | 0.5570199 | 0.82947821 | 39.75398177 | -86.20965016 |
| Lake County, Hanover township | 0.74515869 | 0.82813341 | 41.39226019 | -87.46430955 |
| Madison County, Anderson township | 0.52582033 | 0.82544379 | 40.09104951 | -85.6577316 |
| Howard County, Clay township | 0.6584185 | 0.82544379 | 40.53310056 | -86.13212579 |
| Jefferson County, Madison township | 0.67778375 | 0.82436794 | 38.74609258 | -85.41083972 |
| Porter County, Morgan township | 0.64349112 | 0.81925767 | 41.42812465 | -87.005883 |
| Tippecanoe County, Perry township | 0.62479828 | 0.81253362 | 40.4046103 | -86.80687536 |
| Tippecanoe County, Sheffield township | 0.47875202 | 0.80742335 | 40.3802394 | -86.78018257 |
| Tippecanoe County, Wabash township | 0.6640667 | 0.80150619 | 40.48195126 | -86.94627514 |
| Lake County, Hobart township | 0.73547606 | 0.79989242 | 41.56009717 | -87.29225292 |
| Marion County, Center township | 0.57813341 | 0.79827864 | 39.79144689 | -86.11568803 |
| Grant County, Washington township | 0.55043034 | 0.79800968 | 40.58395022 | -85.64577809 |
| Dearborn County, Center township | 0.6781872 | 0.79424422 | 39.06000285 | -84.89527451 |
| Montgomery County, Union township | 0.26533082 | 0.79263045 | 40.02916512 | -86.88481976 |
| Porter County, Porter township | 0.63501883 | 0.79020979 | 41.41427342 | -87.12873744 |
| Grant County, Pleasant township | 0.612695 | 0.79020979 | 40.58637377 | -85.6753877 |
| Dearborn County, Lawrenceburg township | 0.73318989 | 0.78832706 | 39.1017037 | -84.85949242 |
| Marshall County, North township | 0.51613771 | 0.78563744 | 41.37772429 | -86.31619759 |
| Wayne County, Wayne township | 0.51842388 | 0.78456159 | 39.8236875 | -84.89553454 |
| St. Joseph County, Olive township | 0.60301237 | 0.77138246 | 41.70462685 | -86.48383179 |
| DeKalb County, Butler township | 0.53066165 | 0.7654653 | 41.30088644 | -85.08589173 |
| Tippecanoe County, Tippecanoe township | 0.42415277 | 0.76277569 | 40.47769139 | -86.88706319 |
| Clark County, Jeffersonville township | 0.58230231 | 0.76089295 | 38.2803249 | -85.73947272 |
| Dearborn County, Harrison township | 0.56993007 | 0.75981711 | 39.2346179 | -84.8389793 |
| Marshall County, Center township | 0.5718128 | 0.7584723 | 41.33328857 | -86.32232169 |
| Elkhart County, Locke township | 0.71005917 | 0.75658956 | 41.44350061 | -86.00400358 |
| Lake County, Eagle Creek township | 0.53200646 | 0.75605164 | 41.359915 | -87.31705126 |
| Dearborn County, Miller township | 0.62278107 | 0.75470683 | 39.2124655 | -84.84240337 |
| Grant County, Mill township | 0.66043572 | 0.7506724 | 40.49034131 | -85.59957678 |
| Tippecanoe County, Union township | 0.37331899 | 0.74986552 | 40.3909616 | -86.92800777 |
| Marion County, Warren township | 0.45911781 | 0.74905863 | 39.81131604 | -86.08332265 |
| Lake County, Winfield township | 0.63824637 | 0.74771382 | 41.42459526 | -87.27878587 |
| Porter County, Union township | 0.63824637 | 0.74556213 | 41.43859038 | -87.15594298 |
| Lake County, Cedar Creek township | 0.60018827 | 0.74475524 | 41.31949338 | -87.38248411 |
| Elkhart County, Elkhart township | 0.67563206 | 0.74018289 | 41.54034601 | -85.80801285 |
| LaGrange County, Van Buren township | 0.51062399 | 0.73668639 | 41.75280776 | -85.65516645 |
| LaPorte County, Coolspring township | 0.70521786 | 0.73534158 | 41.66205737 | -86.82390758 |
| Kosciusko County, Harrison township | 0.52461001 | 0.7288865 | 41.23420015 | -85.91283334 |
| Lake County, West Creek township | 0.47928994 | 0.72431415 | 41.34777224 | -87.46971077 |
| Delaware County, Center township | 0.55997848 | 0.72027972 | 40.18231846 | -85.40610317 |
| Jefferson County, Smyrna township | 0.57369554 | 0.7197418 | 38.77320695 | -85.46534233 |
| Marion County, Decatur township | 0.41164605 | 0.71651425 | 39.71188854 | -86.24497094 |
| Grant County, Van Buren township | 0.39725659 | 0.71382464 | 40.57663811 | -85.55696138 |
| LaGrange County, Lima township | 0.53980635 | 0.71247983 | 41.75536837 | -85.41104518 |
| Madison County, Union township | 0.50363098 | 0.71032813 | 40.08862257 | -85.62832714 |
| LaPorte County, Michigan township | 0.68370091 | 0.70898332 | 41.73306651 | -86.87541684 |
| Dearborn County, Hogan township | 0.52662722 | 0.70548682 | 39.06259994 | -84.92423763 |
| DeKalb County, Keyser township | 0.62157074 | 0.69849381 | 41.34781531 | -85.10890113 |
| Noble County, Allen township | 0.52824099 | 0.69741797 | 41.42959096 | -85.2782323 |
| Ripley County, Adams township | 0.56078537 | 0.6958042 | 39.28512016 | -85.1551027 |
| Hamilton County, Clay township | 0.46153846 | 0.69419042 | 39.92921405 | -86.15601164 |
| Elkhart County, Jackson township | 0.53173749 | 0.69150081 | 41.51813689 | -85.81124346 |
| Allen County, Perry township | 0.57853685 | 0.68746638 | 41.2419703 | -85.18582144 |
| Clark County, Silver Creek township | 0.55675094 | 0.68719742 | 38.39359944 | -85.75263158 |
| Vanderburgh County, Center township | 0.34776762 | 0.68665949 | 38.01457909 | -87.53872611 |
| Marion County, Perry township | 0.5091447 | 0.68504572 | 39.70499978 | -86.15716957 |
| Kosciusko County, Plain township | 0.54263045 | 0.6845078 | 41.27624087 | -85.87655199 |
| Wabash County, Noble township | 0.52286175 | 0.68208714 | 40.75612635 | -85.83081422 |
| LaPorte County, Wills township | 0.5591716 | 0.68020441 | 41.68684879 | -86.54706019 |
| Miami County, Peru township | 0.58633674 | 0.67939753 | 40.77497728 | -86.0683705 |
| Clinton County, Washington township | 0.49381388 | 0.6788596 | 40.27831635 | -86.61456296 |
| Floyd County, New Albany township | 0.45481442 | 0.67859064 | 38.30968171 | -85.82244733 |
| Clark County, Utica township | 0.46853147 | 0.67832168 | 38.36661202 | -85.69830497 |
| St. Joseph County, Liberty township | 0.62170522 | 0.67751479 | 41.517839 | -86.38785002 |
| LaPorte County, Clinton township | 0.52272727 | 0.67348037 | 41.44399431 | -86.91299101 |
| Noble County, Wayne township | 0.56401291 | 0.67159763 | 41.45178437 | -85.27483766 |
| Madison County, Lafayette township | 0.30043034 | 0.67052179 | 40.1378338 | -85.6807264 |
| LaPorte County, Center township | 0.60623991 | 0.66783217 | 41.63766464 | -86.79673064 |
| Miami County, Washington township | 0.47686928 | 0.66568047 | 40.75044516 | -86.04179275 |
| Elkhart County, York township | 0.61457773 | 0.66218397 | 41.75523155 | -85.68527027 |
| Ripley County, Washington township | 0.58418505 | 0.66164605 | 39.03881297 | -85.16265516 |
| Porter County, Washington township | 0.66030124 | 0.66030124 | 41.44612326 | -86.94300341 |
| Elkhart County, Clinton township | 0.5957504 | 0.65922539 | 41.57996614 | -85.74147797 |
| Dearborn County, Washington township | 0.51559978 | 0.65922539 | 39.04303552 | -84.95658411 |
| Franklin County, Ray township | 0.48951049 | 0.65788058 | 39.31487332 | -85.23899011 |
| Ripley County, Brown township | 0.55970952 | 0.6570737 | 39.01411195 | -85.13701804 |
| Delaware County, Salem township | 0.3485745 | 0.65438408 | 40.10592702 | -85.56628341 |
| Jefferson County, Milton township | 0.49112426 | 0.65411512 | 38.70640601 | -85.21218776 |
| Delaware County, Hamilton township | 0.54639591 | 0.65330823 | 40.24635829 | -85.36674904 |
| Fayette County, Connersville township | 0.36067778 | 0.65250134 | 39.64225609 | -85.13075029 |
| White County, Union township | 0.60274341 | 0.65196342 | 40.73783076 | -86.76362978 |
| Vigo County, Harrison township | 0.49946208 | 0.64954276 | 39.4403594 | -87.43030319 |
| Vanderburgh County, Pigeon township | 0.50968263 | 0.64792899 | 37.97409653 | -87.60107142 |
| Tipton County, Liberty township | 0.36202259 | 0.64470145 | 40.39760031 | -86.12105333 |
| Ripley County, Johnson township | 0.5957504 | 0.64147391 | 39.06350879 | -85.18830969 |
| Jefferson County, Shelby township | 0.48332437 | 0.63743948 | 38.82977661 | -85.3401332 |
| DeKalb County, Grant township | 0.54774072 | 0.63555675 | 41.4314376 | -85.03522085 |
| LaPorte County, Springfield township | 0.56750941 | 0.63555675 | 41.75100667 | -86.81223032 |
| Marion County, Franklin township | 0.2972028 | 0.63394298 | 39.65367466 | -86.07551849 |
| Ripley County, Laughery township | 0.52850995 | 0.63286713 | 39.26802382 | -85.2165416 |
| Kosciusko County, Prairie township | 0.54168908 | 0.63125336 | 41.27861181 | -85.90646004 |
| Jackson County, Jackson township | 0.44109736 | 0.63044648 | 38.96233556 | -85.84799503 |
| Dearborn County, Caesar Creek township | 0.6070468 | 0.6274879 | 38.98940574 | -85.11139834 |
| LaPorte County, Hudson township | 0.55849919 | 0.62587413 | 41.73131094 | -86.54104767 |
| Dearborn County, Logan township | 0.48466918 | 0.62452932 | 39.2372282 | -84.86801014 |
| LaPorte County, Kankakee township | 0.55016138 | 0.62372243 | 41.66904001 | -86.61025654 |
| LaPorte County, Cass township | 0.41070468 | 0.62345347 | 41.42176102 | -86.91586856 |
| DeKalb County, Union township | 0.48507262 | 0.62345347 | 41.36744424 | -85.07553833 |
| Dearborn County, Manchester township | 0.42711135 | 0.62318451 | 39.10949979 | -84.94643404 |
| Noble County, Orange township | 0.52299623 | 0.62318451 | 41.48149543 | -85.36136066 |
| DeKalb County, Jackson township | 0.49623453 | 0.62291555 | 41.32050852 | -85.05254474 |
| Clinton County, Center township | 0.51896181 | 0.62237762 | 40.27168448 | -86.52603095 |
| Whitley County, Richland township | 0.47875202 | 0.62157074 | 41.11896995 | -85.59729143 |
| St. Joseph County, Lincoln township | 0.54209252 | 0.62103281 | 41.48012405 | -86.48397323 |
| Howard County, Liberty township | 0.48278644 | 0.62049489 | 40.47705808 | -85.99038898 |
| St. Joseph County, Greene township | 0.54841313 | 0.62049489 | 41.56678575 | -86.44184593 |
| Jefferson County, Hanover township | 0.56939215 | 0.619688 | 38.72887183 | -85.47176496 |
| White County, Honey Creek township | 0.48977945 | 0.619688 | 40.7442978 | -86.85276444 |
| Fulton County, Rochester township | 0.44701452 | 0.61646046 | 41.07752181 | -86.20667293 |
| LaPorte County, Hanna township | 0.43249059 | 0.61592254 | 41.40884436 | -86.73588887 |
| Whitley County, Cleveland township | 0.51317913 | 0.61565358 | 41.101638 | -85.66021658 |
| Grant County, Monroe township | 0.49946208 | 0.61565358 | 40.48545223 | -85.54044451 |
| Tipton County, Prairie township | 0.35718128 | 0.61538462 | 40.40221798 | -86.18015137 |
| Elkhart County, Union township | 0.43181818 | 0.61377084 | 41.46571289 | -86.00083723 |
| DeKalb County, Richland township | 0.51237224 | 0.61350188 | 41.39474212 | -85.13194118 |
| Boone County, Center township | 0.33754707 | 0.61323292 | 40.04520768 | -86.49662286 |
| Dearborn County, Clay township | 0.57235073 | 0.6105433 | 39.00901028 | -85.07911738 |
| Jennings County, Center township | 0.58902636 | 0.6105433 | 39.01446465 | -85.66481374 |
| Wells County, Jefferson township | 0.48345885 | 0.61027434 | 40.88196481 | -85.18072792 |
| Jefferson County, Saluda township | 0.54074771 | 0.60946746 | 38.64266057 | -85.51340401 |
| Delaware County, Washington township | 0.36646046 | 0.60866057 | 40.35235708 | -85.56001054 |
| Kosciusko County, Scott township | 0.47337278 | 0.60866057 | 41.42128873 | -86.00716794 |
| Scott County, Lexington township | 0.53173749 | 0.60731576 | 38.67944977 | -85.68320536 |
| Hamilton County, Noblesville township | 0.44055944 | 0.60623991 | 40.0507249 | -85.99064347 |
| Kosciusko County, Turkey Creek township | 0.52528241 | 0.60623991 | 41.39749659 | -85.70753847 |
| Marshall County, German township | 0.50336202 | 0.60516407 | 41.44817663 | -86.06397231 |
| Clark County, Charlestown township | 0.3809844 | 0.60516407 | 38.43071283 | -85.66018058 |
| Carroll County, Jefferson township | 0.51707907 | 0.60435718 | 40.69122967 | -86.73972972 |
| Blackford County, Licking township | 0.48157612 | 0.60301237 | 40.44844123 | -85.36641721 |
| Steuben County, Jackson township | 0.45467994 | 0.60301237 | 41.6166475 | -85.09745288 |
| Kosciusko County, Clay township | 0.45320065 | 0.60220549 | 41.16521842 | -85.89251669 |
| Hendricks County, Washington township | 0.3683432 | 0.60166756 | 39.76310424 | -86.32680864 |
| Union County, Union township | 0.50161377 | 0.6013986 | 39.56953073 | -84.81652862 |
| Jefferson County, Monroe township | 0.45763852 | 0.60032275 | 38.83226989 | -85.36902038 |
| Jennings County, Vernon township | 0.44835933 | 0.60005379 | 38.98743737 | -85.61005617 |
| Scott County, Vienna township | 0.51506186 | 0.59978483 | 38.71123819 | -85.79546438 |
| Steuben County, Pleasant township | 0.48413125 | 0.5979021 | 41.63371005 | -85.03392843 |
| Ripley County, Center township | 0.51156536 | 0.59736417 | 39.11540167 | -85.26866741 |
| Marion County, Pike township | 0.47283486 | 0.59682625 | 39.91391024 | -86.24711082 |
| Decatur County, Washington township | 0.48601399 | 0.59494352 | 39.33482004 | -85.4716635 |
| LaPorte County, New Durham township | 0.59440559 | 0.59440559 | 41.52864697 | -86.8413681 |
| Grant County, Liberty township | 0.3359333 | 0.59386767 | 40.47543614 | -85.69151097 |
| Bartholomew County, German township | 0.27998924 | 0.59279182 | 39.26250144 | -85.9528408 |
| Madison County, Richland township | 0.32302313 | 0.59037117 | 40.13541217 | -85.65130196 |
| Jefferson County, Republican township | 0.52313072 | 0.58956428 | 38.69434042 | -85.59352913 |
| Johnson County, Pleasant township | 0.41312534 | 0.58687466 | 39.63380876 | -86.10781529 |
| Jasper County, Marion township | 0.38999462 | 0.58687466 | 40.94529769 | -87.15753211 |
| LaPorte County, Scipio township | 0.5380581 | 0.5866057 | 41.59103296 | -86.77250244 |
| Jennings County, Geneva township | 0.33674018 | 0.58552985 | 39.03663797 | -85.66164965 |
| Ohio County, Randolph township | 0.52487897 | 0.58499193 | 39.01309429 | -84.87312619 |
| Floyd County, Lafayette township | 0.25968263 | 0.5831092 | 38.33423961 | -85.84806838 |
| LaGrange County, Clay township | 0.49300699 | 0.58203335 | 41.64435316 | -85.42785497 |
| Decatur County, Saltcreek township | 0.48762776 | 0.58176439 | 39.32739089 | -85.38439597 |
| St. Joseph County, Union township | 0.5648198 | 0.58149543 | 41.51331408 | -86.32780065 |
| Whitley County, Thorncreek township | 0.5056482 | 0.58068854 | 41.20039451 | -85.49456595 |
| Miami County, Deer Creek township | 0.49394836 | 0.57988166 | 40.57749851 | -86.1259519 |
| Grant County, Fairmount township | 0.43114578 | 0.57934373 | 40.42621979 | -85.63885637 |
| Steuben County, Steuben township | 0.50188273 | 0.57826789 | 41.60894427 | -85.00738881 |
| Franklin County, Whitewater township | 0.54424422 | 0.57799892 | 39.32322772 | -84.82526206 |
| Allen County, Maumee township | 0.44405594 | 0.57799892 | 41.14451263 | -84.83811946 |
| Marshall County, Bourbon township | 0.49610005 | 0.57719204 | 41.31722466 | -86.1127478 |
| LaPorte County, Pleasant township | 0.53093061 | 0.5739645 | 41.58231417 | -86.65223735 |
| Whitley County, Union township | 0.51371705 | 0.57342657 | 41.12639918 | -85.41500751 |
| Wells County, Lancaster township | 0.42953201 | 0.57288865 | 40.7463553 | -85.17141616 |
| LaGrange County, Johnson township | 0.40102205 | 0.5718128 | 41.52838276 | -85.38461292 |
| LaGrange County, Greenfield township | 0.34454008 | 0.57154384 | 41.75038561 | -85.3508588 |
| Dearborn County, Kelso township | 0.50551372 | 0.57019903 | 39.28932982 | -84.94833078 |
| Newton County, Lincoln township | 0.35623991 | 0.56993007 | 41.17792461 | -87.27915422 |
| Vanderburgh County, Knight township | 0.38407746 | 0.56885422 | 37.99236245 | -87.54128149 |
| White County, Liberty township | 0.48547606 | 0.56858526 | 40.82886568 | -86.78177564 |
| Marion County, Washington township | 0.47928994 | 0.5683163 | 39.83814048 | -86.1388846 |
| Johnson County, Franklin township | 0.31455083 | 0.56724045 | 39.49604194 | -86.06774838 |
| Wabash County, Waltz township | 0.42280796 | 0.56697149 | 40.73393176 | -85.83400207 |
| Cass County, Jackson township | 0.47538999 | 0.56670253 | 40.5821144 | -86.18519956 |
| Whitley County, Columbia township | 0.50430339 | 0.56589564 | 41.11652714 | -85.56746144 |
| Franklin County, Highland township | 0.45589026 | 0.56455083 | 39.31148553 | -84.94493547 |
| Union County, Center township | 0.45373857 | 0.56293706 | 39.59691106 | -84.87142888 |
| Marshall County, Polk township | 0.46180742 | 0.5626681 | 41.40447405 | -86.37308444 |
| Putnam County, Warren township | 0.17267348 | 0.56239914 | 39.52988891 | -86.80058026 |
| Jennings County, Sand Creek township | 0.36793975 | 0.56186122 | 39.05638544 | -85.62949662 |
| Boone County, Perry township | 0.42899408 | 0.56159225 | 39.94519506 | -86.36154686 |
| Miami County, Clay township | 0.457773 | 0.56159225 | 40.57285496 | -86.06671194 |
| Franklin County, Butler township | 0.40586337 | 0.55997848 | 39.32944273 | -85.14843691 |
| Decatur County, Marion township | 0.4316837 | 0.55970952 | 39.28553749 | -85.41998685 |
| Steuben County, Salem township | 0.3986014 | 0.55944056 | 41.59445515 | -85.10091152 |
| Porter County, Pleasant township | 0.43370091 | 0.55809575 | 41.3169536 | -87.02010294 |
| LaGrange County, Newbury township | 0.48682087 | 0.55809575 | 41.65666616 | -85.57812311 |
| White County, Monon township | 0.48184508 | 0.55728886 | 40.8310242 | -86.81152405 |
| Kosciusko County, Monroe township | 0.49623453 | 0.5570199 | 41.17306917 | -85.71014483 |
| Carroll County, Deer Creek township | 0.49529317 | 0.55648198 | 40.6158512 | -86.62979802 |
| LaPorte County, Washington township | 0.47579344 | 0.55621302 | 41.53785376 | -86.65816684 |
| Scott County, Johnson township | 0.48641743 | 0.55513717 | 38.70644391 | -85.73775907 |
| Starke County, Oregon township | 0.45346961 | 0.55459925 | 41.36900699 | -86.49902674 |
| Porter County, Boone township | 0.50712749 | 0.5535234 | 41.3232435 | -87.1099929 |
| Kosciusko County, Etna township | 0.46987628 | 0.55271651 | 41.26347433 | -85.99935594 |
| Adams County, Wabash township | 0.4232114 | 0.55190963 | 40.59280994 | -84.95497176 |
| Scott County, Jennings township | 0.47848306 | 0.55164067 | 38.70884441 | -85.76661078 |
| Grant County, Jefferson township | 0.43975256 | 0.55137171 | 40.46326915 | -85.54371059 |
| Madison County, Green township | 0.32732652 | 0.55110274 | 39.99454022 | -85.84996978 |
| Morgan County, Brown township | 0.38609467 | 0.55083378 | 39.6100053 | -86.37693961 |
| Wabash County, Pleasant township | 0.49825175 | 0.5500269 | 41.00743328 | -85.88498838 |
| Elkhart County, Benton township | 0.4605971 | 0.5500269 | 41.5133424 | -85.75123215 |
| Howard County, Ervin township | 0.41554599 | 0.54895105 | 40.52009706 | -86.25360329 |
| Dearborn County, Sparta township | 0.49233459 | 0.54895105 | 39.05588196 | -85.10139732 |
| Rush County, Rushville township | 0.32786444 | 0.54868209 | 39.6230335 | -85.42942414 |
| Marion County, Lawrence township | 0.43988704 | 0.54841313 | 39.91993128 | -86.03860431 |
| Wayne County, Center township | 0.28792361 | 0.5478752 | 39.87319586 | -84.94723775 |
| Wabash County, Lagro township | 0.46153846 | 0.54679935 | 40.81792506 | -85.76181441 |
| Monroe County, Van Buren township | 0.17872512 | 0.54572351 | 39.1550606 | -86.58417226 |
| Allen County, Eel River township | 0.464766 | 0.54384077 | 41.24450249 | -85.21569101 |
| Allen County, Cedar Creek township | 0.51721356 | 0.542227 | 41.22404988 | -84.97679482 |
| LaGrange County, Bloomfield township | 0.45225928 | 0.54142012 | 41.64186952 | -85.39780769 |
| Boone County, Washington township | 0.26519634 | 0.54115116 | 40.16066434 | -86.54076539 |
| Bartholomew County, Columbus township | 0.3809844 | 0.54034427 | 39.21578238 | -85.92994808 |
| Franklin County, Springfield township | 0.46369016 | 0.54007531 | 39.43660841 | -84.837178 |
| Elkhart County, Middlebury township | 0.52434104 | 0.53926842 | 41.62438408 | -85.7349649 |
| Kosciusko County, Washington township | 0.47108661 | 0.53899946 | 41.19767891 | -85.73676626 |
| Fulton County, Richland township | 0.41958042 | 0.5387305 | 41.10202577 | -86.23343077 |
| Kosciusko County, Lake township | 0.46637977 | 0.53846154 | 41.04707392 | -85.81899745 |
| Kosciusko County, Van Buren township | 0.46046261 | 0.53711673 | 41.4293045 | -85.82414551 |
| Adams County, Washington township | 0.4908553 | 0.53630984 | 40.7480008 | -84.93073839 |
| Ripley County, Delaware township | 0.44566971 | 0.53630984 | 39.13504752 | -85.23637293 |
| Pulaski County, Monroe township | 0.45158687 | 0.53604088 | 41.0401356 | -86.6035617 |
| Wabash County, Chester township | 0.48789672 | 0.53577192 | 41.0224845 | -85.79240407 |
| Jasper County, Carpenter township | 0.34521248 | 0.53550296 | 40.76332388 | -87.12026469 |
| Ripley County, Otter Creek township | 0.44136633 | 0.53496503 | 39.09825003 | -85.329936 |
| Carroll County, Madison township | 0.49004841 | 0.53335126 | 40.52700625 | -86.64151682 |
| DeKalb County, Fairfield township | 0.42159763 | 0.53335126 | 41.52532328 | -85.08128816 |
| Marshall County, Tippecanoe township | 0.31603012 | 0.53173749 | 41.27280127 | -86.11899474 |
| Tippecanoe County, Washington township | 0.46799354 | 0.53146853 | 40.49346332 | -86.79538273 |
| LaPorte County, Galena township | 0.48655191 | 0.53119957 | 41.71350427 | -86.60428559 |
| Monroe County, Richland township | 0.26869285 | 0.53066165 | 39.17725689 | -86.58128308 |
| Miami County, Pipe Creek township | 0.49354492 | 0.53039268 | 40.73289964 | -86.10428253 |
| Jackson County, Redding township | 0.28980635 | 0.52985476 | 38.98451332 | -85.84488904 |
| Kosciusko County, Jefferson township | 0.41837009 | 0.5295858 | 41.41894101 | -85.97719592 |
| White County, Big Creek township | 0.39873588 | 0.52877891 | 40.67334226 | -86.80199349 |
| Kosciusko County, Jackson township | 0.47068316 | 0.52850995 | 41.10405912 | -85.69004348 |
| Jefferson County, Graham township | 0.42980097 | 0.52824099 | 38.76814124 | -85.67061792 |
| Hendricks County, Guilford township | 0.37116729 | 0.52770307 | 39.67658852 | -86.36802058 |
| Ripley County, Franklin township | 0.47928994 | 0.52716514 | 39.12745418 | -85.14936988 |
| Blackford County, Jackson township | 0.37547068 | 0.5260893 | 40.39658324 | -85.28448754 |
| Starke County, Center township | 0.48050027 | 0.52582033 | 41.28676745 | -86.60083201 |
| DeKalb County, Smithfield township | 0.40115654 | 0.52528241 | 41.52275901 | -85.05130418 |
| Clinton County, Perry township | 0.33754707 | 0.52474449 | 40.25390675 | -86.58798288 |
| Dubois County, Bainbridge township | 0.35718128 | 0.52313072 | 38.37535946 | -86.94577144 |
| Adams County, Monroe township | 0.4549489 | 0.52286175 | 40.6593188 | -84.94459898 |
| Fayette County, Harrison township | 0.25174825 | 0.52259279 | 39.66441916 | -85.12739443 |
| Miami County, Harrison township | 0.46140398 | 0.52178591 | 40.61257665 | -86.00122506 |
| Switzerland County, Pleasant township | 0.52178591 | 0.52178591 | 38.92801132 | -85.17921627 |
| Howard County, Honey Creek township | 0.42334588 | 0.52071006 | 40.4290077 | -86.23620984 |
| Jasper County, Keener township | 0.45669715 | 0.52017214 | 41.13140483 | -87.25476576 |
| Allen County, Jackson township | 0.43141474 | 0.51990317 | 41.0114653 | -84.85918778 |
| Franklin County, Bath township | 0.41810113 | 0.51855836 | 39.48091561 | -84.83030331 |
| Madison County, Adams township | 0.302851 | 0.51855836 | 40.04668753 | -85.66415328 |
| Delaware County, Yorktown town | 0.39698763 | 0.51721356 | 40.16755658 | -85.49768213 |
| Miami County, Butler township | 0.42428725 | 0.51559978 | 40.74810689 | -86.01210247 |
| Allen County, Milan township | 0.48534158 | 0.51559978 | 41.17710263 | -84.95391349 |
| Wells County, Harrison township | 0.43343195 | 0.51425498 | 40.72417794 | -85.17480939 |
| Dearborn County, Jackson township | 0.43894567 | 0.51075847 | 39.25017818 | -85.01319557 |
| Clark County, Monroe township | 0.25779989 | 0.51022055 | 38.59796819 | -85.78218206 |
| Carroll County, Tippecanoe township | 0.45642819 | 0.51022055 | 40.57580585 | -86.69493577 |
| Carroll County, Clay township | 0.43491124 | 0.50968263 | 40.48038898 | -86.61776722 |
| Kosciusko County, Tippecanoe township | 0.48278644 | 0.5091447 | 41.35067397 | -85.68411475 |
| Hamilton County, Delaware township | 0.47633136 | 0.50860678 | 39.98415847 | -85.99995898 |
| Starke County, Davis township | 0.41016676 | 0.50779989 | 41.37566285 | -86.58893548 |
| Hamilton County, Westfield Washington township | 0.33916084 | 0.50645508 | 40.04017511 | -86.14072946 |
| Putnam County, Greencastle township | 0.38327058 | 0.50591716 | 39.64735525 | -86.87415581 |
| Clinton County, Jackson township | 0.32867133 | 0.50591716 | 40.24948004 | -86.52898149 |
| Whitley County, Smith township | 0.46288327 | 0.50537924 | 41.25456159 | -85.33519065 |
| Noble County, Swan township | 0.41810113 | 0.50511027 | 41.31106725 | -85.20546839 |
| Boone County, Zionsville town | 0.27541689 | 0.50484131 | 39.95829828 | -86.24106123 |
| Madison County, Monroe township | 0.40640129 | 0.50457235 | 40.27335025 | -85.69092428 |
| Vanderburgh County, Perry township | 0.33189887 | 0.50322754 | 37.97606192 | -87.62969177 |
| Delaware County, Niles township | 0.38327058 | 0.50268962 | 40.37691485 | -85.31733944 |
| LaPorte County, Johnson township | 0.43020441 | 0.50268962 | 41.48457578 | -86.54400421 |
| Miami County, Erie township | 0.39147391 | 0.50242066 | 40.77030614 | -86.00896964 |
| Monroe County, Bloomington township | 0.19123185 | 0.50215169 | 39.17504639 | -86.55221924 |
| Grant County, Sims township | 0.48184508 | 0.50215169 | 40.55161681 | -85.80022864 |
| Noble County, Jefferson township | 0.4084185 | 0.50161377 | 41.43460607 | -85.33813938 |
| LaGrange County, Clearspring township | 0.38179129 | 0.50134481 | 41.5999501 | -85.43456401 |
| Clinton County, Union township | 0.43437332 | 0.50107585 | 40.31831296 | -86.54965135 |
| Monroe County, Perry township | 0.34642281 | 0.50080689 | 39.10845967 | -86.56090787 |
| Tippecanoe County, Shelby township | 0.30016138 | 0.50026896 | 40.46396522 | -87.00830426 |
| Dearborn County, York township | 0.41769769 | 0.49946208 | 39.17596408 | -84.93626539 |
| Marshall County, Union township | 0.42361485 | 0.49946208 | 41.26890247 | -86.36141228 |
| Fulton County, Aubbeenaubbee township | 0.44244217 | 0.49838623 | 41.16456225 | -86.46624198 |
| LaPorte County, Noble township | 0.43114578 | 0.49838623 | 41.50426126 | -86.81423664 |
| Grant County, Richland township | 0.40478752 | 0.49811727 | 40.57380771 | -85.79703843 |
| Jay County, Wayne township | 0.41635288 | 0.49784831 | 40.43763085 | -84.97909972 |
| Harrison County, Harrison township | 0.27138246 | 0.49784831 | 38.22226878 | -86.12449063 |
| Starke County, Washington township | 0.45669715 | 0.49704142 | 41.32900423 | -86.56493474 |
| Howard County, Jackson township | 0.42684239 | 0.49596557 | 40.48980887 | -85.86893748 |
| Madison County, Fall Creek township | 0.36874664 | 0.4943518 | 40.02933543 | -85.72612212 |
| Madison County, Pipe Creek township | 0.39443249 | 0.49408284 | 40.260772 | -85.8120399 |
| Adams County, Root township | 0.44002152 | 0.49327595 | 40.86145429 | -84.94306242 |
| Pulaski County, Salem township | 0.38058096 | 0.49327595 | 40.92420618 | -86.85954423 |
| Jasper County, Union township | 0.2880581 | 0.49112426 | 41.12934945 | -87.22487771 |
| Jasper County, Wheatfield township | 0.45118343 | 0.49004841 | 41.20789259 | -87.06419292 |
| Boone County, Worth township | 0.28321678 | 0.49004841 | 39.96964835 | -86.38793564 |
| Hendricks County, Brown township | 0.3331092 | 0.48816568 | 39.85866131 | -86.40282135 |
| White County, Prairie township | 0.38246369 | 0.48816568 | 40.61098966 | -86.86994593 |
| Noble County, Sparta township | 0.39779451 | 0.48762776 | 41.39265423 | -85.64763702 |
| LaGrange County, Eden township | 0.41568047 | 0.48762776 | 41.55050863 | -85.65467366 |
| Jennings County, Campbell township | 0.33647122 | 0.4873588 | 39.00472669 | -85.54894716 |
| Kosciusko County, Seward township | 0.40989779 | 0.48655191 | 41.1675854 | -85.92237829 |
| Ripley County, Shelby township | 0.33485745 | 0.48574502 | 39.00959016 | -85.34299589 |
| Wabash County, Liberty township | 0.39214632 | 0.48547606 | 40.72915129 | -85.77465114 |
| Carroll County, Burlington township | 0.43222162 | 0.4852071 | 40.48477319 | -86.37804359 |
| White County, Lincoln township | 0.43908015 | 0.48466918 | 40.7513249 | -86.64187469 |
| Miami County, Jefferson township | 0.37399139 | 0.48466918 | 40.79717836 | -86.06525407 |
| Miami County, Jackson township | 0.45051103 | 0.48305541 | 40.63007001 | -85.93881558 |
| Wells County, Union township | 0.47821409 | 0.48305541 | 40.86484161 | -85.24354412 |
| Union County, Brownsville township | 0.38542227 | 0.48278644 | 39.67117427 | -84.94874156 |
| Allen County, Springfield township | 0.37130178 | 0.48251748 | 41.19928213 | -84.95042912 |
| Marshall County, Green township | 0.44513179 | 0.48224852 | 41.17777558 | -86.34369539 |
| Hendricks County, Lincoln township | 0.39537386 | 0.48063475 | 39.83421236 | -86.37646822 |
| Pulaski County, Harrison township | 0.4020979 | 0.4795589 | 41.01571065 | -86.57669934 |
| Wells County, Rockcreek township | 0.39766003 | 0.4795589 | 40.75141749 | -85.2307398 |
| Allen County, Jefferson township | 0.4493007 | 0.47902098 | 41.03884972 | -84.91522348 |
| Pulaski County, Beaver township | 0.4140667 | 0.47821409 | 40.91990431 | -86.7999681 |
| Fulton County, Henry township | 0.42912856 | 0.47740721 | 41.01451768 | -85.97437877 |
| White County, Cass township | 0.43719742 | 0.4760624 | 40.88241307 | -86.59446542 |
| Howard County, Monroe township | 0.40478752 | 0.47552448 | 40.45349375 | -86.26273463 |
| Vigo County, Lost Creek township | 0.25255514 | 0.47552448 | 39.47873181 | -87.3373885 |
| Hancock County, Buck Creek township | 0.29074771 | 0.47471759 | 39.81943437 | -85.90440213 |
| Franklin County, Brookville township | 0.42616998 | 0.4739107 | 39.42226464 | -84.9279277 |
| Wabash County, Paw Paw township | 0.39093599 | 0.47364174 | 40.86948815 | -85.84457857 |
| Fayette County, Waterloo township | 0.27030662 | 0.47337278 | 39.66186898 | -85.09818227 |
| Blackford County, Harrison township | 0.34655729 | 0.47229693 | 40.55432894 | -85.29062173 |
| Ripley County, Jackson township | 0.4429801 | 0.47175901 | 39.164745 | -85.32012015 |
| Vigo County, Honey Creek township | 0.4190425 | 0.47149005 | 39.3474672 | -87.38253357 |
| Delaware County, Liberty township | 0.25739645 | 0.47095212 | 40.19701394 | -85.31448653 |
| Vigo County, Otter Creek township | 0.25255514 | 0.47014524 | 39.52519033 | -87.36127233 |
| Clinton County, Madison township | 0.35906401 | 0.46987628 | 40.30272069 | -86.6411612 |
| Starke County, California township | 0.37573964 | 0.46960732 | 41.24232207 | -86.60676904 |
| LaPorte County, Lincoln township | 0.38636364 | 0.46933835 | 41.54903089 | -86.504961 |
| Pulaski County, Indian Creek township | 0.43020441 | 0.46880043 | 40.99569962 | -86.60947634 |
| Floyd County, Georgetown township | 0.25685853 | 0.46853147 | 38.31680389 | -85.90854863 |
| Pulaski County, Tippecanoe township | 0.40801506 | 0.46853147 | 41.14234495 | -86.4692486 |
| Jefferson County, Lancaster township | 0.38690156 | 0.46853147 | 38.84217566 | -85.48458878 |
| Starke County, North Bend township | 0.42011834 | 0.46772458 | 41.25789988 | -86.51403286 |
| Jennings County, Columbia township | 0.4042496 | 0.46718666 | 39.15499882 | -85.46846928 |
| LaGrange County, Milford township | 0.38555675 | 0.4669177 | 41.54056199 | -85.26123768 |
| Morgan County, Monroe township | 0.2711135 | 0.4669177 | 39.61673989 | -86.46464602 |
| Noble County, Green township | 0.41299086 | 0.4669177 | 41.27925103 | -85.36171185 |
| Johnson County, White River township | 0.3161646 | 0.46611081 | 39.61624167 | -86.16932206 |
| Fulton County, Newcastle township | 0.34965035 | 0.46584185 | 41.0951224 | -86.14390406 |
| Carroll County, Monroe township | 0.39402905 | 0.46557289 | 40.54481318 | -86.57934077 |
| Knox County, Vincennes township | 0.19311458 | 0.46530393 | 38.68502386 | -87.51913066 |
| Howard County, Union township | 0.34938139 | 0.46503497 | 40.47235154 | -85.93124136 |
| Fayette County, Jennings township | 0.39174287 | 0.464766 | 39.59282789 | -85.07908671 |
| Gibson County, Johnson township | 0.1520979 | 0.46422808 | 38.17207633 | -87.54948967 |
| Adams County, Blue Creek township | 0.42038731 | 0.46395912 | 40.74018717 | -84.84179306 |
| Noble County, Noble township | 0.44136633 | 0.46369016 | 41.28917401 | -85.48129091 |
| Delaware County, Union township | 0.39698763 | 0.46261431 | 40.31288561 | -85.35680973 |
| Miami County, Perry township | 0.40182894 | 0.46153846 | 40.98996133 | -85.9477437 |
| Delaware County, Harrison township | 0.3957773 | 0.45992469 | 40.2858124 | -85.56976624 |
| Clinton County, Michigan township | 0.39846692 | 0.45911781 | 40.31163542 | -86.46107586 |
| Hendricks County, Center township | 0.27299623 | 0.45884884 | 39.7543842 | -86.50553907 |
| Shelby County, Addison township | 0.35745024 | 0.45857988 | 39.54135953 | -85.79580439 |
| Franklin County, Metamora township | 0.42590102 | 0.45723507 | 39.4402499 | -85.1317367 |
| Pulaski County, Franklin township | 0.35543303 | 0.45669715 | 41.08457295 | -86.59763962 |
| Marshall County, Walnut township | 0.35502959 | 0.45615923 | 41.2598022 | -86.24174825 |
| LaPorte County, Prairie township | 0.39039806 | 0.4556213 | 41.34215998 | -86.7446793 |
| Carroll County, Jackson township | 0.38542227 | 0.45454545 | 40.56702285 | -86.57639369 |
| Henry County, Spiceland township | 0.21167294 | 0.45427649 | 39.84722696 | -85.42599881 |
| Jasper County, Kankakee township | 0.41325982 | 0.45346961 | 41.22802393 | -87.03144665 |
| Steuben County, Jamestown township | 0.35718128 | 0.4521248 | 41.74209156 | -84.98642229 |
| Clark County, Washington township | 0.41043572 | 0.45158687 | 38.6032193 | -85.57739165 |
| Grant County, Green township | 0.2647929 | 0.45158687 | 40.46285563 | -85.81296965 |
| Vigo County, Sugar Creek township | 0.20091447 | 0.45104895 | 39.50700976 | -87.4223763 |
| Pulaski County, Van Buren township | 0.37385691 | 0.45104895 | 40.99349357 | -86.579665 |
| Clark County, Owen township | 0.39618074 | 0.45077999 | 38.4899266 | -85.56446613 |
| Noble County, Elkhart township | 0.41312534 | 0.45077999 | 41.46922081 | -85.48461488 |
| Adams County, St. Marys township | 0.43343195 | 0.45051103 | 40.75973654 | -84.80864192 |
| Gibson County, Patoka township | 0.23372781 | 0.4499731 | 38.3537619 | -87.5865045 |
| Cass County, Boone township | 0.42724583 | 0.44970414 | 40.88020371 | -86.56470216 |
| Wayne County, Boston township | 0.3119957 | 0.44889726 | 39.77676849 | -84.87315106 |
| Allen County, Monroe township | 0.38542227 | 0.44862829 | 41.00885043 | -84.82942946 |
| Jay County, Greene township | 0.33781603 | 0.44835933 | 40.44278178 | -85.03815297 |
| Whitley County, Washington township | 0.42267348 | 0.44835933 | 41.06969084 | -85.54422342 |
| White County, Princeton township | 0.34440559 | 0.44835933 | 40.73901334 | -87.09332979 |
| Noble County, Washington township | 0.4020979 | 0.44755245 | 41.32362062 | -85.62753426 |
| Jennings County, Spencer township | 0.32840237 | 0.4464766 | 39.01688225 | -85.69378522 |
| Henry County, Franklin township | 0.18988704 | 0.4464766 | 39.86691815 | -85.39341863 |
| Warrick County, Ohio township | 0.21812803 | 0.44593868 | 37.9601515 | -87.40077208 |
| Whitley County, Etna-Troy township | 0.40008069 | 0.44486283 | 41.22750354 | -85.55099672 |
| Johnson County, Needham township | 0.28079613 | 0.44459387 | 39.49371133 | -86.03857249 |
| Clark County, Union township | 0.25968263 | 0.44432491 | 38.4846972 | -85.76894081 |
| Union County, Harrison township | 0.37143626 | 0.44378698 | 39.66858243 | -84.91953304 |
| Daviess County, Washington township | 0.21637977 | 0.44351802 | 38.65861167 | -87.14361954 |
| LaPorte County, Union township | 0.3781603 | 0.44163529 | 41.49558234 | -86.69411468 |
| Starke County, Jackson township | 0.37331899 | 0.44136633 | 41.33998486 | -86.71471722 |
| Parke County, Adams township | 0.2232383 | 0.4408284 | 39.76138767 | -87.21457618 |
| Noble County, Perry township | 0.41070468 | 0.44029048 | 41.52099339 | -85.56794213 |
| Ohio County, Union township | 0.43975256 | 0.43975256 | 38.99613803 | -84.93440143 |
| Greene County, Wright township | 0.13972566 | 0.43867671 | 39.15547791 | -87.20013111 |
| Jennings County, Bigger township | 0.3479021 | 0.43840775 | 38.98010575 | -85.52319113 |
| Adams County, French township | 0.38891877 | 0.43760086 | 40.66448654 | -85.00383377 |
| Clinton County, Ross township | 0.37762238 | 0.43652501 | 40.41376161 | -86.62655361 |
| Union County, Liberty township | 0.34171598 | 0.43625605 | 39.64901597 | -84.95215263 |
| Carroll County, Adams township | 0.36525013 | 0.43625605 | 40.72911085 | -86.64480895 |
| Tippecanoe County, Lauramie township | 0.28066165 | 0.43571813 | 40.29569794 | -86.85072785 |
| Huntington County, Salamonie township | 0.40935987 | 0.43544917 | 40.69740329 | -85.38906721 |
| Adams County, Hartford township | 0.41500807 | 0.4351802 | 40.64231499 | -85.00727532 |
| Jasper County, Walker township | 0.36713287 | 0.43410436 | 41.1634294 | -87.06983185 |
| Clinton County, Owen township | 0.37735342 | 0.43410436 | 40.36050441 | -86.51421041 |
| Shelby County, Brandywine township | 0.27353416 | 0.43222162 | 39.58810756 | -85.81871353 |
| Fayette County, Jackson township | 0.39429801 | 0.43195266 | 39.59538245 | -85.10827024 |
| Fulton County, Liberty township | 0.37452932 | 0.43195266 | 40.9465589 | -86.25492974 |
| Warren County, Liberty township | 0.17388381 | 0.4316837 | 40.3028314 | -87.26701093 |
| Newton County, Lake township | 0.30648198 | 0.4316837 | 41.16783636 | -87.46133655 |
| DeKalb County, Wilmington township | 0.38192577 | 0.4316837 | 41.44067644 | -84.88201337 |
| Carroll County, Democrat township | 0.39080151 | 0.43114578 | 40.47153447 | -86.49939334 |
| DeKalb County, Franklin township | 0.36847768 | 0.42953201 | 41.44328003 | -84.91195619 |
| White County, Jackson township | 0.40545992 | 0.42845616 | 40.77134046 | -86.60921956 |
| Dubois County, Patoka township | 0.1788596 | 0.42764927 | 38.30874998 | -86.95400858 |
| Jackson County, Vernon township | 0.3006993 | 0.42684239 | 38.73818281 | -85.85008329 |
| Morgan County, Washington township | 0.20790748 | 0.42684239 | 39.4347004 | -86.42980602 |
| LaGrange County, Springfield township | 0.39376009 | 0.4260355 | 41.64901663 | -85.21414554 |
| Delaware County, Monroe township | 0.28536848 | 0.42549758 | 40.13548318 | -85.38327353 |
| Clark County, Oregon township | 0.38488435 | 0.42495966 | 38.51453803 | -85.59006364 |
| White County, West Point township | 0.31966111 | 0.42388381 | 40.66815441 | -87.04232278 |
| Steuben County, Fremont township | 0.23345885 | 0.42307692 | 41.71731314 | -84.95985844 |
| Madison County, Duck Creek township | 0.29451318 | 0.422539 | 40.30753241 | -85.83519028 |
| LaPorte County, Dewey township | 0.37896719 | 0.422539 | 41.33069956 | -86.8973969 |
| Shelby County, Marion township | 0.23937601 | 0.42200108 | 39.56353909 | -85.79265745 |
| Decatur County, Sandcreek township | 0.35476062 | 0.42038731 | 39.15991898 | -85.5265132 |
| Whitley County, Jefferson township | 0.40962883 | 0.42011834 | 41.08201854 | -85.4216755 |
| Noble County, York township | 0.37869822 | 0.41958042 | 41.42235435 | -85.4613156 |
| Jennings County, Montgomery township | 0.38138784 | 0.41931146 | 38.87900475 | -85.65484053 |
| Miami County, Allen township | 0.3577192 | 0.41823561 | 40.93967156 | -86.16559728 |
| Adams County, Kirkland township | 0.38327058 | 0.41742873 | 40.80009989 | -85.01281474 |
| Jay County, Pike township | 0.24610005 | 0.4168908 | 40.39329596 | -84.98597417 |
| Clinton County, Warren township | 0.39107047 | 0.41662184 | 40.39595937 | -86.3900541 |
| Johnson County, Clark township | 0.1753631 | 0.41662184 | 39.53808192 | -86.03242995 |
| Clinton County, Forest township | 0.35906401 | 0.41608392 | 40.41364141 | -86.32792974 |
| Wells County, Liberty township | 0.36928456 | 0.41554599 | 40.72923847 | -85.23411476 |
| Wayne County, Abington township | 0.35233997 | 0.41393222 | 39.74023721 | -84.96773293 |
| DeKalb County, Spencer township | 0.39093599 | 0.4133943 | 41.28800358 | -84.93646975 |
| Wells County, Chester township | 0.34776762 | 0.41285637 | 40.57650674 | -85.28727265 |
| Fountain County, Logan township | 0.32409898 | 0.41258741 | 40.30078015 | -87.23746886 |
| Franklin County, Laurel township | 0.35180204 | 0.41231845 | 39.44532746 | -85.18998637 |
| White County, Round Grove township | 0.34736417 | 0.41231845 | 40.64382903 | -87.01545053 |
| Vigo County, Riley township | 0.22485207 | 0.41204949 | 39.4322694 | -87.31353536 |
| Clark County, Bethlehem township | 0.3894567 | 0.41043572 | 38.52689855 | -85.4717984 |
| Scott County, Finley township | 0.31118881 | 0.40855299 | 38.66927198 | -85.83051943 |
| Wells County, Nottingham township | 0.37197418 | 0.40855299 | 40.6031077 | -85.07334075 |
| Hamilton County, Wayne township | 0.27824099 | 0.40667025 | 40.00164493 | -85.93809555 |
| DeKalb County, Concord township | 0.38273265 | 0.40586337 | 41.33236644 | -84.92947689 |
| Carroll County, Carrollton township | 0.39429801 | 0.40371167 | 40.55364069 | -86.39863584 |
| Pulaski County, Rich Grove township | 0.3844809 | 0.40290479 | 41.11773609 | -86.74399218 |
| Vanderburgh County, Scott township | 0.2626412 | 0.40263583 | 38.10542674 | -87.55715783 |
| Steuben County, Millgrove township | 0.33458849 | 0.40263583 | 41.73526877 | -85.17035129 |
| Adams County, Union township | 0.36941904 | 0.40182894 | 40.85103282 | -84.82428156 |
| Parke County, Washington township | 0.19661108 | 0.40155998 | 39.78360228 | -87.21185714 |
| Jennings County, Lovett township | 0.36148467 | 0.40129102 | 38.89387501 | -85.5649129 |
| Crawford County, Jennings township | 0.27124798 | 0.40048413 | 38.24056352 | -86.35393336 |
| Huntington County, Jefferson township | 0.35395374 | 0.40021517 | 40.68017976 | -85.45167006 |
| Franklin County, Fairfield township | 0.37022593 | 0.39994621 | 39.49132481 | -84.94683377 |
| Wayne County, Webster township | 0.29182356 | 0.39967725 | 39.89535604 | -84.94381452 |
| Kosciusko County, Franklin township | 0.37493276 | 0.39967725 | 41.10568214 | -85.9915788 |
| Steuben County, Scott township | 0.3535503 | 0.39967725 | 41.67293162 | -84.96687045 |
| Vermillion County, Highland township | 0.14792899 | 0.39940828 | 40.11697023 | -87.49771405 |
| Greene County, Taylor township | 0.18141474 | 0.39913932 | 38.91247845 | -86.93726795 |
| Henry County, Henry township | 0.25268962 | 0.39806347 | 39.88909018 | -85.39013328 |
| Jennings County, Marion township | 0.29639591 | 0.39752555 | 38.85925125 | -85.68690845 |
| Jay County, Bearcreek township | 0.32947821 | 0.39698763 | 40.56804997 | -84.92884725 |
| Blackford County, Washington township | 0.35314685 | 0.39698763 | 40.49528815 | -85.38933599 |
| Allen County, Madison township | 0.36605702 | 0.39618074 | 40.93056596 | -84.96236729 |
| Jay County, Jackson township | 0.36740183 | 0.39564282 | 40.48454719 | -85.0017604 |
| Adams County, Preble township | 0.37856374 | 0.39456697 | 40.86662348 | -85.00246589 |
| Tipton County, Jefferson township | 0.33055406 | 0.39429801 | 40.28662039 | -86.13644245 |
| Starke County, Wayne township | 0.34991931 | 0.39349112 | 41.2266773 | -86.69946421 |
| Steuben County, Otsego township | 0.30365788 | 0.3929532 | 41.60377377 | -84.9473571 |
| Fulton County, Wayne township | 0.36188811 | 0.39268424 | 40.94017244 | -86.46643801 |
| Franklin County, Salt Creek township | 0.30688542 | 0.39187735 | 39.38136446 | -85.22906132 |
| Hamilton County, Fall Creek township | 0.34763314 | 0.39107047 | 39.93743622 | -85.9768056 |
| Cass County, Deer Creek township | 0.38784293 | 0.38784293 | 40.56678349 | -86.27712763 |
| Pulaski County, Jefferson township | 0.32409898 | 0.38757396 | 41.07111737 | -86.71997671 |
| Franklin County, Blooming Grove township | 0.35637439 | 0.38676708 | 39.4769011 | -85.0376303 |
| Vanderburgh County, German township | 0.12022593 | 0.38622916 | 38.02049885 | -87.62463388 |
| Montgomery County, Franklin township | 0.12735342 | 0.38596019 | 40.04493161 | -86.79371203 |
| Washington County, Washington township | 0.28671329 | 0.38488435 | 38.60402191 | -86.13103441 |
| Rush County, Jackson township | 0.12694997 | 0.38407746 | 39.64520431 | -85.42616098 |
| Fulton County, Union township | 0.35435718 | 0.38300161 | 41.07121444 | -86.41857718 |
| DeKalb County, Newville township | 0.36282948 | 0.3816568 | 41.28019025 | -84.84684289 |
| Crawford County, Union township | 0.29827864 | 0.380312 | 38.23402542 | -86.55764943 |
| Delaware County, Delaware township | 0.29316837 | 0.37977407 | 40.26353528 | -85.30450547 |
| DeKalb County, Stafford township | 0.33297472 | 0.37923615 | 41.43544831 | -84.82213439 |
| Fayette County, Columbia township | 0.26089295 | 0.37896719 | 39.55614404 | -85.17332312 |
| Lawrence County, Shawswick township | 0.17576654 | 0.37869822 | 38.83547296 | -86.50871693 |
| Tipton County, Madison township | 0.26519634 | 0.37842926 | 40.28772239 | -85.8678492 |
| Huntington County, Wayne township | 0.34615385 | 0.3781603 | 40.67270165 | -85.63280674 |
| Jasper County, Hanging Grove township | 0.27487897 | 0.3781603 | 40.93060677 | -86.94892194 |
| Decatur County, Clay township | 0.25551372 | 0.3781603 | 39.34218812 | -85.55894886 |
| Pulaski County, Cass township | 0.34830554 | 0.37762238 | 41.12421869 | -86.83360407 |
| Randolph County, White River township | 0.28752017 | 0.37735342 | 40.16647495 | -84.96139032 |
| Decatur County, Clinton township | 0.21624529 | 0.37708445 | 39.38162423 | -85.49429778 |
| Union County, Harmony township | 0.32409898 | 0.37708445 | 39.55779586 | -84.93661171 |
| Cass County, Adams township | 0.36121571 | 0.37627757 | 40.89985543 | -86.23129402 |
| Crawford County, Liberty township | 0.29827864 | 0.37600861 | 38.37369907 | -86.33637029 |
| Jasper County, Jordan township | 0.25511027 | 0.37547068 | 40.83415641 | -87.17141789 |
| Tipton County, Cicero township | 0.29316837 | 0.37358795 | 40.28430496 | -86.10694166 |
| Madison County, Van Buren township | 0.35556751 | 0.37358795 | 40.31529804 | -85.65500022 |
| Noble County, Albion township | 0.37224314 | 0.37224314 | 41.39768224 | -85.43470563 |
| Tippecanoe County, Randolph township | 0.16514255 | 0.37062937 | 40.2978396 | -86.8802565 |
| Miami County, Union township | 0.35368478 | 0.36982249 | 40.9174656 | -86.16868855 |
| Adams County, Jefferson township | 0.35960194 | 0.36955352 | 40.62673401 | -84.82965144 |
| Floyd County, Franklin township | 0.22834857 | 0.36928456 | 38.22335284 | -85.86343747 |
| Madison County, Stony Creek township | 0.25645508 | 0.36928456 | 40.0610958 | -85.84051371 |
| Wells County, Jackson township | 0.34534696 | 0.36874664 | 40.63333831 | -85.4286507 |
| Cass County, Jefferson township | 0.36780527 | 0.36820871 | 40.81356142 | -86.57358784 |
| Carroll County, Rock Creek township | 0.33015062 | 0.36820871 | 40.66027529 | -86.62392764 |
| Starke County, Railroad township | 0.34117805 | 0.36632598 | 41.2396432 | -86.87897289 |
| Dubois County, Marion township | 0.11982249 | 0.36525013 | 38.39330413 | -86.88549284 |
| Hancock County, Sugar Creek township | 0.17563206 | 0.36417429 | 39.77742621 | -85.93993335 |
| Montgomery County, Coal Creek township | 0.14376009 | 0.36363636 | 40.1772459 | -87.07416166 |
| Putnam County, Madison township | 0.14739107 | 0.36309844 | 39.64949081 | -86.9034197 |
| Jackson County, Hamilton township | 0.14846692 | 0.36282948 | 38.99161974 | -85.93179113 |
| Bartholomew County, Rock Creek township | 0.24354492 | 0.36229155 | 39.132575 | -85.73602531 |
| Miami County, Richland township | 0.33458849 | 0.35825713 | 40.90115719 | -85.96037877 |
| Jay County, Penn township | 0.31320602 | 0.3577192 | 40.56131848 | -85.10975191 |
| Bartholomew County, Wayne township | 0.2739376 | 0.35664336 | 39.14923996 | -85.9392121 |
| Hamilton County, Adams township | 0.30930608 | 0.35610543 | 40.10675375 | -86.13153743 |
| Jay County, Richland township | 0.29491662 | 0.35502959 | 40.36684644 | -85.19931032 |
| Pulaski County, White Post township | 0.3189887 | 0.35476062 | 41.03960192 | -86.90482714 |
| Hancock County, Center township | 0.19728349 | 0.3542227 | 39.78773571 | -85.79036636 |
| Wayne County, New Garden township | 0.15559441 | 0.3542227 | 39.91492229 | -84.91108082 |
| Clinton County, Johnson township | 0.31239914 | 0.35207101 | 40.32483626 | -86.33999057 |
| Clay County, Brazil township | 0.29034427 | 0.35180204 | 39.53098209 | -87.12476831 |
| Jay County, Knox township | 0.30123722 | 0.35180204 | 40.4111921 | -85.19257095 |
| Hamilton County, Jackson township | 0.24905863 | 0.35018827 | 40.11963742 | -86.01073862 |
| Jasper County, Gillam township | 0.31307154 | 0.35018827 | 41.04173202 | -86.93466785 |
| Clay County, Posey township | 0.18356643 | 0.34911243 | 39.5108477 | -87.15671975 |
| Gibson County, Union township | 0.1633943 | 0.34884346 | 38.28513734 | -87.56541748 |
| Decatur County, Fugit township | 0.20091447 | 0.3485745 | 39.3742108 | -85.40697149 |
| Vanderburgh County, Union township | 0.16003228 | 0.34696073 | 37.93162447 | -87.63474356 |
| Clinton County, Kirklin township | 0.27810651 | 0.34642281 | 40.19163502 | -86.35802573 |
| Allen County, Scipio township | 0.32611619 | 0.34480904 | 41.25539447 | -84.82050185 |
| Warren County, Washington township | 0.15115654 | 0.34427111 | 40.28265361 | -87.29926722 |
| Spencer County, Carter township | 0.19674556 | 0.34427111 | 38.19561229 | -86.9390158 |
| Wayne County, Jackson township | 0.24300699 | 0.34346423 | 39.84682013 | -85.15904571 |
| Hancock County, Jackson township | 0.11498117 | 0.34319527 | 39.82488683 | -85.69614292 |
| Randolph County, Wayne township | 0.22942442 | 0.34319527 | 40.19777825 | -84.80739897 |
| Benton County, Gilboa township | 0.30580958 | 0.34319527 | 40.67234218 | -87.10170249 |
| Harrison County, Posey township | 0.18477676 | 0.34238838 | 38.18373023 | -85.92687221 |
| Shelby County, Shelby township | 0.20172136 | 0.34185046 | 39.49700066 | -85.80209248 |
| Benton County, Center township | 0.27595481 | 0.33835395 | 40.57156032 | -87.2638722 |
| Newton County, Jefferson township | 0.22202797 | 0.33781603 | 40.76157198 | -87.42034555 |
| Madison County, Boone township | 0.29209252 | 0.3359333 | 40.31771904 | -85.68449885 |
| DeKalb County, Troy township | 0.30876815 | 0.3359333 | 41.45762953 | -84.81859571 |
| Parke County, Reserve township | 0.14766003 | 0.33512641 | 39.79381742 | -87.35851935 |
| Wayne County, Harrison township | 0.22054868 | 0.33512641 | 39.84427714 | -85.12975834 |
| Randolph County, Washington township | 0.25080689 | 0.3337816 | 40.05307325 | -84.94915671 |
| Decatur County, Adams township | 0.19607316 | 0.33324368 | 39.38408179 | -85.52341051 |
| Jasper County, Barkley township | 0.25806885 | 0.33216783 | 40.96336299 | -87.09511947 |
| Montgomery County, Walnut township | 0.10516407 | 0.33189887 | 40.02272368 | -86.79657041 |
| Crawford County, Whiskey Run township | 0.25766541 | 0.33082302 | 38.37143158 | -86.30763221 |
| Hamilton County, White River township | 0.22565896 | 0.3302851 | 40.11729285 | -85.98131066 |
| Benton County, Oak Grove township | 0.27192039 | 0.32920925 | 40.54728352 | -87.23696082 |
| Orange County, French Lick township | 0.22498655 | 0.32759548 | 38.55357129 | -86.60388061 |
| Benton County, Pine township | 0.2767617 | 0.32625067 | 40.5674487 | -87.20456703 |
| Benton County, Bolivar township | 0.2732652 | 0.32625067 | 40.49247986 | -87.09433578 |
| Crawford County, Sterling township | 0.285234 | 0.32571275 | 38.35827022 | -86.42550093 |
| Hendricks County, Liberty township | 0.25134481 | 0.32571275 | 39.6189711 | -86.49388512 |
| Jay County, Jefferson township | 0.25739645 | 0.32436794 | 40.37370522 | -85.01890662 |
| Dubois County, Harbison township | 0.1422808 | 0.32356105 | 38.44196823 | -86.93751934 |
| Pike County, Washington township | 0.1788596 | 0.32248521 | 38.48917897 | -87.28039233 |
| Orange County, Orleans township | 0.21705218 | 0.32167832 | 38.65123915 | -86.44528723 |
| Jay County, Wabash township | 0.20629371 | 0.32167832 | 40.5628476 | -84.86969811 |
| Warrick County, Boon township | 0.14174287 | 0.32140936 | 38.04292187 | -87.30445455 |
| Clark County, Carr township | 0.15599785 | 0.31925767 | 38.42056284 | -85.80699816 |
| Warren County, Warren township | 0.15371167 | 0.3189887 | 40.32300118 | -87.23473642 |
| Benton County, Grant township | 0.28348575 | 0.3189887 | 40.51301781 | -87.39058649 |
| Shelby County, Noble township | 0.14147391 | 0.31845078 | 39.43819336 | -85.63350251 |
| Dubois County, Jackson township | 0.1584185 | 0.31791286 | 38.30236011 | -86.86782012 |
| Lawrence County, Marion township | 0.22902098 | 0.31441635 | 38.72005114 | -86.46542594 |
| Boone County, Sugar Creek township | 0.18719742 | 0.31307154 | 40.16508534 | -86.59969433 |
| Sullivan County, Hamilton township | 0.12641205 | 0.31307154 | 39.16974897 | -87.40369069 |
| Henry County, Wayne township | 0.18235611 | 0.31118881 | 39.78810371 | -85.52363236 |
| Boone County, Marion township | 0.23816568 | 0.31011296 | 40.16034297 | -86.24319744 |
| Randolph County, Jackson township | 0.23977945 | 0.31011296 | 40.22255761 | -84.83334629 |
| Shelby County, Liberty township | 0.23050027 | 0.30903712 | 39.46036808 | -85.63031028 |
| Newton County, Grant township | 0.26627219 | 0.30903712 | 40.77569522 | -87.29867689 |
| Crawford County, Ohio township | 0.27218935 | 0.30903712 | 38.24731353 | -86.44000363 |
| Posey County, Black township | 0.12775686 | 0.30903712 | 37.92487385 | -87.86607909 |
| Benton County, Richland township | 0.2175901 | 0.30876815 | 40.69487858 | -87.42841425 |
| Tippecanoe County, Wayne township | 0.18571813 | 0.30823023 | 40.397312 | -87.01672409 |
| Clay County, Dick Johnson township | 0.137305 | 0.30769231 | 39.533059 | -87.15399235 |
| Spencer County, Clay township | 0.16796665 | 0.30742335 | 38.10891636 | -86.97863057 |
| Bartholomew County, Sand Creek township | 0.19755245 | 0.30661646 | 39.14451241 | -85.88114923 |
| Randolph County, Franklin township | 0.25013448 | 0.3063475 | 40.28503536 | -85.03258532 |
| Brown County, Washington township | 0.19150081 | 0.30580958 | 39.19006567 | -86.16852269 |
| Jasper County, Newton township | 0.21732114 | 0.30527165 | 40.92720211 | -87.21991291 |
| Daviess County, Madison township | 0.25416891 | 0.30446477 | 38.85009847 | -87.00346375 |
| Jasper County, Milroy township | 0.27192039 | 0.3041958 | 40.84382932 | -86.99005223 |
| Randolph County, Ward township | 0.2584723 | 0.30365788 | 40.27988236 | -84.97366275 |
| Franklin County, Posey township | 0.26613771 | 0.30338892 | 39.47254261 | -85.24493327 |
| Morgan County, Jefferson township | 0.20064551 | 0.302851 | 39.43693962 | -86.45897045 |
| Decatur County, Jackson township | 0.25013448 | 0.30258203 | 39.1450758 | -85.61677822 |
| Washington County, Gibson township | 0.23413125 | 0.29962345 | 38.69856717 | -85.91397235 |
| Floyd County, Greenville township | 0.17374933 | 0.29935449 | 38.34134382 | -85.93420095 |
| Putnam County, Cloverdale township | 0.16608392 | 0.29881657 | 39.50768549 | -86.80341616 |
| Owen County, Washington township | 0.17724583 | 0.29800968 | 39.28134492 | -86.77345161 |
| Newton County, Colfax township | 0.19109736 | 0.2972028 | 41.06674892 | -87.29289458 |
| Fountain County, Shawnee township | 0.11565358 | 0.29693384 | 40.25633875 | -87.24292865 |
| Bartholomew County, Clay township | 0.18639053 | 0.29666487 | 39.20867428 | -85.84277763 |
| Randolph County, Green township | 0.19244217 | 0.29585799 | 40.29525894 | -85.15045584 |
| Henry County, Fall Creek township | 0.12170522 | 0.29532006 | 40.06156939 | -85.572763 |
| Wayne County, Franklin township | 0.10623991 | 0.29478214 | 39.91232139 | -84.88177459 |
| Washington County, Franklin township | 0.19782141 | 0.29451318 | 38.63908208 | -86.00967604 |
| Perry County, Oil township | 0.21274879 | 0.29370629 | 38.2051654 | -86.4744689 |
| Johnson County, Union township | 0.12452932 | 0.29316837 | 39.50299293 | -86.15528731 |
| Clinton County, Sugar Creek township | 0.28321678 | 0.29263045 | 40.25141682 | -86.26053447 |
| Perry County, Troy township | 0.18908015 | 0.29236148 | 37.95859179 | -86.7661964 |
| Crawford County, Boone township | 0.26775148 | 0.29047875 | 38.18073827 | -86.44868424 |
| Greene County, Stockton township | 0.19257665 | 0.29020979 | 39.04029563 | -87.15555308 |
| Tipton County, Wildcat township | 0.25524476 | 0.29020979 | 40.40577386 | -85.9406673 |
| Steuben County, Richland township | 0.21987628 | 0.28859602 | 41.54898024 | -84.83440359 |
| Martin County, Perry township | 0.14443249 | 0.2880581 | 38.88601513 | -86.88211974 |
| Vigo County, Linton township | 0.17374933 | 0.2880581 | 39.32525219 | -87.38518388 |
| Morgan County, Ray township | 0.15761162 | 0.28698225 | 39.39922253 | -86.55229376 |
| Bartholomew County, Haw Creek township | 0.12103281 | 0.28671329 | 39.26321677 | -85.68803984 |
| Clay County, Van Buren township | 0.17105971 | 0.28644432 | 39.52889835 | -87.09554596 |
| Hancock County, Vernon township | 0.17832168 | 0.28536848 | 39.9105389 | -85.92122748 |
| Orange County, Southeast township | 0.24852071 | 0.28402367 | 38.46245431 | -86.32462612 |
| Posey County, Marrs township | 0.11040882 | 0.28267886 | 37.93747698 | -87.72056241 |
| Warrick County, Hart township | 0.17011834 | 0.28214094 | 38.19839972 | -87.28602149 |
| Henry County, Dudley township | 0.17993545 | 0.28187197 | 39.81007237 | -85.25357393 |
| Spencer County, Ohio township | 0.13219473 | 0.28160301 | 37.80224154 | -87.07374089 |
| Jackson County, Washington township | 0.21785906 | 0.28052717 | 38.90054077 | -85.91516478 |
| Washington County, Posey township | 0.2169177 | 0.2802582 | 38.46018408 | -86.29585361 |
| Warren County, Pine township | 0.21880043 | 0.27998924 | 40.44431699 | -87.36904964 |
| Vermillion County, Helt township | 0.11498117 | 0.27998924 | 39.79583988 | -87.38785676 |
| Fountain County, Van Buren township | 0.14443249 | 0.27945132 | 40.12301975 | -87.25926739 |
| Benton County, Parish Grove township | 0.18370091 | 0.27891339 | 40.57970038 | -87.38250298 |
| Hendricks County, Middle township | 0.22243141 | 0.27622378 | 39.86314616 | -86.46149429 |
| Newton County, Beaver township | 0.21073158 | 0.27622378 | 40.94345666 | -87.45838237 |
| Randolph County, Greensfork township | 0.14806347 | 0.27407208 | 40.05048106 | -84.91979264 |
| Delaware County, Perry township | 0.13501883 | 0.27272727 | 40.13049423 | -85.32444906 |
| Bartholomew County, Clifty township | 0.16460463 | 0.27138246 | 39.24104176 | -85.69120575 |
| Benton County, Union township | 0.22337278 | 0.27057558 | 40.65840858 | -87.2232334 |
| Monroe County, Clear Creek township | 0.08364712 | 0.26976869 | 39.06185481 | -86.53767331 |
| Lawrence County, Indian Creek township | 0.1514255 | 0.26923077 | 38.85988759 | -86.53475332 |
| Washington County, Pierce township | 0.20104895 | 0.26654115 | 38.55501895 | -86.07943092 |
| Sullivan County, Curry township | 0.1873319 | 0.26654115 | 39.19196355 | -87.40105172 |
| Shelby County, Hanover township | 0.19889726 | 0.26627219 | 39.66722184 | -85.68918392 |
| Parke County, Wabash township | 0.13394298 | 0.2654653 | 39.77159957 | -87.36119351 |
| Parke County, Penn township | 0.1514255 | 0.26519634 | 39.80787442 | -87.23847451 |
| Harrison County, Blue River township | 0.19419042 | 0.26492738 | 38.36687647 | -86.2501614 |
| Vigo County, Fayette township | 0.14456697 | 0.26412049 | 39.53123472 | -87.44896241 |
| Newton County, McClellan township | 0.17011834 | 0.26385153 | 41.03240142 | -87.4476206 |
| Washington County, Madison township | 0.23399677 | 0.26304465 | 38.48009436 | -86.26412515 |
| Hendricks County, Union township | 0.1422808 | 0.26277569 | 39.89202273 | -86.54658874 |
| Morgan County, Adams township | 0.13313609 | 0.26250672 | 39.56342351 | -86.64876992 |
| Morgan County, Madison township | 0.21866595 | 0.26250672 | 39.57873897 | -86.26302826 |
| Clay County, Jackson township | 0.18881119 | 0.26062399 | 39.50668835 | -87.09829116 |
| Johnson County, Blue River township | 0.16366326 | 0.26008607 | 39.35826835 | -86.02783396 |
| Jay County, Noble township | 0.21274879 | 0.26008607 | 40.47418354 | -84.8835986 |
| Rush County, Richland township | 0.14416353 | 0.25981711 | 39.49973381 | -85.29992155 |
| Boone County, Clinton township | 0.22202797 | 0.25981711 | 40.16943552 | -86.36102506 |
| Wayne County, Green township | 0.13582571 | 0.2584723 | 39.94742046 | -85.02492619 |
| Montgomery County, Wayne township | 0.15008069 | 0.2563206 | 40.08629073 | -87.05580965 |
| Orange County, Jackson township | 0.18934911 | 0.25389995 | 38.48917292 | -86.67003367 |
| Newton County, Washington township | 0.1274879 | 0.25389995 | 40.89698024 | -87.43395704 |
| Harrison County, Webster township | 0.14201183 | 0.25363098 | 38.15340114 | -86.10480704 |
| Greene County, Grant township | 0.16420118 | 0.25363098 | 39.03821936 | -87.12652704 |
| Montgomery County, Madison township | 0.16325982 | 0.25228618 | 40.16456544 | -86.89727973 |
| Johnson County, Nineveh township | 0.17186659 | 0.25147929 | 39.36060139 | -86.05695492 |
| Washington County, Howard township | 0.21516945 | 0.25094137 | 38.53746956 | -86.14003824 |
| Knox County, Johnson township | 0.10247445 | 0.25013448 | 38.63860279 | -87.49542102 |
| Knox County, Palmyra township | 0.13528779 | 0.24986552 | 38.68103471 | -87.46134214 |
| Perry County, Clark township | 0.12103281 | 0.24959656 | 38.22280635 | -86.70394277 |
| Daviess County, Elmore township | 0.15330823 | 0.24905863 | 38.85220345 | -87.03241137 |
| Rush County, Union township | 0.13864981 | 0.24798279 | 39.64272384 | -85.39694617 |
| Warrick County, Anderson township | 0.15734266 | 0.24663798 | 37.93389428 | -87.34616978 |
| Hendricks County, Marion township | 0.10919849 | 0.24636902 | 39.76102898 | -86.59343324 |
| Benton County, Hickory Grove township | 0.20979021 | 0.24583109 | 40.49880944 | -87.51178858 |
| Posey County, Harmony township | 0.14281872 | 0.24529317 | 38.13058667 | -87.92998888 |
| Washington County, Jackson township | 0.18854223 | 0.24529317 | 38.42192124 | -86.09750388 |
| Warrick County, Campbell township | 0.10798817 | 0.24502421 | 38.09945969 | -87.47115647 |
| Gibson County, White River township | 0.1035503 | 0.24475524 | 38.39819547 | -87.58139441 |
| Posey County, Robb township | 0.13824637 | 0.24421732 | 38.18770753 | -87.7791003 |
| Greene County, Fairplay township | 0.12600861 | 0.24341044 | 39.03194986 | -87.03945887 |
| Crawford County, Johnson township | 0.16299086 | 0.24260355 | 38.28062441 | -86.58063486 |
| Shelby County, Moral township | 0.11511565 | 0.24206563 | 39.61267081 | -85.84478735 |
| Daviess County, Veale township | 0.13986014 | 0.2415277 | 38.59198914 | -87.15170841 |
| Bartholomew County, Ohio township | 0.1457773 | 0.24125874 | 39.15394041 | -85.99728251 |
| Gibson County, Center township | 0.154922 | 0.24098978 | 38.34381754 | -87.44268776 |
| Putnam County, Washington township | 0.14819796 | 0.24045186 | 39.47819021 | -87.01343115 |
| Martin County, Center township | 0.16433566 | 0.24018289 | 38.67977287 | -86.82050334 |
| Crawford County, Patoka township | 0.19284562 | 0.23991393 | 38.32501387 | -86.57492142 |
| Warren County, Prairie township | 0.18975256 | 0.23910705 | 40.46856411 | -87.3959671 |
| Hendricks County, Eel River township | 0.13811189 | 0.2380312 | 39.92084199 | -86.63175153 |
| Dubois County, Boone township | 0.11390533 | 0.23587951 | 38.4440886 | -86.96630389 |
| Dubois County, Columbia township | 0.14053254 | 0.23534158 | 38.51355314 | -86.69600068 |
| Wayne County, Washington township | 0.14631522 | 0.23507262 | 39.8024892 | -85.16575288 |
| Daviess County, Barr township | 0.15895643 | 0.23480366 | 38.65444702 | -87.08587291 |
| Bartholomew County, Harrison township | 0.13717052 | 0.23426573 | 39.22048721 | -85.98807117 |
| Madison County, Jackson township | 0.14174287 | 0.23399677 | 40.12287297 | -85.77219142 |
| Steuben County, York township | 0.17912856 | 0.23345885 | 41.68732434 | -84.87321563 |
| Dubois County, Ferdinand township | 0.16299086 | 0.23292093 | 38.23361414 | -86.84741935 |
| Bartholomew County, Camp Atterbury UT | 0.14846692 | 0.23184508 | 39.33841663 | -86.06001209 |
| Harrison County, Spencer township | 0.20629371 | 0.23184508 | 38.25136562 | -86.20755266 |
| Greene County, Jefferson township | 0.13609467 | 0.23184508 | 39.1165556 | -86.9703604 |
| Morgan County, Clay township | 0.12775686 | 0.23130715 | 39.54342302 | -86.38584219 |
| Putnam County, Marion township | 0.15330823 | 0.23130715 | 39.64090756 | -86.78637456 |
| Hendricks County, Franklin township | 0.10597095 | 0.23130715 | 39.63221495 | -86.66935746 |
| Vermillion County, Clinton township | 0.1253362 | 0.23103819 | 39.6645428 | -87.43310849 |
| Warren County, Medina township | 0.18087682 | 0.23076923 | 40.47025957 | -87.09711846 |
| Orange County, Stampers Creek township | 0.16258741 | 0.22996235 | 38.48690712 | -86.35046855 |
| Washington County, Monroe township | 0.14792899 | 0.22996235 | 38.68578261 | -86.03243904 |
| Orange County, Paoli township | 0.17509414 | 0.22996235 | 38.54028055 | -86.45981797 |
| Putnam County, Monroe township | 0.16299086 | 0.22915546 | 39.69176757 | -86.86851486 |
| Wayne County, Perry township | 0.15976331 | 0.22915546 | 39.94998801 | -85.05425189 |
| Morgan County, Harrison township | 0.15761162 | 0.22727273 | 39.55654773 | -86.26603133 |
| Clay County, Lewis township | 0.1035503 | 0.22727273 | 39.17768805 | -87.19743036 |
| Henry County, Liberty township | 0.08015062 | 0.22619688 | 39.92844639 | -85.32489744 |
| Parke County, Florida township | 0.13017751 | 0.22565896 | 39.66051242 | -87.3745396 |
| Fountain County, Troy township | 0.10166756 | 0.22565896 | 40.12710479 | -87.31820787 |
| Sullivan County, Haddon township | 0.08176439 | 0.22538999 | 38.9678069 | -87.39836337 |
| Randolph County, Monroe township | 0.15263583 | 0.22485207 | 40.17931958 | -85.10847711 |
| Washington County, Vernon township | 0.19741797 | 0.22404519 | 38.56884262 | -86.25227629 |
| Knox County, Busseron township | 0.11767079 | 0.22377622 | 38.88096275 | -87.43785092 |
| Morgan County, Baker township | 0.15613233 | 0.22377622 | 39.3503901 | -86.49983881 |
| Brown County, Van Buren township | 0.15519096 | 0.2232383 | 39.16557294 | -86.14249126 |
| Fayette County, Orange township | 0.15922539 | 0.2232383 | 39.54155548 | -85.26415555 |
| Knox County, Vigo township | 0.13999462 | 0.2232383 | 38.90704402 | -87.17177896 |
| Warren County, Adams township | 0.18585261 | 0.22296934 | 40.45838833 | -87.24791196 |
| Knox County, Decker township | 0.07557827 | 0.22270038 | 38.52951446 | -87.53719117 |
| Greene County, Cass township | 0.15236686 | 0.22189349 | 38.91459891 | -86.96623844 |
| Montgomery County, Ripley township | 0.11780527 | 0.22135557 | 40.01333283 | -86.97588739 |
| Perry County, Union township | 0.17025282 | 0.22081764 | 38.02984499 | -86.52609596 |
| Fountain County, Richland township | 0.16352878 | 0.22081764 | 40.20571106 | -87.15987476 |
| Greene County, Washington township | 0.1225121 | 0.2197418 | 38.98963307 | -87.07394784 |
| Washington County, Polk township | 0.18558365 | 0.21732114 | 38.52348877 | -85.96728842 |
| Rush County, Orange township | 0.13367402 | 0.21705218 | 39.4579354 | -85.60116321 |
| Martin County, Lost River township | 0.14308768 | 0.21705218 | 38.53575045 | -86.69317071 |
| Greene County, Richland township | 0.09709521 | 0.21651425 | 39.02561934 | -86.9524058 |
| Vermillion County, Eugene township | 0.09548144 | 0.21651425 | 39.95741335 | -87.45736549 |
| Morgan County, Jackson township | 0.14497041 | 0.21543841 | 39.37674228 | -86.26085386 |
| Jackson County, Brownstown township | 0.13824637 | 0.21490048 | 38.92743299 | -85.96996991 |
| Harrison County, Morgan township | 0.15895643 | 0.21436256 | 38.39973802 | -86.10050966 |
| Daviess County, Van Buren township | 0.16944594 | 0.21436256 | 38.80357858 | -86.98002898 |
| Posey County, Point township | 0.07369554 | 0.2140936 | 37.88611325 | -87.95676288 |
| Harrison County, Jackson township | 0.16541151 | 0.2140936 | 38.26894837 | -86.14719745 |
| Boone County, Jefferson township | 0.11915008 | 0.2140936 | 40.04964907 | -86.55545422 |
| Washington County, Jefferson township | 0.12116729 | 0.21382464 | 38.65300335 | -86.18270707 |
| Lawrence County, Bono township | 0.15075309 | 0.2119419 | 38.71331875 | -86.3787962 |
| Montgomery County, Sugar Creek township | 0.13932222 | 0.2119419 | 40.19605261 | -86.71468499 |
| Harrison County, Washington township | 0.1697149 | 0.21086606 | 38.14043401 | -86.22236279 |
| Hendricks County, Clay township | 0.12318451 | 0.21086606 | 39.71442137 | -86.56996718 |
| Posey County, Bethel township | 0.13636364 | 0.21005917 | 38.22325743 | -87.97765863 |
| Dubois County, Madison township | 0.11726735 | 0.20979021 | 38.40179391 | -87.00055864 |
| Lawrence County, Marshall township | 0.14591178 | 0.20925229 | 38.95088102 | -86.55214519 |
| Orange County, Greenfield township | 0.18504572 | 0.20898332 | 38.40712819 | -86.47719682 |
| Hancock County, Green township | 0.10059172 | 0.20898332 | 39.94300881 | -85.76820962 |
| Warren County, Mound township | 0.18060785 | 0.20898332 | 40.13519219 | -87.4361089 |
| Shelby County, Sugar Creek township | 0.13206025 | 0.20871436 | 39.59286476 | -85.87712923 |
| Shelby County, Hendricks township | 0.12802582 | 0.20817644 | 39.52394638 | -85.85730806 |
| Parke County, Jackson township | 0.15747714 | 0.20817644 | 39.66424828 | -87.10831511 |
| Knox County, Steen township | 0.13017751 | 0.20790748 | 38.64465002 | -87.26179557 |
| Benton County, York township | 0.13757396 | 0.20790748 | 40.67465448 | -87.4608025 |
| Shelby County, Van Buren township | 0.11753631 | 0.20736955 | 39.62768117 | -85.75398424 |
| Bartholomew County, Flat Rock township | 0.11861216 | 0.20629371 | 39.27996032 | -85.89158195 |
| Hancock County, Brown township | 0.10112964 | 0.20548682 | 39.88900558 | -85.65724298 |
| Henry County, Greensboro township | 0.06777838 | 0.20548682 | 39.87187432 | -85.45203222 |
| Spencer County, Harrison township | 0.14174287 | 0.20333513 | 38.16915016 | -86.88440427 |
| Monroe County, Bean Blossom township | 0.1584185 | 0.20306616 | 39.27264146 | -86.65701652 |
| Fountain County, Cain township | 0.10597095 | 0.2027972 | 40.11890716 | -87.20033364 |
| Boone County, Jackson township | 0.11888112 | 0.20199032 | 39.94304507 | -86.62884625 |
| Rush County, Center township | 0.06777838 | 0.20172136 | 39.78564584 | -85.4943562 |
| Newton County, Iroquois township | 0.18074233 | 0.20145239 | 40.82422514 | -87.35273347 |
| Newton County, Jackson township | 0.17993545 | 0.20091447 | 40.93741349 | -87.36894352 |
| Jackson County, Driftwood township | 0.16191501 | 0.20064551 | 38.84804355 | -86.0978797 |
| Parke County, Greene township | 0.1155191 | 0.20064551 | 39.79960371 | -87.12113229 |
| Lawrence County, Spice Valley township | 0.09467456 | 0.19983862 | 38.8199189 | -86.59837526 |
| Boone County, Harrison township | 0.12762238 | 0.19903174 | 39.97637609 | -86.47608246 |
| Monroe County, Benton township | 0.11888112 | 0.19795589 | 39.21276137 | -86.45918429 |
| Putnam County, Russell township | 0.09965035 | 0.19688004 | 39.85785454 | -86.99545638 |
| Jay County, Madison township | 0.13394298 | 0.19688004 | 40.38552299 | -84.89746457 |
| Jackson County, Grassy Fork township | 0.15922539 | 0.19607316 | 38.76510336 | -85.90474252 |
| Lawrence County, Pleasant Run township | 0.11350188 | 0.1958042 | 38.91977963 | -86.43918954 |
| Brown County, Hamblen township | 0.14308768 | 0.19526627 | 39.23674044 | -86.19156092 |
| Wayne County, Clay township | 0.12547068 | 0.19526627 | 39.92782428 | -85.05764366 |
| Greene County, Center township | 0.12721893 | 0.19499731 | 39.03063855 | -86.71749729 |
| Clark County, Wood township | 0.10892953 | 0.19445939 | 38.49895583 | -85.94156123 |
| Vigo County, Pierson township | 0.09521248 | 0.19445939 | 39.29898623 | -87.32955843 |
| Fayette County, Fairview township | 0.14523938 | 0.19284562 | 39.65238548 | -85.24758405 |
| Morgan County, Green township | 0.14631522 | 0.19230769 | 39.51444166 | -86.30122273 |
| Greene County, Stafford township | 0.11511565 | 0.19096288 | 38.99380338 | -87.13195983 |
| Brown County, Jackson township | 0.12426036 | 0.19096288 | 39.24814628 | -86.33696613 |
| Shelby County, Jackson township | 0.10812265 | 0.19042496 | 39.35122847 | -85.94048236 |
| Martin County, Halbert township | 0.14739107 | 0.18988704 | 38.67545723 | -86.7627588 |
| Pike County, Patoka township | 0.14147391 | 0.18908015 | 38.35182262 | -87.2387526 |
| Dubois County, Cass township | 0.09709521 | 0.18854223 | 38.26222125 | -86.93077776 |
| Tippecanoe County, Jackson township | 0.14483593 | 0.18665949 | 40.28832765 | -87.06025232 |
| Fountain County, Davis township | 0.16594944 | 0.18639053 | 40.29250609 | -87.11931763 |
| Gibson County, Barton township | 0.11161915 | 0.18585261 | 38.20247094 | -87.34342121 |
| Greene County, Highland township | 0.0809575 | 0.18423884 | 39.11443602 | -86.94130924 |
| Spencer County, Grass township | 0.08700914 | 0.18396988 | 38.00000137 | -87.0208646 |
| Johnson County, Hensley township | 0.15169446 | 0.18343195 | 39.396646 | -86.22871384 |
| Rush County, Ripley township | 0.05177515 | 0.18316299 | 39.76592916 | -85.52687198 |
| Gibson County, Columbia township | 0.11807423 | 0.18289403 | 38.33574103 | -87.32766263 |
| Gibson County, Montgomery township | 0.06186122 | 0.18181818 | 38.38769355 | -87.75663315 |
| Owen County, Jennings township | 0.12735342 | 0.18181818 | 39.46542956 | -86.83827019 |
| Dubois County, Hall township | 0.09359871 | 0.18181818 | 38.35599117 | -86.68701899 |
| Owen County, Marion township | 0.14039806 | 0.18181818 | 39.31852012 | -86.97450535 |
| Owen County, Jackson township | 0.15290479 | 0.18154922 | 39.46757339 | -86.86745938 |
| Randolph County, Union township | 0.14160839 | 0.18074233 | 40.06080876 | -85.03726159 |
| Daviess County, Bogard township | 0.13407746 | 0.18047337 | 38.80988964 | -87.06681943 |
| Pike County, Monroe township | 0.16648736 | 0.17993545 | 38.242821 | -87.28074058 |
| Harrison County, Franklin township | 0.12856374 | 0.1788596 | 38.21529779 | -86.03847924 |
| Randolph County, Stoney Creek township | 0.10785368 | 0.17616998 | 40.15715186 | -85.11186295 |
| Warrick County, Skelton township | 0.12076385 | 0.17590102 | 38.07913405 | -87.18458718 |
| Dubois County, Jefferson township | 0.112695 | 0.17563206 | 38.31159661 | -86.69266707 |
| Parke County, Raccoon township | 0.1451049 | 0.1753631 | 39.68854123 | -87.13485352 |
| Clay County, Washington township | 0.1366326 | 0.17482517 | 39.36293226 | -86.96895158 |
| Wayne County, Dalton township | 0.08162991 | 0.17455621 | 39.97981595 | -85.13887447 |
| Rush County, Noble township | 0.12372243 | 0.17455621 | 39.59589816 | -85.37429315 |
| Bartholomew County, Jackson township | 0.13044648 | 0.17428725 | 39.06987926 | -86.06754587 |
| Orange County, Northwest township | 0.10462614 | 0.17401829 | 38.60015587 | -86.65583199 |
| Vigo County, Prairieton township | 0.11323292 | 0.17348037 | 39.35349643 | -87.47001017 |
| Knox County, Washington township | 0.10032275 | 0.1732114 | 38.74967728 | -87.48244567 |
| Orange County, Orangeville township | 0.11471221 | 0.17240452 | 38.68011857 | -86.52896405 |
| Perry County, Leopold township | 0.15169446 | 0.17213556 | 38.07645087 | -86.54899772 |
| Monroe County, Indian Creek township | 0.08526089 | 0.17052179 | 39.07066955 | -86.65375642 |
| Shelby County, Union township | 0.10946746 | 0.16998386 | 39.53174217 | -85.67907487 |
| Daviess County, Harrison township | 0.06118881 | 0.16944594 | 38.58991209 | -87.12286033 |
| Pike County, Lockhart township | 0.08754707 | 0.16890802 | 38.23666017 | -87.1946014 |
| Warren County, Steuben township | 0.09978483 | 0.16863905 | 40.24631047 | -87.42276987 |
| Putnam County, Jefferson township | 0.10220549 | 0.16863905 | 39.54993226 | -86.76852134 |
| Washington County, Brown township | 0.13273265 | 0.16837009 | 38.6598731 | -86.26924715 |
| Warrick County, Greer township | 0.10260893 | 0.16810113 | 38.18228387 | -87.37473573 |
| Lawrence County, Guthrie township | 0.10677784 | 0.16756321 | 38.78214185 | -86.39888934 |
| Parke County, Union township | 0.13932222 | 0.16594944 | 39.70867129 | -87.10281718 |
| Spencer County, Luce township | 0.06939215 | 0.16487359 | 37.90351704 | -87.23443868 |
| Knox County, Harrison township | 0.07611619 | 0.16379774 | 38.62855165 | -87.35105658 |
| Vermillion County, Vermillion township | 0.06468531 | 0.16379774 | 39.86451378 | -87.40921737 |
| Warrick County, Lane township | 0.1161915 | 0.16352878 | 38.19430167 | -87.22862815 |
| Jackson County, Carr township | 0.12560516 | 0.16352878 | 38.81060122 | -86.19063577 |
| Harrison County, Heth township | 0.13004303 | 0.16245293 | 38.11365374 | -86.16805139 |
| Sullivan County, Gill township | 0.08055406 | 0.16164605 | 39.02421618 | -87.56725814 |
| Morgan County, Gregg township | 0.09400215 | 0.16110812 | 39.55015124 | -86.47346856 |
| Henry County, Prairie township | 0.09023669 | 0.16110812 | 39.9777797 | -85.37697161 |
| Harrison County, Boone township | 0.12358795 | 0.16083916 | 37.96895884 | -86.04297299 |
| Parke County, Liberty township | 0.08270576 | 0.1605702 | 39.88065841 | -87.31843444 |
| Montgomery County, Brown township | 0.10328133 | 0.1605702 | 39.88636881 | -87.08075751 |
| Clay County, Harrison township | 0.11955352 | 0.16003228 | 39.28459317 | -87.12565066 |
| Putnam County, Clinton township | 0.10731576 | 0.15895643 | 39.71824225 | -86.92427491 |
| Jackson County, Owen township | 0.13125336 | 0.1584185 | 38.87485478 | -86.15271265 |
| Perry County, Tobin township | 0.1105433 | 0.15788058 | 38.00765079 | -86.5289568 |
| Wayne County, Jefferson township | 0.10381926 | 0.15788058 | 39.88860694 | -85.12302507 |
| Rush County, Anderson township | 0.12506724 | 0.15761162 | 39.47030765 | -85.48135628 |
| Pike County, Logan township | 0.09696073 | 0.1570737 | 38.42458612 | -87.31712508 |
| Warrick County, Owen township | 0.10973642 | 0.1570737 | 38.1720921 | -87.23128339 |
| Clay County, Cass township | 0.15330823 | 0.15680473 | 39.45175744 | -86.95782357 |
| Parke County, Howard township | 0.13152232 | 0.15653577 | 39.93288351 | -87.10461081 |
| Jackson County, Salt Creek township | 0.13636364 | 0.15599785 | 39.03705539 | -86.2185918 |
| Shelby County, Washington township | 0.13017751 | 0.154922 | 39.36392578 | -85.82091052 |
| Gibson County, Wabash township | 0.08552985 | 0.15357719 | 38.24548235 | -87.97522715 |
| Sullivan County, Turman township | 0.10328133 | 0.15277031 | 39.10911415 | -87.49878267 |
| Morgan County, Ashland township | 0.09803658 | 0.15277031 | 39.4990093 | -86.68661273 |
| Sullivan County, Jefferson township | 0.05863367 | 0.15250134 | 39.00411052 | -87.27701871 |
| Owen County, Wayne township | 0.10866057 | 0.15223238 | 39.34360679 | -86.70666364 |
| Sullivan County, Cass township | 0.10220549 | 0.1514255 | 39.09499897 | -87.29537417 |
| Henry County, Jefferson township | 0.0739645 | 0.1514255 | 39.98026516 | -85.40632209 |
| Owen County, Morgan township | 0.10825713 | 0.15088757 | 39.36081222 | -86.93980159 |
| Rush County, Walker township | 0.07450242 | 0.15034965 | 39.60825865 | -85.52028854 |
| Perry County, Anderson township | 0.1105433 | 0.14954276 | 38.11181612 | -86.71799915 |
| Pike County, Madison township | 0.11820871 | 0.1492738 | 38.49325272 | -87.33801957 |
| Daviess County, Steele township | 0.1218397 | 0.14900484 | 38.749516 | -87.16172203 |
| Martin County, Mitcheltree township | 0.08122647 | 0.14873588 | 38.74205401 | -86.75430581 |
| Lawrence County, Perry township | 0.08741259 | 0.14873588 | 38.90870062 | -86.58687845 |
| Greene County, Beech Creek township | 0.09682625 | 0.14766003 | 39.141633 | -86.70326226 |
| Harrison County, Taylor township | 0.12977407 | 0.14685315 | 38.07283217 | -85.9420913 |
| Vigo County, Prairie Creek township | 0.08741259 | 0.14604626 | 39.33127954 | -87.47263387 |
| Fountain County, Jackson township | 0.09965035 | 0.1457773 | 39.95717997 | -87.13124725 |
| Warren County, Jordan township | 0.11202259 | 0.1457773 | 40.3614496 | -87.46853176 |
| Owen County, Clay township | 0.11027434 | 0.14550834 | 39.23260707 | -86.72093311 |
| Fayette County, Posey township | 0.08364712 | 0.14523938 | 39.78790468 | -85.25689773 |
| Jackson County, Pershing township | 0.12829478 | 0.14416353 | 38.96590128 | -86.16963033 |
| Steuben County, Clear Lake township | 0.11767079 | 0.14416353 | 41.70689848 | -84.83962552 |
| Putnam County, Floyd township | 0.0929263 | 0.14335664 | 39.70100471 | -86.68998431 |
| Monroe County, Washington township | 0.112695 | 0.14281872 | 39.26382924 | -86.54060957 |
| Posey County, Smith township | 0.0626681 | 0.14093599 | 38.16355659 | -87.75289106 |
| Vigo County, Nevins township | 0.09601937 | 0.14066703 | 39.58777054 | -87.29476697 |
| Spencer County, Huff township | 0.12264658 | 0.1401291 | 38.05813884 | -86.89817858 |
| Sullivan County, Jackson township | 0.09467456 | 0.13770845 | 39.18791882 | -87.34286602 |
| Parke County, Sugar Creek township | 0.10785368 | 0.13770845 | 39.89261064 | -87.16886441 |
| Spencer County, Jackson township | 0.11363636 | 0.13743948 | 38.11313131 | -87.0359433 |
| Orange County, Northeast township | 0.09763314 | 0.1366326 | 38.66668223 | -86.35580344 |
| Fountain County, Millcreek township | 0.10247445 | 0.1344809 | 39.96338656 | -87.21944538 |
| Spencer County, Hammond township | 0.08781603 | 0.13367402 | 38.01586181 | -86.9322931 |
| Greene County, Jackson township | 0.09400215 | 0.13313609 | 38.98624092 | -86.72317915 |
| Monroe County, Salt Creek township | 0.11161915 | 0.13286713 | 39.16389213 | -86.40692676 |
| Montgomery County, Clark township | 0.09090909 | 0.13286713 | 39.8808115 | -86.69625858 |
| Warren County, Kent township | 0.12345347 | 0.13232921 | 40.20786151 | -87.46027581 |
| Owen County, Taylor township | 0.08257127 | 0.13179129 | 39.46112146 | -86.77989702 |
| Warrick County, Pigeon township | 0.08606778 | 0.13125336 | 38.1374348 | -87.06189864 |
| Knox County, Widner township | 0.10758472 | 0.12937063 | 38.82642167 | -87.29833334 |
| Owen County, Franklin township | 0.0739645 | 0.12910167 | 39.23910002 | -86.80821431 |
| Vanderburgh County, Armstrong township | 0.08848843 | 0.12802582 | 38.10937126 | -87.61449979 |
| Warren County, Pike township | 0.08821947 | 0.12802582 | 40.22207227 | -87.39592692 |
| Clay County, Sugar Ridge township | 0.10059172 | 0.12775686 | 39.34913557 | -87.08831084 |
| Owen County, Harrison township | 0.09588488 | 0.1274879 | 39.42803286 | -86.63687862 |
| Pike County, Marion township | 0.04558903 | 0.12668101 | 38.3699107 | -87.17856521 |
| Hancock County, Brandywine township | 0.06952663 | 0.12614309 | 39.74337313 | -85.79667931 |
| Sullivan County, Fairbanks township | 0.08270576 | 0.12506724 | 39.17376525 | -87.46186518 |
| Putnam County, Franklin township | 0.08795051 | 0.12237762 | 39.78273334 | -86.88652963 |
| Hancock County, Blue River township | 0.07530931 | 0.119688 | 39.71157766 | -85.68281146 |
| Montgomery County, Scott township | 0.08499193 | 0.11753631 | 39.94032686 | -86.89612752 |
| Rush County, Posey township | 0.07288865 | 0.11726735 | 39.63288413 | -85.54626791 |
| Fountain County, Fulton township | 0.06455083 | 0.11672942 | 39.97762805 | -87.42529977 |
| Monroe County, Polk township | 0.07235073 | 0.11646046 | 39.02397824 | -86.33755547 |
| Putnam County, Jackson township | 0.09279182 | 0.11457773 | 39.79417087 | -86.73709345 |
| Daviess County, Reeve township | 0.07275417 | 0.11350188 | 38.55510509 | -86.95256027 |
| Martin County, Rutherford township | 0.10328133 | 0.11323292 | 38.54657035 | -86.8372591 |
| Pike County, Jefferson township | 0.05890264 | 0.1105433 | 38.48301785 | -87.19396352 |
| Owen County, Montgomery township | 0.07087144 | 0.10919849 | 39.3457813 | -86.73579981 |
| Henry County, Stoney Creek township | 0.07638515 | 0.10866057 | 40.05392555 | -85.21687653 |
| Owen County, Jefferson township | 0.08041958 | 0.10785368 | 39.25401233 | -87.0119307 |
| Henry County, Blue River township | 0.07719204 | 0.10785368 | 39.97028224 | -85.28893242 |
| Posey County, Robinson township | 0.05217859 | 0.10731576 | 38.04857837 | -87.70804985 |
| Henry County, Harrison township | 0.05728886 | 0.10650888 | 39.94086577 | -85.47156033 |
| Owen County, Lafayette township | 0.07436794 | 0.1048951 | 39.31427588 | -86.91624248 |
| Rush County, Washington township | 0.06750941 | 0.1048951 | 39.70424839 | -85.32864118 |
| Gibson County, Washington township | 0.04975793 | 0.10166756 | 38.41645099 | -87.52125379 |
| Posey County, Center township | 0.06508876 | 0.1013986 | 38.12488113 | -87.84392842 |
| Clay County, Perry township | 0.06750941 | 0.09924691 | 39.35329942 | -87.14661171 |
| Fountain County, Wabash township | 0.06804734 | 0.09359871 | 40.08873611 | -87.41197331 |
| Greene County, Smith township | 0.0683163 | 0.09090909 | 39.08263545 | -87.1210876 |
| Posey County, Lynn township | 0.04733728 | 0.07611619 | 38.01567289 | -87.88489161 |
| Pike County, Clay township | 0.04626143 | 0.07073696 | 38.47508671 | -87.3982688 |
