## Supplementary material for "Mapping long-distance jump dispersal risk by leveraging transport infrastructure data and sampling bias correction: a case study with the invasive spotted lanternfly (*Lycorma delicatula*)": S5 Kentucky risk

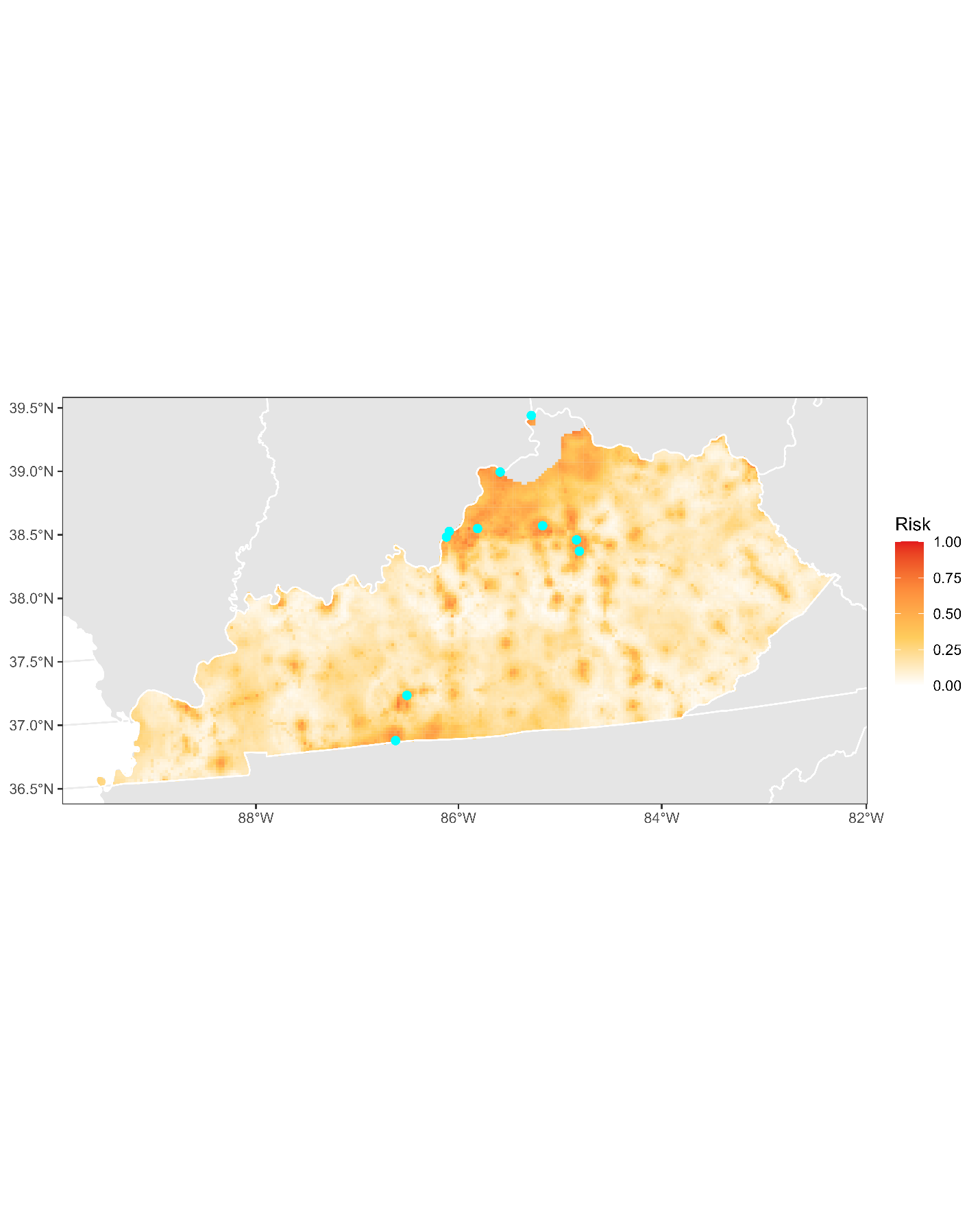

**Figure S5.1:** Projected risk of colonization by *L. delicatula* jump dispersal for Kentucky. Risk projections are based on an ensemble random forest model including distances from existing *L. delicatula* populations. Note that areas falling less than 20 km from existing *L. delicatula* populations are not included in the map because any colonization there would be categorized as diffusive spread based on our methodology (regardless of the actual source of the resulting population). Darker, redder colors indicate increased risk of colonization by jump dispersal. Cyan points show the top ten highest risk locations in the state.

**Table S5.1:** Ranked list of Kentucky county subdivisions sorted by the maximum projected risk of colonization by jump dispersal occurring there. For each subdivision, we identified the 2.5 x 2.5 km grid cell centroid included there with the highest projected risk of colonization by jump dispersal and report that location and risk value.

| **County, subdivision** | **Median risk value** | **Maximum risk value** | **Latitude of maximum risk** | **Longitude of maximum risk** |
| --- | --- | --- | --- | --- |
| Jefferson County, Louisville West CCD | 0.62210866 | 0.8211404 | 38.26294253 | -85.79993435 |
| Simpson County, Franklin CCD | 0.41541151 | 0.79451318 | 36.66464081 | -86.55703335 |
| Fayette County, Lexington-Fayette Northwest CCD | 0.33189887 | 0.75363098 | 38.08222899 | -84.52176917 |
| Warren County, Bowling Green CCD | 0.48144164 | 0.75201721 | 37.01093686 | -86.39913082 |
| Jefferson County, North Dixie CCD | 0.49354492 | 0.74798279 | 38.22097494 | -85.83477312 |
| Carroll County, Prestonville CCD | 0.61807423 | 0.73776224 | 38.68424523 | -85.21547391 |
| Jefferson County, East Jefferson CCD | 0.48466918 | 0.73749328 | 38.26088301 | -85.51006541 |
| Franklin County, Frankfort CCD | 0.50981711 | 0.73614847 | 38.22475558 | -84.84783734 |
| Fayette County, Lexington-Fayette Southeast CCD | 0.20414201 | 0.73614847 | 37.9909712 | -84.5070907 |
| Boone County, Hebron CCD | 0.59332975 | 0.72996235 | 39.0990915 | -84.83051603 |
| Woodford County, Midway CCD | 0.43289403 | 0.71893491 | 38.17259374 | -84.76875009 |
| Campbell County, Alexandria CCD | 0.68746638 | 0.71597633 | 38.92626304 | -84.41739521 |
| Jefferson County, Central Jefferson CCD | 0.57450242 | 0.70118343 | 38.15979839 | -85.64044617 |
| Scott County, Georgetown CCD | 0.39133943 | 0.67859064 | 38.26473638 | -84.55123547 |
| Boyd County, Ashland CCD | 0.40572889 | 0.67778375 | 38.47396803 | -82.62185172 |
| McCracken County, Paducah CCD | 0.37600861 | 0.67428725 | 37.07738307 | -88.61365807 |
| Jefferson County, Louisville East CCD | 0.52918236 | 0.67374933 | 38.25574432 | -85.71390678 |
| Trimble County, Milton CCD | 0.56535772 | 0.67374933 | 38.72392651 | -85.41406737 |
| Shelby County, Shelbyville CCD | 0.48493814 | 0.67267348 | 38.21381759 | -85.22674082 |
| Fayette County, Lexington-Fayette Central CCD | 0.60099516 | 0.67132867 | 38.05739671 | -84.49668026 |
| Greenup County, Flatwoods-Russell CCD | 0.37076385 | 0.66487359 | 38.50229898 | -82.67509662 |
| Trimble County, Bedford CCD | 0.50161377 | 0.6605702 | 38.6424443 | -85.25084835 |
| Simpson County, East Simpson CCD | 0.38784293 | 0.65653577 | 36.70683248 | -86.52332186 |
| Jefferson County, South Jefferson CCD | 0.41581495 | 0.65438408 | 38.13276108 | -85.58633329 |
| Jefferson County, Southeast Jefferson CCD | 0.56051641 | 0.6514255 | 38.22144327 | -85.57372259 |
| Fayette County, Lexington-Fayette Southwest CCD | 0.39281872 | 0.64739107 | 38.06277001 | -84.55379188 |
| Carroll County, Carrollton CCD | 0.64631522 | 0.64631522 | 38.68171617 | -85.18665054 |
| Fayette County, Lexington-Fayette Northeast CCD | 0.32006455 | 0.64308768 | 38.0741367 | -84.43608577 |
| Woodford County, Versailles CCD | 0.29074771 | 0.64281872 | 38.15307051 | -84.80074669 |
| Hardin County, Elizabethtown CCD | 0.22740721 | 0.64093599 | 37.68631971 | -85.87964892 |
| Boone County, Burlington CCD | 0.59951587 | 0.6401291 | 39.03524665 | -84.86972086 |
| Oldham County, Pewee Valley CCD | 0.48386229 | 0.62022593 | 38.3692605 | -85.46544443 |
| Jessamine County, Keene CCD | 0.34534696 | 0.62022593 | 37.94937265 | -84.54253314 |
| Christian County, Hopkinsville CCD | 0.13313609 | 0.6183432 | 36.8674595 | -87.46972768 |
| Franklin County, Bald Knob CCD | 0.46342119 | 0.61807423 | 38.25471306 | -84.93038974 |
| Henry County, Eminence CCD | 0.50901022 | 0.60946746 | 38.44549328 | -85.30895388 |
| Clark County, Winchester CCD | 0.18343195 | 0.60785368 | 38.00523527 | -84.18628173 |
| Henry County, Campbellsburg CCD | 0.46772458 | 0.60328133 | 38.54875477 | -85.20637732 |
| Oldham County, La Grange CCD | 0.49045186 | 0.60059172 | 38.38895627 | -85.43353118 |
| Jefferson County, Northeast Jefferson CCD | 0.48117267 | 0.59951587 | 38.33229329 | -85.55791639 |
| Campbell County, Grants Lick CCD | 0.48628295 | 0.59870898 | 38.90140938 | -84.39205121 |
| Scott County, Sadieville CCD | 0.37789134 | 0.59736417 | 38.2841943 | -84.51912405 |
| Jefferson County, Pond Creek CCD | 0.3781603 | 0.59521248 | 38.16945133 | -85.75499421 |
| Greenup County, South Shore CCD | 0.40048413 | 0.59413663 | 38.73503887 | -82.95479077 |
| Kenton County, Piner-Visalia CCD | 0.46799354 | 0.59386767 | 38.82634258 | -84.55047263 |
| Todd County, Guthrie CCD | 0.23897257 | 0.59332975 | 36.64302445 | -87.15518808 |
| Calloway County, Murray CCD | 0.43571813 | 0.59090909 | 36.61131395 | -88.31995976 |
| Jefferson County, Louisville South CCD | 0.52918236 | 0.59010221 | 38.19879796 | -85.83784992 |
| Grant County, Williamstown-Dry Ridge CCD | 0.43571813 | 0.58821947 | 38.696157 | -84.60020204 |
| Allen County, Scottsville CCD | 0.45400753 | 0.58445401 | 36.69959297 | -86.1551627 |
| Daviess County, Whitesville-Philpot CCD | 0.09400215 | 0.58418505 | 37.78003402 | -87.07642856 |
| Jefferson County, Louisville Central CCD | 0.57261969 | 0.57719204 | 38.25815042 | -85.74258077 |
| Whitley County, Williamsburg CCD | 0.12721893 | 0.57557827 | 36.721308 | -84.16168133 |
| Allen County, Adolphus CCD | 0.42980097 | 0.57530931 | 36.67970111 | -86.18616761 |
| Henderson County, Henderson CCD | 0.47592792 | 0.57315761 | 37.8165991 | -87.5902268 |
| Boyle County, Danville CCD | 0.23184508 | 0.57154384 | 37.63573579 | -84.79288123 |
| Scott County, Stamping Ground CCD | 0.27756859 | 0.56939215 | 38.31170025 | -84.57295237 |
| Whitley County, Corbin CCD | 0.26869285 | 0.5683163 | 36.93992452 | -84.09836103 |
| Shelby County, Simpsonville CCD | 0.41958042 | 0.56697149 | 38.22385649 | -85.34131792 |
| Nelson County, Bardstown CCD | 0.2105971 | 0.56670253 | 37.78795688 | -85.49090863 |
| Hopkins County, Madisonville CCD | 0.19526627 | 0.56347499 | 37.34009423 | -87.50121919 |
| Madison County, Richmond CCD | 0.34077461 | 0.56347499 | 37.74512605 | -84.28574295 |
| Jefferson County, Louisville Airport CCD | 0.51102743 | 0.56105433 | 38.12029081 | -85.70394603 |
| Christian County, LaFayette CCD | 0.10866057 | 0.55433029 | 36.70989018 | -87.45931864 |
| Warren County, Woodburn CCD | 0.2534965 | 0.55298548 | 36.90665564 | -86.49794563 |
| Pendleton County, Falmouth CCD | 0.50927918 | 0.54545455 | 38.60272262 | -84.32290116 |
| Pendleton County, Goforth-Locust Grove CCD | 0.47310382 | 0.54518558 | 38.69399975 | -84.33747244 |
| Warren County, Sand Hill-Girkin CCD | 0.20804196 | 0.54491662 | 37.03088034 | -86.36804525 |
| Bullitt County, Shepherdsville Southeast CCD | 0.23466918 | 0.54115116 | 37.98483439 | -85.69402187 |
| Henry County, Pleasureville CCD | 0.47485207 | 0.54061323 | 38.36807025 | -84.94229458 |
| Bullitt County, Shepherdsville Northwest CCD | 0.27192039 | 0.54034427 | 38.05376866 | -85.71327638 |
| Allen County, Holland CCD | 0.38743948 | 0.53550296 | 36.67739749 | -86.15808249 |
| Henry County, New Castle CCD | 0.44701452 | 0.535234 | 38.48731067 | -85.27370769 |
| Logan County, Adairville CCD | 0.34319527 | 0.53335126 | 36.66638206 | -86.86884807 |
| Adair County, Columbia CCD | 0.29491662 | 0.5330823 | 37.10054191 | -85.33092008 |
| Pendleton County, Butler CCD | 0.447149 | 0.53281334 | 38.73553734 | -84.3015518 |
| Mercer County, Harrodsburg CCD | 0.11229155 | 0.52824099 | 37.77649972 | -84.85802062 |
| Harrison County, Berry CCD | 0.31993007 | 0.52555137 | 38.5584504 | -84.33000039 |
| Taylor County, Campbellsville CCD | 0.2345347 | 0.5239376 | 37.32723061 | -85.35561676 |
| Pulaski County, Somerset CCD | 0.24004841 | 0.52286175 | 37.05814991 | -84.62271508 |
| Lyon County, Eddyville CCD | 0.27151694 | 0.52178591 | 37.10624044 | -88.01248789 |
| Montgomery County, Mount Sterling CCD | 0.26815492 | 0.52151694 | 38.06857568 | -83.9150941 |
| Mason County, Maysville CCD | 0.26923077 | 0.51963421 | 38.6409137 | -83.79073064 |
| Daviess County, Owensboro CCD | 0.44392146 | 0.51855836 | 37.78420355 | -87.13349198 |
| Shelby County, Christianburg CCD | 0.45642819 | 0.51855836 | 38.33149665 | -85.0350273 |
| Boyd County, Catlettsburg CCD | 0.21732114 | 0.51775148 | 38.42666983 | -82.60134104 |
| Shelby County, Harrisonville CCD | 0.32436794 | 0.51640667 | 38.15424525 | -85.0615539 |
| Caldwell County, Princeton CCD | 0.20225928 | 0.51210328 | 37.12092507 | -87.8968913 |
| Warren County, Smiths Grove CCD | 0.21880043 | 0.50806885 | 37.04170404 | -86.22403656 |
| Anderson County, Lawrenceburg CCD | 0.29599247 | 0.50779989 | 38.07489383 | -84.92856343 |
| Jessamine County, Nicholasville CCD | 0.21490048 | 0.50645508 | 37.88561907 | -84.58137648 |
| Knox County, Barbourville CCD | 0.12856374 | 0.50268962 | 36.87315589 | -83.88073215 |
| Knox County, Corbin CCD | 0.32356105 | 0.50080689 | 36.95927079 | -84.06670091 |
| Bracken County, Foster CCD | 0.42630446 | 0.49946208 | 38.77155701 | -84.20794119 |
| Washington County, Willisburg CCD | 0.14766003 | 0.49677246 | 37.87128206 | -85.16139487 |
| Barren County, Glasgow CCD | 0.2528241 | 0.4965035 | 37.01846359 | -85.94189205 |
| Laurel County, London CCD | 0.42442173 | 0.49220011 | 37.14194014 | -84.09461024 |
| Owen County, Owenton CCD | 0.38300161 | 0.4887036 | 38.58177726 | -84.82233839 |
| Owen County, New Liberty CCD | 0.43706294 | 0.48789672 | 38.58034493 | -85.05589514 |
| Christian County, Oak Grove-Pembroke CCD | 0.17065627 | 0.48655191 | 36.68364774 | -87.40560512 |
| Carter County, Grayson CCD | 0.18074233 | 0.48574502 | 38.35027664 | -82.93614907 |
| Daviess County, Utica-Stanley CCD | 0.08902636 | 0.48493814 | 37.76406877 | -87.16468669 |
| Owen County, Monterey-New Columbus CCD | 0.31334051 | 0.48413125 | 38.38502987 | -84.88154574 |
| Bell County, Middlesborough CCD | 0.15895643 | 0.48359333 | 36.60684435 | -83.69635658 |
| Logan County, Russellville CCD | 0.26492738 | 0.48251748 | 36.82613818 | -86.90617095 |
| Madison County, Berea CCD | 0.16541151 | 0.48090371 | 37.56802455 | -84.31384945 |
| Jefferson County, Floyds Fork CCD | 0.26331361 | 0.48063475 | 38.2313155 | -85.42727118 |
| Grant County, Corinth CCD | 0.35395374 | 0.47660032 | 38.49152078 | -84.57391168 |
| Boyd County, Cannonsburg CCD | 0.10758472 | 0.47498655 | 38.36978497 | -82.6993188 |
| Hardin County, Radcliff CCD | 0.13636364 | 0.46853147 | 37.82649182 | -85.94689759 |
| Grant County, Crittenden CCD | 0.43652501 | 0.46718666 | 38.78205679 | -84.55745552 |
| Rowan County, Morehead CCD | 0.15814954 | 0.46369016 | 38.17723804 | -83.43226022 |
| Christian County, Fort Campbell CCD | 0.25511027 | 0.45911781 | 36.6412101 | -87.43880932 |
| Bracken County, Brooksville CCD | 0.31038193 | 0.45884884 | 38.6359994 | -84.20070137 |
| Madison County, Waco-Bybee CCD | 0.15545992 | 0.45831092 | 37.72024404 | -84.26084754 |
| Pike County, Millard CCD | 0.14133943 | 0.45669715 | 37.42357452 | -82.49531396 |
| Henderson County, Spottsville CCD | 0.10381926 | 0.45077999 | 37.95187846 | -87.60360663 |
| Warren County, Alvaton CCD | 0.22162453 | 0.44513179 | 36.91989243 | -86.3823759 |
| Hart County, Munfordville CCD | 0.23507262 | 0.44136633 | 37.28468308 | -85.90580649 |
| Bracken County, Augusta CCD | 0.34521248 | 0.43437332 | 38.72713202 | -83.98109498 |
| Franklin County, Peaks Mill CCD | 0.28321678 | 0.4316837 | 38.29381839 | -84.8663599 |
| Jefferson County, South Dixie CCD | 0.28644432 | 0.42980097 | 38.09028482 | -85.88181931 |
| Graves County, Hickory-Lowes CCD | 0.15438408 | 0.42630446 | 36.81044149 | -88.64001313 |
| Laurel County, North Corbin CCD | 0.30392684 | 0.42630446 | 36.96206188 | -84.09482072 |
| Graves County, Mayfield CCD | 0.19822485 | 0.4260355 | 36.76594734 | -88.64438741 |
| Hardin County, Fort Knox CCD | 0.11847768 | 0.42334588 | 37.91756899 | -85.96332122 |
| Muhlenberg County, Central City CCD | 0.22565896 | 0.42146315 | 37.29142068 | -87.13515171 |
| Pike County, Pikeville CCD | 0.26600323 | 0.41850457 | 37.49613391 | -82.53970303 |
| Laurel County, East Bernstadt CCD | 0.10086068 | 0.41823561 | 37.16686062 | -84.11925715 |
| Boyd County, Ashland South CCD | 0.26304465 | 0.4168908 | 38.43604261 | -82.68721865 |
| Jessamine County, Wilmore CCD | 0.30123722 | 0.4168908 | 37.86614392 | -84.61329827 |
| Bourbon County, Paris CCD | 0.14954276 | 0.41554599 | 38.19332817 | -84.27207677 |
| Spencer County, West Spencer CCD | 0.16702528 | 0.41500807 | 38.04404445 | -85.33848984 |
| Fulton County, Fulton CCD | 0.07961269 | 0.41285637 | 36.51250262 | -88.89524414 |
| Monroe County, Fountain Run CCD | 0.3218128 | 0.4112426 | 36.68327931 | -85.958558 |
| Mason County, Mays Lick CCD | 0.22754169 | 0.4112426 | 38.51103867 | -83.84168316 |
| Madison County, Boonesborough-White Hall CCD | 0.20185584 | 0.41070468 | 37.83914997 | -84.32856146 |
| Cumberland County, Burkesville CCD | 0.17724583 | 0.41016676 | 36.79012881 | -85.37527231 |
| Greenup County, Greenup CCD | 0.11296396 | 0.40855299 | 38.73198342 | -82.9260339 |
| Bullitt County, Lebanon Junction CCD | 0.16003228 | 0.40828402 | 37.82961244 | -85.71574749 |
| Clark County, Ford-Boonesborough CCD | 0.15115654 | 0.40693921 | 37.98586383 | -84.21835933 |
| McCracken County, St. Johns-Freemont CCD | 0.18329747 | 0.40640129 | 37.00717634 | -88.56371527 |
| Warren County, Hadley-Browning CCD | 0.16998386 | 0.40640129 | 36.89330809 | -86.61347432 |
| Lawrence County, Louisa CCD | 0.21503497 | 0.40505648 | 38.11123891 | -82.60097148 |
| Perry County, Hazard CCD | 0.20225928 | 0.40102205 | 37.27644823 | -83.21184796 |
| Whitley County, Saxton CCD | 0.23184508 | 0.40048413 | 36.71853217 | -84.13364923 |
| Allen County, Settle CCD | 0.18908015 | 0.39967725 | 36.78605815 | -86.11534174 |
| Garrard County, Lancaster CCD | 0.11215707 | 0.39913932 | 37.61452798 | -84.56573078 |
| Clay County, Manchester CCD | 0.23641743 | 0.39887036 | 37.13014121 | -83.75285002 |
| Anderson County, Northwest Anderson CCD | 0.19123185 | 0.39887036 | 38.07748314 | -84.95714043 |
| Perry County, Viper CCD | 0.10758472 | 0.39752555 | 37.22623705 | -83.1631073 |
| Madison County, Kirksville CCD | 0.12990855 | 0.38864981 | 37.70358969 | -84.32119199 |
| Hart County, Horse Cave CCD | 0.24583109 | 0.38811189 | 37.1761215 | -85.94913803 |
| Laurel County, Bush CCD | 0.14900484 | 0.38757396 | 37.09208337 | -84.04536578 |
| Henderson County, Corydon CCD | 0.07853685 | 0.38757396 | 37.92677729 | -87.89468728 |
| Calloway County, Shiloh CCD | 0.19876278 | 0.38407746 | 36.62993981 | -88.26143061 |
| Grayson County, Caneyville CCD | 0.14994621 | 0.38353954 | 37.42168074 | -86.48922326 |
| Rowan County, Hilda-Triplett CCD | 0.13555675 | 0.37977407 | 38.18603138 | -83.51794891 |
| Logan County, Auburn CCD | 0.18988704 | 0.37977407 | 36.89550507 | -86.64165408 |
| Harrison County, Berry West CCD | 0.29854761 | 0.37923615 | 38.54176322 | -84.39102 |
| Todd County, Trenton CCD | 0.20871436 | 0.37278107 | 36.64715917 | -87.21139213 |
| Spencer County, Taylorsville CCD | 0.19150081 | 0.37170522 | 38.04154405 | -85.30991261 |
| Crittenden County, Marion CCD | 0.25336202 | 0.37116729 | 37.33415255 | -88.0737271 |
| Lincoln County, Stanford CCD | 0.17415277 | 0.36982249 | 37.53127955 | -84.63615064 |
| Allen County, Halfway CCD | 0.26452394 | 0.3690156 | 36.71332013 | -86.32374842 |
| Harrison County, Cynthiana CCD | 0.23722431 | 0.36874664 | 38.39802863 | -84.29741804 |
| Lewis County, Tollesboro CCD | 0.19230769 | 0.36874664 | 38.55149791 | -83.57210331 |
| Estill County, Winston-West Irvine CCD | 0.11027434 | 0.36067778 | 37.69242904 | -83.97680366 |
| Hopkins County, Dawson Springs CCD | 0.15572889 | 0.35960194 | 37.19636296 | -87.68890465 |
| Hart County, Hardyville CCD | 0.18881119 | 0.35879505 | 37.26013181 | -85.88052739 |
| Laurel County, Colony CCD | 0.17025282 | 0.35610543 | 37.15027351 | -84.17918558 |
| Union County, Morganfield CCD | 0.12479828 | 0.35529855 | 37.67131689 | -88.09499948 |
| Ohio County, Beaver Dam CCD | 0.22001076 | 0.35502959 | 37.37918533 | -86.809719 |
| Marshall County, Benton CCD | 0.09924691 | 0.35395374 | 36.85957151 | -88.35124956 |
| Muhlenberg County, Greenville CCD | 0.21530393 | 0.35314685 | 37.22685637 | -87.17141997 |
| Wayne County, Monticello CCD | 0.28604088 | 0.34776762 | 36.83289909 | -84.85652835 |
| Bullitt County, Mount Washington CCD | 0.2802582 | 0.34722969 | 38.02433755 | -85.63063336 |
| Wolfe County, Campton CCD | 0.19257665 | 0.34696073 | 37.74084051 | -83.56457527 |
| Powell County, Stanton CCD | 0.11229155 | 0.34669177 | 37.7995492 | -83.69918941 |
| Laurel County, Keavy CCD | 0.11484669 | 0.34642281 | 36.96484641 | -84.12294263 |
| Floyd County, Risner CCD | 0.12641205 | 0.34588488 | 37.63156952 | -82.74601782 |
| Russell County, Jamestown CCD | 0.2175901 | 0.34454008 | 36.98656357 | -85.06188739 |
| Livingston County, Smithland CCD | 0.1316568 | 0.34346423 | 37.04991406 | -88.53098948 |
| Pulaski County, Burnside CCD | 0.11780527 | 0.3429263 | 36.98902865 | -84.60472378 |
| Butler County, Welchs Creek CCD | 0.21247983 | 0.34023669 | 37.25732477 | -86.68184487 |
| Mercer County, Salvisa-McAfee CCD | 0.15626681 | 0.34023669 | 37.93417565 | -84.86302621 |
| Clinton County, Albany CCD | 0.26331361 | 0.3394298 | 36.67878234 | -85.13533495 |
| Harlan County, Harlan CCD | 0.10758472 | 0.3394298 | 36.86228911 | -83.34009746 |
| Lyon County, Southeast Lyon CCD | 0.2345347 | 0.33754707 | 37.08213267 | -87.98658281 |
| Monroe County, Mount Hermon CCD | 0.24179666 | 0.33647122 | 36.75056804 | -85.69368546 |
| Barren County, Cave City CCD | 0.19661108 | 0.3359333 | 37.13410075 | -85.98338542 |
| Nicholas County, Headquarters CCD | 0.20575578 | 0.33566434 | 38.37589777 | -84.06805546 |
| Laurel County, Greenmount-Langnau CCD | 0.11081226 | 0.33512641 | 37.16128373 | -84.06286342 |
| Johnson County, Oil Springs CCD | 0.11430877 | 0.33297472 | 37.77339502 | -82.80724461 |
| Floyd County, Martin-Allen CCD | 0.11013986 | 0.33189887 | 37.56529812 | -82.75793422 |
| Whitley County, Siler CCD | 0.12291555 | 0.3302851 | 36.73788983 | -84.10209162 |
| Barren County, Tracy CCD | 0.23924153 | 0.32947821 | 36.79658215 | -85.97176758 |
| Johnson County, Paintsville CCD | 0.15250134 | 0.32920925 | 37.7703051 | -82.77886769 |
| Wayne County, Mill Springs CCD | 0.1612426 | 0.3267886 | 36.86286319 | -84.9375501 |
| Calloway County, Hazel-Lynn Grove CCD | 0.17711135 | 0.32651963 | 36.61490376 | -88.37619974 |
| Cumberland County, Kettle-Peytonsburg CCD | 0.22350726 | 0.32571275 | 36.78763603 | -85.3471718 |
| Calloway County, Kirksey CCD | 0.15290479 | 0.32329209 | 36.63714781 | -88.37393825 |
| Hancock County, Hawesville CCD | 0.10812265 | 0.32302313 | 37.93639181 | -86.76898228 |
| Monroe County, Tompkinsville CCD | 0.25416891 | 0.32302313 | 36.70620085 | -85.69980008 |
| Marshall County, Calvert City CCD | 0.14900484 | 0.32302313 | 36.97967236 | -88.48114265 |
| Rockcastle County, Conway CCD | 0.11309844 | 0.32302313 | 37.4130483 | -84.33833279 |
| Marion County, Lebanon CCD | 0.15303927 | 0.32275417 | 37.56605804 | -85.26369605 |
| Breathitt County, Taulbee CCD | 0.11941904 | 0.32248521 | 37.61270451 | -83.41314507 |
| Mercer County, Burgin CCD | 0.08364712 | 0.32221625 | 37.85196466 | -84.70216268 |
| Hardin County, Cecilia CCD | 0.0648198 | 0.3211404 | 37.6688669 | -85.93959277 |
| Estill County, Ravenna CCD | 0.08176439 | 0.31979559 | 37.68961115 | -83.94841106 |
| Logan County, Lewisburg CCD | 0.2345347 | 0.31764389 | 36.94144828 | -86.94904558 |
| McCracken County, Grahamville-Heath CCD | 0.10368478 | 0.31710597 | 37.13045353 | -88.75085254 |
| Warren County, Greencastle CCD | 0.14712211 | 0.31683701 | 37.08202009 | -86.44702897 |
| Rockcastle County, Mount Vernon CCD | 0.08068854 | 0.31683701 | 37.39090782 | -84.34182208 |
| Caldwell County, Fredonia CCD | 0.20656267 | 0.31629909 | 37.1284722 | -88.01010317 |
| Barren County, Hiseville CCD | 0.18867671 | 0.31629909 | 37.06283627 | -85.93589565 |
| Floyd County, Prestonsburg CCD | 0.14685315 | 0.31603012 | 37.67884864 | -82.7664007 |
| Clay County, Sibert-Hima CCD | 0.14792899 | 0.31441635 | 37.13300518 | -83.78102232 |
| Metcalfe County, Summer Shade CCD | 0.14497041 | 0.31280258 | 36.99456439 | -85.65992541 |
| Wayne County, Zula-Powersburg CCD | 0.23157612 | 0.31172673 | 36.81594503 | -84.91602468 |
| Todd County, Elkton CCD | 0.15747714 | 0.31118881 | 36.82284199 | -87.16232235 |
| Montgomery County, Jeffersonville CCD | 0.20925229 | 0.30957504 | 37.97156961 | -83.8441727 |
| Ballard County, Wickliffe CCD | 0.15061861 | 0.30661646 | 36.9707399 | -89.07903658 |
| Butler County, Morgantown CCD | 0.20790748 | 0.30500269 | 37.21509935 | -86.71571552 |
| Floyd County, Betsy Layne-Stanville CCD | 0.24018289 | 0.30446477 | 37.60326603 | -82.6933711 |
| McCreary County, Stearns-Whitley City CCD | 0.11767079 | 0.30311996 | 36.72926018 | -84.47358968 |
| Robertson County, Mount Olivet CCD | 0.1958042 | 0.30123722 | 38.49212055 | -84.10735271 |
| Marshall County, Fairdealing CCD | 0.13071544 | 0.29962345 | 36.78197438 | -88.18896897 |
| Powell County, Clay City CCD | 0.14201183 | 0.29935449 | 37.84449628 | -83.92301879 |
| Clinton County, Cumberland City CCD | 0.22243141 | 0.29774072 | 36.72312069 | -85.12888903 |
| Muhlenberg County, Drakesboro CCD | 0.23318989 | 0.29558903 | 37.21852509 | -87.05814861 |
| Casey County, Liberty CCD | 0.20925229 | 0.29397526 | 37.311243 | -84.92797069 |
| Estill County, Irvine CCD | 0.11040882 | 0.29370629 | 37.71455697 | -83.97319104 |
| Lyon County, Land Between the Lakes CCD | 0.21597633 | 0.29343733 | 37.04512801 | -88.10445701 |
| Carlisle County, Bardwell CCD | 0.24475524 | 0.29289941 | 36.92623375 | -89.08315922 |
| Lincoln County, Waynesburg CCD | 0.21288327 | 0.29236148 | 37.3762376 | -84.65995021 |
| Magoffin County, Salyersville CCD | 0.10852609 | 0.29182356 | 37.73762931 | -83.10277835 |
| Boyle County, Junction City CCD | 0.11242604 | 0.29047875 | 37.59143192 | -84.79960722 |
| Rockcastle County, Livingston CCD | 0.08902636 | 0.28967187 | 37.34115644 | -84.29224742 |
| Pulaski County, Nancy CCD | 0.14093599 | 0.28967187 | 37.06346614 | -84.67907166 |
| Leslie County, Hyden CCD | 0.15317375 | 0.2894029 | 37.15865502 | -83.37552674 |
| Nicholas County, Carlisle CCD | 0.18961807 | 0.28671329 | 38.35097353 | -84.04301998 |
| Taylor County, Mannsville CCD | 0.1570737 | 0.28644432 | 37.41089837 | -85.28619759 |
| Russell County, Russell Springs CCD | 0.21032813 | 0.28536848 | 37.055616 | -85.0803008 |
| Webster County, Providence CCD | 0.16083916 | 0.28536848 | 37.40220583 | -87.75169798 |
| Adair County, Cane Valley CCD | 0.21866595 | 0.28483055 | 37.12520893 | -85.35596394 |
| Wayne County, Coopersville CCD | 0.19566971 | 0.28321678 | 36.76641699 | -84.86644995 |
| Lawrence County, Blaine CCD | 0.24515869 | 0.28321678 | 38.02564613 | -82.8489057 |
| Breathitt County, Jackson CCD | 0.20763852 | 0.28267886 | 37.56553378 | -83.39236867 |
| Adair County, Glens Fork CCD | 0.1880043 | 0.28214094 | 37.07084558 | -85.24948332 |
| Fulton County, Kentucky Bend CCD | 0.26465842 | 0.2802582 | 36.52683447 | -89.54366169 |
| Nelson County, Coxs Creek CCD | 0.13851533 | 0.27891339 | 37.85692531 | -85.50992231 |
| Fleming County, Flemingsburg CCD | 0.18558365 | 0.27864443 | 38.41112765 | -83.74176706 |
| Letcher County, Jenkins CCD | 0.19822485 | 0.27729962 | 37.17421542 | -82.68430603 |
| Graves County, Farmington CCD | 0.13609467 | 0.2767617 | 36.69400244 | -88.56646695 |
| Fleming County, Hillsboro CCD | 0.15277031 | 0.27568585 | 38.35825777 | -83.66327083 |
| Calloway County, New Concord CCD | 0.16931146 | 0.27568585 | 36.54277966 | -88.29869255 |
| Adair County, Gradyville CCD | 0.18316299 | 0.27487897 | 37.08335591 | -85.3905211 |
| Webster County, Sebree CCD | 0.1070468 | 0.27434104 | 37.60870899 | -87.49905431 |
| Meade County, Battletown-Payneville CCD | 0.16164605 | 0.27272727 | 38.17399433 | -86.36269106 |
| Hopkins County, Nebo CCD | 0.15949435 | 0.27272727 | 37.37998152 | -87.75416869 |
| Crittenden County, West Crittenden CCD | 0.20091447 | 0.27245831 | 37.43774799 | -88.2916236 |
| Letcher County, Blackey CCD | 0.12305003 | 0.27245831 | 37.13874358 | -82.97752601 |
| Butler County, Rochester CCD | 0.19190425 | 0.27138246 | 37.2058296 | -86.88828944 |
| Larue County, Magnolia-Upton CCD | 0.09709521 | 0.2711135 | 37.46214207 | -85.8816058 |
| Grayson County, Leitchfield CCD | 0.1697149 | 0.27084454 | 37.47249099 | -86.28190503 |
| Trigg County, Land Between the Lakes CCD | 0.19795589 | 0.27003765 | 36.77644875 | -88.10443682 |
| Washington County, Springfield CCD | 0.13004303 | 0.26949973 | 37.69652078 | -85.21589308 |
| Letcher County, Whitesburg CCD | 0.1873319 | 0.267617 | 37.12342351 | -82.83683609 |
| Bath County, Owingsville CCD | 0.13528779 | 0.267617 | 38.10430345 | -83.82214322 |
| Crittenden County, South Crittenden CCD | 0.1880043 | 0.26734804 | 37.29154393 | -88.10683982 |
| Bell County, Pineville CCD | 0.0866057 | 0.26707907 | 36.79250039 | -83.75127555 |
| Bath County, Sharpsburg CCD | 0.19419042 | 0.26681011 | 38.20415113 | -83.92175159 |
| Union County, Sturgis CCD | 0.18195266 | 0.26627219 | 37.55083177 | -87.96452206 |
| Russell County, Eli CCD | 0.17953201 | 0.26573427 | 37.07264922 | -85.02065093 |
| Hopkins County, Hanson CCD | 0.0957504 | 0.26573427 | 37.40675172 | -87.49357981 |
| Elliott County, Isonville CCD | 0.18154922 | 0.26492738 | 38.05999556 | -82.95889863 |
| Lewis County, Garrison CCD | 0.12399139 | 0.26465842 | 38.60153065 | -83.18348751 |
| Breckinridge County, McDaniels-Hudson CCD | 0.12721893 | 0.26465842 | 37.79844323 | -86.15247348 |
| Casey County, Dunnville CCD | 0.1781872 | 0.26412049 | 37.20045144 | -84.9444968 |
| Pulaski County, Science Hill CCD | 0.16137708 | 0.26412049 | 37.21586912 | -84.62717465 |
| Carlisle County, Arlington CCD | 0.17670791 | 0.26385153 | 36.78781977 | -89.0108902 |
| McCracken County, Camelia CCD | 0.11538462 | 0.26358257 | 36.96960858 | -88.68118322 |
| Adair County, White Oak CCD | 0.20804196 | 0.26358257 | 37.06579537 | -85.19308149 |
| Ohio County, Centertown CCD | 0.14107047 | 0.26331361 | 37.3358443 | -87.12984511 |
| Lee County, Beattyville CCD | 0.13259817 | 0.26304465 | 37.57547202 | -83.70773756 |
| Meade County, Brandenburg CCD | 0.12143626 | 0.26277569 | 37.97823637 | -86.15727606 |
| Metcalfe County, Edmonton CCD | 0.17684239 | 0.26223776 | 36.99213826 | -85.63173871 |
| Carlisle County, Cunningham CCD | 0.17011834 | 0.2619688 | 36.86854204 | -88.86141492 |
| Clay County, Big Creek CCD | 0.17025282 | 0.26143088 | 37.14363273 | -83.66466966 |
| Boyle County, Perryville CCD | 0.0718128 | 0.26116192 | 37.61881817 | -84.85303631 |
| Grayson County, Millerstown CCD | 0.1958042 | 0.25981711 | 37.37226906 | -86.15179017 |
| Knox County, Gray CCD | 0.16245293 | 0.25874126 | 36.95366895 | -84.0104676 |
| Livingston County, Salem CCD | 0.16514255 | 0.2584723 | 37.3728459 | -88.32695675 |
| Bourbon County, Millersburg CCD | 0.10919849 | 0.25793437 | 38.23484697 | -84.23636384 |
| Ohio County, McHenry-Rockport CCD | 0.19338354 | 0.25739645 | 37.38561747 | -86.89483255 |
| Martin County, Threeforks CCD | 0.06535772 | 0.25739645 | 37.76412027 | -82.31694604 |
| Graves County, Symsonia CCD | 0.1105433 | 0.25712749 | 36.76421864 | -88.61620382 |
| Estill County, Wagersville-South Irvine CCD | 0.10543303 | 0.2563206 | 37.67030088 | -83.98041412 |
| Carter County, Willard CCD | 0.14497041 | 0.2563206 | 38.28093755 | -82.9194305 |
| Ballard County, Barlow CCD | 0.18854223 | 0.25551372 | 37.02008842 | -89.15977813 |
| Lewis County, Vanceburg CCD | 0.16487359 | 0.25389995 | 38.59433549 | -83.33094451 |
| Meade County, Flaherty CCD | 0.1309844 | 0.25201721 | 37.83118892 | -86.00392933 |
| Ohio County, Horse Branch CCD | 0.18074233 | 0.24959656 | 37.38834228 | -86.63673324 |
| Breckinridge County, Hardinsburg CCD | 0.14402905 | 0.24878967 | 37.77679658 | -86.4434157 |
| Lawrence County, Adams CCD | 0.15505648 | 0.24852071 | 38.0481164 | -82.64157152 |
| Edmonson County, Mammoth Cave CCD | 0.15788058 | 0.24825175 | 37.30339284 | -86.13230606 |
| Carter County, Carter CCD | 0.16648736 | 0.24798279 | 38.45386798 | -83.06353202 |
| Fleming County, Ewing CCD | 0.16137708 | 0.24744486 | 38.47246946 | -83.90644614 |
| Casey County, Middleburg CCD | 0.2140936 | 0.24448628 | 37.25388599 | -84.79328047 |
| Knott County, Beaver Creek CCD | 0.14752555 | 0.24394836 | 37.32844548 | -82.8578626 |
| Hickman County, Columbus CCD | 0.19876278 | 0.24394836 | 36.77045362 | -89.0975499 |
| Harlan County, Cumberland CCD | 0.0957504 | 0.2436794 | 36.98098226 | -82.97655049 |
| Carter County, Grahn CCD | 0.14604626 | 0.24233459 | 38.32429663 | -83.11563965 |
| Barren County, Rocky Hill CCD | 0.13569123 | 0.24206563 | 37.00097627 | -86.00128586 |
| Muhlenberg County, Penrod CCD | 0.2084454 | 0.24206563 | 37.16778837 | -86.97860288 |
| Clark County, Southeast Clark CCD | 0.14308768 | 0.24206563 | 37.85854849 | -84.06529612 |
| Fulton County, Hickman CCD | 0.15747714 | 0.24179666 | 36.51917709 | -89.40310571 |
| Martin County, Inez CCD | 0.09830554 | 0.2415277 | 37.88099066 | -82.55606399 |
| Graves County, Fancy Farm CCD | 0.15115654 | 0.24125874 | 36.77279616 | -88.75713466 |
| Taylor County, Saloma CCD | 0.10745024 | 0.24125874 | 37.39373634 | -85.34605766 |
| Greenup County, Tygarts Valley CCD | 0.10368478 | 0.24072082 | 38.47596573 | -83.05960625 |
| Hart County, Hammonville CCD | 0.15277031 | 0.24045186 | 37.33931324 | -85.75511572 |
| Johnson County, Flatgap CCD | 0.11538462 | 0.24045186 | 37.99065757 | -82.94222447 |
| Knott County, Hindman CCD | 0.19257665 | 0.23964497 | 37.33151745 | -82.88607432 |
| Lawrence County, Clifford CCD | 0.12399139 | 0.23910705 | 38.0670719 | -82.6090568 |
| Pulaski County, Eubank CCD | 0.15505648 | 0.23910705 | 37.26016876 | -84.62038002 |
| Hart County, Bonnieville CCD | 0.1422808 | 0.23830016 | 37.33141252 | -85.92809002 |
| Wolfe County, Hazel Green CCD | 0.13058096 | 0.23722431 | 37.80134624 | -83.49651568 |
| Lawrence County, Webbville CCD | 0.16796665 | 0.23668639 | 38.12015888 | -82.89004789 |
| Crittenden County, East Crittenden CCD | 0.17213556 | 0.23641743 | 37.24522619 | -88.08320785 |
| Knott County, Mousie CCD | 0.08364712 | 0.23587951 | 37.35361395 | -82.88215507 |
| Hopkins County, Nortonville CCD | 0.10758472 | 0.23561054 | 37.20278197 | -87.45981448 |
| Edmonson County, Brownsville North CCD | 0.13407746 | 0.23561054 | 37.20162862 | -86.26016524 |
| Hancock County, Lewisport CCD | 0.08821947 | 0.23534158 | 37.92278851 | -86.88607244 |
| Muhlenberg County, Rosewood CCD | 0.19970414 | 0.23480366 | 37.09357164 | -87.1872234 |
| Martin County, Warfield CCD | 0.1668908 | 0.23480366 | 37.8431054 | -82.41813636 |
| Trigg County, Cadiz CCD | 0.12049489 | 0.23426573 | 36.88519371 | -87.723527 |
| Butler County, Sugar Grove CCD | 0.14241528 | 0.23318989 | 37.11973371 | -86.64199495 |
| Wayne County, Gregory CCD | 0.09938139 | 0.23318989 | 36.78334699 | -84.80698172 |
| Barren County, Temple Hill CCD | 0.18154922 | 0.23157612 | 36.94480784 | -85.86633883 |
| Butler County, Jetson CCD | 0.13959118 | 0.23157612 | 37.23293586 | -86.65632024 |
| Muhlenberg County, Bremen CCD | 0.12560516 | 0.23103819 | 37.33998946 | -87.18657089 |
| Green County, Greensburg CCD | 0.16433566 | 0.23023131 | 37.31251019 | -85.44369346 |
| Henderson County, Hebbardsville-Robards CCD | 0.07651963 | 0.22969338 | 37.65513443 | -87.52243667 |
| Wolfe County, Malaga CCD | 0.14120495 | 0.22969338 | 37.71289251 | -83.51153978 |
| Trigg County, Canton CCD | 0.17697687 | 0.2288865 | 36.83380942 | -87.95640824 |
| Elliott County, Sandy Hook CCD | 0.13864981 | 0.22861754 | 38.08718805 | -83.21544673 |
| Pike County, South Williamson CCD | 0.10516407 | 0.22781065 | 37.64733649 | -82.28084991 |
| Trigg County, Cerulean CCD | 0.16379774 | 0.22619688 | 36.90239338 | -87.97744473 |
| Graves County, Wingo-Water Valley CCD | 0.09494352 | 0.22592792 | 36.65984595 | -88.73973446 |
| Pike County, Dorton CCD | 0.12977407 | 0.22538999 | 37.25632655 | -82.61205811 |
| Owsley County, Sturgeon CCD | 0.15357719 | 0.22458311 | 37.44847488 | -83.78639664 |
| Letcher County, Colson CCD | 0.14631522 | 0.22377622 | 37.21488879 | -82.84929661 |
| Anderson County, Southwest Anderson CCD | 0.12130178 | 0.22377622 | 37.96153198 | -84.91672698 |
| Lincoln County, Hustonville CCD | 0.06885422 | 0.2232383 | 37.54712714 | -84.80632518 |
| Menifee County, West Menifee CCD | 0.1457773 | 0.22189349 | 37.96300386 | -83.75869457 |
| Carter County, Olive Hill CCD | 0.14698763 | 0.22135557 | 38.33331078 | -83.2014617 |
| Green County, Gresham CCD | 0.16325982 | 0.22081764 | 37.17204147 | -85.37785502 |
| Woodford County, South Woodford CCD | 0.12372243 | 0.22081764 | 38.01228562 | -84.735384 |
| Hickman County, Clinton CCD | 0.12775686 | 0.22001076 | 36.67490754 | -88.99310497 |
| Todd County, Kirkmansville CCD | 0.15438408 | 0.22001076 | 36.97209616 | -87.05922317 |
| Pulaski County, Shopville CCD | 0.12143626 | 0.2197418 | 37.15817641 | -84.49288884 |
| Clay County, Burning Springs CCD | 0.16083916 | 0.21893491 | 37.16084113 | -83.83374626 |
| Jackson County, Annville CCD | 0.13932222 | 0.21893491 | 37.33273316 | -83.97779608 |
| Morgan County, Elkfork-Ophir CCD | 0.15868747 | 0.21893491 | 37.99979315 | -83.027637 |
| Clay County, Oneida CCD | 0.1851802 | 0.21812803 | 37.27056431 | -83.58612504 |
| Russell County, Bryan CCD | 0.17079075 | 0.21732114 | 36.99167517 | -85.11822057 |
| Knox County, Girdler CCD | 0.0718128 | 0.21651425 | 36.92025492 | -83.90162389 |
| Menifee County, Frenchburg CCD | 0.16016676 | 0.21597633 | 37.98701327 | -83.55182094 |
| Breathitt County, Oakdale CCD | 0.10381926 | 0.21463152 | 37.59059298 | -83.41691976 |
| Greenup County, Wurtland CCD | 0.13797741 | 0.21436256 | 38.53370831 | -82.75705217 |
| Bourbon County, North Middletown CCD | 0.08364712 | 0.2140936 | 38.18766584 | -83.98256944 |
| Perry County, Krypton CCD | 0.14349112 | 0.21382464 | 37.34874734 | -83.25684663 |
| Jackson County, McKee CCD | 0.16352878 | 0.21328671 | 37.42406732 | -83.99169447 |
| Nelson County, Boston CCD | 0.1048951 | 0.21274879 | 37.77069572 | -85.5510261 |
| Breckinridge County, Irvington CCD | 0.11726735 | 0.21247983 | 37.87646399 | -86.28626873 |
| Clay County, Ashers Fork CCD | 0.13206025 | 0.2119419 | 37.04063561 | -83.538689 |
| Pike County, Mayflower CCD | 0.09332975 | 0.21167294 | 37.58131223 | -82.49527195 |
| Laurel County, Camp Ground CCD | 0.08741259 | 0.20898332 | 37.04501294 | -84.02433542 |
| Knott County, Emmalena CCD | 0.09306079 | 0.2084454 | 37.35585686 | -83.11185142 |
| Edmonson County, Brownsville CCD | 0.12587413 | 0.20710059 | 37.18171431 | -86.2913508 |
| Hardin County, West Point CCD | 0.20710059 | 0.20710059 | 37.98411126 | -85.95423362 |
| Breathitt County, Rousseau CCD | 0.07772996 | 0.20683163 | 37.56257978 | -83.36404739 |
| Garrard County, Bryantsville CCD | 0.08162991 | 0.20602474 | 37.82716842 | -84.67709137 |
| Breckinridge County, Cloverport CCD | 0.1007262 | 0.20548682 | 37.87226391 | -86.51753546 |
| Union County, Uniontown CCD | 0.08189887 | 0.20467994 | 37.88422432 | -87.92816884 |
| Owsley County, Cowcreek CCD | 0.16258741 | 0.20145239 | 37.31770629 | -83.60694185 |
| Lincoln County, Crab Orchard CCD | 0.10570199 | 0.20091447 | 37.52594413 | -84.579442 |
| Leslie County, Mozelle CCD | 0.1105433 | 0.19849381 | 37.01559607 | -83.51427773 |
| McCreary County, Pine Knot CCD | 0.10785368 | 0.19822485 | 36.65196399 | -84.37176046 |
| Harlan County, Poor Fork CCD | 0.0866057 | 0.19795589 | 36.89759373 | -83.24837181 |
| Bath County, Salt Lick CCD | 0.11807423 | 0.19768693 | 38.15632029 | -83.66826726 |
| Floyd County, Grethel-Teaberry CCD | 0.09803658 | 0.19741797 | 37.53076512 | -82.64876779 |
| McLean County, Island CCD | 0.11444325 | 0.197149 | 37.44897828 | -87.14495496 |
| Casey County, Clementsville CCD | 0.14631522 | 0.19688004 | 37.37355937 | -85.11949075 |
| Ohio County, Hartford CCD | 0.15922539 | 0.1958042 | 37.43216102 | -86.91776687 |
| Magoffin County, West Magoffin CCD | 0.10005379 | 0.19499731 | 37.74665341 | -83.18791849 |
| Perry County, Dice CCD | 0.13811189 | 0.19472835 | 37.33977046 | -83.17215337 |
| Pike County, Sidney CCD | 0.10839161 | 0.1936525 | 37.72316489 | -82.35350126 |
| Harlan County, Cawood CCD | 0.09386767 | 0.19311458 | 36.7840445 | -83.23926368 |
| Garrard County, Paint Lick CCD | 0.05459925 | 0.19311458 | 37.57348368 | -84.37057098 |
| Nelson County, Bloomfield CCD | 0.13622916 | 0.19203873 | 37.89344252 | -85.15812867 |
| Morgan County, Grassy Creek CCD | 0.15761162 | 0.19176977 | 37.90897459 | -83.4492274 |
| Harlan County, Wallins Creek CCD | 0.08364712 | 0.19150081 | 36.82985328 | -83.45978252 |
| Marion County, St. Mary CCD | 0.08821947 | 0.19069392 | 37.57107432 | -85.32048296 |
| Morgan County, Cannel City CCD | 0.14214632 | 0.19042496 | 37.7615615 | -83.32986464 |
| Nelson County, New Haven CCD | 0.08284024 | 0.18988704 | 37.66227786 | -85.59516136 |
| Morgan County, Wrigley CCD | 0.13071544 | 0.18827327 | 37.99741726 | -83.43408862 |
| Grayson County, Short Creek CCD | 0.12157074 | 0.18692846 | 37.60147581 | -86.49481793 |
| Madison County, Cottonburg-Valley View CCD | 0.07342657 | 0.18665949 | 37.75604344 | -84.39946173 |
| Harlan County, Pine Mountain CCD | 0.08351264 | 0.18639053 | 36.88736558 | -83.36440048 |
| Edmonson County, Sunfish CCD | 0.11565358 | 0.18585261 | 37.23740119 | -86.42704525 |
| Adair County, Casey Creek CCD | 0.15357719 | 0.18558365 | 37.25321884 | -85.28033157 |
| Trigg County, Fort Campbell CCD | 0.13313609 | 0.18531469 | 36.63872648 | -87.72241433 |
| Pike County, Fedscreek CCD | 0.10812265 | 0.18423884 | 37.4104968 | -82.18078371 |
| Webster County, Clay CCD | 0.1542496 | 0.18370091 | 37.51910415 | -87.82466565 |
| Letcher County, Eolia CCD | 0.10462614 | 0.18343195 | 37.01600803 | -82.88450837 |
| Hardin County, Sonora CCD | 0.06159225 | 0.18343195 | 37.53105477 | -85.9008989 |
| Ohio County, Fordsville CCD | 0.13690156 | 0.18343195 | 37.54812197 | -86.67391953 |
| Perry County, Daisy CCD | 0.11457773 | 0.18235611 | 37.0463754 | -83.16556031 |
| Breathitt County, Haddix CCD | 0.0809575 | 0.18208714 | 37.51835889 | -83.37161867 |
| Lee County, St. Helens CCD | 0.11242604 | 0.18154922 | 37.57259142 | -83.67940007 |
| Ohio County, Heflin-Beda CCD | 0.10933297 | 0.1810113 | 37.49877854 | -86.90959446 |
| Marion County, Bradfordsville CCD | 0.12291555 | 0.18047337 | 37.43493049 | -85.05304204 |
| Larue County, Hodgenville CCD | 0.05782679 | 0.18047337 | 37.63242787 | -85.77197959 |
| Knott County, Carr Fork CCD | 0.13636364 | 0.17966649 | 37.28425336 | -82.86571054 |
| Metcalfe County, Center CCD | 0.11659494 | 0.17778375 | 37.09570162 | -85.53160582 |
| Morgan County, West Liberty CCD | 0.15277031 | 0.17590102 | 37.86016799 | -83.19706637 |
| Harlan County, Alva CCD | 0.0626681 | 0.17563206 | 36.66913493 | -83.42981103 |
| Casey County, Dividing Ridge CCD | 0.07974718 | 0.17563206 | 37.39987146 | -84.91471436 |
| Pike County, Raccoon CCD | 0.09803658 | 0.17509414 | 37.53081943 | -82.44681727 |
| Breathitt County, Canoe CCD | 0.10381926 | 0.17428725 | 37.33400871 | -83.54675228 |
| Perry County, Buckhorn CCD | 0.15008069 | 0.17428725 | 37.28685668 | -83.52596902 |
| Pulaski County, Mount Victory CCD | 0.12210866 | 0.17374933 | 37.06958677 | -84.50660268 |
| Lewis County, Laurel-Petersville CCD | 0.09104357 | 0.1732114 | 38.49852 | -83.49357931 |
| Jackson County, Sandgap CCD | 0.11430877 | 0.1732114 | 37.3882059 | -84.08371343 |
| Floyd County, Wayland CCD | 0.06791286 | 0.17186659 | 37.48310866 | -82.8303213 |
| McLean County, Calhoun CCD | 0.09817106 | 0.17105971 | 37.63092651 | -87.4964995 |
| Bell County, Kentucky Ridge CCD | 0.09319527 | 0.17052179 | 36.72324353 | -83.73415131 |
| Webster County, Dixon CCD | 0.10516407 | 0.16756321 | 37.58192792 | -87.76035068 |
| Green County, Donansburg CCD | 0.13676708 | 0.16729424 | 37.22874965 | -85.51284405 |
| Pike County, Long Fork CCD | 0.09991931 | 0.16675632 | 37.24048812 | -82.67238668 |
| Green County, Summersville CCD | 0.12883271 | 0.16675632 | 37.36914188 | -85.57898746 |
| Hardin County, Summit CCD | 0.07073696 | 0.16648736 | 37.60563397 | -86.26439456 |
| Grayson County, Clarkson CCD | 0.13636364 | 0.16648736 | 37.51230249 | -86.21926786 |
| Graves County, Sedalia CCD | 0.10476062 | 0.16594944 | 36.51601811 | -88.58407985 |
| Pike County, Robinson Creek CCD | 0.10233997 | 0.16433566 | 37.38256192 | -82.53160349 |
| Owsley County, Booneville CCD | 0.12103281 | 0.16379774 | 37.46773539 | -83.75443879 |
| McCreary County, Co-operative CCD | 0.09440559 | 0.16325982 | 36.61770296 | -84.71811796 |
| Ballard County, La Center CCD | 0.11995697 | 0.16299086 | 37.15037825 | -89.0908357 |
| Pike County, Phelps CCD | 0.12318451 | 0.1605702 | 37.53643815 | -82.09942952 |
| Breathitt County, Hardshell CCD | 0.07934373 | 0.15949435 | 37.53157051 | -83.28290874 |
| Knox County, Trosper CCD | 0.06401291 | 0.1570737 | 36.82605217 | -83.85986666 |
| Breckinridge County, Glen Dean-Vanzant CCD | 0.11081226 | 0.15626681 | 37.61033226 | -86.60860939 |
| Marion County, New Market CCD | 0.09306079 | 0.15303927 | 37.45773867 | -85.30812886 |
| Lee County, Southwest Lee CCD | 0.11753631 | 0.15061861 | 37.5312302 | -83.71510535 |
| Christian County, Crofton CCD | 0.08391608 | 0.14900484 | 37.055306 | -87.27724835 |
| Whitley County, Cumberland Falls CCD | 0.09037117 | 0.14819796 | 36.77388383 | -84.23881422 |
| Johnson County, Nippa CCD | 0.09615385 | 0.14792899 | 37.85866487 | -82.76295535 |
| Larue County, Buffalo CCD | 0.04357181 | 0.14604626 | 37.48973775 | -85.67685998 |
| Hancock County, Patesville CCD | 0.09655729 | 0.14550834 | 37.78314301 | -86.81695959 |
| Leslie County, Cutshin CCD | 0.09817106 | 0.14308768 | 37.13061979 | -83.32297593 |
| Bullitt County, Fort Knox CCD | 0.06993007 | 0.1422808 | 37.96987063 | -85.78281532 |
| Whitley County, Pearl CCD | 0.10435718 | 0.13986014 | 36.59386518 | -84.01128916 |
| McCreary County, Parkers Lake CCD | 0.07894029 | 0.13905325 | 36.84270958 | -84.48456069 |
| Knox County, Bryants Store CCD | 0.07799892 | 0.13609467 | 36.83172453 | -83.91599292 |
| McLean County, Sacramento CCD | 0.08902636 | 0.13582571 | 37.3864784 | -87.20967359 |
| Knox County, Dewitt CCD | 0.07261969 | 0.1344809 | 36.93874251 | -83.64148522 |
| Marion County, Loretto CCD | 0.08956428 | 0.1344809 | 37.71403901 | -85.41501979 |
| Perry County, Defiance-Vicco CCD | 0.10220549 | 0.13313609 | 37.22021518 | -83.10674322 |
| Floyd County, Wheelwright-Weeksbury CCD | 0.10005379 | 0.13017751 | 37.38236721 | -82.73315112 |
| Floyd County, McDowell CCD | 0.05944056 | 0.12990855 | 37.37614571 | -82.67670419 |
| Pike County, Ashcamp CCD | 0.08297472 | 0.12641205 | 37.2248139 | -82.5315685 |
| Hopkins County, St. Charles CCD | 0.07894029 | 0.12426036 | 37.14604108 | -87.6089552 |
| Magoffin County, Royalton CCD | 0.05890264 | 0.12264658 | 37.65005674 | -82.91598782 |
| Carter County, Upper Tygart CCD | 0.08068854 | 0.12022593 | 38.2670033 | -83.21308106 |
| Bell County, Tejay CCD | 0.05554061 | 0.119688 | 36.79146895 | -83.52328922 |
| Marion County, Gravel Switch CCD | 0.06078537 | 0.11861216 | 37.55594599 | -85.15014535 |
| Harlan County, Upper Clover Fork CCD | 0.08391608 | 0.11403981 | 36.94285584 | -83.04043259 |
| McLean County, Livermore CCD | 0.08418505 | 0.11350188 | 37.55586045 | -87.07477212 |
| Pike County, Elkhorn City CCD | 0.0809575 | 0.11188811 | 37.34464455 | -82.39461222 |
| Pike County, Hellier CCD | 0.08956428 | 0.10516407 | 37.35416376 | -82.4792147 |
| Pike County, McCarr CCD | 0.08122647 | 0.1048951 | 37.6123768 | -82.17178658 |
| Bell County, Pruden-Fonde CCD | 0.07907477 | 0.09682625 | 36.65182029 | -83.91663025 |
| Marion County, Raywick CCD | 0.04948897 | 0.08041958 | 37.49234864 | -85.44674843 |
