## Supplementary material for "Mapping long-distance jump dispersal risk by leveraging transport infrastructure data and sampling bias correction: a case study with the invasive spotted lanternfly (*Lycorma delicatula*)": S6 Maryland risk

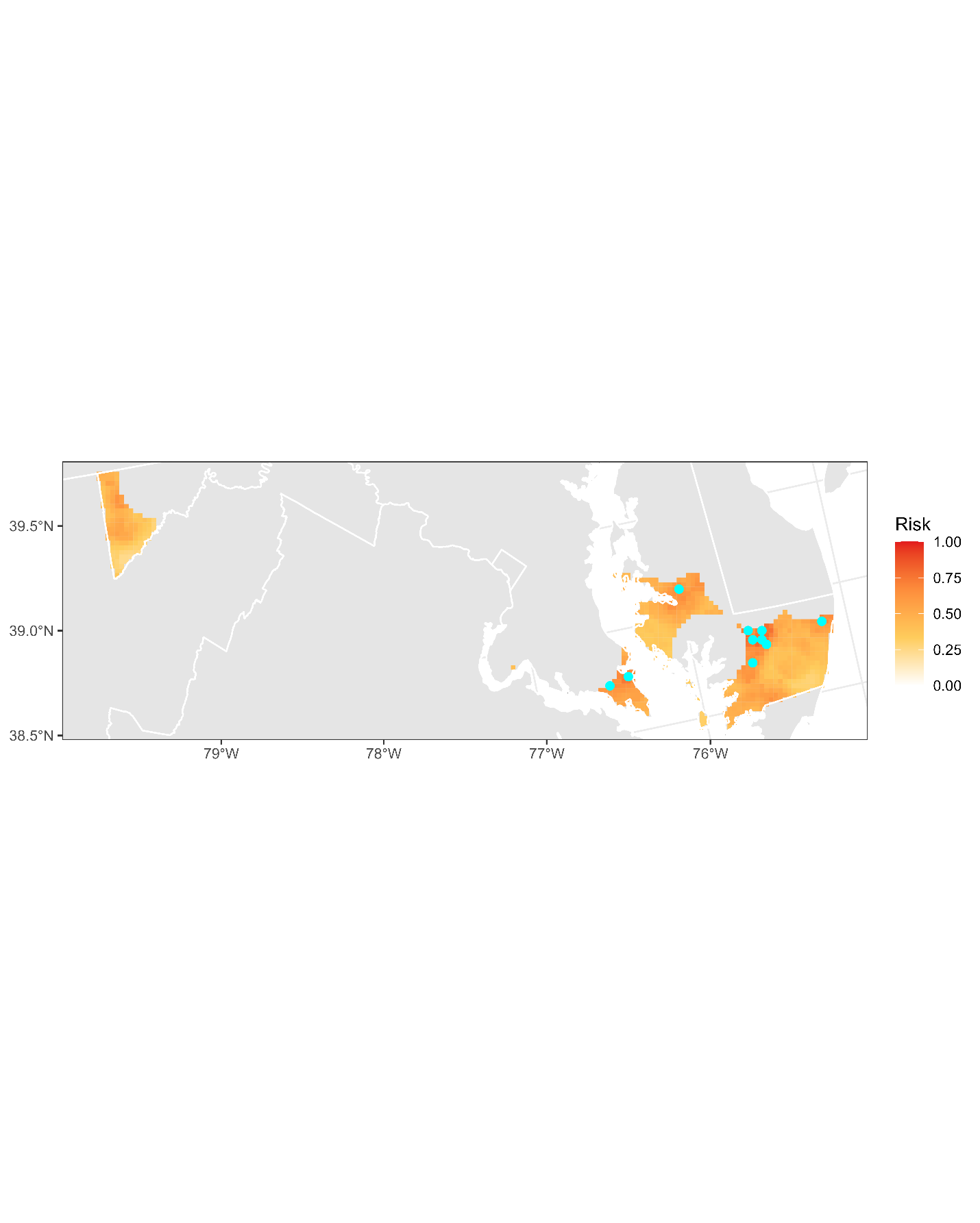

**Figure S6.1:** Projected risk of colonization by *L. delicatula* jump dispersal for Maryland. Risk projections are based on an ensemble random forest model including distances from existing *L. delicatula* populations. Note that areas falling less than 20 km from existing *L. delicatula* populations are not included in the map because any colonization there would be categorized as diffusive spread based on our methodology (regardless of the actual source of the resulting population). Darker, redder colors indicate increased risk of colonization by jump dispersal. Cyan points show the top ten highest risk locations in the state.

**Table S6.1:** Ranked list of Maryland county subdivisions sorted by the maximum projected risk of colonization by jump dispersal occurring there. For each subdivision, we identified the 2.5 x 2.5 km grid cell centroid included there with the highest projected risk of colonization by jump dispersal and report that location and risk value.

| **County, subdivision** | **Median risk value** | **Maximum risk value** | **Latitude of maximum risk** | **Longitude of maximum risk** |
| --- | --- | --- | --- | --- |
| Wicomico County, District 5, Parsons | 0.75295858 | 0.81871974 | 38.35220458 | -75.54156557 |
| Wicomico County, District 8, Nutters | 0.76358257 | 0.80930608 | 38.30862518 | -75.55389669 |
| Wicomico County, District 16, Fruitland | 0.77380312 | 0.80177515 | 38.31813576 | -75.61029833 |
| St. Mary's County, District 8, Bay | 0.67711135 | 0.78052717 | 38.2693411 | -76.42045818 |
| Wicomico County, District 9, Salisbury | 0.73318989 | 0.73937601 | 38.36646919 | -75.62622135 |
| Worcester County, District 3, Sinepuxent | 0.42469069 | 0.69956966 | 38.33325998 | -75.16255355 |
| Worcester County, District 4, Western | 0.43464228 | 0.69849381 | 38.28207139 | -75.53186939 |
| St. Mary's County, District 2, Valley Lee | 0.57100592 | 0.690156 | 38.24386411 | -76.54516603 |
| Talbot County, District 3, Trappe | 0.55513717 | 0.66568047 | 38.63323619 | -75.99561879 |
| Somerset County, District 15, East Princess Anne | 0.62157074 | 0.66460463 | 38.20916731 | -75.64098261 |
| Dorchester County, District 14, Linkwood | 0.56051641 | 0.66352878 | 38.58495098 | -75.97939379 |
| Wicomico County, District 13, Camden | 0.64281872 | 0.65599785 | 38.33992844 | -75.60415054 |
| Dorchester County, District 7, Cambridge | 0.52649274 | 0.64954276 | 38.5552831 | -76.07646443 |
| Worcester County, District 1, Southern | 0.3894567 | 0.64792899 | 38.01582522 | -75.57751295 |
| Dorchester County, District 15, Hurlock | 0.57853685 | 0.63986014 | 38.63634865 | -75.87615286 |
| Dorchester County, District 2, East New Market | 0.57261969 | 0.63743948 | 38.60676189 | -75.97333359 |
| Somerset County, District 1, West Princess Anne | 0.6161915 | 0.63717052 | 38.19684053 | -75.70342922 |
| Garrett County, District 6, Sang Run | 0.4817106 | 0.63071544 | 39.57043659 | -79.38345662 |
| Somerset County, District 13, Westover | 0.52528241 | 0.62775686 | 38.14851651 | -75.6874922 |
| St. Mary's County, District 3, Leonardtown | 0.61699839 | 0.61861216 | 38.23107816 | -76.60748749 |
| Wicomico County, District 6, Dennis | 0.48480366 | 0.61296396 | 38.31609645 | -75.46313404 |
| St. Mary's County, District 1, St. Inigoes | 0.52071006 | 0.60758472 | 38.22111183 | -76.40403464 |
| Worcester County, District 2, Central | 0.42912856 | 0.59897795 | 38.33810819 | -75.19073642 |
| Dorchester County, District 12, Williamsburg | 0.59494352 | 0.59494352 | 38.67996012 | -75.86395869 |
| Somerset County, District 3, Brinkleys | 0.50941366 | 0.57772996 | 38.05659683 | -75.68378814 |
| Dorchester County, District 13, Bucktown | 0.48130715 | 0.577461 | 38.5459864 | -76.01982011 |
| Calvert County, District 1, Solomons Island | 0.51465842 | 0.57719204 | 38.33940119 | -76.43100118 |
| Garrett County, District 14, West Oakland | 0.49273803 | 0.56374395 | 39.41665883 | -79.41914162 |
| Somerset County, District 4, Dublin | 0.45965573 | 0.56132329 | 38.06891384 | -75.62145711 |
| Garrett County, District 2, Friendsville | 0.47364174 | 0.55675094 | 39.66606636 | -79.42091011 |
| Garrett County, District 5, Accident | 0.51492738 | 0.54760624 | 39.61437264 | -79.37323296 |
| Caroline County, District 4, Preston | 0.54572351 | 0.54572351 | 38.68932839 | -75.92068701 |
| Dorchester County, District 17, Salem | 0.48063475 | 0.54545455 | 38.53199262 | -75.93488064 |
| Wicomico County, District 4, Pittsburg | 0.51237224 | 0.53335126 | 38.38145135 | -75.44454912 |
| Wicomico County, District 15, Hebron | 0.52985476 | 0.52985476 | 38.3977428 | -75.67654567 |
| Garrett County, District 16, Mountain Lake Park | 0.41312534 | 0.52985476 | 39.40890088 | -79.36142818 |
| Garrett County, District 7, East Oakland | 0.47660032 | 0.52635826 | 39.41278319 | -79.39028336 |
| Dorchester County, District 3, Vienna | 0.44123185 | 0.5112964 | 38.54912233 | -75.90049759 |
| Wicomico County, District 14, Willards | 0.49731038 | 0.50645508 | 38.37186479 | -75.38812041 |
| Somerset County, District 8, Lawsons | 0.48856912 | 0.50080689 | 38.01006418 | -75.81444932 |
| Garrett County, District 10, Deer Park | 0.40909091 | 0.4873588 | 39.42697672 | -79.32746207 |
| Somerset County, District 7, Crisfield | 0.46906939 | 0.48601399 | 37.99295787 | -75.84860105 |
| Talbot County, District 2, St. Michaels | 0.46732114 | 0.48574502 | 38.70472145 | -76.15376328 |
| Wicomico County, District 1, Barren Creek | 0.4704142 | 0.48305541 | 38.48202908 | -75.77121307 |
| Dorchester County, District 8, Neck | 0.43316299 | 0.48144164 | 38.57379846 | -76.18979617 |
| Somerset County, District 6, Fairmount | 0.43302851 | 0.47821409 | 38.0801766 | -75.82437867 |
| Dorchester County, District 9, Church Creek | 0.38461538 | 0.47417967 | 38.51628678 | -76.1168227 |
| Somerset County, District 12, Asbury | 0.4669177 | 0.47122109 | 37.96645609 | -75.82656203 |
| Talbot County, District 5, Bay Hundred | 0.43329747 | 0.46207639 | 38.68869239 | -76.33608755 |
| Garrett County, District 1, Swanton | 0.37062937 | 0.4521248 | 39.44504342 | -79.29347898 |
| Somerset County, District 2, St. Peters | 0.42092523 | 0.42092523 | 38.14558686 | -75.80617421 |
| Dorchester County, District 1, Fork | 0.40667025 | 0.40667025 | 38.5350464 | -75.8155774 |
| Charles County, District 3, Nanjemoy | 0.40344271 | 0.40344271 | 38.42511353 | -77.11606317 |
| Dorchester County, District 16, Madison | 0.36995697 | 0.37977407 | 38.46468831 | -76.21972522 |
| Dorchester County, District 4, Taylors Island | 0.35691232 | 0.3781603 | 38.49571857 | -76.27036933 |
| Dorchester County, District 5, Lakes | 0.35449166 | 0.37762238 | 38.38537497 | -76.15282896 |
| Garrett County, District 8, Red House | 0.28684777 | 0.36578806 | 39.21893961 | -79.46480002 |
| Dorchester County, District 6, Hoopers Island | 0.35785368 | 0.3633674 | 38.32914241 | -76.22727286 |
| Garrett County, District 13, Kitzmiller | 0.32208176 | 0.3577192 | 39.43722732 | -79.23575644 |
| Somerset County, District 10, Smith Island | 0.31522324 | 0.33539537 | 38.09110335 | -76.02736135 |
| Dorchester County, District 10, Straits | 0.33149543 | 0.33458849 | 38.18766069 | -76.05967777 |
