## Supplementary material for "Mapping long-distance jump dispersal risk by leveraging transport infrastructure data and sampling bias correction: a case study with the invasive spotted lanternfly (*Lycorma delicatula*)": S7 Massachusetts risk

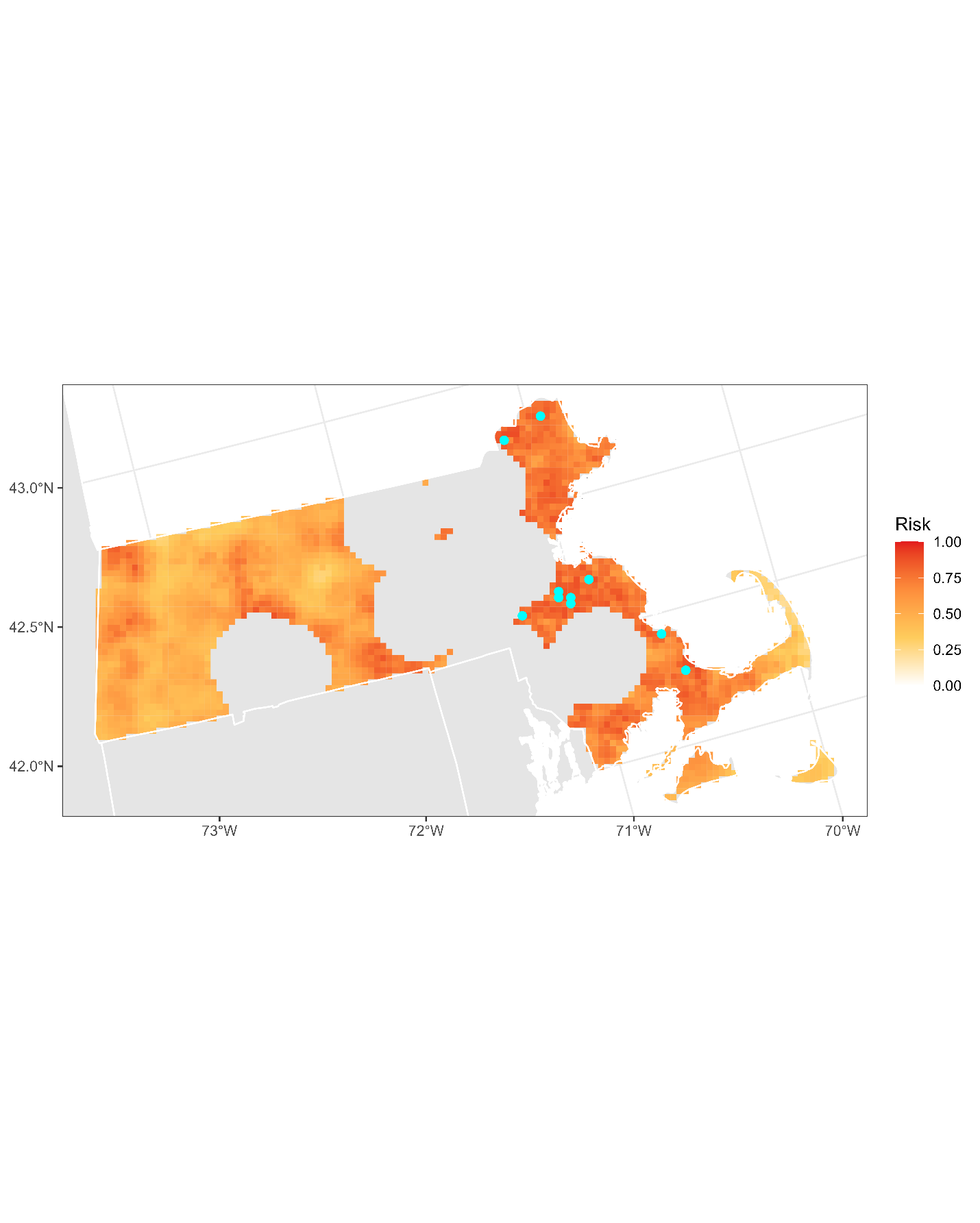

**Figure S7.1:** Projected risk of colonization by *L. delicatula* jump dispersal for Massachusetts. Risk projections are based on an ensemble random forest model including distances from existing *L. delicatula* populations. Note that areas falling less than 20 km from existing *L. delicatula* populations are not included in the map because any colonization there would be categorized as diffusive spread based on our methodology (regardless of the actual source of the resulting population). Darker, redder colors indicate increased risk of colonization by jump dispersal. Cyan points show the top ten highest risk locations in the state.

**Table S7.1:** Ranked list of Massachusetts county subdivisions sorted by the maximum projected risk of colonization by jump dispersal occurring there. For each subdivision, we identified the 2.5 x 2.5 km grid cell centroid included there with the highest projected risk of colonization by jump dispersal and report that location and risk value.

| **County, subdivision** | **Median risk value** | **Maximum risk value** | **Latitude of maximum risk** | **Longitude of maximum risk** |
| --- | --- | --- | --- | --- |
| Essex County, Haverhill city | 0.84077461 | 0.90989779 | 42.76639682 | -71.13843646 |
| Norfolk County, Randolph Town city | 0.86538462 | 0.90909091 | 42.17429203 | -71.07056354 |
| Norfolk County, Avon town | 0.83337816 | 0.90290479 | 42.11951534 | -71.02755228 |
| Plymouth County, Hingham town | 0.78133405 | 0.89214632 | 42.18838316 | -70.90761534 |
| Norfolk County, Stoughton town | 0.82423346 | 0.88972566 | 42.15270322 | -71.07846146 |
| Barnstable County, Bourne town | 0.78442711 | 0.88676708 | 41.7709812 | -70.55834842 |
| Essex County, Newburyport city | 0.77192039 | 0.88246369 | 42.81795455 | -70.92840984 |
| Plymouth County, Plymouth town | 0.69903174 | 0.88111888 | 41.92390022 | -70.62726741 |
| Norfolk County, Holbrook town | 0.85018827 | 0.87681549 | 42.14110015 | -71.01964236 |
| Norfolk County, Walpole town | 0.80594406 | 0.87627757 | 42.12255221 | -71.2785482 |
| Plymouth County, Brockton city | 0.83700914 | 0.87547068 | 42.08214497 | -71.07273925 |
| Norfolk County, Weymouth Town city | 0.79424422 | 0.87493276 | 42.17845466 | -70.97438874 |
| Essex County, Danvers town | 0.80957504 | 0.87385691 | 42.57456108 | -70.98672152 |
| Hampshire County, Amherst Town city | 0.77165143 | 0.87170522 | 42.35266424 | -72.48872071 |
| Plymouth County, Pembroke town | 0.78509952 | 0.87143626 | 42.09442253 | -70.78456217 |
| Barnstable County, Sandwich town | 0.73964497 | 0.87009145 | 41.75914734 | -70.4999365 |
| Essex County, Saugus town | 0.77192039 | 0.86955352 | 42.43918306 | -71.00493273 |
| Essex County, Methuen Town city | 0.84722969 | 0.86713287 | 42.77217951 | -71.1681194 |
| Essex County, Peabody city | 0.81038193 | 0.8655191 | 42.52555297 | -70.97308961 |
| Middlesex County, Woburn city | 0.81737493 | 0.86471221 | 42.48977049 | -71.14473159 |
| Plymouth County, Rockland town | 0.8148198 | 0.86363636 | 42.12364578 | -70.93145045 |
| Bristol County, Dartmouth town | 0.71247983 | 0.86256052 | 41.6531075 | -71.00944547 |
| Hampshire County, Hadley town | 0.74878967 | 0.86202259 | 42.39074339 | -72.57012547 |
| Norfolk County, Canton town | 0.7408553 | 0.86175363 | 42.14268713 | -71.14517625 |
| Bristol County, Fairhaven town | 0.7767617 | 0.85906401 | 41.65722255 | -70.91398087 |
| Worcester County, Charlton town | 0.75578268 | 0.85798817 | 42.10151752 | -72.0097665 |
| Plymouth County, Hanson town | 0.85691232 | 0.85691232 | 42.08455862 | -70.85126318 |
| Norfolk County, Sharon town | 0.79747176 | 0.85637439 | 42.13263678 | -71.21187185 |
| Suffolk County, Revere city | 0.81508876 | 0.8542227 | 42.433366 | -70.97540561 |
| Bristol County, Norton town | 0.80204411 | 0.85180204 | 41.99154132 | -71.20017296 |
| Plymouth County, Norwell town | 0.77756859 | 0.85018827 | 42.12185163 | -70.80595618 |
| Norfolk County, Quincy city | 0.79478214 | 0.84669177 | 42.23905902 | -71.0468404 |
| Norfolk County, Braintree Town city | 0.75336202 | 0.845078 | 42.18427005 | -71.00380783 |
| Berkshire County, Adams town | 0.68881119 | 0.84427111 | 42.65058512 | -73.11266489 |
| Essex County, Amesbury Town city | 0.76627219 | 0.84265734 | 42.82378542 | -70.95809728 |
| Plymouth County, Abington town | 0.81091985 | 0.8415815 | 42.12947054 | -70.96084285 |
| Plymouth County, Hanover town | 0.77541689 | 0.84050565 | 42.09040131 | -70.88063242 |
| Bristol County, New Bedford city | 0.7506724 | 0.83781603 | 41.68461666 | -70.93529216 |
| Essex County, Lynnfield town | 0.77689618 | 0.83727811 | 42.5097766 | -71.01061543 |
| Hampshire County, Hatfield town | 0.71651425 | 0.83351264 | 42.37995668 | -72.63684513 |
| Essex County, Salisbury town | 0.78227542 | 0.83324368 | 42.83371407 | -70.89069794 |
| Middlesex County, Winchester town | 0.83216783 | 0.83216783 | 42.46238465 | -71.12309055 |
| Essex County, Gloucester city | 0.70965573 | 0.83136095 | 42.6141137 | -70.71799408 |
| Essex County, Merrimac town | 0.73722431 | 0.83082302 | 42.82960959 | -70.98778973 |
| Norfolk County, Norfolk town | 0.8302851 | 0.8302851 | 42.10669611 | -71.31578817 |
| Essex County, Lynn city | 0.72296934 | 0.82894029 | 42.50396027 | -70.98105781 |
| Worcester County, Southbridge Town city | 0.79370629 | 0.82813341 | 42.06380706 | -72.0544137 |
| Bristol County, Fall River city | 0.76869285 | 0.82732652 | 41.74102365 | -71.10277876 |
| Berkshire County, Williamstown town | 0.6570737 | 0.82705756 | 42.73174373 | -73.18016283 |
| Essex County, Georgetown town | 0.78752017 | 0.82409898 | 42.72159696 | -71.0277331 |
| Middlesex County, Wakefield town | 0.77272727 | 0.82248521 | 42.51558626 | -71.04017799 |
| Essex County, Salem city | 0.79827864 | 0.81925767 | 42.51389634 | -70.9139722 |
| Essex County, North Andover town | 0.76143088 | 0.81791286 | 42.68999353 | -71.10295515 |
| Essex County, Nahant town | 0.81764389 | 0.81764389 | 42.44913058 | -70.93791307 |
| Barnstable County, Mashpee town | 0.75013448 | 0.81737493 | 41.66705531 | -70.50284797 |
| Plymouth County, Rochester town | 0.61941904 | 0.81683701 | 41.76907056 | -70.77884381 |
| Middlesex County, Stoneham town | 0.77998924 | 0.81468531 | 42.45659426 | -71.0935437 |
| Worcester County, Webster town | 0.8006993 | 0.81226466 | 42.02463771 | -71.84833414 |
| Suffolk County, Boston city | 0.71369016 | 0.8119957 | 42.39018582 | -70.99131451 |
| Berkshire County, North Adams city | 0.72095212 | 0.8119957 | 42.69936518 | -73.12783223 |
| Worcester County, Sturbridge town | 0.72189349 | 0.81145777 | 42.13431655 | -72.06112692 |
| Plymouth County, Kingston town | 0.79142012 | 0.809844 | 42.01799132 | -70.74980875 |
| Essex County, Marblehead town | 0.75322754 | 0.80957504 | 42.48647128 | -70.89241471 |
| Essex County, Newbury town | 0.72996235 | 0.80715438 | 42.76892338 | -70.91477128 |
| Plymouth County, Wareham town | 0.76143088 | 0.80607854 | 41.75145993 | -70.69118 |
| Worcester County, Dudley town | 0.73318989 | 0.80392684 | 42.03586227 | -71.90719042 |
| Norfolk County, Foxborough town | 0.75605164 | 0.80338892 | 42.06208621 | -71.20601649 |
| Bristol County, Westport town | 0.68141474 | 0.80231307 | 41.63733729 | -71.04649716 |
| Essex County, Swampscott town | 0.80150619 | 0.80150619 | 42.46488468 | -70.90040351 |
| Middlesex County, Reading town | 0.79693384 | 0.80123722 | 42.5055851 | -71.10724709 |
| Bristol County, Easton town | 0.69311458 | 0.80123722 | 42.04476051 | -71.11787786 |
| Essex County, Topsfield town | 0.73480366 | 0.80016138 | 42.61192526 | -70.94116714 |
| Hampshire County, Belchertown town | 0.72027972 | 0.79854761 | 42.34170024 | -72.42950354 |
| Plymouth County, Mattapoisett town | 0.64335664 | 0.79800968 | 41.66711413 | -70.84769463 |
| Berkshire County, Pittsfield city | 0.67011834 | 0.79289941 | 42.46041113 | -73.20803914 |
| Barnstable County, Falmouth town | 0.65532544 | 0.78967187 | 41.64759353 | -70.63549842 |
| Essex County, Wenham town | 0.73292093 | 0.78913394 | 42.60609232 | -70.91157182 |
| Bristol County, Acushnet town | 0.77057558 | 0.78644432 | 41.67878907 | -70.90608813 |
| Plymouth County, Duxbury town | 0.73399677 | 0.7859064 | 42.06699079 | -70.76318496 |
| Essex County, Rowley town | 0.72485207 | 0.78133405 | 42.74148862 | -70.89313403 |
| Franklin County, Whately town | 0.65344271 | 0.77918236 | 42.42333135 | -72.62196322 |
| Norfolk County, Cohasset town | 0.76438946 | 0.77918236 | 42.24142837 | -70.82486463 |
| Plymouth County, Scituate town | 0.66191501 | 0.77810651 | 42.21399833 | -70.80342647 |
| Bristol County, Mansfield town | 0.75658956 | 0.77783755 | 42.03472544 | -71.1844839 |
| Essex County, Middleton town | 0.75591716 | 0.77703066 | 42.62357109 | -71.00037272 |
| Essex County, West Newbury town | 0.75188273 | 0.77649274 | 42.77475693 | -70.94443654 |
| Worcester County, Harvard town | 0.76277569 | 0.77514793 | 42.49949434 | -71.55284562 |
| Barnstable County, Barnstable Town city | 0.67348037 | 0.77407208 | 41.69002125 | -70.27457934 |
| Plymouth County, Marion town | 0.65196342 | 0.7732652 | 41.74164811 | -70.75756799 |
| Plymouth County, Hull town | 0.7239107 | 0.7711135 | 42.27470094 | -70.87576625 |
| Essex County, Boxford town | 0.67563206 | 0.76976869 | 42.72740444 | -71.05738756 |
| Norfolk County, Milton town | 0.76734804 | 0.76734804 | 42.26064823 | -71.03892288 |
| Middlesex County, Melrose city | 0.75605164 | 0.76600323 | 42.44499346 | -71.03446478 |
| Essex County, Groveland town | 0.76519634 | 0.76600323 | 42.76480144 | -71.01177071 |
| Plymouth County, Marshfield town | 0.71906939 | 0.76519634 | 42.13756107 | -70.76859282 |
| Worcester County, Spencer town | 0.71113502 | 0.76143088 | 42.28584913 | -72.00788893 |
| Middlesex County, Boxborough town | 0.76143088 | 0.76143088 | 42.49380139 | -71.52324742 |
| Middlesex County, Malden city | 0.7471759 | 0.75981711 | 42.42920112 | -71.07193317 |
| Hampshire County, Pelham town | 0.59951587 | 0.75901022 | 42.36886169 | -72.45161926 |
| Franklin County, Greenfield city | 0.68491124 | 0.75901022 | 42.58060418 | -72.59944689 |
| Essex County, Hamilton town | 0.71516945 | 0.75874126 | 42.64927544 | -70.89556364 |
| Middlesex County, North Reading town | 0.74287251 | 0.75551372 | 42.5861856 | -71.04590284 |
| Essex County, Ipswich town | 0.6908284 | 0.75255514 | 42.67670616 | -70.91717305 |
| Essex County, Beverly city | 0.71086606 | 0.75255514 | 42.57866339 | -70.88998662 |
| Hampden County, Palmer Town city | 0.71046261 | 0.74825175 | 42.22199681 | -72.2821209 |
| Plymouth County, Whitman town | 0.74825175 | 0.74825175 | 42.08048793 | -70.94731596 |
| Franklin County, Sunderland town | 0.59817106 | 0.74690694 | 42.46125434 | -72.57739387 |
| Berkshire County, Clarksburg town | 0.64631522 | 0.74610005 | 42.72642015 | -73.15033573 |
| Barnstable County, Yarmouth town | 0.57961269 | 0.74045186 | 41.66251823 | -70.25352475 |
| Franklin County, Deerfield town | 0.68719742 | 0.73883808 | 42.51553256 | -72.62183447 |
| Essex County, Rockport town | 0.71893491 | 0.73722431 | 42.64549143 | -70.64267959 |
| Berkshire County, Dalton town | 0.64954276 | 0.73614847 | 42.47681602 | -73.17104891 |
| Worcester County, Templeton town | 0.68195266 | 0.7288865 | 42.55192994 | -72.07202544 |
| Bristol County, Somerset town | 0.72861754 | 0.72861754 | 41.70941611 | -71.17691629 |
| Plymouth County, Carver town | 0.53711673 | 0.72054868 | 41.80632467 | -70.73370227 |
| Hampshire County, Ware town | 0.57315761 | 0.71947283 | 42.24365917 | -72.27458878 |
| Worcester County, Oakham town | 0.60718128 | 0.71301775 | 42.33473031 | -72.02220166 |
| Middlesex County, Everett city | 0.70333513 | 0.70333513 | 42.40180539 | -71.05033966 |
| Berkshire County, Richmond town | 0.58714363 | 0.69822485 | 42.41113553 | -73.31890501 |
| Berkshire County, Hinsdale town | 0.577461 | 0.69741797 | 42.44977402 | -73.14861817 |
| Worcester County, Rutland town | 0.58472297 | 0.69176977 | 42.32914546 | -71.99263548 |
| Hampden County, Brimfield town | 0.57907477 | 0.68719742 | 42.10214435 | -72.13524403 |
| Essex County, Essex town | 0.64537386 | 0.68396988 | 42.60427734 | -70.78520465 |
| Suffolk County, Chelsea city | 0.68047337 | 0.68047337 | 42.39599893 | -71.02082462 |
| Worcester County, New Braintree town | 0.48695535 | 0.67455621 | 42.31865644 | -72.05938451 |
| Worcester County, Winchendon town | 0.51089295 | 0.67186659 | 42.61690912 | -72.0491104 |
| Worcester County, Athol town | 0.58633674 | 0.66917698 | 42.57416325 | -72.19072402 |
| Franklin County, Montague town | 0.57288865 | 0.66810113 | 42.59683823 | -72.56224935 |
| Berkshire County, Lee town | 0.6457773 | 0.66675632 | 42.30836407 | -73.25879328 |
| Worcester County, Douglas town | 0.60032275 | 0.66487359 | 42.05100309 | -71.744739 |
| Franklin County, Gill town | 0.5866057 | 0.66460463 | 42.64021714 | -72.54727416 |
| Worcester County, Warren town | 0.60180204 | 0.66272189 | 42.23259909 | -72.21549812 |
| Worcester County, Brookfield town | 0.58512641 | 0.66245293 | 42.16709651 | -72.11253563 |
| Berkshire County, Lanesborough town | 0.55244755 | 0.66003228 | 42.50385514 | -73.19349713 |
| Worcester County, West Brookfield town | 0.53281334 | 0.65949435 | 42.25425787 | -72.20794413 |
| Berkshire County, Lenox town | 0.56764389 | 0.65761162 | 42.40584604 | -73.28920485 |
| Worcester County, Uxbridge town | 0.65330823 | 0.65330823 | 42.06132683 | -71.67819896 |
| Essex County, Manchester-by-the-Sea town | 0.62560516 | 0.65169446 | 42.58269385 | -70.79324038 |
| Worcester County, East Brookfield town | 0.53967187 | 0.65115654 | 42.17761089 | -72.04593967 |
| Berkshire County, Cheshire town | 0.51990317 | 0.65061861 | 42.5255775 | -73.18621929 |
| Franklin County, Shutesbury town | 0.46584185 | 0.64523938 | 42.41769865 | -72.46626102 |
| Berkshire County, Hancock town | 0.5232652 | 0.6401291 | 42.43814671 | -73.34138951 |
| Dukes County, Tisbury town | 0.63528779 | 0.63636364 | 41.43599709 | -70.61982011 |
| Dukes County, West Tisbury town | 0.55540613 | 0.63340506 | 41.44188818 | -70.64889963 |
| Dukes County, Oak Bluffs town | 0.61646046 | 0.6253362 | 41.43009945 | -70.59074545 |
| Worcester County, North Brookfield town | 0.55675094 | 0.62291555 | 42.29700457 | -72.0669917 |
| Hampshire County, Williamsburg town | 0.55406132 | 0.61995697 | 42.3908266 | -72.6961274 |
| Worcester County, Phillipston town | 0.53926842 | 0.61995697 | 42.58472337 | -72.12374895 |
| Franklin County, Shelburne town | 0.51896181 | 0.61888112 | 42.59149409 | -72.65889484 |
| Berkshire County, Washington town | 0.45992469 | 0.61780527 | 42.41165648 | -73.19286768 |
| Berkshire County, Stockbridge town | 0.51936525 | 0.61753631 | 42.27021611 | -73.30289262 |
| Franklin County, Conway town | 0.51344809 | 0.61081226 | 42.52641128 | -72.68123019 |
| Berkshire County, Great Barrington town | 0.54679935 | 0.59655729 | 42.21033205 | -73.35414077 |
| Hampden County, Holland town | 0.55459925 | 0.59655729 | 42.08049388 | -72.14280411 |
| Franklin County, Buckland town | 0.49488973 | 0.59413663 | 42.61319293 | -72.77784665 |
| Franklin County, Orange town | 0.50470683 | 0.59171598 | 42.62858164 | -72.23493386 |
| Dukes County, Edgartown town | 0.50443787 | 0.59090909 | 41.3870161 | -70.60666329 |
| Franklin County, Charlemont town | 0.46584185 | 0.59010221 | 42.62947866 | -72.74067346 |
| Hampshire County, Goshen town | 0.57073696 | 0.59010221 | 42.42877749 | -72.77769024 |
| Dukes County, Chilmark town | 0.52501345 | 0.57853685 | 41.38313032 | -70.70177945 |
| Berkshire County, Florida town | 0.45831092 | 0.57799892 | 42.66162017 | -73.0457594 |
| Franklin County, Leverett town | 0.49731038 | 0.56750941 | 42.44485655 | -72.4884153 |
| Franklin County, Wendell town | 0.49704142 | 0.56643357 | 42.59662491 | -72.43587344 |
| Hampshire County, Chesterfield town | 0.49973104 | 0.56293706 | 42.42327483 | -72.87408514 |
| Hampshire County, Northampton city | 0.55728886 | 0.55728886 | 42.35286648 | -72.74058687 |
| Berkshire County, Sheffield town | 0.49677246 | 0.55567509 | 42.13460814 | -73.31652717 |
| Worcester County, Royalston town | 0.50941366 | 0.55164067 | 42.64471252 | -72.19762548 |
| Berkshire County, Savoy town | 0.37358795 | 0.55137171 | 42.61818327 | -73.06042709 |
| Barnstable County, Dennis town | 0.49327595 | 0.54975793 | 41.71511254 | -70.1709021 |
| Hampshire County, Middlefield town | 0.48090371 | 0.54841313 | 42.3362287 | -73.02941896 |
| Hampden County, Blandford town | 0.46637977 | 0.5478752 | 42.20626568 | -72.9475248 |
| Worcester County, Hubbardston town | 0.46718666 | 0.54706832 | 42.49746213 | -72.02799214 |
| Barnstable County, Harwich town | 0.41594944 | 0.54679935 | 41.70310385 | -70.11260259 |
| Hampshire County, Cummington town | 0.4943518 | 0.54626143 | 42.47206719 | -72.88903529 |
| Berkshire County, New Ashford town | 0.46342119 | 0.54384077 | 42.61779051 | -73.18684094 |
| Franklin County, Leyden town | 0.3548951 | 0.54276493 | 42.65113176 | -72.60676505 |
| Berkshire County, Tyringham town | 0.4464766 | 0.53765465 | 42.27603574 | -73.2067442 |
| Hampden County, Wales town | 0.52232383 | 0.53711673 | 42.0809917 | -72.26824482 |
| Hampden County, Granville town | 0.44782141 | 0.53657881 | 42.04900961 | -72.96913701 |
| Hampden County, Russell town | 0.44620764 | 0.53496503 | 42.16843232 | -72.86612984 |
| Franklin County, Ashfield town | 0.45790748 | 0.53415815 | 42.5155655 | -72.74806292 |
| Hampden County, Monson town | 0.52582033 | 0.53254438 | 42.04319313 | -72.31273916 |
| Franklin County, Erving town | 0.46355568 | 0.53200646 | 42.58042698 | -72.47310042 |
| Worcester County, Barre town | 0.46422808 | 0.53012372 | 42.38918826 | -72.06612738 |
| Worcester County, Hardwick town | 0.45938677 | 0.52662722 | 42.38421385 | -72.16252138 |
| Franklin County, Northfield town | 0.44271114 | 0.52420656 | 42.62379462 | -72.458073 |
| Berkshire County, West Stockbridge town | 0.4493007 | 0.52232383 | 42.35123426 | -73.37024319 |
| Berkshire County, Otis town | 0.47377622 | 0.52178591 | 42.19494068 | -73.1396564 |
| Franklin County, Bernardston town | 0.38972566 | 0.52097902 | 42.66190694 | -72.53977953 |
| Hampshire County, Worthington town | 0.47686928 | 0.51882733 | 42.40695566 | -72.9111168 |
| Berkshire County, Becket town | 0.4556213 | 0.51748252 | 42.31986717 | -73.06636783 |
| Berkshire County, Egremont town | 0.48399677 | 0.51640667 | 42.19914561 | -73.42056403 |
| Berkshire County, Windsor town | 0.40411512 | 0.51640667 | 42.48787842 | -73.10432009 |
| Middlesex County, Pepperell town | 0.51533082 | 0.51533082 | 42.68960229 | -71.57942218 |
| Berkshire County, Monterey town | 0.44513179 | 0.51371705 | 42.17275426 | -73.27252105 |
| Hampden County, Tolland town | 0.42711135 | 0.51291017 | 42.05436985 | -72.99866108 |
| Franklin County, New Salem town | 0.44822485 | 0.51075847 | 42.54742262 | -72.29492946 |
| Franklin County, Warwick town | 0.47458311 | 0.51022055 | 42.69402679 | -72.33869837 |
| Hampden County, Chester town | 0.45965573 | 0.50806885 | 42.249666 | -72.93287347 |
| Berkshire County, Sandisfield town | 0.44055944 | 0.50430339 | 42.17323028 | -73.14691423 |
| Hampshire County, Huntington town | 0.41984938 | 0.50242066 | 42.24428944 | -72.9032717 |
| Berkshire County, Alford town | 0.45481442 | 0.50107585 | 42.24259851 | -73.40620907 |
| Hampshire County, Westhampton town | 0.42832168 | 0.49327595 | 42.35828093 | -72.77022223 |
| Hampshire County, Plainfield town | 0.457773 | 0.49246907 | 42.49377146 | -72.88166559 |
| Berkshire County, Peru town | 0.45293168 | 0.49193115 | 42.41205515 | -73.06682868 |
| Hampden County, Southwick town | 0.48897257 | 0.48897257 | 42.0708815 | -72.8364124 |
| Franklin County, Heath town | 0.42899408 | 0.4873588 | 42.66200907 | -72.79276999 |
| Berkshire County, New Marlborough town | 0.40371167 | 0.48628295 | 42.12931998 | -73.28694488 |
| Franklin County, Colrain town | 0.3655191 | 0.47848306 | 42.65117921 | -72.73324054 |
| Worcester County, Petersham town | 0.35785368 | 0.47364174 | 42.5308296 | -72.20590768 |
| Hampden County, Westfield city | 0.46664874 | 0.46664874 | 42.11965631 | -72.85126073 |
| Dukes County, Gosnold town | 0.4190425 | 0.46046261 | 41.46538693 | -70.76526634 |
| Barnstable County, Orleans town | 0.30836471 | 0.44862829 | 41.79259981 | -69.98429699 |
| Dukes County, Aquinnah town | 0.43948359 | 0.44782141 | 41.33606724 | -70.81266538 |
| Berkshire County, Mount Washington town | 0.42092523 | 0.44190425 | 42.1122421 | -73.44922034 |
| Barnstable County, Brewster town | 0.40371167 | 0.44136633 | 41.73063109 | -70.13361408 |
| Barnstable County, Chatham town | 0.31845078 | 0.43598709 | 41.68504163 | -70.02519079 |
| Franklin County, Rowe town | 0.36820871 | 0.4338354 | 42.67820318 | -72.88209901 |
| Franklin County, Hawley town | 0.38084992 | 0.41097364 | 42.55888562 | -72.85952881 |
| Nantucket County, Nantucket town | 0.35341581 | 0.40693921 | 41.27980884 | -70.08445129 |
| Barnstable County, Provincetown town | 0.35664336 | 0.40640129 | 42.06199451 | -70.16561947 |
| Franklin County, Monroe town | 0.29558903 | 0.34588488 | 42.72141815 | -72.99391047 |
| Barnstable County, Eastham town | 0.2880581 | 0.32786444 | 41.81411139 | -69.97610316 |
| Barnstable County, Truro town | 0.29115116 | 0.30016138 | 42.04997866 | -70.10703127 |
| Barnstable County, Wellfleet town | 0.23977945 | 0.29989242 | 41.93979734 | -70.02274272 |
