## Supplementary material for "Mapping long-distance jump dispersal risk by leveraging transport infrastructure data and sampling bias correction: a case study with the invasive spotted lanternfly (*Lycorma delicatula*)": S8 Michigan risk

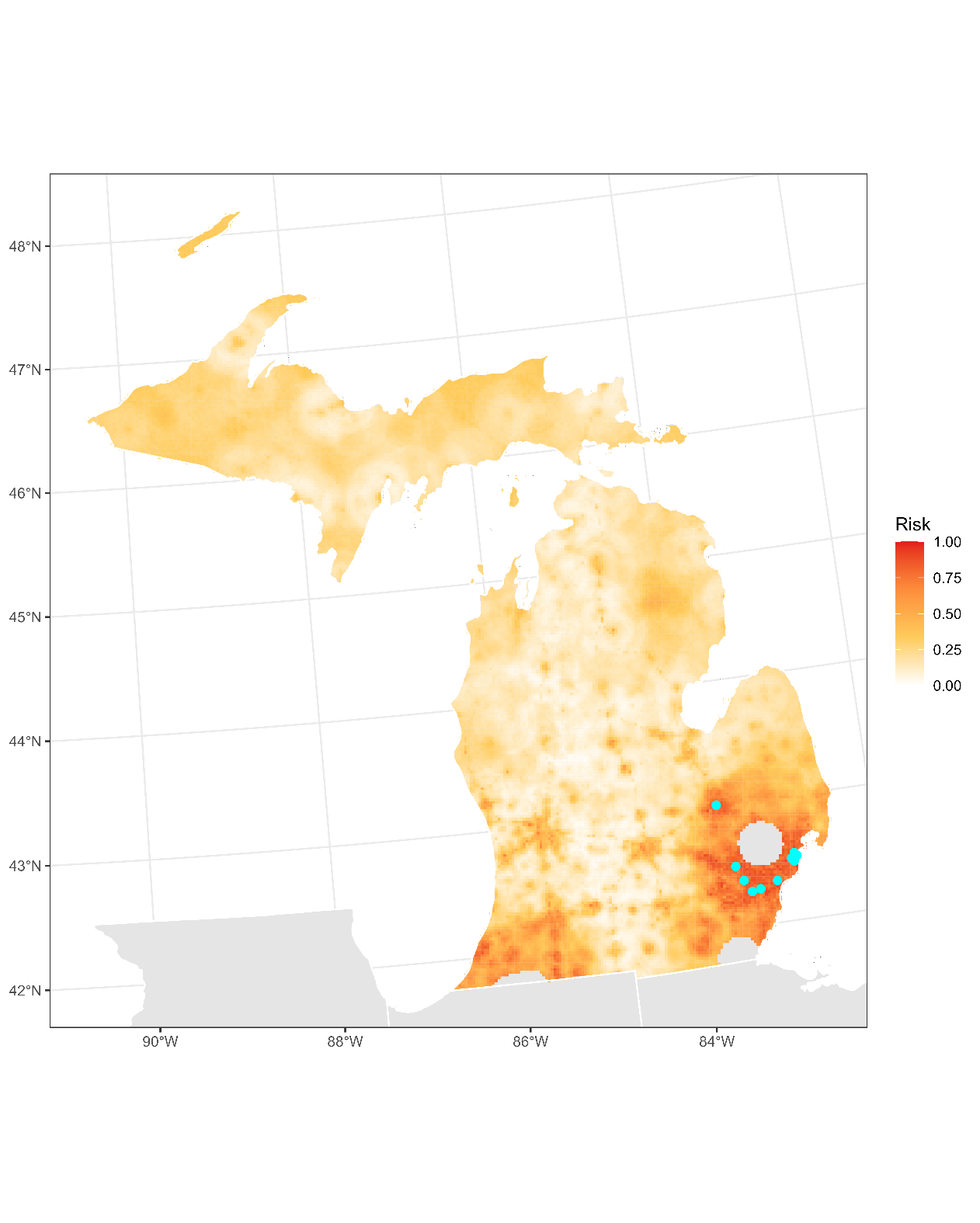

**Figure S8.1:** Projected risk of colonization by *L. delicatula* jump dispersal for Michigan. Risk projections are based on an ensemble random forest model including distances from existing *L. delicatula* populations. Note that areas falling less than 20 km from existing *L. delicatula* populations are not included in the map because any colonization there would be categorized as diffusive spread based on our methodology (regardless of the actual source of the resulting population). Darker, redder colors indicate increased risk of colonization by jump dispersal. Cyan points show the top ten highest risk locations in the state.

**Table S8.1:** Ranked list of Michigan county subdivisions sorted by the maximum projected risk of colonization by jump dispersal occurring there. For each subdivision, we identified the 2.5 x 2.5 km grid cell centroid included there with the highest projected risk of colonization by jump dispersal and report that location and risk value.

| **County, subdivision** | **Median risk value** | **Maximum risk value** | **Latitude of maximum risk** | **Longitude of maximum risk** |
| --- | --- | --- | --- | --- |
| Oakland County, Lyon charter township | 0.83889188 | 0.94029048 | 42.4917323 | -83.57159183 |
| Macomb County, Clinton charter township | 0.82046799 | 0.93006993 | 42.53968985 | -82.9139732 |
| Wayne County, Inkster city | 0.86767079 | 0.92173211 | 42.28791624 | -83.33113822 |
| Genesee County, Flint city | 0.74273803 | 0.91742873 | 43.00004859 | -83.69710897 |
| Wayne County, Wayne city | 0.87305003 | 0.91608392 | 42.274638 | -83.42590828 |
| Wayne County, Detroit city | 0.81643357 | 0.91447015 | 42.33641898 | -83.137371 |
| Macomb County, Roseville city | 0.84709521 | 0.91393222 | 42.49845 | -82.95269346 |
| Macomb County, St. Clair Shores city | 0.84144701 | 0.91204949 | 42.51446531 | -82.88785084 |
| Wayne County, Plymouth charter township | 0.80446477 | 0.90667025 | 42.37211473 | -83.50064007 |
| Macomb County, Eastpointe city | 0.81065089 | 0.90613233 | 42.47323466 | -82.92657256 |
| Oakland County, Ferndale city | 0.8485745 | 0.90344271 | 42.44719376 | -83.11673607 |
| Wayne County, Redford charter township | 0.86767079 | 0.90263583 | 42.39573433 | -83.28053635 |
| Oakland County, Novi city | 0.81710597 | 0.90155998 | 42.48588563 | -83.51091084 |
| Washtenaw County, Northfield township | 0.73668639 | 0.89967725 | 42.35100047 | -83.75088104 |
| Wayne County, Taylor city | 0.80957504 | 0.89913932 | 42.25681484 | -83.24450058 |
| Wayne County, Grosse Pointe Woods city | 0.89537386 | 0.8986014 | 42.42586584 | -82.90467184 |
| Wayne County, Northville township | 0.8267886 | 0.89564282 | 42.39721091 | -83.52693066 |
| Macomb County, Center Line city | 0.89268424 | 0.89268424 | 42.48240274 | -83.01750615 |
| Livingston County, Brighton township | 0.71247983 | 0.89080151 | 42.55633178 | -83.77620162 |
| Wayne County, Melvindale city | 0.86565358 | 0.88757396 | 42.27297047 | -83.17995024 |
| Macomb County, Sterling Heights city | 0.8070199 | 0.88569123 | 42.57406973 | -83.03117011 |
| Wayne County, Livonia city | 0.85476062 | 0.88353954 | 42.41056503 | -83.43201278 |
| Wayne County, Dearborn city | 0.84911243 | 0.88327058 | 42.33943237 | -83.16762913 |
| Livingston County, Genoa township | 0.80863367 | 0.88138784 | 42.56205517 | -83.8369663 |
| Livingston County, Oceola township | 0.65061861 | 0.880312 | 42.61211932 | -83.88995374 |
| Wayne County, Van Buren charter township | 0.8183163 | 0.87842926 | 42.21696469 | -83.52862511 |
| Macomb County, Shelby charter township | 0.83055406 | 0.87762238 | 42.66270354 | -83.01447493 |
| Genesee County, Flint charter township | 0.79800968 | 0.87681549 | 42.96429502 | -83.79673353 |
| Wayne County, Allen Park city | 0.8211404 | 0.87573964 | 42.2759737 | -83.21018264 |
| Wayne County, Dearborn Heights city | 0.84104357 | 0.87009145 | 42.3292569 | -83.29275959 |
| Washtenaw County, Ypsilanti charter township | 0.77824099 | 0.86982249 | 42.20644303 | -83.65348182 |
| Oakland County, Southfield city | 0.77353416 | 0.86632598 | 42.46221643 | -83.26828973 |
| Wayne County, Canton charter township | 0.81091985 | 0.86417429 | 42.34407215 | -83.44409387 |
| St. Joseph County, Constantine township | 0.55271651 | 0.86202259 | 41.84408008 | -85.67213195 |
| Wayne County, River Rouge city | 0.85745024 | 0.86202259 | 42.2669429 | -83.11949328 |
| Oakland County, Farmington Hills city | 0.8154922 | 0.8598709 | 42.47706265 | -83.41990856 |
| Oakland County, Walled Lake city | 0.85960194 | 0.85960194 | 42.52729024 | -83.47252987 |
| Wayne County, Westland city | 0.81495428 | 0.85718128 | 42.33816719 | -83.38355243 |
| Wayne County, Garden City city | 0.85018827 | 0.85664336 | 42.31304395 | -83.35733587 |
| Oakland County, Northville city | 0.85556751 | 0.85556751 | 42.43861532 | -83.48860821 |
| Macomb County, Warren city | 0.79424422 | 0.8542227 | 42.48544466 | -83.04782267 |
| Washtenaw County, Pittsfield charter township | 0.76627219 | 0.85233997 | 42.21222112 | -83.71393142 |
| Wayne County, Hamtramck city | 0.85018827 | 0.85018827 | 42.39682807 | -83.06443279 |
| Oakland County, Pleasant Ridge city | 0.84803658 | 0.84803658 | 42.47236998 | -83.14291616 |
| Macomb County, Macomb township | 0.78644432 | 0.84642281 | 42.63137491 | -82.9275115 |
| Genesee County, Burton city | 0.79074771 | 0.84373319 | 42.99425612 | -83.63597009 |
| Washtenaw County, Ann Arbor charter township | 0.75726197 | 0.84319527 | 42.32306772 | -83.69425132 |
| Oakland County, Commerce charter township | 0.80217859 | 0.84131253 | 42.59967323 | -83.52121531 |
| Washtenaw County, Scio township | 0.71180742 | 0.84077461 | 42.29591066 | -83.88370991 |
| Berrien County, Royalton township | 0.66917698 | 0.8394298 | 42.03812377 | -86.43823784 |
| Livingston County, Green Oak township | 0.74636902 | 0.83916084 | 42.50910263 | -83.7536958 |
| Genesee County, Grand Blanc charter township | 0.7175901 | 0.83647122 | 42.94697272 | -83.6134175 |
| Oakland County, Milford charter township | 0.7943787 | 0.83647122 | 42.53607698 | -83.56360467 |
| Berrien County, Benton charter township | 0.7027972 | 0.83566434 | 42.08034847 | -86.40185568 |
| Wayne County, Romulus city | 0.74852071 | 0.83055406 | 42.22736616 | -83.40373091 |
| Monroe County, Frenchtown township | 0.7084454 | 0.82840237 | 41.9969362 | -83.35352711 |
| Macomb County, Chesterfield township | 0.76035503 | 0.82517483 | 42.66644815 | -82.82792185 |
| Macomb County, Washington charter township | 0.69217321 | 0.82275417 | 42.73223139 | -83.03234943 |
| Oakland County, Wixom city | 0.81078537 | 0.81629909 | 42.51098354 | -83.53724823 |
| Livingston County, Brighton city | 0.80123722 | 0.81603012 | 42.51197293 | -83.7840553 |
| Macomb County, Harrison charter township | 0.75578268 | 0.8154922 | 42.59998966 | -82.84062672 |
| Wayne County, Lincoln Park city | 0.80446477 | 0.81495428 | 42.25081749 | -83.18405205 |
| St. Joseph County, Fabius township | 0.59978483 | 0.8119957 | 41.95515128 | -85.65566209 |
| Livingston County, Marion township | 0.54653039 | 0.81038193 | 42.59560321 | -83.95465238 |
| Oakland County, White Lake charter township | 0.69002152 | 0.80742335 | 42.62184704 | -83.51720068 |
| Wayne County, Grosse Pointe city | 0.80580958 | 0.80580958 | 42.38156921 | -82.91306627 |
| Oakland County, Madison Heights city | 0.80527165 | 0.80527165 | 42.49150733 | -83.10846367 |
| Washtenaw County, Ann Arbor city | 0.74139322 | 0.80446477 | 42.30665528 | -83.75872768 |
| Macomb County, Fraser city | 0.80338892 | 0.80338892 | 42.54275614 | -82.9443102 |
| Macomb County, Mount Clemens city | 0.80311996 | 0.80311996 | 42.6030733 | -82.87098685 |
| Oakland County, Huntington Woods city | 0.80150619 | 0.80150619 | 42.47538262 | -83.17323376 |
| Livingston County, Howell city | 0.77057558 | 0.80096826 | 42.5927724 | -83.92424926 |
| Wayne County, Riverview city | 0.71785906 | 0.80016138 | 42.1813554 | -83.16614977 |
| Washtenaw County, York charter township | 0.64658419 | 0.79908553 | 42.09561878 | -83.67319377 |
| Oakland County, Highland charter township | 0.70185584 | 0.79881657 | 42.6335083 | -83.63881202 |
| Livingston County, Hamburg township | 0.59467456 | 0.79774072 | 42.49265657 | -83.81833084 |
| Oakland County, Hazel Park city | 0.7972028 | 0.7972028 | 42.46935028 | -83.11260119 |
| Genesee County, Mundy township | 0.70252824 | 0.7950511 | 42.95854941 | -83.73561797 |
| Wayne County, Grosse Pointe Farms city | 0.79451318 | 0.79451318 | 42.40371727 | -82.90887039 |
| Wayne County, Trenton city | 0.7436794 | 0.79424422 | 42.15920541 | -83.17025058 |
| Berrien County, Lincoln charter township | 0.76385153 | 0.7915546 | 42.02036138 | -86.50177969 |
| Washtenaw County, Superior charter township | 0.71570737 | 0.79074771 | 42.2642145 | -83.55088392 |
| St. Joseph County, Mottville township | 0.60543303 | 0.78913394 | 41.7996549 | -85.67870527 |
| Monroe County, Monroe city | 0.74220011 | 0.78832706 | 41.93048462 | -83.36561135 |
| Wayne County, Woodhaven city | 0.77461001 | 0.78509952 | 42.12090581 | -83.2387747 |
| Washtenaw County, Salem township | 0.68934911 | 0.78402367 | 42.40013369 | -83.55723186 |
| Lenawee County, Raisin township | 0.61081226 | 0.77972028 | 41.92182036 | -83.97943306 |
| Wayne County, Sumpter township | 0.60677784 | 0.77837547 | 42.16971386 | -83.50639647 |
| St. Joseph County, White Pigeon township | 0.59117805 | 0.77649274 | 41.79722945 | -85.6485826 |
| Monroe County, Ash township | 0.68464228 | 0.77541689 | 42.01611782 | -83.31936669 |
| Monroe County, Berlin charter township | 0.63797741 | 0.77030662 | 41.99396742 | -83.32340926 |
| Oakland County, Oxford charter township | 0.71005917 | 0.77030662 | 42.82585023 | -83.29399822 |
| Macomb County, Lenox township | 0.6640667 | 0.76869285 | 42.73600181 | -82.84560989 |
| Wayne County, Highland Park city | 0.76815492 | 0.76815492 | 42.39985871 | -83.09471404 |
| Berrien County, Sodus township | 0.63017751 | 0.76788596 | 42.06036088 | -86.43518275 |
| Wayne County, Southgate city | 0.74744486 | 0.76734804 | 42.20651302 | -83.19224786 |
| Genesee County, Grand Blanc city | 0.76681011 | 0.76681011 | 42.92478543 | -83.61742013 |
| Oakland County, South Lyon city | 0.76627219 | 0.76627219 | 42.45320412 | -83.64022193 |
| Cass County, Pokagon township | 0.6366326 | 0.7654653 | 41.9708639 | -86.14211126 |
| St. Joseph County, Sturgis township | 0.62775686 | 0.76465842 | 41.80473561 | -85.46453895 |
| Genesee County, Swartz Creek city | 0.76035503 | 0.76035503 | 42.96715714 | -83.82729512 |
| St. Joseph County, Florence township | 0.52286175 | 0.7584723 | 41.84165297 | -85.64199039 |
| Wayne County, Ecorse city | 0.74556213 | 0.75551372 | 42.24780827 | -83.1538317 |
| Wayne County, Flat Rock city | 0.70817644 | 0.75497579 | 42.09875516 | -83.24284891 |
| Lenawee County, Madison charter township | 0.68396988 | 0.75470683 | 41.86093578 | -84.05108407 |
| Monroe County, Monroe charter township | 0.69472835 | 0.75174825 | 41.88914247 | -83.37365476 |
| Wayne County, Brownstown charter township | 0.6873319 | 0.74959656 | 42.16520852 | -83.23061851 |
| Oakland County, Springfield charter township | 0.65357719 | 0.7493276 | 42.75783408 | -83.52351524 |
| Genesee County, Davison township | 0.654922 | 0.74878967 | 42.985514 | -83.54428108 |
| Monroe County, Milan township | 0.49811727 | 0.74852071 | 42.07345536 | -83.67712866 |
| St. Joseph County, Lockport township | 0.51291017 | 0.74825175 | 41.97493487 | -85.6221635 |
| Genesee County, Fenton charter township | 0.6457773 | 0.74825175 | 42.82253296 | -83.72886786 |
| Lenawee County, Adrian city | 0.7380312 | 0.74744486 | 41.88590452 | -84.07737949 |
| Livingston County, Howell township | 0.62829478 | 0.74636902 | 42.63997743 | -83.94689844 |
| Washtenaw County, Bridgewater township | 0.51802044 | 0.74583109 | 42.07983215 | -83.98274786 |
| Cass County, Wayne township | 0.6218397 | 0.74314147 | 42.0130017 | -86.10558539 |
| Genesee County, Mount Morris township | 0.6457773 | 0.74314147 | 43.05020188 | -83.75033153 |
| Genesee County, Genesee charter township | 0.64254976 | 0.74179666 | 43.08592484 | -83.65055871 |
| Berrien County, St. Joseph charter township | 0.70037655 | 0.74098978 | 42.08484145 | -86.46239772 |
| Lenawee County, Palmyra township | 0.54572351 | 0.73964497 | 41.89965227 | -83.98326119 |
| Wayne County, Grosse Ile township | 0.63878429 | 0.73964497 | 42.13705589 | -83.17434878 |
| Macomb County, New Baltimore city | 0.70010758 | 0.73910705 | 42.70455725 | -82.75863102 |
| Washtenaw County, Webster township | 0.62103281 | 0.73910705 | 42.37890951 | -83.80755622 |
| Washtenaw County, Ypsilanti city | 0.68235611 | 0.73883808 | 42.25077616 | -83.64557948 |
| Berrien County, Bertrand township | 0.60731576 | 0.7380312 | 41.80450611 | -86.31792162 |
| Lenawee County, Adrian township | 0.61847768 | 0.73587951 | 41.92463593 | -84.00954695 |
| Kalamazoo County, Schoolcraft township | 0.5767886 | 0.73507262 | 42.15511046 | -85.62588412 |
| Washtenaw County, Augusta charter township | 0.62036041 | 0.73345885 | 42.15921428 | -83.63117197 |
| St. Clair County, Casco township | 0.47337278 | 0.73345885 | 42.74575113 | -82.71967986 |
| Wayne County, Huron charter township | 0.62641205 | 0.73292093 | 42.15793661 | -83.38561592 |
| Washtenaw County, Lodi township | 0.46395912 | 0.73130715 | 42.19580123 | -83.77830063 |
| Genesee County, Atlas township | 0.62641205 | 0.73103819 | 42.94114167 | -83.55233268 |
| Cass County, Dowagiac city | 0.72646584 | 0.73023131 | 41.99309213 | -86.13896493 |
| Berrien County, Oronoko charter township | 0.56522324 | 0.72646584 | 41.94242314 | -86.35980589 |
| Lenawee County, Clinton township | 0.65976331 | 0.72431415 | 42.07700981 | -83.95256811 |
| St. Joseph County, Three Rivers city | 0.71893491 | 0.72350726 | 41.93050565 | -85.62878086 |
| Monroe County, London township | 0.55244755 | 0.72108661 | 42.07056176 | -83.64696419 |
| Genesee County, Clayton charter township | 0.49515869 | 0.72081764 | 42.97001214 | -83.85785924 |
| Washtenaw County, Lima township | 0.45467994 | 0.71947283 | 42.3209266 | -83.9101096 |
| St. Joseph County, Sturgis city | 0.68491124 | 0.71812803 | 41.8022601 | -85.43442158 |
| Macomb County, Richmond city | 0.65532544 | 0.71624529 | 42.79317223 | -82.7415551 |
| Lapeer County, Metamora township | 0.47686928 | 0.71113502 | 42.895356 | -83.31219628 |
| Macomb County, Ray township | 0.62802582 | 0.70871436 | 42.72000254 | -82.9106698 |
| Wayne County, Grosse Pointe Park city | 0.7084454 | 0.7084454 | 42.38463507 | -82.94333425 |
| St. Joseph County, Fawn River township | 0.53012372 | 0.70521786 | 41.77757274 | -85.40767684 |
| Cass County, Silver Creek township | 0.62157074 | 0.7049489 | 41.99540343 | -86.16918862 |
| Wayne County, Rockwood city | 0.70467994 | 0.70467994 | 42.07660498 | -83.24692054 |
| Kalamazoo County, Texas charter township | 0.54895105 | 0.70414201 | 42.1575422 | -85.65616061 |
| Wayne County, Plymouth city | 0.70360409 | 0.70360409 | 42.36917885 | -83.47035346 |
| Van Buren County, Antwerp township | 0.64066703 | 0.70360409 | 42.19421023 | -85.83461118 |
| Kalamazoo County, Prairie Ronde township | 0.44809037 | 0.70225928 | 42.13532164 | -85.65946818 |
| Berrien County, Benton Harbor city | 0.65519096 | 0.70118343 | 42.12707531 | -86.42600582 |
| Washtenaw County, Manchester township | 0.48251748 | 0.69956966 | 42.08264746 | -84.01293004 |
| Cass County, Howard township | 0.59628833 | 0.69849381 | 41.83981058 | -86.19110462 |
| Oakland County, Oak Park city | 0.69795589 | 0.69795589 | 42.453224 | -83.17734967 |
| St. Joseph County, Sherman township | 0.52232383 | 0.69634212 | 41.82446622 | -85.43105839 |
| Washtenaw County, Dexter city | 0.69634212 | 0.69634212 | 42.34026275 | -83.87594414 |
| Kent County, Walker city | 0.39806347 | 0.69526627 | 43.02180547 | -85.77411424 |
| Oakland County, Brandon charter township | 0.56952663 | 0.69472835 | 42.81554557 | -83.41999035 |
| Berrien County, Baroda township | 0.57369554 | 0.69311458 | 41.97588666 | -86.50784205 |
| Ottawa County, Holland charter township | 0.37681549 | 0.69230769 | 42.79834768 | -86.08549051 |
| Monroe County, La Salle township | 0.67563206 | 0.68961807 | 41.87288308 | -83.46787303 |
| Oakland County, Groveland township | 0.5204411 | 0.6880043 | 42.79631699 | -83.45451784 |
| Monroe County, Dundee township | 0.55137171 | 0.68746638 | 41.95975662 | -83.66664818 |
| Kalamazoo County, Portage city | 0.61027434 | 0.68531469 | 42.22420705 | -85.64622525 |
| Livingston County, Hartland township | 0.62856374 | 0.68504572 | 42.6257425 | -83.79481202 |
| Wayne County, Gibraltar city | 0.68316299 | 0.68316299 | 42.09576022 | -83.21269185 |
| Genesee County, Fenton city | 0.61753631 | 0.68262507 | 42.80034628 | -83.73282626 |
| Oakland County, Rose township | 0.58082302 | 0.67966649 | 42.70004547 | -83.62686749 |
| Berrien County, Berrien township | 0.58848843 | 0.67939753 | 41.94015657 | -86.32959885 |
| Calhoun County, Battle Creek city | 0.35798817 | 0.6788596 | 42.32275516 | -85.2014186 |
| Wayne County, Wyandotte city | 0.66272189 | 0.6788596 | 42.20350587 | -83.16204636 |
| St. Joseph County, Burr Oak township | 0.48318989 | 0.67805272 | 41.82198284 | -85.40093373 |
| Kalamazoo County, Kalamazoo city | 0.61175363 | 0.67805272 | 42.26377394 | -85.57894481 |
| Kent County, Grand Rapids city | 0.41984938 | 0.67482517 | 42.94782179 | -85.69215427 |
| Cass County, LaGrange township | 0.54922001 | 0.67348037 | 41.9662217 | -86.08168889 |
| Lenawee County, Tecumseh township | 0.65761162 | 0.67240452 | 42.00483732 | -83.90380062 |
| Macomb County, Bruce township | 0.57934373 | 0.66863905 | 42.82088541 | -83.01561757 |
| St. Clair County, Columbus township | 0.40048413 | 0.66863905 | 42.81221165 | -82.70683195 |
| Berrien County, Lake charter township | 0.53496503 | 0.66648736 | 41.97811161 | -86.53807026 |
| Berrien County, Watervliet township | 0.49246907 | 0.6640667 | 42.18016467 | -86.23493387 |
| Livingston County, Tyrone township | 0.57261969 | 0.66245293 | 42.78103553 | -83.76725901 |
| Kent County, Byron township | 0.23641743 | 0.66110812 | 42.83417866 | -85.67822617 |
| Oakland County, Holly township | 0.5831092 | 0.65976331 | 42.81675691 | -83.66788324 |
| Berrien County, Niles township | 0.56508876 | 0.65949435 | 41.82673438 | -86.31483855 |
| Kalamazoo County, Oshtemo charter township | 0.51008607 | 0.65922539 | 42.29330644 | -85.66660677 |
| Monroe County, Exeter township | 0.56576116 | 0.6570737 | 42.00580049 | -83.44389599 |
| Genesee County, Richfield township | 0.57235073 | 0.6570737 | 43.05207727 | -83.53218425 |
| Berrien County, Buchanan township | 0.55150619 | 0.65545992 | 41.83352244 | -86.40531738 |
| Berrien County, New Buffalo township | 0.51533082 | 0.65465304 | 41.78897785 | -86.71591263 |
| Berrien County, St. Joseph city | 0.65438408 | 0.65438408 | 42.08707732 | -86.49267165 |
| Berrien County, Niles city | 0.65169446 | 0.65169446 | 41.84440095 | -86.25142512 |
| Genesee County, Mount Morris city | 0.65169446 | 0.65169446 | 43.11390722 | -83.70779824 |
| Genesee County, Gaines township | 0.40909091 | 0.65115654 | 42.94496268 | -83.83123017 |
| Lenawee County, Rome township | 0.47579344 | 0.65088757 | 41.91365516 | -84.13380781 |
| Monroe County, Raisinville township | 0.58418505 | 0.65034965 | 41.89797059 | -83.49396438 |
| Macomb County, Richmond township | 0.54451318 | 0.6492738 | 42.81532742 | -82.73727928 |
| Van Buren County, Keeler township | 0.54316837 | 0.64604626 | 42.13109731 | -86.18060853 |
| Lenawee County, Dover township | 0.45602474 | 0.64120495 | 41.89148272 | -84.13758659 |
| Berrien County, Chikaming township | 0.55446477 | 0.63878429 | 41.84690819 | -86.58632727 |
| Washtenaw County, Saline city | 0.63555675 | 0.63555675 | 42.17363192 | -83.78220775 |
| Ingham County, Lansing city | 0.5112964 | 0.63555675 | 42.742244 | -84.57897067 |
| Berrien County, Buchanan city | 0.63421194 | 0.63421194 | 41.8312668 | -86.37515582 |
| Lapeer County, Attica township | 0.58176439 | 0.63394298 | 43.01341322 | -83.13476042 |
| Van Buren County, Paw Paw township | 0.52797203 | 0.63313609 | 42.20133813 | -85.92551938 |
| Oakland County, Addison township | 0.54868209 | 0.63313609 | 42.81386374 | -83.17210782 |
| Berrien County, Bainbridge township | 0.47417967 | 0.63152232 | 42.09806299 | -86.33823384 |
| Genesee County, Vienna charter township | 0.38690156 | 0.63044648 | 43.13610381 | -83.70381374 |
| Jackson County, Jackson city | 0.50215169 | 0.62802582 | 42.22954186 | -84.38696449 |
| Lenawee County, Macon township | 0.53079613 | 0.62802582 | 42.0463322 | -83.86591699 |
| Cass County, Niles city | 0.62775686 | 0.62775686 | 41.86433617 | -86.21814846 |
| Lenawee County, Franklin township | 0.46369016 | 0.62721893 | 42.0604742 | -84.01675634 |
| St. Joseph County, Park township | 0.43652501 | 0.62587413 | 42.06623466 | -85.63913983 |
| Kent County, Wyoming city | 0.45266272 | 0.62452932 | 42.85642077 | -85.67489118 |
| St. Clair County, Ira township | 0.42711135 | 0.62426036 | 42.69832962 | -82.69783457 |
| Genesee County, Forest township | 0.49556213 | 0.62264658 | 43.14376583 | -83.54664692 |
| Monroe County, Milan city | 0.6218397 | 0.6218397 | 42.07634195 | -83.70729562 |
| Kalamazoo County, Pavilion township | 0.41231845 | 0.62076385 | 42.19220873 | -85.52836964 |
| Lenawee County, Tecumseh city | 0.61915008 | 0.62022593 | 42.00767081 | -83.93394723 |
| Macomb County, Armada township | 0.53711673 | 0.61699839 | 42.81477654 | -82.95469585 |
| St. Joseph County, Flowerfield township | 0.48090371 | 0.61323292 | 41.99958315 | -85.64905949 |
| Lapeer County, Imlay township | 0.50995159 | 0.61323292 | 43.03256341 | -83.10003522 |
| Muskegon County, Muskegon city | 0.50833782 | 0.61161915 | 43.23747809 | -86.23965697 |
| Washtenaw County, Saline township | 0.43773534 | 0.61081226 | 42.1514631 | -83.78611239 |
| Genesee County, Davison city | 0.60812265 | 0.60812265 | 43.0269585 | -83.50564197 |
| Isabella County, Union charter township | 0.37036041 | 0.60381926 | 43.61350138 | -84.74607603 |
| Washtenaw County, Dexter township | 0.46772458 | 0.60247445 | 42.34310389 | -83.90623498 |
| Livingston County, Handy township | 0.49166218 | 0.60247445 | 42.64843329 | -84.03818015 |
| Lapeer County, Lapeer township | 0.50188273 | 0.60086068 | 43.02543803 | -83.25699711 |
| St. Clair County, St. Clair township | 0.4873588 | 0.59682625 | 42.88512662 | -82.53727629 |
| Kalamazoo County, Kalamazoo charter township | 0.48655191 | 0.59601937 | 42.28354502 | -85.54527592 |
| St. Clair County, Kimball township | 0.54679935 | 0.59467456 | 42.95789372 | -82.58523063 |
| Berrien County, Three Oaks township | 0.54410974 | 0.59332975 | 41.82467301 | -86.58932507 |
| Van Buren County, Porter township | 0.45333513 | 0.59279182 | 42.15213998 | -85.87139555 |
| Berrien County, Weesaw township | 0.47767617 | 0.58714363 | 41.88916514 | -86.55013385 |
| Kent County, Cascade charter township | 0.38353954 | 0.5866057 | 42.91337889 | -85.54243568 |
| Eaton County, Windsor charter township | 0.28725121 | 0.58606778 | 42.68096311 | -84.65094159 |
| Lapeer County, Elba township | 0.49018289 | 0.58552985 | 43.01812464 | -83.41392615 |
| Gratiot County, Pine River township | 0.21032813 | 0.58526089 | 43.40271594 | -84.65587489 |
| Lapeer County, Hadley township | 0.47498655 | 0.58122647 | 42.88503266 | -83.43831244 |
| Cass County, Jefferson township | 0.56024744 | 0.58015062 | 41.85509589 | -86.09748545 |
| St. Clair County, Clay township | 0.51533082 | 0.57988166 | 42.64149859 | -82.5849104 |
| Van Buren County, Decatur township | 0.50874126 | 0.57961269 | 42.1545143 | -85.9016792 |
| Lapeer County, Imlay City city | 0.56670253 | 0.57961269 | 43.02953123 | -83.06947576 |
| Ottawa County, Georgetown charter township | 0.35449166 | 0.57638515 | 42.89070911 | -85.82455205 |
| Calhoun County, Emmett charter township | 0.33700914 | 0.57557827 | 42.31767092 | -85.14075661 |
| Berrien County, Hagar township | 0.46369016 | 0.57477138 | 42.1693011 | -86.38956984 |
| Calhoun County, Pennfield charter township | 0.10610543 | 0.57477138 | 42.339883 | -85.13727473 |
| Berrien County, Galien township | 0.46705218 | 0.57450242 | 41.80022651 | -86.56216682 |
| Kalamazoo County, Alamo township | 0.2950511 | 0.5739645 | 42.35998356 | -85.65666203 |
| Cass County, Calvin township | 0.53698225 | 0.57235073 | 41.88546329 | -85.91012489 |
| Livingston County, Putnam township | 0.4704142 | 0.56858526 | 42.4568373 | -83.9171656 |
| St. Clair County, Port Huron charter township | 0.52138246 | 0.56777838 | 42.97056892 | -82.48936406 |
| Lapeer County, Arcadia township | 0.49946208 | 0.56750941 | 43.06379531 | -83.18758324 |
| Berrien County, Pipestone township | 0.4943518 | 0.56535772 | 42.02223596 | -86.2265222 |
| Lapeer County, Lapeer city | 0.50578268 | 0.56508876 | 43.03140772 | -83.3181314 |
| Lenawee County, Ogden township | 0.35933297 | 0.56455083 | 41.81098432 | -83.99854947 |
| Kent County, Kentwood city | 0.4077461 | 0.56293706 | 42.86889834 | -85.54919428 |
| Muskegon County, Muskegon Heights city | 0.55580958 | 0.56132329 | 43.1929401 | -86.24599633 |
| Cass County, Penn township | 0.52662722 | 0.56105433 | 41.93227059 | -85.93387905 |
| Shiawassee County, Owosso city | 0.50699301 | 0.55755783 | 43.00094653 | -84.19422996 |
| Lenawee County, Fairfield township | 0.33512641 | 0.55675094 | 41.81379508 | -84.02861648 |
| Jackson County, Columbia township | 0.39833244 | 0.55459925 | 42.07997769 | -84.2280333 |
| Kalamazoo County, Comstock charter township | 0.33674018 | 0.55298548 | 42.25886659 | -85.51830703 |
| Isabella County, Mount Pleasant city | 0.52259279 | 0.55298548 | 43.59389868 | -84.78062981 |
| St. Clair County, Mussey township | 0.43531469 | 0.55271651 | 42.99822212 | -82.9820248 |
| Jackson County, Blackman charter township | 0.33297472 | 0.55190963 | 42.25987657 | -84.4740716 |
| Lapeer County, Almont township | 0.50053792 | 0.55110274 | 42.96301379 | -83.08201472 |
| St. Clair County, Marysville city | 0.53456159 | 0.55110274 | 42.92626521 | -82.49809127 |
| Genesee County, Argentine township | 0.46463152 | 0.5500269 | 42.83114372 | -83.82036382 |
| Livingston County, Deerfield township | 0.49300699 | 0.54948897 | 42.74238846 | -83.83606685 |
| Cass County, Porter township | 0.4556213 | 0.5478752 | 41.78467465 | -85.77234101 |
| Lapeer County, Dryden township | 0.48318989 | 0.54491662 | 42.96604274 | -83.11254527 |
| Lenawee County, Ridgeway township | 0.45185584 | 0.54464766 | 41.99914929 | -83.84351476 |
| St. Joseph County, Colon township | 0.50511027 | 0.5443787 | 41.92801976 | -85.32370121 |
| Cass County, Newberg township | 0.47982786 | 0.54249597 | 41.92753122 | -85.87350401 |
| Berrien County, Coloma charter township | 0.42630446 | 0.54249597 | 42.16477356 | -86.3289592 |
| Gratiot County, Alma city | 0.42065627 | 0.54249597 | 43.38048179 | -84.65956669 |
| Hillsdale County, Hillsdale city | 0.4993276 | 0.53819258 | 41.91291063 | -84.6232101 |
| Lapeer County, Marathon township | 0.46369016 | 0.53200646 | 43.15714858 | -83.45070296 |
| Jackson County, Norvell township | 0.44728349 | 0.53093061 | 42.07721258 | -84.19784367 |
| Sanilac County, Marlette township | 0.3746638 | 0.53066165 | 43.30174226 | -83.080431 |
| Ingham County, Meridian charter township | 0.41635288 | 0.52985476 | 42.73145901 | -84.45701508 |
| Van Buren County, Hartford township | 0.47229693 | 0.5274341 | 42.19548778 | -86.14086712 |
| Lapeer County, Oregon township | 0.464766 | 0.52339968 | 43.06544754 | -83.43636511 |
| Ingham County, East Lansing city | 0.49811727 | 0.52259279 | 42.71195983 | -84.49121059 |
| Hillsdale County, Hillsdale township | 0.39833244 | 0.5204411 | 41.91557593 | -84.65334533 |
| Berrien County, Watervliet city | 0.51990317 | 0.51990317 | 42.18245403 | -86.26524183 |
| St. Clair County, Fort Gratiot charter township | 0.45831092 | 0.51963421 | 43.03702904 | -82.4762522 |
| Lenawee County, Cambridge township | 0.42980097 | 0.51882733 | 42.0577982 | -84.23179116 |
| Ingham County, Lansing charter township | 0.51748252 | 0.51748252 | 42.68975433 | -84.49491839 |
| Jackson County, Leoni township | 0.3189887 | 0.51667563 | 42.24900105 | -84.35297659 |
| St. Clair County, Clyde township | 0.34911243 | 0.51640667 | 43.00220537 | -82.57655539 |
| Van Buren County, Hamilton township | 0.4669177 | 0.51613771 | 42.07968856 | -86.09609979 |
| St. Clair County, Port Huron city | 0.5112964 | 0.51586875 | 42.96421323 | -82.42835783 |
| Lapeer County, Mayfield township | 0.46100054 | 0.51559978 | 43.06979776 | -83.24874554 |
| Van Buren County, Waverly township | 0.37036041 | 0.51452394 | 42.24579686 | -85.91906564 |
| Lenawee County, Blissfield township | 0.48816568 | 0.50995159 | 41.86615994 | -83.8667307 |
| Ottawa County, Hudsonville city | 0.48009683 | 0.50672405 | 42.87084887 | -85.85843676 |
| Muskegon County, Roosevelt Park city | 0.50672405 | 0.50672405 | 43.19523013 | -86.27675108 |
| Kent County, Alpine township | 0.30742335 | 0.50591716 | 43.05905997 | -85.67545276 |
| Saginaw County, Saginaw city | 0.40828402 | 0.50591716 | 43.45054619 | -83.92871978 |
| Monroe County, Summerfield township | 0.48682087 | 0.50537924 | 41.90191499 | -83.76870415 |
| Sanilac County, Marlette city | 0.50268962 | 0.50268962 | 43.32392609 | -83.07623292 |
| Van Buren County, Almena township | 0.3161646 | 0.50188273 | 42.25849909 | -85.79451416 |
| Roscommon County, Higgins township | 0.13959118 | 0.50026896 | 44.50397969 | -84.59713754 |
| Allegan County, Fillmore township | 0.26102743 | 0.49811727 | 42.75151564 | -86.06133631 |
| Lenawee County, Woodstock township | 0.36309844 | 0.49677246 | 42.06055532 | -84.26197374 |
| Genesee County, Thetford township | 0.35960194 | 0.49354492 | 43.14960135 | -83.60791106 |
| Berrien County, Bridgman city | 0.49273803 | 0.49273803 | 41.93585374 | -86.57431698 |
| Livingston County, Cohoctah township | 0.37587413 | 0.49193115 | 42.69000226 | -84.00002492 |
| St. Joseph County, Nottawa township | 0.45454545 | 0.49139322 | 41.89850293 | -85.51142774 |
| Kalamazoo County, Cooper charter township | 0.3655191 | 0.49058634 | 42.33288737 | -85.5992711 |
| Van Buren County, South Haven charter township | 0.32880581 | 0.48924153 | 42.36265314 | -86.27063983 |
| Newaygo County, Fremont city | 0.4887036 | 0.4887036 | 43.46382372 | -85.95765208 |
| Branch County, Matteson township | 0.32894029 | 0.48601399 | 41.9477165 | -85.29012194 |
| Midland County, Midland city | 0.37654653 | 0.48601399 | 43.6091491 | -84.18276327 |
| Lapeer County, North Branch township | 0.46005917 | 0.48305541 | 43.17469523 | -83.166811 |
| Calhoun County, Albion city | 0.45925229 | 0.48251748 | 42.2617335 | -84.75027598 |
| St. Clair County, Riley township | 0.40263583 | 0.48224852 | 42.92922638 | -82.74635699 |
| Kent County, Plainfield charter township | 0.39053254 | 0.48063475 | 43.0343815 | -85.64813376 |
| Kent County, Grandville city | 0.46355568 | 0.47982786 | 42.90815834 | -85.76003637 |
| Allegan County, Gun Plain township | 0.25389995 | 0.47928994 | 42.42666573 | -85.64669821 |
| Ingham County, Leroy township | 0.36430877 | 0.47875202 | 42.65960845 | -84.15992359 |
| Otsego County, Bagley township | 0.27622378 | 0.47767617 | 44.98552197 | -84.67491611 |
| Van Buren County, Lawrence township | 0.41460463 | 0.47740721 | 42.16394083 | -86.02283431 |
| Cass County, Marcellus township | 0.41447015 | 0.47660032 | 41.99419415 | -85.86380863 |
| St. Clair County, Cottrellville township | 0.40465304 | 0.47552448 | 42.67947705 | -82.51551113 |
| Berrien County, New Buffalo city | 0.47498655 | 0.47498655 | 41.79115366 | -86.74606674 |
| Ottawa County, Wright township | 0.16473911 | 0.47229693 | 43.04645716 | -85.80147332 |
| Kent County, Ada township | 0.25376547 | 0.47122109 | 42.95292958 | -85.47442364 |
| Oscoda County, Comins township | 0.40545992 | 0.47014524 | 44.68835788 | -84.12005665 |
| Muskegon County, North Muskegon city | 0.39967725 | 0.46933835 | 43.23976976 | -86.27043143 |
| Kent County, Gaines charter township | 0.32853685 | 0.46853147 | 42.84665904 | -85.55257032 |
| Allegan County, Holland city | 0.41823561 | 0.46611081 | 42.75384737 | -86.0918888 |
| Lapeer County, Goodland township | 0.40263583 | 0.46584185 | 43.07691368 | -83.09168176 |
| Shiawassee County, Owosso charter township | 0.32961269 | 0.46557289 | 42.97596448 | -84.16746699 |
| Gratiot County, Emerson township | 0.10866057 | 0.46503497 | 43.35290423 | -84.60171964 |
| Saginaw County, Buena Vista charter township | 0.28778913 | 0.46503497 | 43.4477055 | -83.89793848 |
| Livingston County, Conway township | 0.3302851 | 0.46449704 | 42.69562218 | -84.06092519 |
| Muskegon County, Muskegon charter township | 0.26573427 | 0.46449704 | 43.23287313 | -86.17811422 |
| Eaton County, Lansing city | 0.46342119 | 0.46342119 | 42.68362452 | -84.6814122 |
| Lenawee County, Deerfield township | 0.40935987 | 0.46315223 | 41.90477965 | -83.79880093 |
| Ottawa County, Holland city | 0.41931146 | 0.46261431 | 42.7760972 | -86.08869068 |
| Ingham County, Alaiedon township | 0.25201721 | 0.46207639 | 42.68161798 | -84.40352732 |
| Oscoda County, Mentor township | 0.33889188 | 0.46207639 | 44.67992739 | -84.02600136 |
| Oscoda County, Big Creek township | 0.26748252 | 0.46180742 | 44.66887774 | -84.15534241 |
| St. Clair County, Wales township | 0.40828402 | 0.46100054 | 42.9735467 | -82.73778883 |
| St. Clair County, Brockway township | 0.41043572 | 0.46073158 | 43.13801718 | -82.79948366 |
| Van Buren County, Arlington township | 0.3218128 | 0.45938677 | 42.27509856 | -86.0068485 |
| Clinton County, East Lansing city | 0.45938677 | 0.45938677 | 42.78128167 | -84.51057991 |
| Eaton County, Delta charter township | 0.3485745 | 0.45911781 | 42.70050328 | -84.6168064 |
| Cass County, Volinia township | 0.41823561 | 0.45911781 | 42.00365442 | -85.98468074 |
| Bay County, Bangor charter township | 0.23117267 | 0.45696611 | 43.62544317 | -83.86632836 |
| Jackson County, Napoleon township | 0.33579882 | 0.45589026 | 42.18243945 | -84.36415105 |
| St. Clair County, East China township | 0.43114578 | 0.45481442 | 42.74591094 | -82.50247952 |
| Ottawa County, Zeeland charter township | 0.30433029 | 0.45454545 | 42.83346748 | -85.95673084 |
| Clinton County, St. Johns city | 0.38999462 | 0.45454545 | 43.01149005 | -84.56522142 |
| Kalamazoo County, Brady township | 0.37399139 | 0.45400753 | 42.1008806 | -85.5115076 |
| St. Clair County, St. Clair city | 0.44109736 | 0.45266272 | 42.81234992 | -82.48942286 |
| Clinton County, Bath charter township | 0.25107585 | 0.45239376 | 42.77315517 | -84.41906608 |
| Lapeer County, Burnside township | 0.38515331 | 0.4521248 | 43.16865351 | -83.10556267 |
| Oceana County, Golden township | 0.22969338 | 0.45051103 | 43.65445101 | -86.39894534 |
| Kalamazoo County, Parchment city | 0.45024207 | 0.45024207 | 42.33044178 | -85.56892031 |
| Livingston County, Unadilla township | 0.3246369 | 0.44943518 | 42.46812267 | -84.03855381 |
| Calhoun County, Springfield city | 0.42146315 | 0.44916622 | 42.32528666 | -85.23175292 |
| St. Clair County, China township | 0.38300161 | 0.44835933 | 42.81551636 | -82.51986039 |
| Shiawassee County, Durand city | 0.44755245 | 0.44755245 | 42.9147642 | -83.99179354 |
| St. Clair County, Berlin township | 0.37856374 | 0.44674556 | 42.97605289 | -82.98623334 |
| Ottawa County, Tallmadge charter township | 0.26667563 | 0.4464766 | 43.02659816 | -85.83544122 |
| Lapeer County, Deerfield township | 0.43544917 | 0.4464766 | 43.2088793 | -83.28521608 |
| Ingham County, Delhi charter township | 0.26788596 | 0.44566971 | 42.66754943 | -84.49862381 |
| Oscoda County, Elmer township | 0.32974718 | 0.44217321 | 44.69115338 | -84.15141362 |
| St. Clair County, Emmett township | 0.36444325 | 0.44190425 | 42.98594126 | -82.85988498 |
| Kent County, Caledonia township | 0.31280258 | 0.44190425 | 42.82196296 | -85.5253791 |
| St. Clair County, Marine City city | 0.42791824 | 0.43975256 | 42.70162112 | -82.51117004 |
| St. Joseph County, Mendon township | 0.36659494 | 0.43948359 | 42.03423619 | -85.52155394 |
| Sanilac County, Lamotte township | 0.28146853 | 0.43948359 | 43.41570472 | -83.09014315 |
| Shiawassee County, Caledonia charter township | 0.26022055 | 0.43921463 | 42.99538629 | -84.13304927 |
| Clinton County, Watertown charter township | 0.17307692 | 0.43786982 | 42.77247042 | -84.66680751 |
| Sanilac County, Croswell city | 0.43706294 | 0.43706294 | 43.27758833 | -82.61620045 |
| Sanilac County, Buel township | 0.34104357 | 0.4351802 | 43.25855918 | -82.65118382 |
| Saginaw County, Saginaw charter township | 0.3415815 | 0.43464228 | 43.45337969 | -83.95950362 |
| Washtenaw County, Sharon township | 0.32235073 | 0.43410436 | 42.17415747 | -84.02782287 |
| Sanilac County, Lexington township | 0.36471221 | 0.43302851 | 43.25542115 | -82.62054406 |
| Jackson County, Summit township | 0.34171598 | 0.43275955 | 42.20735372 | -84.39067983 |
| Ottawa County, Allendale charter township | 0.24744486 | 0.43222162 | 42.9868556 | -85.90331833 |
| Muskegon County, Dalton township | 0.19029048 | 0.42953201 | 43.35113181 | -86.25459643 |
| Ottawa County, Ferrysburg city | 0.42872512 | 0.42872512 | 43.07931466 | -86.23110533 |
| Washtenaw County, Freedom township | 0.31038193 | 0.42738031 | 42.17134532 | -83.99760048 |
| Clinton County, Bingham township | 0.29585799 | 0.42711135 | 42.98658362 | -84.53832022 |
| Bay County, Bay City city | 0.3683432 | 0.4260355 | 43.60322334 | -83.87028861 |
| St. Joseph County, Leonidas township | 0.33795051 | 0.42442173 | 41.99964187 | -85.37387561 |
| Allegan County, Plainwell city | 0.42388381 | 0.42388381 | 42.44889426 | -85.64337269 |
| St. Clair County, Lynn township | 0.35099516 | 0.42388381 | 43.1472829 | -82.89127872 |
| Kent County, Grand Rapids charter township | 0.38757396 | 0.42307692 | 42.96031761 | -85.56629401 |
| Jackson County, Grass Lake charter township | 0.32907477 | 0.42173211 | 42.25743133 | -84.19788428 |
| Hillsdale County, Cambria township | 0.16635288 | 0.42146315 | 41.89072418 | -84.62683375 |
| Branch County, Bronson township | 0.2760893 | 0.41958042 | 41.8785873 | -85.27021265 |
| Hillsdale County, Adams township | 0.1993007 | 0.41796665 | 41.9324243 | -84.5894418 |
| Lapeer County, Burlington township | 0.35650888 | 0.41769769 | 43.26040542 | -83.11948064 |
| Wexford County, Cadillac city | 0.34211942 | 0.41581495 | 44.254185 | -85.39795525 |
| Ingham County, Locke township | 0.21005917 | 0.41447015 | 42.70399879 | -84.15229405 |
| Kent County, East Grand Rapids city | 0.41393222 | 0.41393222 | 42.94052229 | -85.60028708 |
| Isabella County, Chippewa township | 0.2140936 | 0.41366326 | 43.61085109 | -84.7151886 |
| Genesee County, Clio city | 0.41312534 | 0.41312534 | 43.18338564 | -83.72649044 |
| Grand Traverse County, Garfield charter township | 0.34709521 | 0.41204949 | 44.7249689 | -85.64178017 |
| Livingston County, Iosco township | 0.2380312 | 0.41016676 | 42.58186393 | -84.04972115 |
| Shiawassee County, Vernon township | 0.33512641 | 0.40909091 | 42.93696245 | -83.98791183 |
| Clinton County, DeWitt charter township | 0.22754169 | 0.40828402 | 42.80349055 | -84.50687223 |
| Grand Traverse County, Traverse City city | 0.36874664 | 0.40801506 | 44.74484131 | -85.60689855 |
| Oscoda County, Clinton township | 0.36148467 | 0.40801506 | 44.79129913 | -84.00611455 |
| Otsego County, Livingston township | 0.33230231 | 0.40505648 | 45.09973836 | -84.68758578 |
| Genesee County, Flushing city | 0.4042496 | 0.4042496 | 43.05880048 | -83.84213241 |
| Muskegon County, Fruitport charter township | 0.19876278 | 0.40398063 | 43.18833845 | -86.18449298 |
| Bay County, Hampton charter township | 0.16137708 | 0.40290479 | 43.60036822 | -83.83944108 |
| Lapeer County, Rich township | 0.32920925 | 0.4020979 | 43.25026185 | -83.24629538 |
| Muskegon County, Norton Shores city | 0.30997848 | 0.40182894 | 43.19064287 | -86.21524363 |
| Sanilac County, Flynn township | 0.30957504 | 0.40182894 | 43.29260656 | -82.98840419 |
| Hillsdale County, Fayette township | 0.24233459 | 0.40182894 | 41.95728491 | -84.6159559 |
| Ontonagon County, Matchwood township | 0.33391608 | 0.40155998 | 46.57006328 | -89.49556595 |
| Sanilac County, Watertown township | 0.30688542 | 0.40048413 | 43.41339262 | -82.84010219 |
| Tuscola County, Millington township | 0.2049489 | 0.39940828 | 43.22666824 | -83.46914713 |
| Branch County, Coldwater city | 0.38111888 | 0.3986014 | 41.94701111 | -85.0151453 |
| Sanilac County, Speaker township | 0.31871974 | 0.39725659 | 43.16018601 | -82.79520507 |
| Allegan County, Allegan city | 0.35825713 | 0.39698763 | 42.53245171 | -85.8464659 |
| Saginaw County, Birch Run township | 0.22538999 | 0.39671867 | 43.25574465 | -83.77590818 |
| Ottawa County, Jamestown charter township | 0.23184508 | 0.39618074 | 42.85098044 | -85.89230199 |
| Branch County, Kinderhook township | 0.19029048 | 0.39591178 | 41.76428428 | -84.98292013 |
| Otsego County, Gaylord city | 0.3951049 | 0.3951049 | 45.00782864 | -84.67114113 |
| Washtenaw County, Sylvan township | 0.29626143 | 0.39349112 | 42.29624336 | -84.12975675 |
| Ottawa County, Grand Haven city | 0.31925767 | 0.39322216 | 43.05705115 | -86.23427152 |
| Shiawassee County, Burns township | 0.3542227 | 0.39241528 | 42.84535277 | -83.97290747 |
| Montcalm County, Eureka township | 0.26533082 | 0.39160839 | 43.20483125 | -85.24901599 |
| Jackson County, Sandstone township | 0.14550834 | 0.39053254 | 42.2625778 | -84.50435125 |
| Sanilac County, Maple Valley township | 0.36081226 | 0.39026358 | 43.17865993 | -82.97887782 |
| Arenac County, Lincoln township | 0.16850457 | 0.39026358 | 43.97308574 | -83.96168301 |
| Midland County, Homer township | 0.21516945 | 0.38864981 | 43.64243869 | -84.30242098 |
| Shiawassee County, Perry city | 0.38864981 | 0.38864981 | 42.82331252 | -84.22471265 |
| Allegan County, Hopkins township | 0.18289403 | 0.387305 | 42.67607896 | -85.67100532 |
| Montcalm County, Greenville city | 0.387305 | 0.387305 | 43.1825882 | -85.25250554 |
| Tuscola County, Caro city | 0.33727811 | 0.387305 | 43.49007411 | -83.38937752 |
| Van Buren County, South Haven city | 0.38703604 | 0.38703604 | 42.40713798 | -86.26439191 |
| St. Clair County, Kenockee township | 0.34440559 | 0.38703604 | 42.99570778 | -82.73350062 |
| Calhoun County, Marshall city | 0.3239645 | 0.38676708 | 42.27744781 | -84.93205636 |
| Alcona County, Mitchell township | 0.33109198 | 0.38569123 | 44.71026496 | -83.86124643 |
| Delta County, Escanaba city | 0.28725121 | 0.38542227 | 45.75354681 | -87.06259827 |
| Clinton County, Greenbush township | 0.12681549 | 0.38515331 | 43.03370821 | -84.56151896 |
| Sanilac County, Worth township | 0.33082302 | 0.38488435 | 43.23325464 | -82.62488488 |
| Wexford County, Haring charter township | 0.22404519 | 0.38461538 | 44.27647556 | -85.3944583 |
| Kent County, Rockford city | 0.38434642 | 0.38434642 | 43.11849074 | -85.57330418 |
| Washtenaw County, Lyndon township | 0.21382464 | 0.38407746 | 42.39876188 | -84.01974237 |
| Kent County, Cannon township | 0.17482517 | 0.3838085 | 43.09378866 | -85.54599823 |
| Ottawa County, Crockery township | 0.17132867 | 0.38353954 | 43.0923313 | -86.10510416 |
| Ottawa County, Zeeland city | 0.38327058 | 0.38327058 | 42.81592322 | -86.02112703 |
| Tuscola County, Watertown township | 0.23292093 | 0.38327058 | 43.23999604 | -83.37307249 |
| Montcalm County, Montcalm township | 0.22942442 | 0.38300161 | 43.20735482 | -85.27974123 |
| Ingham County, Williamstown township | 0.15572889 | 0.38138784 | 42.76770193 | -84.35806884 |
| Allegan County, Wayland city | 0.38084992 | 0.38084992 | 42.67364899 | -85.64050161 |
| Ogemaw County, Horton township | 0.24502421 | 0.37896719 | 44.24582172 | -84.22949995 |
| Iosco County, Oscoda charter township | 0.21772458 | 0.37842926 | 44.43516171 | -83.33925163 |
| Delta County, Wells township | 0.14254976 | 0.37789134 | 45.77595676 | -87.05958209 |
| Shiawassee County, Corunna city | 0.33136095 | 0.37681549 | 42.97318178 | -84.13688757 |
| Montmorency County, Loud township | 0.35153308 | 0.37600861 | 44.86659038 | -84.08845109 |
| Branch County, Coldwater township | 0.30016138 | 0.37573964 | 41.90003618 | -84.99200176 |
| Sanilac County, Custer township | 0.26035503 | 0.37520172 | 43.43557288 | -82.83581955 |
| Alcona County, Millen township | 0.20710059 | 0.37493276 | 44.65424027 | -83.74397766 |
| St. Clair County, Greenwood township | 0.28012372 | 0.37305003 | 43.10963847 | -82.74259585 |
| Saginaw County, Carrollton township | 0.37143626 | 0.37143626 | 43.47559636 | -83.95558094 |
| Sanilac County, Elmer township | 0.28509952 | 0.37116729 | 43.41648023 | -82.87082245 |
| Saginaw County, Kochville township | 0.28886498 | 0.37089833 | 43.52003191 | -83.94772794 |
| Alcona County, Curtis township | 0.27945132 | 0.36982249 | 44.59321115 | -83.81874491 |
| Crawford County, Grayling charter township | 0.10718128 | 0.36928456 | 44.60380945 | -84.70736851 |
| Ontonagon County, Bergland township | 0.31105433 | 0.36820871 | 46.57313389 | -89.56042913 |
| Lenawee County, Seneca township | 0.32087144 | 0.36767079 | 41.80001563 | -84.12261054 |
| Grand Traverse County, Blair township | 0.29841313 | 0.36767079 | 44.68033322 | -85.64865475 |
| Van Buren County, Bangor township | 0.26573427 | 0.36686391 | 42.30666442 | -86.12506258 |
| Marquette County, Powell township | 0.29263045 | 0.36659494 | 46.84309589 | -87.83152243 |
| Allegan County, Otsego township | 0.18558365 | 0.36659494 | 42.42909344 | -85.67709375 |
| Clare County, Grant township | 0.15021517 | 0.36605702 | 43.84129138 | -84.77118627 |
| Tuscola County, Koylton township | 0.30123722 | 0.36578806 | 43.3521697 | -83.1334358 |
| Alcona County, Harrisville city | 0.36578806 | 0.36578806 | 44.65759945 | -83.29729321 |
| Eaton County, Grand Ledge city | 0.27864443 | 0.36471221 | 42.75820799 | -84.76197483 |
| Keweenaw County, Grant township | 0.30876815 | 0.36444325 | 47.45887972 | -87.88646033 |
| Allegan County, Leighton township | 0.15250134 | 0.36363636 | 42.69588493 | -85.63716223 |
| Jackson County, Rives township | 0.15613233 | 0.36363636 | 42.36812885 | -84.42526537 |
| Jackson County, Hanover township | 0.1880043 | 0.36282948 | 42.14892429 | -84.49249416 |
| Kent County, Lowell charter township | 0.1958042 | 0.36202259 | 42.92573004 | -85.41661139 |
| Sanilac County, Sanilac township | 0.31522324 | 0.36202259 | 43.42962327 | -82.55500195 |
| Ottawa County, Park township | 0.22216245 | 0.36094675 | 42.78074088 | -86.14982127 |
| Sanilac County, Fremont township | 0.31253362 | 0.36094675 | 43.24264395 | -82.71678347 |
| Marquette County, Marquette city | 0.33620226 | 0.36094675 | 46.56795413 | -87.40882459 |
| Kalamazoo County, Galesburg city | 0.3598709 | 0.3598709 | 42.29588859 | -85.42058999 |
| Genesee County, Flushing charter township | 0.26143088 | 0.3598709 | 43.08099919 | -83.83819437 |
| Charlevoix County, St. James township | 0.33539537 | 0.3598709 | 45.7353817 | -85.5484678 |
| Branch County, Noble township | 0.26707907 | 0.35960194 | 41.76757571 | -85.287275 |
| Alpena County, Alpena city | 0.29047875 | 0.35960194 | 45.079138 | -83.44156218 |
| Allegan County, Wayland township | 0.15034965 | 0.35933297 | 42.65141366 | -85.64383884 |
| Midland County, Lincoln township | 0.22108661 | 0.35852609 | 43.66467283 | -84.29859891 |
| Dickinson County, Iron Mountain city | 0.30782679 | 0.35852609 | 45.81545187 | -88.0544948 |
| Schoolcraft County, Seney township | 0.3148198 | 0.35825713 | 46.33876184 | -86.03913133 |
| Keweenaw County, Eagle Harbor township | 0.33257127 | 0.35798817 | 47.96666592 | -88.85553301 |
| Ottawa County, Coopersville city | 0.3041958 | 0.3577192 | 43.05598858 | -85.9241834 |
| Montmorency County, Albert township | 0.25618612 | 0.35745024 | 44.89447682 | -84.14738902 |
| Ottawa County, Spring Lake township | 0.29922001 | 0.35718128 | 43.07701437 | -86.20040383 |
| Saginaw County, Thomas township | 0.16568047 | 0.35664336 | 43.39518231 | -84.06353492 |
| Gratiot County, St. Louis city | 0.35610543 | 0.35610543 | 43.40004552 | -84.62508581 |
| Van Buren County, Pine Grove township | 0.1309844 | 0.35529855 | 42.33234703 | -85.87577118 |
| Houghton County, Houghton city | 0.34911243 | 0.3542227 | 47.11263804 | -88.58710276 |
| Bay County, Portsmouth charter township | 0.2619688 | 0.35395374 | 43.57815018 | -83.8434086 |
| Charlevoix County, Peaine township | 0.32288865 | 0.35368478 | 45.71053749 | -85.52009594 |
| Ontonagon County, Greenland township | 0.28119957 | 0.35341581 | 46.75356275 | -89.0852546 |
| Allegan County, Allegan township | 0.24448628 | 0.35314685 | 42.55468849 | -85.84319932 |
| Ogemaw County, West Branch township | 0.24852071 | 0.35260893 | 44.2680795 | -84.22561996 |
| Clinton County, Eagle township | 0.09252286 | 0.35233997 | 42.79999428 | -84.72419519 |
| Kalamazoo County, Charleston township | 0.26573427 | 0.35207101 | 42.29340133 | -85.39026166 |
| Saginaw County, Zilwaukee township | 0.23641743 | 0.35207101 | 43.4949782 | -83.92085224 |
| Oceana County, Leavitt township | 0.29558903 | 0.35126412 | 43.70067077 | -86.11063506 |
| Gogebic County, Marenisco township | 0.28994083 | 0.35099516 | 46.50408175 | -89.53462628 |
| Bay County, Frankenlust township | 0.22996235 | 0.35099516 | 43.5422508 | -83.94379762 |
| Alger County, Burt township | 0.3246369 | 0.3507262 | 46.65753814 | -86.05619711 |
| Luce County, McMillan township | 0.27138246 | 0.35045724 | 46.59623118 | -85.2476566 |
| Tuscola County, Kingston township | 0.28160301 | 0.35018827 | 43.41873397 | -83.12087558 |
| Calhoun County, Albion township | 0.12668101 | 0.34830554 | 42.23689133 | -84.72359808 |
| Iron County, Bates township | 0.22377622 | 0.34830554 | 46.41665221 | -88.69777619 |
| Tuscola County, Novesta township | 0.23856912 | 0.34803658 | 43.58314331 | -83.18389229 |
| Keweenaw County, Houghton township | 0.32544379 | 0.34749866 | 48.00672985 | -88.75113844 |
| Chippewa County, Whitefish township | 0.30957504 | 0.34669177 | 46.54629044 | -85.19042055 |
| Iosco County, Au Sable charter township | 0.27756859 | 0.34642281 | 44.41292303 | -83.34343232 |
| Allegan County, Casco township | 0.21490048 | 0.34480904 | 42.42938127 | -86.26126496 |
| Allegan County, Dorr township | 0.18585261 | 0.34480904 | 42.72055319 | -85.66434378 |
| Baraga County, Covington township | 0.26896181 | 0.34480904 | 46.43390759 | -88.59825556 |
| Washtenaw County, Chelsea city | 0.34454008 | 0.34454008 | 42.31003596 | -84.0350938 |
| Calhoun County, Sheridan township | 0.1394567 | 0.34346423 | 42.26437022 | -84.780567 |
| Bay County, Williams charter township | 0.24448628 | 0.34346423 | 43.60636702 | -84.15190019 |
| Mecosta County, Big Rapids charter township | 0.23561054 | 0.34265734 | 43.67494087 | -85.48807652 |
| Shiawassee County, Bennington township | 0.15236686 | 0.34185046 | 42.95375963 | -84.17129314 |
| Ontonagon County, Carp Lake township | 0.31105433 | 0.34185046 | 46.78780331 | -89.80086674 |
| Montmorency County, Rust township | 0.2619688 | 0.3415815 | 44.85813418 | -83.99415089 |
| Oceana County, Hart township | 0.24811727 | 0.34050565 | 43.67674092 | -86.39580514 |
| Bay County, Essexville city | 0.34050565 | 0.34050565 | 43.62258702 | -83.83547099 |
| Kent County, Algoma township | 0.16594944 | 0.34023669 | 43.1407399 | -85.56992151 |
| Lenawee County, Rollin township | 0.267617 | 0.33916084 | 41.97732391 | -84.33724048 |
| Eaton County, Eaton Rapids city | 0.31401291 | 0.33916084 | 42.50064635 | -84.64972428 |
| Gratiot County, Arcada township | 0.14846692 | 0.33862292 | 43.36090967 | -84.69402791 |
| Calhoun County, Bedford charter township | 0.19203873 | 0.33835395 | 42.35254489 | -85.28899407 |
| Ingham County, Mason city | 0.309844 | 0.33835395 | 42.57332813 | -84.45258807 |
| Iron County, Hematite township | 0.23318989 | 0.33754707 | 46.41142459 | -88.60077871 |
| St. Clair County, Burtchville township | 0.27850995 | 0.33754707 | 43.08451172 | -82.49805228 |
| Mecosta County, Big Rapids city | 0.29370629 | 0.33431953 | 43.69720892 | -85.48463854 |
| Jackson County, Spring Arbor township | 0.21584185 | 0.33324368 | 42.24038539 | -84.50803095 |
| Baraga County, L'Anse township | 0.23426573 | 0.33243679 | 46.74170446 | -88.43278545 |
| Chippewa County, Drummond township | 0.31159225 | 0.33136095 | 45.99188008 | -83.49532459 |
| Hillsdale County, Somerset township | 0.26223776 | 0.33109198 | 42.00223753 | -84.36368181 |
| Calhoun County, Tekonsha township | 0.17011834 | 0.33082302 | 42.10241151 | -84.99057533 |
| Saginaw County, Bridgeport charter township | 0.25389995 | 0.3302851 | 43.35601056 | -83.88294263 |
| Ontonagon County, Ontonagon township | 0.27568585 | 0.33001614 | 46.68262476 | -89.4844314 |
| Kalamazoo County, Richland township | 0.17522862 | 0.32894029 | 42.39465934 | -85.52849917 |
| Shiawassee County, Perry township | 0.22525551 | 0.32840237 | 42.8427449 | -84.19038702 |
| Bay County, Fraser township | 0.22081764 | 0.32840237 | 43.8146156 | -83.95832506 |
| Houghton County, Duncan township | 0.25551372 | 0.32813341 | 46.50660149 | -88.68780469 |
| Houghton County, Hancock city | 0.32544379 | 0.32813341 | 47.13517028 | -88.58454787 |
| Jackson County, Liberty township | 0.23318989 | 0.32813341 | 42.16025325 | -84.36787113 |
| Hillsdale County, Wheatland township | 0.23830016 | 0.32759548 | 41.98005616 | -84.36739245 |
| Gogebic County, Wakefield township | 0.25739645 | 0.32732652 | 46.70207472 | -89.90686568 |
| Allegan County, Laketown township | 0.17845616 | 0.3267886 | 42.71398838 | -86.15935116 |
| Tuscola County, Elkland township | 0.21557289 | 0.32571275 | 43.60534117 | -83.17971049 |
| Grand Traverse County, Long Lake township | 0.26707907 | 0.32544379 | 44.75700237 | -85.76412587 |
| Otsego County, Otsego Lake township | 0.19042496 | 0.32490586 | 44.87400438 | -84.69375355 |
| Barry County, Thornapple township | 0.232383 | 0.32436794 | 42.72808894 | -85.47784561 |
| Branch County, Sherwood township | 0.22243141 | 0.32436794 | 41.99212896 | -85.28328772 |
| Gogebic County, Wakefield city | 0.30957504 | 0.32436794 | 46.47690982 | -89.92766364 |
| Clare County, Clare city | 0.27918236 | 0.32409898 | 43.81903663 | -84.77486421 |
| Lenawee County, Hudson township | 0.29276493 | 0.32383002 | 41.84425099 | -84.35951139 |
| Gogebic County, Watersmeet township | 0.29935449 | 0.32383002 | 46.19601065 | -89.24071081 |
| Benzie County, Benzonia township | 0.24341044 | 0.32383002 | 44.62446804 | -86.10177923 |
| Clare County, Hayes township | 0.25551372 | 0.32275417 | 44.06917382 | -84.79646474 |
| Alger County, Munising township | 0.2760893 | 0.32087144 | 46.41816487 | -86.51614962 |
| Muskegon County, Laketon township | 0.1542496 | 0.32006455 | 43.28431235 | -86.26410359 |
| Van Buren County, Covert township | 0.2534965 | 0.31979559 | 42.24916631 | -86.25588604 |
| Dickinson County, West Branch township | 0.26613771 | 0.31979559 | 46.08398053 | -87.63359598 |
| Saginaw County, James township | 0.23776224 | 0.31871974 | 43.39237341 | -84.0327729 |
| Houghton County, Franklin township | 0.1845078 | 0.31871974 | 47.13340458 | -88.55186168 |
| Saginaw County, Chesaning township | 0.23991393 | 0.31737493 | 43.15083623 | -84.10611184 |
| Iron County, Iron River city | 0.31535772 | 0.31710597 | 46.09681298 | -88.63592332 |
| Houghton County, Adams township | 0.11672942 | 0.31683701 | 47.11439527 | -88.61977954 |
| Kalamazoo County, Wakeshma township | 0.25 | 0.31656805 | 42.08347457 | -85.29982492 |
| Van Buren County, Gobles city | 0.31656805 | 0.31656805 | 42.3545786 | -85.87252336 |
| Arenac County, Moffatt township | 0.18948359 | 0.31656805 | 44.12342504 | -84.12445944 |
| Houghton County, Portage charter township | 0.14618074 | 0.31656805 | 47.08834319 | -88.55699203 |
| Iron County, Iron River township | 0.22149005 | 0.31629909 | 46.40102893 | -88.82957474 |
| Muskegon County, Ravenna township | 0.12399139 | 0.31522324 | 43.19426715 | -85.96603935 |
| Sanilac County, Washington township | 0.2711135 | 0.31468531 | 43.36940742 | -82.62948617 |
| Tuscola County, Indianfields township | 0.26627219 | 0.31387843 | 43.42347637 | -83.40169023 |
| Schoolcraft County, Germfask township | 0.2288865 | 0.31360947 | 46.28681582 | -85.9493355 |
| Dickinson County, Breitung charter township | 0.13232921 | 0.31334051 | 45.81356807 | -88.02246987 |
| Shiawassee County, Antrim township | 0.22417967 | 0.31307154 | 42.78439821 | -84.04552131 |
| Marquette County, Marquette charter township | 0.15236686 | 0.31307154 | 46.54751795 | -87.44412059 |
| Oceana County, Elbridge township | 0.2739376 | 0.31280258 | 43.68302531 | -86.17580884 |
| Shiawassee County, Venice township | 0.21920387 | 0.31280258 | 42.95633585 | -83.95346311 |
| Ogemaw County, Rose township | 0.28886498 | 0.3119957 | 44.47951964 | -84.0616328 |
| Shiawassee County, Woodhull township | 0.18074233 | 0.31172673 | 42.80937149 | -84.32004032 |
| Lenawee County, Medina township | 0.2612964 | 0.31172673 | 41.70846594 | -84.35166083 |
| Hillsdale County, Pittsford township | 0.27528241 | 0.31145777 | 41.84697763 | -84.38960694 |
| Van Buren County, Geneva township | 0.24098978 | 0.31145777 | 42.40255042 | -86.20358213 |
| Schoolcraft County, Doyle township | 0.19661108 | 0.31118881 | 46.31634052 | -86.04251918 |
| Mason County, Ludington city | 0.30769231 | 0.31118881 | 43.95106373 | -86.45126206 |
| Ottawa County, Blendon township | 0.24663798 | 0.31091985 | 42.94235175 | -85.90985154 |
| Menominee County, Menominee city | 0.28415815 | 0.31091985 | 45.11231741 | -87.62711407 |
| Van Buren County, Bloomingdale township | 0.24018289 | 0.31065089 | 42.33472087 | -85.90613341 |
| Kent County, Solon township | 0.14120495 | 0.31065089 | 43.20749163 | -85.55976037 |
| Muskegon County, Whitehall city | 0.31065089 | 0.31065089 | 43.40253102 | -86.34079129 |
| Sanilac County, Elk township | 0.26896181 | 0.31065089 | 43.25815017 | -82.87000331 |
| Delta County, Masonville township | 0.14671867 | 0.31011296 | 45.92860654 | -86.97431225 |
| Emmet County, Bear Creek township | 0.20091447 | 0.31011296 | 45.34933712 | -84.96657253 |
| Houghton County, Quincy township | 0.29343733 | 0.31011296 | 47.15770416 | -88.58199123 |
| Ontonagon County, Haight township | 0.27434104 | 0.309844 | 46.3790899 | -89.28694434 |
| Alger County, Au Train township | 0.27420656 | 0.30876815 | 46.31710447 | -86.69338491 |
| Leelanau County, Leland township | 0.23534158 | 0.30849919 | 45.14058122 | -86.05734396 |
| Montmorency County, Avery township | 0.24394836 | 0.30796127 | 44.95573099 | -84.07258549 |
| Schoolcraft County, Manistique township | 0.1993007 | 0.30769231 | 45.96709518 | -86.22456109 |
| Alger County, Munising city | 0.30096826 | 0.30769231 | 46.40464928 | -86.64843423 |
| Ontonagon County, Rockland township | 0.25658956 | 0.30769231 | 46.75519998 | -89.11776601 |
| St. Clair County, Grant township | 0.2711135 | 0.30742335 | 43.097132 | -82.62030177 |
| Gogebic County, Ironwood charter township | 0.26949973 | 0.30742335 | 46.68385337 | -90.00643836 |
| Emmet County, Petoskey city | 0.25094137 | 0.30715438 | 45.37167158 | -84.96287432 |
| Oceana County, Weare township | 0.24677246 | 0.30661646 | 43.76590884 | -86.38322386 |
| Ogemaw County, Hill township | 0.27017214 | 0.3063475 | 44.40427605 | -83.97979615 |
| Manistee County, Bear Lake township | 0.22619688 | 0.30554061 | 44.42357935 | -86.13123407 |
| Grand Traverse County, Mayfield township | 0.22081764 | 0.30554061 | 44.59350712 | -85.6937481 |
| Marquette County, Wells township | 0.26882733 | 0.30554061 | 46.05955541 | -87.60430063 |
| Alpena County, Alpena township | 0.13636364 | 0.30446477 | 45.05981996 | -83.47724923 |
| Schoolcraft County, Manistique city | 0.3041958 | 0.3041958 | 45.96940401 | -86.2566034 |
| Menominee County, Holmes township | 0.26600323 | 0.30365788 | 45.48243132 | -87.77321781 |
| Gratiot County, Bethany township | 0.1035503 | 0.30258203 | 43.39736792 | -84.59429914 |
| Leelanau County, Leelanau township | 0.26896181 | 0.30258203 | 45.13178245 | -85.64273736 |
| Baraga County, Spurr township | 0.26331361 | 0.30231307 | 46.54532208 | -88.1619431 |
| Ontonagon County, Interior township | 0.26492738 | 0.30177515 | 46.34362734 | -89.03071731 |
| Branch County, Girard township | 0.14416353 | 0.30123722 | 41.9914088 | -85.00813646 |
| Roscommon County, Gerrish township | 0.15519096 | 0.30123722 | 44.5093375 | -84.65972168 |
| Oceana County, Colfax township | 0.20199032 | 0.30096826 | 43.74756668 | -86.13515895 |
| Menominee County, Stephenson township | 0.267617 | 0.30096826 | 45.40540521 | -87.62239441 |
| Baraga County, Arvon township | 0.24771382 | 0.30043034 | 46.90148537 | -88.05376079 |
| Allegan County, Ganges township | 0.13864981 | 0.29989242 | 42.51606606 | -86.21828299 |
| Ionia County, Berlin township | 0.08687466 | 0.29935449 | 42.87571442 | -85.08360813 |
| Lenawee County, Morenci city | 0.29935449 | 0.29935449 | 41.71965404 | -84.22779765 |
| Marquette County, Michigamme township | 0.26586875 | 0.29908553 | 46.67462276 | -88.04850837 |
| Genesee County, Montrose charter township | 0.22189349 | 0.29881657 | 43.1475993 | -83.82636502 |
| Tuscola County, Fremont township | 0.22538999 | 0.29881657 | 43.39831593 | -83.3750521 |
| Barry County, Hastings city | 0.27756859 | 0.29854761 | 42.64644899 | -85.30510357 |
| Oceana County, Newfield township | 0.26976869 | 0.29827864 | 43.61153969 | -86.12355937 |
| Ogemaw County, Cumming township | 0.27259279 | 0.29827864 | 44.38484484 | -84.01497431 |
| Isabella County, Isabella township | 0.2084454 | 0.29800968 | 43.64102782 | -84.80420159 |
| Chippewa County, Raber township | 0.23372781 | 0.29800968 | 46.0271339 | -84.1387973 |
| Marquette County, Ishpeming city | 0.23305541 | 0.29800968 | 46.49411317 | -87.67923959 |
| Alger County, Grand Island township | 0.27756859 | 0.29800968 | 46.49664006 | -86.66798718 |
| Presque Isle County, Rogers City city | 0.285234 | 0.29774072 | 45.40973176 | -83.83015395 |
| Montmorency County, Briley township | 0.20091447 | 0.29747176 | 44.96134026 | -84.1355474 |
| Muskegon County, Fruitland township | 0.23937601 | 0.2972028 | 43.37797812 | -86.3130969 |
| Leelanau County, Solon township | 0.23587951 | 0.29666487 | 44.77690809 | -85.72926375 |
| Sanilac County, Moore township | 0.26223776 | 0.29666487 | 43.42263399 | -82.93227124 |
| Grand Traverse County, Grant township | 0.2352071 | 0.29585799 | 44.5711939 | -85.69716269 |
| Calhoun County, Marshall township | 0.13582571 | 0.29558903 | 42.32704404 | -84.98561097 |
| Ogemaw County, Ogemaw township | 0.20414201 | 0.29558903 | 44.29310751 | -84.25291469 |
| Iron County, Stambaugh township | 0.23722431 | 0.29532006 | 46.31782479 | -88.96849941 |
| Bay County, Pinconning township | 0.20199032 | 0.29532006 | 43.84249832 | -84.01630224 |
| Marquette County, Negaunee township | 0.1873319 | 0.2950511 | 46.53109838 | -87.54410773 |
| Menominee County, Spalding township | 0.18370091 | 0.29343733 | 45.69272823 | -87.52177047 |
| Luce County, Columbus township | 0.23789672 | 0.29343733 | 46.48142039 | -85.8216505 |
| Schoolcraft County, Hiawatha township | 0.25739645 | 0.29316837 | 46.25210533 | -86.40995399 |
| Mackinac County, Garfield township | 0.19876278 | 0.29316837 | 46.07852139 | -85.59155954 |
| Grand Traverse County, Green Lake township | 0.22996235 | 0.29209252 | 44.68519946 | -85.7114818 |
| Gogebic County, Bessemer township | 0.25497579 | 0.29209252 | 46.31210712 | -89.78056927 |
| Ottawa County, Olive township | 0.15169446 | 0.29209252 | 42.86743173 | -86.10648083 |
| Gogebic County, Ironwood city | 0.27434104 | 0.29128564 | 46.46567335 | -90.18884497 |
| Sanilac County, Bridgehampton township | 0.2288865 | 0.29047875 | 43.4390645 | -82.64715412 |
| Houghton County, Laird township | 0.2471759 | 0.29047875 | 46.76608888 | -88.88778361 |
| Dickinson County, Breen township | 0.19123185 | 0.29020979 | 46.03909931 | -87.63926638 |
| Schoolcraft County, Thompson township | 0.20481442 | 0.29020979 | 46.20725732 | -86.41646758 |
| Presque Isle County, Rogers township | 0.25511027 | 0.28994083 | 45.38455319 | -83.80256204 |
| Clare County, Frost township | 0.22633136 | 0.28994083 | 44.0887981 | -84.761674 |
| Mackinac County, Hendricks township | 0.25685853 | 0.28994083 | 46.16035405 | -85.18856063 |
| Presque Isle County, Krakow township | 0.21866595 | 0.28913394 | 45.33899703 | -83.55346927 |
| Ontonagon County, Stannard township | 0.25618612 | 0.28832706 | 46.66839808 | -89.19212689 |
| Allegan County, Martin township | 0.12506724 | 0.2880581 | 42.54024613 | -85.66049293 |
| Saginaw County, Zilwaukee city | 0.28725121 | 0.28725121 | 43.47276183 | -83.92478728 |
| Chippewa County, Trout Lake township | 0.20871436 | 0.28671329 | 46.19483274 | -85.05276911 |
| Menominee County, Lake township | 0.26654115 | 0.28644432 | 45.44148795 | -87.84242165 |
| Mason County, Pere Marquette charter township | 0.24031737 | 0.28617536 | 43.9733672 | -86.44812531 |
| Chippewa County, Sault Ste. Marie city | 0.2626412 | 0.28617536 | 46.47715283 | -84.35189163 |
| Newaygo County, Dayton township | 0.22592792 | 0.28563744 | 43.48609539 | -85.95437721 |
| Ontonagon County, McMillan township | 0.26223776 | 0.28536848 | 46.53974792 | -89.33571208 |
| Oceana County, Shelby township | 0.24744486 | 0.28509952 | 43.60761211 | -86.37427419 |
| Benzie County, Joyfield township | 0.26331361 | 0.28483055 | 44.57981915 | -86.10833988 |
| Alcona County, Caledonia township | 0.25524476 | 0.28483055 | 44.82656241 | -83.64858816 |
| Presque Isle County, Bismarck township | 0.2373588 | 0.28483055 | 45.21189456 | -83.89834909 |
| Wexford County, Wexford township | 0.22404519 | 0.28483055 | 44.50426017 | -85.70739296 |
| Kent County, Courtland township | 0.15868747 | 0.28429263 | 43.20503362 | -85.52902492 |
| Cheboygan County, Burt township | 0.09279182 | 0.28429263 | 45.50154422 | -84.61920995 |
| Oceana County, Grant township | 0.21086606 | 0.28402367 | 43.49392226 | -86.35908034 |
| Sanilac County, Forester township | 0.21221087 | 0.28375471 | 43.47712347 | -82.57698148 |
| Mackinac County, St. Ignace township | 0.20925229 | 0.28375471 | 46.05135129 | -84.68684029 |
| Ogemaw County, Goodar township | 0.26237224 | 0.28348575 | 44.42653378 | -83.97582502 |
| Ingham County, Vevay township | 0.17334588 | 0.28294782 | 42.54841456 | -84.42589711 |
| Crawford County, Beaver Creek township | 0.11995697 | 0.28267886 | 44.58152214 | -84.711109 |
| Huron County, Oliver township | 0.20535234 | 0.28214094 | 43.80817516 | -83.17286203 |
| Leelanau County, Glen Arbor township | 0.24569661 | 0.28214094 | 45.03351088 | -86.13711205 |
| Alpena County, Ossineke township | 0.20938677 | 0.28187197 | 44.8690324 | -83.86442023 |
| Oceana County, Hart city | 0.28133405 | 0.28133405 | 43.6967668 | -86.36167755 |
| Houghton County, Elm River township | 0.21893491 | 0.28106509 | 46.82852657 | -88.78283552 |
| Isabella County, Coe township | 0.13636364 | 0.28079613 | 43.5192267 | -84.69906377 |
| Montcalm County, Maple Valley township | 0.13851533 | 0.28079613 | 43.30387627 | -85.35810676 |
| Wexford County, Clam Lake township | 0.13044648 | 0.28052717 | 44.23189536 | -85.4014499 |
| Menominee County, Cedarville township | 0.25013448 | 0.28052717 | 45.39339108 | -87.43152527 |
| Hillsdale County, Wright township | 0.20239376 | 0.27998924 | 41.79989693 | -84.36691623 |
| Chippewa County, Kinross charter township | 0.17616998 | 0.27998924 | 46.25884133 | -84.45566916 |
| Delta County, Fairbanks township | 0.21422808 | 0.27972028 | 45.56646709 | -86.63736851 |
| Presque Isle County, Metz township | 0.26492738 | 0.27945132 | 45.27305577 | -83.82296361 |
| Eaton County, Charlotte city | 0.17052179 | 0.27864443 | 42.56092586 | -84.8249241 |
| Marquette County, Forsyth township | 0.21153846 | 0.27864443 | 46.17177153 | -87.59004665 |
| Oscoda County, Greenwood township | 0.2260624 | 0.27837547 | 44.70226204 | -84.27686738 |
| Otsego County, Elmira township | 0.17038731 | 0.27837547 | 45.08274677 | -84.75448667 |
| Chippewa County, Bay Mills township | 0.16164605 | 0.27837547 | 46.43856579 | -84.97952583 |
| Ontonagon County, Bohemia township | 0.26169984 | 0.27810651 | 46.83699588 | -88.9455563 |
| Sanilac County, Greenleaf township | 0.16393222 | 0.27783755 | 43.59627011 | -83.08726219 |
| Mackinac County, Newton township | 0.22202797 | 0.27783755 | 46.03619305 | -85.63066043 |
| Oceana County, Crystal township | 0.22350726 | 0.27729962 | 43.74988487 | -86.16615976 |
| Barry County, Hastings charter township | 0.18625605 | 0.27703066 | 42.62170866 | -85.2780791 |
| Oceana County, Ferry township | 0.24744486 | 0.2767617 | 43.63845638 | -86.18223109 |
| Leelanau County, Cleveland township | 0.2534965 | 0.2767617 | 44.95001839 | -85.92609034 |
| Ottawa County, Polkton charter township | 0.13797741 | 0.27649274 | 43.01147894 | -85.93070969 |
| Leelanau County, Elmwood charter township | 0.22807961 | 0.27622378 | 44.79437155 | -85.662915 |
| Menominee County, Daggett township | 0.27057558 | 0.27622378 | 45.48903659 | -87.51556075 |
| Presque Isle County, Belknap township | 0.24341044 | 0.27622378 | 45.36512495 | -83.83830742 |
| Iosco County, Plainfield township | 0.22619688 | 0.27568585 | 44.50703423 | -83.86611933 |
| Tuscola County, Dayton township | 0.25201721 | 0.27541689 | 43.40858622 | -83.24798275 |
| Sanilac County, Evergreen township | 0.2197418 | 0.27541689 | 43.52056289 | -83.00755202 |
| Eaton County, Walton township | 0.11323292 | 0.27514793 | 42.45770681 | -84.93400242 |
| Shiawassee County, Shiawassee township | 0.20952125 | 0.27514793 | 42.87599119 | -84.0606087 |
| Chippewa County, Detour township | 0.24690694 | 0.27514793 | 46.02432599 | -84.10680885 |
| Alcona County, Alcona township | 0.19956966 | 0.27487897 | 44.77911675 | -83.62541282 |
| Mackinac County, Clark township | 0.23265196 | 0.27487897 | 45.97675834 | -84.34262221 |
| Menominee County, Mellen township | 0.23547606 | 0.27380312 | 45.38300471 | -87.62520559 |
| Ogemaw County, Logan township | 0.25605164 | 0.27353416 | 44.29300116 | -83.99961276 |
| Allegan County, Cheshire township | 0.21732114 | 0.2732652 | 42.45297979 | -85.98116478 |
| Alger County, Rock River township | 0.19889726 | 0.2732652 | 46.30775229 | -86.88988416 |
| Menominee County, Ingallston township | 0.22552448 | 0.27299623 | 45.36896751 | -87.40260553 |
| Alcona County, Mikado township | 0.19445939 | 0.27272727 | 44.53135661 | -83.63918365 |
| Dickinson County, Kingsford city | 0.23964497 | 0.27245831 | 45.79489415 | -88.08919378 |
| Chippewa County, Hulbert township | 0.20925229 | 0.27245831 | 46.27231299 | -85.17019313 |
| Ogemaw County, Mills township | 0.22646584 | 0.27165143 | 44.22624706 | -84.01147156 |
| Ogemaw County, Klacking township | 0.24825175 | 0.27165143 | 44.41833636 | -84.13592071 |
| Presque Isle County, Presque Isle township | 0.20172136 | 0.27165143 | 45.33310941 | -83.49020225 |
| Branch County, Batavia township | 0.17079075 | 0.27138246 | 41.92995193 | -85.07895889 |
| Montmorency County, Montmorency township | 0.21221087 | 0.27138246 | 45.15070036 | -83.97359716 |
| Chippewa County, Chippewa township | 0.23076923 | 0.27138246 | 46.39896617 | -85.05143721 |
| Mackinac County, Hudson township | 0.22834857 | 0.2711135 | 46.16545509 | -85.25275871 |
| Mackinac County, Moran township | 0.22619688 | 0.27084454 | 46.13026125 | -85.09598025 |
| Wexford County, Hanover township | 0.16110812 | 0.27057558 | 44.50183074 | -85.67606243 |
| Schoolcraft County, Inwood township | 0.22404519 | 0.27030662 | 46.18935618 | -86.48403197 |
| Manistee County, Springdale township | 0.23910705 | 0.26923077 | 44.46119855 | -86.03072451 |
| Muskegon County, Egelston township | 0.1788596 | 0.26923077 | 43.22823936 | -86.11657974 |
| Grand Traverse County, East Bay township | 0.17832168 | 0.26896181 | 44.73992529 | -85.54402356 |
| Muskegon County, Montague township | 0.22377622 | 0.26896181 | 43.42480969 | -86.33764373 |
| Montmorency County, Hillman township | 0.22417967 | 0.26869285 | 45.10611334 | -83.98162398 |
| Menominee County, Nadeau township | 0.22202797 | 0.26869285 | 45.51742592 | -87.60831024 |
| Alger County, Mathias township | 0.2084454 | 0.26869285 | 46.23393834 | -86.80259244 |
| Oceana County, Benona township | 0.24112426 | 0.26842388 | 43.63441704 | -86.43304092 |
| Eaton County, Eaton township | 0.11672942 | 0.26842388 | 42.55829845 | -84.79450189 |
| Eaton County, Bellevue township | 0.10731576 | 0.26788596 | 42.44325616 | -85.02868342 |
| Gladwin County, Gladwin township | 0.21073158 | 0.26788596 | 44.01202466 | -84.39621513 |
| Clinton County, Olive township | 0.10166756 | 0.267617 | 42.89503172 | -84.52257539 |
| Tuscola County, Wells township | 0.20589026 | 0.267617 | 43.41754181 | -83.34019924 |
| Arenac County, Deep River township | 0.13071544 | 0.267617 | 43.99815308 | -83.98876251 |
| Ingham County, Stockbridge township | 0.20118343 | 0.26734804 | 42.43491784 | -84.16758124 |
| Clare County, Harrison city | 0.24852071 | 0.26707907 | 44.02464437 | -84.8038263 |
| Muskegon County, Montague city | 0.25242066 | 0.26681011 | 43.40479961 | -86.37164313 |
| Presque Isle County, Case township | 0.22592792 | 0.26681011 | 45.24555441 | -84.0207307 |
| Iron County, Mansfield township | 0.21678322 | 0.26654115 | 46.22885148 | -88.16701282 |
| Ogemaw County, Foster township | 0.17052179 | 0.26600323 | 44.4461745 | -84.19448819 |
| Manistee County, Pleasanton township | 0.23830016 | 0.26519634 | 44.44589638 | -86.12796994 |
| Arenac County, Standish township | 0.14779451 | 0.26492738 | 43.95084874 | -83.96563235 |
| Otsego County, Corwith township | 0.15303927 | 0.26465842 | 45.12205143 | -84.68380859 |
| Newaygo County, Sheridan charter township | 0.19271114 | 0.26412049 | 43.46618235 | -85.98851859 |
| Genesee County, Montrose city | 0.26385153 | 0.26385153 | 43.17550759 | -83.88372719 |
| Sanilac County, Minden township | 0.2267348 | 0.26358257 | 43.60993574 | -82.77065314 |
| Huron County, Fairhaven township | 0.18370091 | 0.26358257 | 43.80689444 | -83.39343614 |
| Ogemaw County, Churchill township | 0.24098978 | 0.26277569 | 44.29582253 | -84.03078211 |
| Crawford County, Grayling city | 0.26250672 | 0.26250672 | 44.64838702 | -84.69988012 |
| Iron County, Crystal Falls township | 0.21274879 | 0.26250672 | 46.17064354 | -88.33599689 |
| Arenac County, Standish city | 0.26169984 | 0.26169984 | 43.9759142 | -83.99270446 |
| Eaton County, Benton township | 0.1344809 | 0.26143088 | 42.61964792 | -84.72278525 |
| Emmet County, Little Traverse township | 0.20710059 | 0.26143088 | 45.43344948 | -84.88828698 |
| Ingham County, White Oak township | 0.15734266 | 0.26143088 | 42.59302731 | -84.17134958 |
| Sanilac County, Argyle township | 0.18773534 | 0.26143088 | 43.51750664 | -82.97677896 |
| Alger County, Limestone township | 0.18531469 | 0.26143088 | 46.33019318 | -86.88678408 |
| Keweenaw County, Sherman township | 0.17711135 | 0.26143088 | 47.31438782 | -88.13538395 |
| Bay County, Monitor charter township | 0.16460463 | 0.26116192 | 43.58669082 | -83.93592933 |
| Mason County, Riverton township | 0.23843464 | 0.26116192 | 43.83052375 | -86.34272086 |
| Gogebic County, Erwin township | 0.24260355 | 0.26062399 | 46.41788249 | -90.12806731 |
| Ionia County, Ionia city | 0.20131791 | 0.26035503 | 42.98686033 | -85.06594333 |
| Marquette County, Republic township | 0.22538999 | 0.26035503 | 46.49475365 | -88.07021672 |
| Iosco County, Reno township | 0.21826251 | 0.25927918 | 44.3038918 | -83.87096304 |
| Alcona County, Hawes township | 0.18975256 | 0.25927918 | 44.73167218 | -83.60227007 |
| Clinton County, Lansing city | 0.25901022 | 0.25901022 | 42.78666373 | -84.57160104 |
| Leelanau County, Suttons Bay township | 0.22471759 | 0.25901022 | 45.03998716 | -85.62497966 |
| Saginaw County, Taymouth township | 0.17348037 | 0.2584723 | 43.30588641 | -83.82938635 |
| Sanilac County, Delaware township | 0.23265196 | 0.2584723 | 43.62276028 | -82.67390706 |
| Iosco County, Baldwin township | 0.21261431 | 0.2584723 | 44.32694863 | -83.39128399 |
| Benzie County, Gilmore township | 0.24529317 | 0.2584723 | 44.61141569 | -86.23064205 |
| Gladwin County, Clement township | 0.20952125 | 0.25820334 | 44.09830049 | -84.34985703 |
| Crawford County, South Branch township | 0.13797741 | 0.25820334 | 44.52626006 | -84.5933638 |
| Benzie County, Homestead township | 0.20629371 | 0.25820334 | 44.66443517 | -86.03237059 |
| Allegan County, Overisel township | 0.14362561 | 0.25766541 | 42.74683073 | -86.00023752 |
| Allegan County, Trowbridge township | 0.15465304 | 0.25739645 | 42.50543324 | -85.78885837 |
| Oceana County, Pentwater township | 0.24731038 | 0.25712749 | 43.79046382 | -86.41110086 |
| Benzie County, Crystal Lake township | 0.23883808 | 0.25712749 | 44.65144856 | -86.16132311 |
| Leelanau County, Kasson township | 0.21503497 | 0.25685853 | 44.78653737 | -85.85511837 |
| Ionia County, Portland township | 0.07813341 | 0.2563206 | 42.86019861 | -84.90021893 |
| Grand Traverse County, Peninsula township | 0.13152232 | 0.2563206 | 44.78702508 | -85.56853054 |
| Alpena County, Green township | 0.20925229 | 0.25605164 | 45.05802354 | -83.70192957 |
| Newaygo County, Beaver township | 0.18908015 | 0.25578268 | 43.67137459 | -86.02097579 |
| Calhoun County, Athens township | 0.2175901 | 0.25551372 | 42.08095975 | -85.26959319 |
| Midland County, Greendale township | 0.17751479 | 0.25551372 | 43.60017776 | -84.59166292 |
| Shiawassee County, Middlebury township | 0.14900484 | 0.25497579 | 43.00923336 | -84.28601941 |
| Ottawa County, Grand Haven charter township | 0.09776762 | 0.25470683 | 43.03478834 | -86.23743567 |
| Allegan County, Manlius township | 0.12170522 | 0.25443787 | 42.59579028 | -86.08373351 |
| Montcalm County, Reynolds township | 0.1633943 | 0.25443787 | 43.42510364 | -85.46408135 |
| Otsego County, Chester township | 0.17616998 | 0.25443787 | 45.02477576 | -84.60429378 |
| Antrim County, Kearney township | 0.16541151 | 0.25443787 | 44.98266824 | -85.18671352 |
| Menominee County, Harris township | 0.13071544 | 0.25443787 | 45.98222134 | -87.45236284 |
| Midland County, Larkin charter township | 0.15949435 | 0.25416891 | 43.68415149 | -84.26387163 |
| Benzie County, Almira township | 0.22270038 | 0.25416891 | 44.71955992 | -85.86522455 |
| Gogebic County, Bessemer city | 0.25403443 | 0.25416891 | 46.48260934 | -90.05725124 |
| Huron County, Sherman township | 0.21933835 | 0.25389995 | 43.73054201 | -82.62127769 |
| Charlevoix County, Wilson township | 0.13353954 | 0.25363098 | 45.19302764 | -84.9923911 |
| Kalamazoo County, Climax township | 0.18047337 | 0.25309306 | 42.24397465 | -85.33644528 |
| Otsego County, Hayes township | 0.10892953 | 0.25309306 | 45.01315767 | -84.73420039 |
| Muskegon County, Whitehall township | 0.22377622 | 0.25255514 | 43.4002552 | -86.30994149 |
| Keweenaw County, Allouez township | 0.16164605 | 0.25255514 | 47.39310035 | -88.32404363 |
| Newaygo County, Denver township | 0.20118343 | 0.25228618 | 43.60453519 | -86.03075558 |
| Mackinac County, Brevort township | 0.20145239 | 0.25228618 | 46.0027753 | -84.922328 |
| Delta County, Maple Ridge township | 0.18854223 | 0.25201721 | 46.12071095 | -87.14266198 |
| Hillsdale County, Jefferson township | 0.14671867 | 0.25174825 | 41.88537433 | -84.56658902 |
| Dickinson County, Felch township | 0.19203873 | 0.25174825 | 46.18552682 | -87.81538529 |
| Marquette County, Turin township | 0.21140398 | 0.25174825 | 46.20252898 | -87.35893627 |
| Baraga County, Baraga township | 0.21221087 | 0.25174825 | 46.61907139 | -88.67530202 |
| Midland County, Midland charter township | 0.18316299 | 0.25147929 | 43.59246105 | -84.24833379 |
| Tuscola County, Almer township | 0.16608392 | 0.25147929 | 43.5122748 | -83.38526797 |
| Osceola County, Evart city | 0.25147929 | 0.25147929 | 43.90490466 | -85.26385212 |
| Benzie County, Frankfort city | 0.25121033 | 0.25121033 | 44.63374416 | -86.2274017 |
| Leelanau County, Centerville township | 0.21570737 | 0.25121033 | 44.89578651 | -85.80671145 |
| Presque Isle County, Ocqueoc township | 0.21570737 | 0.25121033 | 45.47991068 | -84.10742198 |
| Montcalm County, Bloomer township | 0.18343195 | 0.25094137 | 43.17137173 | -84.84979549 |
| Iosco County, Tawas City city | 0.25094137 | 0.25094137 | 44.27204444 | -83.52827882 |
| Presque Isle County, Posen township | 0.20252824 | 0.25094137 | 45.26729554 | -83.75973536 |
| Iosco County, Wilber township | 0.22525551 | 0.2506724 | 44.41135439 | -83.56597251 |
| Delta County, Garden township | 0.18961807 | 0.25040344 | 46.14451273 | -86.4904893 |
| Iron County, Gaastra city | 0.25040344 | 0.25040344 | 46.0501444 | -88.60876168 |
| Presque Isle County, Pulawski township | 0.21382464 | 0.25013448 | 45.40108236 | -83.73512097 |
| Isabella County, Lincoln township | 0.11538462 | 0.24986552 | 43.54676927 | -84.75709045 |
| Tuscola County, Vassar city | 0.2493276 | 0.2493276 | 43.3715801 | -83.56760635 |
| Allegan County, Otsego city | 0.24878967 | 0.24878967 | 42.45374427 | -85.7041851 |
| Antrim County, Forest Home township | 0.17079075 | 0.24878967 | 44.98521524 | -85.21825007 |
| Benzie County, Blaine township | 0.23964497 | 0.24852071 | 44.55982364 | -86.14298857 |
| Gladwin County, Secord township | 0.21960732 | 0.24852071 | 44.07330217 | -84.3226048 |
| Mason County, Amber township | 0.21113502 | 0.24825175 | 43.94655209 | -86.38906028 |
| Luce County, Lakefield township | 0.20629371 | 0.24825175 | 46.32447256 | -85.84588426 |
| Luce County, Pentland township | 0.18908015 | 0.24825175 | 46.31997315 | -85.48824445 |
| Marquette County, Champion township | 0.17280796 | 0.24825175 | 46.75311977 | -87.84271586 |
| Manistee County, Cleon township | 0.21207639 | 0.24771382 | 44.47643029 | -85.93343522 |
| Ionia County, Odessa township | 0.08499193 | 0.24744486 | 42.78936668 | -85.12823318 |
| Ingham County, Williamston city | 0.22942442 | 0.2471759 | 42.6928688 | -84.27793098 |
| Mason County, Sherman township | 0.18477676 | 0.2471759 | 44.04664214 | -86.21753482 |
| Benzie County, Lake township | 0.22054868 | 0.24690694 | 44.67377731 | -86.15805791 |
| Mackinac County, Portage township | 0.21073158 | 0.24663798 | 46.23481866 | -85.85968058 |
| Mason County, Summit township | 0.23910705 | 0.24663798 | 43.85735363 | -86.40166724 |
| Cheboygan County, Forest township | 0.19782141 | 0.24663798 | 45.35717467 | -84.25799983 |
| Alpena County, Long Rapids township | 0.17509414 | 0.24610005 | 45.08030551 | -83.69782758 |
| Midland County, Ingersoll township | 0.14792899 | 0.24556213 | 43.56469273 | -84.19047394 |
| Jackson County, Waterloo township | 0.18854223 | 0.24502421 | 42.32120916 | -84.15624491 |
| Eaton County, Oneida charter township | 0.13125336 | 0.24502421 | 42.73335147 | -84.73510133 |
| Saginaw County, Tittabawassee township | 0.18719742 | 0.24502421 | 43.56191268 | -84.15963058 |
| Leelanau County, Empire township | 0.18020441 | 0.24502421 | 44.86067686 | -85.93949936 |
| Shiawassee County, Hazelton township | 0.17173211 | 0.24475524 | 43.08953662 | -83.93004717 |
| Huron County, Colfax township | 0.18343195 | 0.24475524 | 43.81520802 | -83.01409048 |
| Emmet County, Resort township | 0.19755245 | 0.24475524 | 45.3519381 | -84.99827341 |
| Osceola County, Marion township | 0.16863905 | 0.24448628 | 44.12257318 | -85.16624295 |
| Marquette County, Ely township | 0.14806347 | 0.24448628 | 46.25094507 | -87.77480771 |
| Marquette County, Ishpeming township | 0.10866057 | 0.24448628 | 46.49608177 | -87.71158497 |
| Chippewa County, Pickford township | 0.17186659 | 0.24394836 | 46.08300277 | -84.25883041 |
| Huron County, Verona township | 0.18692846 | 0.2436794 | 43.78995173 | -82.98744263 |
| Clinton County, DeWitt city | 0.24341044 | 0.24341044 | 42.83108594 | -84.56422193 |
| Presque Isle County, North Allis township | 0.2119419 | 0.24341044 | 45.41576591 | -84.15108045 |
| Manistee County, Onekama township | 0.20131791 | 0.24314147 | 44.40589855 | -86.19709835 |
| Branch County, Quincy township | 0.11484669 | 0.24260355 | 41.93924728 | -84.9246649 |
| Marquette County, Humboldt township | 0.20629371 | 0.24260355 | 46.32988584 | -87.96001048 |
| Shiawassee County, New Haven township | 0.18289403 | 0.24206563 | 43.12862709 | -84.10996749 |
| Chippewa County, Soo township | 0.15949435 | 0.24206563 | 46.45476773 | -84.35585897 |
| Gratiot County, North Shade township | 0.15465304 | 0.24206563 | 43.16874771 | -84.81910241 |
| Benzie County, Weldon township | 0.20723507 | 0.24125874 | 44.59746261 | -86.0422829 |
| Hillsdale County, Jonesville city | 0.24072082 | 0.24072082 | 41.98480181 | -84.67265467 |
| Branch County, Bronson city | 0.24018289 | 0.24018289 | 41.87100283 | -85.17979261 |
| Presque Isle County, Allis township | 0.21624529 | 0.24018289 | 45.3097756 | -84.23420914 |
| Presque Isle County, Onaway city | 0.24018289 | 0.24018289 | 45.35439977 | -84.22632623 |
| Barry County, Rutland charter township | 0.16783217 | 0.23991393 | 42.62673069 | -85.3390172 |
| Huron County, Port Austin township | 0.23318989 | 0.23991393 | 44.04032255 | -83.00246089 |
| Marquette County, Ewing township | 0.22189349 | 0.23991393 | 46.04136014 | -87.31520456 |
| Emmet County, Harbor Springs city | 0.23964497 | 0.23964497 | 45.44128693 | -84.98350752 |
| Charlevoix County, Boyne City city | 0.232383 | 0.23937601 | 45.2153542 | -84.98871011 |
| Saginaw County, Frankenmuth township | 0.18881119 | 0.23856912 | 43.322358 | -83.76398832 |
| Sanilac County, Marion township | 0.2267348 | 0.23856912 | 43.58463995 | -82.74417503 |
| Manistee County, Manistee city | 0.23856912 | 0.23856912 | 44.23426776 | -86.31662947 |
| Dickinson County, Norway city | 0.20736955 | 0.23776224 | 45.78352791 | -87.89712487 |
| Presque Isle County, Moltke township | 0.1936525 | 0.23749328 | 45.39315475 | -83.8975802 |
| Dickinson County, Sagola township | 0.19378698 | 0.23749328 | 46.19890991 | -88.04081622 |
| Kalamazoo County, Ross township | 0.11740183 | 0.23722431 | 42.33283865 | -85.32276906 |
| Presque Isle County, Bearinger township | 0.18128026 | 0.23722431 | 45.46039708 | -84.14312954 |
| Newaygo County, Wilcox township | 0.17697687 | 0.23695535 | 43.5807163 | -85.72154839 |
| Otsego County, Dover township | 0.18047337 | 0.23668639 | 45.04708251 | -84.60049315 |
| Ogemaw County, Richland township | 0.19499731 | 0.23641743 | 44.22342158 | -83.98033494 |
| Benzie County, Platte township | 0.20535234 | 0.23641743 | 44.72669623 | -85.95954635 |
| Eaton County, Hamlin township | 0.14470145 | 0.23587951 | 42.49797823 | -84.61933517 |
| Gladwin County, Buckeye township | 0.11323292 | 0.23561054 | 43.98704007 | -84.3689804 |
| Alger County, Onota township | 0.1873319 | 0.23507262 | 46.49377923 | -86.96198077 |
| Emmet County, Wawatam township | 0.13986014 | 0.2345347 | 45.76067868 | -84.73641825 |
| Huron County, Huron township | 0.21221087 | 0.23426573 | 44.02494825 | -82.84744929 |
| Branch County, Ovid township | 0.14443249 | 0.23426573 | 41.87783987 | -84.99550784 |
| Menominee County, Menominee township | 0.18060785 | 0.23426573 | 45.17947777 | -87.61871084 |
| Marquette County, Tilden township | 0.12721893 | 0.23399677 | 46.24703303 | -87.7103545 |
| Kent County, Spencer township | 0.14766003 | 0.23372781 | 43.279122 | -85.33080459 |
| Mecosta County, Mecosta township | 0.15599785 | 0.23372781 | 43.63287496 | -85.52587114 |
| Eaton County, Eaton Rapids township | 0.13528779 | 0.23345885 | 42.52551232 | -84.67647712 |
| Mason County, Custer township | 0.18881119 | 0.23345885 | 43.95973977 | -86.26148438 |
| Tuscola County, Vassar township | 0.20629371 | 0.23318989 | 43.3493791 | -83.57164854 |
| Manistee County, Arcadia township | 0.20199032 | 0.23292093 | 44.49517559 | -86.1841192 |
| Iosco County, Grant township | 0.19876278 | 0.23292093 | 44.31465831 | -83.74227225 |
| Kent County, Oakfield township | 0.13394298 | 0.232383 | 43.18762634 | -85.31393867 |
| Huron County, Rubicon township | 0.21315223 | 0.23211404 | 43.94592146 | -82.7365488 |
| Iosco County, East Tawas city | 0.22713825 | 0.23211404 | 44.28839519 | -83.46187588 |
| Newaygo County, Garfield township | 0.18464228 | 0.23184508 | 43.45908477 | -85.89592544 |
| Huron County, Pointe Aux Barques township | 0.23184508 | 0.23184508 | 44.05947202 | -82.96717392 |
| Chippewa County, Rudyard township | 0.17643895 | 0.23184508 | 46.22224899 | -84.5597978 |
| Tuscola County, Tuscola township | 0.15357719 | 0.23184508 | 43.37449761 | -83.59833967 |
| Saginaw County, Richland township | 0.15519096 | 0.23157612 | 43.40911913 | -84.21738262 |
| Schoolcraft County, Mueller township | 0.20037655 | 0.23130715 | 45.98598837 | -85.86538986 |
| Saginaw County, Frankenmuth city | 0.22162453 | 0.23130715 | 43.34168425 | -83.72928313 |
| Houghton County, Schoolcraft township | 0.16231845 | 0.23103819 | 47.19385973 | -88.4132917 |
| Manistee County, Manistee township | 0.17348037 | 0.23076923 | 44.27660744 | -86.27899169 |
| Alpena County, Wilson township | 0.15761162 | 0.23050027 | 45.05512026 | -83.67042043 |
| Alpena County, Sanborn township | 0.19741797 | 0.23023131 | 44.87575497 | -83.44772883 |
| Charlevoix County, Boyne Valley township | 0.18908015 | 0.23023131 | 45.19042631 | -84.96076553 |
| Branch County, Gilead township | 0.16581495 | 0.22996235 | 41.77966425 | -85.16346192 |
| Iosco County, Tawas township | 0.21113502 | 0.22996235 | 44.27497602 | -83.55941859 |
| Hillsdale County, Reading township | 0.14523938 | 0.22996235 | 41.81258381 | -84.76167193 |
| Otsego County, Charlton township | 0.20145239 | 0.22969338 | 44.96391888 | -84.42286219 |
| Tuscola County, Arbela township | 0.16393222 | 0.22969338 | 43.22200099 | -83.65717975 |
| Huron County, Sand Beach township | 0.2105971 | 0.22942442 | 43.78119209 | -82.6742756 |
| Newaygo County, Sherman township | 0.18074233 | 0.22915546 | 43.48135472 | -85.89263074 |
| Gladwin County, Butman township | 0.17939753 | 0.22915546 | 44.07878394 | -84.38477113 |
| Arenac County, Whitney township | 0.21032813 | 0.22861754 | 44.12224424 | -83.59213331 |
| Cheboygan County, Cheboygan city | 0.21732114 | 0.22834857 | 45.64706425 | -84.46509138 |
| Alpena County, Wellington township | 0.21113502 | 0.22834857 | 45.11413689 | -83.81986881 |
| Grand Traverse County, Paradise township | 0.14806347 | 0.22781065 | 44.58126806 | -85.53691374 |
| Houghton County, Stanton township | 0.1422808 | 0.22781065 | 47.16121382 | -88.64739124 |
| Houghton County, Calumet charter township | 0.15788058 | 0.22781065 | 47.24073193 | -88.44078909 |
| Allegan County, Clyde township | 0.15344271 | 0.2267348 | 42.59345717 | -86.05325092 |
| Huron County, Sigel township | 0.1753631 | 0.2267348 | 43.84733479 | -82.8818981 |
| Jackson County, Henrietta township | 0.13905325 | 0.22646584 | 42.39883624 | -84.26615316 |
| Calhoun County, Leroy township | 0.13111888 | 0.22619688 | 42.23643314 | -85.24554372 |
| Huron County, Sebewaing township | 0.17563206 | 0.22565896 | 43.74321589 | -83.43669425 |
| Benzie County, Inland township | 0.19593868 | 0.22565896 | 44.67967472 | -85.93478874 |
| Charlevoix County, Melrose township | 0.13959118 | 0.22565896 | 45.21275186 | -84.95707414 |
| Oceana County, Claybanks township | 0.21301775 | 0.22512103 | 43.52522337 | -86.47956086 |
| Antrim County, Central Lake township | 0.18208714 | 0.22512103 | 45.05726085 | -85.27060123 |
| Montcalm County, Richland township | 0.14564282 | 0.22458311 | 43.40157846 | -84.9058862 |
| Huron County, Harbor Beach city | 0.22458311 | 0.22458311 | 43.84776248 | -82.66118823 |
| Mackinac County, Marquette township | 0.18908015 | 0.22404519 | 46.04329127 | -84.59077967 |
| Delta County, Nahma township | 0.19324906 | 0.22377622 | 46.06377122 | -86.63176766 |
| Calhoun County, Convis township | 0.09870898 | 0.22350726 | 42.3492522 | -84.98207909 |
| Ottawa County, Robinson township | 0.1338085 | 0.22350726 | 43.02322333 | -86.08404676 |
| Iosco County, Sherman township | 0.20898332 | 0.22350726 | 44.22279686 | -83.72729536 |
| Grand Traverse County, Acme township | 0.14120495 | 0.2232383 | 44.75977288 | -85.50910542 |
| Menominee County, Meyer township | 0.1429532 | 0.2232383 | 45.71914122 | -87.58283934 |
| Clinton County, Dallas township | 0.12076385 | 0.22296934 | 43.00525609 | -84.75257885 |
| Sanilac County, Wheatland township | 0.20723507 | 0.22296934 | 43.58775086 | -82.7749678 |
| Tuscola County, Juniata township | 0.14456697 | 0.22270038 | 43.42938227 | -83.46319184 |
| Huron County, Lincoln township | 0.17401829 | 0.22270038 | 43.88181585 | -83.00132838 |
| Huron County, Gore township | 0.21315223 | 0.22270038 | 43.96811976 | -82.73219893 |
| Ogemaw County, Edwards township | 0.18840775 | 0.22270038 | 44.22633107 | -84.26452448 |
| Charlevoix County, Bay township | 0.17751479 | 0.22243141 | 45.3098621 | -85.03732408 |
| Allegan County, Watson township | 0.17186659 | 0.22216245 | 42.51801442 | -85.66381735 |
| Leelanau County, Bingham township | 0.19311458 | 0.22189349 | 44.93319965 | -85.70531547 |
| Bay County, Mount Forest township | 0.10005379 | 0.22162453 | 43.84531331 | -84.04726658 |
| Huron County, Meade township | 0.15909091 | 0.22135557 | 43.85961245 | -83.00558518 |
| Cheboygan County, Benton township | 0.17267348 | 0.22135557 | 45.62472799 | -84.46897154 |
| Huron County, Bloomfield township | 0.18786982 | 0.22108661 | 43.86644512 | -82.84667714 |
| Sanilac County, Austin township | 0.17200108 | 0.22108661 | 43.62231217 | -82.89389128 |
| Kent County, Vergennes township | 0.10086068 | 0.22081764 | 42.94796845 | -85.41318768 |
| Montcalm County, Pine township | 0.14806347 | 0.22081764 | 43.29885782 | -85.29656646 |
| Barry County, Irving township | 0.12264658 | 0.21920387 | 42.72313111 | -85.41680825 |
| Wexford County, Springville township | 0.18598709 | 0.21920387 | 44.41502888 | -85.72100188 |
| Newaygo County, Newaygo city | 0.19809037 | 0.21893491 | 43.40738979 | -85.80999391 |
| Mecosta County, Green charter township | 0.16890802 | 0.21893491 | 43.7912302 | -85.53285666 |
| Missaukee County, Caldwell township | 0.11188811 | 0.21893491 | 44.35304679 | -85.22419792 |
| Emmet County, Littlefield township | 0.14268424 | 0.21893491 | 45.44787901 | -84.78932847 |
| Emmet County, Bliss township | 0.18222162 | 0.21893491 | 45.75676863 | -84.96340695 |
| Allegan County, Valley township | 0.154922 | 0.21839699 | 42.55944567 | -85.90411827 |
| Missaukee County, Forest township | 0.1570737 | 0.21839699 | 44.35050478 | -85.192955 |
| Hillsdale County, Amboy township | 0.15720818 | 0.21732114 | 41.70523528 | -84.56560205 |
| Manistee County, Maple Grove township | 0.17132867 | 0.21732114 | 44.41421728 | -86.00601453 |
| Barry County, Woodland township | 0.08714363 | 0.21705218 | 42.76714109 | -85.13174059 |
| Kent County, Cedar Springs city | 0.21705218 | 0.21705218 | 43.22974364 | -85.55636894 |
| Oceana County, Otto township | 0.19002152 | 0.21678322 | 43.54932824 | -86.19505051 |
| Cheboygan County, Waverly township | 0.15559441 | 0.21678322 | 45.37948953 | -84.25406483 |
| Alcona County, Harrisville township | 0.16191501 | 0.21624529 | 44.6605875 | -83.32860072 |
| Tuscola County, Columbia township | 0.14954276 | 0.21597633 | 43.65437473 | -83.45310518 |
| Iosco County, Alabaster township | 0.20548682 | 0.21570737 | 44.16962467 | -83.64207281 |
| Huron County, Hume township | 0.16231845 | 0.21570737 | 44.02116442 | -83.03772801 |
| Crawford County, Lovells township | 0.14308768 | 0.21543841 | 44.69096927 | -84.40620947 |
| Iron County, Crystal Falls city | 0.19042496 | 0.21490048 | 46.10326522 | -88.34379756 |
| Midland County, Jerome township | 0.11229155 | 0.21463152 | 43.72553544 | -84.47647509 |
| Houghton County, Torch Lake township | 0.14093599 | 0.21463152 | 47.10731993 | -88.48908298 |
| Saginaw County, Spaulding township | 0.12829478 | 0.2140936 | 43.3616909 | -83.94442431 |
| Mason County, Eden township | 0.18719742 | 0.21355568 | 43.81906541 | -86.18752473 |
| Marquette County, Skandia township | 0.15761162 | 0.21328671 | 46.26161018 | -87.22123801 |
| Mason County, Hamlin township | 0.17186659 | 0.21301775 | 43.9979181 | -86.47611065 |
| Marquette County, Negaunee city | 0.18719742 | 0.21301775 | 46.51460662 | -87.64404502 |
| Eaton County, Sunfield township | 0.14066703 | 0.21274879 | 42.76202106 | -85.07069405 |
| Van Buren County, Columbia township | 0.1873319 | 0.21247983 | 42.34881481 | -86.08835003 |
| Eaton County, Potterville city | 0.21247983 | 0.21247983 | 42.62229231 | -84.75323165 |
| Clare County, Summerfield township | 0.16272189 | 0.21247983 | 44.1189704 | -84.85133357 |
| Clare County, Hatton township | 0.12721893 | 0.21140398 | 43.9774815 | -84.780122 |
| Delta County, Escanaba township | 0.10328133 | 0.21113502 | 45.8207805 | -87.05354362 |
| Antrim County, Mancelona township | 0.09951587 | 0.21086606 | 44.90546278 | -85.07151571 |
| Tuscola County, Akron township | 0.16030124 | 0.2105971 | 43.70174163 | -83.47578026 |
| Huron County, Dwight township | 0.19284562 | 0.2105971 | 44.01505309 | -82.97572924 |
| Missaukee County, Reeder township | 0.14981173 | 0.2105971 | 44.32821617 | -85.19652199 |
| Houghton County, Osceola township | 0.16581495 | 0.2105971 | 47.12984995 | -88.48649459 |
| Allegan County, Fennville city | 0.21032813 | 0.21032813 | 42.59811627 | -86.11421815 |
| Cheboygan County, Koehler township | 0.14591178 | 0.21005917 | 45.44336171 | -84.46819387 |
| Charlevoix County, Charlevoix city | 0.20979021 | 0.20979021 | 45.30545469 | -85.26276665 |
| Marquette County, Chocolay charter township | 0.12990855 | 0.20979021 | 46.49646968 | -87.3529439 |
| Midland County, Mount Haley township | 0.14483593 | 0.20952125 | 43.55625289 | -84.34853041 |
| Ionia County, Portland city | 0.18020441 | 0.20898332 | 42.85758767 | -84.8696621 |
| Saginaw County, St. Charles township | 0.15277031 | 0.20871436 | 43.22026137 | -84.12521627 |
| Huron County, Paris township | 0.1732114 | 0.2084454 | 43.70179203 | -82.78421104 |
| Cheboygan County, Nunda township | 0.13717052 | 0.20790748 | 45.22557544 | -84.56999702 |
| Iron County, Mastodon township | 0.13891877 | 0.20790748 | 46.05835323 | -88.34898921 |
| Gratiot County, Elba township | 0.13232921 | 0.20763852 | 43.13398346 | -84.42030782 |
| Arenac County, Turner township | 0.18665949 | 0.20763852 | 44.13671893 | -83.7745276 |
| Clinton County, Ovid township | 0.0979021 | 0.20736955 | 43.00339895 | -84.47339568 |
| Kalkaska County, Kalkaska township | 0.1070468 | 0.20710059 | 44.75442373 | -85.1597722 |
| Ionia County, Ionia township | 0.16272189 | 0.20683163 | 42.98428764 | -85.03532416 |
| Wexford County, Selma township | 0.12210866 | 0.20629371 | 44.25916455 | -85.46036992 |
| Iosco County, Burleigh township | 0.17805272 | 0.20629371 | 44.22855902 | -83.7895494 |
| Ottawa County, Port Sheldon township | 0.11484669 | 0.20548682 | 42.93883641 | -86.15814977 |
| Arenac County, Sims township | 0.1993007 | 0.20548682 | 44.05554743 | -83.63132776 |
| Benzie County, Colfax township | 0.19217321 | 0.20548682 | 44.53621836 | -85.8293705 |
| Lake County, Eden township | 0.17052179 | 0.20521786 | 44.10764285 | -85.83065446 |
| Huron County, Bingham township | 0.17872512 | 0.20521786 | 43.68888476 | -82.88104813 |
| Clinton County, Bengal township | 0.07127488 | 0.2049489 | 42.96973654 | -84.6032131 |
| Calhoun County, Burlington township | 0.16110812 | 0.2049489 | 42.11778613 | -85.1719938 |
| Ingham County, Wheatfield township | 0.12399139 | 0.2049489 | 42.67615821 | -84.3426119 |
| Roscommon County, Roscommon township | 0.16487359 | 0.2049489 | 44.22769498 | -84.80179235 |
| Houghton County, Hancock township | 0.16823561 | 0.2049489 | 47.18199912 | -88.61214314 |
| Chippewa County, Superior township | 0.12641205 | 0.20467994 | 46.36088372 | -84.86203045 |
| Allegan County, Lee township | 0.15922539 | 0.20467994 | 42.43309136 | -86.01479134 |
| Montcalm County, Bushnell township | 0.17307692 | 0.20467994 | 43.16214856 | -85.00686373 |
| Barry County, Prairieville township | 0.1422808 | 0.20441097 | 42.48356093 | -85.51503911 |
| Roscommon County, Richfield township | 0.13596019 | 0.20441097 | 44.31228872 | -84.47122635 |
| Montcalm County, Fairplain township | 0.15088757 | 0.20414201 | 43.17497715 | -85.16037278 |
| Montmorency County, Vienna township | 0.17159763 | 0.20360409 | 44.95844436 | -84.35987413 |
| Allegan County, Heath township | 0.13179129 | 0.20360409 | 42.6800925 | -86.00990687 |
| Jackson County, Concord township | 0.09467456 | 0.20333513 | 42.23424139 | -84.69332116 |
| Bay County, Garfield township | 0.1218397 | 0.20333513 | 43.82307916 | -84.05118065 |
| Arenac County, Au Gres township | 0.14120495 | 0.20333513 | 44.06135261 | -83.69342069 |
| Arenac County, Clayton township | 0.13501883 | 0.20333513 | 44.15950094 | -84.02330707 |
| Menominee County, Gourley township | 0.13071544 | 0.20333513 | 45.55222225 | -87.44323033 |
| Marquette County, Sands township | 0.1492738 | 0.20333513 | 46.49851542 | -87.38528054 |
| Oceana County, Greenwood township | 0.17065627 | 0.20306616 | 43.54469986 | -86.13323053 |
| Marquette County, West Branch township | 0.15586337 | 0.20306616 | 46.40252154 | -87.30014921 |
| Newaygo County, Everett township | 0.1788596 | 0.20252824 | 43.51149111 | -85.70072814 |
| Wexford County, Manton city | 0.20199032 | 0.20199032 | 44.41273848 | -85.40470548 |
| Mason County, Sheridan township | 0.16245293 | 0.20145239 | 43.99510171 | -86.13061764 |
| Gladwin County, Bourret township | 0.15451856 | 0.20145239 | 44.09280216 | -84.2876757 |
| Cheboygan County, Inverness township | 0.1281603 | 0.20118343 | 45.60239291 | -84.4728491 |
| Allegan County, Salem township | 0.12036041 | 0.20091447 | 42.6928891 | -85.88459078 |
| Kent County, Sparta township | 0.10785368 | 0.20091447 | 43.15048957 | -85.69276742 |
| Roscommon County, Denton township | 0.15882195 | 0.20091447 | 44.26429347 | -84.70084684 |
| Newaygo County, Big Prairie township | 0.13999462 | 0.20064551 | 43.52889664 | -85.63559611 |
| Huron County, Winsor township | 0.17079075 | 0.20064551 | 43.81718961 | -83.26560868 |
| Muskegon County, Moorland township | 0.1042227 | 0.20037655 | 43.2141653 | -85.93202639 |
| Cheboygan County, Wilmot township | 0.1486014 | 0.20037655 | 45.23363324 | -84.66488488 |
| Dickinson County, Norway township | 0.10274341 | 0.20037655 | 45.78160713 | -87.86511979 |
| Cheboygan County, Mackinaw township | 0.14093599 | 0.20037655 | 45.75801333 | -84.70453313 |
| Montcalm County, Sidney township | 0.1253362 | 0.19983862 | 43.22200451 | -85.18406086 |
| Clare County, Franklin township | 0.16998386 | 0.19983862 | 44.10841402 | -84.72686317 |
| Kent County, Bowne township | 0.09494352 | 0.19956966 | 42.85153405 | -85.33514863 |
| Lake County, Peacock township | 0.15586337 | 0.19956966 | 44.06065858 | -85.80621088 |
| Huron County, Chandler township | 0.16487359 | 0.19956966 | 43.87479734 | -83.16025631 |
| Newaygo County, Norwich township | 0.13609467 | 0.1993007 | 43.68232786 | -85.58091901 |
| Kalkaska County, Rapid River township | 0.14254976 | 0.1993007 | 44.7767319 | -85.1561705 |
| Clare County, Greenwood township | 0.13179129 | 0.19876278 | 44.05216644 | -84.86232652 |
| Saginaw County, Maple Grove township | 0.16958042 | 0.19768693 | 43.16744126 | -84.04092751 |
| Lake County, Newkirk township | 0.15922539 | 0.19768693 | 44.10524434 | -85.79950218 |
| Isabella County, Denver township | 0.13461538 | 0.19741797 | 43.65534111 | -84.7078139 |
| Hillsdale County, Moscow township | 0.11323292 | 0.19741797 | 42.03528555 | -84.48067565 |
| Jackson County, Parma township | 0.12587413 | 0.19661108 | 42.27331209 | -84.62549323 |
| Kent County, Nelson township | 0.11484669 | 0.19634212 | 43.22728474 | -85.52562367 |
| Muskegon County, Blue Lake township | 0.15277031 | 0.19472835 | 43.44023506 | -86.2418914 |
| Clare County, Sheridan township | 0.09185046 | 0.19472835 | 43.85824671 | -84.70550536 |
| Missaukee County, Lake township | 0.14806347 | 0.19472835 | 44.30846859 | -85.2313093 |
| Ingham County, Bunker Hill township | 0.13972566 | 0.19419042 | 42.44596903 | -84.28897121 |
| Mason County, Logan township | 0.17348037 | 0.19419042 | 43.81675193 | -86.15649178 |
| Gladwin County, Grim township | 0.13004303 | 0.19419042 | 44.04554147 | -84.26430571 |
| Muskegon County, Holton township | 0.13622916 | 0.1936525 | 43.40639431 | -86.09082382 |
| Hillsdale County, Ransom township | 0.12130178 | 0.1936525 | 41.7442151 | -84.49821464 |
| Montcalm County, Evergreen township | 0.10086068 | 0.19311458 | 43.21177734 | -85.06116153 |
| Roscommon County, Nester township | 0.16581495 | 0.19284562 | 44.17598876 | -84.46286606 |
| Newaygo County, Brooks township | 0.15599785 | 0.19257665 | 43.44952021 | -85.77249787 |
| Gladwin County, Gladwin city | 0.19029048 | 0.19257665 | 43.97567579 | -84.49695018 |
| Antrim County, Echo township | 0.13959118 | 0.19257665 | 45.05218319 | -85.20745893 |
| Gladwin County, Hay township | 0.1486014 | 0.19230769 | 43.98429962 | -84.33793849 |
| Shiawassee County, Fairfield township | 0.14981173 | 0.19203873 | 43.084097 | -84.36653411 |
| Muskegon County, Casnovia township | 0.09924691 | 0.19203873 | 43.21179444 | -85.90127487 |
| Mecosta County, Colfax township | 0.14523938 | 0.19150081 | 43.69224624 | -85.42273482 |
| Charlevoix County, Hudson township | 0.15182894 | 0.19123185 | 45.12737594 | -84.74697614 |
| Manistee County, Marilla township | 0.15626681 | 0.19096288 | 44.40711869 | -85.9121227 |
| Cheboygan County, Grant township | 0.14833244 | 0.19096288 | 45.49385489 | -84.26608912 |
| Tuscola County, Elmwood township | 0.14873588 | 0.18988704 | 43.61135263 | -83.24135627 |
| Wexford County, Cedar Creek township | 0.13340506 | 0.18988704 | 44.38794232 | -85.37693907 |
| Mecosta County, Grant township | 0.16554599 | 0.18934911 | 43.7590498 | -85.41234764 |
| Cheboygan County, Hebron township | 0.10691232 | 0.18934911 | 45.63825234 | -84.6280571 |
| Newaygo County, Lincoln township | 0.14362561 | 0.18854223 | 43.57281974 | -85.91034239 |
| Cheboygan County, Mullett township | 0.12963959 | 0.18854223 | 45.46840727 | -84.49606007 |
| Muskegon County, White River township | 0.14497041 | 0.18827327 | 43.40706097 | -86.40249701 |
| Missaukee County, Holland township | 0.16083916 | 0.1880043 | 44.19102268 | -84.902629 |
| Osceola County, Rose Lake township | 0.07665412 | 0.1880043 | 44.07340004 | -85.39472517 |
| Mason County, Victory township | 0.15868747 | 0.18773534 | 44.02893234 | -86.28301212 |
| Midland County, Lee township | 0.15895643 | 0.18773534 | 43.6034556 | -84.37179149 |
| Saginaw County, Swan Creek township | 0.15330823 | 0.18746638 | 43.37576621 | -84.09817281 |
| Shiawassee County, Rush township | 0.13367402 | 0.18719742 | 43.06756991 | -84.18275843 |
| Bay County, Merritt township | 0.10798817 | 0.18719742 | 43.50577707 | -83.79368707 |
| Roscommon County, Lake township | 0.10005379 | 0.18692846 | 44.36398716 | -84.8108406 |
| Gratiot County, Wheeler township | 0.11672942 | 0.18692846 | 43.40881467 | -84.46742339 |
| Newaygo County, Goodwell township | 0.14523938 | 0.18639053 | 43.5758657 | -85.65973295 |
| Chippewa County, Dafter township | 0.119688 | 0.18585261 | 46.41549509 | -84.42816033 |
| Calhoun County, Marengo township | 0.10919849 | 0.18558365 | 42.25524306 | -84.93560055 |
| Charlevoix County, Charlevoix township | 0.16191501 | 0.18531469 | 45.30798577 | -85.29445809 |
| Branch County, Union township | 0.14685315 | 0.18504572 | 42.04862498 | -85.15215299 |
| Montcalm County, Pierson township | 0.12331899 | 0.18477676 | 43.33855124 | -85.50858434 |
| Roscommon County, Backus township | 0.13246369 | 0.1845078 | 44.31499972 | -84.50242377 |
| Saginaw County, Brady township | 0.14187735 | 0.18423884 | 43.15641039 | -84.16742851 |
| Lake County, Webber township | 0.1605702 | 0.18316299 | 43.94920967 | -85.82294423 |
| Emmet County, Readmond township | 0.15814954 | 0.18316299 | 45.60548423 | -85.05303503 |
| Isabella County, Deerfield township | 0.1183432 | 0.18316299 | 43.60439494 | -84.90417322 |
| Delta County, Ford River township | 0.1105433 | 0.18289403 | 45.71293979 | -87.13251871 |
| Antrim County, Banks township | 0.13717052 | 0.18289403 | 45.17648126 | -85.34755872 |
| Menominee County, Faithorn township | 0.1246638 | 0.18289403 | 45.59448975 | -87.75937058 |
| Kent County, Grattan township | 0.11027434 | 0.18262507 | 43.03941647 | -85.43012664 |
| Crawford County, Maple Forest township | 0.10193653 | 0.18262507 | 44.85170399 | -84.69751355 |
| Wexford County, Liberty township | 0.10543303 | 0.18235611 | 44.45733559 | -85.39769736 |
| Ionia County, Ronald township | 0.15949435 | 0.18235611 | 43.11767922 | -85.01399262 |
| Arenac County, Au Gres city | 0.18208714 | 0.18208714 | 44.03911988 | -83.69746279 |
| Ionia County, North Plains township | 0.13259817 | 0.18208714 | 43.10990159 | -84.92197342 |
| Newaygo County, Troy township | 0.15411512 | 0.18154922 | 43.76285331 | -86.03890863 |
| Huron County, Sheridan township | 0.16030124 | 0.18154922 | 43.74860722 | -83.0268277 |
| Barry County, Orangeville township | 0.11592254 | 0.18128026 | 42.52554767 | -85.47786324 |
| Cheboygan County, Walker township | 0.14766003 | 0.1810113 | 45.30124947 | -84.39638498 |
| Cheboygan County, Aloha township | 0.15250134 | 0.1810113 | 45.4656875 | -84.46432398 |
| Kent County, Lowell city | 0.18047337 | 0.18047337 | 42.9429787 | -85.35196056 |
| Gratiot County, North Star township | 0.09319527 | 0.18020441 | 43.21416916 | -84.5625143 |
| Bay County, Kawkawlin township | 0.11847768 | 0.18020441 | 43.73134072 | -84.03589669 |
| Osceola County, Richmond township | 0.15613233 | 0.18020441 | 43.87786059 | -85.48808897 |
| Lake County, Ellsworth township | 0.11350188 | 0.17966649 | 44.05099875 | -85.6817045 |
| Huron County, McKinley township | 0.16581495 | 0.17966649 | 43.86161022 | -83.25726687 |
| Chippewa County, Sugar Island township | 0.09991931 | 0.17939753 | 46.34579186 | -84.1466248 |
| Antrim County, Torch Lake township | 0.13986014 | 0.17912856 | 45.1094823 | -85.35823008 |
| Hillsdale County, Allen township | 0.10032275 | 0.1788596 | 41.94307509 | -84.71002499 |
| Saginaw County, Fremont township | 0.15680473 | 0.17805272 | 43.38412528 | -84.19045203 |
| Osceola County, Reed City city | 0.17805272 | 0.17805272 | 43.88033107 | -85.51912632 |
| Saginaw County, Brant township | 0.15088757 | 0.17805272 | 43.27855866 | -84.27106854 |
| Arenac County, Mason township | 0.14174287 | 0.17778375 | 44.09511384 | -83.81363339 |
| Kalkaska County, Bear Lake township | 0.10879505 | 0.17778375 | 44.69175412 | -84.94717698 |
| Wexford County, Antioch township | 0.12775686 | 0.17724583 | 44.41260311 | -85.68971213 |
| Mecosta County, Austin township | 0.13313609 | 0.17643895 | 43.6229605 | -85.40218306 |
| Mason County, Free Soil township | 0.14187735 | 0.17643895 | 44.09124704 | -86.21109717 |
| Osceola County, Highland township | 0.10193653 | 0.17643895 | 44.12766011 | -85.22851972 |
| Huron County, Lake township | 0.14066703 | 0.17643895 | 43.94444537 | -83.17859809 |
| Branch County, Bethel township | 0.11161915 | 0.17616998 | 41.84626084 | -85.15311148 |
| Newaygo County, White Cloud city | 0.17616998 | 0.17616998 | 43.56086037 | -85.75580456 |
| Hillsdale County, Camden township | 0.12157074 | 0.17616998 | 41.79039599 | -84.76524657 |
| Manistee County, Stronach township | 0.11511565 | 0.17563206 | 44.2319827 | -86.2854023 |
| Emmet County, Center township | 0.10462614 | 0.17563206 | 45.61742761 | -84.92203049 |
| Arenac County, Adams township | 0.11377084 | 0.1753631 | 44.05108467 | -84.07404043 |
| Newaygo County, Croton township | 0.13421194 | 0.17482517 | 43.38798603 | -85.56338377 |
| Cheboygan County, Munro township | 0.11215707 | 0.17455621 | 45.61591195 | -84.6318822 |
| Clare County, Surrey township | 0.10328133 | 0.17428725 | 43.87144297 | -84.86052478 |
| Calhoun County, Homer township | 0.10973642 | 0.17401829 | 42.15337435 | -84.79850313 |
| Gladwin County, Sage township | 0.14214632 | 0.17401829 | 44.02018375 | -84.48939326 |
| Gladwin County, Sherman township | 0.15008069 | 0.17374933 | 44.13957745 | -84.56380838 |
| Delta County, Cornell township | 0.14281872 | 0.17374933 | 45.97611815 | -87.35610599 |
| Osceola County, Lincoln township | 0.15196342 | 0.17374933 | 43.9422119 | -85.44667243 |
| Allegan County, Saugatuck township | 0.14254976 | 0.17348037 | 42.67179693 | -86.19621403 |
| Emmet County, Cross Village township | 0.1605702 | 0.17348037 | 45.64759653 | -85.01382114 |
| Newaygo County, Grant township | 0.1161915 | 0.1732114 | 43.36046194 | -85.78581355 |
| Clare County, Hamilton township | 0.11403981 | 0.1732114 | 44.03896747 | -84.70689511 |
| Houghton County, Chassell township | 0.11403981 | 0.1732114 | 47.04505103 | -88.59475689 |
| Ionia County, Otisco township | 0.11403981 | 0.1732114 | 43.11586326 | -85.26296069 |
| Ingham County, Leslie township | 0.11067778 | 0.17294244 | 42.43742746 | -84.44447311 |
| Manistee County, Brown township | 0.15223238 | 0.17294244 | 44.3143265 | -86.17878186 |
| Saginaw County, Chapin township | 0.14349112 | 0.17294244 | 43.12852772 | -84.35899081 |
| Saginaw County, Blumfield township | 0.13394298 | 0.17267348 | 43.39471951 | -83.81354568 |
| Midland County, Warren township | 0.10892953 | 0.17240452 | 43.73094164 | -84.53833737 |
| Gratiot County, Washington township | 0.11861216 | 0.17213556 | 43.1919446 | -84.56622614 |
| Shiawassee County, Sciota township | 0.11511565 | 0.17213556 | 42.87323714 | -84.27818104 |
| Huron County, Brookfield township | 0.14254976 | 0.17186659 | 43.68990702 | -83.35229543 |
| Hillsdale County, Scipio township | 0.12789134 | 0.17186659 | 42.00699195 | -84.66904069 |
| Delta County, Bark River township | 0.12694997 | 0.17132867 | 45.72126763 | -87.26033106 |
| Ionia County, Lyons township | 0.11726735 | 0.17105971 | 43.02098227 | -84.93632066 |
| Cheboygan County, Ellis township | 0.12157074 | 0.17105971 | 45.29522494 | -84.59017697 |
| Midland County, Geneva township | 0.14133943 | 0.17079075 | 43.64466 | -84.58420904 |
| Delta County, Gladstone city | 0.15559441 | 0.17079075 | 45.84107346 | -87.01851332 |
| Lake County, Pleasant Plains township | 0.12694997 | 0.17052179 | 43.88474041 | -85.86400966 |
| Wexford County, South Branch township | 0.09763314 | 0.17025282 | 44.17453092 | -85.82060522 |
| Branch County, Algansee township | 0.0718128 | 0.16998386 | 41.89485681 | -84.93172147 |
| Charlevoix County, Hayes township | 0.09817106 | 0.16998386 | 45.30291617 | -85.2310776 |
| Mason County, Meade township | 0.13071544 | 0.16917698 | 44.0866214 | -86.14878338 |
| Crawford County, Frederic township | 0.10825713 | 0.16917698 | 44.85436505 | -84.72896992 |
| Saginaw County, Albee township | 0.12130178 | 0.16890802 | 43.22559885 | -83.93722152 |
| Ottawa County, Chester township | 0.0809575 | 0.16863905 | 43.18953432 | -85.90455387 |
| Tuscola County, Ellington township | 0.15438408 | 0.16863905 | 43.50633164 | -83.32369872 |
| Osceola County, Middle Branch township | 0.08620226 | 0.16863905 | 44.03091749 | -85.14938944 |
| Roscommon County, Lyon township | 0.10328133 | 0.16756321 | 44.49767982 | -84.78862987 |
| Lake County, Sauble township | 0.13986014 | 0.16729424 | 44.00568407 | -85.97178868 |
| Delta County, Brampton township | 0.09870898 | 0.16702528 | 45.93073831 | -87.00636101 |
| Delta County, Baldwin township | 0.10610543 | 0.16702528 | 46.07164045 | -87.08442274 |
| Clinton County, Duplain township | 0.12721893 | 0.16675632 | 43.10904249 | -84.39341123 |
| Charlevoix County, Evangeline township | 0.10731576 | 0.1662184 | 45.23768187 | -84.98502665 |
| Osceola County, Hersey township | 0.11928456 | 0.1662184 | 43.89517242 | -85.4225528 |
| Newaygo County, Ensley township | 0.11094675 | 0.16568047 | 43.365728 | -85.56678074 |
| Antrim County, Custer township | 0.12923615 | 0.16568047 | 44.88572297 | -85.10663806 |
| Huron County, Grant township | 0.1401291 | 0.16541151 | 43.76074728 | -83.15036182 |
| Emmet County, Friendship township | 0.10664336 | 0.16541151 | 45.51866863 | -85.09952814 |
| Barry County, Barry township | 0.11403981 | 0.16487359 | 42.49836647 | -85.42040537 |
| Alcona County, Greenbush township | 0.14779451 | 0.16460463 | 44.57457287 | -83.37664656 |
| Missaukee County, Butterfield township | 0.13313609 | 0.1640667 | 44.25784852 | -84.89165323 |
| Alpena County, Maple Ridge township | 0.13636364 | 0.16379774 | 45.19939018 | -83.51527995 |
| Mason County, Grant township | 0.13219473 | 0.16325982 | 44.14274438 | -86.29819828 |
| Kalkaska County, Excelsior township | 0.14658419 | 0.16325982 | 44.74672357 | -85.06549998 |
| Allegan County, Monterey township | 0.08956428 | 0.16299086 | 42.60154767 | -85.86713762 |
| Charlevoix County, Chandler township | 0.09696073 | 0.16299086 | 45.21930267 | -84.76355409 |
| Mecosta County, Morton township | 0.09924691 | 0.16272189 | 43.60787187 | -85.21671866 |
| Gratiot County, Sumner township | 0.0661646 | 0.16245293 | 43.34132918 | -84.72846949 |
| Arenac County, Arenac township | 0.1070468 | 0.16218397 | 44.01472444 | -83.92273779 |
| Mason County, Branch township | 0.13717052 | 0.16164605 | 43.9059199 | -86.14357163 |
| Ingham County, Aurelius township | 0.09171598 | 0.16164605 | 42.55383447 | -84.4867069 |
| Manistee County, Norman township | 0.12614309 | 0.16137708 | 44.2333919 | -86.00115775 |
| Charlevoix County, South Arm township | 0.10220549 | 0.16137708 | 45.1191609 | -85.19664626 |
| Newaygo County, Ashland township | 0.10852609 | 0.16110812 | 43.36286043 | -85.81662766 |
| Osceola County, Cedar township | 0.10301237 | 0.16030124 | 43.93972414 | -85.41560954 |
| Missaukee County, Clam Union township | 0.11067778 | 0.16003228 | 44.17655217 | -84.99973555 |
| Barry County, Baltimore township | 0.11753631 | 0.15949435 | 42.59443567 | -85.22062003 |
| Huron County, Caseville township | 0.13394298 | 0.15949435 | 43.90902882 | -83.27987474 |
| Manistee County, Dickson township | 0.12022593 | 0.15949435 | 44.30266419 | -86.02250662 |
| Ionia County, Sebewa township | 0.08727811 | 0.15922539 | 42.78424485 | -85.06716729 |
| Tuscola County, Wisner township | 0.11538462 | 0.15922539 | 43.62168981 | -83.58465392 |
| Manistee County, Filer charter township | 0.1366326 | 0.15895643 | 44.21423283 | -86.35104058 |
| Gratiot County, New Haven township | 0.12103281 | 0.15868747 | 43.21321 | -84.81184225 |
| Tuscola County, Fairgrove township | 0.11915008 | 0.15868747 | 43.56554574 | -83.46947354 |
| Ionia County, Easton township | 0.08122647 | 0.1584185 | 43.01165795 | -85.09303472 |
| Jackson County, Pulaski township | 0.1042227 | 0.15814954 | 42.11528341 | -84.62071377 |
| Newaygo County, Barton township | 0.10220549 | 0.1570737 | 43.74914255 | -85.57068803 |
| Lake County, Lake township | 0.12694997 | 0.15680473 | 43.85200074 | -86.02584504 |
| Emmet County, Springvale township | 0.09494352 | 0.15680473 | 45.38352079 | -84.8323114 |
| Barry County, Yankee Springs township | 0.11161915 | 0.15680473 | 42.68115023 | -85.45412717 |
| Clinton County, Lebanon township | 0.09965035 | 0.15680473 | 43.10205947 | -84.82997508 |
| Missaukee County, West Branch township | 0.14241528 | 0.15680473 | 44.40969769 | -85.08843397 |
| Ionia County, Danby township | 0.07557827 | 0.15626681 | 42.83274971 | -84.84271126 |
| Kalkaska County, Coldsprings township | 0.13179129 | 0.15626681 | 44.78875165 | -85.02679284 |
| Midland County, Jasper township | 0.10059172 | 0.15626681 | 43.54762377 | -84.50654749 |
| Osceola County, Osceola township | 0.10247445 | 0.15599785 | 43.90237423 | -85.23281296 |
| Isabella County, Vernon township | 0.09090909 | 0.15572889 | 43.74963564 | -84.75493193 |
| Gladwin County, Grout township | 0.07208176 | 0.15572889 | 43.97837897 | -84.52799452 |
| Alcona County, Haynes township | 0.13636364 | 0.15572889 | 44.70209836 | -83.28886833 |
| Emmet County, Carp Lake township | 0.1218397 | 0.15572889 | 45.67922192 | -84.84713028 |
| Eaton County, Carmel township | 0.07947821 | 0.15545992 | 42.54133511 | -84.85893196 |
| Dickinson County, Waucedah township | 0.11672942 | 0.15545992 | 45.93476051 | -87.78169571 |
| Alcona County, Gustin township | 0.1366326 | 0.15519096 | 44.64725527 | -83.42670758 |
| Muskegon County, Sullivan township | 0.10745024 | 0.154922 | 43.20364648 | -86.08903557 |
| Lake County, Dover township | 0.13300161 | 0.154922 | 44.09314235 | -85.64377393 |
| Emmet County, West Traverse township | 0.1274879 | 0.154922 | 45.44388453 | -85.01525265 |
| Missaukee County, Aetna township | 0.12923615 | 0.15330823 | 44.26566315 | -84.985227 |
| Antrim County, Elk Rapids township | 0.13636364 | 0.15303927 | 44.86390319 | -85.3971769 |
| Delta County, Bay de Noc township | 0.10018827 | 0.15303927 | 45.80360957 | -86.79762249 |
| Mecosta County, Aetna township | 0.07409898 | 0.15277031 | 43.49435559 | -85.48463632 |
| Barry County, Hope township | 0.12452932 | 0.15250134 | 42.58477781 | -85.37633134 |
| Kalkaska County, Orange township | 0.11188811 | 0.15250134 | 44.64289812 | -85.17774505 |
| Missaukee County, Norwich township | 0.11417429 | 0.15250134 | 44.42682962 | -85.02228136 |
| Newaygo County, Merrill township | 0.14147391 | 0.15250134 | 43.70170777 | -85.82859617 |
| Missaukee County, Riverside township | 0.12210866 | 0.15223238 | 44.21678702 | -85.21432183 |
| Calhoun County, Clarendon township | 0.09736417 | 0.15223238 | 42.15599946 | -84.82874891 |
| Charlevoix County, Norwood township | 0.12372243 | 0.15223238 | 45.22366511 | -85.37208715 |
| Muskegon County, Cedar Creek township | 0.12977407 | 0.1514255 | 43.31963468 | -86.13452414 |
| Wexford County, Cherry Grove township | 0.05796127 | 0.1514255 | 44.23687306 | -85.46384433 |
| Tuscola County, Gilford township | 0.11565358 | 0.15115654 | 43.48843417 | -83.60892318 |
| Isabella County, Nottawa township | 0.12412587 | 0.15115654 | 43.64627978 | -84.86601293 |
| Newaygo County, Home township | 0.10946746 | 0.15088757 | 43.74386541 | -85.79095005 |
| Bay County, Beaver township | 0.10516407 | 0.15088757 | 43.73415056 | -84.06681147 |
| Charlevoix County, Marion township | 0.12641205 | 0.15088757 | 45.28311579 | -85.26636126 |
| Barry County, Carlton township | 0.0837816 | 0.15061861 | 42.69090231 | -85.29820999 |
| Gladwin County, Bentley township | 0.11000538 | 0.15034965 | 43.86481575 | -84.26408809 |
| Midland County, Coleman city | 0.15008069 | 0.15008069 | 43.75856539 | -84.59647905 |
| Missaukee County, McBain city | 0.15008069 | 0.15008069 | 44.19450394 | -85.2178748 |
| Calhoun County, Lee township | 0.06186122 | 0.14981173 | 42.4106876 | -84.91074369 |
| Roscommon County, Markey township | 0.10718128 | 0.14981173 | 44.39259209 | -84.61596906 |
| Gratiot County, Fulton township | 0.12157074 | 0.14954276 | 43.15817996 | -84.69635358 |
| Midland County, Porter township | 0.11309844 | 0.14954276 | 43.54491774 | -84.47569915 |
| Ingham County, Onondaga township | 0.10220549 | 0.1492738 | 42.47310144 | -84.5926134 |
| Hillsdale County, Woodbridge township | 0.12856374 | 0.14873588 | 41.7629258 | -84.70866868 |
| Kalkaska County, Springfield township | 0.10435718 | 0.14873588 | 44.58107593 | -85.25119664 |
| Lake County, Elk township | 0.13824637 | 0.14819796 | 44.1664824 | -86.01103797 |
| Isabella County, Wise township | 0.08055406 | 0.14792899 | 43.78349241 | -84.62370579 |
| Lake County, Cherry Valley township | 0.10086068 | 0.14792899 | 43.96909724 | -85.7885123 |
| Mackinac County, Mackinac Island city | 0.14766003 | 0.14766003 | 45.86441624 | -84.62159498 |
| Saginaw County, Jonesfield township | 0.1225121 | 0.14739107 | 43.4396206 | -84.30591393 |
| Lake County, Sweetwater township | 0.11040882 | 0.14712211 | 43.98339076 | -85.97508284 |
| Saginaw County, Marion township | 0.13206025 | 0.14712211 | 43.23687294 | -84.30938628 |
| Newaygo County, Lilley township | 0.11672942 | 0.14658419 | 43.74626429 | -85.82193731 |
| Clare County, Winterfield township | 0.1007262 | 0.14658419 | 44.15168394 | -84.97221351 |
| Kalkaska County, Oliver township | 0.11377084 | 0.14631522 | 44.6720564 | -84.98222452 |
| Clare County, Garfield township | 0.10529855 | 0.1457773 | 43.83993225 | -85.02280247 |
| Branch County, California township | 0.10825713 | 0.14550834 | 41.79565253 | -84.82542465 |
| Cheboygan County, Beaugrand township | 0.12493276 | 0.14497041 | 45.63556239 | -84.59623484 |
| Clare County, Lincoln township | 0.12910167 | 0.14443249 | 43.9853703 | -84.87329792 |
| Lake County, Chase township | 0.11484669 | 0.14443249 | 43.84314139 | -85.61906141 |
| Clare County, Arthur township | 0.07598171 | 0.14416353 | 43.90275645 | -84.69809758 |
| Missaukee County, Pioneer township | 0.10718128 | 0.14389457 | 44.43455499 | -85.11610356 |
| Ionia County, Belding city | 0.14389457 | 0.14389457 | 43.0910968 | -85.23576719 |
| Ionia County, Orange township | 0.11592254 | 0.14362561 | 42.89537326 | -85.04949943 |
| Tuscola County, Denmark township | 0.0929263 | 0.14362561 | 43.39670036 | -83.59430463 |
| Gladwin County, Billings township | 0.12130178 | 0.14335664 | 43.87032276 | -84.32605976 |
| Missaukee County, Richland township | 0.10018827 | 0.14281872 | 44.24917622 | -85.33554976 |
| Eaton County, Roxand township | 0.07046799 | 0.14201183 | 42.75169548 | -84.94862821 |
| Montcalm County, Winfield township | 0.06374395 | 0.14201183 | 43.42262231 | -85.43325184 |
| Wexford County, Colfax township | 0.09655729 | 0.14174287 | 44.35082235 | -85.47770072 |
| Mackinac County, Bois Blanc township | 0.12318451 | 0.14174287 | 45.77837245 | -84.40987283 |
| Osceola County, Evart township | 0.11135019 | 0.14093599 | 43.88263385 | -85.26737267 |
| Ionia County, Orleans township | 0.10287789 | 0.14093599 | 43.03645091 | -85.1201456 |
| Isabella County, Sherman township | 0.07907477 | 0.14066703 | 43.67895104 | -84.98606462 |
| Midland County, Mills township | 0.12318451 | 0.14066703 | 43.77585533 | -84.27945121 |
| Midland County, Hope township | 0.13152232 | 0.14039806 | 43.82583688 | -84.33370655 |
| Osceola County, Le Roy township | 0.08579882 | 0.13986014 | 44.05857379 | -85.49155774 |
| Kent County, Tyrone township | 0.06078537 | 0.13959118 | 43.26176358 | -85.67601765 |
| Allegan County, Saugatuck city | 0.13905325 | 0.13905325 | 42.64954782 | -86.19937281 |
| Newaygo County, Monroe township | 0.11807423 | 0.13878429 | 43.72158761 | -85.79429055 |
| Calhoun County, Fredonia township | 0.09090909 | 0.13851533 | 42.23821678 | -84.99971598 |
| Gladwin County, Beaverton city | 0.13797741 | 0.13797741 | 43.88396357 | -84.48103248 |
| Kalkaska County, Clearwater township | 0.06670253 | 0.13717052 | 44.78183047 | -85.21905109 |
| Marquette County, Richmond township | 0.10032275 | 0.13690156 | 46.46569498 | -87.58508606 |
| Clare County, Freeman township | 0.11403981 | 0.1366326 | 43.95125284 | -85.00478811 |
| Charlevoix County, East Jordan city | 0.13609467 | 0.13609467 | 45.15870106 | -85.12618945 |
| Cheboygan County, Tuscarora township | 0.08203335 | 0.13582571 | 45.38720882 | -84.606581 |
| Delta County, Ensign township | 0.08969876 | 0.13555675 | 45.87949042 | -86.91636538 |
| Montcalm County, Stanton city | 0.13555675 | 0.13555675 | 43.30330879 | -85.07769671 |
| Eaton County, Chester township | 0.08579882 | 0.13474987 | 42.63540766 | -84.90549838 |
| Barry County, Maple Grove township | 0.05648198 | 0.1344809 | 42.59190064 | -85.19016838 |
| Gratiot County, Lafayette township | 0.10126412 | 0.1344809 | 43.37842344 | -84.37886011 |
| Saginaw County, Lakefield township | 0.112695 | 0.1344809 | 43.37294607 | -84.31732804 |
| Wexford County, Henderson township | 0.09064013 | 0.13421194 | 44.18474526 | -85.66130971 |
| Roscommon County, Au Sable township | 0.11242604 | 0.13421194 | 44.50727002 | -84.37432689 |
| Osceola County, Burdell township | 0.09467456 | 0.13394298 | 44.08579365 | -85.55036373 |
| Kalkaska County, Garfield township | 0.09467456 | 0.13367402 | 44.51079472 | -84.94513133 |
| Lake County, Yates township | 0.08687466 | 0.13340506 | 43.83298488 | -85.77756624 |
| Isabella County, Fremont township | 0.09306079 | 0.13286713 | 43.46830605 | -84.8950399 |
| Wexford County, Greenwood township | 0.11417429 | 0.13286713 | 44.4920396 | -85.55076291 |
| Chippewa County, Bruce township | 0.10745024 | 0.13125336 | 46.25911498 | -84.19482693 |
| Eaton County, Vermontville township | 0.08445401 | 0.1309844 | 42.62355365 | -85.03088548 |
| Montcalm County, Cato township | 0.06993007 | 0.13044648 | 43.43235973 | -85.27564543 |
| Osceola County, Orient township | 0.09561592 | 0.13044648 | 43.89983654 | -85.2017761 |
| Missaukee County, Enterprise township | 0.09265734 | 0.12990855 | 44.3770725 | -84.96703765 |
| Gladwin County, Tobacco township | 0.08015062 | 0.12963959 | 43.8313109 | -84.39564841 |
| Antrim County, Milton township | 0.10045724 | 0.12963959 | 44.95318968 | -85.38304731 |
| Osceola County, Sherman township | 0.0746369 | 0.12937063 | 44.09568242 | -85.39123646 |
| Grand Traverse County, Union township | 0.0866057 | 0.12856374 | 44.61845097 | -85.43584087 |
| Hillsdale County, Litchfield township | 0.07046799 | 0.12829478 | 42.01229468 | -84.72939808 |
| Newaygo County, Bridgeton township | 0.09736417 | 0.12829478 | 43.35247687 | -85.97399458 |
| Kalkaska County, Blue Lake township | 0.09642281 | 0.12829478 | 44.78095724 | -84.93248797 |
| Calhoun County, Newton township | 0.059844 | 0.12802582 | 42.22882798 | -85.15466193 |
| Cheboygan County, Mentor township | 0.07544379 | 0.12802582 | 45.36488341 | -84.61039891 |
| Grand Traverse County, Whitewater township | 0.10193653 | 0.1274879 | 44.7275063 | -85.38687641 |
| Gratiot County, Hamilton township | 0.11081226 | 0.12721893 | 43.22286091 | -84.40527 |
| Missaukee County, Bloomfield township | 0.06939215 | 0.12694997 | 44.4275118 | -85.30734747 |
| Montcalm County, Ferris township | 0.08472297 | 0.12668101 | 43.37672646 | -84.8787086 |
| Gratiot County, Seville township | 0.08405056 | 0.12641205 | 43.44082797 | -84.83702423 |
| Antrim County, Chestonia township | 0.1035503 | 0.12614309 | 44.95009078 | -85.0642352 |
| Kalkaska County, Boardman township | 0.09870898 | 0.12560516 | 44.65051753 | -85.27188505 |
| Antrim County, Star township | 0.08526089 | 0.1253362 | 45.02896748 | -84.92343777 |
| Ingham County, Ingham township | 0.09037117 | 0.12452932 | 42.55693503 | -84.27014418 |
| Montcalm County, Crystal township | 0.07961269 | 0.12426036 | 43.21583591 | -84.84255489 |
| Barry County, Johnstown township | 0.08714363 | 0.12345347 | 42.44141641 | -85.24484919 |
| Clare County, Redding township | 0.08230231 | 0.12291555 | 44.04806657 | -85.08362019 |
| Wexford County, Slagle township | 0.06535772 | 0.12210866 | 44.33063527 | -85.79707986 |
| Midland County, Edenville township | 0.10462614 | 0.12130178 | 43.78954842 | -84.43420028 |
| Jackson County, Springport township | 0.059844 | 0.11995697 | 42.41183597 | -84.6642937 |
| Jackson County, Tompkins township | 0.07517483 | 0.11995697 | 42.37354778 | -84.4859175 |
| Bay County, Gibson township | 0.09037117 | 0.119688 | 43.9815494 | -84.05475507 |
| Lake County, Pinora township | 0.09077461 | 0.11861216 | 43.97437195 | -85.56752172 |
| Mecosta County, Wheatland township | 0.07356105 | 0.1183432 | 43.62758908 | -85.18228309 |
| Grand Traverse County, Fife Lake township | 0.10287789 | 0.11807423 | 44.58864263 | -85.34525876 |
| Clinton County, Victor township | 0.10247445 | 0.11753631 | 42.86198295 | -84.40410518 |
| Antrim County, Warner township | 0.10435718 | 0.11753631 | 45.07098179 | -84.88447909 |
| Isabella County, Broomfield township | 0.08821947 | 0.11699839 | 43.63444378 | -84.99326091 |
| Barry County, Castleton township | 0.05473373 | 0.11699839 | 42.61412224 | -85.18668856 |
| Ionia County, Campbell township | 0.04841313 | 0.11672942 | 42.84902486 | -85.3045795 |
| Montcalm County, Home township | 0.07880581 | 0.1161915 | 43.40678068 | -84.96748962 |
| Emmet County, McKinley township | 0.08821947 | 0.11377084 | 45.63188863 | -84.82280633 |
| Arenac County, Omer city | 0.11296396 | 0.11296396 | 44.05350066 | -83.85269135 |
| Emmet County, Maple River township | 0.08687466 | 0.11215707 | 45.47285844 | -84.8173167 |
| Montcalm County, Douglass township | 0.07988166 | 0.11188811 | 43.31097944 | -85.16999242 |
| Mecosta County, Chippewa township | 0.08068854 | 0.11188811 | 43.76623544 | -85.22298481 |
| Osceola County, Hartwick township | 0.06791286 | 0.11161915 | 44.06082431 | -85.23914373 |
| Gratiot County, Newark township | 0.07719204 | 0.11027434 | 43.22751861 | -84.71605547 |
| Mecosta County, Deerfield township | 0.0443787 | 0.11027434 | 43.53391354 | -85.41601384 |
| Osceola County, Sylvan township | 0.06239914 | 0.11027434 | 43.92210633 | -85.19823324 |
| Branch County, Butler township | 0.05823023 | 0.10973642 | 42.00583652 | -84.91406324 |
| Antrim County, Helena township | 0.07315761 | 0.10973642 | 44.91825693 | -85.22901974 |
| Mecosta County, Martiny township | 0.09265734 | 0.10892953 | 43.65238907 | -85.20967058 |
| Mecosta County, Sheridan township | 0.07423346 | 0.10892953 | 43.67210601 | -85.17521056 |
| Clinton County, Essex township | 0.09709521 | 0.10758472 | 43.0612892 | -84.61908211 |
| Emmet County, Pleasantview township | 0.07140936 | 0.10758472 | 45.48074671 | -84.91258507 |
| Wexford County, Boon township | 0.07315761 | 0.1070468 | 44.33837943 | -85.60616155 |
| Isabella County, Gilmore township | 0.08606778 | 0.10462614 | 43.78502765 | -84.90608339 |
| Calhoun County, Eckford township | 0.08391608 | 0.10381926 | 42.22521892 | -84.84829916 |
| Antrim County, Jordan township | 0.07517483 | 0.10274341 | 45.10632241 | -85.03867718 |
| Hillsdale County, Litchfield city | 0.10220549 | 0.10220549 | 42.03712884 | -84.75599227 |
| Charlevoix County, Eveline township | 0.0683163 | 0.10220549 | 45.23336287 | -85.21023011 |
| Mecosta County, Fork township | 0.07988166 | 0.10112964 | 43.78341242 | -85.15748888 |
| Montcalm County, Day township | 0.07880581 | 0.09817106 | 43.31781342 | -84.98183992 |
| Ionia County, Boston township | 0.05406132 | 0.09628833 | 42.93796033 | -85.29074231 |
| Eaton County, Brookfield township | 0.07235073 | 0.09601937 | 42.4472581 | -84.81249163 |
| Mecosta County, Hinton township | 0.04007531 | 0.09601937 | 43.54867053 | -85.31990637 |
| Gratiot County, Ithaca city | 0.09373319 | 0.09440559 | 43.30576253 | -84.57838553 |
| Isabella County, Rolland township | 0.05096826 | 0.09037117 | 43.47610705 | -84.98753723 |
| Gladwin County, Beaverton township | 0.06320602 | 0.08956428 | 43.88667001 | -84.51203444 |
| Ionia County, Keene township | 0.04061323 | 0.08472297 | 43.02185221 | -85.21558216 |
| Clinton County, Riley township | 0.05554061 | 0.08284024 | 42.90842159 | -84.67539957 |
| Montcalm County, Belvidere township | 0.05661646 | 0.08284024 | 43.42221455 | -85.15235556 |
| Isabella County, Coldwater township | 0.05486821 | 0.08149543 | 43.7979873 | -85.06097473 |
| Barry County, Assyria township | 0.05217859 | 0.08015062 | 42.50048157 | -85.17365442 |
| Calhoun County, Clarence township | 0.0450511 | 0.07611619 | 42.40284591 | -84.81967143 |
| Clinton County, Westphalia township | 0.05863367 | 0.07557827 | 42.86929025 | -84.7438229 |
| Eaton County, Kalamo township | 0.0535234 | 0.07315761 | 42.58168942 | -85.0683841 |
| Mecosta County, Millbrook township | 0.03079613 | 0.05675094 | 43.46671382 | -85.14528478 |
