## Supplementary material for "Mapping long-distance jump dispersal risk by leveraging transport infrastructure data and sampling bias correction: a case study with the invasive spotted lanternfly (*Lycorma delicatula*)": S9 New York risk

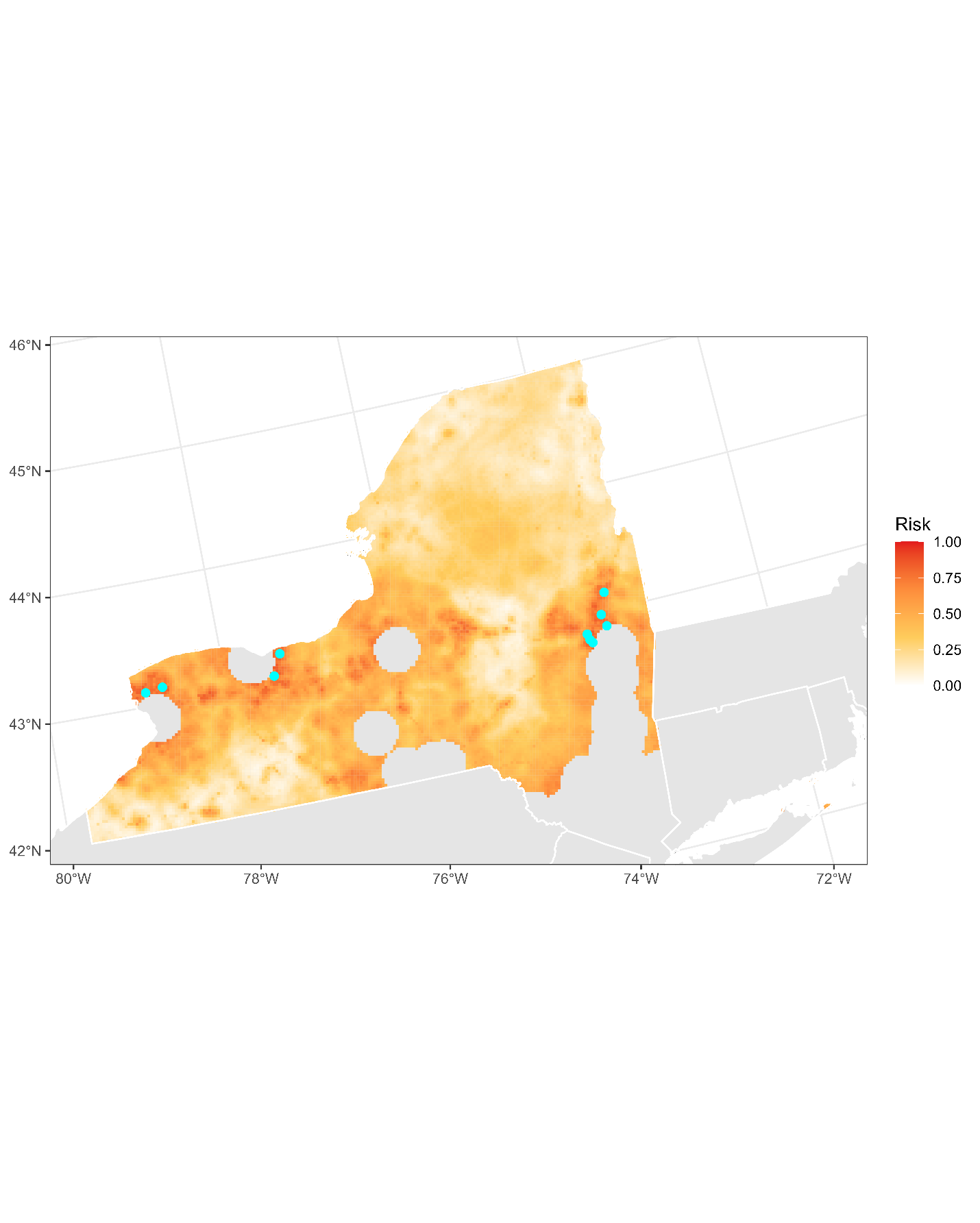

**Figure S9.1:** Projected risk of colonization by *L. delicatula* jump dispersal for New York. Risk projections are based on an ensemble random forest model including distances from existing *L. delicatula* populations. Note that areas falling less than 20 km from existing *L. delicatula* populations are not included in the map because any colonization there would be categorized as diffusive spread based on our methodology (regardless of the actual source of the resulting population). Darker, redder colors indicate increased risk of colonization by jump dispersal. Cyan points show the top ten highest risk locations in the state.

**Table S9.1:** Ranked list of New York county subdivisions sorted by the maximum projected risk of colonization by jump dispersal occurring there. For each subdivision, we identified the 2.5 x 2.5 km grid cell centroid included there with the highest projected risk of colonization by jump dispersal and report that location and risk value.

| **County, subdivision** | **Median risk value** | **Maximum risk value** | **Latitude of maximum risk** | **Longitude of maximum risk** |
| --- | --- | --- | --- | --- |
| Monroe County, Perinton town | 0.81387843 | 0.87896719 | 43.05365788 | -77.46467082 |
| Schenectady County, Glenville town | 0.69351802 | 0.87735342 | 42.85113504 | -73.99427389 |
| Schenectady County, Schenectady city | 0.78563744 | 0.86767079 | 42.80244705 | -73.97846267 |
| Niagara County, Lockport town | 0.74341044 | 0.86713287 | 43.13351492 | -78.70081524 |
| Niagara County, Wheatfield town | 0.79263045 | 0.85395374 | 43.11365728 | -78.8942198 |
| Schenectady County, Rotterdam town | 0.75833782 | 0.84884346 | 42.77553297 | -73.9556114 |
| Saratoga County, Malta town | 0.7767617 | 0.84749866 | 42.97776045 | -73.7947753 |
| Saratoga County, Halfmoon town | 0.81737493 | 0.84696073 | 42.88031958 | -73.76339848 |
| Wayne County, Ontario town | 0.70575578 | 0.84480904 | 43.22061393 | -77.35603221 |
| Saratoga County, Wilton town | 0.74744486 | 0.8394298 | 43.14673512 | -73.70749116 |
| Genesee County, Batavia town | 0.73507262 | 0.83808499 | 43.01963869 | -78.19638151 |
| Monroe County, Penfield town | 0.77138246 | 0.83808499 | 43.13714159 | -77.41041736 |
| Niagara County, Lewiston town | 0.71839699 | 0.83458849 | 43.16170812 | -78.91355818 |
| Niagara County, Niagara town | 0.73318989 | 0.83405056 | 43.12165768 | -78.95499074 |
| Saratoga County, Clifton Park town | 0.74394836 | 0.83055406 | 42.90727143 | -73.78620947 |
| Saratoga County, Saratoga Springs city | 0.68235611 | 0.82947821 | 43.05343318 | -73.83337474 |
| Wayne County, Walworth town | 0.66594944 | 0.82894029 | 43.17669966 | -77.3680823 |
| Saratoga County, Ballston town | 0.76223776 | 0.82786444 | 42.94456243 | -73.86897665 |
| Niagara County, Tuscarora Nation Reservation | 0.74610005 | 0.82490586 | 43.14368239 | -78.94947924 |
| Monroe County, Pittsford town | 0.68262507 | 0.8246369 | 43.05799199 | -77.49494169 |
| Niagara County, Niagara Falls city | 0.73762776 | 0.82329209 | 43.08159671 | -78.99637513 |
| Albany County, Guilderland town | 0.65021517 | 0.82248521 | 42.75376063 | -73.96267371 |
| Genesee County, Batavia city | 0.78778913 | 0.81091985 | 42.99348038 | -78.17183528 |
| Monroe County, Mendon town | 0.67845616 | 0.809844 | 42.97879782 | -77.57927209 |
| Monroe County, Webster town | 0.78456159 | 0.80957504 | 43.22497431 | -77.38636971 |
| Niagara County, Pendleton town | 0.77689618 | 0.80876815 | 43.11554829 | -78.73677719 |
| Monroe County, Rush town | 0.66379774 | 0.80688542 | 43.01366256 | -77.66410817 |
| Saratoga County, Milton town | 0.67307692 | 0.80527165 | 43.05860298 | -73.86339534 |
| Schenectady County, Princetown town | 0.57127488 | 0.8041958 | 42.76402952 | -74.02247863 |
| Ontario County, Farmington town | 0.64093599 | 0.80258203 | 42.97040204 | -77.3616584 |
| Livingston County, Avon town | 0.64537386 | 0.80204411 | 42.88188441 | -77.69951689 |
| Niagara County, Cambria town | 0.72565896 | 0.80150619 | 43.14160519 | -78.76158115 |
| Ontario County, Victor town | 0.7493276 | 0.79532006 | 42.99671037 | -77.38588836 |
| Oneida County, Whitestown town | 0.68020441 | 0.7950511 | 43.11414552 | -75.3026425 |
| Niagara County, Lockport city | 0.70629371 | 0.79424422 | 43.15552929 | -78.6952215 |
| Montgomery County, Amsterdam town | 0.70145239 | 0.79263045 | 42.920296 | -74.13000208 |
| Livingston County, Lima town | 0.66702528 | 0.7859064 | 42.93487992 | -77.59113802 |
| Niagara County, Wilson town | 0.55110274 | 0.78402367 | 43.2337178 | -78.76968578 |
| Erie County, Eden town | 0.58741259 | 0.77918236 | 42.67936156 | -78.87783729 |
| Oneida County, New Hartford town | 0.59117805 | 0.77595481 | 43.08745307 | -75.2791726 |
| Wayne County, Macedon town | 0.65572889 | 0.77407208 | 43.10647238 | -77.35584141 |
| Cortland County, Cortland city | 0.70346961 | 0.77299623 | 42.61071681 | -76.1765832 |
| Monroe County, Henrietta town | 0.76035503 | 0.77218935 | 43.01795032 | -77.69437395 |
| Saratoga County, Stillwater town | 0.62224314 | 0.7711135 | 42.92385108 | -73.74911787 |
| Schenectady County, Niskayuna town | 0.77084454 | 0.77084454 | 42.83570009 | -73.90446644 |
| Oneida County, Westmoreland town | 0.53765465 | 0.76976869 | 43.12862373 | -75.39311623 |
| Ontario County, Geneva city | 0.69795589 | 0.76707907 | 42.86926695 | -76.98128779 |
| Chautauqua County, Dunkirk city | 0.75026896 | 0.76681011 | 42.46662043 | -79.33490153 |
| Erie County, Hamburg town | 0.60758472 | 0.76385153 | 42.7013683 | -78.87233647 |
| Genesee County, Pembroke town | 0.55177515 | 0.76358257 | 43.01426163 | -78.32331193 |
| Chautauqua County, Dunkirk town | 0.7232383 | 0.75874126 | 42.4705183 | -79.36502737 |
| Onondaga County, Elbridge town | 0.63959118 | 0.7584723 | 43.02083589 | -76.46662204 |
| Oneida County, Marcy town | 0.5626681 | 0.75336202 | 43.13115865 | -75.26580926 |
| Onondaga County, Lysander town | 0.55728886 | 0.75309306 | 43.16044083 | -76.3317188 |
| Saratoga County, Corinth town | 0.39685315 | 0.75309306 | 43.20067786 | -73.75326389 |
| Cayuga County, Sennett town | 0.63757396 | 0.7471759 | 42.99892724 | -76.47291429 |
| Saratoga County, Northumberland town | 0.57772996 | 0.74690694 | 43.18506869 | -73.66305467 |
| Saratoga County, Greenfield town | 0.51358257 | 0.74636902 | 43.11357778 | -73.78191661 |
| Montgomery County, Amsterdam city | 0.74166218 | 0.74260355 | 42.93049068 | -74.18997604 |
| Oneida County, Rome city | 0.59601937 | 0.74260355 | 43.19902179 | -75.40335517 |
| Montgomery County, Florida town | 0.612695 | 0.7415277 | 42.93557777 | -74.21996965 |
| Ontario County, Phelps town | 0.58741259 | 0.7345347 | 42.94394501 | -77.02329653 |
| Fulton County, Gloversville city | 0.70158687 | 0.73426573 | 43.06991894 | -74.33518552 |
| Cayuga County, Throop town | 0.59817106 | 0.73157612 | 42.96876881 | -76.57599264 |
| Cayuga County, Aurelius town | 0.53657881 | 0.72942442 | 42.92948046 | -76.61865248 |
| Onondaga County, Van Buren town | 0.56401291 | 0.72834857 | 43.13853386 | -76.33806673 |
| Cayuga County, Owasco town | 0.54679935 | 0.72754169 | 42.91585323 | -76.5281897 |
| Chemung County, Horseheads town | 0.67805272 | 0.72700377 | 42.1627608 | -76.83457258 |
| Steuben County, Erwin town | 0.41312534 | 0.7267348 | 42.15453731 | -77.08548789 |
| Onondaga County, Skaneateles town | 0.56508876 | 0.72619688 | 42.99435767 | -76.44273533 |
| Cayuga County, Auburn city | 0.71799354 | 0.72538999 | 42.92494497 | -76.58849425 |
| Erie County, Aurora town | 0.61188811 | 0.72377622 | 42.73912249 | -78.64430528 |
| Erie County, Clarence town | 0.64739107 | 0.72377622 | 43.08137521 | -78.65127789 |
| Erie County, Orchard Park town | 0.68074233 | 0.72270038 | 42.70727499 | -78.74602369 |
| Wayne County, Arcadia town | 0.55204411 | 0.71812803 | 43.04498308 | -77.08952998 |
| Rensselaer County, Schaghticoke town | 0.48413125 | 0.71732114 | 42.88648547 | -73.66644031 |
| Rensselaer County, Grafton town | 0.57235073 | 0.71516945 | 42.81757484 | -73.40446883 |
| Erie County, Newstead town | 0.61323292 | 0.71328671 | 42.99905517 | -78.5467994 |
| Cayuga County, Mentz town | 0.63528779 | 0.71328671 | 43.04358785 | -76.61762258 |
| Genesee County, Le Roy town | 0.59628833 | 0.7119419 | 42.98606976 | -77.95410586 |
| Fulton County, Johnstown town | 0.48722431 | 0.71032813 | 43.03136582 | -74.37917306 |
| Oneida County, Kirkland town | 0.60476062 | 0.70952125 | 43.08008962 | -75.37625439 |
| Erie County, Boston town | 0.61377084 | 0.70925229 | 42.69333479 | -78.81194377 |
| Onondaga County, Clay town | 0.62775686 | 0.70817644 | 43.19503612 | -76.25849733 |
| Erie County, Grand Island town | 0.68719742 | 0.70710059 | 43.0595722 | -79.00186668 |
| Onondaga County, Camillus town | 0.6366326 | 0.70710059 | 43.02441228 | -76.33956064 |
| Chemung County, Elmira town | 0.68128026 | 0.70656267 | 42.0836119 | -76.76342916 |
| Cayuga County, Brutus town | 0.60287789 | 0.7049489 | 43.02540015 | -76.49681417 |
| Niagara County, Newfane town | 0.50403443 | 0.70387305 | 43.25169687 | -78.73367061 |
| Cortland County, Preble town | 0.50053792 | 0.70387305 | 42.72012931 | -76.14479006 |
| Erie County, Cattaraugus Reservation | 0.52649274 | 0.70333513 | 42.57120547 | -79.09164153 |
| Cortland County, Cortlandville town | 0.55379236 | 0.70306616 | 42.58420182 | -76.15294847 |
| Oswego County, Schroeppel town | 0.5894298 | 0.70252824 | 43.27458721 | -76.33020891 |
| Ontario County, Canandaigua town | 0.55190963 | 0.70225928 | 42.94409028 | -77.33744692 |
| Chemung County, Big Flats town | 0.6013986 | 0.70118343 | 42.14979898 | -76.90034719 |
| Oswego County, Hastings town | 0.53684777 | 0.70091447 | 43.29065968 | -76.13568632 |
| Ontario County, West Bloomfield town | 0.53039268 | 0.70037655 | 42.93056885 | -77.56091508 |
| Oneida County, Utica city | 0.58149543 | 0.70010758 | 43.09474469 | -75.18207016 |
| Oswego County, Volney town | 0.57719204 | 0.69849381 | 43.30569007 | -76.38445748 |
| Ontario County, Canandaigua city | 0.6958042 | 0.6958042 | 42.8914565 | -77.28907928 |
| Chemung County, Baldwin town | 0.49058634 | 0.69553523 | 42.12290203 | -76.72136766 |
| Oswego County, West Monroe town | 0.57208176 | 0.69526627 | 43.28601021 | -76.10540187 |
| Sullivan County, Thompson town | 0.64913932 | 0.69284562 | 41.73060451 | -74.69354645 |
| Rensselaer County, Petersburgh town | 0.53630984 | 0.68988704 | 42.79056022 | -73.38183633 |
| Sullivan County, Fallsburg town | 0.64201183 | 0.68934911 | 41.75238751 | -74.68681399 |
| Ontario County, Geneva town | 0.59655729 | 0.68908015 | 42.89119539 | -76.97516864 |
| Monroe County, Clarkson town | 0.62668101 | 0.68881119 | 43.23630847 | -77.95063441 |
| Montgomery County, Mohawk town | 0.56858526 | 0.6880043 | 42.9558577 | -74.33998823 |
| Sullivan County, Bethel town | 0.61377084 | 0.68773534 | 41.68011294 | -74.80220368 |
| Saratoga County, Charlton town | 0.55231307 | 0.68719742 | 42.91132404 | -73.94310662 |
| Fulton County, Mayfield town | 0.50887574 | 0.68692846 | 43.08158788 | -74.26807391 |
| Onondaga County, Marcellus town | 0.50968263 | 0.68639053 | 43.00250963 | -76.34589356 |
| Oswego County, Fulton city | 0.65922539 | 0.68531469 | 43.31027535 | -76.41476935 |
| Montgomery County, Glen town | 0.51519634 | 0.68477676 | 42.92900093 | -74.31694539 |
| Monroe County, Wheatland town | 0.59332975 | 0.68154922 | 43.02650494 | -77.75491679 |
| Steuben County, Corning city | 0.67939753 | 0.67939753 | 42.15011426 | -77.05563456 |
| Steuben County, Corning town | 0.54491662 | 0.67859064 | 42.14568436 | -77.02578502 |
| Chemung County, Chemung town | 0.53617536 | 0.67482517 | 42.01275746 | -76.59700548 |
| Otsego County, Oneonta city | 0.5979021 | 0.67482517 | 42.45157882 | -75.03312176 |
| Washington County, Kingsbury town | 0.34521248 | 0.67428725 | 43.31046936 | -73.58966306 |
| Cayuga County, Fleming town | 0.542227 | 0.67374933 | 42.89849288 | -76.56459401 |
| Monroe County, Sweden town | 0.62264658 | 0.67267348 | 43.21854516 | -77.986845 |
| Genesee County, Stafford town | 0.53335126 | 0.66998386 | 43.03327397 | -78.13002643 |
| Columbia County, Canaan town | 0.54599247 | 0.66998386 | 42.36176868 | -73.42961367 |
| Chemung County, Erin town | 0.53604088 | 0.66998386 | 42.14479598 | -76.71523234 |
| Erie County, Wales town | 0.59279182 | 0.66675632 | 42.74480856 | -78.5179047 |
| Rensselaer County, Pittstown town | 0.542227 | 0.6662184 | 42.82284099 | -73.4343547 |
| Cattaraugus County, Olean city | 0.57719204 | 0.66218397 | 42.09652169 | -78.43442582 |
| Chemung County, Elmira city | 0.64739107 | 0.66218397 | 42.11448309 | -76.81694475 |
| Chemung County, Southport town | 0.52339968 | 0.66030124 | 42.07516231 | -76.85894737 |
| Yates County, Milo town | 0.49179666 | 0.65303927 | 42.63250937 | -77.07840186 |
| Monroe County, Hamlin town | 0.62049489 | 0.6514255 | 43.32425309 | -77.92724328 |
| Orleans County, Murray town | 0.60435718 | 0.6492738 | 43.25735347 | -78.10257915 |
| Warren County, Queensbury town | 0.39483593 | 0.64766003 | 43.28258043 | -73.69442831 |
| Sullivan County, Liberty town | 0.55164067 | 0.64766003 | 41.76726388 | -74.77540321 |
| Sullivan County, Neversink town | 0.5232652 | 0.6457773 | 41.83454226 | -74.6302861 |
| Oswego County, Granby town | 0.56589564 | 0.6457773 | 43.25727217 | -76.36685316 |
| Ontario County, Hopewell town | 0.56750941 | 0.64281872 | 42.89583 | -77.16230242 |
| Genesee County, Elba town | 0.48144164 | 0.64093599 | 43.06361637 | -78.18488692 |
| Oneida County, Floyd town | 0.43934911 | 0.63878429 | 43.18938015 | -75.34297198 |
| Livingston County, Geneseo town | 0.5478752 | 0.63824637 | 42.8422378 | -77.74148158 |
| Fulton County, Perth town | 0.61861216 | 0.63797741 | 43.01619129 | -74.28904531 |
| Saratoga County, Moreau town | 0.55648198 | 0.63717052 | 43.27737004 | -73.66432264 |
| Washington County, Fort Edward town | 0.45481442 | 0.63690156 | 43.28346604 | -73.56679539 |
| Saratoga County, Saratoga town | 0.46799354 | 0.63636364 | 43.09799052 | -73.69181492 |
| Niagara County, Royalton town | 0.48332437 | 0.63636364 | 43.09931741 | -78.6153115 |
| Erie County, Evans town | 0.54491662 | 0.63232921 | 42.64733488 | -78.97936025 |
| Oneida County, Vernon town | 0.4527972 | 0.63206025 | 43.06991831 | -75.60050612 |
| Rensselaer County, Hoosick town | 0.48440022 | 0.63179129 | 42.83931864 | -73.39722431 |
| Orleans County, Kendall town | 0.59064013 | 0.63125336 | 43.28872472 | -77.99975051 |
| Erie County, Brant town | 0.54330285 | 0.6309844 | 42.5892547 | -79.05605382 |
| Orleans County, Ridgeway town | 0.54774072 | 0.62990855 | 43.23289818 | -78.42382837 |
| Chautauqua County, Sheridan town | 0.51291017 | 0.62937063 | 42.48082037 | -79.26930241 |
| Orleans County, Gaines town | 0.51667563 | 0.62910167 | 43.26154145 | -78.13297924 |
| Erie County, Holland town | 0.56105433 | 0.62883271 | 42.65683923 | -78.54035093 |
| Livingston County, Livonia town | 0.49878967 | 0.62829478 | 42.83796207 | -77.7112898 |
| Livingston County, Caledonia town | 0.54948897 | 0.62802582 | 42.97336633 | -77.8633135 |
| Seneca County, Fayette town | 0.45373857 | 0.62775686 | 42.86481551 | -76.95113225 |
| Orleans County, Clarendon town | 0.56455083 | 0.6274879 | 43.20497838 | -78.05340713 |
| Genesee County, Alexander town | 0.43491124 | 0.6274879 | 42.93999998 | -78.27985796 |
| Oneida County, Verona town | 0.49408284 | 0.62721893 | 43.12318321 | -75.64770092 |
| Erie County, North Collins town | 0.48332437 | 0.62587413 | 42.60532174 | -78.83400023 |
| Tioga County, Spencer town | 0.57140936 | 0.6253362 | 42.23162158 | -76.53518509 |
| Monroe County, Riga town | 0.53039268 | 0.6253362 | 43.09172344 | -77.89465503 |
| Orleans County, Shelby town | 0.53254438 | 0.62479828 | 43.20677372 | -78.3991196 |
| Cortland County, Homer town | 0.42482517 | 0.6218397 | 42.63259858 | -76.17023261 |
| Wayne County, Lyons town | 0.45992469 | 0.61780527 | 43.0580571 | -77.02293632 |
| Chautauqua County, Pomfret town | 0.3041958 | 0.61780527 | 42.44069811 | -79.31012405 |
| Seneca County, Waterloo town | 0.50268962 | 0.61403981 | 42.88674239 | -76.94500391 |
| Genesee County, Darien town | 0.48547606 | 0.612695 | 42.93865826 | -78.43691161 |
| Suffolk County, Southold town | 0.48708983 | 0.61242604 | 41.11150248 | -72.38358943 |
| Chemung County, Veteran town | 0.49488973 | 0.61135019 | 42.20207786 | -76.79250342 |
| Erie County, Concord town | 0.54679935 | 0.61081226 | 42.57123814 | -78.74908168 |
| Erie County, Colden town | 0.58647122 | 0.60973642 | 42.66906706 | -78.63085006 |
| Erie County, Alden town | 0.56253362 | 0.60919849 | 42.95096289 | -78.52776196 |
| Columbia County, New Lebanon town | 0.56979559 | 0.60866057 | 42.44273058 | -73.49720069 |
| Ontario County, East Bloomfield town | 0.51237224 | 0.6070468 | 42.93954658 | -77.46430093 |
| Fulton County, Broadalbin town | 0.50645508 | 0.60650888 | 43.0445418 | -74.1849269 |
| Cattaraugus County, Allegany town | 0.19257665 | 0.60623991 | 42.07865416 | -78.46992234 |
| Wyoming County, Bennington town | 0.45992469 | 0.60597095 | 42.78469753 | -78.47644537 |
| Madison County, Oneida city | 0.58714363 | 0.60570199 | 43.07468242 | -75.63066118 |
| Columbia County, Chatham town | 0.53496503 | 0.60435718 | 42.44797276 | -73.52693237 |
| Fulton County, Johnstown city | 0.60408822 | 0.60408822 | 42.98775931 | -74.39307803 |
| Rensselaer County, Berlin town | 0.49623453 | 0.6013986 | 42.61663699 | -73.43966321 |
| Ontario County, Manchester town | 0.53227542 | 0.60059172 | 42.97484638 | -77.23474108 |
| Wyoming County, Sheldon town | 0.51923077 | 0.60032275 | 42.75860106 | -78.45187673 |
| Yates County, Jerusalem town | 0.37009145 | 0.59924691 | 42.63692996 | -77.10846449 |
| Rensselaer County, Nassau town | 0.55110274 | 0.59817106 | 42.60015258 | -73.47666431 |
| Oneida County, Sangerfield town | 0.40667025 | 0.5979021 | 42.92228938 | -75.39259662 |
| Greene County, Windham town | 0.5626681 | 0.59736417 | 42.32857194 | -74.16244378 |
| Chenango County, Norwich town | 0.51438946 | 0.59655729 | 42.50715689 | -75.51772997 |
| Schenectady County, Duanesburg town | 0.45239376 | 0.59655729 | 42.74737637 | -74.05941895 |
| Oneida County, Marshall town | 0.51613771 | 0.5957504 | 42.99747372 | -75.43291199 |
| Oswego County, Palermo town | 0.47902098 | 0.59467456 | 43.36223592 | -76.30473251 |
| Rensselaer County, Stephentown town | 0.53442711 | 0.59306079 | 42.5948978 | -73.44687118 |
| Onondaga County, Tully town | 0.50699301 | 0.5922539 | 42.8076661 | -76.1192831 |
| Chautauqua County, Jamestown city | 0.43948359 | 0.59090909 | 42.09096437 | -79.24006831 |
| Warren County, Glens Falls city | 0.57557827 | 0.58902636 | 43.32091807 | -73.64989665 |
| Wyoming County, Arcade town | 0.4458042 | 0.58848843 | 42.53051484 | -78.44789746 |
| Schoharie County, Carlisle town | 0.5112964 | 0.58768155 | 42.75463679 | -74.37253471 |
| Chenango County, Bainbridge town | 0.51788596 | 0.58687466 | 42.29143072 | -75.45722548 |
| Oswego County, Constantia town | 0.48830016 | 0.58687466 | 43.27202016 | -76.01457314 |
| Madison County, Eaton town | 0.44553523 | 0.5866057 | 42.85883421 | -75.56930707 |
| Orleans County, Carlton town | 0.51291017 | 0.58418505 | 43.30971462 | -78.15182614 |
| Chemung County, Van Etten town | 0.52864443 | 0.58364712 | 42.16595929 | -76.55377195 |
| Oswego County, Oswego city | 0.56347499 | 0.58203335 | 43.46009454 | -76.49818289 |
| Oswego County, Scriba town | 0.53119957 | 0.58176439 | 43.47745212 | -76.46146342 |
| Herkimer County, Little Falls city | 0.5809575 | 0.5809575 | 43.04082426 | -74.85084822 |
| Tioga County, Barton town | 0.48372781 | 0.58041958 | 42.00368097 | -76.53748647 |
| Herkimer County, Frankfort town | 0.38474987 | 0.57988166 | 43.0315528 | -75.07504779 |
| Yates County, Starkey town | 0.51882733 | 0.57961269 | 42.54455202 | -76.94646252 |
| Orleans County, Albion town | 0.4971759 | 0.57934373 | 43.24372716 | -78.16915594 |
| Chemung County, Ashland town | 0.52124798 | 0.57934373 | 42.03982297 | -76.77565499 |
| Wayne County, Palmyra town | 0.49408284 | 0.57880581 | 43.06701706 | -77.24078502 |
| Greene County, Durham town | 0.55513717 | 0.57853685 | 42.3503423 | -74.15548713 |
| Rensselaer County, Sand Lake town | 0.57450242 | 0.57853685 | 42.62189357 | -73.46946514 |
| Columbia County, Hillsdale town | 0.5 | 0.57719204 | 42.19840417 | -73.54621492 |
| Erie County, Marilla town | 0.50712749 | 0.57692308 | 42.78879597 | -78.50666017 |
| Cayuga County, Genoa town | 0.5112964 | 0.57665412 | 42.67488689 | -76.5968667 |
| Oswego County, Oswego town | 0.54774072 | 0.57638515 | 43.39431841 | -76.51713818 |
| Genesee County, Oakfield town | 0.47471759 | 0.57611619 | 43.07609834 | -78.27585681 |
| Dutchess County, Amenia town | 0.50268962 | 0.57557827 | 41.85677755 | -73.56424618 |
| Genesee County, Alabama town | 0.47686928 | 0.5718128 | 43.03625559 | -78.3176074 |
| Chemung County, Catlin town | 0.47807961 | 0.57154384 | 42.21103899 | -76.85222483 |
| Onondaga County, LaFayette town | 0.48224852 | 0.57019903 | 42.8560829 | -76.13659955 |
| Cayuga County, Ledyard town | 0.4908553 | 0.56966111 | 42.67941879 | -76.62691663 |
| Seneca County, Junius town | 0.34911243 | 0.56858526 | 42.95698334 | -76.95678784 |
| Columbia County, Ghent town | 0.54088219 | 0.5683163 | 42.34450656 | -73.59239594 |
| Herkimer County, Herkimer town | 0.31522324 | 0.56804734 | 43.02666125 | -75.04495052 |
| Ulster County, Olive town | 0.5387305 | 0.56697149 | 41.98027651 | -74.27316116 |
| Dutchess County, North East town | 0.50860678 | 0.56697149 | 41.89499527 | -73.52054983 |
| Washington County, Greenwich town | 0.43262507 | 0.56670253 | 43.0883895 | -73.50449733 |
| Madison County, Madison town | 0.46839699 | 0.5661646 | 42.85405758 | -75.53925041 |
| Cattaraugus County, Ashford town | 0.46355568 | 0.56562668 | 42.48905781 | -78.64525448 |
| Erie County, Sardinia town | 0.51102743 | 0.56428187 | 42.53871506 | -78.50810598 |
| Schoharie County, Cobleskill town | 0.53321678 | 0.56428187 | 42.6775296 | -74.46001902 |
| Oneida County, Lee town | 0.39402905 | 0.56401291 | 43.27903746 | -75.47407849 |
| Livingston County, West Sparta town | 0.30096826 | 0.56374395 | 42.56592832 | -77.72146251 |
| Chenango County, Afton town | 0.48951049 | 0.56374395 | 42.25254668 | -75.50009829 |
| Cayuga County, Venice town | 0.50833782 | 0.56293706 | 42.69679092 | -76.59064309 |
| Oneida County, Camden town | 0.48776224 | 0.5626681 | 43.29531971 | -75.72265911 |
| Oswego County, Sandy Creek town | 0.46180742 | 0.56213018 | 43.60666559 | -76.10618106 |
| Washington County, Easton town | 0.41487359 | 0.56159225 | 43.0988735 | -73.56452388 |
| Genesee County, Bethany town | 0.43625605 | 0.56132329 | 42.94950862 | -78.1833265 |
| Sullivan County, Delaware town | 0.51062399 | 0.56078537 | 41.79460869 | -75.07759006 |
| Onondaga County, Otisco town | 0.47095212 | 0.56024744 | 42.83883324 | -76.17307043 |
| Cayuga County, Moravia town | 0.51909629 | 0.55648198 | 42.71781348 | -76.42785713 |
| Albany County, Westerlo town | 0.54061323 | 0.55648198 | 42.52451428 | -74.09967678 |
| Oswego County, Richland town | 0.50645508 | 0.55648198 | 43.58475309 | -76.11264258 |
| Genesee County, Pavilion town | 0.43544917 | 0.55540613 | 42.94211773 | -77.9657345 |
| Madison County, Hamilton town | 0.39497041 | 0.55513717 | 42.83220006 | -75.54582099 |
| Oneida County, Vienna town | 0.48843464 | 0.55459925 | 43.25155646 | -75.73576373 |
| Otsego County, Milford town | 0.15505648 | 0.55459925 | 42.46849976 | -74.99657392 |
| Albany County, Rensselaerville town | 0.51237224 | 0.5535234 | 42.45269932 | -74.21695889 |
| Seneca County, Ovid town | 0.5091447 | 0.55271651 | 42.66655354 | -76.69322066 |
| Seneca County, Romulus town | 0.46584185 | 0.55190963 | 42.68455068 | -76.81345448 |
| Greene County, Cairo town | 0.5478752 | 0.55029586 | 42.31837338 | -74.10299232 |
| Genesee County, Tonawanda Reservation | 0.54814416 | 0.54922001 | 43.0788712 | -78.46356016 |
| Broome County, Barker town | 0.43060785 | 0.54868209 | 42.29700939 | -75.92399311 |
| Chenango County, Oxford town | 0.42805272 | 0.5478752 | 42.46346587 | -75.53081492 |
| Tompkins County, Lansing town | 0.53321678 | 0.5478752 | 42.6091771 | -76.61551397 |
| Ontario County, Bristol town | 0.49838623 | 0.5478752 | 42.82111537 | -77.43376512 |
| Columbia County, Copake town | 0.5 | 0.54733728 | 42.17667109 | -73.55334647 |
| Cattaraugus County, Portville town | 0.25147929 | 0.54706832 | 42.06632442 | -78.38021646 |
| Columbia County, Taghkanic town | 0.54464766 | 0.54464766 | 42.16538305 | -73.61970552 |
| Wayne County, Williamson town | 0.45831092 | 0.54464766 | 43.2733522 | -77.24685507 |
| Greene County, Jewett town | 0.47579344 | 0.54410974 | 42.26326228 | -74.18328765 |
| Otsego County, Oneonta town | 0.41043572 | 0.54330285 | 42.43952972 | -75.09949449 |
| Oswego County, Hannibal town | 0.48036579 | 0.54330285 | 43.27558636 | -76.48803476 |
| Chenango County, Preston town | 0.43329747 | 0.54303389 | 42.46824182 | -75.56070204 |
| Schoharie County, Richmondville town | 0.48413125 | 0.54276493 | 42.66075489 | -74.49680766 |
| Ontario County, Gorham town | 0.44271114 | 0.54276493 | 42.84754907 | -77.14425853 |
| Schuyler County, Hector town | 0.40344271 | 0.542227 | 42.51371726 | -76.89255773 |
| Greene County, Greenville town | 0.53765465 | 0.54195804 | 42.37856008 | -74.05231097 |
| Albany County, Knox town | 0.47982786 | 0.54115116 | 42.66026995 | -74.08749957 |
| Madison County, Lenox town | 0.48076923 | 0.54034427 | 43.13268359 | -75.70806836 |
| Madison County, Lebanon town | 0.42227004 | 0.54034427 | 42.81511622 | -75.58242598 |
| Cattaraugus County, Yorkshire town | 0.42791824 | 0.53953739 | 42.51672884 | -78.51370932 |
| Columbia County, Austerlitz town | 0.46328671 | 0.53899946 | 42.33928007 | -73.56270354 |
| Chautauqua County, Hanover town | 0.3725121 | 0.53819258 | 42.53113412 | -79.13261899 |
| Delaware County, Deposit town | 0.46853147 | 0.53792361 | 42.00544785 | -75.38702655 |
| Schoharie County, Conesville town | 0.45145239 | 0.53711673 | 42.37066503 | -74.27447743 |
| Delaware County, Masonville town | 0.38219473 | 0.53550296 | 42.28182667 | -75.39762137 |
| Delaware County, Sidney town | 0.36229155 | 0.53496503 | 42.30366009 | -75.39105623 |
| Oswego County, Minetto town | 0.52904788 | 0.53469607 | 43.38975609 | -76.48678237 |
| Chautauqua County, Ellicott town | 0.33647122 | 0.53442711 | 42.10904291 | -79.20476381 |
| Ontario County, Seneca town | 0.46342119 | 0.53442711 | 42.90008058 | -77.03550974 |
| Wyoming County, Covington town | 0.43786982 | 0.53388919 | 42.81868339 | -78.06093025 |
| Genesee County, Bergen town | 0.49502421 | 0.5330823 | 43.09595841 | -77.92496935 |
| Madison County, Lincoln town | 0.47982786 | 0.5330823 | 43.05464417 | -75.79452871 |
| Oneida County, Western town | 0.32759548 | 0.53227542 | 43.29129293 | -75.40696062 |
| Broome County, Colesville town | 0.43948359 | 0.53200646 | 42.23269602 | -75.66207098 |
| Oneida County, Annsville town | 0.40263583 | 0.53200646 | 43.29057507 | -75.69240085 |
| Schoharie County, Schoharie town | 0.48184508 | 0.53146853 | 42.68421016 | -74.3634089 |
| Madison County, Sullivan town | 0.44862829 | 0.53119957 | 43.08124256 | -75.8181867 |
| Ulster County, Woodstock town | 0.51882733 | 0.53093061 | 42.08911307 | -74.23868065 |
| Oswego County, Mexico town | 0.49677246 | 0.53039268 | 43.46718881 | -76.24242165 |
| Chenango County, Greene town | 0.42388381 | 0.53012372 | 42.26402651 | -75.71520945 |
| Steuben County, Bath town | 0.32571275 | 0.52985476 | 42.3781743 | -77.36662242 |
| Tompkins County, Groton town | 0.45696611 | 0.52931684 | 42.58541358 | -76.30920151 |
| Yates County, Benton town | 0.40559441 | 0.52904788 | 42.67636061 | -77.06626297 |
| Cayuga County, Ira town | 0.44055944 | 0.52877891 | 43.25366776 | -76.4943435 |
| Wyoming County, Gainesville town | 0.44432491 | 0.52850995 | 42.66904913 | -78.13137068 |
| Seneca County, Covert town | 0.45360409 | 0.52850995 | 42.61822458 | -76.67556322 |
| Schoharie County, Seward town | 0.42092523 | 0.52824099 | 42.70936938 | -74.51295068 |
| Cayuga County, Niles town | 0.47821409 | 0.52797203 | 42.79265139 | -76.46917583 |
| Chenango County, Coventry town | 0.4168908 | 0.52770307 | 42.25928736 | -75.68539939 |
| Onondaga County, Spafford town | 0.46328671 | 0.52716514 | 42.89300284 | -76.37749797 |
| Niagara County, Hartland town | 0.44271114 | 0.52662722 | 43.23545943 | -78.61196518 |
| Cayuga County, Conquest town | 0.36861216 | 0.52635826 | 43.08742252 | -76.60511796 |
| Rensselaer County, Poestenkill town | 0.51048951 | 0.52555137 | 42.70886022 | -73.44062324 |
| Suffolk County, East Hampton town | 0.50295858 | 0.52447552 | 41.04999832 | -71.94030785 |
| Ulster County, Shandaken town | 0.45225928 | 0.5239376 | 42.07241379 | -74.27521244 |
| Herkimer County, Schuyler town | 0.2760893 | 0.52313072 | 43.05339301 | -75.06830897 |
| Wyoming County, Warsaw town | 0.49246907 | 0.52232383 | 42.73915514 | -78.14432017 |
| Ontario County, Richmond town | 0.46261431 | 0.52178591 | 42.84274463 | -77.58463864 |
| Schoharie County, Esperance town | 0.49731038 | 0.52097902 | 42.72273798 | -74.31963311 |
| Cayuga County, Montezuma town | 0.4253631 | 0.52071006 | 42.98236404 | -76.6665342 |
| Cayuga County, Scipio town | 0.46100054 | 0.52071006 | 42.77075015 | -76.47544653 |
| Albany County, Berne town | 0.47686928 | 0.52017214 | 42.61671853 | -74.10151343 |
| Broome County, Sanford town | 0.46167294 | 0.51990317 | 42.04646405 | -75.49926957 |
| Oswego County, Parish town | 0.42065627 | 0.51936525 | 43.44864201 | -76.12096768 |
| Cattaraugus County, Olean town | 0.27097902 | 0.51909629 | 42.09241037 | -78.40451477 |
| Broome County, Lisle town | 0.41917698 | 0.51882733 | 42.3766382 | -75.99439668 |
| Livingston County, Conesus town | 0.44620764 | 0.51882733 | 42.76351863 | -77.66860409 |
| Chautauqua County, Arkwright town | 0.3154922 | 0.51855836 | 42.41477198 | -79.28536505 |
| Sullivan County, Fremont town | 0.43181818 | 0.51855836 | 41.81641312 | -75.07097092 |
| Onondaga County, Fabius town | 0.43275955 | 0.51855836 | 42.84678389 | -76.07641527 |
| Chenango County, Norwich city | 0.5182894 | 0.5182894 | 42.52900288 | -75.51118131 |
| Seneca County, Seneca Falls town | 0.47095212 | 0.51775148 | 42.91718751 | -76.8422097 |
| Chenango County, Sherburne town | 0.44432491 | 0.51748252 | 42.6867265 | -75.49520191 |
| Livingston County, York town | 0.4281872 | 0.51586875 | 42.90320847 | -77.85060546 |
| Madison County, Georgetown town | 0.40882195 | 0.51506186 | 42.81701289 | -75.73918886 |
| Yates County, Torrey town | 0.44620764 | 0.5147929 | 42.63222056 | -76.92203328 |
| Madison County, Cazenovia town | 0.45548682 | 0.51237224 | 42.92338865 | -75.83351923 |
| Oswego County, Albion town | 0.41016676 | 0.51183432 | 43.55352321 | -76.05829669 |
| Delaware County, Hancock town | 0.41554599 | 0.5112964 | 41.973993 | -75.33423197 |
| Sullivan County, Rockland town | 0.45373857 | 0.5112964 | 41.8861068 | -74.80097634 |
| Oswego County, Orwell town | 0.43033889 | 0.50995159 | 43.59266824 | -76.01492663 |
| Schuyler County, Cayuta town | 0.47686928 | 0.50995159 | 42.30257439 | -76.70206925 |
| Otsego County, Otego town | 0.31118881 | 0.50968263 | 42.42258051 | -75.13600202 |
| Cayuga County, Locke town | 0.43302851 | 0.50941366 | 42.69134016 | -76.40408972 |
| Cayuga County, Sterling town | 0.38273265 | 0.50941366 | 43.38603434 | -76.61451231 |
| Chenango County, Smithville town | 0.44163529 | 0.5091447 | 42.41405134 | -75.79586066 |
| Greene County, Ashland town | 0.43141474 | 0.5091447 | 42.32711068 | -74.28831599 |
| Orleans County, Barre town | 0.44862829 | 0.50887574 | 43.20390773 | -78.21107004 |
| Schoharie County, Sharon town | 0.38569123 | 0.50511027 | 42.75798527 | -74.52911638 |
| Chenango County, Guilford town | 0.37627757 | 0.50484131 | 42.45868308 | -75.50093192 |
| Montgomery County, Root town | 0.34454008 | 0.50484131 | 42.90558454 | -74.45083282 |
| Schoharie County, Middleburgh town | 0.40223238 | 0.50457235 | 42.66746257 | -74.40022446 |
| Sullivan County, Callicoon town | 0.47068316 | 0.50430339 | 41.79683446 | -74.95269411 |
| Madison County, Nelson town | 0.45077999 | 0.50430339 | 42.86548037 | -75.7562412 |
| Niagara County, Porter town | 0.43275955 | 0.50403443 | 43.22172479 | -79.02421937 |
| Oswego County, New Haven town | 0.48251748 | 0.50403443 | 43.43258672 | -76.31594176 |
| Greene County, Hunter town | 0.44849381 | 0.50322754 | 42.11595283 | -74.26141621 |
| Onondaga County, Onondaga town | 0.47310382 | 0.50295858 | 42.91838666 | -76.24431381 |
| Broome County, Triangle town | 0.4190425 | 0.50215169 | 42.32355566 | -75.94744306 |
| Schoharie County, Fulton town | 0.42146315 | 0.50107585 | 42.62891076 | -74.44392054 |
| Steuben County, Campbell town | 0.37009145 | 0.50080689 | 42.18527775 | -77.13920018 |
| Chautauqua County, Villenova town | 0.36081226 | 0.50053792 | 42.36520805 | -79.08016811 |
| Warren County, Lake Luzerne town | 0.33700914 | 0.50053792 | 43.27120171 | -73.76183249 |
| Steuben County, Caton town | 0.3655191 | 0.50026896 | 42.07552731 | -77.01406861 |
| Wyoming County, Perry town | 0.39470145 | 0.5 | 42.79670793 | -78.06669841 |
| Onondaga County, Pompey town | 0.47149005 | 0.49946208 | 42.89373777 | -75.93676202 |
| Dutchess County, Washington town | 0.47807961 | 0.49892415 | 41.80723553 | -73.673906 |
| Tompkins County, Caroline town | 0.44486283 | 0.4965035 | 42.31361355 | -76.32481302 |
| Orleans County, Yates town | 0.43988704 | 0.49623453 | 43.31679916 | -78.37060187 |
| Schuyler County, Orange town | 0.36955352 | 0.49596557 | 42.32096429 | -76.97734479 |
| Dutchess County, Stanford town | 0.47417967 | 0.49542765 | 41.91588826 | -73.63857947 |
| Seneca County, Tyre town | 0.35476062 | 0.49542765 | 42.97847073 | -76.79350019 |
| Ulster County, Denning town | 0.46987628 | 0.49515869 | 41.92343822 | -74.47807639 |
| Cortland County, Scott town | 0.37547068 | 0.49515869 | 42.72476566 | -76.17482971 |
| Oswego County, Redfield town | 0.39994621 | 0.4943518 | 43.49407829 | -75.82179563 |
| Cattaraugus County, Persia town | 0.44822485 | 0.49300699 | 42.45931868 | -78.93265392 |
| Broome County, Nanticoke town | 0.46207639 | 0.49300699 | 42.28450852 | -75.99006127 |
| Schoharie County, Summit town | 0.40572889 | 0.49220011 | 42.55677559 | -74.56102173 |
| Oneida County, Florence town | 0.41258741 | 0.49220011 | 43.40178799 | -75.81762166 |
| Cattaraugus County, Otto town | 0.4316837 | 0.49112426 | 42.38531963 | -78.88889719 |
| Schoharie County, Broome town | 0.43249059 | 0.49058634 | 42.47955537 | -74.2398051 |
| Wyoming County, Java town | 0.38676708 | 0.48977945 | 42.66653616 | -78.44424989 |
| Jefferson County, Watertown city | 0.43571813 | 0.48897257 | 43.96988159 | -75.93454072 |
| Erie County, Collins town | 0.40290479 | 0.48897257 | 42.52532696 | -78.91624542 |
| Greene County, Lexington town | 0.41420118 | 0.48789672 | 42.14278948 | -74.28416926 |
| Clinton County, Plattsburgh city | 0.35180204 | 0.48789672 | 44.6924807 | -73.44654091 |
| Schuyler County, Dix town | 0.44109736 | 0.48762776 | 42.37331855 | -76.86929344 |
| Niagara County, Somerset town | 0.45266272 | 0.4873588 | 43.36346742 | -78.5476542 |
| Wayne County, Marion town | 0.42926304 | 0.4873588 | 43.19870743 | -77.20440254 |
| Otsego County, Unadilla town | 0.33122647 | 0.48682087 | 42.32549379 | -75.38448698 |
| Seneca County, Lodi town | 0.44970414 | 0.48628295 | 42.61432492 | -76.80186202 |
| Dutchess County, Pine Plains town | 0.4612695 | 0.48628295 | 41.97063811 | -73.55825534 |
| Tioga County, Candor town | 0.47552448 | 0.48601399 | 42.29173366 | -76.33108596 |
| Chautauqua County, Cherry Creek town | 0.31818182 | 0.48574502 | 42.3432027 | -79.08557664 |
| Fulton County, Bleecker town | 0.1697149 | 0.48547606 | 43.11352489 | -74.32122731 |
| Cortland County, Marathon town | 0.42993545 | 0.48493814 | 42.45157716 | -76.03504541 |
| Cortland County, Virgil town | 0.39819796 | 0.48359333 | 42.54044328 | -76.16564771 |
| Delaware County, Walton town | 0.36040882 | 0.4817106 | 42.09040359 | -75.2057069 |
| Cayuga County, Cato town | 0.42334588 | 0.48117267 | 43.09569574 | -76.50816464 |
| Wyoming County, Middlebury town | 0.43087682 | 0.48117267 | 42.84485555 | -78.08537243 |
| Washington County, Salem town | 0.40102205 | 0.48063475 | 43.19256009 | -73.31072593 |
| Cattaraugus County, Dayton town | 0.3929532 | 0.47982786 | 42.36124475 | -79.05010399 |
| Wayne County, Sodus town | 0.3894567 | 0.47982786 | 43.16774559 | -76.99228397 |
| Schuyler County, Catharine town | 0.4133943 | 0.4795589 | 42.30707979 | -76.73196531 |
| Saratoga County, Galway town | 0.44782141 | 0.47928994 | 43.05228927 | -73.96057768 |
| Chautauqua County, Cattaraugus Reservation | 0.45898332 | 0.47902098 | 42.54919356 | -79.09706248 |
| Cattaraugus County, Perrysburg town | 0.38891877 | 0.47794513 | 42.51528423 | -79.01208366 |
| Delaware County, Andes town | 0.38017751 | 0.47794513 | 42.07529123 | -74.83614025 |
| Schuyler County, Reading town | 0.44109736 | 0.47794513 | 42.39969524 | -76.89312892 |
| Chenango County, Smyrna town | 0.36632598 | 0.47767617 | 42.69629086 | -75.55516808 |
| Madison County, Fenner town | 0.44499731 | 0.47579344 | 43.00616774 | -75.7774027 |
| Delaware County, Middletown town | 0.42173211 | 0.47444863 | 42.16129516 | -74.5284105 |
| Cattaraugus County, Freedom town | 0.40182894 | 0.47202797 | 42.50442236 | -78.42342948 |
| Yates County, Barrington town | 0.36740183 | 0.47149005 | 42.57536574 | -77.00041523 |
| Ontario County, South Bristol town | 0.39416353 | 0.47149005 | 42.75527459 | -77.45165627 |
| Montgomery County, Charleston town | 0.33768155 | 0.47122109 | 42.7814768 | -74.39554105 |
| Oneida County, Ava town | 0.3929532 | 0.4704142 | 43.41986307 | -75.4947885 |
| Cayuga County, Sempronius town | 0.40290479 | 0.46960732 | 42.77433396 | -76.34884478 |
| Livingston County, Groveland town | 0.40061861 | 0.46933835 | 42.75012277 | -77.73479079 |
| Schoharie County, Wright town | 0.43047337 | 0.46906939 | 42.70752964 | -74.22992003 |
| Oneida County, Augusta town | 0.37963959 | 0.46826251 | 42.9367013 | -75.48283774 |
| Cattaraugus County, Leon town | 0.28362023 | 0.46745562 | 42.33924077 | -79.05552172 |
| Madison County, Stockbridge town | 0.35583647 | 0.46745562 | 43.02618312 | -75.6136454 |
| Yates County, Potter town | 0.32423346 | 0.4669177 | 42.68520101 | -77.12642858 |
| Oneida County, Paris town | 0.309844 | 0.46557289 | 43.00004686 | -75.30584847 |
| Cayuga County, Springport town | 0.37143626 | 0.464766 | 42.86374315 | -76.63734773 |
| Steuben County, Hornby town | 0.35099516 | 0.46422808 | 42.18950242 | -77.0136978 |
| Delaware County, Colchester town | 0.38004303 | 0.46342119 | 41.96904955 | -75.02452137 |
| Cattaraugus County, Salamanca city | 0.3000269 | 0.46288327 | 42.15519654 | -78.69827038 |
| Cattaraugus County, Machias town | 0.36767079 | 0.46180742 | 42.42060206 | -78.47597061 |
| Oneida County, Deerfield town | 0.2197418 | 0.46153846 | 43.14330481 | -75.19879868 |
| Ontario County, Canadice town | 0.41043572 | 0.46019365 | 42.75492756 | -77.60830199 |
| Oswego County, Amboy town | 0.42361485 | 0.45992469 | 43.38932056 | -75.88477352 |
| Wayne County, Butler town | 0.38421194 | 0.45615923 | 43.20222831 | -76.76172452 |
| Tompkins County, Newfield town | 0.43827327 | 0.45589026 | 42.29806214 | -76.67217706 |
| Cattaraugus County, East Otto town | 0.36229155 | 0.4556213 | 42.45722081 | -78.7466064 |
| Delaware County, Tompkins town | 0.37681549 | 0.4556213 | 42.06610701 | -75.33768302 |
| Cortland County, Truxton town | 0.35126412 | 0.4556213 | 42.77647826 | -76.06554424 |
| Delaware County, Hamden town | 0.34117805 | 0.45535234 | 42.2138356 | -74.98085579 |
| Columbia County, Ancram town | 0.42307692 | 0.45508338 | 41.99236606 | -73.55114635 |
| Genesee County, Byron town | 0.38864981 | 0.45346961 | 43.0510715 | -78.09395005 |
| Lewis County, Lewis town | 0.41527703 | 0.45266272 | 43.42943847 | -75.55539584 |
| Wyoming County, Attica town | 0.37869822 | 0.4521248 | 42.85204664 | -78.30267056 |
| Oswego County, Williamstown town | 0.43760086 | 0.45104895 | 43.4863076 | -75.91939326 |
| Wayne County, Huron town | 0.32920925 | 0.45024207 | 43.20717024 | -76.94969009 |
| Steuben County, Wayland town | 0.3119957 | 0.4499731 | 42.57071911 | -77.59530861 |
| Seneca County, Varick town | 0.37062937 | 0.44970414 | 42.79903745 | -76.96949394 |
| Madison County, Smithfield town | 0.39779451 | 0.44970414 | 42.99670118 | -75.7171364 |
| Clinton County, Plattsburgh town | 0.24798279 | 0.44862829 | 44.67065981 | -73.45396867 |
| Oswego County, Boylston town | 0.40061861 | 0.44835933 | 43.61457625 | -76.00843307 |
| Madison County, DeRuyter town | 0.36135019 | 0.44782141 | 42.81872139 | -75.89595991 |
| Washington County, Cambridge town | 0.35556751 | 0.44620764 | 42.94735722 | -73.48794863 |
| Saratoga County, Providence town | 0.36242604 | 0.44620764 | 43.07921377 | -73.98352269 |
| Chenango County, North Norwich town | 0.40613233 | 0.44593868 | 42.64302688 | -75.50833498 |
| Schoharie County, Jefferson town | 0.39402905 | 0.44459387 | 42.4695857 | -74.5883961 |
| Chenango County, Pharsalia town | 0.39053254 | 0.44271114 | 42.63736854 | -75.76112701 |
| Jefferson County, Ellisburg town | 0.29854761 | 0.44002152 | 43.71623626 | -76.0738113 |
| Cortland County, Harford town | 0.38125336 | 0.43921463 | 42.40917588 | -76.20364951 |
| Ulster County, Hardenburgh town | 0.40317375 | 0.43786982 | 42.0691524 | -74.5260225 |
| Wyoming County, Wethersfield town | 0.36511565 | 0.4373319 | 42.67739735 | -78.19166187 |
| Steuben County, Urbana town | 0.3774879 | 0.43706294 | 42.40454571 | -77.23479132 |
| Chenango County, New Berlin town | 0.29679935 | 0.43464228 | 42.54126358 | -75.44479501 |
| Fulton County, Northampton town | 0.38300161 | 0.43222162 | 43.13681587 | -74.18689167 |
| Cattaraugus County, Great Valley town | 0.14093599 | 0.43195266 | 42.17718033 | -78.69275427 |
| Schuyler County, Montour town | 0.38851533 | 0.43087682 | 42.36884441 | -76.83936003 |
| Chenango County, Pitcher town | 0.37869822 | 0.43033889 | 42.62022925 | -75.79757599 |
| Otsego County, Worcester town | 0.31213018 | 0.43033889 | 42.58666897 | -74.74024037 |
| Yates County, Middlesex town | 0.37762238 | 0.42980097 | 42.7334947 | -77.30101524 |
| Wayne County, Rose town | 0.3331092 | 0.4281872 | 43.18523218 | -76.95584547 |
| Wayne County, Wolcott town | 0.34776762 | 0.42791824 | 43.22415783 | -76.75550572 |
| Wyoming County, Orangeville town | 0.36982249 | 0.42738031 | 42.7255187 | -78.21039059 |
| Washington County, White Creek town | 0.35946746 | 0.42630446 | 42.98095177 | -73.28648488 |
| Cortland County, Solon town | 0.30217859 | 0.4260355 | 42.57025982 | -76.06302856 |
| Schoharie County, Gilboa town | 0.39564282 | 0.42549758 | 42.46458649 | -74.55857686 |
| Greene County, Prattsville town | 0.36632598 | 0.42549758 | 42.36405336 | -74.3706607 |
| Delaware County, Stamford town | 0.30043034 | 0.42495966 | 42.39920215 | -74.57908915 |
| Broome County, Maine town | 0.42469069 | 0.42469069 | 42.25476092 | -76.09229077 |
| Cortland County, Willet town | 0.37224314 | 0.42442173 | 42.41569144 | -75.95172442 |
| Delaware County, Davenport town | 0.20118343 | 0.42415277 | 42.44668178 | -75.00327856 |
| Chenango County, German town | 0.38999462 | 0.42092523 | 42.48434332 | -75.80640204 |
| Cayuga County, Summerhill town | 0.36498117 | 0.42065627 | 42.62918881 | -76.29658468 |
| Greene County, Halcott town | 0.37977407 | 0.42038731 | 42.19483029 | -74.45535774 |
| Chautauqua County, Poland town | 0.15505648 | 0.41958042 | 42.10510974 | -79.17480297 |
| St. Lawrence County, Potsdam town | 0.21005917 | 0.41958042 | 44.66991806 | -74.97902074 |
| Cayuga County, Victory town | 0.37802582 | 0.41931146 | 43.19321249 | -76.70115409 |
| Livingston County, Leicester town | 0.35341581 | 0.41850457 | 42.7671601 | -77.85543272 |
| Chenango County, Plymouth town | 0.35287789 | 0.41769769 | 42.56040741 | -75.56447857 |
| Schoharie County, Blenheim town | 0.39093599 | 0.41742873 | 42.48638182 | -74.55173085 |
| Cortland County, Cincinnatus town | 0.38555675 | 0.41742873 | 42.52501926 | -75.91966633 |
| Wyoming County, Pike town | 0.33902636 | 0.41662184 | 42.60730221 | -78.17867606 |
| Delaware County, Roxbury town | 0.36309844 | 0.41662184 | 42.36239004 | -74.49659225 |
| Chenango County, Otselic town | 0.3154922 | 0.41554599 | 42.64209266 | -75.7911049 |
| Washington County, Jackson town | 0.38811189 | 0.41554599 | 43.07787819 | -73.44448907 |
| Schuyler County, Tyrone town | 0.33781603 | 0.41447015 | 42.48325042 | -76.99473284 |
| Tioga County, Richford town | 0.38596019 | 0.41447015 | 42.4045482 | -76.17374427 |
| Jefferson County, Watertown town | 0.30096826 | 0.4133943 | 43.99649722 | -75.95857164 |
| Chenango County, Lincklaen town | 0.35933297 | 0.4112426 | 42.6686753 | -75.81462073 |
| Broome County, Windsor town | 0.37573964 | 0.41070468 | 42.05601864 | -75.55868676 |
| Hamilton County, Indian Lake town | 0.28456159 | 0.41070468 | 43.77148317 | -74.26860672 |
| Wayne County, Savannah town | 0.35341581 | 0.41043572 | 43.06616642 | -76.76872243 |
| Washington County, Argyle town | 0.35449166 | 0.40855299 | 43.25121532 | -73.5138629 |
| Chenango County, McDonough town | 0.37237762 | 0.40613233 | 42.45304118 | -75.75305392 |
| Steuben County, Pulteney town | 0.37493276 | 0.40559441 | 42.49664042 | -77.2407857 |
| Hamilton County, Lake Pleasant town | 0.31737493 | 0.40344271 | 43.48779673 | -74.36010604 |
| Herkimer County, Webb town | 0.32840237 | 0.40236686 | 43.84488509 | -74.85327574 |
| Warren County, Lake George town | 0.30379236 | 0.40236686 | 43.4994152 | -73.75024123 |
| Jefferson County, Lorraine town | 0.30957504 | 0.40155998 | 43.70689488 | -76.01287776 |
| Wyoming County, Eagle town | 0.3598709 | 0.40129102 | 42.59365096 | -78.24461411 |
| Hamilton County, Inlet town | 0.36498117 | 0.40048413 | 43.76445738 | -74.78269592 |
| Steuben County, Bradford town | 0.36229155 | 0.39940828 | 42.36502483 | -77.12096416 |
| Lewis County, Osceola town | 0.3119957 | 0.39940828 | 43.52840758 | -75.74796108 |
| Cattaraugus County, Farmersville town | 0.29693384 | 0.39940828 | 42.35056006 | -78.46274642 |
| Wyoming County, Castile town | 0.26412049 | 0.39913932 | 42.68264445 | -78.0653386 |
| Jefferson County, Alexandria town | 0.24852071 | 0.39887036 | 44.24707927 | -75.94767915 |
| Wayne County, Galen town | 0.30755783 | 0.39806347 | 43.07109752 | -76.95631743 |
| Hamilton County, Arietta town | 0.29464766 | 0.39456697 | 43.84878532 | -74.59594508 |
| Washington County, Whitehall town | 0.26291017 | 0.39429801 | 43.55067807 | -73.38166028 |
| Washington County, Hebron town | 0.29585799 | 0.39376009 | 43.22488951 | -73.36351795 |
| Hamilton County, Long Lake town | 0.30957504 | 0.39268424 | 43.85878175 | -74.65680482 |
| Cortland County, Lapeer town | 0.36040882 | 0.39187735 | 42.41249471 | -76.07769287 |
| Jefferson County, Adams town | 0.24623453 | 0.39133943 | 43.8868978 | -75.99125963 |
| Steuben County, Hornellsville town | 0.18692846 | 0.39080151 | 42.30309197 | -77.63584679 |
| Steuben County, Wayne town | 0.36847768 | 0.39026358 | 42.41327624 | -77.13886666 |
| Chautauqua County, Busti town | 0.11403981 | 0.38838085 | 42.09879342 | -79.29998855 |
| Montgomery County, Canajoharie town | 0.24798279 | 0.38703604 | 42.89883689 | -74.54774407 |
| Cortland County, Taylor town | 0.32356105 | 0.38596019 | 42.56875272 | -75.9068148 |
| Tioga County, Berkshire town | 0.345078 | 0.38434642 | 42.29979537 | -76.23520822 |
| Herkimer County, Little Falls town | 0.23749328 | 0.38273265 | 43.02393024 | -74.88773524 |
| Livingston County, Springwater town | 0.30782679 | 0.38138784 | 42.66711742 | -77.63190536 |
| Livingston County, Sparta town | 0.29854761 | 0.380312 | 42.67141043 | -77.66201779 |
| Livingston County, North Dansville town | 0.28577192 | 0.37977407 | 42.56164968 | -77.69139219 |
| Herkimer County, German Flatts town | 0.23614847 | 0.37869822 | 43.01194534 | -74.95468435 |
| Warren County, Warrensburg town | 0.27165143 | 0.37869822 | 43.48281969 | -73.78764411 |
| Steuben County, Prattsburgh town | 0.2471759 | 0.37869822 | 42.54488377 | -77.25880548 |
| Livingston County, Mount Morris town | 0.27810651 | 0.3781603 | 42.72323119 | -77.86708721 |
| Montgomery County, Palatine town | 0.28321678 | 0.37681549 | 42.9106116 | -74.48083471 |
| Hamilton County, Wells town | 0.25968263 | 0.37358795 | 43.52637249 | -74.3158146 |
| Washington County, Fort Ann town | 0.21732114 | 0.37143626 | 43.40883325 | -73.49312262 |
| Herkimer County, Ohio town | 0.27568585 | 0.37009145 | 43.67705325 | -74.8102709 |
| Saratoga County, Edinburg town | 0.28698225 | 0.3690156 | 43.17020403 | -74.11263673 |
| Saratoga County, Hadley town | 0.26035503 | 0.3690156 | 43.29191254 | -73.88230108 |
| Cattaraugus County, New Albion town | 0.32100592 | 0.36874664 | 42.31532354 | -78.87526433 |
| Tompkins County, Dryden town | 0.36793975 | 0.36793975 | 42.5324303 | -76.26186603 |
| Oneida County, Trenton town | 0.19768693 | 0.36793975 | 43.22342328 | -75.26922665 |
| Jefferson County, Pamelia town | 0.25053792 | 0.36767079 | 44.03564152 | -75.91486185 |
| Cortland County, Cuyler town | 0.31522324 | 0.36498117 | 42.68748212 | -75.93462551 |
| Ontario County, Naples town | 0.30823023 | 0.36256052 | 42.65446387 | -77.38517228 |
| Lewis County, West Turin town | 0.29370629 | 0.36148467 | 43.59006891 | -75.41115312 |
| Delaware County, Harpersfield town | 0.26438946 | 0.36094675 | 42.4091835 | -74.63867876 |
| Chenango County, Columbus town | 0.24636902 | 0.35852609 | 42.72082907 | -75.42206666 |
| Madison County, Brookfield town | 0.2415277 | 0.35798817 | 42.85673347 | -75.41245447 |
| Lewis County, Leyden town | 0.29612695 | 0.35664336 | 43.5414996 | -75.39416486 |
| Allegany County, Amity town | 0.23345885 | 0.35637439 | 42.22237171 | -78.02999936 |
| Delaware County, Franklin town | 0.26062399 | 0.35610543 | 42.37893318 | -75.14931075 |
| Allegany County, Centerville town | 0.30661646 | 0.35556751 | 42.50989285 | -78.2974386 |
| Cattaraugus County, Franklinville town | 0.18934911 | 0.35556751 | 42.32858131 | -78.46834894 |
| Allegany County, Wellsville town | 0.22646584 | 0.35368478 | 42.1177087 | -77.93329526 |
| Oneida County, Forestport town | 0.29666487 | 0.35287789 | 43.54644982 | -75.13800658 |
| Jefferson County, Worth town | 0.28509952 | 0.35260893 | 43.70532319 | -75.85406535 |
| Cortland County, Freetown town | 0.309844 | 0.35233997 | 42.49064765 | -75.99234996 |
| Herkimer County, Manheim town | 0.19150081 | 0.35233997 | 43.09640867 | -74.77018246 |
| Oneida County, Boonville town | 0.25941366 | 0.35180204 | 43.53918514 | -75.23575422 |
| Warren County, Thurman town | 0.22700377 | 0.35018827 | 43.48800361 | -73.81784937 |
| Lewis County, Lowville town | 0.27178591 | 0.34991931 | 43.7843665 | -75.47934786 |
| Chautauqua County, Chautauqua Lake UT | 0.31845078 | 0.34965035 | 42.2101785 | -79.4587416 |
| Warren County, Bolton town | 0.21395912 | 0.34884346 | 43.52120195 | -73.74303447 |
| Lewis County, Lyonsdale town | 0.31710597 | 0.34749866 | 43.64121092 | -75.29991035 |
| Otsego County, Laurens town | 0.23749328 | 0.34722969 | 42.51961809 | -75.16909419 |
| Hamilton County, Morehouse town | 0.30231307 | 0.34696073 | 43.39510055 | -74.77159114 |
| Chautauqua County, Chautauqua town | 0.21624529 | 0.34642281 | 42.26194704 | -79.50826151 |
| St. Lawrence County, Gouverneur town | 0.21086606 | 0.345078 | 44.3339908 | -75.47122798 |
| St. Lawrence County, Fine town | 0.30580958 | 0.34480904 | 44.09807914 | -74.99802885 |
| Lewis County, Croghan town | 0.19311458 | 0.34480904 | 44.03989676 | -75.20871521 |
| Lewis County, Greig town | 0.29827864 | 0.34265734 | 43.68011131 | -75.25605191 |
| Delaware County, Delhi town | 0.26681011 | 0.34265734 | 42.23564583 | -74.97416653 |
| St. Lawrence County, Pitcairn town | 0.29451318 | 0.34238838 | 44.15420315 | -75.20529676 |
| Lewis County, Diana town | 0.23856912 | 0.33969876 | 44.05691133 | -75.17135698 |
| Steuben County, Lindley town | 0.22081764 | 0.33835395 | 42.08880346 | -77.10353636 |
| Steuben County, Thurston town | 0.21839699 | 0.33727811 | 42.26861713 | -77.24078597 |
| Delaware County, Bovina town | 0.29895105 | 0.33674018 | 42.24968989 | -74.78225298 |
| Jefferson County, Cape Vincent town | 0.30365788 | 0.33647122 | 44.10079785 | -76.34371589 |
| Cattaraugus County, Mansfield town | 0.30123722 | 0.33620226 | 42.34322088 | -78.74413924 |
| Herkimer County, Danube town | 0.10919849 | 0.3359333 | 43.01405433 | -74.82757836 |
| Lewis County, Watson town | 0.26492738 | 0.33405056 | 43.96450752 | -75.16803049 |
| Yates County, Italy town | 0.25363098 | 0.33297472 | 42.58874636 | -77.246796 |
| Otsego County, Butternuts town | 0.29922001 | 0.32947821 | 42.41283144 | -75.35816868 |
| Allegany County, Alfred town | 0.2260624 | 0.32786444 | 42.27631745 | -77.76733243 |
| Steuben County, Hornell city | 0.21987628 | 0.32705756 | 42.3293247 | -77.65994795 |
| Essex County, North Hudson town | 0.19284562 | 0.32571275 | 44.02152825 | -73.7050238 |
| Cattaraugus County, Hinsdale town | 0.17832168 | 0.32517483 | 42.21597224 | -78.31091919 |
| Jefferson County, Champion town | 0.22754169 | 0.3246369 | 43.96635579 | -75.61570174 |
| Jefferson County, Brownville town | 0.19244217 | 0.32409898 | 44.00118646 | -75.98916218 |
| Chautauqua County, Kiantone town | 0.19956966 | 0.32409898 | 42.06503639 | -79.21545929 |
| St. Lawrence County, Clifton town | 0.26573427 | 0.32383002 | 44.07836305 | -74.87579914 |
| Otsego County, Morris town | 0.28442711 | 0.32194728 | 42.51717981 | -75.29528369 |
| St. Lawrence County, Ogdensburg city | 0.31334051 | 0.32167832 | 44.72107433 | -75.44866832 |
| Cattaraugus County, Carrollton town | 0.15519096 | 0.32006455 | 42.03313117 | -78.63609088 |
| Warren County, Johnsburg town | 0.20925229 | 0.31818182 | 43.73062447 | -74.02581721 |
| Cattaraugus County, Little Valley town | 0.25013448 | 0.31710597 | 42.25928933 | -78.7961726 |
| Essex County, Schroon town | 0.23843464 | 0.31629909 | 43.78170414 | -73.78475946 |
| Otsego County, Cherry Valley town | 0.12694997 | 0.31495428 | 42.80476837 | -74.67203789 |
| Steuben County, Wheeler town | 0.20938677 | 0.31387843 | 42.39574447 | -77.33069179 |
| St. Lawrence County, Colton town | 0.21584185 | 0.31387843 | 44.0803983 | -74.74669649 |
| Steuben County, Tuscarora town | 0.23614847 | 0.31145777 | 42.00557544 | -77.31230959 |
| Jefferson County, Hounsfield town | 0.22081764 | 0.30930608 | 43.90332102 | -76.40145378 |
| Steuben County, Woodhull town | 0.27057558 | 0.30796127 | 42.00993854 | -77.3421173 |
| St. Lawrence County, Massena town | 0.15384615 | 0.30392684 | 44.93279833 | -74.89551721 |
| Jefferson County, Henderson town | 0.25228618 | 0.30338892 | 43.86297391 | -76.2856119 |
| Chautauqua County, Carroll town | 0.08983324 | 0.30311996 | 42.07916891 | -79.15021321 |
| Allegany County, Friendship town | 0.14873588 | 0.30204411 | 42.23076189 | -78.08989201 |
| Jefferson County, Lyme town | 0.23843464 | 0.30204411 | 44.06501244 | -76.25820979 |
| St. Lawrence County, Morristown town | 0.24394836 | 0.30043034 | 44.49903418 | -75.6461701 |
| Jefferson County, Rodman town | 0.23009683 | 0.29989242 | 43.86029367 | -75.96725455 |
| Franklin County, Malone town | 0.20629371 | 0.29989242 | 44.85939618 | -74.30134989 |
| Essex County, Minerva town | 0.24744486 | 0.29908553 | 43.91768417 | -74.3174057 |
| Chautauqua County, Ellery town | 0.16742873 | 0.29854761 | 42.20630367 | -79.42872304 |
| Cattaraugus County, Ellicottville town | 0.22969338 | 0.29800968 | 42.33513203 | -78.68407218 |
| Franklin County, Tupper Lake town | 0.18558365 | 0.29747176 | 44.24209713 | -74.46974168 |
| Allegany County, Caneadea town | 0.24220011 | 0.2972028 | 42.42319162 | -78.16419931 |
| Allegany County, Angelica town | 0.16890802 | 0.29693384 | 42.27048671 | -78.04849408 |
| St. Lawrence County, Hammond town | 0.23641743 | 0.29666487 | 44.48662794 | -75.71444332 |
| Lewis County, Turin town | 0.2774341 | 0.29585799 | 43.63864022 | -75.42816549 |
| Warren County, Stony Creek town | 0.2027972 | 0.29397526 | 43.44959826 | -73.86239553 |
| Cattaraugus County, Napoli town | 0.19809037 | 0.29343733 | 42.24131907 | -78.83165899 |
| Allegany County, Rushford town | 0.24475524 | 0.29343733 | 42.40539093 | -78.19993202 |
| Lewis County, Denmark town | 0.23937601 | 0.29343733 | 43.910168 | -75.69670359 |
| Jefferson County, Orleans town | 0.18437332 | 0.28994083 | 44.27370715 | -75.97181341 |
| Chautauqua County, Westfield town | 0.13367402 | 0.28913394 | 42.33957072 | -79.58267951 |
| Herkimer County, Salisbury town | 0.0894298 | 0.28913394 | 43.1182353 | -74.76334065 |
| St. Lawrence County, Edwards town | 0.22565896 | 0.28859602 | 44.2466407 | -75.2086833 |
| Saratoga County, Day town | 0.19176977 | 0.28752017 | 43.30222679 | -73.94256256 |
| Jefferson County, Wilna town | 0.14766003 | 0.28752017 | 43.98825912 | -75.60904411 |
| Jefferson County, Le Ray town | 0.20441097 | 0.28563744 | 44.0339505 | -75.75528473 |
| Lewis County, Montague town | 0.2380312 | 0.28536848 | 43.73968345 | -75.78001869 |
| Steuben County, Troupsburg town | 0.2056213 | 0.28456159 | 42.00539686 | -77.46727637 |
| Wyoming County, Genesee Falls town | 0.20239376 | 0.28456159 | 42.59476171 | -78.08833848 |
| St. Lawrence County, Oswegatchie town | 0.18558365 | 0.28429263 | 44.68202831 | -75.49309232 |
| Oneida County, Bridgewater town | 0.19903174 | 0.28294782 | 42.92966322 | -75.29573841 |
| Cattaraugus County, Allegany Reservation | 0.15357719 | 0.28214094 | 42.15114594 | -78.66831873 |
| Otsego County, Richfield town | 0.14147391 | 0.28214094 | 42.85912273 | -75.00198963 |
| St. Lawrence County, Piercefield town | 0.18773534 | 0.28187197 | 44.13102502 | -74.63415392 |
| Essex County, Willsboro town | 0.16568047 | 0.28133405 | 44.36070375 | -73.39721382 |
| Chautauqua County, Charlotte town | 0.19257665 | 0.28106509 | 42.33303782 | -79.18113941 |
| St. Lawrence County, Fowler town | 0.2119419 | 0.27998924 | 44.3072692 | -75.44726913 |
| Lewis County, Martinsburg town | 0.2471759 | 0.27864443 | 43.77474995 | -75.41843616 |
| Lewis County, New Bremen town | 0.20172136 | 0.27837547 | 43.8867123 | -75.2561636 |
| Allegany County, Hume town | 0.23910705 | 0.27810651 | 42.49327046 | -78.17713609 |
| Allegany County, Almond town | 0.08848843 | 0.2767617 | 42.29826386 | -77.76151297 |
| Steuben County, Jasper town | 0.17227004 | 0.27514793 | 42.11936231 | -77.46764617 |
| Clinton County, Champlain town | 0.172539 | 0.27461001 | 45.00334484 | -73.37289801 |
| Jefferson County, Theresa town | 0.22861754 | 0.27434104 | 44.25015773 | -75.81829497 |
| Clinton County, Beekmantown town | 0.15559441 | 0.27434104 | 44.75794723 | -73.42422879 |
| Otsego County, Maryland town | 0.17482517 | 0.27380312 | 42.54800781 | -74.78369638 |
| Chautauqua County, Ripley town | 0.11256052 | 0.27218935 | 42.24854515 | -79.75910972 |
| Cattaraugus County, Lyndon town | 0.14631522 | 0.27138246 | 42.34234095 | -78.40270606 |
| Herkimer County, Litchfield town | 0.18370091 | 0.2711135 | 43.00735728 | -75.20887187 |
| Franklin County, Bellmont town | 0.23413125 | 0.27084454 | 44.76964304 | -74.03830035 |
| Washington County, Hampton town | 0.2436794 | 0.27057558 | 43.56186691 | -73.31391841 |
| Jefferson County, Antwerp town | 0.19607316 | 0.27003765 | 44.23594363 | -75.72623516 |
| Cattaraugus County, Conewango town | 0.18961807 | 0.26896181 | 42.25733056 | -78.95167919 |
| Essex County, Newcomb town | 0.21274879 | 0.26896181 | 43.85369751 | -74.20989864 |
| Chautauqua County, Portland town | 0.19889726 | 0.26788596 | 42.37591667 | -79.5119782 |
| Jefferson County, Rutland town | 0.21853147 | 0.26788596 | 44.01203792 | -75.76189978 |
| Hamilton County, Hope town | 0.22283486 | 0.26734804 | 43.34320903 | -74.18378875 |
| Franklin County, Duane town | 0.22700377 | 0.26681011 | 44.62035626 | -74.24980469 |
| Allegany County, Belfast town | 0.16675632 | 0.26492738 | 42.34475489 | -78.09128321 |
| Clinton County, Schuyler Falls town | 0.16487359 | 0.26492738 | 44.69170849 | -73.57684313 |
| Steuben County, Rathbone town | 0.22081764 | 0.26385153 | 42.10633755 | -77.37807047 |
| Washington County, Granville town | 0.22135557 | 0.26358257 | 43.35013022 | -73.28953716 |
| Franklin County, Santa Clara town | 0.18343195 | 0.26358257 | 44.59186958 | -74.35632669 |
| Washington County, Dresden town | 0.17805272 | 0.26250672 | 43.60477108 | -73.42744606 |
| Herkimer County, Russia town | 0.14685315 | 0.2619688 | 43.24280916 | -75.10480704 |
| Otsego County, Pittsfield town | 0.21584185 | 0.26169984 | 42.57784357 | -75.24549573 |
| Clinton County, Altona town | 0.2119419 | 0.26089295 | 44.87074513 | -73.67903937 |
| Washington County, Hartford town | 0.20683163 | 0.26062399 | 43.40357563 | -73.46297692 |
| Steuben County, Cohocton town | 0.15115654 | 0.25981711 | 42.56209658 | -77.53518317 |
| Jefferson County, Clayton town | 0.20077999 | 0.25954814 | 44.21396148 | -76.18219671 |
| Clinton County, Ellenburg town | 0.21463152 | 0.25954814 | 44.83006608 | -73.98571292 |
| Franklin County, Burke town | 0.19956966 | 0.25927918 | 44.91483486 | -74.21804697 |
| Clinton County, Dannemora town | 0.20857988 | 0.2584723 | 44.73233105 | -73.95319671 |
| St. Lawrence County, Clare town | 0.21597633 | 0.25712749 | 44.27311579 | -74.9429496 |
| Cattaraugus County, Coldspring town | 0.13474987 | 0.25605164 | 42.11737912 | -78.92436489 |
| Steuben County, Cameron town | 0.15653577 | 0.25551372 | 42.15453751 | -77.39608432 |
| Franklin County, Franklin town | 0.1753631 | 0.25470683 | 44.62690287 | -74.15039076 |
| St. Lawrence County, Louisville town | 0.10516407 | 0.25443787 | 44.91088761 | -74.90250075 |
| Steuben County, West Union town | 0.15115654 | 0.25363098 | 42.00021954 | -77.74737324 |
| Allegany County, Scio town | 0.19257665 | 0.25363098 | 42.14387996 | -77.95744408 |
| Essex County, North Elba town | 0.11726735 | 0.25363098 | 44.3224019 | -74.12109303 |
| Franklin County, Bangor town | 0.2021248 | 0.25363098 | 44.85273371 | -74.40109209 |
| Essex County, St. Armand town | 0.14201183 | 0.25336202 | 44.36608469 | -74.10673858 |
| St. Lawrence County, Rossie town | 0.22727273 | 0.25255514 | 44.27305775 | -75.52210365 |
| St. Lawrence County, Lisbon town | 0.13555675 | 0.25228618 | 44.74300364 | -75.44188184 |
| Steuben County, Addison town | 0.197149 | 0.25174825 | 42.11079329 | -77.25272468 |
| St. Lawrence County, De Peyster town | 0.18961807 | 0.24986552 | 44.51412584 | -75.44796658 |
| Herkimer County, Warren town | 0.10731576 | 0.24959656 | 42.88095341 | -74.99524455 |
| St. Lawrence County, Canton town | 0.16487359 | 0.24878967 | 44.60192969 | -75.12989063 |
| Steuben County, Avoca town | 0.13259817 | 0.24798279 | 42.38252687 | -77.39659328 |
| Livingston County, Nunda town | 0.18477676 | 0.24744486 | 42.60917789 | -77.86605717 |
| Warren County, Horicon town | 0.22243141 | 0.2471759 | 43.70072125 | -73.7155064 |
| Franklin County, Brighton town | 0.2027972 | 0.24690694 | 44.44696736 | -74.17714049 |
| Otsego County, Decatur town | 0.21221087 | 0.24636902 | 42.66395395 | -74.65318288 |
| Oneida County, Remsen town | 0.14550834 | 0.24636902 | 43.39829068 | -75.21553987 |
| Otsego County, Otsego town | 0.10328133 | 0.24583109 | 42.70851779 | -74.92254393 |
| Warren County, Chester town | 0.21597633 | 0.24529317 | 43.7472949 | -73.98833662 |
| Allegany County, Grove town | 0.13528779 | 0.24475524 | 42.49514277 | -77.86503051 |
| Livingston County, Portage town | 0.18316299 | 0.24421732 | 42.59990817 | -77.96215809 |
| Fulton County, Caroga town | 0.18477676 | 0.24394836 | 43.21423869 | -74.51111996 |
| St. Lawrence County, Macomb town | 0.19419042 | 0.24341044 | 44.51140582 | -75.57787187 |
| Otsego County, Middlefield town | 0.1344809 | 0.24314147 | 42.68669637 | -74.92929513 |
| St. Lawrence County, De Kalb town | 0.16541151 | 0.24206563 | 44.50930677 | -75.4171821 |
| Clinton County, Clinton town | 0.21812803 | 0.24206563 | 44.91750191 | -73.95659283 |
| Delaware County, Kortright town | 0.19445939 | 0.24045186 | 42.39733658 | -74.7050789 |
| Montgomery County, Minden town | 0.12157074 | 0.23964497 | 42.93564418 | -74.63090951 |
| Allegany County, Independence town | 0.2119419 | 0.23856912 | 42.09650381 | -77.78388075 |
| Franklin County, Waverly town | 0.17912856 | 0.23856912 | 44.54142995 | -74.46984422 |
| Warren County, Hague town | 0.19809037 | 0.2380312 | 43.75566381 | -73.63314234 |
| St. Lawrence County, Russell town | 0.17401829 | 0.23749328 | 44.31232717 | -75.1882206 |
| Livingston County, Ossian town | 0.18719742 | 0.23749328 | 42.54824614 | -77.75738197 |
| Franklin County, Chateaugay town | 0.21140398 | 0.23722431 | 44.88408789 | -74.03285524 |
| Fulton County, Ephratah town | 0.19150081 | 0.23614847 | 42.98607221 | -74.52014792 |
| Steuben County, Canisteo town | 0.12358795 | 0.23587951 | 42.27685566 | -77.61176379 |
| Essex County, Westport town | 0.13286713 | 0.23399677 | 44.18629871 | -73.45633911 |
| Franklin County, Fort Covington town | 0.19768693 | 0.23265196 | 44.96047949 | -74.49615021 |
| Hamilton County, Benson town | 0.19069392 | 0.23265196 | 43.33002774 | -74.37887991 |
| Essex County, Ticonderoga town | 0.18706294 | 0.232383 | 43.8601436 | -73.43767085 |
| St. Lawrence County, Norfolk town | 0.10623991 | 0.23211404 | 44.88403032 | -74.87856293 |
| St. Lawrence County, Pierrepont town | 0.17294244 | 0.23103819 | 44.62118952 | -74.96207217 |
| Lewis County, Pinckney town | 0.17912856 | 0.23103819 | 43.79294455 | -75.8278168 |
| Chautauqua County, Gerry town | 0.1401291 | 0.23050027 | 42.18290907 | -79.24868203 |
| Allegany County, Willing town | 0.21032813 | 0.23050027 | 42.08306159 | -77.8494132 |
| Clinton County, Mooers town | 0.1929801 | 0.23023131 | 44.9694807 | -73.58038682 |
| Cattaraugus County, Salamanca town | 0.17751479 | 0.2288865 | 42.13321311 | -78.70378302 |
| Otsego County, Edmeston town | 0.16272189 | 0.2288865 | 42.70154985 | -75.30214545 |
| Delaware County, Meredith town | 0.17563206 | 0.22834857 | 42.39814618 | -74.98685401 |
| Franklin County, Constable town | 0.15599785 | 0.22781065 | 44.9199347 | -74.24892867 |
| Clinton County, Saranac town | 0.17495966 | 0.22781065 | 44.68346862 | -73.9369739 |
| Franklin County, Harrietstown town | 0.14389457 | 0.22700377 | 44.36467079 | -74.23640236 |
| St. Lawrence County, Parishville town | 0.17348037 | 0.2267348 | 44.53079119 | -74.8290441 |
| St. Lawrence County, Stockholm town | 0.11713287 | 0.22646584 | 44.77657503 | -74.78300231 |
| Franklin County, Dickinson town | 0.16796665 | 0.22619688 | 44.69281262 | -74.55043722 |
| Fulton County, Stratford town | 0.14053254 | 0.22538999 | 43.2560842 | -74.62481056 |
| Otsego County, Roseboom town | 0.16393222 | 0.22458311 | 42.73435792 | -74.66260079 |
| Herkimer County, Columbia town | 0.18289403 | 0.22458311 | 42.95625486 | -75.03510264 |
| Allegany County, Alma town | 0.19956966 | 0.22431415 | 42.08223878 | -78.00454742 |
| Allegany County, Bolivar town | 0.19903174 | 0.22296934 | 42.00700855 | -78.11697709 |
| Fulton County, Oppenheim town | 0.11457773 | 0.22189349 | 43.09144658 | -74.74007867 |
| Washington County, Putnam town | 0.17146315 | 0.22027972 | 43.80602897 | -73.39170568 |
| Chautauqua County, Stockton town | 0.18316299 | 0.22001076 | 42.34872739 | -79.3013972 |
| Essex County, Chesterfield town | 0.10960194 | 0.21920387 | 44.43140309 | -73.40555774 |
| Allegany County, Genesee town | 0.15545992 | 0.21893491 | 42.01000662 | -78.30180455 |
| Chautauqua County, Clymer town | 0.13152232 | 0.21732114 | 42.00540679 | -79.63110281 |
| Jefferson County, Philadelphia town | 0.17697687 | 0.21705218 | 44.1435226 | -75.72214552 |
| Chautauqua County, Mina town | 0.10005379 | 0.21516945 | 42.14518605 | -79.6597903 |
| Essex County, Moriah town | 0.12479828 | 0.21328671 | 44.05025583 | -73.4700514 |
| Montgomery County, St. Johnsville town | 0.16756321 | 0.21247983 | 43.00108694 | -74.61028679 |
| Lewis County, Harrisburg town | 0.17374933 | 0.21140398 | 43.84730816 | -75.58792554 |
| Franklin County, St. Regis Mohawk Reservation | 0.18033889 | 0.21086606 | 44.973681 | -74.71976194 |
| Herkimer County, Newport town | 0.1309844 | 0.20952125 | 43.21118429 | -75.05119161 |
| St. Lawrence County, Hopkinton town | 0.16756321 | 0.20898332 | 44.61030716 | -74.60945938 |
| Allegany County, Burns town | 0.12210866 | 0.2084454 | 42.46043452 | -77.78072673 |
| Allegany County, Andover town | 0.14954276 | 0.20548682 | 42.12270224 | -77.80796616 |
| Franklin County, Bombay town | 0.16191501 | 0.20548682 | 44.94362278 | -74.53418065 |
| Otsego County, New Lisbon town | 0.14497041 | 0.2049489 | 42.61666007 | -75.20227902 |
| Oneida County, Steuben town | 0.14712211 | 0.2049489 | 43.29868154 | -75.30958368 |
| Essex County, Wilmington town | 0.10193653 | 0.2049489 | 44.42363055 | -73.76432646 |
| Allegany County, Allen town | 0.13232921 | 0.20252824 | 42.36252428 | -78.05555388 |
| Franklin County, Westville town | 0.15330823 | 0.20252824 | 44.89649718 | -74.38680644 |
| Allegany County, New Hudson town | 0.13959118 | 0.20145239 | 42.33948538 | -78.21699119 |
| Steuben County, Dansville town | 0.10543303 | 0.19661108 | 42.51774821 | -77.70311594 |
| Franklin County, Brandon town | 0.1570737 | 0.1958042 | 44.69453306 | -74.42014994 |
| Otsego County, Westford town | 0.16218397 | 0.19445939 | 42.61839078 | -74.79323106 |
| Cattaraugus County, Randolph town | 0.13986014 | 0.19230769 | 42.16934509 | -78.97342152 |
| Clinton County, Black Brook town | 0.14066703 | 0.19230769 | 44.44545689 | -73.75702578 |
| Chautauqua County, Ellington town | 0.11511565 | 0.19123185 | 42.25122882 | -79.07715844 |
| Franklin County, Moira town | 0.15075309 | 0.19096288 | 44.84599714 | -74.50081446 |
| Otsego County, Hartwick town | 0.15155998 | 0.19069392 | 42.64796677 | -74.97270828 |
| Clinton County, Chazy town | 0.11242604 | 0.190156 | 44.8287061 | -73.43265029 |
| Chautauqua County, Sherman town | 0.09965035 | 0.18988704 | 42.15570084 | -79.5645556 |
| Chautauqua County, North Harmony town | 0.1274879 | 0.18908015 | 42.15839442 | -79.40929522 |
| Essex County, Jay town | 0.16083916 | 0.18746638 | 44.41321242 | -73.70309374 |
| Essex County, Lewis town | 0.11135019 | 0.18665949 | 44.33817065 | -73.53424467 |
| Cattaraugus County, South Valley town | 0.10462614 | 0.18585261 | 42.02542069 | -78.91618606 |
| Essex County, Keene town | 0.14766003 | 0.18585261 | 44.07965703 | -73.91097586 |
| Cattaraugus County, Ischua town | 0.1309844 | 0.18558365 | 42.20226125 | -78.37646117 |
| St. Lawrence County, Brasher town | 0.11323292 | 0.18477676 | 44.95177738 | -74.72680695 |
| Allegany County, Cuba town | 0.13757396 | 0.18181818 | 42.21183176 | -78.28096419 |
| Herkimer County, Fairfield town | 0.09978483 | 0.18128026 | 43.08236386 | -74.96445801 |
| St. Lawrence County, Hermon town | 0.14981173 | 0.18047337 | 44.3488178 | -75.27346865 |
| Essex County, Essex town | 0.12210866 | 0.18047337 | 44.2900143 | -73.38888729 |
| Allegany County, West Almond town | 0.08189887 | 0.17939753 | 42.30177099 | -77.94709793 |
| Clinton County, Peru town | 0.11807423 | 0.17778375 | 44.5825746 | -73.61372326 |
| Essex County, Crown Point town | 0.10623991 | 0.1732114 | 44.00140502 | -73.4543373 |
| Allegany County, Wirt town | 0.13192577 | 0.17267348 | 42.09482923 | -78.09420686 |
| St. Lawrence County, Lawrence town | 0.13340506 | 0.17079075 | 44.7666176 | -74.72129489 |
| Otsego County, Burlington town | 0.08929532 | 0.17052179 | 42.75252744 | -75.19224974 |
| Otsego County, Plainfield town | 0.14497041 | 0.16863905 | 42.82775426 | -75.23224823 |
| Steuben County, Fremont town | 0.09306079 | 0.16702528 | 42.45238105 | -77.56471168 |
| St. Lawrence County, Waddington town | 0.1514255 | 0.16648736 | 44.85035537 | -75.246385 |
| Herkimer County, Winfield town | 0.13434642 | 0.16594944 | 42.88842084 | -75.182181 |
| Allegany County, Ward town | 0.13259817 | 0.16460463 | 42.20973474 | -77.94018748 |
| Chautauqua County, French Creek town | 0.12210866 | 0.15895643 | 42.00923381 | -79.66104623 |
| Cattaraugus County, Red House town | 0.10731576 | 0.15895643 | 42.11527097 | -78.73922908 |
| Clinton County, Au Sable town | 0.11874664 | 0.1584185 | 44.5397478 | -73.49843463 |
| Allegany County, Granger town | 0.11484669 | 0.15519096 | 42.45457253 | -78.06263239 |
| Chautauqua County, Harmony town | 0.09373319 | 0.15465304 | 42.00431002 | -79.44624606 |
| Essex County, Elizabethtown town | 0.08620226 | 0.15465304 | 44.2282383 | -73.70041321 |
| Herkimer County, Norway town | 0.09494352 | 0.14954276 | 43.22322229 | -74.98406538 |
| Cattaraugus County, Humphrey town | 0.09064013 | 0.14900484 | 42.25292574 | -78.58067354 |
| Steuben County, Howard town | 0.08983324 | 0.14497041 | 42.29018061 | -77.54600328 |
| St. Lawrence County, Madrid town | 0.11309844 | 0.14389457 | 44.79188894 | -75.16740213 |
| Steuben County, Greenwood town | 0.09938139 | 0.14174287 | 42.10563623 | -77.68844937 |
| Allegany County, Oil Springs Reservation | 0.14039806 | 0.14039806 | 42.237941 | -78.30527608 |
| Otsego County, Exeter town | 0.1133674 | 0.12775686 | 42.79133591 | -75.148894 |
| Otsego County, Springfield town | 0.09884346 | 0.12694997 | 42.85139805 | -74.81515257 |
| Steuben County, Hartsville town | 0.09144701 | 0.12614309 | 42.25921054 | -77.64756203 |
| Allegany County, Clarksville town | 0.0887574 | 0.11726735 | 42.12396591 | -78.30353748 |
| Herkimer County, Stark town | 0.07920925 | 0.11027434 | 42.89998638 | -74.83156418 |
| Allegany County, Birdsall town | 0.07988166 | 0.10812265 | 42.39381776 | -77.95402667 |
