## Supplementary material for "Mapping long-distance jump dispersal risk by leveraging transport infrastructure data and sampling bias correction: a case study with the invasive spotted lanternfly (*Lycorma delicatula*)": S10 North Carolina risk

*Supplemental material S10.* Mapping long-distance jump dispersal risk by leveraging transport infrastructure data and sampling bias correction: a case study with the invasive spotted lanternfly (*Lycorma delicatula*)
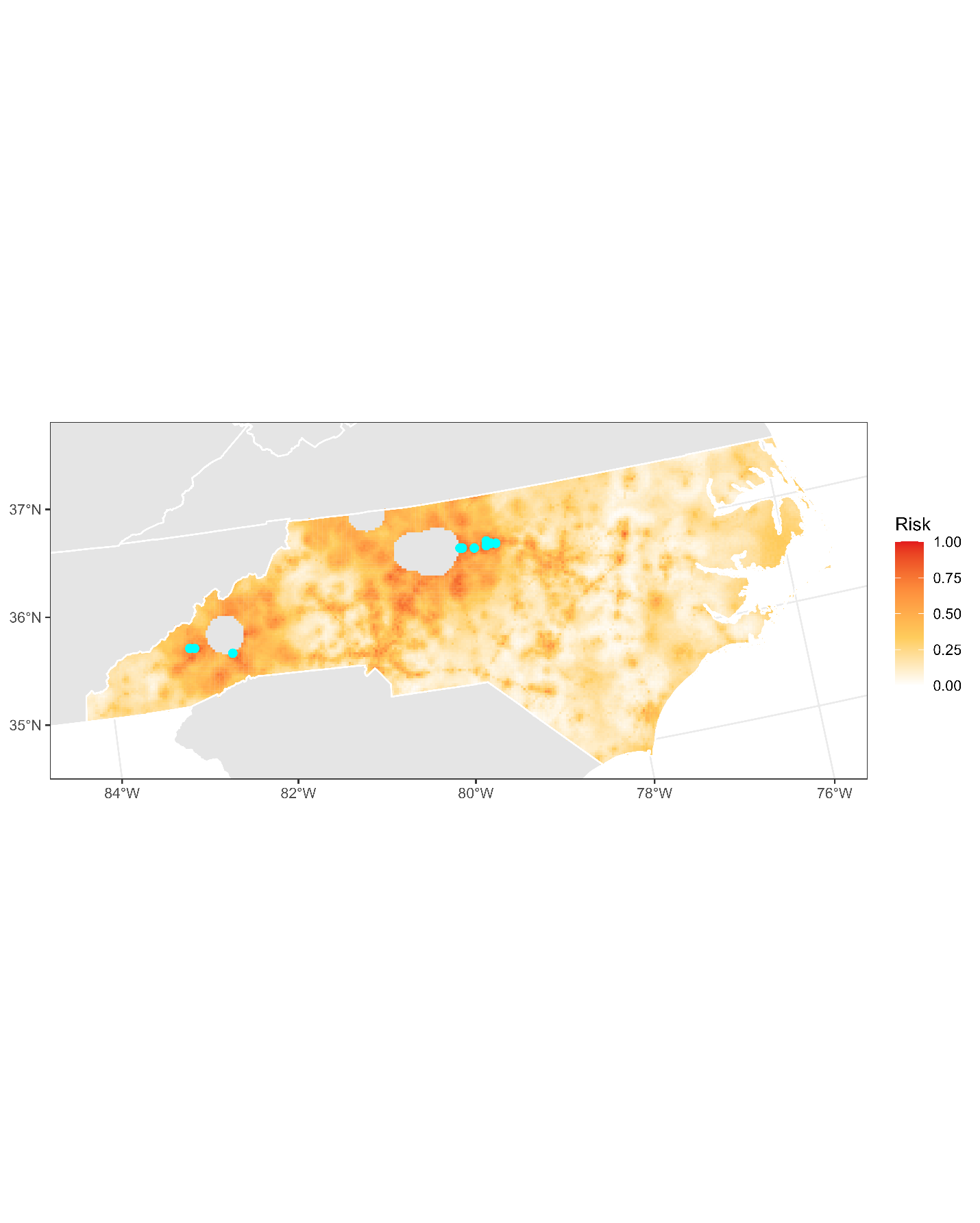

**Figure S10.1:** Projected risk of colonization by *L. delicatula* jump dispersal for North Carolina. Risk projections are based on an ensemble random forest model including distances from existing *L. delicatula* populations. Note that areas falling less than 20 km from existing *L. delicatula* populations are not included in the map because any colonization there would be categorized as diffusive spread based on our methodology (regardless of the actual source of the resulting population). Darker, redder colors indicate increased risk of colonization by jump dispersal. Cyan points show the top ten highest risk locations in the state.

**Table S10.1:** Ranked list of North Carolina county subdivisions sorted by the maximum projected risk of colonization by jump dispersal occurring there. For each subdivision, we identified the 2.5 x 2.5 km grid cell centroid included there with the highest projected risk of colonization by jump dispersal and report that location and risk value.

| **County, subdivision** | **Median risk value** | **Maximum risk value** | **Latitude of maximum risk** | **Longitude of maximum risk** |
| --- | --- | --- | --- | --- |
| Guilford County, Rock Creek township | 0.70629371 | 0.88407746 | 36.0616383 | -79.56431936 |
| Alamance County, Township 10, Melville | 0.74636902 | 0.87654653 | 36.0707663 | -79.30635547 |
| Guilford County, Gilmer township | 0.71328671 | 0.83485745 | 36.08457288 | -79.72989866 |
| Guilford County, Jefferson township | 0.76062399 | 0.83189887 | 36.08076644 | -79.70229515 |
| Alamance County, Township 12, Burlington | 0.75389995 | 0.8302851 | 36.10830443 | -79.41182615 |
| Alamance County, Township 13, Haw River | 0.74583109 | 0.82840237 | 36.07855606 | -79.36151779 |
| Alamance County, Township 6, Graham | 0.77891339 | 0.82759548 | 36.06433555 | -79.42155389 |
| Haywood County, Waynesville township | 0.6958042 | 0.82598171 | 35.53836461 | -82.97175481 |
| Haywood County, Ivy Hill township | 0.63179129 | 0.82087144 | 35.54445595 | -83.02686506 |
| Buncombe County, Asheville city | 0.81710597 | 0.81710597 | 35.44448836 | -82.53904109 |
| Haywood County, Clyde township | 0.76923077 | 0.81603012 | 35.53530931 | -82.94420299 |
| Randolph County, Asheboro township | 0.7021248 | 0.81468531 | 35.75455449 | -79.8011801 |
| Davie County, Mocksville township | 0.67616998 | 0.81360947 | 35.92781366 | -80.58595383 |
| Henderson County, Hoopers Creek township | 0.71705218 | 0.80796127 | 35.39394933 | -82.49191267 |
| Randolph County, Back Creek township | 0.70683163 | 0.8006993 | 35.78034516 | -79.82394502 |
| Nash County, Rocky Mount township | 0.26465842 | 0.79827864 | 35.9384819 | -77.80011398 |
| Buncombe County, Black Mountain township | 0.58445401 | 0.79800968 | 35.60548046 | -82.36980324 |
| Davie County, Farmington township | 0.61215707 | 0.79074771 | 35.98305649 | -80.48957088 |
| Randolph County, Randleman township | 0.76640667 | 0.78994083 | 35.80235077 | -79.81921716 |
| Guilford County, Fentress township | 0.67724583 | 0.78725121 | 35.97837539 | -79.78129549 |
| Alamance County, Township 9, Thompson | 0.55110274 | 0.78671329 | 36.03070081 | -79.34370382 |
| Rowan County, Litaker township | 0.5535234 | 0.77272727 | 35.6232954 | -80.50701209 |
| Buncombe County, Limestone township | 0.67348037 | 0.77192039 | 35.46344216 | -82.50761008 |
| Alamance County, Township 2, Coble | 0.64792899 | 0.76976869 | 36.02809676 | -79.48640874 |
| Rowan County, Locke township | 0.65868747 | 0.76250672 | 35.64895275 | -80.5299761 |
| Randolph County, Cedar Grove township | 0.60852609 | 0.76116192 | 35.69231697 | -79.84282915 |
| Rowan County, Salisbury township | 0.68827327 | 0.75766541 | 35.64170174 | -80.47500931 |
| Henderson County, Mills River township | 0.4817106 | 0.75470683 | 35.42237688 | -82.54296166 |
| Alamance County, Township 3, Boone Station | 0.6042227 | 0.75363098 | 36.05779349 | -79.53673258 |
| Rowan County, Franklin township | 0.61484669 | 0.75121033 | 35.69302308 | -80.52094234 |
| Henderson County, Hendersonville township | 0.60193653 | 0.7493276 | 35.36867192 | -82.46837183 |
| Transylvania County, Brevard township | 0.54532006 | 0.74045186 | 35.26422196 | -82.73887416 |
| Davidson County, Boone township | 0.53899946 | 0.73749328 | 35.75929173 | -80.33769582 |
| Rowan County, Gold Hill township | 0.51237224 | 0.73587951 | 35.59036044 | -80.42913812 |
| Davidson County, Lexington township | 0.61027434 | 0.73184508 | 35.81802286 | -80.26887984 |
| Alamance County, Township 5, Faucette | 0.55970952 | 0.72996235 | 36.12640702 | -79.37935428 |
| Haywood County, Iron Duff township | 0.6048951 | 0.72565896 | 35.56048785 | -82.96795259 |
| Guilford County, Monroe township | 0.56468531 | 0.72485207 | 36.12856605 | -79.72034751 |
| Wilson County, Wilson township | 0.43544917 | 0.72135557 | 35.70581166 | -77.91320871 |
| Rockingham County, Reidsville township | 0.60166756 | 0.72054868 | 36.3226915 | -79.64954023 |
| Haywood County, Jonathan Creek township | 0.6070468 | 0.71785906 | 35.56658084 | -83.02307852 |
| Mecklenburg County, Township 1, Charlotte | 0.41796665 | 0.7175901 | 35.26585574 | -80.88880418 |
| Davidson County, Cotton Grove township | 0.46637977 | 0.71678322 | 35.75562516 | -80.31018047 |
| Davidson County, Conrad Hill township | 0.57060247 | 0.7140936 | 35.81065178 | -80.21381595 |
| Randolph County, New Market township | 0.59534696 | 0.71113502 | 35.8834589 | -79.9151593 |
| Buncombe County, Avery Creek township | 0.70871436 | 0.70871436 | 35.44763821 | -82.56654372 |
| Rowan County, Providence township | 0.55621302 | 0.70548682 | 35.6968672 | -80.37890817 |
| Guilford County, Madison township | 0.52716514 | 0.7027972 | 36.14675226 | -79.68794091 |
| Davidson County, Tyro township | 0.5119688 | 0.70252824 | 35.78131784 | -80.33311799 |
| Madison County, Township 3, Mars Hill | 0.6070468 | 0.70199032 | 35.80766873 | -82.55889265 |
| Alamance County, Township 7, Albright | 0.55782679 | 0.70064551 | 36.0164888 | -79.40370536 |
| Randolph County, Level Cross township | 0.61027434 | 0.69983862 | 35.84636028 | -79.80975321 |
| McDowell County, Marion township | 0.52783755 | 0.6993007 | 35.69597935 | -81.98729086 |
| Davidson County, Thomasville township | 0.63569123 | 0.697149 | 35.85815671 | -80.06222836 |
| Transylvania County, Dunns Rock township | 0.56858526 | 0.69661108 | 35.23587502 | -82.68785793 |
| Davie County, Shady Grove township | 0.612695 | 0.69661108 | 35.97214503 | -80.4067673 |
| Randolph County, Trinity township | 0.58230231 | 0.6958042 | 35.86520756 | -79.94739904 |
| Randolph County, Providence township | 0.56656805 | 0.6936525 | 35.83877668 | -79.75471405 |
| Haywood County, Beaverdam township | 0.60785368 | 0.69230769 | 35.5482248 | -82.85772723 |
| Orange County, Cheeks township | 0.55849919 | 0.69096288 | 36.08492872 | -79.2462927 |
| Cabarrus County, Township 4, Kannapolis | 0.4873588 | 0.6880043 | 35.45421467 | -80.59791689 |
| McDowell County, Old Fort township | 0.49919311 | 0.68719742 | 35.61475994 | -82.25560013 |
| Davie County, Clarksville township | 0.42993545 | 0.68370091 | 35.95344953 | -80.60903669 |
| Davie County, Calahaln township | 0.47364174 | 0.68289403 | 35.90938154 | -80.6180503 |
| Randolph County, Franklinville township | 0.52797203 | 0.67724583 | 35.7947713 | -79.76420891 |
| Edgecombe County, Township 12, Rocky Mount | 0.2175901 | 0.66890802 | 35.91656391 | -77.80543451 |
| Madison County, Township 4, Beech Glenn | 0.61847768 | 0.66756321 | 35.82346406 | -82.4996783 |
| Durham County, Durham township | 0.50901022 | 0.66433566 | 35.98956473 | -78.89801206 |
| Buncombe County, Fairview township | 0.51398601 | 0.66379774 | 35.52025151 | -82.41322347 |
| Guilford County, Clay township | 0.59306079 | 0.66325982 | 35.9889692 | -79.69382562 |
| Surry County, Elkin township | 0.57127488 | 0.66218397 | 36.25361036 | -80.83180603 |
| Iredell County, Statesville township | 0.56764389 | 0.66003228 | 35.80569638 | -80.86541373 |
| Rockingham County, Williamsburg township | 0.48386229 | 0.65922539 | 36.31886707 | -79.62185372 |
| Buncombe County, Upper Hominy township | 0.52985476 | 0.65545992 | 35.49162533 | -82.75526495 |
| Davidson County, Alleghany township | 0.47283486 | 0.65465304 | 35.50956762 | -80.16389781 |
| Iredell County, Cool Springs township | 0.41742873 | 0.65330823 | 35.82064521 | -80.80585874 |
| Transylvania County, Boyd township | 0.457773 | 0.65277031 | 35.2769906 | -82.65267018 |
| Jackson County, Sylva township | 0.60650888 | 0.65196342 | 35.38252039 | -83.19457934 |
| Guilford County, Sumner township | 0.64201183 | 0.6514255 | 35.96016165 | -79.81360775 |
| Haywood County, Pigeon township | 0.58902636 | 0.65088757 | 35.48493631 | -82.89675077 |
| Henderson County, Clear Creek township | 0.56239914 | 0.65061861 | 35.38760821 | -82.43695448 |
| Henderson County, Blue Ridge township | 0.55567509 | 0.65034965 | 35.31491728 | -82.39388672 |
| Mecklenburg County, Township 9, Deweese | 0.48292093 | 0.64954276 | 35.48993566 | -80.87223707 |
| Rockingham County, New Bethel township | 0.5211135 | 0.6492738 | 36.31657999 | -79.93601918 |
| Catawba County, Hickory township | 0.41366326 | 0.64766003 | 35.73258509 | -81.35999236 |
| Cabarrus County, Township 12, Concord | 0.49865519 | 0.64712211 | 35.43217238 | -80.60239876 |
| Guilford County, Greene township | 0.52998924 | 0.64604626 | 35.99949867 | -79.60633138 |
| Iredell County, Bethany township | 0.50941366 | 0.6457773 | 35.82774169 | -80.86098612 |
| Transylvania County, Catheys Creek township | 0.5500269 | 0.64147391 | 35.19785512 | -82.75042847 |
| Mitchell County, Grassy Creek township | 0.54357181 | 0.64039806 | 35.88249921 | -82.03736228 |
| Jackson County, Scott Creek township | 0.57100592 | 0.63797741 | 35.41775096 | -83.10456332 |
| Haywood County, Crabtree township | 0.57154384 | 0.63743948 | 35.59554007 | -82.87762214 |
| Rockingham County, Huntsville township | 0.59844002 | 0.63528779 | 36.32033171 | -79.96372061 |
| Nash County, Stony Creek township | 0.35166756 | 0.6344809 | 35.94272828 | -77.82756486 |
| Randolph County, Grant township | 0.50268962 | 0.63421194 | 35.70295846 | -79.75569653 |
| Jackson County, Webster township | 0.54061323 | 0.63394298 | 35.33825198 | -83.2020328 |
| Yadkin County, North Knobs township | 0.60086068 | 0.63394298 | 36.23157293 | -80.8362671 |
| Alamance County, Township 11, Pleasant Grove | 0.45589026 | 0.63232921 | 36.14450054 | -79.34686687 |
| Durham County, Triangle township | 0.39456697 | 0.62829478 | 35.96360195 | -78.87550023 |
| Wake County, Swift Creek township | 0.380312 | 0.62802582 | 35.75976866 | -78.72339272 |
| Yadkin County, North Liberty township | 0.57409898 | 0.62802582 | 36.13690325 | -80.6282078 |
| Yancey County, Cane River township | 0.54464766 | 0.62668101 | 35.9118716 | -82.48384906 |
| Rockingham County, Simpsonville township | 0.45696611 | 0.62668101 | 36.3303211 | -79.70492167 |
| Caswell County, Pelham township | 0.46557289 | 0.62641205 | 36.52719147 | -79.49024414 |
| Iredell County, Chambersburg township | 0.53765465 | 0.62426036 | 35.773002 | -80.78719827 |
| Cumberland County, Cross Creek township | 0.44217321 | 0.62130178 | 35.03871183 | -78.91543603 |
| Robeson County, Lumberton township | 0.36578806 | 0.62103281 | 34.62090147 | -79.00880739 |
| Union County, Monroe township | 0.27245831 | 0.62103281 | 34.98437027 | -80.46901936 |
| Wake County, Raleigh township | 0.45992469 | 0.62049489 | 35.78341414 | -78.57608464 |
| Cleveland County, Cleveland | 0.20521786 | 0.61780527 | 35.31150429 | -81.60951474 |
| Cumberland County, Pearces Mill township | 0.48063475 | 0.61753631 | 35.03070887 | -78.86106639 |
| Davie County, Jerusalem township | 0.54276493 | 0.61699839 | 35.85449463 | -80.54435851 |
| Rowan County, China Grove township | 0.47848306 | 0.6161915 | 35.58284295 | -80.54350717 |
| Rockingham County, Mayo township | 0.40909091 | 0.61511565 | 36.40833939 | -79.94482033 |
| Rockingham County, Leaksville township | 0.51640667 | 0.61484669 | 36.51386599 | -79.72198523 |
| Buncombe County, Ivy township | 0.47821409 | 0.61188811 | 35.79819734 | -82.4760019 |
| Rockingham County, Madison township | 0.50779989 | 0.6105433 | 36.37182485 | -80.00971848 |
| Yadkin County, South Liberty township | 0.53980635 | 0.61027434 | 36.09283968 | -80.6372321 |
| McDowell County, Nebo township | 0.45104895 | 0.61027434 | 35.68610847 | -81.90459452 |
| Wilkes County, Edwards township | 0.47054868 | 0.60892953 | 36.23865313 | -80.89169706 |
| Davie County, Fulton township | 0.54424422 | 0.60677784 | 35.86928186 | -80.48470248 |
| Alamance County, Township 4, Morton | 0.49569661 | 0.60543303 | 36.13803094 | -79.46217406 |
| Chatham County, Matthews township | 0.51385153 | 0.60462614 | 35.73835382 | -79.52162089 |
| Cabarrus County, Township 5, New Gilead | 0.37735342 | 0.60247445 | 35.49468837 | -80.56151213 |
| Davidson County, Silver Hill township | 0.49690694 | 0.60247445 | 35.7886308 | -80.2184301 |
| Davidson County, Reedy Creek township | 0.51398601 | 0.60247445 | 35.86941669 | -80.31478014 |
| Gaston County, South Point township | 0.21315223 | 0.60193653 | 35.23933759 | -81.03435705 |
| Yadkin County, Boonville township | 0.51237224 | 0.59897795 | 36.24293885 | -80.74865699 |
| McDowell County, Crooked Creek township | 0.45696611 | 0.59736417 | 35.58944038 | -82.23206952 |
| Jackson County, Cullowhee township | 0.4190425 | 0.59682625 | 35.33524977 | -83.17454391 |
| Stanly County, Harris township | 0.47417967 | 0.59628833 | 35.47291006 | -80.22792694 |
| Chatham County, Albright township | 0.50860678 | 0.59386767 | 35.78234163 | -79.51199273 |
| Yancey County, Price Creek township | 0.52286175 | 0.59359871 | 35.89916137 | -82.37319357 |
| Buncombe County, Sandy Mush township | 0.42401829 | 0.59332975 | 35.6145813 | -82.84621322 |
| Yadkin County, South Fall Creek township | 0.47471759 | 0.59306079 | 36.15893427 | -80.62369171 |
| Iredell County, Olin township | 0.42966649 | 0.5922539 | 35.89032642 | -80.82009949 |
| Transylvania County, Little River township | 0.46933835 | 0.5922539 | 35.25174255 | -82.62911226 |
| Iredell County, Coddle Creek township | 0.42079075 | 0.59171598 | 35.59663823 | -80.82269276 |
| Madison County, Township 7, Ebbs Chapel | 0.53066165 | 0.59037117 | 35.91503306 | -82.51151871 |
| Wake County, Neuse township | 0.43881119 | 0.58983324 | 35.87529868 | -78.58317956 |
| Mecklenburg County, Township 10, Lemley | 0.35314685 | 0.58929532 | 35.46788376 | -80.87664185 |
| Randolph County, Liberty township | 0.50295858 | 0.5887574 | 35.89334791 | -79.65784343 |
| Avery County, Ingalls township | 0.46503497 | 0.58848843 | 35.92339714 | -82.00152352 |
| Ashe County, West Jefferson township | 0.44809037 | 0.58768155 | 36.38101491 | -81.48928148 |
| Haywood County, Fines Creek township | 0.4817106 | 0.5866057 | 35.68023825 | -83.03171839 |
| Catawba County, Newton township | 0.345078 | 0.58552985 | 35.69331297 | -81.22656673 |
| Durham County, Oak Grove township | 0.24260355 | 0.58391608 | 36.00752874 | -78.86546885 |
| Wake County, St. Matthews township | 0.3781603 | 0.58015062 | 35.83139147 | -78.59335662 |
| Yadkin County, Forbush township | 0.49542765 | 0.57907477 | 36.10767085 | -80.57740641 |
| Wake County, St. Marys township | 0.3485745 | 0.57853685 | 35.69152248 | -78.56900685 |
| Montgomery County, Eldorado township | 0.29800968 | 0.57853685 | 35.48754308 | -80.16850816 |
| Stokes County, Beaver Island township | 0.48574502 | 0.57772996 | 36.35355409 | -80.042144 |
| Gaston County, Gastonia township | 0.32208176 | 0.5739645 | 35.27536629 | -81.13951154 |
| Yancey County, Jacks Creek township | 0.41863905 | 0.57235073 | 35.9212585 | -82.36919894 |
| Lenoir County, Kinston township | 0.42778375 | 0.57154384 | 35.26918093 | -77.59410692 |
| Wake County, Cary township | 0.49757934 | 0.57046799 | 35.80772541 | -78.74075227 |
| Buncombe County, Broad River township | 0.45320065 | 0.56966111 | 35.57377552 | -82.29115741 |
| Mecklenburg County, Township 7, Crab Orchard | 0.40438408 | 0.56939215 | 35.31066889 | -80.71134338 |
| Rowan County, Unity township | 0.49784831 | 0.56858526 | 35.72589484 | -80.598961 |
| Madison County, Township 10, Grapevine | 0.48641743 | 0.56777838 | 35.87712869 | -82.57473681 |
| Wake County, Leesville township | 0.36081226 | 0.56777838 | 35.8736054 | -78.72561485 |
| Rutherford County, Cool Spring township | 0.48964497 | 0.56750941 | 35.32609132 | -81.91531202 |
| Yadkin County, Deep Creek township | 0.50874126 | 0.56670253 | 36.10001289 | -80.69254178 |
| McDowell County, Glenwood township | 0.44324906 | 0.56643357 | 35.62971355 | -81.99956455 |
| Mitchell County, Snow Creek township | 0.50753093 | 0.56347499 | 35.95854043 | -82.1080789 |
| Henderson County, Crab Creek township | 0.45884884 | 0.56320602 | 35.30865981 | -82.53507688 |
| Haywood County, Cecil township | 0.4288596 | 0.56293706 | 35.44681421 | -82.95942114 |
| McDowell County, North Cove township | 0.42711135 | 0.56159225 | 35.83832567 | -82.04553809 |
| Surry County, South Westfield township | 0.4739107 | 0.56132329 | 36.44168049 | -80.53696552 |
| Yancey County, Burnsville township | 0.49246907 | 0.56051641 | 35.91806397 | -82.34153299 |
| Surry County, Marsh township | 0.46624529 | 0.55997848 | 36.29413106 | -80.79514136 |
| Rowan County, Morgan township | 0.40882195 | 0.55970952 | 35.5794037 | -80.34676969 |
| Jackson County, Dillsboro township | 0.53173749 | 0.55970952 | 35.36637326 | -83.25330701 |
| Rockingham County, Wentworth township | 0.49959656 | 0.55944056 | 36.40010463 | -79.71824064 |
| Randolph County, Tabernacle township | 0.49273803 | 0.5591716 | 35.79542343 | -79.93396342 |
| Davidson County, Arcadia township | 0.53657881 | 0.55836471 | 35.93181047 | -80.27342137 |
| Wilkes County, Wilkesboro township | 0.48655191 | 0.55809575 | 36.13789633 | -81.1674521 |
| Stokes County, Yadkin township | 0.40882195 | 0.5570199 | 36.30246814 | -80.33792818 |
| Moore County, Township 8, Sandhills | 0.17953201 | 0.55675094 | 35.13548388 | -79.42770805 |
| Watauga County, Brushy Fork township | 0.39241528 | 0.55675094 | 36.228077 | -81.71762169 |
| New Hanover County, Cape Fear township | 0.29854761 | 0.55675094 | 34.28121555 | -77.9706312 |
| Iredell County, Barringer township | 0.44324906 | 0.55675094 | 35.69192232 | -80.85999502 |
| Mecklenburg County, Township 12, Paw Creek | 0.31414739 | 0.55621302 | 35.25789241 | -81.0026419 |
| Madison County, Township 1, North Marshall | 0.47525551 | 0.55567509 | 35.79809831 | -82.67334768 |
| Orange County, Hillsborough township | 0.45024207 | 0.55540613 | 36.06132301 | -79.08086192 |
| Rutherford County, Rutherfordton township | 0.52138246 | 0.55540613 | 35.40224039 | -81.98539241 |
| Davidson County, Emmons township | 0.51802044 | 0.55486821 | 35.74807984 | -80.08550614 |
| Wake County, Marks Creek township | 0.30473373 | 0.55486821 | 35.80276826 | -78.40125041 |
| Randolph County, Columbia township | 0.43840775 | 0.55433029 | 35.74220308 | -79.54909108 |
| Cabarrus County, Township 1, Harrisburg | 0.3302851 | 0.5535234 | 35.32913545 | -80.67952056 |
| Wilkes County, Traphill township | 0.48050027 | 0.54841313 | 36.35590952 | -80.92496373 |
| Rowan County, Steele township | 0.39591178 | 0.54841313 | 35.68181977 | -80.60795351 |
| Wake County, Cedar Fork township | 0.41810113 | 0.5478752 | 35.81979524 | -78.82312546 |
| Surry County, Westfield township | 0.4943518 | 0.54760624 | 36.54816821 | -80.48631999 |
| Mecklenburg County, Township 15, Huntersville | 0.38864981 | 0.54626143 | 35.42023762 | -80.85802339 |
| Polk County, Saluda township | 0.41151157 | 0.54545455 | 35.28642259 | -82.34296481 |
| Union County, Vance township | 0.39349112 | 0.54545455 | 35.07941199 | -80.67375858 |
| Alamance County, Township 1, Patterson | 0.47794513 | 0.54545455 | 35.89615768 | -79.51539771 |
| Yancey County, Crabtree township | 0.47767617 | 0.54491662 | 35.86103545 | -82.23895822 |
| Wake County, Meredith township | 0.49542765 | 0.54464766 | 35.80368942 | -78.71330036 |
| Transylvania County, Eastatoe township | 0.44459387 | 0.54410974 | 35.1536076 | -82.75812024 |
| Orange County, Bingham township | 0.29451318 | 0.54410974 | 36.01899384 | -79.2610196 |
| Wilkes County, New Castle township | 0.39160839 | 0.54410974 | 36.21661325 | -80.89613979 |
| Guilford County, Washington township | 0.47014524 | 0.54303389 | 36.13528511 | -79.60507705 |
| Surry County, Mount Airy township | 0.53267886 | 0.54249597 | 36.44529271 | -80.56474227 |
| Alleghany County, Cherry Lane township | 0.49959656 | 0.542227 | 36.40702937 | -80.97163736 |
| Rockingham County, Ruffin township | 0.39214632 | 0.53926842 | 36.53104584 | -79.51800058 |
| Cabarrus County, Township 2, Poplar Tent | 0.40667025 | 0.53899946 | 35.41012949 | -80.60687804 |
| Wake County, Wake Forest township | 0.33916084 | 0.53899946 | 35.93707957 | -78.54040095 |
| Yadkin County, South Knobs township | 0.51156536 | 0.53846154 | 36.19103796 | -80.87287938 |
| Yancey County, Pensacola township | 0.45360409 | 0.53711673 | 35.84537055 | -82.29825587 |
| Alleghany County, Gap Civil township | 0.46342119 | 0.53630984 | 36.48716705 | -81.06958262 |
| Surry County, Bryan township | 0.48668639 | 0.53469607 | 36.37794849 | -80.92052122 |
| Cabarrus County, Township 11, Central Cabarrus | 0.36646046 | 0.53146853 | 35.36243861 | -80.58844176 |
| Alleghany County, Whitehead township | 0.47122109 | 0.53146853 | 36.46512407 | -81.07398796 |
| Stanly County, North Albemarle township | 0.34548144 | 0.53093061 | 35.35168246 | -80.16875125 |
| Mecklenburg County, Township 13, Morning Star | 0.37668101 | 0.53066165 | 35.12710221 | -80.69217989 |
| Lee County, Township 6, West Sanford | 0.34534696 | 0.53039268 | 35.48595384 | -79.18148674 |
| Stokes County, Sauratown township | 0.45185584 | 0.53012372 | 36.30954205 | -80.05153605 |
| Stokes County, Big Creek township | 0.46301775 | 0.52985476 | 36.54090141 | -80.43070077 |
| Rutherford County, Logan Store township | 0.27864443 | 0.52904788 | 35.41775495 | -81.92637483 |
| Alamance County, Township 8, Newlin | 0.45535234 | 0.52877891 | 35.9388754 | -79.33567377 |
| Rutherford County, Gilkey township | 0.46503497 | 0.52797203 | 35.42104737 | -81.95384205 |
| Orange County, Chapel Hill township | 0.32060247 | 0.52797203 | 35.87763484 | -79.06545048 |
| Ashe County, Jefferson township | 0.45427649 | 0.52797203 | 36.3996752 | -81.45719849 |
| Stokes County, Meadows township | 0.43948359 | 0.52770307 | 36.31346803 | -80.25016218 |
| Yancey County, South Toe township | 0.45857988 | 0.5274341 | 35.85781049 | -82.2113197 |
| Haywood County, White Oak township | 0.50134481 | 0.52716514 | 35.63295228 | -83.01170572 |
| Henderson County, Green River township | 0.36229155 | 0.52689618 | 35.26750956 | -82.37436387 |
| Iredell County, Davidson township | 0.39752555 | 0.52501345 | 35.57813743 | -80.85459248 |
| McDowell County, Higgins township | 0.47202797 | 0.52339968 | 35.66731891 | -81.93626682 |
| Davidson County, Hampton township | 0.52339968 | 0.52339968 | 35.94281499 | -80.35616172 |
| Yancey County, Egypt township | 0.42415277 | 0.52232383 | 35.94335514 | -82.36520199 |
| Stanly County, Ridenhour township | 0.31118881 | 0.52205487 | 35.45456785 | -80.25991846 |
| Haywood County, East Fork township | 0.43464228 | 0.52097902 | 35.45357847 | -82.81801136 |
| Surry County, Pilot township | 0.4316837 | 0.5204411 | 36.39039292 | -80.49056565 |
| Forsyth County, Lewisville township | 0.51963421 | 0.51963421 | 36.05276673 | -80.50355121 |
| Davidson County, Healing Spring township | 0.40801506 | 0.51909629 | 35.66743095 | -80.15902425 |
| Transylvania County, Gloucester township | 0.47364174 | 0.51721356 | 35.16904223 | -82.89517858 |
| Rutherford County, Sulphur Springs township | 0.25699301 | 0.51533082 | 35.30399864 | -81.91940661 |
| Polk County, Columbus township | 0.36793975 | 0.51506186 | 35.2482433 | -82.20981841 |
| Surry County, Long Hill township | 0.45320065 | 0.51291017 | 36.41603915 | -80.51375769 |
| Davidson County, Jackson Hill township | 0.43410436 | 0.51183432 | 35.61596874 | -80.1133507 |
| Haywood County, Cataloochee township | 0.40371167 | 0.50995159 | 35.76172505 | -83.15851501 |
| Caswell County, Dan River township | 0.34480904 | 0.5091447 | 36.53368629 | -79.37434316 |
| Caldwell County, Lower Creek township | 0.37385691 | 0.50833782 | 35.93635258 | -81.54669968 |
| Vance County, Henderson township | 0.33324368 | 0.50833782 | 36.28000353 | -78.40307345 |
| Rutherford County, Union township | 0.44136633 | 0.50699301 | 35.30729302 | -81.94683271 |
| Henderson County, Edneyville township | 0.44916622 | 0.50618612 | 35.40015458 | -82.35056154 |
| Cumberland County, Seventy-First township | 0.32813341 | 0.50403443 | 35.13860942 | -78.97731545 |
| Halifax County, Weldon township | 0.22417967 | 0.50403443 | 36.42957802 | -77.59402419 |
| Alleghany County, Glade Creek township | 0.45306616 | 0.50322754 | 36.502211 | -81.00952932 |
| Wayne County, Goldsboro township | 0.38004303 | 0.50268962 | 35.38104309 | -78.01922674 |
| Catawba County, Clines township | 0.28321678 | 0.50242066 | 35.70844411 | -81.16717094 |
| Johnston County, Smithfield township | 0.23614847 | 0.50188273 | 35.47902459 | -78.36362502 |
| Wilkes County, Walnut Grove township | 0.37197418 | 0.50026896 | 36.42452243 | -81.11058572 |
| Swain County, Charleston township | 0.29693384 | 0.50026896 | 35.55957186 | -83.16467902 |
| Mecklenburg County, Township 2, Berryhill | 0.36605702 | 0.49919311 | 35.23231809 | -80.97963706 |
| Wilkes County, North Wilkesboro township | 0.48063475 | 0.49811727 | 36.15994739 | -81.16309306 |
| Alleghany County, Prathers Creek township | 0.46705218 | 0.49784831 | 36.48940197 | -81.26871666 |
| Harnett County, Averasboro township | 0.39887036 | 0.49623453 | 35.26613551 | -78.61055226 |
| Cumberland County, Manchester township | 0.19540075 | 0.49542765 | 35.16059283 | -78.97238434 |
| Transylvania County, Hogback township | 0.33674018 | 0.49381388 | 35.12478409 | -82.902784 |
| Rowan County, Atwell township | 0.27218935 | 0.49327595 | 35.63774243 | -80.61693562 |
| Lenoir County, Neuse township | 0.39833244 | 0.49300699 | 35.24725982 | -77.59943991 |
| Wake County, White Oak township | 0.38273265 | 0.49220011 | 35.73193385 | -78.84317601 |
| Scotland County, Stewartsville township | 0.17025282 | 0.49112426 | 34.77562544 | -79.44991418 |
| Wake County, House Creek township | 0.39147391 | 0.49112426 | 35.84356326 | -78.67573201 |
| Randolph County, Union township | 0.31132329 | 0.49112426 | 35.60050213 | -79.83423352 |
| Jackson County, Qualla township | 0.36632598 | 0.4908553 | 35.51831454 | -83.19972074 |
| Iredell County, Shiloh township | 0.28133405 | 0.48977945 | 35.77570869 | -80.98445336 |
| Rutherford County, Green Hill township | 0.44566971 | 0.4887036 | 35.38342468 | -82.01692725 |
| Franklin County, Louisburg township | 0.22270038 | 0.48843464 | 36.08389287 | -78.30665834 |
| Caldwell County, Lovelady township | 0.31468531 | 0.48682087 | 35.80903606 | -81.4298903 |
| McDowell County, Montford Cove township | 0.38878429 | 0.48601399 | 35.61089776 | -82.03118999 |
| Stokes County, Quaker Gap township | 0.40317375 | 0.48574502 | 36.45658052 | -80.30559672 |
| Orange County, Eno township | 0.2288865 | 0.48547606 | 36.05736638 | -79.05330017 |
| Wilkes County, Reddies River township | 0.43652501 | 0.48466918 | 36.19238454 | -81.24187045 |
| Jackson County, River township | 0.37977407 | 0.48413125 | 35.31010772 | -83.15079628 |
| Wilkes County, Union township | 0.38622916 | 0.48197956 | 36.35709383 | -81.29476163 |
| Polk County, Tryon township | 0.43639053 | 0.47982786 | 35.25147031 | -82.2372369 |
| Madison County, Township 5, Walnut | 0.40989779 | 0.47928994 | 35.84854774 | -82.72084727 |
| Stanly County, South Albemarle township | 0.32490586 | 0.47875202 | 35.33335428 | -80.20070627 |
| Ashe County, Old Fields township | 0.26385153 | 0.47848306 | 36.35895528 | -81.49355793 |
| Wilkes County, Moravian Falls township | 0.35906401 | 0.47794513 | 36.12276987 | -81.22718479 |
| Jackson County, Greens Creek township | 0.39725659 | 0.47740721 | 35.32210051 | -83.26072492 |
| Ashe County, Peak Creek township | 0.44540075 | 0.47740721 | 36.38258395 | -81.31821606 |
| Burke County, Morganton township | 0.32961269 | 0.47713825 | 35.72908585 | -81.6991906 |
| Wilkes County, Rock Creek township | 0.45131791 | 0.47686928 | 36.25973924 | -81.05804902 |
| Yadkin County, North Buck Shoals township | 0.43410436 | 0.47633136 | 36.11072454 | -80.77552592 |
| Cumberland County, Carvers Creek township | 0.1993007 | 0.47525551 | 35.13462579 | -78.95009061 |
| Rutherford County, Chimney Rock township | 0.42348037 | 0.47364174 | 35.51668785 | -82.18908309 |
| Randolph County, Coleridge township | 0.39591178 | 0.47310382 | 35.69821084 | -79.55869272 |
| Alleghany County, Piney Creek township | 0.45104895 | 0.47283486 | 36.53694544 | -81.28785917 |
| Iredell County, Fallstown township | 0.27487897 | 0.47283486 | 35.64782641 | -80.86883247 |
| Ashe County, Pine Swamp township | 0.3063475 | 0.4725659 | 36.35555831 | -81.46576977 |
| Madison County, Township 8, Spring Creek | 0.29881657 | 0.47202797 | 35.68708737 | -82.88988156 |
| Randolph County, Richland township | 0.35045724 | 0.47014524 | 35.59671845 | -79.8068001 |
| Edgecombe County, Township 1, Tarboro | 0.29303389 | 0.47014524 | 35.89997908 | -77.55319494 |
| Johnston County, Banner township | 0.23910705 | 0.46987628 | 35.37188167 | -78.55801179 |
| Davidson County, Yadkin College township | 0.46288327 | 0.46960732 | 35.87674062 | -80.36989638 |
| Wilkes County, Antioch township | 0.39833244 | 0.46880043 | 36.19010527 | -81.04351953 |
| Pitt County, Greenville township | 0.3063475 | 0.46745562 | 35.64196551 | -77.36096085 |
| Mecklenburg County, Township 14, Pineville | 0.46718666 | 0.46718666 | 35.08584946 | -80.89652788 |
| Randolph County, Concord township | 0.40909091 | 0.46718666 | 35.67457652 | -80.04449146 |
| Rutherford County, Colfax township | 0.29948897 | 0.4669177 | 35.31284641 | -81.80560034 |
| Lenoir County, Falling Creek township | 0.30527165 | 0.4669177 | 35.27347726 | -77.62130635 |
| Stokes County, Danbury township | 0.43208714 | 0.46664874 | 36.43089469 | -80.2824631 |
| Catawba County, Jacobs Fork township | 0.20199032 | 0.46664874 | 35.6340242 | -81.29452817 |
| Caldwell County, Lenoir township | 0.3091716 | 0.46611081 | 35.89221517 | -81.55516662 |
| Yancey County, Brush Creek township | 0.38757396 | 0.46584185 | 35.94617689 | -82.1951474 |
| New Hanover County, Wilmington township | 0.39133943 | 0.46530393 | 34.25081005 | -77.92206563 |
| Wilkes County, Mulberry township | 0.37009145 | 0.46207639 | 36.20404803 | -81.15436736 |
| Stokes County, Peters Creek township | 0.41769769 | 0.46180742 | 36.45290828 | -80.27782834 |
| Chatham County, Hickory Mountain township | 0.26882733 | 0.46153846 | 35.74100095 | -79.3794606 |
| Rutherford County, Camp Creek township | 0.34965035 | 0.4612695 | 35.4398456 | -81.92227616 |
| Johnston County, Selma township | 0.13797741 | 0.46019365 | 35.51466551 | -78.29870828 |
| Alleghany County, Cranberry township | 0.41796665 | 0.45696611 | 36.44185411 | -81.24959831 |
| Mitchell County, Poplar township | 0.33620226 | 0.45615923 | 36.08230658 | -82.39662085 |
| Madison County, Township 1, South Marshall | 0.3844809 | 0.45427649 | 35.78221614 | -82.73250607 |
| Madison County, Township 2, Laurel | 0.38905325 | 0.45427649 | 35.97182385 | -82.61437084 |
| Gaston County, Crowders Mountain township | 0.23641743 | 0.45427649 | 35.28923076 | -81.24905205 |
| Moore County, Township 7, McNeill | 0.16433566 | 0.45373857 | 35.17560596 | -79.39083973 |
| Iredell County, Turnersburg township | 0.37210866 | 0.45373857 | 35.93441024 | -80.81120549 |
| Iredell County, Concord township | 0.34279182 | 0.45346961 | 35.81629018 | -80.94810068 |
| Franklin County, Youngsville township | 0.24354492 | 0.45320065 | 36.06468155 | -78.48216561 |
| Mitchell County, Cane Creek township | 0.3718397 | 0.45266272 | 35.99945732 | -82.07224235 |
| Yadkin County, South Buck Shoals township | 0.42751479 | 0.45266272 | 36.0630869 | -80.75681307 |
| Duplin County, Island Creek township | 0.15788058 | 0.4521248 | 34.73717814 | -77.97494218 |
| Ashe County, Chestnut Hill township | 0.43975256 | 0.45185584 | 36.54723969 | -81.37140799 |
| Macon County, Franklin township | 0.33727811 | 0.45131791 | 35.15685133 | -83.40002457 |
| Gaston County, Riverbend township | 0.17966649 | 0.45024207 | 35.36013866 | -81.09481501 |
| Jackson County, Caney Fork township | 0.39954276 | 0.44943518 | 35.34833592 | -83.08833334 |
| Rowan County, Cleveland township | 0.36793975 | 0.44755245 | 35.77356932 | -80.61749196 |
| Pasquotank County, Mount Hermon township | 0.28778913 | 0.44728349 | 36.24709419 | -76.32129221 |
| Watauga County, New River township | 0.35153308 | 0.4464766 | 36.22138007 | -81.66212407 |
| Robeson County, Pembroke township | 0.28187197 | 0.44593868 | 34.69258159 | -79.18851703 |
| Yadkin County, East Bend township | 0.4316837 | 0.44593868 | 36.21057697 | -80.49935163 |
| Ashe County, Obids township | 0.3986014 | 0.44540075 | 36.34874498 | -81.41020098 |
| McDowell County, Dysartsville township | 0.29558903 | 0.44513179 | 35.68280535 | -81.87703384 |
| Wake County, Middle Creek township | 0.23695535 | 0.44513179 | 35.5880865 | -78.79102138 |
| Rutherford County, Morgan township | 0.39349112 | 0.44405594 | 35.51020654 | -82.13406202 |
| Wayne County, Stoney Creek township | 0.20158687 | 0.44378698 | 35.42071047 | -77.981505 |
| Surry County, Rockford township | 0.39618074 | 0.44351802 | 36.25424044 | -80.6610206 |
| Johnston County, Micro township | 0.18504572 | 0.44190425 | 35.56389656 | -78.17388286 |
| Jackson County, Canada township | 0.4020979 | 0.44136633 | 35.20035694 | -82.97365725 |
| Ashe County, Grassy Creek township | 0.42469069 | 0.44109736 | 36.57271187 | -81.39494767 |
| Jackson County, Barkers Creek township | 0.38582571 | 0.43894567 | 35.41064378 | -83.2458805 |
| Pitt County, Winterville township | 0.25470683 | 0.43867671 | 35.55869249 | -77.40993565 |
| Rowan County, Scotch Irish township | 0.35314685 | 0.43840775 | 35.84327552 | -80.63155112 |
| Jackson County, Savannah township | 0.27030662 | 0.43760086 | 35.29996328 | -83.26443066 |
| Avery County, Plumtree township | 0.37278107 | 0.43760086 | 35.99619394 | -82.04456052 |
| Surry County, Dobson township | 0.39564282 | 0.43679398 | 36.37870505 | -80.74945491 |
| Avery County, Banner Elk township | 0.37224314 | 0.43652501 | 36.13432907 | -81.87719206 |
| Caswell County, Anderson township | 0.380312 | 0.43652501 | 36.25829123 | -79.35003946 |
| Rutherford County, High Shoals township | 0.32087144 | 0.43625605 | 35.27529923 | -81.8686692 |
| Stanly County, Big Lick township | 0.20710059 | 0.43544917 | 35.26361597 | -80.35571143 |
| Lenoir County, Vance township | 0.31441635 | 0.43491124 | 35.31731971 | -77.61064618 |
| Randolph County, New Hope township | 0.30876815 | 0.43464228 | 35.58650586 | -80.0630947 |
| Yadkin County, North Fall Creek township | 0.40720818 | 0.43437332 | 36.24345016 | -80.57789518 |
| Caswell County, Locust Hill township | 0.3655191 | 0.43410436 | 36.40181932 | -79.40372255 |
| Caswell County, Yanceyville township | 0.38192577 | 0.43410436 | 36.39793822 | -79.37601955 |
| Lincoln County, Catawba Springs township | 0.23332437 | 0.43222162 | 35.53311912 | -81.03259066 |
| Harnett County, Black River township | 0.34965035 | 0.4316837 | 35.49216587 | -78.75636082 |
| Wilkes County, Lovelace township | 0.34938139 | 0.43141474 | 36.12396513 | -81.05669874 |
| Johnston County, Clayton township | 0.28214094 | 0.43114578 | 35.65732271 | -78.49191775 |
| Ashe County, Walnut Hill township | 0.39429801 | 0.43087682 | 36.48107882 | -81.3843617 |
| Polk County, White Oak township | 0.37762238 | 0.43060785 | 35.34576705 | -82.07995039 |
| Caswell County, Stoney Creek township | 0.40048413 | 0.43060785 | 36.2919165 | -79.42812697 |
| Brunswick County, Town Creek township | 0.15572889 | 0.42845616 | 34.18915028 | -77.96440697 |
| Wayne County, Brogden township | 0.16810113 | 0.42845616 | 35.33716669 | -78.0296616 |
| Gaston County, Dallas township | 0.27299623 | 0.42791824 | 35.32644527 | -81.18566921 |
| Iredell County, Eagle Mills township | 0.35745024 | 0.42684239 | 36.03390776 | -80.70601158 |
| Rowan County, Mount Ulla township | 0.28200646 | 0.42657343 | 35.65565755 | -80.75439841 |
| Jackson County, Hamburg township | 0.38770845 | 0.42549758 | 35.16824772 | -83.09090182 |
| Stanly County, Furr township | 0.28617536 | 0.42469069 | 35.25252989 | -80.44225892 |
| Madison County, Township 9, Sandy Mush | 0.41984938 | 0.42388381 | 35.69996146 | -82.80322426 |
| Person County, Roxboro township | 0.33808499 | 0.42280796 | 36.36489224 | -78.98347264 |
| Wilson County, Black Creek township | 0.12654653 | 0.422539 | 35.69231816 | -77.97321478 |
| Johnston County, Beulah township | 0.1577461 | 0.42227004 | 35.58166821 | -78.14135167 |
| Polk County, Cooper Gap township | 0.37157074 | 0.42200108 | 35.39971805 | -82.15424453 |
| Stokes County, Snow Creek township | 0.37358795 | 0.42200108 | 36.50788164 | -80.18054791 |
| Johnston County, Cleveland township | 0.2563206 | 0.42092523 | 35.5996235 | -78.56194612 |
| Wilkes County, Boomer township | 0.23964497 | 0.41984938 | 36.12622276 | -81.25487668 |
| Rockingham County, Price township | 0.37116729 | 0.41958042 | 36.51833809 | -79.92113282 |
| Johnston County, Wilson Mills township | 0.25242066 | 0.4190425 | 35.57506056 | -78.39778542 |
| Nash County, South Whitakers township | 0.30096826 | 0.41877353 | 36.00423229 | -77.78413393 |
| Orange County, Cedar Grove township | 0.30876815 | 0.41850457 | 36.15085871 | -79.23154014 |
| Richmond County, Wolf Pit township | 0.08848843 | 0.41742873 | 34.93554947 | -79.77932768 |
| Cabarrus County, Township 9, Georgeville | 0.25336202 | 0.41742873 | 35.29660885 | -80.43322143 |
| Burke County, Quaker Meadows township | 0.29612695 | 0.41635288 | 35.77323906 | -81.69082541 |
| Cumberland County, Eastover township | 0.1309844 | 0.41608392 | 35.04867773 | -78.82892085 |
| Surry County, Shoals township | 0.39483593 | 0.41554599 | 36.34634722 | -80.49970062 |
| Mecklenburg County, Township 6, Clear Creek | 0.29935449 | 0.41554599 | 35.21171591 | -80.64707562 |
| Brunswick County, Northwest township | 0.17697687 | 0.41554599 | 34.26346724 | -78.00265646 |
| Chatham County, Williams township | 0.29451318 | 0.41447015 | 35.85566121 | -79.0704012 |
| Mecklenburg County, Township 11, Long Creek | 0.29343733 | 0.41393222 | 35.3099679 | -80.88002234 |
| Chatham County, Baldwin township | 0.2972028 | 0.41366326 | 35.85961449 | -79.09789066 |
| Avery County, Altamont township | 0.38273265 | 0.41151157 | 35.98636517 | -81.96152936 |
| Watauga County, Boone township | 0.4112426 | 0.4112426 | 36.22473176 | -81.68987165 |
| Lenoir County, Southwest township | 0.33364712 | 0.41097364 | 35.23865057 | -77.54506517 |
| Wake County, Holly Springs township | 0.28375471 | 0.40855299 | 35.61407297 | -78.81339351 |
| Onslow County, Jacksonville township | 0.25578268 | 0.40747714 | 34.74802726 | -77.46754607 |
| Burke County, Icard township | 0.20010758 | 0.40747714 | 35.73942486 | -81.41510659 |
| Ashe County, Helton township | 0.39416353 | 0.40720818 | 36.56087476 | -81.48284185 |
| Cabarrus County, Township 10, Midland | 0.30580958 | 0.40693921 | 35.25616745 | -80.46959982 |
| Mitchell County, Bakersville township | 0.38058096 | 0.40613233 | 35.98712035 | -82.15936369 |
| Richmond County, Marks Creek township | 0.16810113 | 0.40613233 | 34.87628128 | -79.67999913 |
| Yancey County, Green Mountain township | 0.32894029 | 0.40451856 | 35.97793606 | -82.27413921 |
| Johnston County, Pine Level township | 0.28348575 | 0.4042496 | 35.53247219 | -78.26622717 |
| Mecklenburg County, Township 8, Mallard Creek | 0.35260893 | 0.40371167 | 35.36903515 | -80.81202822 |
| Madison County, Township 11, Revere-Rice Cove | 0.36256052 | 0.40371167 | 35.90863737 | -82.65381482 |
| New Hanover County, Harnett township | 0.30258203 | 0.40317375 | 34.26429759 | -77.86316504 |
| Jackson County, Cashiers township | 0.26331361 | 0.40317375 | 35.14308489 | -83.0672333 |
| Pitt County, Arthur township | 0.12076385 | 0.40129102 | 35.56736153 | -77.46452323 |
| Chatham County, Bear Creek township | 0.26143088 | 0.40102205 | 35.6465197 | -79.51340719 |
| Cabarrus County, Township 3, Odell | 0.32880581 | 0.40102205 | 35.42448666 | -80.7165137 |
| Hoke County, Raeford township | 0.25820334 | 0.40021517 | 35.00843163 | -79.17484597 |
| Cabarrus County, Township 7, Gold Hill | 0.24852071 | 0.40021517 | 35.4839505 | -80.31016344 |
| Anson County, Wadesboro township | 0.17563206 | 0.39994621 | 34.97318923 | -80.05141048 |
| Sampson County, Plain View township | 0.22270038 | 0.39994621 | 35.26206756 | -78.58330138 |
| Montgomery County, Biscoe township | 0.2647929 | 0.39967725 | 35.29160865 | -79.73130268 |
| Robeson County, Raft Swamp township | 0.32342657 | 0.39806347 | 34.65875317 | -79.11215219 |
| Lee County, Township 2, Jonesboro | 0.28160301 | 0.39806347 | 35.46003422 | -79.15902705 |
| Chatham County, Hadley township | 0.309844 | 0.39806347 | 35.82505815 | -79.33253603 |
| Cumberland County, Rockfish township | 0.29854761 | 0.39752555 | 34.97275878 | -78.93024631 |
| Onslow County, Camp Lejeune UT | 0.22565896 | 0.39671867 | 34.7393529 | -77.413538 |
| Iredell County, Union Grove township | 0.35153308 | 0.39671867 | 36.04816533 | -80.81656829 |
| Halifax County, Roanoke Rapids township | 0.18020441 | 0.39564282 | 36.43815655 | -77.64926346 |
| Surry County, Eldora township | 0.36498117 | 0.39483593 | 36.40484662 | -80.60160418 |
| Vance County, Sandy Creek township | 0.28913394 | 0.39456697 | 36.27589091 | -78.37546813 |
| Burke County, Linville township | 0.32517483 | 0.39402905 | 35.8064116 | -81.96678718 |
| Union County, Marshville township | 0.13649812 | 0.39187735 | 34.97708665 | -80.41453348 |
| Craven County, Township 8 | 0.21516945 | 0.39187735 | 35.09875551 | -77.0997823 |
| Avery County, Newland No. 2 township | 0.37331899 | 0.39080151 | 36.07797519 | -81.97274449 |
| Union County, Goose Creek township | 0.13905325 | 0.39053254 | 35.06862054 | -80.59190348 |
| Montgomery County, Star township | 0.29451318 | 0.38999462 | 35.45708228 | -79.78030487 |
| Lincoln County, Lincolnton township | 0.28429263 | 0.38918774 | 35.46922487 | -81.24213841 |
| Edgecombe County, Township 7, Swift Creek | 0.15734266 | 0.38891877 | 36.02189054 | -77.75132273 |
| Cherokee County, Valleytown township | 0.21180742 | 0.38838085 | 35.20328574 | -83.83921423 |
| Pitt County, Farmville township | 0.21503497 | 0.38784293 | 35.61513288 | -77.62301131 |
| Avery County, Frank township | 0.37493276 | 0.38784293 | 36.08125506 | -82.00045312 |
| Avery County, Pyatte township | 0.35381926 | 0.38703604 | 36.03052801 | -81.95328862 |
| Harnett County, Duke township | 0.30204411 | 0.38676708 | 35.31818658 | -78.65500087 |
| Catawba County, Mountain Creek township | 0.22270038 | 0.38569123 | 35.55517556 | -81.02822719 |
| Pitt County, Belvoir township | 0.10408822 | 0.38542227 | 35.65065651 | -77.41560252 |
| Burke County, Silver Creek township | 0.30473373 | 0.38300161 | 35.72034358 | -81.81363151 |
| Durham County, Lebanon township | 0.21920387 | 0.3816568 | 36.0814153 | -78.90551677 |
| Warren County, Hawtree township | 0.22552448 | 0.38004303 | 36.52253741 | -78.20234154 |
| Hoke County, McLauchlin township | 0.2584723 | 0.38004303 | 35.02648662 | -79.14278327 |
| Craven County, Township 7 | 0.16675632 | 0.37950511 | 35.06358629 | -77.02395236 |
| Caldwell County, Hudson township | 0.3394298 | 0.37869822 | 35.8379107 | -81.48081788 |
| Burke County, Jonas Ridge township | 0.27837547 | 0.37842926 | 35.96099477 | -81.93798174 |
| Bertie County, Windsor township | 0.22081764 | 0.37842926 | 35.98762901 | -76.96095218 |
| Yancey County, Ramseytown township | 0.33620226 | 0.37654653 | 36.06021114 | -82.40061597 |
| Macon County, Ellijay township | 0.27097902 | 0.37573964 | 35.17307462 | -83.34148829 |
| Avery County, Hughes township | 0.37210866 | 0.37547068 | 36.0558941 | -81.97686297 |
| Robeson County, Union township | 0.12923615 | 0.37520172 | 34.68234745 | -79.27458601 |
| Wilkes County, Brushy Mountain township | 0.26223776 | 0.37520172 | 36.08684382 | -81.12081223 |
| Columbus County, Whiteville township | 0.17764927 | 0.37520172 | 34.30483723 | -78.71706113 |
| Madison County, Township 6, Hot Springs | 0.30204411 | 0.37493276 | 35.87995992 | -82.799956 |
| Catawba County, Catawba township | 0.19445939 | 0.37493276 | 35.72005725 | -81.08021292 |
| Vance County, Middleburg township | 0.24583109 | 0.37439484 | 36.38557333 | -78.34953638 |
| Pasquotank County, Elizabeth City township | 0.36740183 | 0.37385691 | 36.27697798 | -76.22728609 |
| Surry County, Siloam township | 0.33458849 | 0.37305003 | 36.28750473 | -80.56881827 |
| Wilkes County, Somers township | 0.33458849 | 0.37170522 | 36.08083351 | -80.89508057 |
| Cherokee County, Murphy township | 0.22512103 | 0.37116729 | 35.08166638 | -83.97007386 |
| Watauga County, Blowing Rock township | 0.25739645 | 0.37116729 | 36.17724349 | -81.67055556 |
| Lenoir County, Woodington township | 0.28146853 | 0.37116729 | 35.19911631 | -77.5829228 |
| Duplin County, Limestone township | 0.24139322 | 0.37062937 | 34.970776 | -77.72268521 |
| Brunswick County, Smithville township | 0.26667563 | 0.37036041 | 33.92553576 | -78.02606809 |
| Ashe County, Hurricane township | 0.26654115 | 0.36982249 | 36.56765346 | -81.5385739 |
| Pitt County, Pactolus township | 0.18477676 | 0.3690156 | 35.63761055 | -77.33364469 |
| Chatham County, Center township | 0.29195804 | 0.36767079 | 35.73169451 | -79.15493509 |
| Martin County, Williamston township | 0.17670791 | 0.36713287 | 35.84332027 | -77.05458702 |
| Wayne County, New Hope township | 0.12547068 | 0.36686391 | 35.35069297 | -77.96994409 |
| Moore County, Township 3, Sheffield | 0.23413125 | 0.36498117 | 35.43037244 | -79.58870755 |
| Watauga County, Stony Fork township | 0.32571275 | 0.36498117 | 36.25203602 | -81.54265376 |
| Vance County, Kittrell township | 0.2288865 | 0.36471221 | 36.2580643 | -78.40824303 |
| Watauga County, Blue Ridge township | 0.14093599 | 0.36282948 | 36.19595481 | -81.63860384 |
| Pasquotank County, Providence township | 0.26062399 | 0.36229155 | 36.32065514 | -76.21567638 |
| Randolph County, Pleasant Grove township | 0.33754707 | 0.35960194 | 35.56237897 | -79.56002246 |
| Dare County, Atlantic township | 0.31253362 | 0.35933297 | 35.99950723 | -75.67117556 |
| Watauga County, Watauga township | 0.26277569 | 0.35906401 | 36.20934496 | -81.74956742 |
| Pasquotank County, Nixonton township | 0.2197418 | 0.35906401 | 36.27236421 | -76.19980767 |
| Brunswick County, Shallotte township | 0.18222162 | 0.35825713 | 33.9204321 | -78.58157716 |
| Duplin County, Smith township | 0.19109736 | 0.35691232 | 34.97504392 | -77.74978625 |
| Pender County, Topsail township | 0.20467994 | 0.35664336 | 34.37897766 | -77.72438067 |
| Montgomery County, Little River township | 0.26143088 | 0.35664336 | 35.50489361 | -79.79825857 |
| Lee County, Township 5, East Sanford | 0.26734804 | 0.35583647 | 35.47807721 | -79.12677106 |
| Wake County, Little River township | 0.24098978 | 0.35556751 | 35.83429085 | -78.30863353 |
| Pender County, Union township | 0.11296396 | 0.35449166 | 34.71943784 | -78.00715559 |
| Duplin County, Rockfish township | 0.14604626 | 0.35368478 | 34.74559332 | -78.02901081 |
| Granville County, Oxford township | 0.24045186 | 0.35260893 | 36.30861303 | -78.59639455 |
| New Hanover County, Masonboro township | 0.29612695 | 0.35126412 | 34.18993489 | -77.8250485 |
| Mecklenburg County, Township 3, Steele Creek | 0.23050027 | 0.35045724 | 35.12200649 | -81.00140735 |
| Randolph County, Brower township | 0.3189887 | 0.34965035 | 35.57770457 | -79.66967397 |
| Cumberland County, Black River township | 0.19069392 | 0.34965035 | 35.24823016 | -78.64284016 |
| Robeson County, Wisharts township | 0.08189887 | 0.34911243 | 34.61293827 | -78.95472202 |
| Harnett County, Neills Creek township | 0.31118881 | 0.34830554 | 35.42626114 | -78.77139615 |
| Johnston County, Pleasant Grove township | 0.2380312 | 0.34830554 | 35.55977776 | -78.59945379 |
| Northampton County, Oconeechee township | 0.10718128 | 0.34830554 | 36.44717884 | -77.56098333 |
| Hyde County, Lake Mattamuskeet UT | 0.3063475 | 0.34803658 | 35.54434941 | -76.07881114 |
| Dare County, Nags Head township | 0.27783755 | 0.34749866 | 35.9729498 | -75.6497782 |
| Person County, Flat River township | 0.29182356 | 0.34696073 | 36.29502753 | -78.97083939 |
| Wake County, Panther Branch township | 0.24381388 | 0.34669177 | 35.66384201 | -78.6887392 |
| Jackson County, Mountain township | 0.30580958 | 0.34588488 | 35.19943193 | -83.16945938 |
| Hyde County, Lake Landing township | 0.31764389 | 0.34588488 | 35.58803612 | -76.06722792 |
| Macon County, Highlands township | 0.21516945 | 0.34561592 | 35.04446121 | -83.19549739 |
| Carteret County, Morehead township | 0.2415277 | 0.345078 | 34.73435866 | -76.82515459 |
| McDowell County, Brackett township | 0.31320602 | 0.34454008 | 35.60105682 | -81.94858024 |
| Ashe County, Piney Creek township | 0.30486821 | 0.34265734 | 36.49130623 | -81.46786165 |
| Onslow County, White Oak township | 0.19849381 | 0.3415815 | 34.74821493 | -77.32717588 |
| Washington County, Plymouth township | 0.18813878 | 0.3415815 | 35.85991277 | -76.73682952 |
| Orange County, Little River township | 0.27877891 | 0.34104357 | 36.14524162 | -79.03340737 |
| Pitt County, Ayden township | 0.17657343 | 0.34077461 | 35.49296934 | -77.42614467 |
| Wake County, Bartons Creek township | 0.16675632 | 0.34023669 | 35.90537891 | -78.63304661 |
| Graham County, Cheoah township | 0.19445939 | 0.3394298 | 35.33621853 | -83.81797375 |
| Bladen County, Colly township | 0.26519634 | 0.33916084 | 34.64440797 | -78.55638973 |
| Mitchell County, Fork Mountain-Little Rock Creek township | 0.28536848 | 0.33889188 | 36.05338423 | -82.14717185 |
| Tyrrell County, Scuppernong township | 0.28536848 | 0.33835395 | 35.9182798 | -76.2649709 |
| Avery County, Montezuma township | 0.32436794 | 0.33835395 | 36.0934749 | -81.91319778 |
| Beaufort County, Washington township | 0.11726735 | 0.33700914 | 35.60694913 | -77.1425193 |
| Bertie County, Merry Hill township | 0.18343195 | 0.33674018 | 35.86440439 | -76.76419089 |
| Cumberland County, Cedar Creek township | 0.09171598 | 0.33620226 | 35.02669793 | -78.83388582 |
| Johnston County, Ingrams township | 0.10946746 | 0.33566434 | 35.46119036 | -78.39606067 |
| Nash County, Oak Level township | 0.197149 | 0.33431953 | 35.97312287 | -77.87717512 |
| Vance County, Dabney township | 0.22807961 | 0.33405056 | 36.33209099 | -78.44797589 |
| Mitchell County, Harrell township | 0.27259279 | 0.33351264 | 36.05985657 | -82.20258847 |
| Brunswick County, Lockwoods Folly township | 0.1838354 | 0.33351264 | 33.97968985 | -78.37409801 |
| Polk County, Green Creek township | 0.2563206 | 0.33297472 | 35.29504104 | -82.03318642 |
| Iredell County, New Hope township | 0.26627219 | 0.33297472 | 36.03674818 | -80.90394296 |
| Avery County, Linville township | 0.30164067 | 0.33243679 | 36.06809691 | -81.88963305 |
| Carteret County, Portsmouth township | 0.31401291 | 0.33189887 | 35.00600523 | -76.1357073 |
| Hyde County, Fairfield township | 0.29693384 | 0.33162991 | 35.62380329 | -76.14313471 |
| Ashe County, Clifton township | 0.17401829 | 0.33162991 | 36.45058536 | -81.50425954 |
| Cabarrus County, Township 6, Rimertown | 0.24986552 | 0.33136095 | 35.50222388 | -80.44727692 |
| Pamlico County, Township 2 | 0.26169984 | 0.33082302 | 35.08011255 | -76.70941576 |
| Avery County, Newland No. 1 township | 0.30258203 | 0.33082302 | 36.10005579 | -81.96862361 |
| Camden County, South Mills township | 0.22001076 | 0.33082302 | 36.42642465 | -76.30254645 |
| Anson County, Lilesville township | 0.1225121 | 0.3302851 | 34.96196367 | -79.96975771 |
| New Hanover County, Federal Point township | 0.23977945 | 0.33001614 | 34.13254448 | -77.89424901 |
| Durham County, Carr township | 0.18934911 | 0.33001614 | 35.92960874 | -78.79799699 |
| Wilson County, Taylors township | 0.24206563 | 0.32894029 | 35.76231491 | -77.98483061 |
| Moore County, Township 4, Ritter | 0.19230769 | 0.32894029 | 35.42653135 | -79.56134747 |
| Avery County, Pineola township | 0.29881657 | 0.32813341 | 36.04601854 | -81.89377522 |
| Tyrrell County, Gum Neck township | 0.27716514 | 0.32786444 | 35.68469831 | -76.09853198 |
| Wilson County, Toisnot township | 0.18961807 | 0.32705756 | 35.83312653 | -77.85409872 |
| Granville County, Fishing Creek township | 0.2056213 | 0.3267886 | 36.30454513 | -78.56876829 |
| Carteret County, Cedar Island township | 0.29639591 | 0.32598171 | 34.96351502 | -76.28791804 |
| Moore County, Township 2, Bensalem | 0.18840775 | 0.32517483 | 35.40836919 | -79.59347877 |
| Lee County, Township 7, Pocket | 0.17469069 | 0.32490586 | 35.423921 | -79.22349332 |
| Craven County, Township 6 | 0.12937063 | 0.3246369 | 34.88807955 | -76.92709031 |
| Sampson County, Little Coharie township | 0.15411512 | 0.32221625 | 34.98362909 | -78.39430062 |
| Granville County, Brassfield township | 0.16379774 | 0.32167832 | 36.02895554 | -78.54746699 |
| Jones County, Township 7, Beaver Creek | 0.25255514 | 0.32140936 | 35.23433651 | -77.51788238 |
| Lenoir County, Sand Hill township | 0.25255514 | 0.32087144 | 35.27817118 | -77.50716845 |
| Carteret County, Newport township | 0.1880043 | 0.32033351 | 34.78710159 | -76.86808335 |
| Ashe County, Horse Creek township | 0.25658956 | 0.32033351 | 36.55234546 | -81.59857261 |
| Wilkes County, Jobs Cabin township | 0.23641743 | 0.31925767 | 36.27915835 | -81.39532978 |
| Scotland County, Laurel Hill township | 0.10126412 | 0.3189887 | 34.79763648 | -79.44514114 |
| Carteret County, Beaufort township | 0.08768155 | 0.31818182 | 34.72926142 | -76.65785595 |
| Gaston County, Cherryville township | 0.1697149 | 0.31791286 | 35.39128108 | -81.34159198 |
| Clay County, Hayesville township | 0.26681011 | 0.31737493 | 35.04532262 | -83.83649155 |
| Iredell County, Sharpesburg township | 0.24072082 | 0.31737493 | 35.9265258 | -80.92605389 |
| Granville County, Dutchville township | 0.25147929 | 0.31656805 | 36.10528056 | -78.75763984 |
| Dare County, East Lake township | 0.26923077 | 0.31656805 | 35.68791629 | -75.98369981 |
| Greene County, Olds township | 0.16890802 | 0.31629909 | 35.57130088 | -77.633706 |
| Caswell County, Milton township | 0.16487359 | 0.31629909 | 36.54006913 | -79.25842235 |
| Alexander County, Taylorsville township | 0.26062399 | 0.31576116 | 35.91735799 | -81.21090313 |
| Wilkes County, Stanton township | 0.27487897 | 0.3154922 | 36.19927674 | -81.29730909 |
| Caldwell County, Little River township | 0.14671867 | 0.31522324 | 35.92617646 | -81.46379979 |
| Lenoir County, Institute township | 0.18141474 | 0.31495428 | 35.34782054 | -77.6597645 |
| Northampton County, Gaston township | 0.15828402 | 0.31334051 | 36.45577251 | -77.61623175 |
| Dare County, Croatan township | 0.25416891 | 0.31226466 | 35.63796219 | -75.82611383 |
| Person County, Olive Hill township | 0.23023131 | 0.31172673 | 36.37282971 | -79.03881644 |
| Chatham County, Gulf township | 0.21543841 | 0.31145777 | 35.57150306 | -79.30374699 |
| Granville County, Tally Ho township | 0.19499731 | 0.31145777 | 36.20697748 | -78.67712562 |
| Lincoln County, North Brook township | 0.11296396 | 0.31118881 | 35.54759826 | -81.50835895 |
| Wilkes County, Beaver Creek township | 0.19486283 | 0.31091985 | 36.10761335 | -81.28689358 |
| Caldwell County, Patterson township | 0.1035503 | 0.31038193 | 35.98386896 | -81.56587681 |
| Pamlico County, Township 4 | 0.25779989 | 0.31038193 | 35.22796434 | -76.50194498 |
| Lee County, Township 4, Deep River | 0.24125874 | 0.31011296 | 35.57388974 | -79.16189077 |
| Chatham County, Oakland township | 0.26506186 | 0.31011296 | 35.62178915 | -79.17948373 |
| Greene County, Carrs township | 0.16527703 | 0.31011296 | 35.59750439 | -77.65567714 |
| Caswell County, Hightowers township | 0.25255514 | 0.309844 | 36.24659041 | -79.26710522 |
| Macon County, Millshoal township | 0.26519634 | 0.30957504 | 35.1952169 | -83.33781093 |
| Alexander County, Wittenburg township | 0.24744486 | 0.30957504 | 35.81740749 | -81.31527294 |
| Lincoln County, Ironton township | 0.20979021 | 0.30930608 | 35.48783166 | -81.21037833 |
| Avery County, Cranberry township | 0.29679935 | 0.30930608 | 36.12541759 | -81.99222476 |
| Lenoir County, Moseley Hall township | 0.16151157 | 0.30849919 | 35.26867286 | -77.7354301 |
| Robeson County, Rowland township | 0.13676708 | 0.30823023 | 34.5062725 | -79.31302544 |
| Wilkes County, Lewis Fork township | 0.25860678 | 0.30823023 | 36.1772214 | -81.30163126 |
| Warren County, Warrenton township | 0.20790748 | 0.30823023 | 36.40041946 | -78.14561049 |
| Wake County, Buckhorn township | 0.1492738 | 0.30742335 | 35.71797091 | -78.90303475 |
| Avery County, Roaring Creek township | 0.26681011 | 0.30715438 | 36.06244558 | -82.03226682 |
| Bladen County, Elizabethtown township | 0.21463152 | 0.30661646 | 34.60452682 | -78.59343659 |
| Edgecombe County, Township 14, Upper Town Creek | 0.15384615 | 0.30661646 | 35.87272612 | -77.81606637 |
| Hyde County, Swan Quarter township | 0.27434104 | 0.30661646 | 35.37542886 | -76.2936732 |
| Lenoir County, Trent township | 0.22942442 | 0.30607854 | 35.12856476 | -77.71282439 |
| Montgomery County, Mount Gilead township | 0.1936525 | 0.30554061 | 35.21548955 | -80.00052109 |
| Johnston County, Boon Hill township | 0.09534696 | 0.30527165 | 35.48028511 | -78.22191919 |
| Person County, Allensville township | 0.21947283 | 0.30527165 | 36.35293799 | -78.90047882 |
| Chatham County, New Hope township | 0.2288865 | 0.30500269 | 35.74183006 | -79.06766375 |
| Jones County, Township 5, Tuckahoe | 0.21651425 | 0.30365788 | 35.01464043 | -77.71212486 |
| Moore County, Township 9, Mineral Springs | 0.2027972 | 0.30338892 | 35.21695495 | -79.52240018 |
| Wilson County, Old Fields township | 0.14335664 | 0.30338892 | 35.76651926 | -78.01222816 |
| Harnett County, Hectors Creek township | 0.23924153 | 0.30311996 | 35.52218286 | -78.80604568 |
| Chowan County, Township 1, Edenton | 0.1993007 | 0.302851 | 36.06503273 | -76.59814546 |
| Onslow County, Stump Sound township | 0.22270038 | 0.30258203 | 34.68224437 | -77.48353579 |
| Currituck County, Poplar Branch township | 0.20656267 | 0.30231307 | 36.09329761 | -75.81760699 |
| Warren County, Roanoke township | 0.24475524 | 0.30231307 | 36.5244959 | -77.91499368 |
| Pasquotank County, Newland township | 0.19176977 | 0.30204411 | 36.40917205 | -76.33586919 |
| Burke County, Drexel township | 0.29074771 | 0.30123722 | 35.76316214 | -81.60808406 |
| Hyde County, Ocracoke township | 0.27864443 | 0.30043034 | 35.07481472 | -76.00458355 |
| Avery County, Minneapolis township | 0.25658956 | 0.30016138 | 36.10333656 | -81.99634014 |
| Bladen County, Cypress Creek township | 0.20333513 | 0.29989242 | 34.69395355 | -78.43313639 |
| Stanly County, Almond township | 0.18692846 | 0.29989242 | 35.3407345 | -80.25542535 |
| Onslow County, Richlands township | 0.20710059 | 0.29935449 | 34.81813451 | -77.4785598 |
| Warren County, Fork township | 0.22027972 | 0.29881657 | 36.23441383 | -78.09958113 |
| Edgecombe County, Township 10, Lower Town Creek | 0.19351802 | 0.29827864 | 35.79902425 | -77.63488252 |
| Mitchell County, Bradshaw township | 0.2506724 | 0.29774072 | 36.02853777 | -82.32148543 |
| Stanly County, Center township | 0.21584185 | 0.29747176 | 35.27446601 | -80.10053666 |
| Burke County, Lower Creek township | 0.2049489 | 0.29693384 | 35.79196139 | -81.65904847 |
| Wilson County, Springhill township | 0.09090909 | 0.29666487 | 35.63492788 | -78.04366715 |
| Franklin County, Franklinton township | 0.18558365 | 0.29612695 | 36.0921446 | -78.36172449 |
| Carteret County, White Oak township | 0.17294244 | 0.29558903 | 34.71286904 | -76.97097394 |
| Nash County, Nashville township | 0.12278107 | 0.29558903 | 35.96807329 | -77.99233445 |
| Wilson County, Gardners township | 0.11807423 | 0.2950511 | 35.74119814 | -77.84788841 |
| Person County, Mount Tirzah township | 0.22512103 | 0.29478214 | 36.3309786 | -78.90550874 |
| Onslow County, Hofmann Forest UT | 0.18692846 | 0.29451318 | 34.82269264 | -77.36508789 |
| Pender County, Rocky Point township | 0.11592254 | 0.29424422 | 34.44841198 | -77.87539982 |
| Johnston County, Elevation township | 0.21315223 | 0.29424422 | 35.43776227 | -78.54280315 |
| Tyrrell County, Columbia township | 0.22834857 | 0.29424422 | 35.91367354 | -76.23762005 |
| Franklin County, Hayesville township | 0.25524476 | 0.29370629 | 36.24571072 | -78.32545931 |
| Harnett County, Lillington township | 0.21288327 | 0.29343733 | 35.38634315 | -78.80870951 |
| Burke County, Lovelady township | 0.25712749 | 0.29316837 | 35.73095666 | -81.52959538 |
| Clay County, Sweetwater township | 0.27407208 | 0.29289941 | 35.04817138 | -83.86390187 |
| Johnston County, Wilders township | 0.12439484 | 0.29289941 | 35.75064347 | -78.35669245 |
| Montgomery County, Troy township | 0.23574502 | 0.29263045 | 35.3401091 | -79.91793098 |
| Hertford County, Ahoskie township | 0.15909091 | 0.29263045 | 36.2897826 | -76.9983955 |
| Wayne County, Fork township | 0.1218397 | 0.2915546 | 35.43331057 | -78.06333867 |
| Watauga County, Laurel Creek township | 0.22081764 | 0.29101668 | 36.20386157 | -81.89247813 |
| Stanly County, Endy township | 0.20602474 | 0.29020979 | 35.31501714 | -80.23264598 |
| Washington County, Lees Mill township | 0.21005917 | 0.29020979 | 35.87277791 | -76.67647677 |
| Alexander County, Ellendale township | 0.17616998 | 0.28913394 | 35.92081519 | -81.23851933 |
| Richmond County, Rockingham township | 0.19903174 | 0.28886498 | 34.93175074 | -79.75213409 |
| Sampson County, North Clinton township | 0.12641205 | 0.2880581 | 34.99734449 | -78.33492227 |
| Macon County, Smithbridge township | 0.22942442 | 0.28778913 | 35.09041446 | -83.41099093 |
| Robeson County, Burnt Swamp township | 0.18854223 | 0.28698225 | 34.6847022 | -79.13436299 |
| Bertie County, Indian Woods township | 0.1958042 | 0.28698225 | 35.92642668 | -77.00503918 |
| Edgecombe County, Township 5, Lower Fishing Creek | 0.0683163 | 0.28671329 | 35.94379229 | -77.54240744 |
| Bladen County, Central township | 0.21785906 | 0.28644432 | 34.64848472 | -78.58342127 |
| Mitchell County, Red Hill township | 0.28402367 | 0.28536848 | 36.0189023 | -82.23838318 |
| Craven County, Township 3 | 0.14954276 | 0.28509952 | 35.22568954 | -77.463526 |
| Onslow County, Swansboro township | 0.24179666 | 0.28483055 | 34.70879582 | -77.22460571 |
| Lenoir County, Pink Hill township | 0.24475524 | 0.28483055 | 35.05422328 | -77.67442466 |
| Person County, Bushy Fork township | 0.22781065 | 0.28348575 | 36.32889939 | -79.04879547 |
| Mecklenburg County, Township 5, Providence | 0.26815492 | 0.28052717 | 35.04603418 | -80.76446477 |
| Lenoir County, Contentnea Neck township | 0.16648736 | 0.27998924 | 35.32632174 | -77.52365497 |
| Dare County, Kinnakeet township | 0.22646584 | 0.27998924 | 35.64649176 | -75.48177027 |
| Clay County, Brasstown township | 0.25201721 | 0.27918236 | 35.0538497 | -83.91872861 |
| Harnett County, Grove township | 0.23157612 | 0.27837547 | 35.48004659 | -78.67435167 |
| Lincoln County, Howards Creek township | 0.11135019 | 0.27783755 | 35.47612094 | -81.29705198 |
| Bladen County, Hollow township | 0.16204949 | 0.27729962 | 34.7208161 | -78.76279906 |
| Durham County, Mangum township | 0.18410436 | 0.2767617 | 36.19921123 | -78.93562684 |
| Cherokee County, Notla township | 0.19607316 | 0.27595481 | 35.02918495 | -84.11758093 |
| Union County, Sandy Ridge township | 0.19661108 | 0.27595481 | 34.99119648 | -80.69149621 |
| Bladen County, Lake Creek township | 0.21355568 | 0.27595481 | 34.66787231 | -78.41115912 |
| Swain County, Forneys Creek UT | 0.18370091 | 0.27568585 | 35.56547671 | -83.64087047 |
| Watauga County, Cove Creek township | 0.19768693 | 0.27541689 | 36.27889022 | -81.76475116 |
| Wilson County, Stantonsburg township | 0.08337816 | 0.27461001 | 35.67542578 | -77.86375821 |
| Jones County, Township 6, Chinquapin | 0.23050027 | 0.27461001 | 35.14234303 | -77.51212363 |
| Harnett County, Anderson Creek township | 0.16756321 | 0.27407208 | 35.24851981 | -78.95263159 |
| Tyrrell County, Alligator township | 0.23130715 | 0.27353416 | 35.84222687 | -76.08520981 |
| Cabarrus County, Township 8, Mount Pleasant | 0.22216245 | 0.2732652 | 35.41043977 | -80.4379767 |
| Avery County, Elk Park township | 0.25484131 | 0.27272727 | 36.14749813 | -81.98810698 |
| Wilson County, Cross Roads township | 0.18934911 | 0.27218935 | 35.65266267 | -78.0110753 |
| Montgomery County, Ophir township | 0.22337278 | 0.27218935 | 35.50549768 | -79.96736866 |
| Carteret County, Atlantic township | 0.25954814 | 0.27218935 | 34.90250822 | -76.33195278 |
| Richmond County, Mineral Springs township | 0.17711135 | 0.2711135 | 35.09347942 | -79.77369958 |
| Macon County, Cartoogechaye township | 0.16769769 | 0.2711135 | 35.1627453 | -83.45489407 |
| Bladen County, Turnbull township | 0.21113502 | 0.27084454 | 34.75836648 | -78.55833304 |
| Avery County, Heaton township | 0.26667563 | 0.27057558 | 36.16300474 | -81.92850868 |
| Clay County, Shooting Creek township | 0.22565896 | 0.27003765 | 35.02230221 | -83.61728256 |
| Robeson County, St. Pauls township | 0.15155998 | 0.27003765 | 34.82282534 | -78.99181907 |
| Pamlico County, Township 5 | 0.13959118 | 0.26923077 | 35.01445795 | -76.72612265 |
| Beaufort County, Chocowinity township | 0.1520979 | 0.26842388 | 35.51056202 | -77.10992687 |
| Avery County, Beech Mountain township | 0.21315223 | 0.26815492 | 36.2071598 | -81.9202294 |
| Pender County, Burgaw township | 0.095078 | 0.26788596 | 34.54891412 | -77.93546 |
| Franklin County, Gold Mine township | 0.1958042 | 0.26681011 | 36.21666243 | -78.13240299 |
| Carteret County, Davis township | 0.16083916 | 0.26627219 | 35.03932177 | -76.4658228 |
| Hertford County, Winton township | 0.1908284 | 0.26627219 | 36.40359008 | -76.99802949 |
| Bladen County, White Oak township | 0.18665949 | 0.26600323 | 34.74682958 | -78.78491011 |
| Carteret County, Sea Level township | 0.23265196 | 0.26600323 | 34.9600007 | -76.40165494 |
| Duplin County, Warsaw township | 0.10395374 | 0.26573427 | 35.00382156 | -78.08041092 |
| Union County, Lanes Creek township | 0.08821947 | 0.2654653 | 34.8191109 | -80.41883967 |
| Halifax County, Littleton township | 0.11377084 | 0.2654653 | 36.43261484 | -77.90865216 |
| Martin County, Jamesville township | 0.1570737 | 0.26465842 | 35.8688897 | -76.79155549 |
| Perquimans County, New Hope township | 0.13824637 | 0.26304465 | 36.22065874 | -76.29959687 |
| Franklin County, Cedar Rock township | 0.19042496 | 0.26277569 | 36.08924589 | -78.19134424 |
| Warren County, River township | 0.21005917 | 0.26277569 | 36.50679844 | -77.94798546 |
| Watauga County, Elk township | 0.16581495 | 0.26250672 | 36.20452488 | -81.5234232 |
| Warren County, Nutbush township | 0.21140398 | 0.26035503 | 36.4431138 | -78.27861549 |
| Camden County, Courthouse township | 0.2288865 | 0.26008607 | 36.34249283 | -76.20986649 |
| Harnett County, Upper Little River township | 0.13932222 | 0.25901022 | 35.41233085 | -78.83102861 |
| Bladen County, Abbotts township | 0.17079075 | 0.2584723 | 34.51083367 | -78.72637211 |
| Martin County, Williams township | 0.1880043 | 0.25820334 | 35.81701104 | -77.03275246 |
| Edgecombe County, Township 4, Deep Creek | 0.13340506 | 0.25820334 | 35.93948016 | -77.51497227 |
| Robeson County, Smyrna township | 0.09991931 | 0.25793437 | 34.58087632 | -79.04560922 |
| Sampson County, Belvoir township | 0.15814954 | 0.25793437 | 34.98774184 | -78.42144276 |
| Watauga County, Shawneehaw township | 0.22969338 | 0.25793437 | 36.19724576 | -81.83698286 |
| Nash County, Bailey township | 0.11659494 | 0.25793437 | 35.78327329 | -78.12184858 |
| Currituck County, Moyock township | 0.19983862 | 0.25739645 | 36.51717358 | -76.16326599 |
| Granville County, Oak Hill township | 0.22350726 | 0.25685853 | 36.45651948 | -78.67636112 |
| Ashe County, Elk township | 0.18504572 | 0.25605164 | 36.27747921 | -81.56615915 |
| Granville County, Sassafras Fork township | 0.21893491 | 0.25551372 | 36.53773683 | -78.4568022 |
| Chatham County, Haw River township | 0.22592792 | 0.25551372 | 35.67985111 | -79.10990415 |
| Perquimans County, Parkville township | 0.13690156 | 0.25551372 | 36.19984969 | -76.44842865 |
| Burke County, Upper Creek township | 0.20965573 | 0.25524476 | 35.81198594 | -81.82462911 |
| Vance County, Williamsboro township | 0.21315223 | 0.25524476 | 36.39380356 | -78.4048275 |
| Ashe County, Pond Mountain township | 0.22700377 | 0.25416891 | 36.5557112 | -81.62643941 |
| Vance County, Townsville township | 0.22619688 | 0.25389995 | 36.53363726 | -78.42909942 |
| Franklin County, Sandy Creek township | 0.19526627 | 0.25363098 | 36.21550901 | -78.27549043 |
| Montgomery County, Rocky Springs township | 0.17200108 | 0.25336202 | 35.26959688 | -79.73602427 |
| Northampton County, Pleasant Hill township | 0.17805272 | 0.25336202 | 36.47337917 | -77.58318459 |
| Moore County, Township 1, Carthage | 0.18908015 | 0.25309306 | 35.29336512 | -79.42134023 |
| Hoke County, Fort Bragg Military Reservation township | 0.08257127 | 0.25228618 | 35.04847935 | -79.13790671 |
| Warren County, Smith Creek township | 0.22243141 | 0.25174825 | 36.50061022 | -78.20758625 |
| Perquimans County, Bethel township | 0.09359871 | 0.25174825 | 36.17799843 | -76.45415868 |
| Harnett County, Barbecue township | 0.14550834 | 0.25147929 | 35.33827791 | -79.10155876 |
| Caldwell County, Mulberry township | 0.12076385 | 0.25147929 | 35.94310452 | -81.60197859 |
| Duplin County, Rose Hill township | 0.19607316 | 0.25121033 | 34.80723747 | -77.98641561 |
| Robeson County, West Howellsville township | 0.08203335 | 0.25094137 | 34.65692642 | -78.94491124 |
| Lee County, Township 1, Greenwood | 0.17267348 | 0.25094137 | 35.40193253 | -79.22836922 |
| Dare County, Hatteras township | 0.22740721 | 0.2506724 | 35.2660387 | -75.52856384 |
| Washington County, Skinnersville township | 0.18639053 | 0.2506724 | 35.89387639 | -76.52834882 |
| Person County, Woodsdale township | 0.17616998 | 0.25013448 | 36.45273796 | -78.96341724 |
| Richmond County, Beaverdam township | 0.12614309 | 0.2493276 | 34.99687616 | -79.57004676 |
| Perquimans County, Hertford township | 0.11955352 | 0.2493276 | 36.18255402 | -76.48161822 |
| Harnett County, Stewarts Creek township | 0.2084454 | 0.24905863 | 35.31640706 | -78.79639751 |
| Washington County, Scuppernong township | 0.19607316 | 0.24825175 | 35.7927914 | -76.46909127 |
| Scotland County, Williamson township | 0.11377084 | 0.2471759 | 34.81308025 | -79.55368157 |
| Union County, Jackson township | 0.15747714 | 0.24690694 | 34.82187363 | -80.78124991 |
| Beaufort County, Richland township | 0.15088757 | 0.24690694 | 35.23328101 | -76.67032173 |
| Montgomery County, Pee Dee township | 0.19445939 | 0.24663798 | 35.27074295 | -80.07320697 |
| Johnston County, O'Neals township | 0.07907477 | 0.24663798 | 35.80407315 | -78.25894181 |
| Greene County, Hookerton township | 0.18827327 | 0.24610005 | 35.36974235 | -77.65444178 |
| Hoke County, Quewhiffle township | 0.09601937 | 0.24556213 | 35.10571803 | -79.37803131 |
| Watauga County, Meat Camp township | 0.20333513 | 0.24502421 | 36.26215475 | -81.62592177 |
| Swain County, Nantahala township | 0.14389457 | 0.24448628 | 35.37085301 | -83.50455184 |
| Harnett County, Johnsonville township | 0.13703604 | 0.24448628 | 35.36421127 | -79.12396321 |
| Hoke County, Stonewall township | 0.18370091 | 0.24341044 | 34.95656164 | -79.13022513 |
| Lee County, Township 3, Cape Fear | 0.15801506 | 0.24314147 | 35.44423447 | -79.04964878 |
| Tyrrell County, South Fork township | 0.2169177 | 0.24314147 | 35.74807697 | -76.33826492 |
| Burke County, Smoky Creek township | 0.20898332 | 0.24287251 | 35.7886015 | -81.6314602 |
| Pamlico County, Township 3 | 0.15519096 | 0.24206563 | 35.12838202 | -76.72535857 |
| Clay County, Hiawassee township | 0.2027972 | 0.2415277 | 35.0396059 | -83.78167701 |
| Granville County, Salem township | 0.21718666 | 0.2415277 | 36.37445125 | -78.5810306 |
| Robeson County, Philadelphus township | 0.0957504 | 0.24125874 | 34.71064629 | -79.15658873 |
| Caldwell County, Johns River township | 0.16487359 | 0.24098978 | 35.9023279 | -81.6380416 |
| Cumberland County, Grays Creek township | 0.13622916 | 0.24045186 | 34.97469973 | -78.78948465 |
| Robeson County, Saddletree township | 0.14967725 | 0.24018289 | 34.6908641 | -79.02120094 |
| Jones County, Township 2, Pollocksville | 0.10166756 | 0.23964497 | 35.06815802 | -77.19200153 |
| Warren County, Sandy Creek township | 0.20817644 | 0.23910705 | 36.32931555 | -78.2770506 |
| Hyde County, Currituck township | 0.18222162 | 0.23883808 | 35.40290752 | -76.45669211 |
| Currituck County, Fruitville township | 0.20467994 | 0.23883808 | 36.47061023 | -75.88779745 |
| Wake County, New Light township | 0.12130178 | 0.23856912 | 35.96717633 | -78.59029166 |
| Edgecombe County, Township 8, Sparta | 0.18370091 | 0.23830016 | 35.79473297 | -77.60749436 |
| Bladen County, Whites Creek township | 0.20064551 | 0.2380312 | 34.54423825 | -78.49545011 |
| Sampson County, Mingo township | 0.15653577 | 0.23722431 | 35.21813726 | -78.59339692 |
| Wayne County, Great Swamp township | 0.11296396 | 0.23695535 | 35.53458966 | -77.98261023 |
| Caswell County, Leasburg township | 0.16594944 | 0.23668639 | 36.4043233 | -79.26030761 |
| Edgecombe County, Township 11, Walnut Creek | 0.15115654 | 0.23641743 | 35.93478082 | -77.63010408 |
| Bertie County, Snakebite township | 0.14039806 | 0.23534158 | 36.02281066 | -77.03767703 |
| Bladen County, Bethel township | 0.12600861 | 0.23507262 | 34.69479772 | -78.74070292 |
| Duplin County, Albertson township | 0.11780527 | 0.23507262 | 35.08897249 | -77.75054129 |
| Sampson County, Franklin township | 0.1542496 | 0.23426573 | 34.64307934 | -78.24902734 |
| Columbus County, Chadbourn township | 0.11995697 | 0.23426573 | 34.32096443 | -78.82476463 |
| Pitt County, Falkland township | 0.11538462 | 0.23399677 | 35.61117833 | -77.45373183 |
| Robeson County, Maxton township | 0.08284024 | 0.23372781 | 34.70826492 | -79.29685817 |
| Duplin County, Wolfscrape township | 0.12237762 | 0.23265196 | 35.17938401 | -78.03889739 |
| Pitt County, Bethel township | 0.10153308 | 0.232383 | 35.82587843 | -77.37220221 |
| Macon County, Sugarfork township | 0.2084454 | 0.23130715 | 35.12284472 | -83.29399961 |
| Avery County, Carey's Flat township | 0.18181818 | 0.23103819 | 36.08355522 | -81.83007713 |
| Martin County, Goose Nest township | 0.16729424 | 0.23103819 | 35.97480483 | -77.30662643 |
| Warren County, Shocco township | 0.18128026 | 0.23076923 | 36.26468867 | -78.14949212 |
| Clay County, Tusquittee township | 0.18235611 | 0.23023131 | 35.06176457 | -83.77813298 |
| Columbus County, Bogue township | 0.15626681 | 0.23023131 | 34.33258526 | -78.599437 |
| Pitt County, Carolina township | 0.09427111 | 0.22996235 | 35.68577012 | -77.3500993 |
| Nash County, Dry Wells township | 0.15277031 | 0.2288865 | 35.79992587 | -78.23151716 |
| Franklin County, Dunn township | 0.14523938 | 0.22861754 | 35.86989109 | -78.24338742 |
| Sampson County, McDaniels township | 0.17980097 | 0.22807961 | 34.8380797 | -78.48401677 |
| Sampson County, Honeycutt township | 0.15935987 | 0.22727273 | 34.96988349 | -78.45365776 |
| Granville County, Walnut Grove township | 0.1936525 | 0.2267348 | 36.3207784 | -78.67929116 |
| Halifax County, Scotland Neck township | 0.19069392 | 0.22646584 | 36.19362363 | -77.39484894 |
| Halifax County, Faucett township | 0.06737493 | 0.22646584 | 36.37671982 | -77.69305417 |
| Bladen County, Carvers Creek township | 0.14981173 | 0.22592792 | 34.43025153 | -78.49360467 |
| Camden County, Shiloh township | 0.17442173 | 0.22565896 | 36.28031872 | -76.11155904 |
| Sampson County, South Clinton township | 0.15034965 | 0.22512103 | 34.97951005 | -78.36716138 |
| Chatham County, Cape Fear township | 0.13125336 | 0.22485207 | 35.64203409 | -79.00517615 |
| Wayne County, Buck Swamp township | 0.11753631 | 0.22485207 | 35.49072248 | -77.99308712 |
| Bertie County, Whites township | 0.14107047 | 0.22350726 | 36.03935473 | -76.71916332 |
| Columbus County, Welches Creek township | 0.14470145 | 0.2232383 | 34.36676085 | -78.67525608 |
| Hertford County, Murfreesboro township | 0.18504572 | 0.22296934 | 36.42127111 | -77.10834528 |
| Craven County, Township 2 | 0.10839161 | 0.22216245 | 35.10738265 | -77.01298033 |
| Ashe County, Laurel township | 0.17764927 | 0.22189349 | 36.51158642 | -81.63492927 |
| Duplin County, Cypress Creek township | 0.15734266 | 0.22162453 | 34.83062404 | -77.70019009 |
| Robeson County, Back Swamp township | 0.14792899 | 0.22135557 | 34.61079882 | -79.09482028 |
| Stanly County, Tyson township | 0.11955352 | 0.22054868 | 35.29298608 | -80.23722483 |
| Warren County, Sixpound township | 0.19903174 | 0.22027972 | 36.5194112 | -78.03099384 |
| Pasquotank County, Salem township | 0.18343195 | 0.21947283 | 36.21481555 | -76.12904952 |
| Bladen County, Frenches Creek township | 0.14873588 | 0.21920387 | 34.54978276 | -78.38244563 |
| Wayne County, Indian Springs township | 0.08821947 | 0.21920387 | 35.22326766 | -78.02848929 |
| Franklin County, Harris township | 0.16810113 | 0.21893491 | 35.98243765 | -78.38753826 |
| Greene County, Ormonds township | 0.15034965 | 0.21839699 | 35.46644024 | -77.54597225 |
| Chowan County, Township 4, Yeopim | 0.12143626 | 0.21678322 | 36.03864433 | -76.57640574 |
| Robeson County, Britts township | 0.09010221 | 0.21651425 | 34.57292981 | -78.99154805 |
| Macon County, Cowee township | 0.13676708 | 0.21624529 | 35.26164025 | -83.32676611 |
| Robeson County, East Howellsville township | 0.08956428 | 0.21597633 | 34.65293333 | -78.91785785 |
| Gates County, Mintonsville township | 0.18235611 | 0.21543841 | 36.34542445 | -76.63986281 |
| Pitt County, Fountain township | 0.15949435 | 0.21463152 | 35.63704792 | -77.61765936 |
| Bladen County, Brown Marsh township | 0.17267348 | 0.21301775 | 34.51660197 | -78.61343311 |
| Duplin County, Kenansville township | 0.09037117 | 0.21301775 | 34.96085845 | -77.95000675 |
| Hoke County, Blue Springs township | 0.10180204 | 0.21274879 | 34.95029583 | -79.24376904 |
| Vance County, Watkins township | 0.20091447 | 0.2119419 | 36.29230307 | -78.48590741 |
| Person County, Holloway township | 0.16917698 | 0.21167294 | 36.44476601 | -78.90802272 |
| Johnston County, Meadow township | 0.14766003 | 0.21140398 | 35.29374695 | -78.49141374 |
| Columbus County, Waccamaw township | 0.18114578 | 0.21113502 | 34.38219405 | -78.47670679 |
| Wilkes County, Elk township | 0.17549758 | 0.2105971 | 36.07721024 | -81.40623936 |
| Franklin County, Cypress Creek township | 0.11672942 | 0.2105971 | 35.95348482 | -78.19512811 |
| Pitt County, Grimesland township | 0.14107047 | 0.2105971 | 35.58508152 | -77.28991603 |
| Moore County, Township 5, Deep River | 0.15317375 | 0.20979021 | 35.50943146 | -79.34570126 |
| Sampson County, South River township | 0.16837009 | 0.20952125 | 34.80790792 | -78.43489877 |
| Caldwell County, North Catawba township | 0.1732114 | 0.20952125 | 35.82262416 | -81.54025201 |
| Watauga County, Bald Mountain township | 0.17939753 | 0.20898332 | 36.25878829 | -81.59816329 |
| Currituck County, Crawford township | 0.18813878 | 0.2084454 | 36.43778171 | -76.09815883 |
| Hoke County, Antioch township | 0.1309844 | 0.20817644 | 34.94637313 | -79.21659713 |
| Rutherford County, Golden Valley township | 0.15667025 | 0.20817644 | 35.56244312 | -81.8150836 |
| Gates County, Hunters Mill township | 0.17899408 | 0.20817644 | 36.40647347 | -76.59523484 |
| Warren County, Judkins township | 0.17455621 | 0.20736955 | 36.42753606 | -78.02451134 |
| Beaufort County, Pantego township | 0.14981173 | 0.20683163 | 35.54403965 | -76.6189009 |
| Columbus County, Bolton township | 0.15586337 | 0.20656267 | 34.24620199 | -78.47990798 |
| Gates County, Holly Grove township | 0.15895643 | 0.20602474 | 36.45470743 | -76.61138509 |
| Richmond County, Black Jack township | 0.15088757 | 0.20575578 | 35.03121461 | -79.81501249 |
| Montgomery County, Uwharrie township | 0.1344809 | 0.20575578 | 35.41367727 | -79.95864442 |
| Halifax County, Halifax township | 0.10852609 | 0.20414201 | 36.3505383 | -77.6708432 |
| Martin County, Hamilton township | 0.11242604 | 0.20387305 | 35.90897325 | -77.03798175 |
| Craven County, Township 1 | 0.11135019 | 0.20333513 | 35.31349015 | -77.15921257 |
| Alexander County, Sugar Loaf township | 0.16541151 | 0.20333513 | 36.02763354 | -81.18920922 |
| Pitt County, Grifton township | 0.11511565 | 0.20306616 | 35.42724021 | -77.44232581 |
| Beaufort County, Bath township | 0.14981173 | 0.20306616 | 35.46614592 | -76.83752216 |
| Jones County, Township 1, White Oak | 0.14429801 | 0.20225928 | 34.89288171 | -77.23538485 |
| Edgecombe County, Township 9, Otter Creek | 0.15599785 | 0.20172136 | 35.75519777 | -77.64559622 |
| Carteret County, Merrimon township | 0.09991931 | 0.20118343 | 34.97825303 | -76.50975974 |
| Edgecombe County, Township 2, Lower Conetoe | 0.1570737 | 0.20118343 | 35.84322212 | -77.48176509 |
| Anson County, Ansonville township | 0.11242604 | 0.20091447 | 35.18635524 | -80.11897842 |
| Caldwell County, Yadkin Valley township | 0.13501883 | 0.20091447 | 35.99916642 | -81.50632145 |
| Graham County, Yellow Creek township | 0.17307692 | 0.20064551 | 35.43621208 | -83.91396548 |
| Cumberland County, Beaver Dam township | 0.12990855 | 0.20064551 | 34.8641425 | -78.50606725 |
| Macon County, Nantahala township | 0.16191501 | 0.20037655 | 35.21110421 | -83.69833646 |
| Pitt County, Chicod township | 0.08472297 | 0.19956966 | 35.54564163 | -77.32807758 |
| Jones County, Township 4, Cypress Creek | 0.17455621 | 0.19903174 | 35.06360606 | -77.44679301 |
| Anson County, Burnsville township | 0.12560516 | 0.19849381 | 35.09467803 | -80.27831622 |
| Sampson County, Newton Grove township | 0.17105971 | 0.19849381 | 35.22517204 | -78.3381743 |
| Northampton County, Rich Square township | 0.12896719 | 0.19849381 | 36.20681925 | -77.33434309 |
| Halifax County, Palmyra township | 0.15317375 | 0.19768693 | 36.15431316 | -77.29028499 |
| Greene County, Snow Hill township | 0.15962883 | 0.19741797 | 35.46170944 | -77.66038901 |
| Caldwell County, Wilson Creek township | 0.18289403 | 0.19741797 | 35.96320607 | -81.76787412 |
| Macon County, Flats township | 0.1753631 | 0.19688004 | 35.01509749 | -83.33978966 |
| Nash County, Mannings township | 0.08902636 | 0.19634212 | 35.94098112 | -78.11271171 |
| Catawba County, Caldwell township | 0.11780527 | 0.19607316 | 35.57608347 | -81.19312537 |
| Montgomery County, Cheek Creek township | 0.17509414 | 0.19553523 | 35.24827133 | -79.90928484 |
| Perquimans County, Belvidere township | 0.13259817 | 0.19526627 | 36.35370324 | -76.55157588 |
| Rutherford County, Duncans Creek township | 0.17751479 | 0.19499731 | 35.41661345 | -81.73001789 |
| Duplin County, Faison township | 0.06804734 | 0.19472835 | 35.16163408 | -78.07128167 |
| Caldwell County, Globe township | 0.1218397 | 0.19445939 | 36.09898361 | -81.77049691 |
| Anson County, Gulledge township | 0.11686391 | 0.19392146 | 34.81522731 | -80.05646067 |
| Moore County, Township 6, Greenwood | 0.10933297 | 0.19284562 | 35.33596329 | -79.24298007 |
| Cherokee County, Shoal Creek township | 0.11000538 | 0.19257665 | 35.05968658 | -84.19640817 |
| Pender County, Holly township | 0.08700914 | 0.19203873 | 34.6727878 | -77.71003868 |
| Caldwell County, Kings Creek township | 0.16850457 | 0.19203873 | 35.96689946 | -81.42763186 |
| Edgecombe County, Township 3, Upper Conetoe | 0.16783217 | 0.19176977 | 35.9308369 | -77.4601113 |
| Watauga County, Beaverdam township | 0.14416353 | 0.19150081 | 36.27008788 | -81.88001727 |
| Columbus County, Lees township | 0.11767079 | 0.19096288 | 34.16235192 | -78.52680033 |
| Union County, Buford township | 0.07046799 | 0.19096288 | 34.82275548 | -80.44602568 |
| Greene County, Jason township | 0.14900484 | 0.19069392 | 35.40875829 | -77.75811866 |
| Nash County, Ferrells township | 0.12641205 | 0.19042496 | 35.90545453 | -78.17808016 |
| Wayne County, Nahunta township | 0.05473373 | 0.19042496 | 35.53037767 | -77.95529759 |
| Warren County, Fishing Creek township | 0.16783217 | 0.18988704 | 36.3440389 | -78.07326272 |
| Halifax County, Conoconnara township | 0.13743948 | 0.18934911 | 36.21551873 | -77.38938993 |
| Nash County, Red Oak township | 0.0866057 | 0.18881119 | 35.99927279 | -77.89934672 |
| Robeson County, Lumber Bridge township | 0.11525013 | 0.18773534 | 34.94469586 | -79.04872858 |
| Nash County, North Whitakers township | 0.05150619 | 0.18692846 | 36.07423072 | -77.79562287 |
| Wayne County, Saulston township | 0.07235073 | 0.18639053 | 35.40805358 | -77.89969837 |
| Ashe County, North Fork township | 0.11592254 | 0.18531469 | 36.41130545 | -81.73955839 |
| Richmond County, Steeles township | 0.14712211 | 0.18531469 | 35.16018847 | -79.92793996 |
| Anson County, Morven township | 0.10112964 | 0.18531469 | 34.81149235 | -80.02929598 |
| Wayne County, Grantham township | 0.09090909 | 0.18504572 | 35.30056006 | -78.23593021 |
| Beaufort County, Long Acre township | 0.13313609 | 0.18504572 | 35.54108245 | -77.01714026 |
| Brunswick County, Waccamaw township | 0.14618074 | 0.18504572 | 34.15825978 | -78.49994072 |
| Carteret County, Stacy township | 0.14766003 | 0.18477676 | 34.89438221 | -76.41859814 |
| Bladen County, Bladenboro township | 0.12641205 | 0.18370091 | 34.54491945 | -78.80242176 |
| Person County, Cunningham township | 0.14281872 | 0.18316299 | 36.44663983 | -79.07922834 |
| Jones County, Township 3, Trenton | 0.10274341 | 0.18289403 | 34.94533522 | -77.41941291 |
| Robeson County, Thompson township | 0.09332975 | 0.18262507 | 34.55662219 | -79.19048008 |
| Catawba County, Bandy's township | 0.09198494 | 0.18262507 | 35.56627521 | -81.47663129 |
| Nash County, Jackson township | 0.07557827 | 0.18235611 | 35.84581034 | -77.93635535 |
| Union County, New Salem township | 0.09870898 | 0.18208714 | 35.09835416 | -80.30559311 |
| Moore County, Township 10, Little River | 0.10691232 | 0.18181818 | 35.21029886 | -79.15820898 |
| Alexander County, Gwaltneys township | 0.14429801 | 0.18128026 | 36.01022745 | -81.05097025 |
| Anson County, White Store township | 0.11013986 | 0.1810113 | 34.8191533 | -80.2240582 |
| Nash County, Coopers township | 0.09279182 | 0.18047337 | 35.88543585 | -77.89835955 |
| Northampton County, Roanoke township | 0.13461538 | 0.18020441 | 36.33816353 | -77.30143048 |
| Alexander County, Millers township | 0.12116729 | 0.17859064 | 35.86283971 | -81.13677648 |
| Harnett County, Buckhorn township | 0.14147391 | 0.17697687 | 35.55217819 | -78.85576963 |
| Gates County, Gatesville township | 0.10879505 | 0.17697687 | 36.46373831 | -76.6665281 |
| Chowan County, Township 3, Upper | 0.15061861 | 0.17643895 | 36.30170726 | -76.65123082 |
| Chowan County, Township 2, Middle | 0.14201183 | 0.17374933 | 36.14417116 | -76.66345557 |
| Wayne County, Pikeville township | 0.09709521 | 0.17348037 | 35.48651275 | -77.96578959 |
| Hertford County, St. Johns township | 0.11296396 | 0.1732114 | 36.3379653 | -77.01477833 |
| Northampton County, Kirby township | 0.15182894 | 0.1732114 | 36.45195459 | -77.1579792 |
| Duplin County, Glisson township | 0.1070468 | 0.17294244 | 35.13638211 | -77.90821818 |
| Bertie County, Roxobel township | 0.14362561 | 0.17294244 | 36.17620405 | -77.28479855 |
| Nash County, Griffins township | 0.07692308 | 0.17267348 | 36.15607511 | -78.03269804 |
| Alexander County, Little River township | 0.11874664 | 0.17132867 | 35.96146977 | -81.20223313 |
| Greene County, Speights Bridge township | 0.11027434 | 0.17132867 | 35.60178525 | -77.68299715 |
| Nash County, Castalia township | 0.13353954 | 0.17052179 | 36.04225235 | -78.0315085 |
| Northampton County, Wiccacanee township | 0.12789134 | 0.16998386 | 36.54387816 | -77.30712366 |
| Macon County, Burningtown township | 0.08512641 | 0.1697149 | 35.22918162 | -83.44395542 |
| Bertie County, Colerain township | 0.08862292 | 0.16863905 | 36.08810366 | -76.87816516 |
| Carteret County, Smyrna township | 0.12412587 | 0.16810113 | 34.77224166 | -76.50631206 |
| Craven County, Township 5 | 0.11578806 | 0.16783217 | 34.89658112 | -76.84050674 |
| Burke County, Upper Fork township | 0.14080151 | 0.16783217 | 35.63742582 | -81.68835584 |
| Columbus County, Fair Bluff township | 0.11403981 | 0.16729424 | 34.32296434 | -78.99127877 |
| Pender County, Grady township | 0.07611619 | 0.16729424 | 34.35552224 | -78.00893204 |
| Pender County, Canetuck township | 0.09440559 | 0.16729424 | 34.43292171 | -78.21402986 |
| Bertie County, Woodville township | 0.13824637 | 0.1662184 | 35.93969666 | -77.08725206 |
| Robeson County, Fairmont township | 0.09655729 | 0.16541151 | 34.5007859 | -79.1190987 |
| Halifax County, Enfield township | 0.09413663 | 0.16541151 | 36.26718155 | -77.71993557 |
| Hertford County, Maneys Neck township | 0.11135019 | 0.16433566 | 36.45619528 | -77.04202952 |
| Sampson County, Westbrook township | 0.15532544 | 0.1640667 | 35.21145801 | -78.39772852 |
| Alexander County, Sharpes township | 0.12829478 | 0.16352878 | 35.90346503 | -81.10046354 |
| Martin County, Robersonville township | 0.08741259 | 0.16272189 | 35.81716863 | -77.31743956 |
| Columbus County, Tatums township | 0.07907477 | 0.16218397 | 34.4029906 | -78.91782589 |
| Anson County, Lanesboro township | 0.11417429 | 0.16218397 | 34.99546955 | -80.21478723 |
| Columbus County, Bug Hill township | 0.11457773 | 0.16191501 | 33.97257183 | -78.62523902 |
| Columbus County, Ransom township | 0.09763314 | 0.16137708 | 34.37116923 | -78.25623832 |
| Robeson County, Parkton township | 0.10610543 | 0.16083916 | 34.94072799 | -79.02156866 |
| Ashe County, Creston township | 0.11175363 | 0.15949435 | 36.39116992 | -81.57268475 |
| Pamlico County, Township 1 | 0.10543303 | 0.15949435 | 35.09807918 | -76.81776206 |
| Carteret County, Straits township | 0.08741259 | 0.15922539 | 34.75490835 | -76.53887404 |
| Greene County, Bull Head township | 0.10381926 | 0.15922539 | 35.52264044 | -77.75889042 |
| Columbus County, South Williams township | 0.12493276 | 0.1584185 | 34.1409513 | -78.83717228 |
| Graham County, Stecoah township | 0.14308768 | 0.15761162 | 35.38541497 | -83.64214512 |
| Craven County, Township 9 | 0.08068854 | 0.15734266 | 35.13826798 | -77.20280949 |
| Sampson County, Piney Grove township | 0.06414739 | 0.15572889 | 35.19495084 | -78.28886581 |
| Carteret County, Harkers Island township | 0.12264658 | 0.15357719 | 34.68096294 | -76.64203777 |
| Gates County, Hall township | 0.10798817 | 0.15303927 | 36.41213265 | -76.90971424 |
| Halifax County, Roseneath township | 0.09682625 | 0.15223238 | 36.1190212 | -77.49913258 |
| Cherokee County, Hothouse township | 0.12842926 | 0.14954276 | 35.01810707 | -84.23066577 |
| Burke County, Lower Fork township | 0.11430877 | 0.14792899 | 35.67149638 | -81.59735411 |
| Halifax County, Brinkleyville township | 0.07100592 | 0.14766003 | 36.3011142 | -77.94054891 |
| Greene County, Shine township | 0.08902636 | 0.14523938 | 35.45260768 | -77.74751812 |
| Hertford County, Harrellsville township | 0.09359871 | 0.14523938 | 36.28885876 | -76.71193064 |
| Sampson County, Dismal township | 0.08176439 | 0.14443249 | 35.15629931 | -78.63573176 |
| Columbus County, Williams township | 0.11592254 | 0.14389457 | 34.18093127 | -78.80047144 |
| Columbus County, Western Prong township | 0.13044648 | 0.14389457 | 34.43677856 | -78.68730771 |
| Carteret County, Harlowe township | 0.07611619 | 0.14389457 | 34.755666 | -76.67925435 |
| Edgecombe County, Township 13, Cokey | 0.08230231 | 0.14389457 | 35.82522267 | -77.65691989 |
| Cherokee County, Beaverdam township | 0.09467456 | 0.14308768 | 35.26740865 | -84.02456266 |
| Carteret County, Marshallberg township | 0.14201183 | 0.14201183 | 34.72847812 | -76.5175239 |
| Sampson County, Taylors Bridge township | 0.06777838 | 0.14147391 | 34.79680807 | -78.21321939 |
| Pender County, Columbia township | 0.07073696 | 0.14066703 | 34.5385812 | -78.16151352 |
| Sampson County, Lisbon township | 0.08714363 | 0.13340506 | 34.76533018 | -78.30462894 |
| Pitt County, Swift Creek township | 0.08243679 | 0.13340506 | 35.37921964 | -77.28434059 |
| Robeson County, Alfordsville township | 0.06562668 | 0.13340506 | 34.68625783 | -79.30166795 |
| Hoke County, Allendale township | 0.10287789 | 0.13152232 | 34.88429587 | -79.25828099 |
| Robeson County, Red Springs township | 0.1105433 | 0.13044648 | 34.82852085 | -79.18654705 |
| Watauga County, North Fork township | 0.11511565 | 0.12883271 | 36.36383097 | -81.72016326 |
| Robeson County, Shannon township | 0.10301237 | 0.12721893 | 34.84658238 | -79.15456196 |
| Pender County, Caswell township | 0.10408822 | 0.12614309 | 34.54274973 | -78.18848707 |
| Johnston County, Bentonville township | 0.07665412 | 0.12560516 | 35.291033 | -78.32280567 |
| Gates County, Haslett township | 0.08122647 | 0.12399139 | 36.50745258 | -76.65514452 |
| Martin County, Beargrass township | 0.10166756 | 0.12237762 | 35.77324038 | -77.04381545 |
| Scotland County, Spring Hill township | 0.07934373 | 0.119688 | 34.81576911 | -79.41322957 |
| Robeson County, Orrum township | 0.07100592 | 0.11941904 | 34.46293372 | -79.01596127 |
| Columbus County, Cerro Gordo township | 0.07544379 | 0.11915008 | 34.31898047 | -78.96433942 |
| Robeson County, Smiths township | 0.07383002 | 0.11780527 | 34.72244016 | -79.23785461 |
| Martin County, Cross Roads township | 0.09494352 | 0.11780527 | 35.83466772 | -77.14222461 |
| Sampson County, Turkey township | 0.04612695 | 0.11753631 | 35.086379 | -78.17350236 |
| Martin County, Griffins township | 0.07867133 | 0.11699839 | 35.70282148 | -76.89093413 |
| Martin County, Poplar Point township | 0.08445401 | 0.11646046 | 35.84345814 | -77.19697972 |
| Halifax County, Butterwood township | 0.08633674 | 0.11592254 | 36.36264416 | -77.89700379 |
| Wilson County, Saratoga township | 0.06791286 | 0.11511565 | 35.67181529 | -77.6943324 |
| Edgecombe County, Township 6, Upper Fishing Creek | 0.04142012 | 0.11323292 | 36.08336315 | -77.70778451 |
| Pender County, Long Creek township | 0.07073696 | 0.11215707 | 34.48305357 | -77.95101084 |
| Duplin County, Magnolia township | 0.05446477 | 0.10812265 | 34.82403086 | -78.09466544 |
| Bertie County, Mitchell township | 0.07557827 | 0.1070468 | 36.23715585 | -76.95452275 |
| Sampson County, Herring township | 0.0683163 | 0.10435718 | 35.17165043 | -78.43513162 |
| Northampton County, Seaboard township | 0.0837816 | 0.10193653 | 36.47803103 | -77.46722672 |
| Gates County, Reynoldson township | 0.06750941 | 0.10059172 | 36.48219348 | -76.92045242 |
| Robeson County, Sterlings township | 0.05823023 | 0.09521248 | 34.44886639 | -79.07480766 |
| Sampson County, Halls township | 0.04760624 | 0.09010221 | 35.15930498 | -78.35351652 |
| Robeson County, Rennert township | 0.07423346 | 0.07772996 | 34.83868854 | -79.10030319 |
| Northampton County, Jackson township | 0.0648198 | 0.07638515 | 36.36422879 | -77.46688521 |
| Robeson County, Gaddys township | 0.03227542 | 0.0739645 | 34.47251844 | -79.23678995 |
| Robeson County, Whitehouse township | 0.04908553 | 0.06051641 | 34.33883723 | -79.0990638 |
