## Supplementary material for "Mapping long-distance jump dispersal risk by leveraging transport infrastructure data and sampling bias correction: a case study with the invasive spotted lanternfly (*Lycorma delicatula*)": S11 Ohio risk

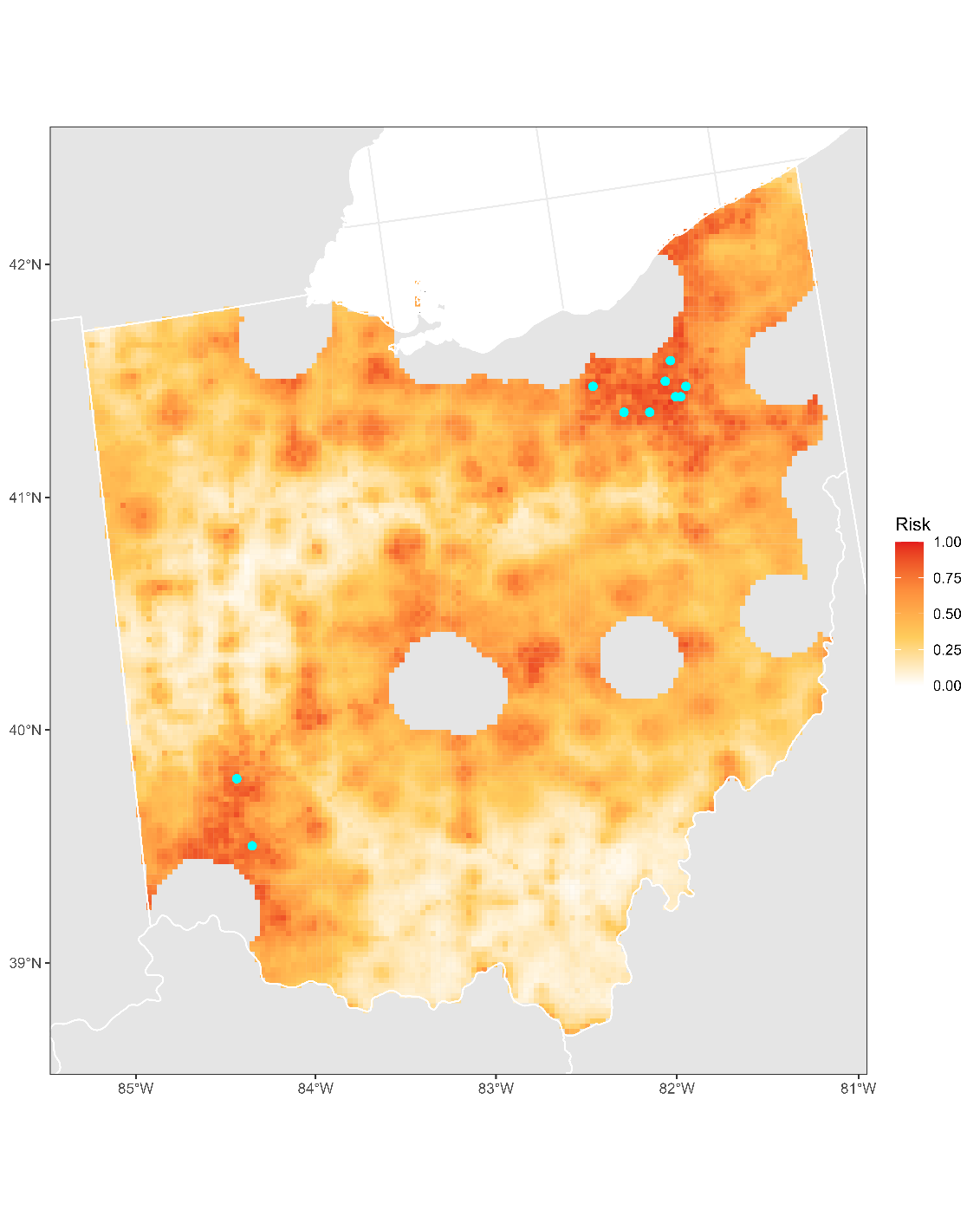

**Figure S11.1:** Projected risk of colonization by *L. delicatula* jump dispersal for Ohio. Risk projections are based on an ensemble random forest model including distances from existing *L. delicatula* populations. Note that areas falling less than 20 km from existing *L. delicatula* populations are not included in the map because any colonization there would be categorized as diffusive spread based on our methodology (regardless of the actual source of the resulting population). Darker, redder colors indicate increased risk of colonization by jump dispersal. Cyan points show the top ten highest risk locations in the state.

**Table S11.1:** Ranked list of Ohio county subdivisions sorted by the maximum projected risk of colonization by jump dispersal occurring there. For each subdivision, we identified the 2.5 x 2.5 km grid cell centroid included there with the highest projected risk of colonization by jump dispersal and report that location and risk value.

| **County, subdivision** | **Median risk value** | **Maximum risk value** | **Latitude of maximum risk** | **Longitude of maximum risk** |
| --- | --- | --- | --- | --- |
| Summit County, Akron city | 0.85287789 | 0.96395912 | 41.06944009 | -81.42973677 |
| Medina County, York township | 0.80782679 | 0.93249059 | 41.1672858 | -81.89542833 |
| Medina County, Wadsworth city | 0.82060247 | 0.93087682 | 41.03696461 | -81.73971339 |
| Warren County, Union township | 0.79424422 | 0.9281872 | 39.39958977 | -84.1936293 |
| Portage County, Brimfield township | 0.76909629 | 0.92711135 | 41.10673463 | -81.36126508 |
| Summit County, Mogadore village | 0.90182894 | 0.92307692 | 41.06602679 | -81.40010035 |
| Summit County, Barberton city | 0.83068854 | 0.91850457 | 41.02018975 | -81.59155331 |
| Summit County, Cuyahoga Falls city | 0.81091985 | 0.91662184 | 41.14245203 | -81.47531967 |
| Summit County, Hudson city | 0.80823023 | 0.91420118 | 41.22732282 | -81.42730958 |
| Montgomery County, Moraine city | 0.78375471 | 0.91070468 | 39.69564598 | -84.2338141 |
| Medina County, Medina township | 0.83647122 | 0.90720818 | 41.18606374 | -81.86127745 |
| Portage County, Aurora city | 0.84077461 | 0.90317375 | 41.33082418 | -81.3448259 |
| Summit County, Fairlawn city | 0.84938139 | 0.90182894 | 41.11521474 | -81.63275327 |
| Medina County, Brunswick Hills township | 0.76855836 | 0.89483593 | 41.22359317 | -81.7929212 |
| Summit County, Copley township | 0.84776762 | 0.8929532 | 41.11857416 | -81.66242144 |
| Summit County, Tallmadge city | 0.84346423 | 0.8929532 | 41.0949138 | -81.4548037 |
| Warren County, Clear Creek township | 0.61148467 | 0.88891877 | 39.54067865 | -84.25919056 |
| Summit County, Green city | 0.80849919 | 0.88811189 | 40.959113 | -81.45260085 |
| Geauga County, Bainbridge township | 0.79989242 | 0.88569123 | 41.35289242 | -81.34020095 |
| Clermont County, Goshen township | 0.7027972 | 0.8838085 | 39.21974916 | -84.19360084 |
| Summit County, Springfield township | 0.82544379 | 0.88300161 | 41.00664618 | -81.473076 |
| Montgomery County, Washington township | 0.78267886 | 0.88246369 | 39.64307864 | -84.15355971 |
| Medina County, Granger township | 0.7704411 | 0.88219473 | 41.14734991 | -81.71727243 |
| Mahoning County, Beaver township | 0.79115116 | 0.88058096 | 40.95283753 | -80.63539461 |
| Stark County, Alliance city | 0.74852071 | 0.87950511 | 40.92125135 | -81.1272532 |
| Licking County, Union township | 0.68020441 | 0.87842926 | 40.01348393 | -82.48206443 |
| Guernsey County, Cambridge township | 0.70871436 | 0.87762238 | 40.00376185 | -81.58791524 |
| Summit County, Lakemore village | 0.84938139 | 0.87735342 | 41.02530854 | -81.43889095 |
| Summit County, Norton city | 0.80688542 | 0.87735342 | 41.04563059 | -81.61666099 |
| Licking County, Heath city | 0.85153308 | 0.87573964 | 39.99139703 | -82.48625209 |
| Clermont County, Williamsburg township | 0.71785906 | 0.87547068 | 39.07306589 | -84.07057862 |
| Licking County, Licking township | 0.69230769 | 0.87439484 | 39.98823154 | -82.45701215 |
| Medina County, Brunswick city | 0.83001614 | 0.86982249 | 41.22691568 | -81.82264249 |
| Lake County, Mentor city | 0.82759548 | 0.86955352 | 41.69426844 | -81.3601597 |
| Stark County, Canton city | 0.81038193 | 0.86793975 | 40.75032488 | -81.40505381 |
| Butler County, Fairfield township | 0.84211942 | 0.86713287 | 39.40744149 | -84.51708751 |
| Medina County, Montville township | 0.7972028 | 0.86713287 | 41.11318734 | -81.81526826 |
| Warren County, Turtlecreek township | 0.70952125 | 0.86605702 | 39.43546539 | -84.33542925 |
| Butler County, West Chester township | 0.86417429 | 0.86605702 | 39.346908 | -84.34976702 |
| Summit County, Bath township | 0.75739645 | 0.86417429 | 41.14064951 | -81.65790954 |
| Portage County, Ravenna township | 0.72189349 | 0.86229155 | 41.16605622 | -81.28809217 |
| Licking County, Newark city | 0.80231307 | 0.85825713 | 40.0765757 | -82.44021076 |
| Montgomery County, West Carrollton city | 0.82517483 | 0.85745024 | 39.65136924 | -84.24107565 |
| Clermont County, Stonelick township | 0.72377622 | 0.85691232 | 39.17271532 | -84.17185789 |
| Lake County, Madison township | 0.77084454 | 0.85610543 | 41.77347023 | -81.06752979 |
| Stark County, Nimishillen township | 0.67267348 | 0.85449166 | 40.82827635 | -81.29815762 |
| Richland County, Mansfield city | 0.5683163 | 0.8542227 | 40.78358392 | -82.51549934 |
| Butler County, Lemon township | 0.8394298 | 0.85341581 | 39.46577041 | -84.41915096 |
| Hamilton County, Whitewater township | 0.7943787 | 0.85260893 | 39.20984925 | -84.81338336 |
| Summit County, Munroe Falls city | 0.84359871 | 0.85207101 | 41.13904909 | -81.44565044 |
| Butler County, Middletown city | 0.74462076 | 0.8507262 | 39.48519653 | -84.38647032 |
| Montgomery County, Miami township | 0.78671329 | 0.84884346 | 39.62647511 | -84.21553768 |
| Summit County, Stow city | 0.79128564 | 0.84749866 | 41.16452119 | -81.4707478 |
| Wayne County, Chippewa township | 0.5887574 | 0.84642281 | 40.98277458 | -81.65981985 |
| Lake County, Painesville city | 0.84521248 | 0.84642281 | 41.72464544 | -81.2312248 |
| Medina County, Sharon township | 0.71705218 | 0.84588488 | 41.13194297 | -81.78112223 |
| Sandusky County, Sandusky township | 0.79639591 | 0.845078 | 41.38839388 | -83.12900164 |
| Lake County, Concord township | 0.79747176 | 0.84454008 | 41.66188005 | -81.27514622 |
| Portage County, Tallmadge city | 0.8415815 | 0.8415815 | 41.11015707 | -81.39091669 |
| Lake County, Perry township | 0.75309306 | 0.84023669 | 41.73978586 | -81.16671552 |
| Warren County, Deerfield township | 0.75443787 | 0.83889188 | 39.36083101 | -84.25899917 |
| Butler County, St. Clair township | 0.69150081 | 0.83808499 | 39.4322722 | -84.54265179 |
| Warren County, Lebanon city | 0.7950511 | 0.83754707 | 39.40235055 | -84.22270242 |
| Cuyahoga County, Strongsville city | 0.79854761 | 0.83754707 | 41.27771092 | -81.87318958 |
| Sandusky County, Fremont city | 0.78711673 | 0.83727811 | 41.36626424 | -83.13306799 |
| Mahoning County, Springfield township | 0.71274879 | 0.8359333 | 40.97487227 | -80.63057354 |
| Butler County, Hamilton city | 0.78550296 | 0.83431953 | 39.38798168 | -84.54969794 |
| Columbiana County, Perry township | 0.77850995 | 0.83324368 | 40.88611237 | -80.83178711 |
| Summit County, Coventry township | 0.80486821 | 0.83162991 | 41.01681416 | -81.56192973 |
| Portage County, Kent city | 0.7471759 | 0.83136095 | 41.12879901 | -81.35665992 |
| Wood County, Bowling Green city | 0.82221625 | 0.83109198 | 41.36931957 | -83.6186668 |
| Portage County, Streetsboro city | 0.75779989 | 0.83082302 | 41.26462118 | -81.3586834 |
| Lake County, Painesville township | 0.77783755 | 0.83082302 | 41.74671832 | -81.2265381 |
| Clark County, Springfield city | 0.73372781 | 0.83055406 | 39.89406687 | -83.72536117 |
| Medina County, Wadsworth township | 0.8006993 | 0.83055406 | 41.05569883 | -81.70558414 |
| Butler County, Liberty township | 0.78725121 | 0.82974718 | 39.37990757 | -84.46246544 |
| Sandusky County, Ballville township | 0.60126412 | 0.82974718 | 41.3380938 | -83.0774746 |
| Summit County, Boston township | 0.58902636 | 0.82894029 | 41.21205975 | -81.49129472 |
| Stark County, Lexington township | 0.74946208 | 0.82840237 | 40.92472758 | -81.15681585 |
| Portage County, Franklin township | 0.7119419 | 0.82813341 | 41.16949582 | -81.31776512 |
| Clermont County, Miami township | 0.82033351 | 0.82625067 | 39.26953534 | -84.24439267 |
| Warren County, Franklin township | 0.76250672 | 0.82625067 | 39.54616312 | -84.3174576 |
| Marion County, Marion township | 0.71100054 | 0.82625067 | 40.57993686 | -83.15573132 |
| Licking County, Granville township | 0.73480366 | 0.82544379 | 40.08290797 | -82.4987638 |
| Warren County, Salem township | 0.73924153 | 0.82383002 | 39.36913432 | -84.11007888 |
| Summit County, Macedonia city | 0.82383002 | 0.82383002 | 41.27827387 | -81.47757198 |
| Warren County, Harlan township | 0.65465304 | 0.82221625 | 39.32208419 | -84.08834534 |
| Stark County, Plain township | 0.73924153 | 0.82140936 | 40.86062528 | -81.38218309 |
| Medina County, Westfield Center village | 0.81979559 | 0.81979559 | 41.03478465 | -81.92202325 |
| Summit County, Twinsburg township | 0.81845078 | 0.81845078 | 41.29353097 | -81.41352381 |
| Butler County, Trenton city | 0.74031737 | 0.81710597 | 39.4906216 | -84.44470221 |
| Clermont County, Batavia township | 0.71516945 | 0.81414739 | 39.07863765 | -84.12845644 |
| Columbiana County, Center township | 0.6936525 | 0.81414739 | 40.78728941 | -80.76230422 |
| Wood County, Center township | 0.6936525 | 0.81360947 | 41.34717509 | -83.62257829 |
| Warren County, Mason city | 0.75309306 | 0.81360947 | 39.34417685 | -84.32071134 |
| Medina County, Liverpool township | 0.72364174 | 0.81334051 | 41.23354002 | -81.88209343 |
| Lake County, Mentor-on-the-Lake city | 0.81334051 | 0.81334051 | 41.71634528 | -81.35551561 |
| Ashtabula County, Austinburg township | 0.6570737 | 0.8119957 | 41.77442841 | -80.88328048 |
| Hamilton County, Harrison township | 0.73103819 | 0.81172673 | 39.23200083 | -84.80995055 |
| Medina County, Lafayette township | 0.74448628 | 0.81145777 | 41.10103404 | -81.90873825 |
| Columbiana County, Elkrun township | 0.58714363 | 0.81118881 | 40.78372204 | -80.73281835 |
| Warren County, Hamilton township | 0.73991393 | 0.81065089 | 39.35807871 | -84.22994134 |
| Stark County, Lake township | 0.70602474 | 0.81038193 | 40.97435347 | -81.38883582 |
| Geauga County, Newbury township | 0.73830016 | 0.80957504 | 41.48186622 | -81.28258443 |
| Mahoning County, Smith township | 0.61457773 | 0.80930608 | 40.91427832 | -81.0681366 |
| Stark County, Canton township | 0.76882733 | 0.80903712 | 40.76896528 | -81.37097425 |
| Stark County, Washington township | 0.73856912 | 0.80742335 | 40.85509515 | -81.14123362 |
| Clark County, Springfield township | 0.72619688 | 0.80688542 | 39.87767373 | -83.78765389 |
| Butler County, Madison township | 0.66460463 | 0.80580958 | 39.44633586 | -84.45181403 |
| Medina County, Harrisville township | 0.66863905 | 0.80580958 | 40.9971924 | -81.99012878 |
| Stark County, Marlboro township | 0.64335664 | 0.80554061 | 40.95371542 | -81.21131083 |
| Medina County, Hinckley township | 0.69701452 | 0.80231307 | 41.24234465 | -81.75871583 |
| Stark County, Jackson township | 0.70387305 | 0.80150619 | 40.88951519 | -81.43672073 |
| Medina County, Westfield township | 0.70320065 | 0.80123722 | 41.03807663 | -81.95167129 |
| Union County, Jerome township | 0.64470145 | 0.80016138 | 40.1125479 | -83.21013881 |
| Ashtabula County, Saybrook township | 0.66352878 | 0.79881657 | 41.7929459 | -80.84856429 |
| Licking County, Madison township | 0.75094137 | 0.79881657 | 40.04494317 | -82.35663134 |
| Montgomery County, Jefferson township | 0.65438408 | 0.79854761 | 39.70114255 | -84.29220451 |
| Hancock County, Findlay city | 0.68854223 | 0.79800968 | 41.08146698 | -83.66932516 |
| Summit County, Richfield township | 0.73991393 | 0.79800968 | 41.21023417 | -81.67406412 |
| Geauga County, Burton township | 0.7232383 | 0.79693384 | 41.46106988 | -81.10381063 |
| Columbiana County, Knox township | 0.59763314 | 0.79639591 | 40.89222833 | -81.07281756 |
| Ashtabula County, Geneva township | 0.69688004 | 0.79532006 | 41.80355643 | -80.93832315 |
| Montgomery County, Dayton city | 0.65868747 | 0.7894029 | 39.71778447 | -84.23017998 |
| Medina County, Medina City township | 0.77891339 | 0.78913394 | 41.13858607 | -81.84048942 |
| Hancock County, Liberty township | 0.62022593 | 0.78886498 | 41.03718663 | -83.67708219 |
| Lake County, Willoughby city | 0.78859602 | 0.78859602 | 41.69769304 | -81.39005961 |
| Ashtabula County, Harpersfield township | 0.62641205 | 0.78617536 | 41.78501455 | -80.97302086 |
| Columbiana County, Fairfield township | 0.64631522 | 0.78563744 | 40.88673455 | -80.64983985 |
| Stark County, Perry township | 0.63636364 | 0.78536848 | 40.775797 | -81.42999929 |
| Clark County, Harmony township | 0.46073158 | 0.78375471 | 39.91331409 | -83.69229905 |
| Hancock County, Allen township | 0.69553523 | 0.78375471 | 41.12574852 | -83.66155841 |
| Portage County, Randolph township | 0.70790748 | 0.78375471 | 41.04885722 | -81.25196128 |
| Delaware County, Delaware City township | 0.71167294 | 0.78187197 | 40.30251964 | -83.08611284 |
| Portage County, Ravenna city | 0.73910705 | 0.78160301 | 41.15915639 | -81.22875495 |
| Clermont County, Jackson township | 0.67482517 | 0.78106509 | 39.0979858 | -84.09587239 |
| Medina County, Guilford township | 0.69055944 | 0.78079613 | 41.02154793 | -81.80345876 |
| Tuscarawas County, Goshen township | 0.66944594 | 0.77998924 | 40.46016156 | -81.43480032 |
| Summit County, Twinsburg city | 0.77568585 | 0.77998924 | 41.31560091 | -81.4089228 |
| Portage County, Suffield township | 0.68625605 | 0.77810651 | 41.04054297 | -81.37506334 |
| Marion County, Claridon township | 0.57867133 | 0.77649274 | 40.59299689 | -83.06316539 |
| Columbiana County, Butler township | 0.5859333 | 0.77649274 | 40.89672593 | -80.92039615 |
| Geauga County, Munson township | 0.70817644 | 0.77514793 | 41.54462251 | -81.23878209 |
| Portage County, Rootstown township | 0.69069392 | 0.7732652 | 41.07091656 | -81.2473258 |
| Licking County, Franklin township | 0.62963959 | 0.77272727 | 40.02286012 | -82.36085944 |
| Montgomery County, German township | 0.64147391 | 0.77165143 | 39.61804493 | -84.36497683 |
| Montgomery County, Kettering city | 0.65008069 | 0.77138246 | 39.66243636 | -84.12072728 |
| Delaware County, Concord township | 0.68181818 | 0.77057558 | 40.19498628 | -83.13552862 |
| Geauga County, Auburn township | 0.67643895 | 0.76976869 | 41.41566047 | -81.29653016 |
| Tuscarawas County, New Philadelphia city | 0.65572889 | 0.76949973 | 40.48221791 | -81.43025964 |
| Clinton County, Wilmington city | 0.67455621 | 0.76869285 | 39.45157887 | -83.80088821 |
| Stark County, Osnaburg township | 0.56414739 | 0.76815492 | 40.80621943 | -81.30276147 |
| Licking County, Newark township | 0.7415277 | 0.767617 | 40.07974524 | -82.46948599 |
| Guernsey County, Jackson township | 0.7175901 | 0.76707907 | 39.98170578 | -81.59237722 |
| Clark County, Moorefield township | 0.654922 | 0.76600323 | 40.01043725 | -83.76493455 |
| Summit County, New Franklin city | 0.69338354 | 0.76519634 | 40.96929487 | -81.54139204 |
| Lake County, Kirtland Hills village | 0.76331361 | 0.76331361 | 41.64325046 | -81.30968477 |
| Stark County, Louisville city | 0.76331361 | 0.76331361 | 40.84688916 | -81.26401257 |
| Geauga County, Middlefield township | 0.6584185 | 0.76250672 | 41.43551891 | -81.07874054 |
| Wayne County, Canaan township | 0.54330285 | 0.76062399 | 40.97510811 | -81.99452802 |
| Seneca County, Tiffin city | 0.75147929 | 0.76035503 | 41.12285952 | -83.17762991 |
| Marion County, Pleasant township | 0.57315761 | 0.76035503 | 40.55481098 | -83.13024156 |
| Seneca County, Hopewell township | 0.65384615 | 0.75954814 | 41.12585967 | -83.20737069 |
| Licking County, Harrison township | 0.68773534 | 0.75927918 | 40.00712262 | -82.63249033 |
| Lake County, Leroy township | 0.63515331 | 0.75927918 | 41.69564514 | -81.17612267 |
| Geauga County, Claridon township | 0.69903174 | 0.75874126 | 41.53769724 | -81.17913021 |
| Butler County, Hanover township | 0.6732114 | 0.75766541 | 39.40130301 | -84.69512845 |
| Montgomery County, Trotwood city | 0.55607854 | 0.75766541 | 39.7481629 | -84.3141856 |
| Delaware County, Brown township | 0.57208176 | 0.75685853 | 40.29645973 | -83.02732889 |
| Sandusky County, Rice township | 0.5295858 | 0.75685853 | 41.40750372 | -83.09507527 |
| Champaign County, Urbana township | 0.62520172 | 0.75658956 | 40.03542709 | -83.79045256 |
| Summit County, Boston Heights village | 0.74569661 | 0.75658956 | 41.23752453 | -81.5164345 |
| Geauga County, Chardon city | 0.74179666 | 0.75658956 | 41.58529933 | -81.19958884 |
| Geauga County, Chardon township | 0.72108661 | 0.75470683 | 41.61428625 | -81.2546141 |
| Clinton County, Union township | 0.59077461 | 0.75443787 | 39.43515109 | -83.86278269 |
| Delaware County, Delaware township | 0.73614847 | 0.75336202 | 40.25829991 | -83.09414613 |
| Geauga County, Middlefield village | 0.72404519 | 0.75094137 | 41.45757966 | -81.07402529 |
| Muskingum County, Hopewell township | 0.65613233 | 0.75013448 | 39.95296314 | -82.13554431 |
| Guernsey County, Center township | 0.63972566 | 0.74986552 | 39.9936191 | -81.50031146 |
| Stark County, Lawrence township | 0.66164605 | 0.74959656 | 40.90309134 | -81.55499542 |
| Clinton County, Liberty township | 0.47767617 | 0.7493276 | 39.52365366 | -83.84783056 |
| Knox County, Mount Vernon city | 0.70118343 | 0.74825175 | 40.37639737 | -82.50289681 |
| Ashtabula County, Plymouth township | 0.55540613 | 0.74744486 | 41.85201266 | -80.77424062 |
| Hancock County, Marion township | 0.65882195 | 0.74287251 | 41.07276888 | -83.58010831 |
| Medina County, Rittman city | 0.74098978 | 0.74098978 | 40.99946968 | -81.80791771 |
| Seneca County, Clinton township | 0.63609467 | 0.73991393 | 41.09772759 | -83.15193666 |
| Ashland County, Montgomery township | 0.55500269 | 0.73937601 | 40.89707704 | -82.31241825 |
| Stark County, Massillon city | 0.57450242 | 0.73830016 | 40.78599324 | -81.51855808 |
| Ashland County, Ashland city | 0.72485207 | 0.7380312 | 40.87498352 | -82.31671092 |
| Ross County, Scioto township | 0.4880312 | 0.73776224 | 39.32172823 | -82.96663047 |
| Portage County, Freedom township | 0.52071006 | 0.73776224 | 41.24393549 | -81.18043723 |
| Portage County, Shalersville township | 0.59709521 | 0.73776224 | 41.20673347 | -81.24913736 |
| Trumbull County, Bazetta township | 0.69889726 | 0.73722431 | 41.31205607 | -80.80033578 |
| Greene County, Sugarcreek township | 0.58848843 | 0.73668639 | 39.65965157 | -84.09155322 |
| Medina County, Chatham township | 0.55190963 | 0.73561054 | 41.06344678 | -81.9769146 |
| Fairfield County, Lancaster City township | 0.71140398 | 0.73507262 | 39.71369153 | -82.62785075 |
| Fairfield County, Pleasant township | 0.61525013 | 0.73480366 | 39.75473524 | -82.59044108 |
| Geauga County, Troy township | 0.66863905 | 0.73480366 | 41.41694634 | -81.11321997 |
| Fairfield County, Walnut township | 0.44553523 | 0.732383 | 39.92513688 | -82.4987996 |
| Wood County, Plain township | 0.6753631 | 0.73184508 | 41.35007218 | -83.65243231 |
| Ashland County, Milton township | 0.54115116 | 0.72996235 | 40.88137631 | -82.37591217 |
| Delaware County, Scioto township | 0.62076385 | 0.72915546 | 40.2643286 | -83.15290435 |
| Pickaway County, Harrison township | 0.6218397 | 0.7288865 | 39.74769877 | -82.94848454 |
| Clermont County, Pierce township | 0.70091447 | 0.72781065 | 39.01778265 | -84.1971839 |
| Delaware County, Liberty township | 0.61915008 | 0.72727273 | 40.19197143 | -83.10617746 |
| Ashtabula County, Ashtabula township | 0.55675094 | 0.72646584 | 41.85557391 | -80.80417869 |
| Mahoning County, Goshen township | 0.64766003 | 0.72538999 | 40.91877036 | -80.9156675 |
| Van Wert County, Pleasant township | 0.62587413 | 0.72377622 | 40.87418395 | -84.57965918 |
| Pickaway County, Circleville city | 0.69123185 | 0.72350726 | 39.61205024 | -82.94351411 |
| Mercer County, Jefferson township | 0.48251748 | 0.7232383 | 40.555899 | -84.54095361 |
| Butler County, Reily township | 0.57208176 | 0.72243141 | 39.40394695 | -84.72422096 |
| Mahoning County, Green township | 0.62990855 | 0.72189349 | 40.9337415 | -80.85182445 |
| Madison County, Deer Creek township | 0.52931684 | 0.72081764 | 39.9505052 | -83.38810697 |
| Brown County, Green township | 0.47875202 | 0.72081764 | 39.03406434 | -83.90070113 |
| Greene County, Bath township | 0.46288327 | 0.72027972 | 39.79245701 | -84.06946334 |
| Wayne County, Rittman city | 0.72027972 | 0.72027972 | 40.97406749 | -81.78275775 |
| Van Wert County, Tully township | 0.43544917 | 0.72001076 | 40.92916825 | -84.69134068 |
| Fairfield County, Berne township | 0.52877891 | 0.71893491 | 39.70112215 | -82.5113322 |
| Wood County, Liberty township | 0.54263045 | 0.7175901 | 41.32792708 | -83.65633213 |
| Ottawa County, Salem township | 0.53617536 | 0.7175901 | 41.52419721 | -83.13445592 |
| Fulton County, Pike township | 0.46812803 | 0.7175901 | 41.589344 | -84.03660141 |
| Seneca County, Fostoria city | 0.61430877 | 0.71732114 | 41.16584834 | -83.38189378 |
| Brown County, Pike township | 0.59144701 | 0.71651425 | 39.0119397 | -83.90440133 |
| Geauga County, Parkman township | 0.60233997 | 0.71651425 | 41.41345845 | -81.08345282 |
| Delaware County, Troy township | 0.64254976 | 0.71597633 | 40.34674 | -83.07806961 |
| Montgomery County, Harrison township | 0.52770307 | 0.71436256 | 39.80357646 | -84.18638507 |
| Seneca County, Liberty township | 0.51371705 | 0.71355568 | 41.17011427 | -83.19930964 |
| Hancock County, Portage township | 0.48843464 | 0.71328671 | 41.0843525 | -83.69906897 |
| Butler County, Ross township | 0.64779451 | 0.71247983 | 39.37650377 | -84.66952283 |
| Fairfield County, Hocking township | 0.60408822 | 0.71167294 | 39.71681695 | -82.65698673 |
| Knox County, Clinton township | 0.70346961 | 0.71113502 | 40.37955285 | -82.53229774 |
| Delaware County, Berkshire township | 0.63770845 | 0.71032813 | 40.23698442 | -82.88857081 |
| Trumbull County, Champion township | 0.6316568 | 0.70925229 | 41.29710579 | -80.8645322 |
| Washington County, Marietta township | 0.41877353 | 0.70898332 | 39.40335768 | -81.47129519 |
| Sandusky County, Green Creek township | 0.60233997 | 0.70790748 | 41.33506282 | -83.04764985 |
| Tuscarawas County, Dover township | 0.46584185 | 0.70736955 | 40.4890208 | -81.48905299 |
| Sandusky County, Washington township | 0.50941366 | 0.70521786 | 41.35014833 | -83.19679919 |
| Columbiana County, Salem township | 0.65398063 | 0.70467994 | 40.83492583 | -80.78226704 |
| Wood County, Portage township | 0.48682087 | 0.70441097 | 41.32503096 | -83.62648732 |
| Pickaway County, Circleville township | 0.69176977 | 0.70360409 | 39.58689421 | -82.91844001 |
| Carroll County, Center township | 0.63084992 | 0.70306616 | 40.57658257 | -81.07920931 |
| Huron County, Townsend township | 0.61726735 | 0.70225928 | 41.25721298 | -82.51550223 |
| Fairfield County, Greenfield township | 0.63676708 | 0.70091447 | 39.74202649 | -82.68201466 |
| Wood County, Henry township | 0.48897257 | 0.70037655 | 41.17003127 | -83.65378193 |
| Madison County, Union township | 0.52770307 | 0.70010758 | 39.91509519 | -83.4836617 |
| Wayne County, Green township | 0.5324099 | 0.70010758 | 40.83828641 | -81.77993125 |
| Ashland County, Orange township | 0.63703604 | 0.69956966 | 40.96335905 | -82.29952416 |
| Allen County, Lima city | 0.55177515 | 0.69956966 | 40.74188385 | -84.11979402 |
| Fulton County, Swan Creek township | 0.55433029 | 0.6993007 | 41.57806924 | -83.91673059 |
| Tuscarawas County, Dover city | 0.65357719 | 0.6993007 | 40.5110795 | -81.48452726 |
| Sandusky County, Riley township | 0.53388919 | 0.69876278 | 41.35718909 | -83.04355848 |
| Portage County, Mantua township | 0.61027434 | 0.69849381 | 41.27981859 | -81.294622 |
| Trumbull County, Southington township | 0.51748252 | 0.69634212 | 41.28212354 | -80.92870139 |
| Licking County, Etna township | 0.63394298 | 0.69607316 | 39.95980955 | -82.61153465 |
| Delaware County, Berlin township | 0.6155191 | 0.6958042 | 40.26521359 | -82.94323415 |
| Columbiana County, Unity township | 0.58431953 | 0.6958042 | 40.88314083 | -80.62031832 |
| Coshocton County, Tuscarawas township | 0.59682625 | 0.69553523 | 40.23904427 | -81.86992909 |
| Defiance County, Richland township | 0.54747176 | 0.69472835 | 41.30122256 | -84.32829043 |
| Wayne County, Congress township | 0.61215707 | 0.69419042 | 40.9849124 | -82.08341547 |
| Sandusky County, Clyde city | 0.68141474 | 0.69419042 | 41.30685616 | -82.9921152 |
| Licking County, Newton township | 0.56508876 | 0.69338354 | 40.12074841 | -82.43179452 |
| Ashland County, Clear Creek township | 0.59601937 | 0.69150081 | 40.90347194 | -82.37163755 |
| Delaware County, Sunbury Village township | 0.67697687 | 0.690156 | 40.23391204 | -82.85921293 |
| Hancock County, Cass township | 0.49677246 | 0.68934911 | 41.09781386 | -83.60594436 |
| Washington County, Belpre city | 0.61202259 | 0.68881119 | 39.28121801 | -81.58474768 |
| Sandusky County, Bellevue city | 0.68881119 | 0.68881119 | 41.26941462 | -82.84725215 |
| Seneca County, Eden township | 0.45884884 | 0.68827327 | 41.07860809 | -83.18569913 |
| Fulton County, Fulton township | 0.49273803 | 0.68746638 | 41.60022763 | -83.912899 |
| Portage County, Atwater township | 0.50995159 | 0.68719742 | 41.04194133 | -81.19272583 |
| Stark County, Pike township | 0.5295858 | 0.68692846 | 40.72484835 | -81.38012622 |
| Coshocton County, Jackson township | 0.57100592 | 0.68504572 | 40.26771296 | -81.92418896 |
| Allen County, Bath township | 0.29343733 | 0.68504572 | 40.76124433 | -84.08644442 |
| Delaware County, Radnor township | 0.58714363 | 0.68396988 | 40.30855222 | -83.14490674 |
| Butler County, Morgan township | 0.57235073 | 0.68370091 | 39.38179676 | -84.72768817 |
| Huron County, Lyme township | 0.46261431 | 0.68370091 | 41.26633025 | -82.8174664 |
| Geauga County, Huntsburg township | 0.59037117 | 0.68262507 | 41.50519464 | -81.09438949 |
| Defiance County, Noble township | 0.55755783 | 0.68235611 | 41.28451456 | -84.39168458 |
| Guernsey County, Valley township | 0.53509952 | 0.68208714 | 39.93759385 | -81.60129298 |
| Greene County, Beavercreek township | 0.57369554 | 0.67993545 | 39.72605392 | -84.08051846 |
| Warren County, Washington township | 0.59722969 | 0.67966649 | 39.40781508 | -84.04462185 |
| Pickaway County, Walnut township | 0.4936794 | 0.67912856 | 39.72254246 | -82.92336149 |
| Madison County, Somerford township | 0.42388381 | 0.67912856 | 39.93721328 | -83.47978538 |
| Lorain County, Huntington township | 0.59252286 | 0.67912856 | 41.13690284 | -82.23531641 |
| Richland County, Jackson township | 0.48951049 | 0.6788596 | 40.88769873 | -82.64663955 |
| Madison County, London city | 0.67105971 | 0.67859064 | 39.88710543 | -83.42905317 |
| Geauga County, Hambden township | 0.64497041 | 0.67670791 | 41.57835557 | -81.13990631 |
| Union County, Paris township | 0.58741259 | 0.67455621 | 40.26013335 | -83.33317973 |
| Huron County, Norwalk township | 0.62708445 | 0.67401829 | 41.23825951 | -82.54951265 |
| Wayne County, Milton township | 0.56764389 | 0.67374933 | 40.97739167 | -81.81237388 |
| Pickaway County, Washington township | 0.52299623 | 0.67348037 | 39.60899155 | -82.91440815 |
| Muskingum County, Falls township | 0.61511565 | 0.67348037 | 39.97178986 | -82.10203363 |
| Lorain County, Penfield township | 0.54733728 | 0.6732114 | 41.17463036 | -82.16720684 |
| Knox County, Miller township | 0.5112964 | 0.67213556 | 40.33221364 | -82.51129919 |
| Greene County, Bellbrook city | 0.67186659 | 0.67186659 | 39.63751761 | -84.09522695 |
| Guernsey County, Richland township | 0.47014524 | 0.67159763 | 39.96817287 | -81.47561273 |
| Montgomery County, Vandalia city | 0.51613771 | 0.66998386 | 39.84785271 | -84.17907449 |
| Huron County, Greenfield township | 0.60785368 | 0.66998386 | 41.09913885 | -82.72764921 |
| Muskingum County, Springfield township | 0.62130178 | 0.66890802 | 39.92764266 | -82.1106351 |
| Muskingum County, Newton township | 0.54088219 | 0.66837009 | 39.87698778 | -82.06086933 |
| Huron County, New Haven township | 0.57907477 | 0.66487359 | 41.02658266 | -82.68080658 |
| Richland County, Cass township | 0.43060785 | 0.66433566 | 40.90980346 | -82.64244894 |
| Huron County, Norwich township | 0.51909629 | 0.66325982 | 41.10224367 | -82.75736062 |
| Richland County, Madison township | 0.56307154 | 0.66272189 | 40.78042366 | -82.48593188 |
| Washington County, Muskingum township | 0.53227542 | 0.66003228 | 39.49154501 | -81.45342596 |
| Pickaway County, Wayne township | 0.41191501 | 0.65761162 | 39.5930029 | -82.97663692 |
| Defiance County, Defiance township | 0.44257665 | 0.65734266 | 41.28178998 | -84.3618288 |
| Sandusky County, Jackson township | 0.48036579 | 0.65680473 | 41.32801734 | -83.20084206 |
| Wayne County, Wooster city | 0.62237762 | 0.65653577 | 40.78860194 | -81.94097676 |
| Seneca County, Seneca township | 0.42052179 | 0.65626681 | 41.05947997 | -83.21944336 |
| Huron County, Norwalk city | 0.59494352 | 0.65572889 | 41.24140615 | -82.57927474 |
| Carroll County, Union township | 0.53846154 | 0.65545992 | 40.55802299 | -81.11327303 |
| Licking County, Hopewell township | 0.4852071 | 0.65384615 | 39.98474096 | -82.21891909 |
| Richland County, Springfield township | 0.49569661 | 0.65357719 | 40.76778337 | -82.57884675 |
| Richland County, Plymouth township | 0.4316837 | 0.65330823 | 40.91292763 | -82.67207728 |
| Union County, Dover township | 0.58472297 | 0.65303927 | 40.23504814 | -83.30774162 |
| Medina County, Seville Village township | 0.62587413 | 0.65277031 | 41.02818005 | -81.86273546 |
| Clark County, Pleasant township | 0.40693921 | 0.65223238 | 39.94306156 | -83.53831277 |
| Wayne County, Wooster township | 0.58068854 | 0.65196342 | 40.80738627 | -81.90702004 |
| Scioto County, Portsmouth city | 0.32275417 | 0.65088757 | 38.73808764 | -82.98355004 |
| Trumbull County, Bristol township | 0.47794513 | 0.65061861 | 41.35970763 | -80.82047298 |
| Licking County, St. Albans township | 0.59090909 | 0.6492738 | 40.05130654 | -82.62420068 |
| Fairfield County, Amanda township | 0.45131791 | 0.6492738 | 39.7230475 | -82.71526625 |
| Licking County, Jersey township | 0.5802851 | 0.64819796 | 40.08586874 | -82.73719461 |
| Richland County, Sharon township | 0.54868209 | 0.64766003 | 40.86871635 | -82.68043779 |
| Madison County, Darby township | 0.51371705 | 0.64739107 | 40.09640869 | -83.27273844 |
| Huron County, Willard city | 0.63313609 | 0.64712211 | 41.05180756 | -82.70630641 |
| Guernsey County, Adams township | 0.64523938 | 0.64523938 | 40.05460648 | -81.63743437 |
| Carroll County, Washington township | 0.51802044 | 0.64443249 | 40.62067299 | -81.06988996 |
| Clermont County, Tate township | 0.49018289 | 0.64362561 | 39.00946304 | -84.11043521 |
| Sandusky County, York township | 0.57584723 | 0.64335664 | 41.27249206 | -82.8770405 |
| Licking County, Mary Ann township | 0.4993276 | 0.64254976 | 40.11119293 | -82.3439314 |
| Morrow County, Gilead township | 0.5415546 | 0.6422808 | 40.54644 | -82.83125213 |
| Logan County, Lake township | 0.55607854 | 0.64201183 | 40.36737303 | -83.73338333 |
| Wood County, Webster township | 0.55944056 | 0.64201183 | 41.38564547 | -83.55501529 |
| Clark County, German township | 0.44136633 | 0.64120495 | 40.01615295 | -83.82354459 |
| Williams County, Bryan city | 0.64120495 | 0.64120495 | 41.47809696 | -84.54191618 |
| Montgomery County, Jackson township | 0.50753093 | 0.64093599 | 39.73419776 | -84.40543086 |
| Hancock County, Washington township | 0.49488973 | 0.64066703 | 41.14961174 | -83.44538727 |
| Knox County, Pleasant township | 0.56697149 | 0.64066703 | 40.37006594 | -82.44410275 |
| Marion County, Waldo township | 0.59709521 | 0.64039806 | 40.48243265 | -83.08334732 |
| Wayne County, Baughman township | 0.47996235 | 0.6401291 | 40.85702842 | -81.74590817 |
| Butler County, Oxford township | 0.55043034 | 0.63851533 | 39.51469862 | -84.70685302 |
| Montgomery County, Riverside city | 0.51412049 | 0.63851533 | 39.82294401 | -84.15348823 |
| Henry County, Liberty township | 0.42092523 | 0.63717052 | 41.41485386 | -84.09677004 |
| Stark County, Paris township | 0.54653039 | 0.63690156 | 40.81099204 | -81.15053942 |
| Noble County, Olive township | 0.57248521 | 0.63636364 | 39.72560718 | -81.52489825 |
| Delaware County, Trenton township | 0.56213018 | 0.63609467 | 40.23083285 | -82.82985759 |
| Montgomery County, Oakwood city | 0.63555675 | 0.63555675 | 39.71225896 | -84.17178084 |
| Clermont County, Monroe township | 0.5274341 | 0.63555675 | 38.99288372 | -84.17188332 |
| Wayne County, Plain township | 0.37721893 | 0.63555675 | 40.79844222 | -82.02962339 |
| Trumbull County, Mecca township | 0.50026896 | 0.63474987 | 41.356152 | -80.79074465 |
| Wood County, Weston township | 0.55083378 | 0.6344809 | 41.38374034 | -83.76800752 |
| Morrow County, Bennington township | 0.4310113 | 0.63421194 | 40.36652595 | -82.83465515 |
| Fairfield County, Rush Creek township | 0.42280796 | 0.63421194 | 39.72004685 | -82.47803708 |
| Lorain County, Wellington township | 0.6070468 | 0.63421194 | 41.15577098 | -82.20127071 |
| Huron County, Clarksfield township | 0.58337816 | 0.63367402 | 41.1750319 | -82.37958125 |
| Trumbull County, Cortland city | 0.63313609 | 0.63313609 | 41.32697436 | -80.73611216 |
| Montgomery County, Huber Heights city | 0.47162453 | 0.63286713 | 39.8672187 | -84.14615542 |
| Wayne County, Wayne township | 0.54693384 | 0.63232921 | 40.85813044 | -81.95732866 |
| Marion County, Big Island township | 0.27084454 | 0.63206025 | 40.58593803 | -83.21476015 |
| Clinton County, Chester township | 0.44499731 | 0.63152232 | 39.48507479 | -83.91349774 |
| Tuscarawas County, York township | 0.42415277 | 0.6309844 | 40.44150802 | -81.46871571 |
| Coshocton County, Coshocton city | 0.61888112 | 0.63071544 | 40.26111305 | -81.86553667 |
| Portage County, Paris township | 0.5654922 | 0.63017751 | 41.13479042 | -81.02116636 |
| Marion County, Richland township | 0.50591716 | 0.62963959 | 40.5487706 | -83.07124568 |
| Portage County, Charlestown township | 0.55500269 | 0.62910167 | 41.14875507 | -81.13977089 |
| Lorain County, Camden township | 0.58526089 | 0.62883271 | 41.20962871 | -82.28177403 |
| Delaware County, Oxford township | 0.58068854 | 0.62802582 | 40.40700208 | -83.00711279 |
| Medina County, Litchfield township | 0.52877891 | 0.62802582 | 41.15507324 | -81.98895601 |
| Tuscarawas County, Warwick township | 0.53173749 | 0.6274879 | 40.4346959 | -81.40996346 |
| Clermont County, Wayne township | 0.51694459 | 0.62694997 | 39.18929514 | -84.11025749 |
| Madison County, Jefferson township | 0.60960194 | 0.62668101 | 39.9667006 | -83.32566048 |
| Tuscarawas County, Lawrence township | 0.50618612 | 0.62479828 | 40.61797073 | -81.43240512 |
| Brown County, Sterling township | 0.55433029 | 0.62452932 | 39.03971518 | -83.95853483 |
| Highland County, Whiteoak township | 0.38286713 | 0.62372243 | 39.02553761 | -83.81396748 |
| Ottawa County, Carroll township | 0.4965035 | 0.62345347 | 41.54633031 | -83.13038087 |
| Champaign County, Salem township | 0.40828402 | 0.62130178 | 40.14320521 | -83.74213099 |
| Portage County, Edinburg township | 0.58566434 | 0.62103281 | 41.06399838 | -81.18807234 |
| Columbiana County, West township | 0.53778913 | 0.61995697 | 40.73439642 | -81.07602053 |
| Trumbull County, Bloomfield township | 0.54061323 | 0.61861216 | 41.47706081 | -80.85604432 |
| Huron County, Wakeman township | 0.57799892 | 0.61861216 | 41.2192345 | -82.37099608 |
| Seneca County, Loudon township | 0.57719204 | 0.6183432 | 41.14371525 | -83.38586816 |
| Pickaway County, Madison township | 0.52649274 | 0.61780527 | 39.74464107 | -82.91932354 |
| Portage County, Palmyra township | 0.56024744 | 0.61753631 | 41.11623847 | -81.05551429 |
| Belmont County, Mead township | 0.48830016 | 0.61753631 | 39.92875464 | -80.76654939 |
| Knox County, Morris township | 0.42307692 | 0.61726735 | 40.42058174 | -82.49448402 |
| Clermont County, Ohio township | 0.51990317 | 0.61646046 | 38.99564935 | -84.2007931 |
| Preble County, Israel township | 0.47122109 | 0.61646046 | 39.58906129 | -84.78390822 |
| Medina County, Homer township | 0.43571813 | 0.6161915 | 41.00699993 | -82.07904335 |
| Union County, Millcreek township | 0.57799892 | 0.61592254 | 40.22310776 | -83.19025708 |
| Delaware County, Harlem township | 0.54397526 | 0.61565358 | 40.13625947 | -82.78758238 |
| Hancock County, Eagle township | 0.26022055 | 0.61565358 | 40.99001883 | -83.65512468 |
| Ashtabula County, Andover township | 0.53899946 | 0.61484669 | 41.57352566 | -80.52906536 |
| Champaign County, Mad River township | 0.34642281 | 0.61350188 | 40.03828244 | -83.81976763 |
| Coshocton County, Keene township | 0.44096288 | 0.61323292 | 40.30193871 | -81.82740392 |
| Carroll County, Brown township | 0.45669715 | 0.61242604 | 40.71583564 | -81.11016071 |
| Washington County, Fearing township | 0.40720818 | 0.61215707 | 39.46609462 | -81.42891525 |
| Licking County, Monroe township | 0.577461 | 0.61161915 | 40.13006129 | -82.72896042 |
| Knox County, Monroe township | 0.49246907 | 0.61161915 | 40.39215599 | -82.43987985 |
| Carroll County, Harrison township | 0.50833782 | 0.61135019 | 40.5835488 | -81.13804669 |
| Fairfield County, Bloom township | 0.57934373 | 0.61108123 | 39.77033836 | -82.73622118 |
| Warren County, Wayne township | 0.49569661 | 0.61027434 | 39.48258164 | -84.12088012 |
| Washington County, Marietta city | 0.5739645 | 0.61027434 | 39.42540453 | -81.46683197 |
| Holmes County, Berlin township | 0.50134481 | 0.61000538 | 40.54469569 | -81.77875007 |
| Miami County, Concord township | 0.34198494 | 0.60973642 | 40.03049866 | -84.20839392 |
| Lorain County, Pittsfield township | 0.58821947 | 0.60946746 | 41.20319032 | -82.22230617 |
| Pickaway County, Jackson township | 0.41016676 | 0.60866057 | 39.61510216 | -82.97262254 |
| Fulton County, York township | 0.4873588 | 0.60839161 | 41.58653578 | -84.0066301 |
| Wayne County, Chester township | 0.41984938 | 0.60812265 | 40.81725036 | -81.99568543 |
| Guernsey County, Wills township | 0.40451856 | 0.60731576 | 39.98682341 | -81.44192273 |
| Perry County, Pike township | 0.53254438 | 0.60570199 | 39.71337518 | -82.21171161 |
| Athens County, York township | 0.43692846 | 0.60381926 | 39.4673347 | -82.229333 |
| Montgomery County, Perry township | 0.44620764 | 0.60274341 | 39.75362212 | -84.3726247 |
| Defiance County, Tiffin township | 0.37573964 | 0.60086068 | 41.34828611 | -84.35079486 |
| Fayette County, Jefferson township | 0.47001076 | 0.60059172 | 39.65260065 | -83.55913992 |
| Coshocton County, Franklin township | 0.5091447 | 0.59978483 | 40.21697561 | -81.87431881 |
| Huron County, Hartland township | 0.49475524 | 0.59924691 | 41.20983981 | -82.4942539 |
| Madison County, Canaan township | 0.54733728 | 0.59897795 | 40.01092815 | -83.31780249 |
| Logan County, Liberty township | 0.47525551 | 0.5979021 | 40.32311149 | -83.74102319 |
| Brown County, Washington township | 0.49193115 | 0.59682625 | 39.00056077 | -83.78879558 |
| Lorain County, Brighton township | 0.57907477 | 0.59628833 | 41.19073823 | -82.31582416 |
| Seneca County, Thompson township | 0.44970414 | 0.59601937 | 41.25037288 | -82.88117914 |
| Stark County, Bethlehem township | 0.40344271 | 0.5957504 | 40.71980145 | -81.53214765 |
| Clinton County, Marion township | 0.44486283 | 0.59548144 | 39.31368787 | -84.00124484 |
| Knox County, Howard township | 0.51533082 | 0.59548144 | 40.40469169 | -82.3474294 |
| Pickaway County, Pickaway township | 0.44540075 | 0.59467456 | 39.56479691 | -82.9224694 |
| Coshocton County, Virginia township | 0.4499731 | 0.59467456 | 40.22357128 | -81.93293573 |
| Tuscarawas County, Sandy township | 0.44244217 | 0.59467456 | 40.61455924 | -81.4029565 |
| Butler County, Milford township | 0.45548682 | 0.59467456 | 39.48191847 | -84.59383304 |
| Clark County, Green township | 0.4908553 | 0.59440559 | 39.86124983 | -83.84991818 |
| Delaware County, Marlboro township | 0.5591716 | 0.59359871 | 40.41004035 | -83.03654818 |
| Wood County, Bloom township | 0.39913932 | 0.59332975 | 41.21431528 | -83.64599569 |
| Clinton County, Adams township | 0.54142012 | 0.5922539 | 39.46859104 | -83.97539524 |
| Delaware County, Thompson township | 0.48910705 | 0.59198494 | 40.38089989 | -83.19174917 |
| Clinton County, Washington township | 0.4316837 | 0.59198494 | 39.39090002 | -83.87024503 |
| Hamilton County, Crosby township | 0.59171598 | 0.59171598 | 39.29319639 | -84.74153581 |
| Belmont County, York township | 0.45051103 | 0.59037117 | 39.8662331 | -80.80977909 |
| Delaware County, Genoa township | 0.59010221 | 0.59010221 | 40.18970857 | -82.86737865 |
| Noble County, Sharon township | 0.52165143 | 0.58902636 | 39.73236363 | -81.58308304 |
| Van Wert County, Union township | 0.52124798 | 0.58821947 | 40.90434794 | -84.66518442 |
| Preble County, Jefferson township | 0.2267348 | 0.58795051 | 39.83800961 | -84.8042719 |
| Fairfield County, Liberty township | 0.47458311 | 0.58741259 | 39.91562839 | -82.61982153 |
| Trumbull County, Johnston township | 0.54733728 | 0.58687466 | 41.38594815 | -80.66218581 |
| Morrow County, Westfield township | 0.55056482 | 0.5866057 | 40.42911105 | -83.00306205 |
| Washington County, Belpre township | 0.28348575 | 0.58633674 | 39.30326817 | -81.5803265 |
| Richland County, Franklin township | 0.45508338 | 0.58606778 | 40.83093688 | -82.53663425 |
| Defiance County, Adams township | 0.4020979 | 0.58606778 | 41.367717 | -84.3172218 |
| Ross County, Green township | 0.37681549 | 0.58526089 | 39.41316671 | -82.97963371 |
| Clark County, Mad River township | 0.49354492 | 0.58526089 | 39.85325508 | -83.99988743 |
| Seneca County, Jackson township | 0.48359333 | 0.58526089 | 41.18502113 | -83.34814277 |
| Seneca County, Pleasant township | 0.43222162 | 0.58499193 | 41.18923889 | -83.16550711 |
| Perry County, Thorn township | 0.47337278 | 0.58472297 | 39.91880514 | -82.44037512 |
| Knox County, Liberty township | 0.49475524 | 0.58472297 | 40.40479563 | -82.557514 |
| Butler County, Wayne township | 0.50860678 | 0.58364712 | 39.51816419 | -84.49940122 |
| Medina County, Spencer village | 0.58364712 | 0.58364712 | 41.10186515 | -82.12089006 |
| Richland County, Weller township | 0.44620764 | 0.58337816 | 40.82461908 | -82.47746065 |
| Noble County, Noble township | 0.45266272 | 0.58337816 | 39.75441696 | -81.57863366 |
| Medina County, Spencer township | 0.52071006 | 0.58337816 | 41.12720315 | -82.14621684 |
| Ashland County, Troy township | 0.5330823 | 0.58284024 | 41.00754829 | -82.29091469 |
| Carroll County, Lee township | 0.49381388 | 0.58257127 | 40.55104539 | -81.05445916 |
| Perry County, Clayton township | 0.4112426 | 0.58230231 | 39.73545017 | -82.20745596 |
| Coshocton County, Lafayette township | 0.54679935 | 0.58041958 | 40.27655271 | -81.80248241 |
| Pickaway County, Scioto township | 0.48574502 | 0.58041958 | 39.70959089 | -83.01483472 |
| Muskingum County, Licking township | 0.51344809 | 0.58041958 | 40.02243065 | -82.15189422 |
| Delaware County, Kingston township | 0.47417967 | 0.58015062 | 40.28425795 | -82.90979099 |
| Ottawa County, Allen township | 0.43477676 | 0.57907477 | 41.56727037 | -83.33986003 |
| Wood County, Perry township | 0.42832168 | 0.57880581 | 41.17174679 | -83.44143121 |
| Lorain County, Rochester township | 0.52877891 | 0.57853685 | 41.12123741 | -82.29904284 |
| Marion County, Grand Prairie township | 0.39080151 | 0.57692308 | 40.64628605 | -83.14368059 |
| Huron County, Ridgefield township | 0.52205487 | 0.57611619 | 41.22556778 | -82.64301867 |
| Seneca County, Big Spring township | 0.31293706 | 0.57530931 | 41.07731778 | -83.39777637 |
| Preble County, Gratis township | 0.41393222 | 0.57530931 | 39.62888837 | -84.48166063 |
| Wood County, Grand Rapids township | 0.50242066 | 0.57450242 | 41.4144619 | -83.8538011 |
| Tuscarawas County, Salem township | 0.47982786 | 0.57423346 | 40.29388036 | -81.55898001 |
| Van Wert County, Ridge township | 0.44069392 | 0.5739645 | 40.87150417 | -84.5499689 |
| Licking County, Liberty township | 0.53254438 | 0.57369554 | 40.14280643 | -82.63689908 |
| Wood County, Milton township | 0.39685315 | 0.57369554 | 41.33944207 | -83.77573581 |
| Ashtabula County, Richmond township | 0.4971759 | 0.57235073 | 41.64689084 | -80.57403664 |
| Morrow County, Franklin township | 0.4168908 | 0.57235073 | 40.54026055 | -82.77228887 |
| Greene County, Xenia city | 0.44123185 | 0.57208176 | 39.6927056 | -83.96745482 |
| Greene County, Spring Valley township | 0.44257665 | 0.57154384 | 39.58766522 | -84.04427914 |
| Fulton County, Clinton township | 0.43464228 | 0.56939215 | 41.531234 | -84.13781482 |
| Licking County, Hartford township | 0.53442711 | 0.56885422 | 40.24365233 | -82.73768454 |
| Mercer County, Center township | 0.27380312 | 0.56885422 | 40.58073849 | -84.56693861 |
| Huron County, Bronson township | 0.47202797 | 0.56885422 | 41.19718881 | -82.58773224 |
| Columbiana County, Wayne township | 0.5683163 | 0.5683163 | 40.70624948 | -80.84026583 |
| Trumbull County, Mesopotamia township | 0.51156536 | 0.5683163 | 41.44706758 | -80.98468705 |
| Union County, Allen township | 0.49596557 | 0.56750941 | 40.27780364 | -83.50954775 |
| Madison County, Paint township | 0.39967725 | 0.56724045 | 39.8458109 | -83.4660516 |
| Ottawa County, Erie township | 0.54518558 | 0.56724045 | 41.5372516 | -83.04064247 |
| Monroe County, Ohio township | 0.43181818 | 0.56643357 | 39.67153442 | -80.88093859 |
| Warren County, Massie township | 0.46073158 | 0.56508876 | 39.44927271 | -84.00817852 |
| Clinton County, Vernon township | 0.51237224 | 0.5648198 | 39.35794578 | -83.99387053 |
| Williams County, Pulaski township | 0.40155998 | 0.56455083 | 41.48078002 | -84.57186216 |
| Tuscarawas County, Union township | 0.45346961 | 0.56428187 | 40.47196246 | -81.34209071 |
| Hardin County, Buck township | 0.26734804 | 0.56374395 | 40.62427611 | -83.59868759 |
| Hancock County, Jackson township | 0.28577192 | 0.56347499 | 40.9871233 | -83.62542228 |
| Knox County, College township | 0.55621302 | 0.56347499 | 40.3857952 | -82.38107843 |
| Ashtabula County, Williamsfield township | 0.51075847 | 0.56293706 | 41.53305481 | -80.56863898 |
| Licking County, McKean township | 0.4965035 | 0.56078537 | 40.12708473 | -82.49038293 |
| Ashtabula County, Morgan township | 0.44526627 | 0.56051641 | 41.70824673 | -80.89766116 |
| Brown County, Perry township | 0.44916622 | 0.56051641 | 39.15597938 | -83.99801607 |
| Union County, Darby township | 0.4880312 | 0.56051641 | 40.19378223 | -83.34498986 |
| Marion County, Prospect township | 0.42280796 | 0.56024744 | 40.46334006 | -83.11683699 |
| Ashland County, Vermillion township | 0.40935987 | 0.56024744 | 40.80229038 | -82.27043644 |
| Ashtabula County, Windsor township | 0.52582033 | 0.55970952 | 41.53881519 | -80.99550007 |
| Columbiana County, Hanover township | 0.51492738 | 0.55944056 | 40.75387078 | -80.86025813 |
| Hardin County, Pleasant township | 0.21463152 | 0.55836471 | 40.64930868 | -83.62436735 |
| Ashtabula County, Trumbull township | 0.38407746 | 0.55809575 | 41.71177878 | -80.92754411 |
| Defiance County, Highland township | 0.36740183 | 0.55728886 | 41.2319946 | -84.30950551 |
| Carroll County, Monroe township | 0.49314147 | 0.55540613 | 40.54292513 | -81.17672426 |
| Montgomery County, Clayton city | 0.45790748 | 0.55406132 | 39.83672857 | -84.29972404 |
| Ashtabula County, Orwell township | 0.50174825 | 0.55190963 | 41.50265629 | -80.88104827 |
| Ross County, Springfield township | 0.35207101 | 0.55164067 | 39.39106865 | -82.98363475 |
| Licking County, Washington township | 0.49300699 | 0.55164067 | 40.20909523 | -82.41493089 |
| Holmes County, Mechanic township | 0.49246907 | 0.55164067 | 40.49172987 | -81.90971166 |
| Trumbull County, Greene township | 0.46167294 | 0.55164067 | 41.49200937 | -80.79168189 |
| Huron County, Richmond township | 0.46893491 | 0.55029586 | 41.05801999 | -82.76569022 |
| Wood County, Troy township | 0.49408284 | 0.54975793 | 41.48462415 | -83.41568615 |
| Ashland County, Ruggles township | 0.50161377 | 0.54948897 | 41.01395373 | -82.35022446 |
| Trumbull County, Farmington township | 0.4704142 | 0.54948897 | 41.42501035 | -80.98942957 |
| Licking County, Perry township | 0.48372781 | 0.54922001 | 40.16459551 | -82.21398767 |
| Ashtabula County, Colebrook township | 0.47296934 | 0.54841313 | 41.5140607 | -80.78687285 |
| Knox County, Hilliar township | 0.46853147 | 0.54841313 | 40.2878487 | -82.72943249 |
| Portage County, Hiram township | 0.52017214 | 0.54760624 | 41.33217914 | -81.16173708 |
| Muskingum County, Muskingum township | 0.54088219 | 0.54733728 | 40.0600853 | -82.08479893 |
| Fairfield County, Madison township | 0.44459387 | 0.54679935 | 39.62845787 | -82.67343884 |
| Ashland County, Sullivan township | 0.42953201 | 0.54679935 | 41.00433524 | -82.26126384 |
| Brown County, Clark township | 0.48816568 | 0.54599247 | 38.97333594 | -83.9695768 |
| Huron County, New London township | 0.49623453 | 0.54599247 | 41.12763967 | -82.35844872 |
| Putnam County, Van Buren township | 0.38407746 | 0.54518558 | 41.08786076 | -83.97065984 |
| Ottawa County, Clay township | 0.51533082 | 0.54491662 | 41.54513021 | -83.34387034 |
| Clermont County, Washington township | 0.48682087 | 0.54410974 | 38.83518912 | -84.1682997 |
| Crawford County, Bucyrus city | 0.52716514 | 0.54410974 | 40.80804946 | -82.96341075 |
| Seneca County, Scipio township | 0.48251748 | 0.54303389 | 41.13291193 | -83.05461715 |
| Muskingum County, Cass township | 0.48655191 | 0.54249597 | 40.10097262 | -82.04689808 |
| Knox County, Harrison township | 0.47364174 | 0.54249597 | 40.38260456 | -82.35168165 |
| Greene County, Miami township | 0.40989779 | 0.54142012 | 39.81699276 | -83.85743264 |
| Richland County, Washington township | 0.3598709 | 0.53980635 | 40.6509967 | -82.54079595 |
| Van Wert County, Liberty township | 0.36202259 | 0.53953739 | 40.81036522 | -84.62007029 |
| Ottawa County, Put-in-Bay township | 0.53496503 | 0.53899946 | 41.65165721 | -82.83623184 |
| Highland County, Clay township | 0.48130715 | 0.5387305 | 39.05335257 | -83.86807654 |
| Harrison County, Moorefield township | 0.44526627 | 0.53765465 | 40.18671968 | -81.22108646 |
| Muskingum County, Jackson township | 0.4845347 | 0.53738569 | 40.16136646 | -82.18468918 |
| Shelby County, Clinton township | 0.4281872 | 0.53657881 | 40.29620072 | -84.16433709 |
| Morgan County, Malta township | 0.45723507 | 0.53657881 | 39.6774859 | -81.89159188 |
| Morrow County, Cardington township | 0.40855299 | 0.53550296 | 40.52741334 | -82.86483932 |
| Wood County, Jackson township | 0.39133943 | 0.53496503 | 41.21225973 | -83.85846844 |
| Fairfield County, Richland township | 0.41823561 | 0.53469607 | 39.76421491 | -82.46968573 |
| Morgan County, Morgan township | 0.49973104 | 0.53415815 | 39.67087281 | -81.83342271 |
| Columbiana County, Franklin township | 0.52124798 | 0.53415815 | 40.71332717 | -80.89918698 |
| Athens County, Athens township | 0.44284562 | 0.53388919 | 39.34080163 | -82.10563227 |
| Carroll County, Augusta township | 0.45172136 | 0.5330823 | 40.71234914 | -81.0806872 |
| Morgan County, Bloom township | 0.38542227 | 0.53200646 | 39.6995496 | -81.88724031 |
| Ottawa County, Benton township | 0.39779451 | 0.53146853 | 41.53021286 | -83.19427587 |
| Ottawa County, Harris township | 0.40680473 | 0.53119957 | 41.50807809 | -83.19832989 |
| Tuscarawas County, Warren township | 0.4795589 | 0.53039268 | 40.51263464 | -81.30354593 |
| Wood County, Freedom township | 0.39362561 | 0.53012372 | 41.36058369 | -83.52909183 |
| Portage County, Nelson township | 0.44513179 | 0.52985476 | 41.34727886 | -81.09757199 |
| Marion County, Green Camp township | 0.34830554 | 0.52931684 | 40.53870715 | -83.19324787 |
| Fayette County, Washington Court House city | 0.50793437 | 0.52904788 | 39.55244114 | -83.45803566 |
| Greene County, Xenia township | 0.35314685 | 0.52904788 | 39.66492898 | -83.91282755 |
| Allen County, Shawnee township | 0.33109198 | 0.52904788 | 40.71973599 | -84.1235149 |
| Perry County, Madison township | 0.41648736 | 0.52850995 | 39.88997664 | -82.17759323 |
| Guernsey County, Knox township | 0.47431415 | 0.52850995 | 40.08002196 | -81.66222027 |
| Trumbull County, Kinsman township | 0.50363098 | 0.52850995 | 41.49257343 | -80.60816708 |
| Holmes County, Hardy township | 0.41393222 | 0.52850995 | 40.53257462 | -81.8714808 |
| Highland County, Liberty township | 0.36363636 | 0.52824099 | 39.21024911 | -83.6354293 |
| Carroll County, East township | 0.47296934 | 0.52797203 | 40.72037756 | -80.95811984 |
| Delaware County, Porter township | 0.48910705 | 0.5274341 | 40.29095134 | -82.75880691 |
| Licking County, Fallsbury township | 0.46180742 | 0.52716514 | 40.18667521 | -82.2097059 |
| Ashtabula County, Cherry Valley township | 0.47686928 | 0.52689618 | 41.63204908 | -80.63858724 |
| Lawrence County, Union township | 0.20642819 | 0.52662722 | 38.43299355 | -82.45415487 |
| Hancock County, Blanchard township | 0.27810651 | 0.52662722 | 41.02367411 | -83.77011346 |
| Seneca County, Bloom township | 0.369688 | 0.52662722 | 41.06654635 | -83.06683431 |
| Brown County, Scott township | 0.46745562 | 0.5260893 | 38.98698125 | -83.8792031 |
| Holmes County, Killbuck township | 0.44782141 | 0.52582033 | 40.49502342 | -81.93913335 |
| Madison County, Monroe township | 0.39967725 | 0.52555137 | 39.9947363 | -83.38028923 |
| Van Wert County, Harrison township | 0.42227004 | 0.52501345 | 40.88483386 | -84.69844266 |
| Holmes County, Walnut Creek township | 0.36444325 | 0.52501345 | 40.54136478 | -81.74931533 |
| Licking County, Bennington township | 0.49986552 | 0.52474449 | 40.22490523 | -82.56159844 |
| Lawrence County, Upper township | 0.28671329 | 0.52447552 | 38.49917855 | -82.64644293 |
| Summit County, Clinton village | 0.52447552 | 0.52447552 | 40.93527238 | -81.63922641 |
| Hancock County, Biglick township | 0.29948897 | 0.52420656 | 41.06693555 | -83.52064264 |
| Licking County, Burlington township | 0.46530393 | 0.52420656 | 40.19966591 | -82.5364441 |
| Huron County, Peru township | 0.47229693 | 0.52232383 | 41.20658098 | -82.6769796 |
| Seneca County, Adams township | 0.42684239 | 0.52097902 | 41.23743866 | -82.97464085 |
| Lucas County, Providence township | 0.4612695 | 0.52071006 | 41.43661479 | -83.84995861 |
| Fayette County, Union township | 0.422539 | 0.52017214 | 39.50233209 | -83.40758866 |
| Miami County, Monroe township | 0.37627757 | 0.51990317 | 39.96131423 | -84.19005676 |
| Tuscarawas County, Sugar Creek township | 0.44002152 | 0.51963421 | 40.52460786 | -81.60218306 |
| Van Wert County, Hoaglin township | 0.41420118 | 0.51936525 | 40.94067542 | -84.56889302 |
| Monroe County, Lee township | 0.38407746 | 0.5182894 | 39.62747985 | -80.89025076 |
| Fayette County, Jasper township | 0.32867133 | 0.5182894 | 39.60836482 | -83.56681096 |
| Noble County, Brookfield township | 0.46167294 | 0.51775148 | 39.75778613 | -81.60773885 |
| Lawrence County, Fayette township | 0.28133405 | 0.51748252 | 38.42353226 | -82.57272005 |
| Coshocton County, Pike township | 0.4873588 | 0.51748252 | 40.18020825 | -82.15109389 |
| Morrow County, Canaan township | 0.29303389 | 0.51721356 | 40.57469933 | -82.88613029 |
| Ashland County, Jackson township | 0.41366326 | 0.51667563 | 40.96932478 | -82.14703873 |
| Ross County, Union township | 0.32504034 | 0.51586875 | 39.37201204 | -83.01664587 |
| Fayette County, Paint township | 0.41783217 | 0.51586875 | 39.66887899 | -83.49698994 |
| Trumbull County, Braceville township | 0.49004841 | 0.51586875 | 41.24505553 | -80.99757118 |
| Logan County, Jefferson township | 0.42643895 | 0.51559978 | 40.34236711 | -83.70776672 |
| Sandusky County, Woodville township | 0.44338354 | 0.51506186 | 41.50085099 | -83.35188339 |
| Portage County, Hiram village | 0.51506186 | 0.51506186 | 41.30664262 | -81.13669181 |
| Pickaway County, Darby township | 0.45024207 | 0.51452394 | 39.79402356 | -83.1778998 |
| Wayne County, Franklin township | 0.27972028 | 0.51398601 | 40.7632302 | -81.91584393 |
| Muskingum County, Jefferson township | 0.51371705 | 0.51371705 | 40.11977708 | -82.01329491 |
| Allen County, American township | 0.39026358 | 0.51371705 | 40.74743657 | -84.17903558 |
| Henry County, Bartlow township | 0.38111888 | 0.51371705 | 41.2151015 | -83.88827501 |
| Geauga County, Montville township | 0.46664874 | 0.51344809 | 41.61900359 | -81.100637 |
| Trumbull County, Gustavus township | 0.48628295 | 0.51291017 | 41.49617188 | -80.63794378 |
| Knox County, Milford township | 0.43840775 | 0.5126412 | 40.3385137 | -82.57006802 |
| Stark County, Tuscarawas township | 0.35758472 | 0.51156536 | 40.81144374 | -81.54355679 |
| Henry County, Napoleon township | 0.38972566 | 0.51156536 | 41.3954799 | -84.13042629 |
| Licking County, Bowling Green township | 0.47444863 | 0.5112964 | 39.95341991 | -82.34430735 |
| Crawford County, Whetstone township | 0.29343733 | 0.51075847 | 40.80499333 | -82.93381348 |
| Hocking County, Falls township | 0.38138784 | 0.51048951 | 39.53069959 | -82.39511699 |
| Scioto County, Washington township | 0.24058634 | 0.51022055 | 38.74112972 | -83.0123117 |
| Portage County, Windham township | 0.46369016 | 0.51022055 | 41.20445367 | -81.0366919 |
| Tuscarawas County, Mill township | 0.43679398 | 0.50995159 | 40.42785655 | -81.35122244 |
| Lorain County, Grafton township | 0.50968263 | 0.50968263 | 41.20252739 | -82.00984866 |
| Greene County, Cedarville township | 0.43343195 | 0.5091447 | 39.74205672 | -83.78109417 |
| Ashtabula County, Hartsgrove township | 0.46005917 | 0.50833782 | 41.60147681 | -80.951406 |
| Union County, Liberty township | 0.37197418 | 0.50753093 | 40.29992668 | -83.50565977 |
| Perry County, Pleasant township | 0.38259817 | 0.50753093 | 39.68483774 | -82.15775486 |
| Ashtabula County, Wayne township | 0.46557289 | 0.50753093 | 41.51821751 | -80.63308626 |
| Richland County, Bloominggrove township | 0.44809037 | 0.50726197 | 40.91933881 | -82.51973236 |
| Montgomery County, Butler township | 0.43410436 | 0.50645508 | 39.91703512 | -84.19737212 |
| Licking County, Hanover township | 0.46799354 | 0.50591716 | 40.02889675 | -82.21037741 |
| Guernsey County, Wheeling township | 0.44782141 | 0.50537924 | 40.21238113 | -81.63549947 |
| Noble County, Wayne township | 0.37789134 | 0.50457235 | 39.92067154 | -81.45543174 |
| Richland County, Butler township | 0.44284562 | 0.50457235 | 40.90983932 | -82.43086752 |
| Wayne County, Sugar Creek township | 0.40962883 | 0.50430339 | 40.81288193 | -81.75483842 |
| Clermont County, Franklin township | 0.44163529 | 0.50403443 | 38.832416 | -84.13945478 |
| Tuscarawas County, Oxford township | 0.4817106 | 0.50403443 | 40.27519545 | -81.59278412 |
| Carroll County, Fox township | 0.49462076 | 0.50403443 | 40.56256833 | -80.96156917 |
| Ashland County, Perry township | 0.37641205 | 0.50322754 | 40.86211546 | -82.19834055 |
| Champaign County, Jackson township | 0.28214094 | 0.50188273 | 40.1188597 | -83.95516661 |
| Knox County, Morgan township | 0.43948359 | 0.50080689 | 40.30379668 | -82.45675582 |
| Union County, York township | 0.39335664 | 0.50053792 | 40.4297339 | -83.45281788 |
| Greene County, Caesarscreek township | 0.41837009 | 0.50053792 | 39.57358484 | -83.89859746 |
| Coshocton County, Oxford township | 0.47579344 | 0.50053792 | 40.27856216 | -81.62210197 |
| Ashland County, Mifflin township | 0.33216783 | 0.50053792 | 40.81190103 | -82.35914515 |
| Miami County, Bethel township | 0.34386767 | 0.50026896 | 39.88935621 | -84.14248563 |
| Morrow County, Peru township | 0.48386229 | 0.5 | 40.41995224 | -82.91474413 |
| Stark County, Sandy township | 0.45051103 | 0.49838623 | 40.68905811 | -81.26678919 |
| Harrison County, Green township | 0.46718666 | 0.49811727 | 40.33514503 | -80.94972128 |
| Morgan County, Meigsville township | 0.43329747 | 0.49704142 | 39.64549559 | -81.80871712 |
| Noble County, Enoch township | 0.42522862 | 0.49704142 | 39.74087281 | -81.46224022 |
| Harrison County, North township | 0.42011834 | 0.4965035 | 40.39674259 | -81.08702806 |
| Harrison County, Cadiz township | 0.37130178 | 0.4965035 | 40.29458523 | -80.9883714 |
| Mahoning County, Berlin township | 0.44486283 | 0.49623453 | 40.98843373 | -80.93104193 |
| Trumbull County, Vernon township | 0.49596557 | 0.49596557 | 41.42284021 | -80.59300218 |
| Morgan County, Manchester township | 0.44809037 | 0.49569661 | 39.73909309 | -81.64127873 |
| Carroll County, Perry township | 0.44351802 | 0.49569661 | 40.46287153 | -81.07308721 |
| Washington County, Warren township | 0.32638515 | 0.49488973 | 39.40675136 | -81.50025415 |
| Williams County, Center township | 0.36027434 | 0.49408284 | 41.45860518 | -84.57547975 |
| Madison County, Range township | 0.43329747 | 0.49381388 | 39.70724102 | -83.43093297 |
| Huron County, Fitchville township | 0.42684239 | 0.49381388 | 41.1371913 | -82.44757765 |
| Auglaize County, St. Marys township | 0.28913394 | 0.49354492 | 40.54510501 | -84.42274226 |
| Lucas County, Richfield township | 0.49300699 | 0.49300699 | 41.66386519 | -83.87139082 |
| Fairfield County, Clearcreek township | 0.4436525 | 0.49273803 | 39.5997749 | -82.82710509 |
| Huron County, Ripley township | 0.39160839 | 0.49273803 | 41.04243718 | -82.61725029 |
| Monroe County, Center township | 0.3964497 | 0.49273803 | 39.77265534 | -81.15771987 |
| Highland County, New Market township | 0.43867671 | 0.49193115 | 39.16890312 | -83.67196525 |
| Crawford County, Lykens township | 0.43598709 | 0.49166218 | 40.94988037 | -83.02781922 |
| Crawford County, Holmes township | 0.3126681 | 0.49139322 | 40.8332134 | -82.98893237 |
| Wood County, Montgomery township | 0.38596019 | 0.49139322 | 41.33844214 | -83.53303097 |
| Holmes County, Clark township | 0.3957773 | 0.49112426 | 40.49722793 | -81.75820394 |
| Marion County, Scott township | 0.25484131 | 0.49112426 | 40.65933785 | -83.05102612 |
| Coshocton County, Linton township | 0.42926304 | 0.49004841 | 40.17830422 | -81.73225509 |
| Perry County, Harrison township | 0.40317375 | 0.48951049 | 39.81077412 | -82.07378846 |
| Seneca County, Reed township | 0.38407746 | 0.48897257 | 41.1115167 | -82.84651047 |
| Ashtabula County, Jefferson township | 0.44755245 | 0.48897257 | 41.71966565 | -80.80321238 |
| Jefferson County, Springfield township | 0.47122109 | 0.48816568 | 40.48940634 | -80.91688341 |
| Logan County, Zane township | 0.46100054 | 0.48789672 | 40.26151428 | -83.57222408 |
| Henry County, Washington township | 0.42670791 | 0.48789672 | 41.41730527 | -83.88369306 |
| Crawford County, Chatfield township | 0.40371167 | 0.48628295 | 40.96285039 | -82.93474647 |
| Coshocton County, Bethlehem township | 0.4133943 | 0.48574502 | 40.31185515 | -81.9154314 |
| Carroll County, Rose township | 0.39537386 | 0.4852071 | 40.57189641 | -81.23093524 |
| Hocking County, Good Hope township | 0.39698763 | 0.4852071 | 39.56546006 | -82.50718636 |
| Champaign County, Union township | 0.35287789 | 0.48493814 | 40.1124384 | -83.65790655 |
| Noble County, Buffalo township | 0.43275955 | 0.48493814 | 39.88673784 | -81.55188002 |
| Morrow County, Harmony township | 0.4020979 | 0.48493814 | 40.47085484 | -82.75521616 |
| Tuscarawas County, Fairfield township | 0.41393222 | 0.48493814 | 40.51949648 | -81.36235225 |
| Union County, Taylor township | 0.39550834 | 0.48466918 | 40.28817005 | -83.38803566 |
| Highland County, Hamer township | 0.41070468 | 0.48440022 | 39.17466058 | -83.72988847 |
| Logan County, Monroe township | 0.36713287 | 0.48386229 | 40.27597835 | -83.71924275 |
| Fulton County, Dover township | 0.40572889 | 0.48359333 | 41.59772677 | -84.12652966 |
| Ottawa County, Bay township | 0.46328671 | 0.48332437 | 41.49299187 | -83.04884541 |
| Monroe County, Summit township | 0.44055944 | 0.48305541 | 39.77612218 | -81.18681247 |
| Ashtabula County, Pierpont township | 0.44055944 | 0.48278644 | 41.76074021 | -80.57945183 |
| Harrison County, German township | 0.44943518 | 0.48251748 | 40.42329359 | -80.93097417 |
| Knox County, Butler township | 0.4429801 | 0.48224852 | 40.3729917 | -82.26350713 |
| Hocking County, Green township | 0.42200108 | 0.4817106 | 39.47376349 | -82.28737402 |
| Scioto County, Porter township | 0.23117267 | 0.48144164 | 38.70376067 | -82.8725244 |
| Ashtabula County, Kingsville township | 0.46315223 | 0.48117267 | 41.866926 | -80.70951599 |
| Jefferson County, Salem township | 0.48117267 | 0.48117267 | 40.46030255 | -80.86286301 |
| Preble County, Monroe township | 0.22646584 | 0.48090371 | 39.84430397 | -84.62515909 |
| Ashtabula County, Lenox township | 0.41554599 | 0.48009683 | 41.68690905 | -80.71842686 |
| Huron County, Fairfield township | 0.36309844 | 0.47982786 | 41.11189109 | -82.63432555 |
| Coshocton County, White Eyes township | 0.35476062 | 0.47902098 | 40.32068664 | -81.79364859 |
| Portage County, Deerfield township | 0.45104895 | 0.47848306 | 41.05007965 | -81.06960029 |
| Darke County, Greenville township | 0.30500269 | 0.47794513 | 40.13494871 | -84.60860001 |
| Putnam County, Liberty township | 0.34077461 | 0.47794513 | 41.09067608 | -84.00041847 |
| Brown County, Huntington township | 0.20199032 | 0.47740721 | 38.66017221 | -83.75825256 |
| Mercer County, Hopewell township | 0.28267886 | 0.47713825 | 40.58341376 | -84.59650831 |
| Knox County, Union township | 0.43275955 | 0.47713825 | 40.39507572 | -82.25922818 |
| Coshocton County, Perry township | 0.46100054 | 0.47660032 | 40.24644186 | -82.1381796 |
| Coshocton County, Adams township | 0.42872512 | 0.47660032 | 40.32604553 | -81.64249482 |
| Ashtabula County, Rome township | 0.43827327 | 0.47660032 | 41.59087777 | -80.86190326 |
| Hocking County, Ward township | 0.36793975 | 0.47633136 | 39.48940838 | -82.22509722 |
| Morrow County, Chester township | 0.39349112 | 0.4760624 | 40.48985173 | -82.72162639 |
| Ashtabula County, Sheffield township | 0.42993545 | 0.47471759 | 41.86334273 | -80.6795779 |
| Huron County, Sherman township | 0.39994621 | 0.4739107 | 41.14646847 | -82.74902061 |
| Crawford County, Bucyrus township | 0.2175901 | 0.47364174 | 40.81109872 | -82.99301057 |
| Seneca County, Venice township | 0.37842926 | 0.47364174 | 41.00094422 | -82.8671785 |
| Wayne County, East Union township | 0.36632598 | 0.47337278 | 40.79746048 | -81.81837937 |
| Noble County, Center township | 0.4105702 | 0.4725659 | 39.76631914 | -81.48685625 |
| Paulding County, Benton township | 0.42455621 | 0.4725659 | 41.06747597 | -84.72953318 |
| Richland County, Mifflin township | 0.36686391 | 0.47175901 | 40.79935309 | -82.45212375 |
| Harrison County, Rumley township | 0.44782141 | 0.47122109 | 40.38975357 | -81.0283496 |
| Montgomery County, Englewood city | 0.40290479 | 0.47095212 | 39.83946504 | -84.32897791 |
| Henry County, Harrison township | 0.36148467 | 0.47068316 | 41.39269411 | -84.10053372 |
| Preble County, Eaton city | 0.46637977 | 0.47068316 | 39.7335506 | -84.64273212 |
| Perry County, Jackson township | 0.31737493 | 0.4704142 | 39.71981267 | -82.26994929 |
| Muskingum County, Monroe township | 0.4704142 | 0.4704142 | 40.13417809 | -81.74110917 |
| Paulding County, Blue Creek township | 0.40088757 | 0.46987628 | 41.05951941 | -84.64020874 |
| Brown County, Eagle township | 0.41258741 | 0.46987628 | 38.99769922 | -83.7598998 |
| Morrow County, Lincoln township | 0.38246369 | 0.46960732 | 40.44205842 | -82.91066403 |
| Logan County, Harrison township | 0.31118881 | 0.46880043 | 40.35097176 | -83.79608698 |
| Union County, Leesburg township | 0.37358795 | 0.46853147 | 40.3043682 | -83.32529414 |
| Knox County, Wayne township | 0.32019903 | 0.46799354 | 40.42688993 | -82.55332415 |
| Ashtabula County, Dorset township | 0.45750403 | 0.46772458 | 41.65409779 | -80.63372052 |
| Miami County, Piqua city | 0.40989779 | 0.46772458 | 40.14396755 | -84.21945064 |
| Van Wert County, York township | 0.2493276 | 0.46772458 | 40.80501787 | -84.56074228 |
| Crawford County, Cranberry township | 0.39698763 | 0.46772458 | 40.97575256 | -82.84163973 |
| Carroll County, Loudon township | 0.44513179 | 0.46664874 | 40.4744075 | -80.98031507 |
| Sandusky County, Scott township | 0.35745024 | 0.46637977 | 41.32670097 | -83.41366919 |
| Perry County, Hopewell township | 0.41110812 | 0.46611081 | 39.86189935 | -82.33201473 |
| Coshocton County, Washington township | 0.42428725 | 0.46584185 | 40.19578974 | -82.08817269 |
| Brown County, Pleasant township | 0.41958042 | 0.46557289 | 38.83494117 | -83.93392211 |
| Highland County, Dodson township | 0.4168908 | 0.46530393 | 39.20536897 | -83.81305481 |
| Fulton County, Amboy township | 0.36578806 | 0.46530393 | 41.66670517 | -83.90138972 |
| Harrison County, Archer township | 0.42119419 | 0.46503497 | 40.31662309 | -80.98370354 |
| Sandusky County, Madison township | 0.41608392 | 0.46449704 | 41.34883859 | -83.40969323 |
| Ashtabula County, Denmark township | 0.39913932 | 0.46395912 | 41.7310154 | -80.70873294 |
| Darke County, Brown township | 0.27245831 | 0.46395912 | 40.20673812 | -84.65680243 |
| Morgan County, Bristol township | 0.4077461 | 0.46369016 | 39.68961671 | -81.79996446 |
| Washington County, Adams township | 0.28321678 | 0.46369016 | 39.49833411 | -81.51141561 |
| Muskingum County, Clay township | 0.422539 | 0.46369016 | 39.80751307 | -82.04464034 |
| Harrison County, Franklin township | 0.3725121 | 0.46288327 | 40.36654478 | -81.21362824 |
| Harrison County, Freeport township | 0.40129102 | 0.46153846 | 40.19017249 | -81.2503504 |
| Brown County, Jackson township | 0.32759548 | 0.4612695 | 38.9448389 | -83.68075797 |
| Clinton County, Green township | 0.24260355 | 0.46100054 | 39.42373199 | -83.7465071 |
| Knox County, Jackson township | 0.41581495 | 0.46046261 | 40.25291505 | -82.19684472 |
| Crawford County, Crestline village | 0.46019365 | 0.46019365 | 40.78340781 | -82.72670432 |
| Hocking County, Marion township | 0.40102205 | 0.45911781 | 39.65379513 | -82.49054494 |
| Union County, Washington township | 0.35664336 | 0.45857988 | 40.45185721 | -83.44890403 |
| Preble County, Somers township | 0.42791824 | 0.45804196 | 39.6227996 | -84.66025119 |
| Richland County, Troy township | 0.35274341 | 0.457773 | 40.72358309 | -82.58724818 |
| Harrison County, Short Creek township | 0.41460463 | 0.45669715 | 40.24699831 | -80.96842219 |
| Paulding County, Auglaize township | 0.37520172 | 0.45642819 | 41.19856823 | -84.43614844 |
| Brown County, Lewis township | 0.42926304 | 0.45615923 | 38.85706615 | -83.93023985 |
| Muskingum County, Brush Creek township | 0.43181818 | 0.45508338 | 39.82631435 | -82.01117365 |
| Fayette County, Wayne township | 0.29370629 | 0.45481442 | 39.52149271 | -83.37462137 |
| Champaign County, Goshen township | 0.3605433 | 0.45481442 | 40.10082486 | -83.54056517 |
| Paulding County, Paulding township | 0.4112426 | 0.45481442 | 41.084348 | -84.66641362 |
| Montgomery County, Clay township | 0.2534965 | 0.45481442 | 39.84491756 | -84.38749231 |
| Belmont County, Wayne township | 0.39308768 | 0.45454545 | 39.87588269 | -81.0765172 |
| Preble County, Jackson township | 0.32947821 | 0.45400753 | 39.8158538 | -84.80773538 |
| Guernsey County, Spencer township | 0.43006993 | 0.45400753 | 39.86805873 | -81.58549458 |
| Muskingum County, Meigs township | 0.3844809 | 0.45373857 | 39.81531265 | -81.71536697 |
| Perry County, Reading township | 0.36350188 | 0.45293168 | 39.74188969 | -82.26571111 |
| Athens County, Canaan township | 0.26371705 | 0.45266272 | 39.33428869 | -82.04771408 |
| Champaign County, Wayne township | 0.36605702 | 0.45185584 | 40.22016987 | -83.60933341 |
| Tuscarawas County, Washington township | 0.36377084 | 0.45131791 | 40.2429984 | -81.50935104 |
| Tuscarawas County, Jefferson township | 0.29397526 | 0.45104895 | 40.42623355 | -81.53198763 |
| Clark County, Bethel township | 0.32598171 | 0.45104895 | 39.91118667 | -83.90092505 |
| Coshocton County, Jefferson township | 0.37170522 | 0.45077999 | 40.32171031 | -82.00348323 |
| Noble County, Marion township | 0.39658419 | 0.45051103 | 39.81882758 | -81.35690536 |
| Miami County, Staunton township | 0.27716514 | 0.44943518 | 40.02773383 | -84.17906679 |
| Licking County, Eden township | 0.40613233 | 0.44862829 | 40.17424184 | -82.30189895 |
| Harrison County, Nottingham township | 0.36471221 | 0.44862829 | 40.23080988 | -81.2119053 |
| Monroe County, Wayne township | 0.39833244 | 0.44835933 | 39.73204334 | -81.19595103 |
| Crawford County, Liberty township | 0.28187197 | 0.44809037 | 40.827106 | -82.9297172 |
| Union County, Union township | 0.36444325 | 0.44782141 | 40.18052001 | -83.43698781 |
| Knox County, Clay township | 0.42092523 | 0.44728349 | 40.26577971 | -82.31420672 |
| Carroll County, Orange township | 0.38703604 | 0.44728349 | 40.48369555 | -81.2493505 |
| Holmes County, Richland township | 0.38784293 | 0.44674556 | 40.50812933 | -82.05684723 |
| Richland County, Sandusky township | 0.41231845 | 0.4464766 | 40.78029666 | -82.69712761 |
| Coshocton County, Tiverton township | 0.36874664 | 0.44540075 | 40.45164196 | -82.15811788 |
| Montgomery County, Union City township | 0.33270576 | 0.44513179 | 39.900408 | -84.25957622 |
| Monroe County, Switzerland township | 0.32261969 | 0.44513179 | 39.84775326 | -80.84357582 |
| Noble County, Jackson township | 0.37399139 | 0.44432491 | 39.66283215 | -81.56734717 |
| Preble County, Washington township | 0.31993007 | 0.44405594 | 39.73088735 | -84.61350949 |
| Harrison County, Washington township | 0.38488435 | 0.44405594 | 40.2342649 | -81.24118683 |
| Brown County, Jefferson township | 0.36296396 | 0.44378698 | 38.87068265 | -83.84000299 |
| Athens County, Bern township | 0.36188811 | 0.44351802 | 39.44695145 | -81.84797195 |
| Washington County, Dunham township | 0.28644432 | 0.4429801 | 39.3253183 | -81.57590264 |
| Coshocton County, Bedford township | 0.41608392 | 0.44244217 | 40.29963598 | -82.00783678 |
| Ashland County, Mohican township | 0.24529317 | 0.44217321 | 40.79584895 | -82.21131068 |
| Madison County, Pleasant township | 0.38972566 | 0.44190425 | 39.7617512 | -83.30253167 |
| Preble County, Dixon township | 0.3359333 | 0.44163529 | 39.69455264 | -84.70816252 |
| Guernsey County, Londonderry township | 0.39456697 | 0.44136633 | 40.16812643 | -81.25492795 |
| Madison County, Oak Run township | 0.41500807 | 0.44109736 | 39.79569953 | -83.41539338 |
| Paulding County, Harrison township | 0.42280796 | 0.4408284 | 41.16143945 | -84.77495704 |
| Guernsey County, Oxford township | 0.37089833 | 0.44055944 | 40.06133693 | -81.3069876 |
| Harrison County, Monroe township | 0.37937063 | 0.44029048 | 40.36999966 | -81.24296551 |
| Hocking County, Perry township | 0.38071544 | 0.43786982 | 39.56218901 | -82.6857515 |
| Paulding County, Emerald township | 0.31226466 | 0.43786982 | 41.24831026 | -84.48853622 |
| Perry County, Bearfield township | 0.34480904 | 0.43706294 | 39.72574019 | -82.12009283 |
| Monroe County, Sunsbury township | 0.34911243 | 0.43679398 | 39.85384637 | -81.08112802 |
| Preble County, Gasper township | 0.41258741 | 0.43652501 | 39.69190469 | -84.67895329 |
| Geauga County, Thompson township | 0.37654653 | 0.43652501 | 41.70026876 | -81.0219619 |
| Madison County, Fairfield township | 0.42227004 | 0.43598709 | 39.84721654 | -83.2576056 |
| Noble County, Seneca township | 0.387305 | 0.43598709 | 39.84087361 | -81.35238129 |
| Huron County, Greenwich village | 0.43598709 | 0.43598709 | 41.03300473 | -82.52821783 |
| Morrow County, Congress township | 0.30621302 | 0.43571813 | 40.55926282 | -82.73867092 |
| Henry County, Flatrock township | 0.34023669 | 0.43544917 | 41.29022321 | -84.20886283 |
| Coshocton County, Newcastle township | 0.4020979 | 0.4351802 | 40.30942668 | -82.09588622 |
| Clinton County, Jefferson township | 0.39564282 | 0.43491124 | 39.30523074 | -83.9141646 |
| Athens County, Dover township | 0.34911243 | 0.43464228 | 39.39142901 | -82.15506065 |
| Fayette County, Marion township | 0.32194728 | 0.43464228 | 39.56275516 | -83.33773688 |
| Athens County, Trimble township | 0.38542227 | 0.43410436 | 39.47321346 | -82.07999666 |
| Muskingum County, Madison township | 0.41312534 | 0.4338354 | 40.13857276 | -81.9796741 |
| Belmont County, Flushing township | 0.4338354 | 0.4338354 | 40.16467473 | -81.2256728 |
| Clark County, Madison township | 0.33969876 | 0.43329747 | 39.84694136 | -83.7037209 |
| Guernsey County, Jefferson township | 0.32974718 | 0.43302851 | 40.08522325 | -81.51156437 |
| Ashtabula County, New Lyme township | 0.3992738 | 0.43302851 | 41.58021653 | -80.7724276 |
| Pickaway County, Saltcreek township | 0.34870898 | 0.43222162 | 39.53349166 | -82.83927186 |
| Tuscarawas County, Rush township | 0.38058096 | 0.43195266 | 40.3616987 | -81.36489889 |
| Portage County, Garrettsville village | 0.4316837 | 0.4316837 | 41.27761403 | -81.08195492 |
| Adams County, Winchester township | 0.33754707 | 0.43087682 | 38.94195223 | -83.65188858 |
| Crawford County, Texas township | 0.36282948 | 0.43060785 | 40.97806286 | -83.08308826 |
| Van Wert County, Willshire township | 0.36780527 | 0.43006993 | 40.7571104 | -84.7789909 |
| Guernsey County, Liberty township | 0.38084992 | 0.43006993 | 40.08860848 | -81.54080072 |
| Champaign County, Rush township | 0.34965035 | 0.42980097 | 40.21726525 | -83.5799523 |
| Holmes County, Monroe township | 0.37520172 | 0.42926304 | 40.53020819 | -82.05249612 |
| Henry County, Freedom township | 0.35099516 | 0.42926304 | 41.43980164 | -84.12291508 |
| Muskingum County, Harrison township | 0.41662184 | 0.42764927 | 39.79440124 | -81.92807439 |
| Huron County, Greenwich township | 0.39241528 | 0.42764927 | 41.0045809 | -82.47312778 |
| Champaign County, Johnson township | 0.20804196 | 0.42738031 | 40.14099427 | -83.9514249 |
| Henry County, Damascus township | 0.3718397 | 0.42711135 | 41.39515178 | -83.88752388 |
| Monroe County, Malaga township | 0.35476062 | 0.42684239 | 39.8352937 | -81.11484907 |
| Crawford County, Auburn township | 0.34696073 | 0.42630446 | 40.98858684 | -82.74849916 |
| Preble County, Lanier township | 0.39443249 | 0.42549758 | 39.72285679 | -84.5258546 |
| Putnam County, Blanchard township | 0.27703066 | 0.42549758 | 41.06571128 | -83.97444561 |
| Defiance County, Washington township | 0.27918236 | 0.42495966 | 41.38671706 | -84.52650345 |
| Fulton County, Franklin township | 0.32194728 | 0.42442173 | 41.59760134 | -84.37013203 |
| Morgan County, Homer township | 0.33136095 | 0.422539 | 39.49202188 | -82.04669397 |
| Brown County, Franklin township | 0.39698763 | 0.422539 | 38.89564865 | -83.86514756 |
| Preble County, Twin township | 0.25968263 | 0.42119419 | 39.74768839 | -84.55153387 |
| Lucas County, Jerusalem township | 0.37600861 | 0.42119419 | 41.6558348 | -83.32379349 |
| Washington County, Aurelius township | 0.33526089 | 0.42038731 | 39.60518228 | -81.4600647 |
| Jefferson County, Wayne township | 0.39308768 | 0.42011834 | 40.36862343 | -80.85238345 |
| Pike County, Scioto township | 0.27084454 | 0.41958042 | 38.99026989 | -83.02637463 |
| Clinton County, Clark township | 0.34454008 | 0.41931146 | 39.24961447 | -83.80557666 |
| Muskingum County, Blue Rock township | 0.39187735 | 0.4190425 | 39.79110638 | -81.89893955 |
| Union County, Claibourne township | 0.36175363 | 0.4190425 | 40.38688357 | -83.25061648 |
| Mercer County, Dublin township | 0.34077461 | 0.4190425 | 40.64989239 | -84.5858019 |
| Defiance County, Hicksville township | 0.36229155 | 0.41877353 | 41.29449022 | -84.75368505 |
| Fayette County, Madison township | 0.37385691 | 0.41823561 | 39.67330792 | -83.31820242 |
| Highland County, Union township | 0.36323292 | 0.41742873 | 39.24389213 | -83.74758756 |
| Ross County, Liberty township | 0.13138784 | 0.41742873 | 39.29352024 | -82.91266963 |
| Paulding County, Crane township | 0.3331092 | 0.4168908 | 41.1978576 | -84.6783716 |
| Pickaway County, Deercreek township | 0.36767079 | 0.41662184 | 39.60211521 | -83.0639507 |
| Allen County, Perry township | 0.19311458 | 0.41635288 | 40.71415751 | -84.06430078 |
| Stark County, Sugar Creek township | 0.25174825 | 0.41527703 | 40.72318319 | -81.56164604 |
| Paulding County, Jackson township | 0.31939215 | 0.41420118 | 41.14284945 | -84.56627565 |
| Fulton County, Chesterfield township | 0.29101668 | 0.41420118 | 41.64205725 | -84.11899435 |
| Putnam County, Ottawa township | 0.32208176 | 0.41366326 | 41.06852566 | -84.00419507 |
| Washington County, Watertown township | 0.25578268 | 0.41312534 | 39.45423624 | -81.52032115 |
| Crawford County, Vernon township | 0.34951587 | 0.41285637 | 40.8970475 | -82.73550522 |
| Franklin County, Pleasant township | 0.4112426 | 0.4112426 | 39.82212127 | -83.23233983 |
| Coshocton County, Monroe township | 0.34803658 | 0.4112426 | 40.44189372 | -82.06988438 |
| Wyandot County, Crane township | 0.30311996 | 0.41070468 | 40.83524496 | -83.22990005 |
| Henry County, Richfield township | 0.302851 | 0.41016676 | 41.26223816 | -83.91045834 |
| Tuscarawas County, Auburn township | 0.34938139 | 0.41016676 | 40.48048186 | -81.61116063 |
| Defiance County, Milford township | 0.30083378 | 0.40935987 | 41.34147823 | -84.77647108 |
| Pike County, Pee Pee township | 0.29101668 | 0.40882195 | 39.11980944 | -82.97362998 |
| Ashtabula County, Conneaut city | 0.26573427 | 0.40855299 | 41.91870881 | -80.57508051 |
| Wyandot County, Sycamore township | 0.29236148 | 0.40801506 | 40.98108465 | -83.11276424 |
| Paulding County, Carryall township | 0.3302851 | 0.40667025 | 41.17833529 | -84.71176438 |
| Morrow County, South Bloomfield township | 0.3267886 | 0.40667025 | 40.39834226 | -82.70875765 |
| Ross County, Concord township | 0.285234 | 0.40640129 | 39.43432833 | -83.18289425 |
| Morgan County, Marion township | 0.32651963 | 0.40586337 | 39.46901188 | -81.84362095 |
| Madison County, Pike township | 0.3655191 | 0.40505648 | 40.08615443 | -83.39394219 |
| Guernsey County, Monroe township | 0.34803658 | 0.40451856 | 40.21755066 | -81.48456302 |
| Defiance County, Delaware township | 0.29908553 | 0.40344271 | 41.29264654 | -84.4812657 |
| Madison County, Stokes township | 0.33351264 | 0.4020979 | 39.72186636 | -83.57679231 |
| Noble County, Jefferson township | 0.35435718 | 0.4020979 | 39.68655219 | -81.38399167 |
| Coshocton County, Clark township | 0.33808499 | 0.4020979 | 40.44758325 | -81.91849485 |
| Morgan County, Deerfield township | 0.32759548 | 0.40129102 | 39.684072 | -81.9497717 |
| Tuscarawas County, Clay township | 0.34534696 | 0.40021517 | 40.36853145 | -81.4235868 |
| Logan County, Union township | 0.16151157 | 0.39994621 | 40.32883923 | -83.7998878 |
| Tuscarawas County, Franklin township | 0.33647122 | 0.39913932 | 40.6061025 | -81.52528982 |
| Guernsey County, Washington township | 0.36793975 | 0.39887036 | 40.1886942 | -81.43051786 |
| Greene County, New Jasper township | 0.30580958 | 0.3986014 | 39.67853512 | -83.821576 |
| Mercer County, Butler township | 0.29612695 | 0.39806347 | 40.51960299 | -84.63673951 |
| Ross County, Huntington township | 0.12856374 | 0.39671867 | 39.30571509 | -83.02860097 |
| Crawford County, Jackson township | 0.37116729 | 0.39671867 | 40.78651208 | -82.75628362 |
| Morgan County, Center township | 0.35798817 | 0.39671867 | 39.65422201 | -81.68804666 |
| Shelby County, Turtle Creek township | 0.10247445 | 0.3964497 | 40.27958663 | -84.22687875 |
| Williams County, Springfield township | 0.3211404 | 0.3964497 | 41.4921775 | -84.44843491 |
| Holmes County, Ripley township | 0.28994083 | 0.39591178 | 40.61525387 | -82.00558754 |
| Morgan County, York township | 0.35502959 | 0.3951049 | 39.75026941 | -81.93676129 |
| Ashtabula County, Monroe township | 0.32154384 | 0.3951049 | 41.80122409 | -80.53975584 |
| Miami County, Huber Heights city | 0.3951049 | 0.3951049 | 39.88657622 | -84.11321862 |
| Noble County, Stock township | 0.35664336 | 0.39483593 | 39.73406026 | -81.40405995 |
| Wayne County, Clinton township | 0.25954814 | 0.39429801 | 40.68476526 | -82.02196273 |
| Jefferson County, Smithfield township | 0.38918774 | 0.39322216 | 40.28048943 | -80.8712131 |
| Pike County, Seal township | 0.28752017 | 0.3929532 | 39.05656212 | -83.01446939 |
| Holmes County, Salt Creek township | 0.28227542 | 0.39268424 | 40.59548101 | -81.82878167 |
| Williams County, Superior township | 0.2267348 | 0.39187735 | 41.59434064 | -84.58373481 |
| Mercer County, Black Creek township | 0.35745024 | 0.39107047 | 40.71277692 | -84.78602104 |
| Hocking County, Laurel township | 0.33243679 | 0.39026358 | 39.54337636 | -82.51134036 |
| Williams County, Jefferson township | 0.2732652 | 0.39026358 | 41.60845324 | -84.49010722 |
| Gallia County, Gallipolis township | 0.34615385 | 0.38918774 | 38.81114153 | -82.20724395 |
| Monroe County, Franklin township | 0.35556751 | 0.38891877 | 39.74583458 | -81.31227987 |
| Holmes County, Prairie township | 0.30486821 | 0.38891877 | 40.61197549 | -81.97611308 |
| Morgan County, Penn township | 0.31522324 | 0.38864981 | 39.58262662 | -81.85087189 |
| Greene County, Ross township | 0.28913394 | 0.38784293 | 39.736322 | -83.72271054 |
| Washington County, Wesley township | 0.34991931 | 0.387305 | 39.44363841 | -81.81898075 |
| Williams County, Brady township | 0.3000269 | 0.38569123 | 41.6003248 | -84.40012235 |
| Union County, Jackson township | 0.34050565 | 0.38461538 | 40.46808717 | -83.38603233 |
| Williams County, Northwest township | 0.17859064 | 0.38461538 | 41.63246087 | -84.76018569 |
| Champaign County, Concord township | 0.19983862 | 0.38434642 | 40.14893189 | -83.80084785 |
| Pickaway County, Muhlenberg township | 0.34077461 | 0.38353954 | 39.72470033 | -83.16061343 |
| Scioto County, New Boston village | 0.38300161 | 0.38300161 | 38.75407698 | -82.92205156 |
| Mercer County, Granville township | 0.15653577 | 0.38219473 | 40.4283049 | -84.62141239 |
| Crawford County, Galion City township | 0.38192577 | 0.38192577 | 40.72328796 | -82.79827447 |
| Miami County, Springcreek township | 0.27353416 | 0.3816568 | 40.16611018 | -84.21578932 |
| Belmont County, Somerset township | 0.29881657 | 0.38111888 | 39.90488616 | -81.13018469 |
| Coshocton County, Mill Creek township | 0.35018827 | 0.38084992 | 40.44428501 | -81.88909367 |
| Belmont County, Washington township | 0.36175363 | 0.38058096 | 39.85835239 | -80.93090747 |
| Auglaize County, Noble township | 0.25484131 | 0.38058096 | 40.56454222 | -84.38956726 |
| Auglaize County, Duchouquet township | 0.25874126 | 0.37950511 | 40.56748024 | -84.17904344 |
| Guernsey County, Madison township | 0.34642281 | 0.37923615 | 40.09707799 | -81.41935229 |
| Coshocton County, Crawford township | 0.30365788 | 0.37789134 | 40.42435236 | -81.71274395 |
| Fulton County, Gorham township | 0.32356105 | 0.37762238 | 41.68355804 | -84.32533443 |
| Ashland County, Green township | 0.26371705 | 0.37708445 | 40.64123457 | -82.24148038 |
| Knox County, Brown township | 0.22538999 | 0.37654653 | 40.46775244 | -82.30522706 |
| Marion County, Tully township | 0.3000269 | 0.37627757 | 40.66313049 | -82.86972362 |
| Morgan County, Windsor township | 0.31226466 | 0.3746638 | 39.59805429 | -81.78840725 |
| Fulton County, Royalton township | 0.33566434 | 0.3746638 | 41.66143299 | -84.08521671 |
| Morrow County, Perry township | 0.29962345 | 0.37439484 | 40.55304025 | -82.67970513 |
| Morgan County, Union township | 0.29881657 | 0.37412587 | 39.61458832 | -81.93369309 |
| Noble County, Beaver township | 0.31441635 | 0.37385691 | 39.8815351 | -81.31418067 |
| Logan County, Rushcreek township | 0.20145239 | 0.37385691 | 40.40875744 | -83.69626935 |
| Hocking County, Salt Creek township | 0.32947821 | 0.37305003 | 39.45174146 | -82.70622244 |
| Athens County, Alexander township | 0.26909629 | 0.3725121 | 39.27783891 | -82.14735206 |
| Brown County, Union township | 0.27891339 | 0.3725121 | 38.82643795 | -83.8474214 |
| Tuscarawas County, Perry township | 0.34077461 | 0.37197418 | 40.25143816 | -81.38763677 |
| Fulton County, German township | 0.33405056 | 0.37170522 | 41.52288342 | -84.29131361 |
| Monroe County, Washington township | 0.32140936 | 0.37062937 | 39.5928843 | -81.16525599 |
| Tuscarawas County, Bucks township | 0.30930608 | 0.36928456 | 40.44307103 | -81.6789006 |
| Paulding County, Latty township | 0.34198494 | 0.36874664 | 41.04611972 | -84.49137957 |
| Putnam County, Pleasant township | 0.21328671 | 0.36820871 | 40.9827307 | -84.04899309 |
| Lawrence County, Perry township | 0.20683163 | 0.36767079 | 38.4487526 | -82.59727744 |
| Wayne County, Paint township | 0.17159763 | 0.36767079 | 40.72125872 | -81.74315669 |
| Logan County, Stokes township | 0.25753093 | 0.36686391 | 40.47586422 | -83.92451972 |
| Shelby County, Orange township | 0.13986014 | 0.36686391 | 40.27128339 | -84.13859554 |
| Pickaway County, Monroe township | 0.33539537 | 0.36632598 | 39.63026028 | -83.11820149 |
| Henry County, Pleasant township | 0.30675094 | 0.36632598 | 41.24590268 | -84.21630417 |
| Highland County, Salem township | 0.33647122 | 0.36605702 | 39.11972366 | -83.85692965 |
| Ross County, Colerain township | 0.28684777 | 0.3655191 | 39.46104204 | -82.79333131 |
| Holmes County, Knox township | 0.24260355 | 0.36444325 | 40.53346865 | -82.08194026 |
| Perry County, Salt Lick township | 0.30970952 | 0.36444325 | 39.64069203 | -82.166288 |
| Allen County, Marion township | 0.2408553 | 0.36363636 | 40.84981697 | -84.31252794 |
| Putnam County, Jennings township | 0.24878967 | 0.36363636 | 40.87470848 | -84.33854054 |
| Wyandot County, Mifflin township | 0.14308768 | 0.36202259 | 40.80880915 | -83.41556737 |
| Washington County, Salem township | 0.25524476 | 0.36094675 | 39.53563862 | -81.44447498 |
| Knox County, Jefferson township | 0.21113502 | 0.36094675 | 40.4834134 | -82.24208586 |
| Logan County, Richland township | 0.23345885 | 0.36013986 | 40.50303683 | -83.7399098 |
| Hancock County, Amanda township | 0.22592792 | 0.35960194 | 40.97839524 | -83.53632967 |
| Morrow County, Washington township | 0.30742335 | 0.35933297 | 40.61275645 | -82.81890471 |
| Shelby County, Franklin township | 0.24475524 | 0.35933297 | 40.31834362 | -84.1606509 |
| Wayne County, Salt Creek township | 0.18988704 | 0.35906401 | 40.72459146 | -81.77266564 |
| Hardin County, Liberty township | 0.26667563 | 0.35879505 | 40.7801239 | -83.8122772 |
| Highland County, Concord township | 0.29854761 | 0.35745024 | 39.08042973 | -83.68705852 |
| Harrison County, Stock township | 0.32315761 | 0.35745024 | 40.35265728 | -81.09630762 |
| Jackson County, Jefferson township | 0.15680473 | 0.35718128 | 38.89667794 | -82.57309448 |
| Putnam County, Palmer township | 0.30231307 | 0.35691232 | 41.1046487 | -84.149247 |
| Guernsey County, Millwood township | 0.3154922 | 0.35637439 | 39.96971578 | -81.29599944 |
| Vinton County, Elk township | 0.19983862 | 0.35637439 | 39.24683029 | -82.47827875 |
| Hocking County, Benton township | 0.28160301 | 0.35583647 | 39.44238018 | -82.61913606 |
| Athens County, Troy township | 0.18558365 | 0.35529855 | 39.23515151 | -81.77137324 |
| Logan County, Perry township | 0.29908553 | 0.3542227 | 40.32788903 | -83.5606138 |
| Logan County, Washington township | 0.24408284 | 0.35395374 | 40.45089539 | -83.8987972 |
| Miami County, Union township | 0.19835933 | 0.35368478 | 39.92529592 | -84.28522682 |
| Logan County, Bloomfield township | 0.13084992 | 0.35260893 | 40.43158827 | -83.93205262 |
| Monroe County, Bethel township | 0.31428187 | 0.35233997 | 39.59980731 | -81.22329975 |
| Gallia County, Springfield township | 0.20145239 | 0.35180204 | 38.86172358 | -82.25638369 |
| Washington County, Palmer township | 0.32302313 | 0.35153308 | 39.4590495 | -81.7566294 |
| Crawford County, Polk township | 0.29774072 | 0.35099516 | 40.72019361 | -82.76871965 |
| Monroe County, Benton township | 0.29263045 | 0.35045724 | 39.5859344 | -81.10722339 |
| Monroe County, Seneca township | 0.33418505 | 0.35045724 | 39.81196593 | -81.2986721 |
| Hancock County, Pleasant township | 0.27756859 | 0.35018827 | 41.09296752 | -83.78831493 |
| Defiance County, Farmer township | 0.23695535 | 0.34938139 | 41.41693002 | -84.61262848 |
| Athens County, Lee township | 0.22525551 | 0.34722969 | 39.21809016 | -82.21791585 |
| Hocking County, Starr township | 0.26048951 | 0.34722969 | 39.4548911 | -82.32060557 |
| Preble County, Harrison township | 0.17697687 | 0.34722969 | 39.83628113 | -84.53736469 |
| Ashtabula County, North Kingsville village | 0.31724045 | 0.34615385 | 41.9074547 | -80.66983505 |
| Lawrence County, Rome township | 0.10623991 | 0.34561592 | 38.44553334 | -82.36417736 |
| Monroe County, Salem township | 0.27649274 | 0.345078 | 39.73761633 | -80.86694897 |
| Logan County, McArthur township | 0.28066165 | 0.345078 | 40.41737044 | -83.78467038 |
| Pickaway County, Perry township | 0.29747176 | 0.34480904 | 39.59206437 | -83.18435394 |
| Jackson County, Wellston city | 0.26277569 | 0.34454008 | 39.09541746 | -82.53611929 |
| Ross County, Twin township | 0.12896719 | 0.34427111 | 39.33387295 | -83.08261599 |
| Richland County, Perry township | 0.23413125 | 0.3429263 | 40.5688887 | -82.61657107 |
| Logan County, Bokescreek township | 0.22861754 | 0.34131253 | 40.41639111 | -83.54509984 |
| Paulding County, Brown township | 0.29518558 | 0.33996772 | 41.15423826 | -84.44343758 |
| Holmes County, Washington township | 0.2049489 | 0.33969876 | 40.63801094 | -82.21198438 |
| Noble County, Elk township | 0.29666487 | 0.33781603 | 39.69831163 | -81.29226882 |
| Perry County, Monroe township | 0.29343733 | 0.33727811 | 39.56148804 | -82.06285402 |
| Wyandot County, Tymochtee township | 0.21705218 | 0.33674018 | 40.99010864 | -83.20180734 |
| Paulding County, Washington township | 0.25645508 | 0.3359333 | 41.01854927 | -84.43550267 |
| Williams County, Montpelier village | 0.33566434 | 0.33566434 | 41.57483565 | -84.61734229 |
| Greene County, Silvercreek township | 0.23991393 | 0.33458849 | 39.66995136 | -83.73407635 |
| Van Wert County, Washington township | 0.2584723 | 0.33458849 | 40.85255207 | -84.3422001 |
| Athens County, Ames township | 0.31038193 | 0.3337816 | 39.44788665 | -82.05527216 |
| Washington County, Ludlow township | 0.28886498 | 0.33243679 | 39.57084717 | -81.16982165 |
| Marion County, Salt Rock township | 0.16742873 | 0.33189887 | 40.6522931 | -83.20276346 |
| Darke County, Allen township | 0.11888112 | 0.33109198 | 40.3396759 | -84.63559475 |
| Mercer County, Liberty township | 0.24623453 | 0.33055406 | 40.62411303 | -84.80005505 |
| Williams County, Madison township | 0.23587951 | 0.3302851 | 41.63332543 | -84.51646387 |
| Gallia County, Green township | 0.18006993 | 0.3302851 | 38.83643496 | -82.2318053 |
| Highland County, Washington township | 0.18961807 | 0.32974718 | 39.14100217 | -83.61784535 |
| Henry County, Ridgeville township | 0.29545455 | 0.32947821 | 41.45912016 | -84.33235864 |
| Pike County, Jackson township | 0.12721893 | 0.32894029 | 39.16400197 | -82.96564929 |
| Washington County, Grandview township | 0.28953739 | 0.32867133 | 39.56389944 | -81.11180632 |
| Henry County, Marion township | 0.29343733 | 0.32625067 | 41.18209298 | -84.01508608 |
| Ross County, Deerfield township | 0.26331361 | 0.32544379 | 39.45342924 | -83.14989853 |
| Van Wert County, Jackson township | 0.26990317 | 0.32490586 | 40.97693111 | -84.47250297 |
| Monroe County, Jackson township | 0.27366864 | 0.32490586 | 39.63804568 | -80.97732253 |
| Tuscarawas County, Wayne township | 0.23399677 | 0.32409898 | 40.57210171 | -81.62263308 |
| Hancock County, Union township | 0.2175901 | 0.32409898 | 40.93510659 | -83.78547926 |
| Darke County, Van Buren township | 0.24314147 | 0.32356105 | 40.08529507 | -84.55694227 |
| Mercer County, Recovery township | 0.15922539 | 0.32329209 | 40.41938584 | -84.77248096 |
| Shelby County, Salem township | 0.14806347 | 0.32248521 | 40.31556723 | -84.13120749 |
| Shelby County, McLean township | 0.19392146 | 0.32060247 | 40.33758633 | -84.36681841 |
| Knox County, Fredericktown village | 0.32033351 | 0.32033351 | 40.47107903 | -82.54493665 |
| Miami County, Washington township | 0.1958042 | 0.31952663 | 40.12457977 | -84.25247703 |
| Putnam County, Union township | 0.19849381 | 0.31871974 | 40.99664557 | -84.1976043 |
| Fayette County, Perry township | 0.18908015 | 0.31791286 | 39.46104986 | -83.44440245 |
| Darke County, Twin township | 0.19136633 | 0.31737493 | 39.99132278 | -84.51248444 |
| Washington County, Waterford township | 0.27487897 | 0.31710597 | 39.58135178 | -81.64318831 |
| Fayette County, Concord township | 0.23224852 | 0.31710597 | 39.51406697 | -83.52393669 |
| Marion County, Bowling Green township | 0.27192039 | 0.31683701 | 40.50640612 | -83.31917495 |
| Shelby County, Washington township | 0.11780527 | 0.31629909 | 40.25744229 | -84.23054244 |
| Athens County, Rome township | 0.19176977 | 0.31441635 | 39.36201181 | -81.89430873 |
| Wyandot County, Eden township | 0.20521786 | 0.31441635 | 40.83225078 | -83.20028007 |
| Belmont County, Warren township | 0.31307154 | 0.31307154 | 39.95937004 | -81.20847896 |
| Monroe County, Green township | 0.25511027 | 0.31280258 | 39.68210687 | -80.96806222 |
| Crawford County, Tod township | 0.2105971 | 0.31280258 | 40.82020519 | -83.08182522 |
| Pike County, Marion township | 0.19688004 | 0.31253362 | 39.01611939 | -82.8452089 |
| Putnam County, Greensburg township | 0.23883808 | 0.31145777 | 41.01880034 | -84.19389204 |
| Monroe County, Perry township | 0.25511027 | 0.31091985 | 39.67755696 | -81.11791313 |
| Adams County, Tiffin township | 0.1753631 | 0.31065089 | 38.79765783 | -83.55923266 |
| Jackson County, Lick township | 0.17778375 | 0.309844 | 39.04170449 | -82.66391501 |
| Holmes County, Paint township | 0.20158687 | 0.309844 | 40.60757132 | -81.73596175 |
| Mercer County, Union township | 0.23372781 | 0.30876815 | 40.71100526 | -84.51582928 |
| Allen County, Monroe township | 0.15263583 | 0.30823023 | 40.87198414 | -84.06774799 |
| Pike County, Newton township | 0.23655191 | 0.30742335 | 39.03749882 | -83.04732057 |
| Greene County, Jefferson township | 0.21826251 | 0.30742335 | 39.57570892 | -83.69094705 |
| Defiance County, Mark township | 0.26949973 | 0.3063475 | 41.28654888 | -84.66407043 |
| Morrow County, Troy township | 0.24946208 | 0.3063475 | 40.66354442 | -82.65887596 |
| Monroe County, Adams township | 0.27030662 | 0.30607854 | 39.78075456 | -81.03676068 |
| Darke County, Monroe township | 0.20629371 | 0.30500269 | 39.98862887 | -84.48316174 |
| Richland County, Monroe township | 0.17348037 | 0.30204411 | 40.70779728 | -82.439546 |
| Scioto County, Clay township | 0.20938677 | 0.30150619 | 38.78532233 | -83.00439586 |
| Williams County, St. Joseph township | 0.25551372 | 0.30123722 | 41.43282378 | -84.79219824 |
| Henry County, Monroe township | 0.2493276 | 0.29908553 | 41.326217 | -84.11181057 |
| Adams County, Wayne township | 0.22364174 | 0.29854761 | 38.89771849 | -83.65942848 |
| Jackson County, Jackson city | 0.23534158 | 0.29693384 | 39.0385813 | -82.63504923 |
| Allen County, Sugar Creek township | 0.20575578 | 0.29693384 | 40.83603782 | -84.1642016 |
| Auglaize County, Moulton township | 0.16729424 | 0.29612695 | 40.55908838 | -84.33046148 |
| Auglaize County, Pusheta township | 0.15357719 | 0.29558903 | 40.54809811 | -84.21227717 |
| Scioto County, Valley township | 0.16218397 | 0.29532006 | 38.94607483 | -83.03429942 |
| Perry County, Monday Creek township | 0.24394836 | 0.29478214 | 39.64109846 | -82.3741473 |
| Scioto County, Harrison township | 0.15223238 | 0.29343733 | 38.83631006 | -82.8485061 |
| Crawford County, Jefferson township | 0.23856912 | 0.29316837 | 40.78960948 | -82.78586551 |
| Scioto County, Green township | 0.11484669 | 0.29289941 | 38.67859514 | -82.8477864 |
| Athens County, Waterloo township | 0.21180742 | 0.29209252 | 39.37259229 | -82.18828887 |
| Perry County, Coal township | 0.24946208 | 0.29209252 | 39.58092604 | -82.23719084 |
| Clark County, Pike township | 0.20669715 | 0.2915546 | 39.9611039 | -83.95200285 |
| Wyandot County, Crawford township | 0.17482517 | 0.29128564 | 40.96371032 | -83.38789101 |
| Mercer County, Washington township | 0.1781872 | 0.29101668 | 40.5470751 | -84.69230285 |
| Knox County, Middlebury township | 0.24273803 | 0.29047875 | 40.50259402 | -82.62909907 |
| Washington County, Fairfield township | 0.20360409 | 0.28913394 | 39.41493483 | -81.76538059 |
| Gallia County, Addison township | 0.21463152 | 0.28886498 | 38.91501547 | -82.12860477 |
| Vinton County, Clinton township | 0.13569123 | 0.28832706 | 39.16166305 | -82.52374923 |
| Adams County, Scott township | 0.16890802 | 0.28778913 | 38.93615875 | -83.59415669 |
| Hocking County, Washington township | 0.19688004 | 0.28698225 | 39.46763687 | -82.43669401 |
| Ashland County, Hanover township | 0.17374933 | 0.28536848 | 40.61914761 | -82.24577964 |
| Allen County, Richland township | 0.17267348 | 0.28536848 | 40.83860144 | -83.95286418 |
| Pike County, Sunfish township | 0.15895643 | 0.28429263 | 39.03644443 | -83.2534497 |
| Wyandot County, Antrim township | 0.12452932 | 0.28375471 | 40.80712931 | -83.17468395 |
| Meigs County, Lebanon township | 0.12264658 | 0.28321678 | 38.94177099 | -81.77025254 |
| Washington County, Liberty township | 0.23076923 | 0.28294782 | 39.55917796 | -81.2614025 |
| Morrow County, North Bloomfield township | 0.25228618 | 0.2824099 | 40.69498768 | -82.74331659 |
| Jackson County, Scioto township | 0.10193653 | 0.28160301 | 39.0099506 | -82.78748462 |
| Hardin County, Washington township | 0.14685315 | 0.28106509 | 40.78789766 | -83.66032954 |
| Ross County, Buckskin township | 0.20952125 | 0.28079613 | 39.36376268 | -83.37274855 |
| Williams County, Bridgewater township | 0.15088757 | 0.28052717 | 41.61919326 | -84.61011929 |
| Mercer County, Franklin township | 0.18894567 | 0.2802582 | 40.51158502 | -84.54811191 |
| Darke County, Liberty township | 0.19244217 | 0.27972028 | 40.05429954 | -84.71080122 |
| Putnam County, Monroe township | 0.20602474 | 0.27837547 | 41.09354325 | -84.27204596 |
| Jackson County, Jackson township | 0.11941904 | 0.27810651 | 39.1614773 | -82.73023665 |
| Crawford County, Sandusky township | 0.22149005 | 0.27595481 | 40.85902663 | -82.80304492 |
| Washington County, Newport township | 0.20831092 | 0.27568585 | 39.38971582 | -81.35548659 |
| Adams County, Liberty township | 0.1732114 | 0.27541689 | 38.77845147 | -83.59182441 |
| Brown County, Byrd township | 0.20010758 | 0.27514793 | 38.8399941 | -83.75720705 |
| Clinton County, Richland township | 0.18544917 | 0.27353416 | 39.44298142 | -83.71366563 |
| Darke County, Adams township | 0.13313609 | 0.27245831 | 40.10744668 | -84.55338824 |
| Darke County, Neave township | 0.19244217 | 0.27138246 | 40.04633572 | -84.62274019 |
| Jackson County, Franklin township | 0.17159763 | 0.27057558 | 38.94084271 | -82.56489522 |
| Putnam County, Monterey township | 0.24690694 | 0.27003765 | 40.94663602 | -84.38697241 |
| Allen County, Amanda township | 0.14766003 | 0.26896181 | 40.75296169 | -84.23828638 |
| Darke County, Butler township | 0.2049489 | 0.26788596 | 39.94088029 | -84.69898233 |
| Hardin County, McDonald township | 0.11417429 | 0.26734804 | 40.52516996 | -83.73608107 |
| Vinton County, Eagle township | 0.19284562 | 0.26438946 | 39.3633837 | -82.72255423 |
| Clinton County, Wilson township | 0.20804196 | 0.26385153 | 39.54491566 | -83.60740623 |
| Shelby County, Dinsmore township | 0.11941904 | 0.26385153 | 40.40691708 | -84.14588326 |
| Knox County, Pike township | 0.1753631 | 0.26358257 | 40.48369209 | -82.45240236 |
| Pike County, Camp Creek township | 0.16648736 | 0.26331361 | 38.97120366 | -83.05919278 |
| Washington County, Independence township | 0.23722431 | 0.2619688 | 39.50819628 | -81.21248788 |
| Williams County, Florence township | 0.21221087 | 0.2619688 | 41.54635827 | -84.80441864 |
| Williams County, Mill Creek township | 0.22646584 | 0.26143088 | 41.62792562 | -84.45644836 |
| Meigs County, Salisbury township | 0.19123185 | 0.26089295 | 38.9779674 | -82.08707676 |
| Ross County, Paxton township | 0.18854223 | 0.26089295 | 39.21924896 | -83.28010967 |
| Jackson County, Milton township | 0.10032275 | 0.26035503 | 39.07333552 | -82.54023763 |
| Darke County, Richland township | 0.16756321 | 0.26008607 | 40.22356424 | -84.5944248 |
| Washington County, Lawrence township | 0.16245293 | 0.25954814 | 39.45584359 | -81.34198536 |
| Champaign County, Harrison township | 0.14039806 | 0.25927918 | 40.21532328 | -83.7894679 |
| Highland County, Penn township | 0.1929801 | 0.25874126 | 39.27949463 | -83.65304684 |
| Adams County, Meigs township | 0.13905325 | 0.25766541 | 38.93777879 | -83.38831692 |
| Jackson County, Coal township | 0.1668908 | 0.25766541 | 39.09856368 | -82.56500338 |
| Wyandot County, Salem township | 0.15223238 | 0.25739645 | 40.89140609 | -83.3404635 |
| Ross County, Harrison township | 0.1788596 | 0.2563206 | 39.36648677 | -82.75155329 |
| Shelby County, Cynthian township | 0.18047337 | 0.25497579 | 40.29328828 | -84.37406276 |
| Adams County, Monroe township | 0.14631522 | 0.25470683 | 38.75343045 | -83.5668138 |
| Allen County, Jackson township | 0.15935987 | 0.25443787 | 40.81645692 | -83.95664221 |
| Marion County, Montgomery township | 0.16272189 | 0.25389995 | 40.57869804 | -83.36632448 |
| Pike County, Beaver township | 0.19284562 | 0.25336202 | 39.03821111 | -82.84118644 |
| Adams County, Franklin township | 0.15720818 | 0.25255514 | 39.01731807 | -83.28622905 |
| Van Wert County, Jennings township | 0.19876278 | 0.25255514 | 40.76937143 | -84.41609377 |
| Highland County, Fairfield township | 0.16178053 | 0.25174825 | 39.33714127 | -83.55459191 |
| Wyandot County, Pitt township | 0.11403981 | 0.25040344 | 40.79100047 | -83.23790404 |
| Pike County, Union township | 0.1457773 | 0.25013448 | 39.02226136 | -82.90294295 |
| Putnam County, Perry township | 0.18665949 | 0.24905863 | 41.07686621 | -84.33526732 |
| Jackson County, Bloomfield township | 0.12399139 | 0.24852071 | 38.98185913 | -82.52784733 |
| Lawrence County, Elizabeth township | 0.11672942 | 0.24663798 | 38.61584216 | -82.68354296 |
| Putnam County, Jackson township | 0.19096288 | 0.24583109 | 40.99940777 | -84.22733353 |
| Fayette County, Green township | 0.154922 | 0.24556213 | 39.44187121 | -83.47732368 |
| Wyandot County, Ridge township | 0.19499731 | 0.24556213 | 40.96960501 | -83.44725908 |
| Shelby County, Perry township | 0.14254976 | 0.24529317 | 40.33213428 | -84.06861214 |
| Darke County, Wabash township | 0.19244217 | 0.24475524 | 40.33433758 | -84.57665633 |
| Scioto County, Rush township | 0.154922 | 0.24448628 | 38.81045668 | -83.02922405 |
| Vinton County, Jackson township | 0.14658419 | 0.24448628 | 39.37298961 | -82.60246903 |
| Hancock County, Orange township | 0.17240452 | 0.24341044 | 40.82440336 | -83.80463258 |
| Darke County, Jackson township | 0.18760086 | 0.24179666 | 40.21993817 | -84.8039169 |
| Highland County, Madison township | 0.19324906 | 0.23964497 | 39.36671454 | -83.40177492 |
| Hardin County, Taylor Creek township | 0.14685315 | 0.23830016 | 40.58580678 | -83.66550486 |
| Hardin County, Hale township | 0.14470145 | 0.23830016 | 40.51822829 | -83.4371479 |
| Scioto County, Morgan township | 0.17011834 | 0.23776224 | 38.94910517 | -83.06314538 |
| Mercer County, Gibson township | 0.11538462 | 0.23641743 | 40.3972242 | -84.77598252 |
| Knox County, Berlin township | 0.2027972 | 0.2345347 | 40.4900201 | -82.51129085 |
| Putnam County, Riley township | 0.16783217 | 0.23292093 | 40.9771163 | -83.98956502 |
| Scioto County, Nile township | 0.09709521 | 0.23157612 | 38.62062909 | -83.15086769 |
| Darke County, Franklin township | 0.15357719 | 0.23157612 | 40.00807572 | -84.45026031 |
| Miami County, Newton township | 0.14174287 | 0.23023131 | 40.01660863 | -84.30001879 |
| Hardin County, Marion township | 0.18652501 | 0.22996235 | 40.71085093 | -83.79413246 |
| Washington County, Barlow township | 0.20118343 | 0.22969338 | 39.35745943 | -81.65830788 |
| Hardin County, Lynn township | 0.14376009 | 0.22969338 | 40.63295842 | -83.68734454 |
| Auglaize County, Union township | 0.06508876 | 0.2288865 | 40.60343096 | -84.08296113 |
| Jackson County, Madison township | 0.16742873 | 0.22754169 | 38.93769656 | -82.53607369 |
| Richland County, Jefferson township | 0.16191501 | 0.22727273 | 40.62574651 | -82.51549838 |
| Crawford County, Dallas township | 0.13932222 | 0.22727273 | 40.70659674 | -83.0724804 |
| Highland County, Brushcreek township | 0.12412587 | 0.22700377 | 39.16477129 | -83.40756827 |
| Allen County, Spencer township | 0.19903174 | 0.22512103 | 40.69746464 | -84.36775318 |
| Pike County, Pebble township | 0.14658419 | 0.22485207 | 39.08472459 | -83.06829374 |
| Hardin County, Jackson township | 0.1161915 | 0.22431415 | 40.79841225 | -83.53796648 |
| Hardin County, Roundhead township | 0.09265734 | 0.22377622 | 40.53660392 | -83.85415279 |
| Darke County, Harrison township | 0.13932222 | 0.22350726 | 39.9656787 | -84.7247994 |
| Hancock County, Madison township | 0.15061861 | 0.22296934 | 40.82927245 | -83.62295305 |
| Wyandot County, Jackson township | 0.14389457 | 0.22189349 | 40.81175316 | -83.44518656 |
| Auglaize County, Washington township | 0.12856374 | 0.22054868 | 40.53693723 | -84.33410734 |
| Auglaize County, Logan township | 0.08808499 | 0.22001076 | 40.66160173 | -84.22342055 |
| Ross County, Franklin township | 0.11686391 | 0.21866595 | 39.1860982 | -82.9616553 |
| Athens County, Lodi township | 0.13864981 | 0.21490048 | 39.26808867 | -82.0605485 |
| Richland County, Worthington township | 0.15048413 | 0.21463152 | 40.56250512 | -82.34705799 |
| Scioto County, Union township | 0.13340506 | 0.21436256 | 38.86373055 | -83.10774582 |
| Miami County, Newberry township | 0.14846692 | 0.21382464 | 40.11065695 | -84.34421361 |
| Gallia County, Cheshire township | 0.1070468 | 0.21274879 | 38.93708301 | -82.12437059 |
| Adams County, Sprigg township | 0.14443249 | 0.21247983 | 38.65442844 | -83.7007348 |
| Ashland County, Lake township | 0.17173211 | 0.21167294 | 40.68218335 | -82.20335992 |
| Auglaize County, Jackson township | 0.15061861 | 0.21113502 | 40.40675897 | -84.38542356 |
| Auglaize County, Goshen township | 0.14739107 | 0.2105971 | 40.54510724 | -83.94273135 |
| Adams County, Oliver township | 0.15330823 | 0.21032813 | 38.92741816 | -83.50757612 |
| Washington County, Decatur township | 0.14981173 | 0.21032813 | 39.35540071 | -81.83639277 |
| Gallia County, Raccoon township | 0.10758472 | 0.20952125 | 38.89660352 | -82.36735782 |
| Adams County, Green township | 0.11323292 | 0.20925229 | 38.61347286 | -83.29836644 |
| Hardin County, Dudley township | 0.11780527 | 0.20736955 | 40.56247667 | -83.4292983 |
| Ross County, Paint township | 0.15989779 | 0.20629371 | 39.24432967 | -83.30518191 |
| Jackson County, Liberty township | 0.09951587 | 0.20575578 | 39.11108143 | -82.68056471 |
| Miami County, Lostcreek township | 0.10341581 | 0.2049489 | 40.04710143 | -84.14606516 |
| Highland County, Paint township | 0.16110812 | 0.20199032 | 39.20603945 | -83.37088387 |
| Hancock County, Delaware township | 0.13179129 | 0.20037655 | 40.8263698 | -83.59331922 |
| Lawrence County, Hamilton township | 0.12506724 | 0.19983862 | 38.57478069 | -82.72031819 |
| Vinton County, Swan township | 0.14766003 | 0.19741797 | 39.36355624 | -82.51549089 |
| Putnam County, Sugar Creek township | 0.12520172 | 0.19634212 | 40.88587606 | -84.21613081 |
| Adams County, Manchester township | 0.19338354 | 0.19338354 | 38.68999255 | -83.60693406 |
| Scioto County, Bloom township | 0.14846692 | 0.19311458 | 38.82394913 | -82.73335142 |
| Mercer County, Marion township | 0.1161915 | 0.19284562 | 40.40080384 | -84.56596879 |
| Miami County, Elizabeth township | 0.15236686 | 0.19230769 | 40.00004548 | -84.1241027 |
| Clinton County, Wayne township | 0.15236686 | 0.19230769 | 39.43432309 | -83.62646387 |
| Pike County, Perry township | 0.13421194 | 0.19069392 | 39.17800746 | -83.31685235 |
| Meigs County, Chester township | 0.09601937 | 0.19069392 | 39.05642872 | -81.98351906 |
| Meigs County, Columbia township | 0.11148467 | 0.18854223 | 39.19601832 | -82.22213797 |
| Gallia County, Guyan township | 0.13421194 | 0.18639053 | 38.61892815 | -82.30233286 |
| Hardin County, Cessna township | 0.1429532 | 0.18477676 | 40.73012541 | -83.76070893 |
| Ross County, Jefferson township | 0.12237762 | 0.18477676 | 39.18976336 | -82.78402831 |
| Gallia County, Greenfield township | 0.1422808 | 0.18370091 | 38.84937057 | -82.5524965 |
| Miami County, Brown township | 0.14039806 | 0.18370091 | 40.15779976 | -84.12764727 |
| Scioto County, Madison township | 0.09978483 | 0.18262507 | 38.86147577 | -82.87329788 |
| Scioto County, Rarden township | 0.09440559 | 0.18208714 | 38.92293814 | -83.24406618 |
| Darke County, Patterson township | 0.07826789 | 0.18208714 | 40.27657044 | -84.43655295 |
| Hardin County, Blanchard township | 0.14093599 | 0.18208714 | 40.74073617 | -83.63846018 |
| Athens County, Carthage township | 0.12426036 | 0.1810113 | 39.27705698 | -81.9405369 |
| Logan County, Miami township | 0.12224314 | 0.18047337 | 40.29876496 | -83.95459535 |
| Wyandot County, Richland township | 0.09198494 | 0.17751479 | 40.83388209 | -83.44124926 |
| Vinton County, Richland township | 0.11686391 | 0.17670791 | 39.25314307 | -82.53616508 |
| Darke County, Wayne township | 0.13609467 | 0.1753631 | 40.23226996 | -84.44374795 |
| Auglaize County, German township | 0.14591178 | 0.17509414 | 40.45106072 | -84.37817474 |
| Darke County, Washington township | 0.14685315 | 0.17428725 | 40.14556707 | -84.72616222 |
| Highland County, Marshall township | 0.15303927 | 0.17401829 | 39.17065181 | -83.46546704 |
| Vinton County, Harrison township | 0.13044648 | 0.17374933 | 39.26880733 | -82.68092483 |
| Allen County, Auglaize township | 0.09198494 | 0.17132867 | 40.71939715 | -83.88292099 |
| Wyandot County, Marseilles township | 0.1492738 | 0.17105971 | 40.71735268 | -83.40174777 |
| Shelby County, Loramie township | 0.1035503 | 0.17025282 | 40.25170896 | -84.41072374 |
| Auglaize County, Salem township | 0.14752555 | 0.16944594 | 40.67803134 | -84.40099614 |
| Highland County, Jackson township | 0.13421194 | 0.16917698 | 39.06884827 | -83.57137833 |
| Vinton County, Wilkesville township | 0.0894298 | 0.16890802 | 39.09528121 | -82.32982 |
| Gallia County, Ohio township | 0.14389457 | 0.16837009 | 38.62812898 | -82.18341654 |
| Hancock County, Van Buren township | 0.12654653 | 0.16568047 | 40.84081406 | -83.74151254 |
| Jackson County, Washington township | 0.0922539 | 0.16514255 | 39.14272939 | -82.5567764 |
| Gallia County, Walnut township | 0.11861216 | 0.1640667 | 38.75158777 | -82.4826349 |
| Vinton County, Knox township | 0.11377084 | 0.16352878 | 39.24659767 | -82.27154894 |
| Lawrence County, Mason township | 0.1035503 | 0.16299086 | 38.67266069 | -82.38011687 |
| Darke County, York township | 0.13555675 | 0.16164605 | 40.29269857 | -84.61322043 |
| Pike County, Mifflin township | 0.10166756 | 0.16110812 | 39.05854924 | -83.24955022 |
| Scioto County, Brush Creek township | 0.10059172 | 0.16003228 | 38.87275015 | -83.19420217 |
| Lawrence County, Aid township | 0.09628833 | 0.1584185 | 38.59387282 | -82.48270483 |
| Jackson County, Hamilton township | 0.10623991 | 0.15734266 | 38.86812405 | -82.72525746 |
| Marion County, Grand township | 0.14712211 | 0.1570737 | 40.67014931 | -83.3800716 |
| Darke County, Mississinawa township | 0.10839161 | 0.15599785 | 40.28641901 | -84.79345774 |
| Vinton County, Brown township | 0.12708445 | 0.15545992 | 39.30672678 | -82.40790189 |
| Lawrence County, Symmes township | 0.10892953 | 0.15519096 | 38.72950859 | -82.48675003 |
| Logan County, Pleasant township | 0.11726735 | 0.15465304 | 40.33737945 | -83.88820235 |
| Lawrence County, Washington township | 0.1274879 | 0.15384615 | 38.83669285 | -82.64293972 |
| Adams County, Jefferson township | 0.12116729 | 0.15277031 | 38.75204462 | -83.33270728 |
| Adams County, Bratton township | 0.11565358 | 0.14873588 | 39.00705541 | -83.4056332 |
| Lawrence County, Lawrence township | 0.10960194 | 0.14766003 | 38.55916442 | -82.57692265 |
| Meigs County, Sutton township | 0.1105433 | 0.14685315 | 39.03436561 | -81.98780293 |
| Scioto County, Vernon township | 0.11323292 | 0.14497041 | 38.71660589 | -82.78228581 |
| Adams County, Brush Creek township | 0.12439484 | 0.1422808 | 38.75795969 | -83.39027263 |
| Shelby County, Van Buren township | 0.11377084 | 0.14201183 | 40.44011578 | -84.26016199 |
| Gallia County, Perry township | 0.10785368 | 0.14066703 | 38.78622444 | -82.38812384 |
| Meigs County, Olive township | 0.10153308 | 0.1401291 | 39.09948714 | -81.76863561 |
| Gallia County, Huntington township | 0.07638515 | 0.13878429 | 38.98490535 | -82.35069966 |
| Pike County, Benton township | 0.08606778 | 0.13690156 | 39.1911775 | -83.22610342 |
| Meigs County, Letart township | 0.11995697 | 0.13528779 | 38.92966458 | -81.860944 |
| Gallia County, Harrison township | 0.10099516 | 0.13501883 | 38.68834965 | -82.31854276 |
| Vinton County, Madison township | 0.10798817 | 0.13474987 | 39.2814662 | -82.38311973 |
| Hardin County, Goshen township | 0.0979021 | 0.13394298 | 40.67604639 | -83.43919403 |
| Gallia County, Morgan township | 0.08243679 | 0.13071544 | 39.00378373 | -82.3176906 |
| Meigs County, Scipio township | 0.05325444 | 0.12990855 | 39.18955975 | -82.1643249 |
| Lawrence County, Windsor township | 0.09884346 | 0.12856374 | 38.51812867 | -82.40905592 |
| Gallia County, Clay township | 0.09359871 | 0.12614309 | 38.76700136 | -82.21564502 |
| Meigs County, Bedford township | 0.10112964 | 0.1253362 | 39.12589527 | -81.99952176 |
| Scioto County, Jefferson township | 0.07880581 | 0.12479828 | 38.93694357 | -82.947776 |
| Champaign County, Adams township | 0.07826789 | 0.12426036 | 40.21021961 | -83.96957735 |
| Auglaize County, Wayne township | 0.08176439 | 0.12426036 | 40.58938711 | -83.93519308 |
| Vinton County, Vinton township | 0.07611619 | 0.11915008 | 39.11735622 | -82.32563646 |
| Shelby County, Jackson township | 0.08445401 | 0.1183432 | 40.39855654 | -84.05745207 |
| Meigs County, Salem township | 0.077461 | 0.11807423 | 39.11415438 | -82.29675536 |
| Meigs County, Orange township | 0.10233997 | 0.11726735 | 39.16674342 | -81.96206064 |
| Lawrence County, Decatur township | 0.08364712 | 0.10892953 | 38.74522958 | -82.63046091 |
| Auglaize County, Clay township | 0.06132329 | 0.1035503 | 40.48712226 | -84.04253964 |
| Meigs County, Rutland township | 0.0570199 | 0.10166756 | 39.00652995 | -82.14048152 |
| Shelby County, Green township | 0.0626681 | 0.09817106 | 40.24077952 | -84.05405038 |
