## Supplementary material for "Mapping long-distance jump dispersal risk by leveraging transport infrastructure data and sampling bias correction: a case study with the invasive spotted lanternfly (*Lycorma delicatula*)": S12 Pennsylvania risk

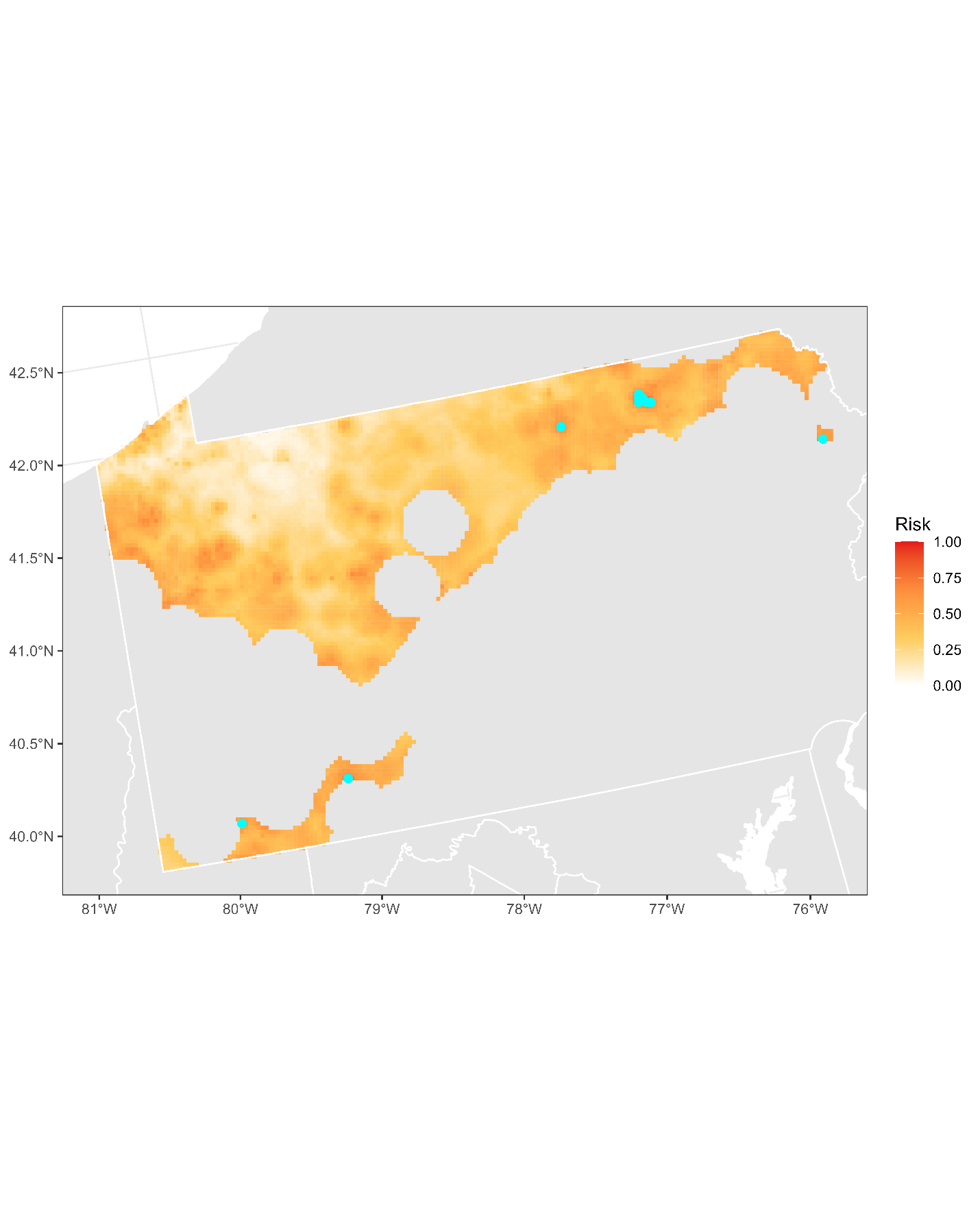

**Figure S12.1:** Projected risk of colonization by *L. delicatula* jump dispersal for Pennsylvania. Risk projections are based on an ensemble random forest model including distances from existing *L. delicatula* populations. Note that areas falling less than 20 km from existing *L. delicatula* populations are not included in the map because any colonization there would be categorized as diffusive spread based on our methodology (regardless of the actual source of the resulting population). Darker, redder colors indicate increased risk of colonization by jump dispersal. Cyan points show the top ten highest risk locations in the state.

**Table S12.1:** Ranked list of Pennsylvania county subdivisions sorted by the maximum projected risk of colonization by jump dispersal occurring there. For each subdivision, we identified the 2.5 x 2.5 km grid cell centroid included there with the highest projected risk of colonization by jump dispersal and report that location and risk value.

| **County, subdivision** | **Median risk value** | **Maximum risk value** | **Latitude of maximum risk** | **Longitude of maximum risk** |
| --- | --- | --- | --- | --- |
| Bradford County, Wysox township | 0.61350188 | 0.77434104 | 41.75757344 | -76.42116744 |
| Bradford County, North Towanda township | 0.6845078 | 0.7493276 | 41.78857444 | -76.47428311 |
| Bradford County, Towanda borough | 0.69956966 | 0.7415277 | 41.78401252 | -76.44462482 |
| Somerset County, Somerset township | 0.61309844 | 0.73964497 | 40.05212263 | -79.09048474 |
| Bradford County, Asylum township | 0.53052717 | 0.70790748 | 41.75299946 | -76.39152573 |
| Tioga County, Covington township | 0.57840237 | 0.70683163 | 41.72064645 | -77.08051268 |
| Pike County, Palmyra township | 0.59803658 | 0.7027972 | 41.35855398 | -75.20911302 |
| Bradford County, Sheshequin township | 0.47660032 | 0.6993007 | 41.8104481 | -76.4680999 |
| Bradford County, Towanda township | 0.61767079 | 0.69526627 | 41.766701 | -76.48046246 |
| Fayette County, German township | 0.54895105 | 0.67105971 | 39.91061649 | -79.90299011 |
| Crawford County, West Mead township | 0.53657881 | 0.66729424 | 41.61769829 | -80.15158018 |
| Clearfield County, Sandy township | 0.53886498 | 0.66568047 | 41.14622621 | -78.7952465 |
| Crawford County, Vernon township | 0.58041958 | 0.66218397 | 41.62879198 | -80.24100771 |
| Mercer County, Springfield township | 0.62775686 | 0.66003228 | 41.16608319 | -80.16243703 |
| Somerset County, Lincoln township | 0.57853685 | 0.65895643 | 40.07408308 | -79.08525038 |
| Venango County, Franklin city | 0.58848843 | 0.65545992 | 41.3965972 | -79.83505847 |
| Mercer County, Sugar Grove township | 0.58337816 | 0.64981173 | 41.46353772 | -80.37006364 |
| Crawford County, West Fallowfield township | 0.52689618 | 0.64685315 | 41.48557242 | -80.36512446 |
| Venango County, Oil City city | 0.63986014 | 0.63986014 | 41.42543403 | -79.70602879 |
| Fayette County, South Union township | 0.58687466 | 0.63905325 | 39.86594417 | -79.73356584 |
| Bradford County, Rome township | 0.49004841 | 0.6366326 | 41.83588921 | -76.33700569 |
| Bradford County, Athens township | 0.51183432 | 0.63636364 | 41.98179918 | -76.54366505 |
| Mercer County, Otter Creek township | 0.57261969 | 0.63555675 | 41.38276694 | -80.26599346 |
| Bradford County, Ulster township | 0.45508338 | 0.63555675 | 41.81500474 | -76.49777091 |
| Venango County, Sugarcreek borough | 0.47135557 | 0.63313609 | 41.40723256 | -79.74086171 |
| Fayette County, Luzerne township | 0.60314685 | 0.6309844 | 39.93260843 | -79.8980146 |
| Clearfield County, Cooper township | 0.38743948 | 0.63044648 | 40.99032625 | -78.16315838 |
| Venango County, Sandycreek township | 0.60879505 | 0.62641205 | 41.34879169 | -79.81557795 |
| Somerset County, Jefferson township | 0.51291017 | 0.61753631 | 40.08197065 | -79.14346928 |
| Fayette County, Georges township | 0.47296934 | 0.61726735 | 39.86974103 | -79.76262431 |
| Centre County, Snow Shoe township | 0.40667025 | 0.61726735 | 41.02677381 | -77.9402646 |
| Mercer County, Pine township | 0.54679935 | 0.61565358 | 41.16237565 | -80.13282399 |
| Clarion County, Clarion township | 0.46557289 | 0.61484669 | 41.19550187 | -79.33259767 |
| Venango County, Barkeyville borough | 0.61377084 | 0.61377084 | 41.1877617 | -79.97472865 |
| Venango County, Frenchcreek township | 0.44795589 | 0.61350188 | 41.37836122 | -79.86984376 |
| Indiana County, Rayne township | 0.53173749 | 0.61161915 | 40.69082351 | -79.11880538 |
| Mercer County, Perry township | 0.47982786 | 0.61108123 | 41.40479588 | -80.26102739 |
| Pike County, Blooming Grove township | 0.59642281 | 0.60623991 | 41.35370344 | -75.17972015 |
| Venango County, Cranberry township | 0.52582033 | 0.60570199 | 41.35560191 | -79.69177143 |
| Venango County, Irwin township | 0.48601399 | 0.60435718 | 41.20602295 | -79.94006431 |
| Crawford County, Woodcock township | 0.35086068 | 0.60301237 | 41.68749676 | -80.16634831 |
| Tioga County, Blossburg borough | 0.56239914 | 0.60274341 | 41.69432551 | -77.05683742 |
| Wayne County, Paupack township | 0.59655729 | 0.59655729 | 41.42880719 | -75.21891128 |
| Bradford County, Sayre borough | 0.59628833 | 0.59628833 | 41.97725223 | -76.51392026 |
| Crawford County, South Shenango township | 0.5154653 | 0.59467456 | 41.56990167 | -80.49926149 |
| Mercer County, Greene township | 0.53093061 | 0.59359871 | 41.4671913 | -80.3998158 |
| Bradford County, Standing Stone township | 0.52259279 | 0.59144701 | 41.74841868 | -76.36188789 |
| Susquehanna County, Franklin township | 0.46880043 | 0.59117805 | 41.8754955 | -75.86052335 |
| Venango County, Rockland township | 0.37412587 | 0.59064013 | 41.32299414 | -79.79101682 |
| Susquehanna County, Bridgewater township | 0.49596557 | 0.58929532 | 41.85834501 | -75.8965516 |
| Greene County, Perry township | 0.58821947 | 0.58821947 | 39.72763781 | -80.06368248 |
| Tioga County, Richmond township | 0.49327595 | 0.58472297 | 41.76886619 | -77.09822285 |
| Crawford County, North Shenango township | 0.54088219 | 0.58364712 | 41.59194367 | -80.49435536 |
| Wyoming County, Washington township | 0.58337816 | 0.58337816 | 41.63988424 | -75.96003163 |
| Wayne County, Honesdale borough | 0.58284024 | 0.58284024 | 41.59597295 | -75.26151441 |
| Bradford County, Athens borough | 0.57772996 | 0.57772996 | 41.95537224 | -76.52010388 |
| Clearfield County, Huston township | 0.50228618 | 0.577461 | 41.20573826 | -78.56671067 |
| Somerset County, Jenner township | 0.53456159 | 0.57692308 | 40.1140477 | -79.04565013 |
| Bradford County, Orwell township | 0.51008607 | 0.57611619 | 41.85316212 | -76.30109723 |
| Mercer County, Findley township | 0.51855836 | 0.57584723 | 41.18810373 | -80.15745391 |
| Greene County, Dunkard township | 0.50537924 | 0.5718128 | 39.73092654 | -79.91367133 |
| Fayette County, Springhill township | 0.51559978 | 0.56401291 | 39.76737305 | -79.84569646 |
| Somerset County, Quemahoning township | 0.52313072 | 0.56132329 | 40.11008461 | -79.01653145 |
| Tioga County, Liberty township | 0.48009683 | 0.56132329 | 41.56734805 | -77.12232429 |
| Crawford County, Meadville city | 0.54357181 | 0.5591716 | 41.63972796 | -80.1465616 |
| Mercer County, Grove City borough | 0.55890264 | 0.55890264 | 41.15866125 | -80.10321406 |
| Susquehanna County, Springville township | 0.51855836 | 0.55890264 | 41.6570484 | -75.92413099 |
| Indiana County, Cherryhill township | 0.45857988 | 0.55863367 | 40.67896034 | -79.03073828 |
| Crawford County, Sadsbury township | 0.49919311 | 0.55648198 | 41.61044176 | -80.27580224 |
| Wayne County, Damascus township | 0.51102743 | 0.55648198 | 41.73406772 | -75.12696734 |
| Erie County, Millcreek township | 0.38717052 | 0.55594406 | 42.09156437 | -80.13552825 |
| Wayne County, Mount Pleasant township | 0.52662722 | 0.55594406 | 41.7824084 | -75.42263786 |
| Wayne County, Oregon township | 0.5260893 | 0.55406132 | 41.63959154 | -75.24842369 |
| Somerset County, Upper Turkeyfoot township | 0.51976869 | 0.55379236 | 39.95167968 | -79.35429316 |
| Clarion County, Highland township | 0.43625605 | 0.55379236 | 41.22138402 | -79.35694672 |
| Clearfield County, Greenwood township | 0.51008607 | 0.55271651 | 40.92437957 | -78.66710936 |
| Clearfield County, DuBois city | 0.55177515 | 0.55244755 | 41.11620996 | -78.74161171 |
| Wayne County, Preston township | 0.50457235 | 0.55137171 | 41.82605203 | -75.40961267 |
| Clearfield County, Bigler township | 0.52541689 | 0.55056482 | 40.83003177 | -78.47762339 |
| Bradford County, Overton township | 0.47740721 | 0.54868209 | 41.62720142 | -76.61239597 |
| Wayne County, Dyberry township | 0.52662722 | 0.54868209 | 41.61778216 | -75.25497108 |
| Bradford County, Monroe township | 0.44459387 | 0.54760624 | 41.71383336 | -76.43354994 |
| Fayette County, Nicholson township | 0.45481442 | 0.54760624 | 39.78936136 | -79.84071335 |
| Clearfield County, Penn township | 0.49623453 | 0.54760624 | 40.94228059 | -78.63224035 |
| Lycoming County, Jackson township | 0.47095212 | 0.54733728 | 41.54544915 | -77.1282825 |
| Susquehanna County, New Milford township | 0.44257665 | 0.54706832 | 41.90031991 | -75.72908964 |
| Tioga County, Ward township | 0.4351802 | 0.54626143 | 41.73369342 | -77.01517381 |
| Bradford County, LeRaysville borough | 0.54545455 | 0.54545455 | 41.83471292 | -76.18242346 |
| Mercer County, Salem township | 0.53133405 | 0.54518558 | 41.43050497 | -80.28578895 |
| Elk County, St. Marys city | 0.37237762 | 0.54276493 | 41.42939107 | -78.54136197 |
| Jefferson County, Washington township | 0.3655191 | 0.54249597 | 41.16819078 | -78.78983926 |
| Somerset County, Stonycreek township | 0.51358257 | 0.542227 | 40.03229247 | -78.94503679 |
| Tioga County, Charleston township | 0.46987628 | 0.542227 | 41.74703506 | -77.25860149 |
| Fayette County, Springfield township | 0.52824099 | 0.54142012 | 39.97364995 | -79.34914683 |
| Butler County, Marion township | 0.46073158 | 0.54115116 | 41.13998657 | -79.95521125 |
| Venango County, Cornplanter township | 0.31764389 | 0.54088219 | 41.4474418 | -79.70088458 |
| Venango County, Victory township | 0.45615923 | 0.54034427 | 41.31232282 | -79.88509347 |
| Butler County, Mercer township | 0.48601399 | 0.53899946 | 41.14747698 | -80.01440299 |
| Bradford County, Pike township | 0.51183432 | 0.53792361 | 41.86119765 | -76.20581646 |
| Crawford County, Hayfield township | 0.48251748 | 0.53765465 | 41.69489313 | -80.22602438 |
| Crawford County, Union township | 0.46745562 | 0.53738569 | 41.59566894 | -80.15659561 |
| Susquehanna County, Auburn township | 0.3781603 | 0.53711673 | 41.66640365 | -75.98327576 |
| Fayette County, Menallen township | 0.53604088 | 0.53604088 | 39.92507041 | -79.81075067 |
| Bradford County, Wyalusing township | 0.45710059 | 0.53496503 | 41.71277311 | -76.27923497 |
| Cambria County, Northern Cambria borough | 0.5126412 | 0.53442711 | 40.6430052 | -78.76671423 |
| Tioga County, Bloss township | 0.44069392 | 0.53415815 | 41.67684578 | -77.09247749 |
| Susquehanna County, Great Bend township | 0.4725659 | 0.53388919 | 41.93926301 | -75.68656005 |
| Somerset County, Shade township | 0.48776224 | 0.53362023 | 40.06020575 | -78.81810955 |
| Fayette County, Wharton township | 0.44055944 | 0.53335126 | 39.83253215 | -79.65145588 |
| Wyoming County, Meshoppen township | 0.45293168 | 0.53281334 | 41.64455691 | -75.98959721 |
| Mercer County, Jackson township | 0.46288327 | 0.53254438 | 41.23584988 | -80.17712143 |
| Crawford County, Conneaut township | 0.48951049 | 0.53254438 | 41.66170043 | -80.50945878 |
| Bradford County, Herrick township | 0.45831092 | 0.53254438 | 41.80942903 | -76.31357158 |
| Wayne County, Lebanon township | 0.49462076 | 0.52985476 | 41.69288273 | -75.29439027 |
| Clearfield County, Ferguson township | 0.49919311 | 0.5295858 | 40.89431511 | -78.61369662 |
| Somerset County, Middlecreek township | 0.5 | 0.52824099 | 39.96975728 | -79.32006846 |
| Crawford County, Greenwood township | 0.3922808 | 0.52635826 | 41.58472612 | -80.25098093 |
| Tioga County, Delmar township | 0.42119419 | 0.52555137 | 41.7294985 | -77.29421797 |
| Wayne County, Manchester township | 0.43302851 | 0.52528241 | 41.79948482 | -75.10716083 |
| Somerset County, Lower Turkeyfoot township | 0.43477676 | 0.52474449 | 39.92970943 | -79.35943635 |
| Jefferson County, Pine Creek township | 0.35180204 | 0.52420656 | 41.14427234 | -78.94830572 |
| Susquehanna County, Montrose borough | 0.52420656 | 0.52420656 | 41.83180441 | -75.87327662 |
| Mercer County, Fairview township | 0.49623453 | 0.52366864 | 41.3570552 | -80.24125589 |
| Clearfield County, Jordan township | 0.46503497 | 0.52205487 | 40.83412118 | -78.50700696 |
| Crawford County, Pine township | 0.44566971 | 0.52071006 | 41.65807155 | -80.47961848 |
| Clearfield County, Brady township | 0.40102205 | 0.51990317 | 41.08617198 | -78.68802245 |
| Bradford County, Ridgebury township | 0.38811189 | 0.51990317 | 41.9999187 | -76.66268283 |
| Elk County, Johnsonburg borough | 0.51990317 | 0.51990317 | 41.49363932 | -78.67865293 |
| Armstrong County, Redbank township | 0.40909091 | 0.51963421 | 41.00791701 | -79.28591202 |
| Tioga County, Wellsboro borough | 0.51748252 | 0.51963421 | 41.75577668 | -77.31799414 |
| Bradford County, Leroy township | 0.47928994 | 0.51855836 | 41.6233685 | -76.73691729 |
| Armstrong County, Mahoning township | 0.47471759 | 0.51775148 | 40.99373163 | -79.35014461 |
| Erie County, Erie city | 0.39268424 | 0.51721356 | 42.12819948 | -80.06536716 |
| Wayne County, Buckingham township | 0.47122109 | 0.51694459 | 41.83342841 | -75.31427115 |
| Mercer County, Coolspring township | 0.51183432 | 0.51640667 | 41.26157111 | -80.20179314 |
| Clearfield County, Morris township | 0.51304465 | 0.51506186 | 40.94645782 | -78.17433053 |
| Fayette County, Henry Clay township | 0.4725659 | 0.51398601 | 39.76168832 | -79.45846309 |
| Tioga County, Sullivan township | 0.45965573 | 0.51237224 | 41.75559137 | -77.00916427 |
| Clearfield County, Union township | 0.48413125 | 0.51210328 | 41.1040811 | -78.65307897 |
| Susquehanna County, Oakland township | 0.45051103 | 0.51183432 | 41.95635638 | -75.65043871 |
| Mercer County, Lake township | 0.45131791 | 0.51075847 | 41.3239413 | -80.15716891 |
| Clearfield County, Beccaria township | 0.46422808 | 0.50941366 | 40.81217466 | -78.4592134 |
| Cambria County, Barr township | 0.45320065 | 0.50833782 | 40.62105065 | -78.77208872 |
| Crawford County, East Fallowfield township | 0.41581495 | 0.50672405 | 41.50394449 | -80.33041499 |
| Bradford County, Albany township | 0.39967725 | 0.50618612 | 41.62635574 | -76.45826867 |
| Cambria County, Susquehanna township | 0.41984938 | 0.50618612 | 40.66495986 | -78.76133643 |
| Crawford County, Summit township | 0.482383 | 0.50511027 | 41.68389814 | -80.32053227 |
| Susquehanna County, Middletown township | 0.45938677 | 0.50511027 | 41.89566166 | -76.1338912 |
| Somerset County, Allegheny township | 0.47565896 | 0.50457235 | 40.03022139 | -78.7652768 |
| Venango County, Scrubgrass township | 0.39456697 | 0.50430339 | 41.18336509 | -79.76240364 |
| Clearfield County, Knox township | 0.49865519 | 0.50376547 | 40.85197709 | -78.47214198 |
| Bradford County, Terry township | 0.46906939 | 0.5034965 | 41.64718698 | -76.29792368 |
| Sullivan County, Fox township | 0.46422808 | 0.50295858 | 41.57510686 | -76.71949175 |
| Bradford County, South Creek township | 0.4140667 | 0.50053792 | 42.00051911 | -76.81763598 |
| Susquehanna County, Jessup township | 0.42119419 | 0.49973104 | 41.81933641 | -75.93891523 |
| Fayette County, Stewart township | 0.46463152 | 0.49919311 | 39.83147032 | -79.4721784 |
| Tioga County, Hamilton township | 0.45589026 | 0.49838623 | 41.68989822 | -77.02718162 |
| Venango County, Mineral township | 0.41877353 | 0.49784831 | 41.34186189 | -79.93936034 |
| Clearfield County, Newburg borough | 0.47902098 | 0.49757934 | 40.8585153 | -78.68337925 |
| Bedford County, Napier township | 0.47444863 | 0.49731038 | 40.10993545 | -78.68572438 |
| Clarion County, Porter township | 0.46073158 | 0.49677246 | 40.99761986 | -79.3796452 |
| Susquehanna County, Thompson township | 0.45629371 | 0.49677246 | 41.86704027 | -75.52158131 |
| Bedford County, Juniata township | 0.44983862 | 0.49596557 | 40.05216796 | -78.7599445 |
| Susquehanna County, Apolacon township | 0.46288327 | 0.49569661 | 41.93008689 | -76.06189512 |
| Crawford County, Beaver township | 0.39322216 | 0.49515869 | 41.76464926 | -80.42511238 |
| Crawford County, Summerhill township | 0.4521248 | 0.49462076 | 41.7132632 | -80.3752694 |
| Susquehanna County, Dimock township | 0.45750403 | 0.4943518 | 41.76626938 | -75.89237672 |
| Somerset County, Indian Lake borough | 0.49408284 | 0.49408284 | 40.0422642 | -78.85250346 |
| Clarion County, Redbank township | 0.43302851 | 0.49381388 | 41.03380389 | -79.31017948 |
| Crawford County, West Shenango township | 0.47727273 | 0.49354492 | 41.52219014 | -80.47928189 |
| Bradford County, Warren township | 0.47417967 | 0.49327595 | 41.90029564 | -76.16357641 |
| Bradford County, Canton township | 0.45320065 | 0.49300699 | 41.61494306 | -76.83181303 |
| Mercer County, Worth township | 0.39887036 | 0.49220011 | 41.29077327 | -80.07316093 |
| Butler County, Cherry township | 0.45938677 | 0.49193115 | 41.06644185 | -79.91120437 |
| Lycoming County, Pine township | 0.45723507 | 0.49193115 | 41.47540477 | -77.27036383 |
| Mercer County, Jackson Center borough | 0.49166218 | 0.49166218 | 41.27618825 | -80.13749034 |
| Tioga County, Morris township | 0.39671867 | 0.49139322 | 41.56739784 | -77.27632556 |
| Westmoreland County, Cook township | 0.47740721 | 0.4908553 | 40.13376233 | -79.19129172 |
| Cambria County, West Carroll township | 0.46826251 | 0.49058634 | 40.61702311 | -78.74277797 |
| Clearfield County, Woodward township | 0.49031737 | 0.49031737 | 40.83966425 | -78.38398533 |
| Sullivan County, Elkland township | 0.48009683 | 0.49031737 | 41.5660912 | -76.66032745 |
| Elk County, Fox township | 0.38555675 | 0.48897257 | 41.24965044 | -78.5557415 |
| Bradford County, Granville township | 0.464766 | 0.48897257 | 41.70639235 | -76.68290631 |
| Jefferson County, Brookville borough | 0.46664874 | 0.4887036 | 41.16411246 | -79.09604432 |
| Susquehanna County, Forest Lake township | 0.42065627 | 0.48762776 | 41.86771495 | -75.95586276 |
| Bradford County, Franklin township | 0.44513179 | 0.48708983 | 41.65360195 | -76.63588174 |
| Venango County, Clintonville borough | 0.48682087 | 0.48682087 | 41.19849771 | -79.88083143 |
| Bradford County, Tuscarora township | 0.3992738 | 0.48601399 | 41.67707589 | -76.19666606 |
| Susquehanna County, Ararat township | 0.47485207 | 0.48601399 | 41.81860522 | -75.50493402 |
| Clearfield County, Bell township | 0.42159763 | 0.48574502 | 40.86255726 | -78.7127864 |
| Lycoming County, Cascade township | 0.35435718 | 0.48547606 | 41.47510682 | -76.96276456 |
| Tioga County, Tioga township | 0.35664336 | 0.4852071 | 41.90911956 | -77.12174517 |
| Mercer County, Wolf Creek township | 0.4373319 | 0.48493814 | 41.19897393 | -80.0635841 |
| Mercer County, French Creek township | 0.43491124 | 0.48413125 | 41.44864586 | -80.06768326 |
| Indiana County, Green township | 0.37694997 | 0.48359333 | 40.69695743 | -78.99608137 |
| Wayne County, Scott township | 0.42926304 | 0.48278644 | 41.99801613 | -75.48259501 |
| Crawford County, Cussewago township | 0.33176439 | 0.48224852 | 41.76468874 | -80.24088289 |
| McKean County, Bradford township | 0.26291017 | 0.48197956 | 41.94522297 | -78.65816031 |
| Clarion County, Knox township | 0.42159763 | 0.48090371 | 41.29512866 | -79.40047164 |
| Susquehanna County, Choconut township | 0.48090371 | 0.48090371 | 41.92542936 | -76.03220403 |
| Clarion County, Toby township | 0.42052179 | 0.48009683 | 41.07907883 | -79.48213327 |
| Clarion County, Sligo borough | 0.47928994 | 0.47928994 | 41.10492985 | -79.50649692 |
| Crawford County, Saegertown borough | 0.47875202 | 0.47875202 | 41.70581882 | -80.13148696 |
| Bradford County, Troy township | 0.44432491 | 0.47848306 | 41.72437824 | -76.80149257 |
| Elk County, Ridgway township | 0.33808499 | 0.47848306 | 41.48958719 | -78.64898392 |
| Crawford County, Oil Creek township | 0.23587951 | 0.47767617 | 41.62733268 | -79.68939699 |
| Bradford County, Burlington township | 0.43786982 | 0.47767617 | 41.75392509 | -76.54593068 |
| Clarion County, Monroe township | 0.46261431 | 0.4760624 | 41.1411862 | -79.4369907 |
| Clarion County, Limestone township | 0.36982249 | 0.4760624 | 41.15152668 | -79.34307545 |
| Venango County, Polk borough | 0.47498655 | 0.47498655 | 41.3638772 | -79.9342951 |
| Susquehanna County, Rush township | 0.38152232 | 0.47444863 | 41.82544752 | -76.12311006 |
| Mercer County, New Lebanon borough | 0.47310382 | 0.47310382 | 41.42662352 | -80.07271289 |
| Lycoming County, McIntyre township | 0.42186659 | 0.4725659 | 41.50143912 | -76.98633086 |
| Clarion County, Paint township | 0.44190425 | 0.47229693 | 41.27313621 | -79.40569841 |
| Jefferson County, Winslow township | 0.34803658 | 0.47175901 | 41.12827756 | -78.83017476 |
| Tioga County, Union township | 0.4429801 | 0.46906939 | 41.65022962 | -76.91462969 |
| Armstrong County, Madison township | 0.42267348 | 0.46853147 | 40.97951408 | -79.41435145 |
| Clarion County, Piney township | 0.43612157 | 0.46799354 | 41.12306261 | -79.47175272 |
| Somerset County, Ursina borough | 0.46772458 | 0.46772458 | 39.81208342 | -79.32707199 |
| Mercer County, Sandy Lake township | 0.43813878 | 0.46745562 | 41.35310022 | -80.02840445 |
| Bradford County, Stevens township | 0.41312534 | 0.46718666 | 41.79099224 | -76.19496857 |
| Mercer County, Mill Creek township | 0.44096288 | 0.46557289 | 41.42289427 | -80.04299503 |
| Lycoming County, McNett township | 0.42980097 | 0.46288327 | 41.56669472 | -76.81432443 |
| Wayne County, Starrucca borough | 0.4373319 | 0.46207639 | 41.87929631 | -75.45582437 |
| Bradford County, Canton borough | 0.46153846 | 0.46153846 | 41.66319143 | -76.84932567 |
| Cambria County, East Carroll township | 0.46073158 | 0.46073158 | 40.63089839 | -78.67876575 |
| Lycoming County, Cogan House township | 0.4112426 | 0.46073158 | 41.4797537 | -77.14613486 |
| Susquehanna County, Silver Lake township | 0.45938677 | 0.45938677 | 41.90673123 | -75.91347928 |
| Tioga County, Rutland township | 0.31925767 | 0.45884884 | 41.83874106 | -76.95537427 |
| Elk County, Horton township | 0.30728886 | 0.45804196 | 41.21792804 | -78.65535605 |
| Butler County, Allegheny township | 0.41393222 | 0.457773 | 41.16136023 | -79.76750947 |
| Erie County, North East borough | 0.457773 | 0.457773 | 42.21207077 | -79.82955268 |
| Crawford County, East Mead township | 0.3598709 | 0.45750403 | 41.63229519 | -80.08694081 |
| Clinton County, West Keating township | 0.44499731 | 0.45723507 | 41.15324497 | -78.02984533 |
| Bradford County, Armenia township | 0.41231845 | 0.45642819 | 41.7507469 | -76.82508205 |
| Clearfield County, Karthaus township | 0.44835933 | 0.45589026 | 41.15743883 | -78.05933708 |
| Clinton County, Leidy township | 0.3302851 | 0.45589026 | 41.46030444 | -77.95048748 |
| Venango County, Clinton township | 0.39456697 | 0.45508338 | 41.24628666 | -79.90031465 |
| Fayette County, Masontown borough | 0.45508338 | 0.45508338 | 39.84464081 | -79.91789836 |
| Clearfield County, Burnside township | 0.35449166 | 0.45481442 | 40.84060147 | -78.7181942 |
| Bradford County, Springfield township | 0.39671867 | 0.45454545 | 41.80293905 | -76.7177963 |
| Clinton County, East Keating township | 0.45400753 | 0.45400753 | 41.25026159 | -77.91299595 |
| Crawford County, Spring township | 0.37345347 | 0.45373857 | 41.77938588 | -80.36039466 |
| Clearfield County, Chest township | 0.35879505 | 0.45266272 | 40.81056039 | -78.66482285 |
| Mercer County, New Vernon township | 0.39887036 | 0.45266272 | 41.37169344 | -80.17687415 |
| Centre County, Burnside township | 0.40182894 | 0.45185584 | 41.21062389 | -77.95387227 |
| Crawford County, Conneautville borough | 0.45158687 | 0.45158687 | 41.75734465 | -80.36535603 |
| Fayette County, Smithfield borough | 0.45158687 | 0.45158687 | 39.80756981 | -79.80668846 |
| Clarion County, Elk township | 0.387305 | 0.45131791 | 41.24075268 | -79.50495971 |
| Clarion County, Millcreek township | 0.32275417 | 0.45077999 | 41.23166635 | -79.26291313 |
| Centre County, Rush township | 0.40828402 | 0.44970414 | 40.97360841 | -78.04546597 |
| Clarion County, Madison township | 0.42442173 | 0.44916622 | 41.00924367 | -79.46816639 |
| Bradford County, Columbia township | 0.39483593 | 0.44862829 | 41.83830611 | -76.80077781 |
| Bedford County, West St. Clair township | 0.38461538 | 0.44809037 | 40.14977484 | -78.64588177 |
| Mercer County, Stoneboro borough | 0.44782141 | 0.44782141 | 41.34225424 | -80.12248493 |
| Susquehanna County, Brooklyn township | 0.43060785 | 0.44620764 | 41.77871297 | -75.82677955 |
| Butler County, Venango township | 0.39349112 | 0.44593868 | 41.1066917 | -79.87151674 |
| Sullivan County, Hillsgrove township | 0.42052179 | 0.44513179 | 41.44382859 | -76.75594077 |
| Bradford County, Litchfield township | 0.42442173 | 0.44486283 | 41.94626096 | -76.46064372 |
| Venango County, Pinegrove township | 0.29989242 | 0.44459387 | 41.38813373 | -79.59243428 |
| Clearfield County, Covington township | 0.42993545 | 0.44459387 | 41.1480427 | -78.15344005 |
| Bradford County, Troy borough | 0.44432491 | 0.44432491 | 41.79004205 | -76.78325485 |
| Susquehanna County, Harmony township | 0.41191501 | 0.44324906 | 41.91547581 | -75.53825165 |
| Somerset County, Addison township | 0.41527703 | 0.4429801 | 39.79011563 | -79.33221351 |
| Butler County, Washington township | 0.38017751 | 0.44244217 | 41.08468396 | -79.8765831 |
| Armstrong County, East Franklin township | 0.42200108 | 0.44136633 | 40.86685513 | -79.59275546 |
| Erie County, McKean township | 0.19903174 | 0.44136633 | 42.02915099 | -80.18066229 |
| Potter County, Wharton township | 0.37036041 | 0.4408284 | 41.48223879 | -77.94479256 |
| Clarion County, Richland township | 0.38703604 | 0.44055944 | 41.18629416 | -79.6092868 |
| Armstrong County, Wayne township | 0.32961269 | 0.44029048 | 40.9277782 | -79.36579534 |
| Mercer County, Deer Creek township | 0.37829478 | 0.44029048 | 41.43034589 | -80.1024339 |
| Clarion County, Beaver township | 0.40169446 | 0.43813878 | 41.21875706 | -79.5101499 |
| Indiana County, East Mahoning township | 0.32947821 | 0.43786982 | 40.77475224 | -79.06830111 |
| McKean County, Bradford city | 0.43786982 | 0.43786982 | 41.96313929 | -78.62277966 |
| Clinton County, Beech Creek township | 0.36229155 | 0.43625605 | 41.24603362 | -77.88347415 |
| Indiana County, Pine township | 0.40828402 | 0.4351802 | 40.60114953 | -78.95865676 |
| Tioga County, Duncan township | 0.42442173 | 0.4338354 | 41.68130785 | -77.27638311 |
| Potter County, Roulette township | 0.38058096 | 0.43195266 | 41.78330845 | -78.1438893 |
| Bradford County, Smithfield township | 0.36067778 | 0.4316837 | 41.8240976 | -76.55712448 |
| Clearfield County, Irvona borough | 0.43141474 | 0.43141474 | 40.77236064 | -78.55276525 |
| Bradford County, Wilmot township | 0.3577192 | 0.43114578 | 41.60346389 | -76.31036343 |
| Clearfield County, Girard township | 0.38596019 | 0.42980097 | 41.18660559 | -78.26587696 |
| Mercer County, Sandy Creek township | 0.35328133 | 0.42980097 | 41.4488546 | -80.25108594 |
| Cambria County, Elder township | 0.41204949 | 0.42953201 | 40.6568928 | -78.70268329 |
| Butler County, Eau Claire borough | 0.42953201 | 0.42953201 | 41.14314651 | -79.80219644 |
| Elk County, Jones township | 0.30957504 | 0.42926304 | 41.62366737 | -78.79988729 |
| Susquehanna County, Jackson township | 0.39698763 | 0.4281872 | 41.8547509 | -75.58731521 |
| Clarion County, Washington township | 0.29357181 | 0.42791824 | 41.32100022 | -79.42488 |
| Bradford County, Wells township | 0.37264658 | 0.42764927 | 41.9830997 | -76.85349192 |
| Clarion County, Licking township | 0.39187735 | 0.42657343 | 41.10878226 | -79.53605246 |
| Clearfield County, Coalport borough | 0.4260355 | 0.4260355 | 40.74633585 | -78.52886372 |
| Centre County, Curtin township | 0.39053254 | 0.42415277 | 41.11519486 | -77.76457782 |
| Lycoming County, Plunketts Creek township | 0.37654653 | 0.42361485 | 41.44831726 | -76.78547576 |
| Jefferson County, Snyder township | 0.31656805 | 0.42334588 | 41.20405799 | -78.71990504 |
| Butler County, Cherry Valley borough | 0.42307692 | 0.42307692 | 41.1651524 | -79.79710272 |
| Clinton County, Noyes township | 0.31979559 | 0.422539 | 41.26796054 | -77.8777757 |
| Clearfield County, New Washington borough | 0.422539 | 0.422539 | 40.81460649 | -78.69420908 |
| Clarion County, Salem township | 0.38999462 | 0.42227004 | 41.23029236 | -79.59896972 |
| Erie County, Harborcreek township | 0.24233459 | 0.42092523 | 42.16104785 | -79.96510403 |
| Bradford County, West Burlington township | 0.39039806 | 0.42038731 | 41.77654926 | -76.69422964 |
| Greene County, Monongahela township | 0.40788058 | 0.42011834 | 39.80065708 | -79.92782201 |
| Jefferson County, Rose township | 0.380312 | 0.41958042 | 41.16806002 | -79.12560197 |
| Jefferson County, Gaskill township | 0.36444325 | 0.41958042 | 40.92659048 | -78.84913466 |
| Cambria County, White township | 0.39483593 | 0.41877353 | 40.72438983 | -78.53431947 |
| Venango County, Canal township | 0.31656805 | 0.41877353 | 41.42992467 | -79.91908037 |
| Erie County, Fairview township | 0.32167832 | 0.41850457 | 42.01080893 | -80.21567207 |
| Bradford County, Windham township | 0.38972566 | 0.41769769 | 41.92336345 | -76.31206127 |
| Erie County, Lawrence Park township | 0.41742873 | 0.41742873 | 42.14650337 | -80.030259 |
| Tioga County, Middlebury township | 0.32033351 | 0.41608392 | 41.83028786 | -77.20511432 |
| Somerset County, Ogle township | 0.39806347 | 0.41473911 | 40.20175658 | -78.69338887 |
| Cambria County, Chest township | 0.37143626 | 0.41366326 | 40.64879869 | -78.64404348 |
| Somerset County, Confluence borough | 0.41312534 | 0.41312534 | 39.81597421 | -79.35608701 |
| Tioga County, Jackson township | 0.31845078 | 0.41097364 | 41.97458468 | -76.94886184 |
| Armstrong County, Perry township | 0.35368478 | 0.41016676 | 41.01412867 | -79.67995075 |
| Clarion County, Ashland township | 0.34279182 | 0.40882195 | 41.2522918 | -79.59380636 |
| Armstrong County, Sugarcreek township | 0.37412587 | 0.40640129 | 40.91467446 | -79.61195335 |
| Armstrong County, Washington township | 0.35381926 | 0.40640129 | 40.90315138 | -79.52352668 |
| Clinton County, Grugan township | 0.32060247 | 0.40559441 | 41.25150488 | -77.60637869 |
| Crawford County, Titusville city | 0.40532544 | 0.40532544 | 41.62351386 | -79.65961481 |
| McKean County, Wetmore township | 0.23641743 | 0.40478752 | 41.64161619 | -78.76470482 |
| Susquehanna County, Lanesboro borough | 0.40357719 | 0.40398063 | 41.96867933 | -75.58462038 |
| Jefferson County, Punxsutawney borough | 0.40088757 | 0.40398063 | 40.9425652 | -78.96693374 |
| Tioga County, Shippen township | 0.36282948 | 0.40344271 | 41.74692126 | -77.41299718 |
| Crawford County, Fairfield township | 0.34023669 | 0.40317375 | 41.51471456 | -80.05257548 |
| Potter County, Pike township | 0.37627757 | 0.40317375 | 41.73758313 | -77.66235065 |
| Indiana County, Banks township | 0.30903712 | 0.40290479 | 40.90061847 | -78.82507247 |
| Jefferson County, Bell township | 0.35825713 | 0.40236686 | 40.93858171 | -78.93747899 |
| McKean County, Foster township | 0.24744486 | 0.39806347 | 41.97125287 | -78.68252011 |
| Venango County, Jackson township | 0.32409898 | 0.39752555 | 41.46641605 | -79.84946629 |
| Indiana County, Washington township | 0.39698763 | 0.39698763 | 40.76461701 | -79.16175808 |
| Susquehanna County, Harford township | 0.39618074 | 0.39618074 | 41.81296221 | -75.75475322 |
| Venango County, President township | 0.18881119 | 0.39564282 | 41.43597485 | -79.61177412 |
| Clarion County, Farmington township | 0.32853685 | 0.39456697 | 41.37141147 | -79.29055578 |
| Jefferson County, Knox township | 0.2654653 | 0.39456697 | 41.13818205 | -79.07181061 |
| Potter County, Galeton borough | 0.39241528 | 0.39241528 | 41.73329524 | -77.63263892 |
| Venango County, Richland township | 0.3302851 | 0.39241528 | 41.19776028 | -79.6980819 |
| Potter County, West Branch township | 0.31011296 | 0.39214632 | 41.71565564 | -77.66815462 |
| Jefferson County, Young township | 0.36444325 | 0.39160839 | 40.9245727 | -79.00171813 |
| Venango County, Utica borough | 0.39080151 | 0.39080151 | 41.43368265 | -79.94879462 |
| Elk County, Jay township | 0.28859602 | 0.39026358 | 41.26752589 | -78.52068352 |
| Butler County, Parker township | 0.35368478 | 0.3894567 | 41.09534697 | -79.78280792 |
| Jefferson County, Perry township | 0.30365788 | 0.38838085 | 40.95051178 | -79.02585314 |
| Crawford County, Cambridge township | 0.3429263 | 0.38811189 | 41.79022278 | -80.08146749 |
| Tioga County, Chatham township | 0.2978752 | 0.38784293 | 41.83023806 | -77.35969365 |
| Crawford County, Wayne township | 0.29868209 | 0.38622916 | 41.51098032 | -80.02282146 |
| Wyoming County, Braintrim township | 0.34965035 | 0.38461538 | 41.64596433 | -76.14375756 |
| Bradford County, Sylvania borough | 0.38461538 | 0.38461538 | 41.80347352 | -76.87231426 |
| Tioga County, Gaines township | 0.35462614 | 0.38407746 | 41.75092537 | -77.59710535 |
| Jefferson County, Eldred township | 0.26869285 | 0.3838085 | 41.2120179 | -79.11498635 |
| Cambria County, Blacklick township | 0.36484669 | 0.38273265 | 40.54525994 | -78.88140087 |
| Venango County, Oakland township | 0.30231307 | 0.3816568 | 41.4732597 | -79.72545609 |
| Clarion County, Foxburg borough | 0.38138784 | 0.38138784 | 41.14994271 | -79.67874883 |
| Indiana County, South Mahoning township | 0.34722969 | 0.37977407 | 40.78265096 | -79.1270926 |
| Bedford County, Lincoln township | 0.35462614 | 0.37762238 | 40.21154409 | -78.60060181 |
| Potter County, Sylvania township | 0.3359333 | 0.37762238 | 41.58261913 | -78.01096859 |
| Jefferson County, Warsaw township | 0.34185046 | 0.37600861 | 41.21417149 | -78.96178493 |
| Venango County, Cherrytree township | 0.28294782 | 0.3746638 | 41.60532139 | -79.69455797 |
| Jefferson County, Henderson township | 0.30446477 | 0.37412587 | 41.03640617 | -78.82223238 |
| Clarion County, Perry township | 0.33983324 | 0.37305003 | 41.0504766 | -79.61062263 |
| McKean County, Liberty township | 0.31777838 | 0.3725121 | 41.81775262 | -78.22755758 |
| Tioga County, Deerfield township | 0.30782679 | 0.37224314 | 41.92226172 | -77.36580314 |
| Potter County, Keating township | 0.28886498 | 0.37116729 | 41.73523461 | -78.12545762 |
| Greene County, Richhill township | 0.34185046 | 0.36686391 | 39.91118708 | -80.44142966 |
| Cameron County, Shippen township | 0.29854761 | 0.36632598 | 41.50213097 | -78.24680322 |
| Elk County, Ridgway borough | 0.36605702 | 0.36605702 | 41.43178177 | -78.72468917 |
| Clearfield County, Lawrence township | 0.34991931 | 0.36578806 | 41.21543913 | -78.47258649 |
| Bradford County, Burlington borough | 0.36525013 | 0.36525013 | 41.78487449 | -76.5990987 |
| Tioga County, Elk township | 0.32719204 | 0.36498117 | 41.6765474 | -77.55520123 |
| Erie County, Union City borough | 0.36471221 | 0.36471221 | 41.89611493 | -79.84167771 |
| Tioga County, Osceola township | 0.30164067 | 0.36363636 | 41.9441805 | -77.35988725 |
| Potter County, Portage township | 0.27272727 | 0.3633674 | 41.56906981 | -78.07597842 |
| Clearfield County, Bloom township | 0.36229155 | 0.36229155 | 41.01625197 | -78.67483789 |
| Crawford County, East Fairfield township | 0.30742335 | 0.36121571 | 41.56621461 | -80.10204156 |
| Elk County, Highland township | 0.23951049 | 0.35933297 | 41.58373365 | -78.84047856 |
| Wyoming County, Windham township | 0.32356105 | 0.35906401 | 41.62873912 | -76.17959874 |
| Greene County, Wayne township | 0.35825713 | 0.35825713 | 39.73153396 | -80.27173491 |
| Clinton County, Colebrook township | 0.35825713 | 0.35825713 | 41.22098911 | -77.55317372 |
| Indiana County, North Mahoning township | 0.26250672 | 0.35798817 | 40.90260368 | -79.00704117 |
| Armstrong County, Bradys Bend township | 0.35637439 | 0.3577192 | 41.00265184 | -79.59138883 |
| Crawford County, Venango township | 0.302851 | 0.3577192 | 41.81225377 | -80.07641451 |
| Lycoming County, Brown township | 0.29747176 | 0.35745024 | 41.48849525 | -77.35911663 |
| Potter County, Hector township | 0.3302851 | 0.35637439 | 41.86915355 | -77.62745052 |
| Potter County, Ulysses township | 0.29424422 | 0.35610543 | 41.81620274 | -77.73415448 |
| Tioga County, Farmington township | 0.30365788 | 0.35233997 | 41.91790453 | -77.33603131 |
| Elk County, Benezette township | 0.30513717 | 0.35180204 | 41.40478705 | -78.36360961 |
| McKean County, Norwich township | 0.31239914 | 0.35099516 | 41.64079552 | -78.42668129 |
| Cameron County, Lumber township | 0.3387574 | 0.35045724 | 41.52936798 | -78.1169481 |
| Tioga County, Elkland borough | 0.34991931 | 0.34991931 | 41.98365728 | -77.3182456 |
| Greene County, Gilmore township | 0.34185046 | 0.34965035 | 39.76819134 | -80.38306516 |
| Erie County, Summit township | 0.2880581 | 0.34938139 | 42.0437691 | -80.11564814 |
| Clarion County, Brady township | 0.34884346 | 0.34884346 | 40.98065722 | -79.59653475 |
| Crawford County, Randolph township | 0.32302313 | 0.34776762 | 41.59157176 | -79.94302633 |
| Armstrong County, Hovey township | 0.32638515 | 0.34615385 | 41.12794184 | -79.68387829 |
| Potter County, Eulalia township | 0.29854761 | 0.34561592 | 41.75300053 | -78.09004743 |
| Erie County, North East township | 0.18988704 | 0.345078 | 42.19003919 | -79.83471268 |
| Greene County, Jackson township | 0.34454008 | 0.34454008 | 39.79020065 | -80.37824624 |
| Jefferson County, Union township | 0.29558903 | 0.34373319 | 41.1939809 | -79.14986567 |
| Greene County, Aleppo township | 0.31387843 | 0.34346423 | 39.81585863 | -80.40249421 |
| Sullivan County, Colley township | 0.34346423 | 0.34346423 | 41.52868568 | -76.26991171 |
| Indiana County, Canoe township | 0.27461001 | 0.34265734 | 40.89862959 | -78.97760077 |
| Jefferson County, Ringgold township | 0.31065089 | 0.3415815 | 40.97418833 | -79.20269036 |
| Potter County, Pleasant Valley township | 0.31320602 | 0.34077461 | 41.85750188 | -78.18648188 |
| Clearfield County, Goshen township | 0.32517483 | 0.33996772 | 41.22507079 | -78.37843654 |
| Forest County, Hickory township | 0.17267348 | 0.33862292 | 41.54871665 | -79.43195588 |
| Cameron County, Portage township | 0.29679935 | 0.33808499 | 41.55548779 | -78.14096315 |
| Lycoming County, McHenry township | 0.29585799 | 0.33781603 | 41.46658717 | -77.36499448 |
| Potter County, Clara township | 0.27286175 | 0.33700914 | 41.82720927 | -78.13256455 |
| Crawford County, Troy township | 0.24757934 | 0.33620226 | 41.65387157 | -79.89796255 |
| McKean County, Sergeant township | 0.27945132 | 0.3359333 | 41.62293929 | -78.46194954 |
| Jefferson County, Big Run borough | 0.33431953 | 0.33431953 | 40.97452276 | -78.8678462 |
| Indiana County, Grant township | 0.30513717 | 0.33431953 | 40.76285329 | -78.98013844 |
| Lycoming County, Cummings township | 0.31334051 | 0.33324368 | 41.38779104 | -77.29395927 |
| Indiana County, West Mahoning township | 0.27514793 | 0.33243679 | 40.86037613 | -79.19954312 |
| Jefferson County, Polk township | 0.28792361 | 0.33189887 | 41.25013419 | -78.89189259 |
| McKean County, Annin township | 0.27555137 | 0.33162991 | 41.86581618 | -78.2460811 |
| Tioga County, Clymer township | 0.29464766 | 0.33136095 | 41.86485699 | -77.59768508 |
| Butler County, Bruin borough | 0.33082302 | 0.33082302 | 41.06574816 | -79.72879553 |
| Potter County, Homer township | 0.28617536 | 0.32920925 | 41.7268658 | -78.06598372 |
| Crawford County, Rockdale township | 0.23964497 | 0.32867133 | 41.80478706 | -80.01665438 |
| Clinton County, Gallagher township | 0.31495428 | 0.3267886 | 41.28672437 | -77.53576593 |
| Crawford County, Hydetown borough | 0.32625067 | 0.32625067 | 41.65315747 | -79.71402724 |
| Greene County, Freeport township | 0.29289941 | 0.32356105 | 39.77183771 | -80.41211772 |
| Jefferson County, McCalmont township | 0.29491662 | 0.32302313 | 41.00846983 | -78.95090837 |
| Jefferson County, Porter township | 0.29101668 | 0.32275417 | 40.95221009 | -79.20795481 |
| Erie County, Greene township | 0.17240452 | 0.32248521 | 42.0766531 | -80.01552989 |
| Erie County, Corry city | 0.27703066 | 0.32248521 | 41.91733641 | -79.65191221 |
| Crawford County, Steuben township | 0.20239376 | 0.32221625 | 41.67589209 | -79.8928618 |
| Jefferson County, Clover township | 0.28362023 | 0.32194728 | 41.12410296 | -79.1362044 |
| Crawford County, Richmond township | 0.24529317 | 0.31925767 | 41.67965707 | -79.92267879 |
| Venango County, Plum township | 0.29236148 | 0.31764389 | 41.55825457 | -79.85879985 |
| Jefferson County, Beaver township | 0.24502421 | 0.31710597 | 41.06603051 | -79.21111966 |
| Indiana County, Montgomery township | 0.28967187 | 0.31683701 | 40.74288613 | -78.8332665 |
| McKean County, Kane borough | 0.31656805 | 0.31656805 | 41.67163775 | -78.81875107 |
| Erie County, Girard borough | 0.3126681 | 0.31387843 | 42.00348695 | -80.34058628 |
| Potter County, Stewardson township | 0.2859064 | 0.31360947 | 41.4872405 | -77.82059576 |
| Clearfield County, Westover borough | 0.31280258 | 0.31280258 | 40.74470212 | -78.6810688 |
| Clinton County, Chapman township | 0.28496503 | 0.31280258 | 41.36484799 | -77.76052699 |
| Jefferson County, Oliver township | 0.24556213 | 0.31145777 | 40.99841895 | -79.04469057 |
| Warren County, Warren city | 0.27891339 | 0.31038193 | 41.82537015 | -79.11951107 |
| Tioga County, Westfield township | 0.27756859 | 0.30903712 | 41.88678489 | -77.59184667 |
| Potter County, Sweden township | 0.28617536 | 0.30876815 | 41.74970108 | -77.90590721 |
| Erie County, LeBoeuf township | 0.19795589 | 0.30849919 | 41.90742742 | -79.93141163 |
| Greene County, Springhill township | 0.2824099 | 0.30607854 | 39.77547736 | -80.44117322 |
| Erie County, Elk Creek township | 0.2049489 | 0.30554061 | 41.86020954 | -80.28068264 |
| Erie County, Union township | 0.20064551 | 0.30446477 | 41.89989268 | -79.87158579 |
| Tioga County, Nelson township | 0.26896181 | 0.30311996 | 41.97928881 | -77.28845059 |
| Potter County, Summit township | 0.27218935 | 0.30204411 | 41.7277624 | -77.91163602 |
| Potter County, Abbott township | 0.26613771 | 0.30150619 | 41.64559257 | -77.65586871 |
| McKean County, Keating township | 0.25416891 | 0.29935449 | 41.81092995 | -78.50683599 |
| Erie County, Conneaut township | 0.19150081 | 0.29854761 | 41.85282614 | -80.40532136 |
| Erie County, Girard township | 0.1873319 | 0.29639591 | 42.02186268 | -80.305613 |
| Potter County, Ulysses borough | 0.29518558 | 0.29612695 | 41.91247024 | -77.77054779 |
| Elk County, Spring Creek township | 0.21624529 | 0.29558903 | 41.44787835 | -78.84330552 |
| Potter County, Harrison township | 0.23628295 | 0.29558903 | 41.91301255 | -77.61578797 |
| Tioga County, Lawrence township | 0.23278644 | 0.29424422 | 41.95293422 | -77.10975895 |
| Potter County, Bingham township | 0.26169984 | 0.29316837 | 41.94291695 | -77.82437372 |
| Tioga County, Brookfield township | 0.25874126 | 0.29289941 | 41.93064155 | -77.58015886 |
| Crawford County, Blooming Valley borough | 0.29209252 | 0.29209252 | 41.6726212 | -80.04703401 |
| McKean County, Hamlin township | 0.23466918 | 0.29209252 | 41.70753822 | -78.74832965 |
| Elk County, Millstone township | 0.25497579 | 0.2880581 | 41.37992995 | -79.01310094 |
| Jefferson County, Heath township | 0.25605164 | 0.28617536 | 41.35397872 | -78.98882356 |
| Potter County, Coudersport borough | 0.27461001 | 0.2859064 | 41.76655956 | -78.02486974 |
| Potter County, Genesee township | 0.24731038 | 0.28483055 | 41.94716157 | -77.85418621 |
| Erie County, Albion borough | 0.28429263 | 0.28429263 | 41.89326042 | -80.36547248 |
| Potter County, Hebron township | 0.25968263 | 0.28375471 | 41.8407909 | -78.06732602 |
| Forest County, Green township | 0.22216245 | 0.28267886 | 41.43738147 | -79.27473591 |
| Potter County, Allegany township | 0.25013448 | 0.27972028 | 41.83322103 | -77.85318856 |
| Clinton County, Renovo borough | 0.27945132 | 0.27945132 | 41.33866214 | -77.73671799 |
| Forest County, Tionesta township | 0.18612157 | 0.27703066 | 41.39085947 | -79.43886577 |
| McKean County, Lafayette township | 0.24731038 | 0.27407208 | 41.76537392 | -78.67234604 |
| McKean County, Smethport borough | 0.27353416 | 0.27353416 | 41.80273666 | -78.44725269 |
| Warren County, Columbus township | 0.17791824 | 0.26815492 | 41.93168652 | -79.58689262 |
| Jefferson County, Barnett township | 0.24892415 | 0.26627219 | 41.33374552 | -79.17726946 |
| Erie County, Concord township | 0.1810113 | 0.2654653 | 41.9029538 | -79.7169051 |
| McKean County, Ceres township | 0.21839699 | 0.26412049 | 41.90973002 | -78.23480711 |
| McKean County, Eldred township | 0.20871436 | 0.26223776 | 41.88648177 | -78.39514088 |
| Forest County, Kingsley township | 0.18033889 | 0.26169984 | 41.45153438 | -79.21008327 |
| Forest County, Barnett township | 0.24825175 | 0.26062399 | 41.39969911 | -79.16135197 |
| Crawford County, Athens township | 0.21920387 | 0.26008607 | 41.71993406 | -79.88265069 |
| McKean County, Lewis Run borough | 0.25685853 | 0.25685853 | 41.87524049 | -78.64484374 |
| Warren County, Pleasant township | 0.09965035 | 0.25658956 | 41.83325265 | -79.17919063 |
| Erie County, Greenfield township | 0.13232921 | 0.25605164 | 42.14597722 | -79.8450229 |
| Erie County, Springfield township | 0.14483593 | 0.25443787 | 41.95557442 | -80.5053237 |
| Potter County, Austin borough | 0.25309306 | 0.25309306 | 41.63908647 | -78.08866995 |
| Forest County, Jenks township | 0.20091447 | 0.2528241 | 41.43577699 | -79.0913775 |
| Erie County, Washington township | 0.15976331 | 0.24986552 | 41.85631677 | -80.066299 |
| Potter County, Oswayo township | 0.2197418 | 0.24878967 | 41.92858586 | -78.04454758 |
| Potter County, Sharon township | 0.22081764 | 0.24878967 | 41.98505418 | -78.12266071 |
| Venango County, Oil Creek township | 0.17670791 | 0.24744486 | 41.59767945 | -79.63501496 |
| Potter County, Oswayo borough | 0.24636902 | 0.24636902 | 41.92438529 | -78.0147315 |
| Erie County, Waterford borough | 0.24341044 | 0.24341044 | 41.93696501 | -79.98615991 |
| Erie County, Wayne township | 0.19472835 | 0.24260355 | 41.92497311 | -79.71172918 |
| Warren County, Conewango township | 0.11780527 | 0.24125874 | 41.85524832 | -79.17385088 |
| Erie County, Waterford township | 0.13932222 | 0.23023131 | 41.98851418 | -80.03587594 |
| Erie County, Platea borough | 0.22579344 | 0.22861754 | 41.95939523 | -80.35055018 |
| Warren County, Glade township | 0.11309844 | 0.22807961 | 41.87330383 | -79.13864816 |
| Potter County, Shinglehouse borough | 0.21705218 | 0.21705218 | 41.97144229 | -78.18802308 |
| Erie County, Elgin borough | 0.21597633 | 0.21597633 | 41.90676048 | -79.74680935 |
| Erie County, Venango township | 0.09601937 | 0.2140936 | 42.00623913 | -79.81596467 |
| Warren County, Deerfield township | 0.13582571 | 0.20952125 | 41.70271832 | -79.39521358 |
| McKean County, Mount Jewett borough | 0.20952125 | 0.20952125 | 41.71737735 | -78.65355721 |
| McKean County, Otto township | 0.16487359 | 0.20710059 | 41.95091728 | -78.53319504 |
| Warren County, Pittsfield township | 0.1366326 | 0.20414201 | 41.84637547 | -79.45316868 |
| Forest County, Harmony township | 0.15250134 | 0.20172136 | 41.57071604 | -79.42671684 |
| McKean County, Hamilton township | 0.1253362 | 0.20172136 | 41.70163754 | -78.87284355 |
| Erie County, Franklin township | 0.14550834 | 0.19849381 | 41.91525177 | -80.17587543 |
| Warren County, Limestone township | 0.16299086 | 0.19634212 | 41.67683128 | -79.3706834 |
| Crawford County, Bloomfield township | 0.16218397 | 0.19607316 | 41.778436 | -79.80759467 |
| Erie County, Amity township | 0.15680473 | 0.19419042 | 41.98421357 | -79.82111376 |
| McKean County, Corydon township | 0.14254976 | 0.19419042 | 41.95545859 | -78.90259604 |
| Forest County, Howe township | 0.14671867 | 0.19176977 | 41.53161298 | -79.12942741 |
| Warren County, Southwest township | 0.14483593 | 0.18692846 | 41.63786331 | -79.59486881 |
| Warren County, Spring Creek township | 0.1344809 | 0.18612157 | 41.80622688 | -79.49351556 |
| Crawford County, Rome township | 0.1457773 | 0.18289403 | 41.69718411 | -79.70370721 |
| McKean County, Eldred borough | 0.18262507 | 0.18262507 | 41.952374 | -78.37836002 |
| Warren County, Triumph township | 0.12157074 | 0.1810113 | 41.68459626 | -79.43026461 |
| Venango County, Allegheny township | 0.13488435 | 0.17966649 | 41.61201613 | -79.57028775 |
| Venango County, Pleasantville borough | 0.17509414 | 0.17509414 | 41.59000999 | -79.57548491 |
| Warren County, Eldred township | 0.13582571 | 0.17401829 | 41.78807602 | -79.52859291 |
| Warren County, Elk township | 0.10597095 | 0.17213556 | 41.96344263 | -78.96236207 |
| Warren County, Mead township | 0.12022593 | 0.16729424 | 41.88949902 | -78.91888385 |
| Warren County, Sheffield township | 0.12963959 | 0.16218397 | 41.73560438 | -78.95676964 |
| Warren County, Watson township | 0.08337816 | 0.15814954 | 41.63922451 | -79.25686129 |
| Erie County, Edinboro borough | 0.15277031 | 0.15277031 | 41.88579541 | -80.12106415 |
| Warren County, Freehold township | 0.13609467 | 0.15277031 | 41.96028916 | -79.45677354 |
| Warren County, Brokenstraw township | 0.11941904 | 0.15250134 | 41.84893497 | -79.29858996 |
| Crawford County, Sparta township | 0.12452932 | 0.14819796 | 41.78142018 | -79.65317932 |
| Warren County, Sugar Grove township | 0.07127488 | 0.13959118 | 41.90463453 | -79.37761882 |
| Warren County, Cherry Grove township | 0.0887574 | 0.12264658 | 41.66353638 | -79.09738284 |
| Warren County, Youngsville borough | 0.11296396 | 0.11296396 | 41.85283831 | -79.32844813 |
| Warren County, Pine Grove township | 0.07342657 | 0.09736417 | 41.91729412 | -79.12793369 |
| Warren County, Farmington township | 0.05433029 | 0.06804734 | 41.91102468 | -79.25278755 |
