## Supplementary material for "Mapping long-distance jump dispersal risk by leveraging transport infrastructure data and sampling bias correction: a case study with the invasive spotted lanternfly (*Lycorma delicatula*)": S13 Rhode Island risk

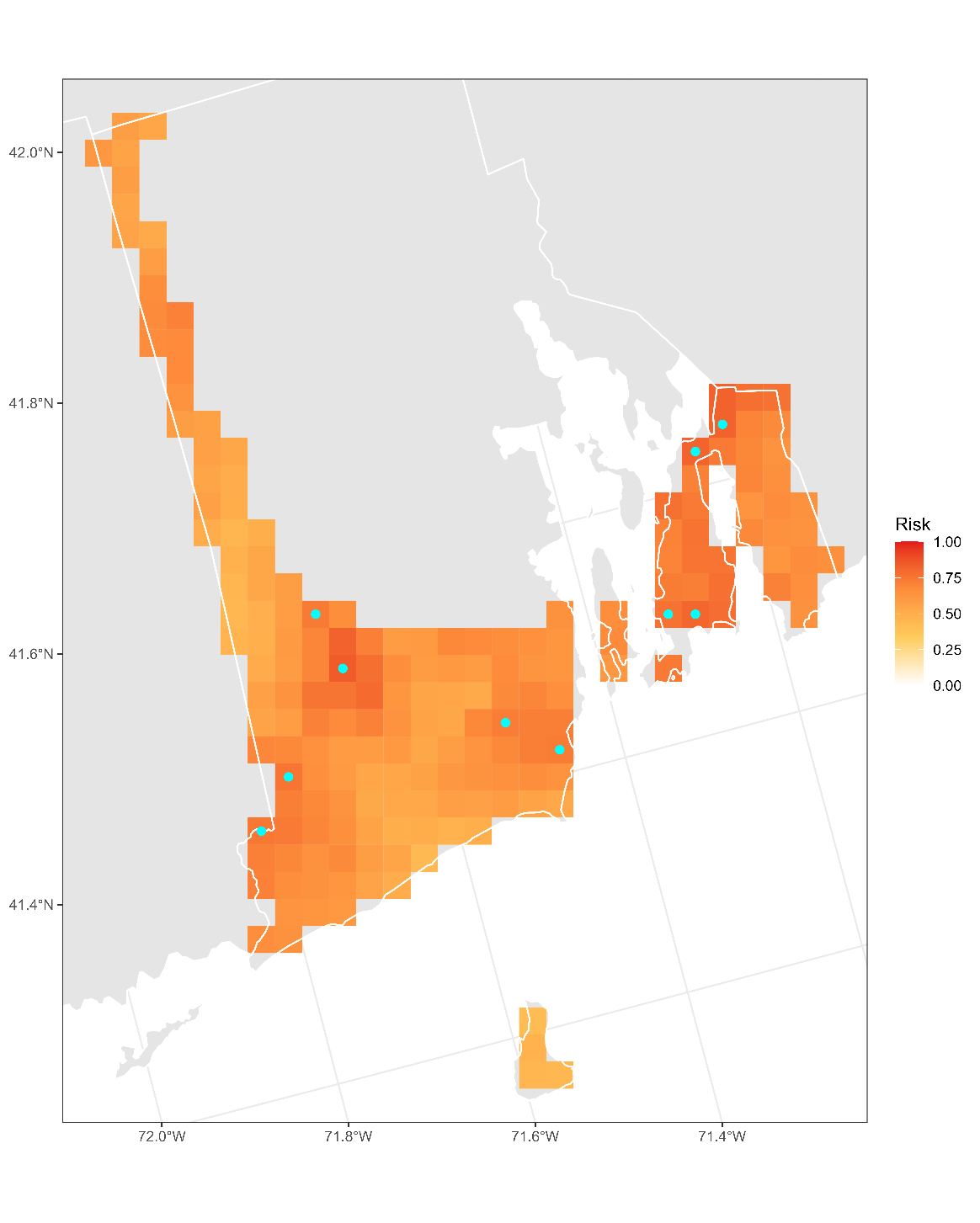

**Figure S13.1:** Projected risk of colonization by *L. delicatula* jump dispersal for Rhode Island. Risk projections are based on an ensemble random forest model including distances from existing *L. delicatula* populations. Note that areas falling less than 20 km from existing *L. delicatula* populations are not included in the map because any colonization there would be categorized as diffusive spread based on our methodology (regardless of the actual source of the resulting population). Darker, redder colors indicate increased risk of colonization by jump dispersal. Cyan points show the top ten highest risk locations in the state.

**Table S13.1:** Ranked list of Rhode Island county subdivisions sorted by the maximum projected risk of colonization by jump dispersal occurring there. For each subdivision, we identified the 2.5 x 2.5 km grid cell centroid included there with the highest projected risk of colonization by jump dispersal and report that location and risk value.

| **County, subdivision** | **Median risk value** | **Maximum risk value** | **Latitude of maximum risk** | **Longitude of maximum risk** |
| --- | --- | --- | --- | --- |
| Washington County, Richmond town | 0.6732114 | 0.84696073 | 41.53022147 | -71.67873504 |
| Newport County, Tiverton town | 0.69123185 | 0.82356105 | 41.64466494 | -71.20034063 |
| Newport County, Portsmouth town | 0.75860678 | 0.81871974 | 41.62883991 | -71.23734491 |
| Newport County, Middletown town | 0.75309306 | 0.80742335 | 41.49932652 | -71.28398559 |
| Newport County, Newport city | 0.7534965 | 0.76815492 | 41.50506725 | -71.31314618 |
| Washington County, Hopkinton town | 0.68289403 | 0.75874126 | 41.45506285 | -71.76761131 |
| Washington County, Westerly town | 0.66003228 | 0.74986552 | 41.41746257 | -71.8119783 |
| Washington County, Exeter town | 0.59736417 | 0.73507262 | 41.57909622 | -71.69266511 |
| Washington County, Narragansett town | 0.6366326 | 0.72969338 | 41.41998934 | -71.46836449 |
| Washington County, South Kingstown town | 0.61646046 | 0.72727273 | 41.45297825 | -71.51897501 |
| Providence County, Foster town | 0.67105971 | 0.71274879 | 41.8451063 | -71.75495734 |
| Newport County, Little Compton town | 0.64900484 | 0.70763852 | 41.5036447 | -71.18874597 |
| Newport County, Jamestown town | 0.66245293 | 0.67294244 | 41.4949377 | -71.37921342 |
| Providence County, Glocester town | 0.56603012 | 0.66379774 | 41.87236558 | -71.77665413 |
| Washington County, North Kingstown town | 0.64470145 | 0.66137708 | 41.51777439 | -71.49590792 |
| Washington County, Charlestown town | 0.53684777 | 0.6514255 | 41.40056219 | -71.72449851 |
| Providence County, Burrillville town | 0.56320602 | 0.61377084 | 41.99175866 | -71.7971543 |
| Kent County, West Greenwich town | 0.51425498 | 0.58230231 | 41.60635819 | -71.71425559 |
| Kent County, Coventry town | 0.50322754 | 0.56186122 | 41.68812693 | -71.77912766 |
| Washington County, New Shoreham town | 0.44701452 | 0.47525551 | 41.18812109 | -71.58176408 |
