## Supplementary material for "Mapping long-distance jump dispersal risk by leveraging transport infrastructure data and sampling bias correction: a case study with the invasive spotted lanternfly (*Lycorma delicatula*)": S14 South Carolina risk

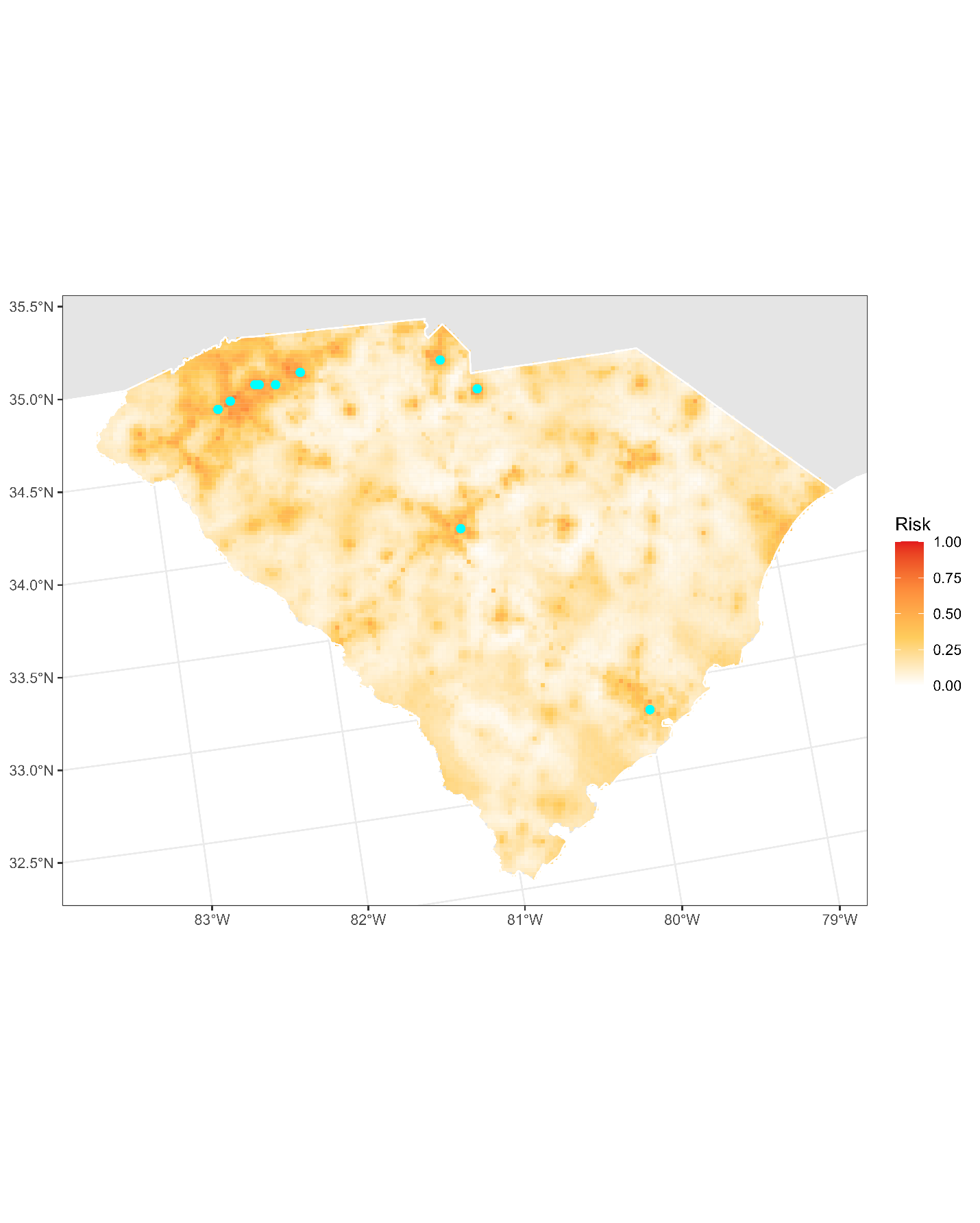

**Figure S14.1:** Projected risk of colonization by *L. delicatula* jump dispersal for South Carolina. Risk projections are based on an ensemble random forest model including distances from existing *L. delicatula* populations. Note that areas falling less than 20 km from existing *L. delicatula* populations are not included in the map because any colonization there would be categorized as diffusive spread based on our methodology (regardless of the actual source of the resulting population). Darker, redder colors indicate increased risk of colonization by jump dispersal. Cyan points show the top ten highest risk locations in the state.

**Table S14.1:** Ranked list of South Carolina county subdivisions sorted by the maximum projected risk of colonization by jump dispersal occurring there. For each subdivision, we identified the 2.5 x 2.5 km grid cell centroid included there with the highest projected risk of colonization by jump dispersal and report that location and risk value.

| **County, subdivision** | **Median risk value** | **Maximum risk value** | **Latitude of maximum risk** | **Longitude of maximum risk** |
| --- | --- | --- | --- | --- |
| Greenville County, Greenville CCD | 0.50416891 | 0.72942442 | 34.86626495 | -82.41798759 |
| Spartanburg County, Spartanburg CCD | 0.41433566 | 0.72431415 | 34.96595423 | -81.92592914 |
| Spartanburg County, Greer CCD | 0.53563744 | 0.70172136 | 34.93227729 | -82.2110719 |
| York County, Rock Hill CCD | 0.23561054 | 0.67859064 | 34.91638659 | -80.98594065 |
| Lancaster County, Lancaster CCD | 0.18477676 | 0.66487359 | 34.73005549 | -80.77163429 |
| Greenville County, Greer CCD | 0.57436794 | 0.60650888 | 34.93550503 | -82.23837785 |
| Spartanburg County, Wellford CCD | 0.44324906 | 0.60193653 | 34.91930271 | -82.10187109 |
| Charleston County, North Charleston CCD | 0.37950511 | 0.60086068 | 32.85442379 | -79.99194945 |
| Richland County, Columbia CCD | 0.39766003 | 0.58230231 | 33.99354559 | -81.02718444 |
| Pickens County, Easley East CCD | 0.34050565 | 0.57530931 | 34.83154 | -82.50767126 |
| Anderson County, Williamston-Pelzer CCD | 0.32974718 | 0.57154384 | 34.6766611 | -82.5349131 |
| Anderson County, Anderson CCD | 0.31710597 | 0.56966111 | 34.53742357 | -82.69794553 |
| Spartanburg County, Inman CCD | 0.39981173 | 0.56804734 | 35.02329846 | -82.02707609 |
| Sumter County, Sumter CCD | 0.27729962 | 0.5626681 | 33.92223827 | -80.32350134 |
| Cherokee County, Gaffney CCD | 0.22565896 | 0.56078537 | 35.06856608 | -81.65538469 |
| Richland County, Dutch Fork CCD | 0.22512103 | 0.55997848 | 34.02614216 | -81.10371304 |
| Oconee County, Walhalla CCD | 0.24731038 | 0.55433029 | 34.75762874 | -83.04889896 |
| Lexington County, West Columbia-Cayce CCD | 0.34185046 | 0.54841313 | 33.97495823 | -81.05837925 |
| Greenville County, Tigerville CCD | 0.42065627 | 0.54599247 | 35.07168309 | -82.43727584 |
| Greenville County, Travelers Rest CCD | 0.40599785 | 0.54357181 | 35.02427148 | -82.41778802 |
| Spartanburg County, Boiling Springs CCD | 0.39080151 | 0.53899946 | 35.04212333 | -81.99569156 |
| Greenville County, Taylors CCD | 0.4020979 | 0.53577192 | 34.91982754 | -82.29696227 |
| Pickens County, Liberty CCD | 0.39631522 | 0.53254438 | 34.78401116 | -82.68295025 |
| Dillon County, Hamer CCD | 0.1640667 | 0.53119957 | 34.46614656 | -79.34961539 |
| Anderson County, Pendleton CCD | 0.2387036 | 0.53066165 | 34.58169555 | -82.69028061 |
| Oconee County, Westminster CCD | 0.13878429 | 0.52985476 | 34.66905289 | -83.06384376 |
| Spartanburg County, Reidville CCD | 0.36498117 | 0.52797203 | 34.85622775 | -82.14119611 |
| Union County, Union CCD | 0.11861216 | 0.52716514 | 34.72709562 | -81.63583545 |
| Florence County, Florence CCD | 0.28711673 | 0.52474449 | 34.19715 | -79.85173114 |
| Berkeley County, Goose Creek-Hanahan CCD | 0.2140936 | 0.52205487 | 33.0056728 | -80.09757143 |
| Pickens County, Pickens CCD | 0.33902636 | 0.51748252 | 34.87564845 | -82.69484887 |
| Pickens County, Clemson CCD | 0.28577192 | 0.51613771 | 34.68577114 | -82.81110805 |
| Greenville County, Simpsonville CCD | 0.30311996 | 0.51291017 | 34.76180427 | -82.2974091 |
| Oconee County, Seneca CCD | 0.28967187 | 0.51183432 | 34.69194253 | -82.8655968 |
| Aiken County, North Augusta CCD | 0.24852071 | 0.5112964 | 33.56493554 | -81.93262515 |
| Pickens County, Easley CCD | 0.40277031 | 0.50753093 | 34.8094164 | -82.51156965 |
| Aiken County, Aiken CCD | 0.23789672 | 0.50295858 | 33.56045699 | -81.71410348 |
| Anderson County, Powdersville-Piedmont CCD | 0.29962345 | 0.49704142 | 34.78095832 | -82.46093814 |
| Greenville County, Highland CCD | 0.37547068 | 0.49596557 | 35.08423355 | -82.35123842 |
| York County, Fort Mill CCD | 0.33700914 | 0.48682087 | 35.00465278 | -80.96855824 |
| Kershaw County, Camden CCD | 0.14685315 | 0.48628295 | 34.25644564 | -80.61588948 |
| Marlboro County, Bennettsville CCD | 0.17025282 | 0.48251748 | 34.60438375 | -79.68219895 |
| Horry County, Myrtle Beach CCD | 0.35476062 | 0.48090371 | 33.70074633 | -78.93474236 |
| Chester County, Chester CCD | 0.08512641 | 0.48036579 | 34.6945192 | -81.19655345 |
| Lexington County, Irmo CCD | 0.36108123 | 0.47633136 | 34.05172623 | -81.12640227 |
| Pickens County, Central CCD | 0.38152232 | 0.47552448 | 34.70790753 | -82.80730177 |
| Greenville County, Piedmont CCD | 0.29935449 | 0.47095212 | 34.72716779 | -82.38702562 |
| Charleston County, Charleston Central CCD | 0.29908553 | 0.46503497 | 32.82481106 | -79.94349267 |
| Spartanburg County, Fairmont Mills CCD | 0.36525013 | 0.46342119 | 34.8906711 | -82.05131925 |
| Laurens County, Laurens CCD | 0.197149 | 0.46019365 | 34.50170022 | -82.0109605 |
| Lexington County, Lexington CCD | 0.267617 | 0.45804196 | 33.97029407 | -81.19723718 |
| Horry County, Conway East CCD | 0.27003765 | 0.45185584 | 33.72276455 | -78.92988993 |
| Greenwood County, Greenwood CCD | 0.28093061 | 0.44889726 | 34.22392829 | -82.14423868 |
| Greenville County, Slater-Marietta CCD | 0.36121571 | 0.44782141 | 35.078026 | -82.49200794 |
| Orangeburg County, Orangeburg CCD | 0.12103281 | 0.44540075 | 33.4941952 | -80.84890671 |
| Spartanburg County, Gramling CCD | 0.26519634 | 0.44540075 | 35.07729934 | -82.10102111 |
| Richland County, Blythewood CCD | 0.19835933 | 0.44217321 | 34.16674334 | -80.9659033 |
| Pickens County, Six Mile CCD | 0.19284562 | 0.44190425 | 34.88187271 | -82.74946261 |
| Spartanburg County, Landrum CCD | 0.35516407 | 0.43571813 | 35.14686407 | -82.11629946 |
| Cherokee County, Macedonia CCD | 0.27595481 | 0.43437332 | 35.05986137 | -81.76884629 |
| Kershaw County, Elgin CCD | 0.10650888 | 0.43410436 | 34.19742196 | -80.68305909 |
| Greenville County, Fountain Inn CCD | 0.23372781 | 0.43410436 | 34.68256144 | -82.20038536 |
| Chester County, Fort Lawn CCD | 0.10650888 | 0.43329747 | 34.70013247 | -80.88905103 |
| Pickens County, North Pickens CCD | 0.20683163 | 0.43329747 | 35.01152806 | -82.69907315 |
| Saluda County, Saluda CCD | 0.15653577 | 0.43329747 | 33.99755105 | -81.7713077 |
| Lancaster County, Van Wyck CCD | 0.17643895 | 0.43006993 | 35.06378835 | -80.90090251 |
| Newberry County, Newberry CCD | 0.21463152 | 0.42980097 | 34.27159906 | -81.61003736 |
| Orangeburg County, Orangeburg West CCD | 0.13986014 | 0.42522862 | 33.47564352 | -80.87994531 |
| Orangeburg County, Santee-Vance CCD | 0.2436794 | 0.42361485 | 33.4880074 | -80.46589882 |
| Darlington County, Darlington CCD | 0.17697687 | 0.42146315 | 34.26327294 | -79.83785244 |
| Williamsburg County, Hemingway CCD | 0.12210866 | 0.41958042 | 33.75352853 | -79.44769178 |
| Laurens County, Clinton CCD | 0.10543303 | 0.41877353 | 34.4598027 | -81.85220418 |
| Darlington County, Hartsville CCD | 0.2105971 | 0.41608392 | 34.38527054 | -80.06234572 |
| Abbeville County, Abbeville CCD | 0.19983862 | 0.40909091 | 34.18650956 | -82.39951125 |
| Spartanburg County, Mayo CCD | 0.30648198 | 0.4077461 | 35.04442037 | -81.82762069 |
| Calhoun County, St. Matthews CCD | 0.09090909 | 0.40344271 | 33.63389141 | -80.87664387 |
| Lexington County, Batesburg-Leesville CCD | 0.15680473 | 0.40048413 | 33.90092653 | -81.54116412 |
| Richland County, Pontiac CCD | 0.22162453 | 0.3986014 | 34.09342866 | -80.92489156 |
| Aiken County, Beech Island CCD | 0.16137708 | 0.39806347 | 33.47307661 | -81.92181629 |
| Horry County, Little River CCD | 0.30957504 | 0.39779451 | 33.86257309 | -78.64998054 |
| Horry County, Conway CCD | 0.17966649 | 0.39752555 | 33.85674901 | -79.06624014 |
| York County, Clover CCD | 0.18854223 | 0.39698763 | 35.09882046 | -81.17396588 |
| Spartanburg County, Cowpens CCD | 0.31253362 | 0.39483593 | 35.01900275 | -81.80440605 |
| Calhoun County, Sandy Run-Staley Crossroads CCD | 0.06804734 | 0.39456697 | 33.80975973 | -81.00760145 |
| Lexington County, Gilbert CCD | 0.13542227 | 0.39349112 | 33.82097776 | -81.44598046 |
| Lee County, Bishopville CCD | 0.13918774 | 0.39160839 | 34.20905887 | -80.26514183 |
| Dorchester County, Summerville CCD | 0.14416353 | 0.39053254 | 33.01683789 | -80.17721924 |
| Greenwood County, Ninety Six CCD | 0.16783217 | 0.38999462 | 34.21741758 | -82.09013707 |
| Anderson County, Belton CCD | 0.19553523 | 0.38622916 | 34.51859587 | -82.53487257 |
| Newberry County, Prosperity CCD | 0.14147391 | 0.38192577 | 34.21724224 | -81.5372017 |
| Georgetown County, Waccamaw Neck CCD | 0.13864981 | 0.38004303 | 33.53650851 | -79.05347299 |
| Spartanburg County, Fingerville CCD | 0.20333513 | 0.37923615 | 35.14360861 | -82.08892347 |
| Dillon County, Dillon CCD | 0.2260624 | 0.37681549 | 34.40398957 | -79.39093434 |
| Berkeley County, Moncks Corner CCD | 0.12802582 | 0.37654653 | 33.19693632 | -80.00389834 |
| Florence County, Danwood CCD | 0.11108123 | 0.37547068 | 34.17888529 | -79.88328613 |
| Lexington County, Chapin CCD | 0.17213556 | 0.3746638 | 34.17125615 | -81.35228211 |
| Florence County, Lake City-Scranton CCD | 0.09480904 | 0.3633674 | 33.8657597 | -79.75521399 |
| Chester County, Richburg CCD | 0.20763852 | 0.36282948 | 34.76540129 | -81.04345764 |
| Beaufort County, St. Helena Island CCD | 0.1810113 | 0.36202259 | 32.4268177 | -80.67466043 |
| Williamsburg County, Kingstree CCD | 0.07880581 | 0.3577192 | 33.67109277 | -79.82353424 |
| Lancaster County, Kershaw CCD | 0.09924691 | 0.35207101 | 34.54690961 | -80.58545709 |
| Darlington County, Lake Swamp CCD | 0.13528779 | 0.35153308 | 34.20470131 | -79.90561657 |
| Cherokee County, Blacksburg CCD | 0.16527703 | 0.35126412 | 35.15870208 | -81.47040703 |
| Spartanburg County, Chesnee CCD | 0.17845616 | 0.35099516 | 35.13943026 | -81.86590073 |
| Charleston County, Mount Pleasant CCD | 0.23076923 | 0.3507262 | 32.81347974 | -79.86407103 |
| Colleton County, Walterboro CCD | 0.1401291 | 0.34884346 | 32.92068522 | -80.6327515 |
| Chesterfield County, Pageland CCD | 0.18639053 | 0.34830554 | 34.77136309 | -80.40066108 |
| Beaufort County, Bluffton CCD | 0.14402905 | 0.34803658 | 32.24192242 | -80.81846533 |
| Cherokee County, Saratt CCD | 0.10220549 | 0.34561592 | 35.01567435 | -81.77708896 |
| Spartanburg County, Woodruff CCD | 0.15626681 | 0.34480904 | 34.72937272 | -82.02504265 |
| Newberry County, Pomaria CCD | 0.15451856 | 0.34373319 | 34.22573184 | -81.42494527 |
| Jasper County, Hardeeville CCD | 0.12264658 | 0.3415815 | 32.27729186 | -81.08182883 |
| Marlboro County, Wallace CCD | 0.12331899 | 0.34131253 | 34.71908717 | -79.85318843 |
| Clarendon County, Summerton CCD | 0.18665949 | 0.33889188 | 33.58745988 | -80.36355228 |
| Dorchester County, St. George CCD | 0.14470145 | 0.33754707 | 33.18957205 | -80.60725002 |
| Charleston County, West Ashley CCD | 0.23426573 | 0.33189887 | 32.79190335 | -80.03192224 |
| Georgetown County, Georgetown CCD | 0.14712211 | 0.33109198 | 33.32561855 | -79.31903369 |
| Dillon County, Little Rock CCD | 0.10126412 | 0.33055406 | 34.47004052 | -79.3766255 |
| Kershaw County, Boykin CCD | 0.12721893 | 0.32894029 | 34.20869803 | -80.59772144 |
| Chesterfield County, Cheraw CCD | 0.12036041 | 0.32759548 | 34.70836446 | -79.93919648 |
| Beaufort County, Beaufort-Port Royal CCD | 0.24045186 | 0.32490586 | 32.43041473 | -80.70104086 |
| Edgefield County, Trenton CCD | 0.15922539 | 0.32221625 | 33.73538438 | -81.84681415 |
| Berkeley County, Wando CCD | 0.12305003 | 0.32167832 | 32.89107091 | -79.92993141 |
| Richland County, Fort Jackson CCD | 0.09064013 | 0.32167832 | 34.0271664 | -80.93781668 |
| Anderson County, Starr CCD | 0.13851533 | 0.32140936 | 34.42361368 | -82.68992322 |
| Kershaw County, Bethune CCD | 0.15962883 | 0.3189887 | 34.42227849 | -80.33263152 |
| Greenwood County, Ware Shoals-Hodges CCD | 0.19311458 | 0.31603012 | 34.27790303 | -82.21750377 |
| Union County, Jonesville CCD | 0.15586337 | 0.31495428 | 34.85099488 | -81.72408757 |
| Greenville County, Woodville CCD | 0.19983862 | 0.31495428 | 34.6101653 | -82.35224336 |
| Abbeville County, Calhoun Falls CCD | 0.17670791 | 0.31414739 | 34.09482127 | -82.58109074 |
| Darlington County, North Hartsville CCD | 0.18921463 | 0.31414739 | 34.38899962 | -80.08936266 |
| Orangeburg County, Springfield CCD | 0.16810113 | 0.31360947 | 33.50602384 | -81.2855019 |
| Oconee County, Oakway CCD | 0.12076385 | 0.31226466 | 34.62172622 | -83.04407339 |
| Aiken County, Aiken Northeast CCD | 0.15223238 | 0.31145777 | 33.66773912 | -81.66693155 |
| Aiken County, Monetta CCD | 0.12641205 | 0.31118881 | 33.76144392 | -81.51214669 |
| Clarendon County, Manning CCD | 0.23292093 | 0.30903712 | 33.58742503 | -80.19858059 |
| Chester County, Chester West CCD | 0.07100592 | 0.30554061 | 34.70144184 | -81.25091173 |
| Barnwell County, Barnwell CCD | 0.14254976 | 0.30500269 | 33.24400146 | -81.36215374 |
| York County, York CCD | 0.14685315 | 0.30500269 | 35.01745524 | -81.24571493 |
| Laurens County, Gray Court CCD | 0.13569123 | 0.30311996 | 34.65720805 | -82.17715695 |
| Clarendon County, Alcolu CCD | 0.07611619 | 0.29827864 | 33.72355631 | -80.19839484 |
| Marion County, Marion CCD | 0.09144701 | 0.29639591 | 34.17600183 | -79.38463003 |
| Charleston County, Johns Island CCD | 0.16944594 | 0.28967187 | 32.72936274 | -80.07183205 |
| Beaufort County, Hilton Head Island CCD | 0.2345347 | 0.2894029 | 32.22400682 | -80.68687147 |
| Sumter County, Privateer CCD | 0.06912318 | 0.2894029 | 33.85969769 | -80.36375491 |
| Dorchester County, Harleyville CCD | 0.08579882 | 0.28886498 | 33.26346111 | -80.48320274 |
| Sumter County, Shaw AFB-Horatio CCD | 0.14873588 | 0.28886498 | 33.94780099 | -80.51163722 |
| Anderson County, Townville CCD | 0.17872512 | 0.2880581 | 34.53079783 | -82.83769101 |
| Jasper County, Ridgeland CCD | 0.13044648 | 0.28778913 | 32.32612125 | -80.93748955 |
| Chesterfield County, McBee CCD | 0.17025282 | 0.28778913 | 34.418606 | -80.30559135 |
| Saluda County, Batesburg-Leesville CCD | 0.13071544 | 0.28483055 | 33.90431415 | -81.56808077 |
| Spartanburg County, Pacolet CCD | 0.16137708 | 0.28483055 | 34.90519567 | -81.79765424 |
| McCormick County, McCormick CCD | 0.14564282 | 0.28402367 | 33.90484721 | -82.3113527 |
| Laurens County, Joanna CCD | 0.12358795 | 0.28402367 | 34.43769236 | -81.85627099 |
| Berkeley County, Bonneau CCD | 0.09763314 | 0.28375471 | 33.21525807 | -79.97276462 |
| Charleston County, McClellanville CCD | 0.17415277 | 0.28133405 | 33.05448207 | -79.48628488 |
| Chesterfield County, Jefferson CCD | 0.17240452 | 0.2802582 | 34.70156011 | -80.38700596 |
| Hampton County, Furman-Scotia CCD | 0.22135557 | 0.27945132 | 32.63596157 | -81.39371627 |
| Charleston County, James Island CCD | 0.20481442 | 0.27729962 | 32.71435906 | -79.96604446 |
| Oconee County, Long Creek CCD | 0.10946746 | 0.27649274 | 34.76368884 | -83.10345728 |
| Colleton County, Cottageville CCD | 0.15599785 | 0.27514793 | 32.94678498 | -80.49133626 |
| Orangeburg County, Elloree CCD | 0.14967725 | 0.27514793 | 33.49528049 | -80.51935532 |
| Chesterfield County, Mount Croghan CCD | 0.21086606 | 0.27434104 | 34.74194386 | -80.18340743 |
| Charleston County, Edisto Island CCD | 0.22834857 | 0.27192039 | 32.56743659 | -80.21315538 |
| Horry County, Loris CCD | 0.1492738 | 0.26949973 | 34.0569468 | -78.88361632 |
| Charleston County, Ravenel-Hollywood CCD | 0.20723507 | 0.26869285 | 32.78474709 | -80.14228591 |
| Spartanburg County, Pauline CCD | 0.12937063 | 0.26815492 | 34.85544402 | -81.94626764 |
| Dillon County, Latta CCD | 0.09709521 | 0.267617 | 34.34181344 | -79.43218708 |
| Saluda County, North Saluda CCD | 0.19607316 | 0.2654653 | 34.15236916 | -81.74278351 |
| Sumter County, Sumter Southeast CCD | 0.03577192 | 0.26465842 | 33.91487832 | -80.26977093 |
| Georgetown County, Andrews CCD | 0.13824637 | 0.26358257 | 33.43057154 | -79.57136546 |
| Edgefield County, Southeast Edgefield CCD | 0.09198494 | 0.26331361 | 33.59037181 | -81.95545059 |
| Richland County, Camp Ground CCD | 0.12278107 | 0.26304465 | 34.10400008 | -81.00576642 |
| Lexington County, Swansea CCD | 0.10301237 | 0.25981711 | 33.73187083 | -81.10517792 |
| Colleton County, Hendersonville CCD | 0.07235073 | 0.25766541 | 32.8023571 | -80.76469998 |
| Florence County, Timmonsville CCD | 0.14470145 | 0.2528241 | 34.13856557 | -79.91942724 |
| Lee County, Stokes Bridge-Cypress Crossroads CCD | 0.14712211 | 0.25201721 | 34.22373021 | -80.20669689 |
| Allendale County, Allendale CCD | 0.18840775 | 0.25013448 | 32.91881956 | -81.47718784 |
| Oconee County, Mountain Rest CCD | 0.13071544 | 0.24959656 | 34.80494762 | -83.06870715 |
| Horry County, Longs CCD | 0.12385691 | 0.24878967 | 33.91873946 | -78.72044159 |
| Chesterfield County, Chesterfield CCD | 0.17563206 | 0.24878967 | 34.71961436 | -80.0205765 |
| Horry County, Aynor CCD | 0.16608392 | 0.24798279 | 33.99054867 | -79.20305848 |
| Charleston County, Kiawah Island-Seabrook Island CCD | 0.13878429 | 0.24610005 | 32.56372683 | -80.18675195 |
| Williamsburg County, Greeleyville CCD | 0.11296396 | 0.24529317 | 33.55759066 | -79.98468473 |
| Spartanburg County, Enoree CCD | 0.07477138 | 0.24529317 | 34.71951948 | -81.94337758 |
| Abbeville County, Due West CCD | 0.17374933 | 0.24421732 | 34.31931186 | -82.37608357 |
| Marion County, Mullins CCD | 0.14039806 | 0.24314147 | 34.20442867 | -79.2674415 |
| Colleton County, Green Pond CCD | 0.17334588 | 0.24206563 | 32.61534576 | -80.39366972 |
| Union County, Carlisle CCD | 0.08149543 | 0.24206563 | 34.59292597 | -81.46639663 |
| Hampton County, Estill CCD | 0.10059172 | 0.2415277 | 32.72798414 | -81.40379915 |
| Allendale County, Fairfax CCD | 0.15922539 | 0.23937601 | 32.75013038 | -81.39969375 |
| Allendale County, Millett CCD | 0.20602474 | 0.23910705 | 33.10615595 | -81.52428624 |
| Sumter County, Shiloh CCD | 0.10153308 | 0.23749328 | 33.97341728 | -80.03674476 |
| Charleston County, Wadmalaw Island CCD | 0.1732114 | 0.23614847 | 32.64116185 | -80.25272001 |
| Marion County, Brittons Neck CCD | 0.07880581 | 0.23614847 | 33.90777647 | -79.41465714 |
| York County, Hickory Grove CCD | 0.09669177 | 0.23561054 | 35.05673048 | -81.37797618 |
| Jasper County, Grays-Tillman CCD | 0.11242604 | 0.23561054 | 32.58964604 | -81.212561 |
| Laurens County, Cross Hill CCD | 0.12641205 | 0.23426573 | 34.25184679 | -82.00097681 |
| Sumter County, Sumter North CCD | 0.10892953 | 0.23318989 | 33.97371184 | -80.36832098 |
| Abbeville County, Donalds CCD | 0.17038731 | 0.232383 | 34.43315649 | -82.38363412 |
| Bamberg County, Denmark CCD | 0.13111888 | 0.23157612 | 33.30479452 | -81.13193855 |
| Oconee County, Salem CCD | 0.15519096 | 0.23103819 | 35.03626059 | -82.91796855 |
| Lexington County, Pelion CCD | 0.13542227 | 0.22915546 | 33.79347105 | -81.23098541 |
| Barnwell County, Kline-Snelling CCD | 0.14981173 | 0.2288865 | 33.1844678 | -81.42789903 |
| Orangeburg County, Neeses CCD | 0.13690156 | 0.2288865 | 33.55152959 | -81.11225311 |
| Georgetown County, Plantersville CCD | 0.11699839 | 0.22754169 | 33.56646693 | -79.10203151 |
| Hampton County, Hampton-Varnville CCD | 0.10381926 | 0.2232383 | 32.8704656 | -81.10531331 |
| Aiken County, Salley CCD | 0.17428725 | 0.22243141 | 33.57926697 | -81.3265434 |
| Fairfield County, Winnsboro North CCD | 0.11215707 | 0.22243141 | 34.5330015 | -81.17212663 |
| Calhoun County, Cameron CCD | 0.08230231 | 0.22189349 | 33.53945571 | -80.5105638 |
| Fairfield County, Ridgeway CCD | 0.08687466 | 0.22162453 | 34.30628958 | -80.99409089 |
| Chester County, Great Falls CCD | 0.09709521 | 0.22162453 | 34.60035209 | -80.9921927 |
| Clarendon County, Manning East CCD | 0.10408822 | 0.2197418 | 33.67571132 | -80.18060532 |
| Anderson County, Honea Path CCD | 0.13259817 | 0.2197418 | 34.45847517 | -82.40686287 |
| Barnwell County, Williston CCD | 0.1451049 | 0.21947283 | 33.46178881 | -81.29385363 |
| Newberry County, Chappells CCD | 0.14497041 | 0.21947283 | 34.17782351 | -81.76571921 |
| Laurens County, Princeton CCD | 0.11915008 | 0.21947283 | 34.35667266 | -82.12036288 |
| Florence County, Pamplico CCD | 0.07423346 | 0.21893491 | 34.15612441 | -79.7217215 |
| Aiken County, New Ellenton CCD | 0.11000538 | 0.21785906 | 33.42095355 | -81.6849048 |
| Berkeley County, Cordesville CCD | 0.13474987 | 0.21678322 | 33.19177038 | -79.64900788 |
| Fairfield County, Winnsboro South CCD | 0.0541958 | 0.21516945 | 34.3388954 | -81.07091396 |
| Orangeburg County, Branchville CCD | 0.13327058 | 0.21274879 | 33.21826497 | -80.82037674 |
| Orangeburg County, Eutawville CCD | 0.16312534 | 0.21221087 | 33.40713364 | -80.37231582 |
| Anderson County, Iva CCD | 0.13421194 | 0.21140398 | 34.33505097 | -82.70521348 |
| Beaufort County, Sheldon CCD | 0.11430877 | 0.20898332 | 32.59608352 | -80.75029919 |
| Colleton County, Smoaks CCD | 0.16083916 | 0.20521786 | 33.1519351 | -80.83324628 |
| Richland County, Hopkins CCD | 0.07235073 | 0.20521786 | 33.92023661 | -80.98620614 |
| Bamberg County, Bamberg CCD | 0.13932222 | 0.20333513 | 33.29430006 | -81.05186797 |
| Florence County, Olanta CCD | 0.09306079 | 0.20306616 | 33.93635486 | -79.93386341 |
| Marlboro County, Blenheim CCD | 0.08526089 | 0.2027972 | 34.5565119 | -79.66452615 |
| Williamsburg County, Lane CCD | 0.12049489 | 0.20172136 | 33.60870618 | -79.86405728 |
| Newberry County, Whitmire CCD | 0.13394298 | 0.19903174 | 34.37012021 | -81.67475199 |
| Lancaster County, Heath Springs CCD | 0.0894298 | 0.19903174 | 34.6054291 | -80.68505179 |
| Greenwood County, Troy CCD | 0.11040882 | 0.19768693 | 34.12299417 | -82.24512942 |
| Marlboro County, McColl CCD | 0.13179129 | 0.19741797 | 34.67713358 | -79.55500488 |
| Marlboro County, Clio CCD | 0.11995697 | 0.197149 | 34.59673232 | -79.62805703 |
| Allendale County, Sycamore CCD | 0.08795051 | 0.197149 | 33.04973022 | -81.26212636 |
| Aiken County, Wagener CCD | 0.12157074 | 0.197149 | 33.65248806 | -81.3676616 |
| Edgefield County, Johnston CCD | 0.12399139 | 0.19661108 | 33.77964227 | -81.83874595 |
| Williamsburg County, Cades CCD | 0.09037117 | 0.19634212 | 33.82166305 | -79.76446863 |
| Chesterfield County, Patrick CCD | 0.14214632 | 0.19634212 | 34.54703642 | -80.08440748 |
| Edgefield County, Edgefield CCD | 0.10032275 | 0.19472835 | 33.76414353 | -81.89654392 |
| Florence County, Sardis CCD | 0.12775686 | 0.19284562 | 34.09823261 | -79.95553091 |
| Saluda County, Ridge Spring CCD | 0.10341581 | 0.190156 | 33.84808314 | -81.66112577 |
| Lee County, Ashwood CCD | 0.09548144 | 0.18961807 | 34.20899747 | -80.43143202 |
| Berkeley County, St. Stephen CCD | 0.11350188 | 0.18854223 | 33.4879952 | -80.13637308 |
| Horry County, Floyds Crossroads CCD | 0.14362561 | 0.18827327 | 34.12489947 | -79.03503238 |
| Union County, Cross Keys CCD | 0.06562668 | 0.18719742 | 34.51810485 | -81.59162837 |
| Orangeburg County, Bowman CCD | 0.09897795 | 0.18612157 | 33.39928084 | -80.64808735 |
| Abbeville County, Antreville-Lowndesville CCD | 0.1013986 | 0.18423884 | 34.21818835 | -82.47679666 |
| Orangeburg County, Holly Hill CCD | 0.14281872 | 0.18396988 | 33.38504765 | -80.37674425 |
| Orangeburg County, North CCD | 0.10731576 | 0.18396988 | 33.59574582 | -81.10378941 |
| Lee County, Lynchburg CCD | 0.11592254 | 0.18154922 | 34.0691086 | -80.07230493 |
| McCormick County, Parksville CCD | 0.12210866 | 0.18047337 | 33.7337359 | -82.20420264 |
| Sumter County, Sumter Southwest CCD | 0.09763314 | 0.18047337 | 33.88519714 | -80.55175321 |
| Bamberg County, Ehrhardt CCD | 0.14066703 | 0.17939753 | 33.11481301 | -80.89504707 |
| Richland County, Horrell Hill CCD | 0.10610543 | 0.1788596 | 33.93961739 | -80.7893038 |
| Edgefield County, Pleasant Lane CCD | 0.09924691 | 0.17859064 | 33.92648166 | -82.11483452 |
| Dorchester County, Reevesville CCD | 0.08284024 | 0.17724583 | 33.16747008 | -80.61160042 |
| Georgetown County, Sampit-North Santee CCD | 0.12009145 | 0.17563206 | 33.36325837 | -79.42082446 |
| Fairfield County, Monticello-Salem Crossroads CCD | 0.08418505 | 0.17482517 | 34.4501269 | -81.41015295 |
| Kershaw County, Camden Northeast CCD | 0.10731576 | 0.1732114 | 34.24197815 | -80.50792163 |
| Dorchester County, Ridgeville CCD | 0.08606778 | 0.17213556 | 33.10528512 | -80.32331388 |
| Aiken County, Windsor CCD | 0.09736417 | 0.17132867 | 33.5248714 | -81.61098612 |
| McCormick County, Mount Carmel CCD | 0.12009145 | 0.1697149 | 33.86697111 | -82.37303935 |
| Sumter County, Sumter Northeast CCD | 0.08606778 | 0.16944594 | 34.01417924 | -80.33248889 |
| Calhoun County, Fort Motte-Lone Star CCD | 0.09682625 | 0.16837009 | 33.63505512 | -80.54650537 |
| Darlington County, Lamar CCD | 0.12224314 | 0.16756321 | 34.11694584 | -80.09011779 |
| Richland County, Eastover CCD | 0.09279182 | 0.16729424 | 33.94379247 | -80.65047779 |
| Hampton County, Yemassee CCD | 0.0985745 | 0.16648736 | 32.70243268 | -80.86552984 |
| Florence County, Coward CCD | 0.08391608 | 0.16433566 | 33.93950365 | -79.79500397 |
| Bamberg County, Olar CCD | 0.11955352 | 0.16379774 | 33.14995142 | -81.16135547 |
| Williamsburg County, Nesmith CCD | 0.07907477 | 0.1584185 | 33.54082641 | -79.54805337 |
| Clarendon County, Paxville CCD | 0.05217859 | 0.15519096 | 33.67196117 | -80.48412974 |
| Orangeburg County, Norway CCD | 0.11377084 | 0.154922 | 33.44097439 | -81.13337068 |
| Colleton County, Lodge CCD | 0.07369554 | 0.15465304 | 33.05200373 | -80.93443544 |
| Aiken County, Jackson CCD | 0.09144701 | 0.15411512 | 33.2029228 | -81.75208857 |
| Williamsburg County, Trio CCD | 0.10839161 | 0.15384615 | 33.4746761 | -79.56204848 |
| Orangeburg County, Cope CCD | 0.0683163 | 0.15034965 | 33.27314303 | -80.8917911 |
| Greenwood County, Kirksey CCD | 0.10381926 | 0.14873588 | 34.04287948 | -81.9560598 |
| Kershaw County, Mount Pisgah CCD | 0.11256052 | 0.14873588 | 34.4184864 | -80.47233092 |
| Marion County, Centenary CCD | 0.06656805 | 0.14685315 | 33.91165802 | -79.44147723 |
| Florence County, Johnsonville CCD | 0.0809575 | 0.14443249 | 33.88574354 | -79.41938436 |
| Berkeley County, Cross CCD | 0.07557827 | 0.14389457 | 33.12369612 | -80.29229061 |
| Union County, Lockhart CCD | 0.0683163 | 0.14335664 | 34.81210131 | -81.59200353 |
| Barnwell County, Blackville CCD | 0.08795051 | 0.14308768 | 33.43967004 | -81.29802597 |
| Sumter County, Rembert CCD | 0.08149543 | 0.14174287 | 34.0581649 | -80.48950119 |
| Clarendon County, Sardinia CCD | 0.077461 | 0.14120495 | 33.84852527 | -80.11769903 |
| Georgetown County, Pleasant Hill-Folly Grove CCD | 0.09817106 | 0.13986014 | 33.58759853 | -79.40045887 |
| Darlington County, Society Hill CCD | 0.07692308 | 0.13232921 | 34.38480344 | -79.89567912 |
| Clarendon County, Turbeville CCD | 0.08647122 | 0.13071544 | 33.881449 | -80.02811013 |
| Lancaster County, Tradesville CCD | 0.08983324 | 0.12775686 | 34.69044497 | -80.47291878 |
| Hampton County, Brunson CCD | 0.0718128 | 0.12668101 | 32.92867185 | -81.20317992 |
| Kershaw County, Westville CCD | 0.08337816 | 0.1218397 | 34.39189605 | -80.78311557 |
| Union County, West Springs CCD | 0.06750941 | 0.11161915 | 34.81014108 | -81.75957469 |
| Lee County, St. Charles CCD | 0.0866057 | 0.10570199 | 34.13919496 | -80.25170708 |
| Williamsburg County, Indiantown CCD | 0.07087144 | 0.1035503 | 33.76230502 | -79.66656667 |
| York County, McConnells CCD | 0.07504034 | 0.10005379 | 34.92914956 | -81.26278227 |
| Dillon County, Lake View CCD | 0.05890264 | 0.09924691 | 34.27281046 | -79.11358448 |
