## Supplementary material for "Mapping long-distance jump dispersal risk by leveraging transport infrastructure data and sampling bias correction: a case study with the invasive spotted lanternfly (*Lycorma delicatula*)": S15 Tennessee risk

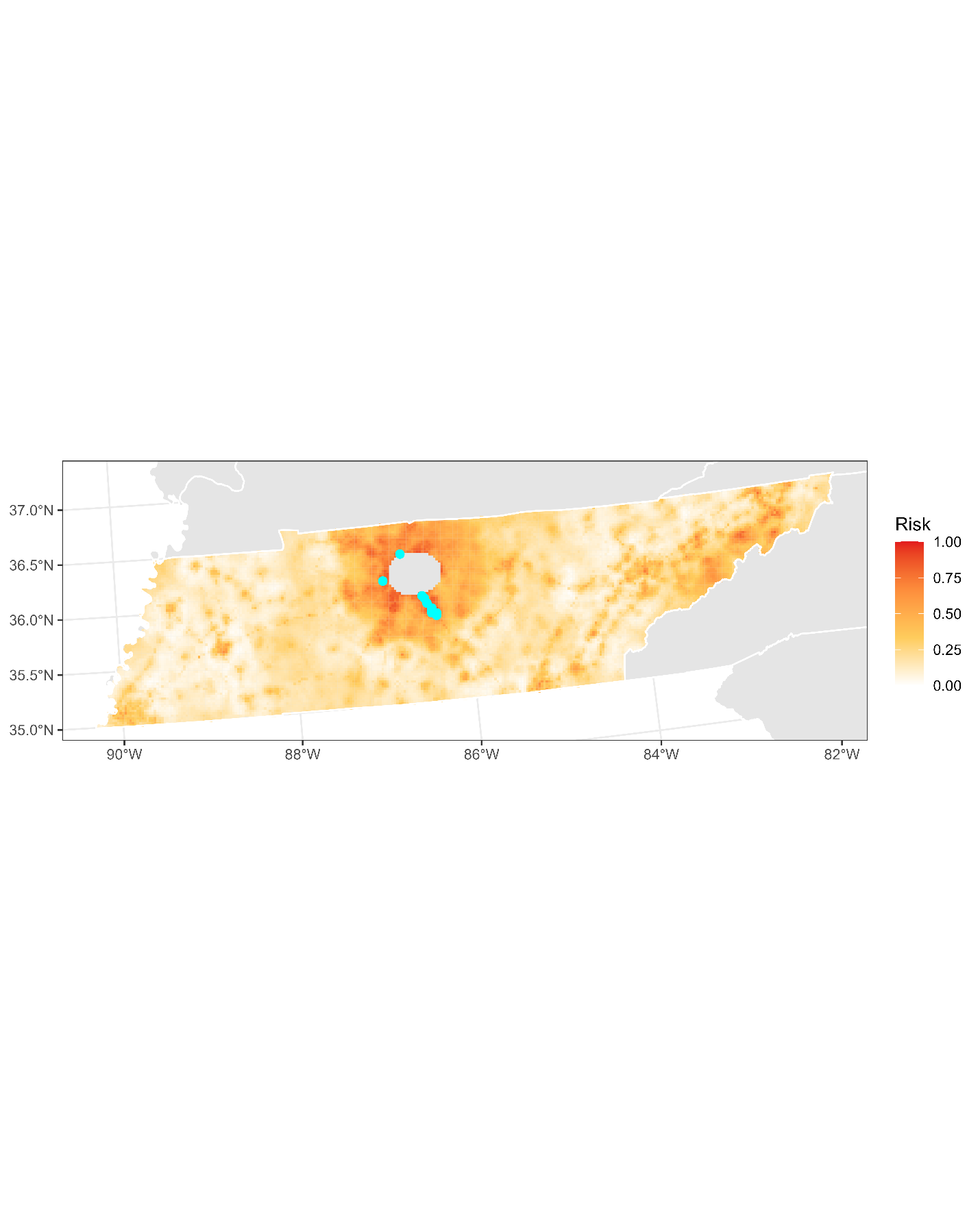

**Figure S15.1:** Projected risk of colonization by *L. delicatula* jump dispersal for Tennessee. Risk projections are based on an ensemble random forest model including distances from existing *L. delicatula* populations. Note that areas falling less than 20 km from existing *L. delicatula* populations are not included in the map because any colonization there would be categorized as diffusive spread based on our methodology (regardless of the actual source of the resulting population). Darker, redder colors indicate increased risk of colonization by jump dispersal. Cyan points show the top ten highest risk locations in the state.

**Table S15.1:** Ranked list of Tennessee county subdivisions sorted by the maximum projected risk of colonization by jump dispersal occurring there. For each subdivision, we identified the 2.5 x 2.5 km grid cell centroid included there with the highest projected risk of colonization by jump dispersal and report that location and risk value.

| **County, subdivision** | **Median risk value** | **Maximum risk value** | **Latitude of maximum risk** | **Longitude of maximum risk** |
| --- | --- | --- | --- | --- |
| Rutherford County, District 18 | 0.87439484 | 0.92684239 | 35.82280886 | -86.40948263 |
| Rutherford County, District 7 | 0.82625067 | 0.90909091 | 35.8005944 | -86.41229444 |
| Rutherford County, District 19 | 0.80876815 | 0.89806347 | 35.89393914 | -86.45664071 |
| Rutherford County, District 3 | 0.72902098 | 0.8951049 | 35.9406028 | -86.47885476 |
| Rutherford County, District 21 | 0.81401291 | 0.88246369 | 35.79608375 | -86.35676196 |
| Rutherford County, District 11 | 0.85946746 | 0.88192577 | 35.96504684 | -86.50388722 |
| Davidson County, District 35 | 0.66164605 | 0.87923615 | 36.1331821 | -86.9333918 |
| Rutherford County, District 13 | 0.87426036 | 0.87869822 | 35.84502287 | -86.40666918 |
| Rutherford County, District 6 | 0.48130715 | 0.87573964 | 35.77387008 | -86.35958818 |
| Sumner County, District 11 | 0.70172136 | 0.87439484 | 36.36250022 | -86.70799967 |
| Sumner County, District 4 | 0.61915008 | 0.87385691 | 36.36929635 | -86.50935323 |
| Sumner County, District 6 | 0.87062937 | 0.87062937 | 36.36706784 | -86.48137397 |
| Rutherford County, District 4 | 0.48587951 | 0.86740183 | 35.84050974 | -86.35110456 |
| Davidson County, District 23 | 0.78832706 | 0.86498117 | 36.10883156 | -86.90816532 |
| Rutherford County, District 15 | 0.83243679 | 0.86067778 | 35.86498249 | -86.3760629 |
| Davidson County, District 1 | 0.69526627 | 0.85233997 | 36.19557437 | -86.86951808 |
| Williamson County, District 8 | 0.79558903 | 0.84884346 | 35.96908216 | -86.84066556 |
| Rutherford County, District 14 | 0.8337816 | 0.84776762 | 35.88944953 | -86.40103733 |
| Williamson County, District 11 | 0.71382464 | 0.84749866 | 35.88233454 | -86.87923048 |
| Williamson County, District 12 | 0.72108661 | 0.84346423 | 35.90241502 | -86.84873375 |
| Williamson County, District 4 | 0.70589026 | 0.84346423 | 35.92248736 | -86.81822088 |
| Williamson County, District 2 | 0.43894567 | 0.83889188 | 35.8558194 | -86.82630382 |
| Davidson County, District 20 | 0.82033351 | 0.8337816 | 36.17335535 | -86.87220864 |
| Wilson County, District 21 | 0.77998924 | 0.83243679 | 36.16909085 | -86.25266218 |
| Williamson County, District 7 | 0.79532006 | 0.83055406 | 36.0070467 | -86.75171155 |
| Rutherford County, District 2 | 0.4281872 | 0.82786444 | 35.90940702 | -86.37041166 |
| Williamson County, District 10 | 0.75470683 | 0.81925767 | 35.94900475 | -86.87119122 |
| Rutherford County, District 16 | 0.80836471 | 0.81845078 | 35.84951 | -86.46224038 |
| Williamson County, District 6 | 0.76573427 | 0.81576116 | 35.98699267 | -86.78227604 |
| Robertson County, District 11 | 0.63770845 | 0.81118881 | 36.41039459 | -87.04126743 |
| Unicoi County, District 2 | 0.59924691 | 0.80580958 | 36.15177402 | -82.41237355 |
| Davidson County, District 10 | 0.73265196 | 0.80500269 | 36.36467578 | -86.73598384 |
| Sumner County, District 5 | 0.76385153 | 0.80446477 | 36.38479715 | -86.42259153 |
| Cheatham County, District 3 | 0.68504572 | 0.80016138 | 36.38607131 | -87.01591341 |
| Greene County, District 3 | 0.72364174 | 0.79666487 | 36.17356405 | -82.80502831 |
| Sumner County, District 3 | 0.63622916 | 0.79639591 | 36.36259119 | -86.42542043 |
| Robertson County, District 6 | 0.67079075 | 0.78994083 | 36.40829046 | -87.01325762 |
| Greene County, District 4 | 0.42025282 | 0.78671329 | 36.15145654 | -82.80890895 |
| Rutherford County, District 12 | 0.78160301 | 0.78160301 | 35.98726067 | -86.50109614 |
| Rutherford County, District 10 | 0.77003765 | 0.77918236 | 35.96948809 | -86.55955206 |
| Davidson County, District 22 | 0.70387305 | 0.77487897 | 36.08873888 | -86.93872941 |
| Sullivan County, District 11 | 0.60906401 | 0.77461001 | 36.52743606 | -82.54337104 |
| Davidson County, District 33 | 0.76788596 | 0.76788596 | 35.97390331 | -86.6152234 |
| Davidson County, District 34 | 0.69688004 | 0.76385153 | 36.0643887 | -86.91351601 |
| Robertson County, District 10 | 0.67052179 | 0.76250672 | 36.4886719 | -86.8904642 |
| Greene County, District 7 | 0.44916622 | 0.76089295 | 36.1766502 | -82.83280694 |
| Davidson County, District 3 | 0.73211404 | 0.75820334 | 36.36684479 | -86.76396962 |
| Sumner County, District 7 | 0.72713825 | 0.75605164 | 36.37594265 | -86.59330089 |
| Rutherford County, District 9 | 0.6457773 | 0.74421732 | 35.9516866 | -86.61798056 |
| Robertson County, District 1 | 0.58499193 | 0.74179666 | 36.60023196 | -86.59349517 |
| Washington County, District 9 | 0.39402905 | 0.74018289 | 36.15495153 | -82.44012794 |
| Robertson County, District 2 | 0.71382464 | 0.73722431 | 36.45134816 | -86.69700727 |
| Rutherford County, District 8 | 0.48708983 | 0.73614847 | 35.80507907 | -86.4678335 |
| Cheatham County, District 2 | 0.59185046 | 0.73614847 | 36.36385178 | -87.01856764 |
| Williamson County, District 9 | 0.5500269 | 0.73561054 | 35.99558863 | -86.89367629 |
| Cheatham County, District 6 | 0.53604088 | 0.73318989 | 36.07914737 | -87.10870767 |
| Sumner County, District 2 | 0.62937063 | 0.73292093 | 36.59580829 | -86.53736148 |
| Washington County, District 4 | 0.70414201 | 0.73157612 | 36.32532541 | -82.35252017 |
| Robertson County, District 9 | 0.67025282 | 0.72996235 | 36.48653262 | -86.86242952 |
| Montgomery County, District 15 | 0.64281872 | 0.72969338 | 36.51177123 | -87.19894958 |
| Robertson County, District 7 | 0.42011834 | 0.72646584 | 36.43471215 | -87.06663782 |
| Washington County, District 5 | 0.4880312 | 0.72431415 | 36.30960895 | -82.41215885 |
| Greene County, District 1 | 0.2049489 | 0.72108661 | 36.1956711 | -82.80114541 |
| Rutherford County, District 20 | 0.6640667 | 0.7197418 | 35.85174382 | -86.49002844 |
| Dickson County, District 6 | 0.53698225 | 0.71812803 | 36.0268946 | -87.31164842 |
| Robertson County, District 3 | 0.63582571 | 0.71785906 | 36.49576995 | -86.69150134 |
| Cheatham County, District 5 | 0.51707907 | 0.71624529 | 36.07496323 | -87.05293087 |
| Sumner County, District 12 | 0.6492738 | 0.71463152 | 36.49139183 | -86.63543977 |
| Knox County, District 1 | 0.4316837 | 0.71140398 | 35.95698306 | -83.94334528 |
| Williamson County, District 5 | 0.4767348 | 0.7049489 | 35.9804773 | -86.69874251 |
| Wilson County, District 7 | 0.51048951 | 0.70414201 | 36.14688579 | -86.25553288 |
| Montgomery County, District 20 | 0.67280796 | 0.69956966 | 36.51588583 | -87.25505802 |
| Hamblen County, District 5 | 0.41527703 | 0.69822485 | 36.20600172 | -83.30912364 |
| Washington County, District 2 | 0.53227542 | 0.69499731 | 36.35060723 | -82.3763206 |
| Maury County, District 3 | 0.59844002 | 0.69284562 | 35.60185943 | -87.02474443 |
| Unicoi County, District 1 | 0.43141474 | 0.69042496 | 36.13285576 | -82.44411419 |
| Williamson County, District 1 | 0.5034965 | 0.690156 | 35.77334205 | -86.9203697 |
| Robertson County, District 5 | 0.61565358 | 0.68934911 | 36.41774333 | -86.84252532 |
| Robertson County, District 4 | 0.66030124 | 0.68881119 | 36.43131412 | -86.72776647 |
| Dickson County, District 7 | 0.47902098 | 0.68827327 | 36.02892949 | -87.33952539 |
| Robertson County, District 8 | 0.49583109 | 0.6880043 | 36.49794918 | -86.71953456 |
| Maury County, District 6 | 0.35570199 | 0.68746638 | 35.75324119 | -86.95079577 |
| Wilson County, District 15 | 0.50026896 | 0.68692846 | 36.23570359 | -86.24403994 |
| Wilson County, District 6 | 0.55123722 | 0.68181818 | 36.26019185 | -86.26909414 |
| Montgomery County, District 3 | 0.56495428 | 0.67724583 | 36.53399273 | -87.19634339 |
| Dickson County, District 11 | 0.65223238 | 0.67563206 | 36.07542163 | -87.36232194 |
| Cheatham County, District 1 | 0.58364712 | 0.6732114 | 36.27286283 | -87.00120968 |
| Wilson County, District 5 | 0.6732114 | 0.6732114 | 36.3358833 | -86.37231707 |
| Washington County, District 6 | 0.52716514 | 0.66541151 | 36.29386261 | -82.47177267 |
| Hamilton County, District 4 | 0.46839699 | 0.66110812 | 35.05176051 | -85.28311744 |
| Davidson County, District 31 | 0.6605702 | 0.6605702 | 35.97610115 | -86.64306149 |
| Maury County, District 11 | 0.51183432 | 0.66003228 | 35.73101306 | -86.95344694 |
| Williamson County, District 3 | 0.64550834 | 0.65949435 | 35.77120872 | -86.89259792 |
| Trousdale County, District 6 | 0.61484669 | 0.65761162 | 36.34900429 | -86.2575999 |
| Maury County, District 1 | 0.56710597 | 0.6570737 | 35.64842698 | -87.04722091 |
| Dickson County, District 10 | 0.55123722 | 0.6570737 | 36.07339213 | -87.33442724 |
| Cocke County, District 7 | 0.59521248 | 0.6570737 | 35.99197358 | -83.20399685 |
| Marshall County, District 9 | 0.20857988 | 0.64954276 | 35.44927193 | -86.79154468 |
| Madison County, District 1 | 0.61323292 | 0.64550834 | 35.61200131 | -88.81184454 |
| Washington County, District 3 | 0.54639591 | 0.64281872 | 36.33170105 | -82.4081539 |
| Sumner County, District 1 | 0.5535234 | 0.6422808 | 36.51124779 | -86.32149382 |
| Cannon County, District 1 | 0.46342119 | 0.63932222 | 35.84208575 | -86.09824274 |
| Marshall County, District 1 | 0.56091985 | 0.63636364 | 35.64279472 | -86.68412955 |
| Knox County, District 9 | 0.30876815 | 0.63555675 | 35.93200945 | -83.91914932 |
| Trousdale County, District 1 | 0.60180204 | 0.63394298 | 36.38421438 | -86.13993093 |
| Cannon County, District 4 | 0.42361485 | 0.63206025 | 35.770811 | -86.05145238 |
| Cocke County, District 1 | 0.43275955 | 0.63206025 | 35.83110616 | -83.17489732 |
| Cocke County, District 4 | 0.51748252 | 0.63179129 | 35.97284332 | -83.235472 |
| Sullivan County, District 1 | 0.1338085 | 0.62345347 | 36.55218826 | -82.16865211 |
| Trousdale County, District 5 | 0.61175363 | 0.6218397 | 36.39340821 | -86.25184263 |
| Hamblen County, District 11 | 0.46369016 | 0.62076385 | 36.18387863 | -83.31285852 |
| Jefferson County, District 3 | 0.36740183 | 0.62022593 | 36.13665832 | -83.29253715 |
| Jefferson County, District 2 | 0.27582033 | 0.61726735 | 36.04081423 | -83.44994232 |
| Trousdale County, District 7 | 0.59924691 | 0.61565358 | 36.39111956 | -86.22386215 |
| Cocke County, District 5 | 0.55311996 | 0.61538462 | 35.83410701 | -83.17865285 |
| Dickson County, District 12 | 0.46906939 | 0.61323292 | 36.04503507 | -87.2533263 |
| Montgomery County, District 4 | 0.33566434 | 0.61188811 | 36.49775001 | -87.31374438 |
| Putnam County, District 2 | 0.57046799 | 0.61135019 | 36.16909883 | -85.46288412 |
| Trousdale County, District 4 | 0.54585799 | 0.61108123 | 36.40641319 | -86.13701715 |
| Montgomery County, District 19 | 0.46839699 | 0.60812265 | 36.56033079 | -87.24987696 |
| Montgomery County, District 2 | 0.54115116 | 0.60731576 | 36.56442144 | -87.30602438 |
| Trousdale County, District 10 | 0.55486821 | 0.60005379 | 36.36662389 | -86.19877915 |
| Trousdale County, District 2 | 0.53442711 | 0.5979021 | 36.40177575 | -86.08105528 |
| Montgomery County, District 21 | 0.52904788 | 0.59682625 | 36.51793328 | -87.28311453 |
| Montgomery County, District 6 | 0.14900484 | 0.59655729 | 36.52403625 | -87.36729319 |
| Cannon County, District 3 | 0.2619688 | 0.59548144 | 35.75093359 | -86.08210578 |
| Montgomery County, District 1 | 0.53953739 | 0.59359871 | 36.58460195 | -87.27536531 |
| Maury County, District 5 | 0.53792361 | 0.59359871 | 35.66644332 | -86.98912591 |
| Greene County, District 2 | 0.3986014 | 0.59306079 | 36.16426679 | -82.72170604 |
| Cocke County, District 6 | 0.57208176 | 0.59171598 | 35.92260503 | -83.18755321 |
| Trousdale County, District 3 | 0.57974718 | 0.58848843 | 36.42397288 | -86.07812348 |
| Sullivan County, District 8 | 0.31629909 | 0.58741259 | 36.50534211 | -82.54734672 |
| Washington County, District 1 | 0.40586337 | 0.5866057 | 36.37906379 | -82.42797333 |
| Dickson County, District 1 | 0.29908553 | 0.58526089 | 36.14211425 | -87.35468785 |
| Greene County, District 5 | 0.39335664 | 0.58364712 | 36.11032371 | -82.84441818 |
| Hamblen County, District 4 | 0.58364712 | 0.58364712 | 36.20302647 | -83.28131511 |
| McMinn County, District 1 | 0.12856374 | 0.58337816 | 35.46067459 | -84.61004992 |
| DeKalb County, District 2 | 0.48130715 | 0.58337816 | 36.0150552 | -86.01928119 |
| Hamilton County, District 5 | 0.33754707 | 0.5831092 | 35.06638762 | -85.19760467 |
| Wilson County, District 12 | 0.41769769 | 0.58015062 | 36.12468033 | -86.2584019 |
| Washington County, District 7 | 0.34211942 | 0.57988166 | 36.40749756 | -82.47966503 |
| McMinn County, District 5 | 0.14066703 | 0.57934373 | 35.4385 | -84.61337116 |
| Smith County, District 4 | 0.45521786 | 0.57826789 | 36.33518207 | -86.08984033 |
| Cheatham County, District 4 | 0.41070468 | 0.57799892 | 36.36805113 | -87.07455613 |
| Cannon County, District 2 | 0.46315223 | 0.57638515 | 35.83976035 | -86.07046999 |
| Madison County, District 5 | 0.48265196 | 0.56885422 | 35.63257765 | -88.78197151 |
| Knox County, District 6 | 0.32638515 | 0.56858526 | 35.91014142 | -84.14803295 |
| Shelby County, District 9 | 0.40559441 | 0.56804734 | 35.1170269 | -89.96741279 |
| Cannon County, District 5 | 0.56724045 | 0.56724045 | 35.81755435 | -86.07338148 |
| Putnam County, District 12 | 0.3598709 | 0.56697149 | 36.19128371 | -85.45977991 |
| Bedford County, District 2 | 0.48144164 | 0.56643357 | 35.5784243 | -86.44032244 |
| Sullivan County, District 5 | 0.23157612 | 0.56589564 | 36.43256837 | -82.30452716 |
| Unicoi County, District 3 | 0.2830823 | 0.56562668 | 36.17068358 | -82.38061703 |
| Warren County, District 4 | 0.35529855 | 0.56508876 | 35.70035354 | -85.75235188 |
| Carter County, District 5 | 0.48897257 | 0.5648198 | 36.27794856 | -82.33276218 |
| Bradley County, District 4 | 0.2739376 | 0.56455083 | 35.14645925 | -84.85199203 |
| Maury County, District 2 | 0.54303389 | 0.56428187 | 35.60396214 | -87.05245818 |
| Washington County, District 10 | 0.33243679 | 0.56401291 | 36.17704683 | -82.43613936 |
| Montgomery County, District 13 | 0.52797203 | 0.56239914 | 36.54423287 | -87.33666701 |
| Macon County, District 6 | 0.50753093 | 0.56159225 | 36.61311066 | -86.19489203 |
| Trousdale County, District 8 | 0.53846154 | 0.56105433 | 36.43322421 | -86.19008669 |
| Marshall County, District 2 | 0.40761162 | 0.55863367 | 35.62275234 | -86.71456573 |
| Putnam County, District 5 | 0.36363636 | 0.55863367 | 36.14691352 | -85.46598651 |
| Putnam County, District 6 | 0.51949973 | 0.55863367 | 36.15183908 | -85.52172108 |
| Putnam County, District 11 | 0.32423346 | 0.55863367 | 36.19375112 | -85.48766244 |
| Dickson County, District 8 | 0.55755783 | 0.55755783 | 36.07744459 | -87.39021813 |
| Hamilton County, District 8 | 0.38757396 | 0.55540613 | 34.99978924 | -85.20700273 |
| Bradley County, District 5 | 0.47283486 | 0.55406132 | 35.14907078 | -84.87947458 |
| Maury County, District 4 | 0.44042496 | 0.55406132 | 35.60605837 | -87.08017347 |
| Loudon County, District 1 | 0.26304465 | 0.55406132 | 35.7327017 | -84.40059757 |
| Putnam County, District 10 | 0.48265196 | 0.5535234 | 36.17647945 | -85.54651292 |
| Montgomery County, District 16 | 0.50161377 | 0.55244755 | 36.54828297 | -87.39280709 |
| Hickman County, District 3 | 0.33405056 | 0.55190963 | 35.93592743 | -87.29401612 |
| Hamblen County, District 2 | 0.55083378 | 0.55083378 | 36.22514829 | -83.27757005 |
| Cocke County, District 2 | 0.37278107 | 0.54814416 | 35.93569404 | -83.10067777 |
| Coffee County, District 12 | 0.31845078 | 0.54599247 | 35.53346021 | -86.16640006 |
| Macon County, District 7 | 0.49811727 | 0.54599247 | 36.4508097 | -86.13118443 |
| Blount County, District 1 | 0.50268962 | 0.54599247 | 35.77979882 | -83.97158847 |
| Wilson County, District 9 | 0.42052179 | 0.54491662 | 36.01739225 | -86.0471166 |
| Hickman County, District 4 | 0.48386229 | 0.54276493 | 35.93388176 | -87.26617453 |
| Jefferson County, District 10 | 0.3809844 | 0.54142012 | 36.13226929 | -83.4629657 |
| Sullivan County, District 9 | 0.47512103 | 0.54061323 | 36.57477076 | -82.56332715 |
| Bradley County, District 3 | 0.46906939 | 0.53846154 | 35.16864646 | -84.84875246 |
| Macon County, District 5 | 0.50484131 | 0.53711673 | 36.52431853 | -86.20650059 |
| Putnam County, District 1 | 0.53267886 | 0.53496503 | 36.14937956 | -85.49385289 |
| Knox County, District 2 | 0.4133943 | 0.53362023 | 36.00409749 | -83.9640234 |
| Carter County, District 3 | 0.31925767 | 0.53362023 | 36.31892388 | -82.29689586 |
| Madison County, District 7 | 0.07853685 | 0.5330823 | 35.61368558 | -88.8396063 |
| Sullivan County, District 3 | 0.51519634 | 0.53227542 | 36.57750312 | -82.19246248 |
| Hamblen County, District 6 | 0.23372781 | 0.53200646 | 36.22812437 | -83.30538657 |
| Smith County, District 2 | 0.47175901 | 0.53039268 | 36.3328555 | -86.0618864 |
| Knox County, District 8 | 0.19271114 | 0.52931684 | 36.00633994 | -83.76615856 |
| Sullivan County, District 10 | 0.33996772 | 0.5274341 | 36.54637592 | -82.51148893 |
| Dickson County, District 2 | 0.27407208 | 0.52662722 | 36.05521235 | -87.39275172 |
| Madison County, District 2 | 0.51317913 | 0.52635826 | 35.63595361 | -88.83750993 |
| Rhea County, District 6 | 0.31737493 | 0.52555137 | 35.47782082 | -85.02726114 |
| Bedford County, District 1 | 0.32167832 | 0.52474449 | 35.50951986 | -86.42103595 |
| Franklin County, District 1 | 0.43652501 | 0.52447552 | 35.16873699 | -86.10201814 |
| Bedford County, District 8 | 0.52259279 | 0.52313072 | 35.55620467 | -86.44311626 |
| Blount County, District 2 | 0.40129102 | 0.52286175 | 35.80194854 | -83.96806527 |
| Bedford County, District 3 | 0.43154922 | 0.52097902 | 35.61618774 | -86.63143726 |
| Putnam County, District 8 | 0.34346423 | 0.5204411 | 36.13942226 | -85.63626404 |
| Cumberland County, District 2 | 0.44956966 | 0.51882733 | 35.97351836 | -85.03972324 |
| Macon County, District 9 | 0.51008607 | 0.51882733 | 36.55481484 | -86.03246085 |
| Maury County, District 8 | 0.20857988 | 0.51802044 | 35.63784275 | -86.90858737 |
| Smith County, District 5 | 0.46382464 | 0.51748252 | 36.16339205 | -85.91503497 |
| Hawkins County, District 1 | 0.35059172 | 0.51667563 | 36.53998225 | -82.65497866 |
| Dickson County, District 5 | 0.40182894 | 0.51667563 | 36.11377232 | -87.27350569 |
| Shelby County, District 8 | 0.3126681 | 0.51613771 | 35.11984848 | -90.02262575 |
| Smith County, District 1 | 0.4527972 | 0.51425498 | 36.36077223 | -85.86027238 |
| Maury County, District 7 | 0.17939753 | 0.5126412 | 35.63038191 | -87.10528843 |
| Sullivan County, District 2 | 0.43867671 | 0.5126412 | 36.57426888 | -82.16456275 |
| Jackson County, District 6 | 0.29962345 | 0.50995159 | 36.33620013 | -85.83531769 |
| Greene County, District 6 | 0.2465035 | 0.50860678 | 36.22322113 | -83.05130601 |
| Macon County, District 3 | 0.41662184 | 0.50779989 | 36.55247516 | -86.0044271 |
| Montgomery County, District 18 | 0.3154922 | 0.50779989 | 36.63312703 | -87.32639138 |
| Hamilton County, District 6 | 0.36188811 | 0.50618612 | 35.03206944 | -85.31368184 |
| Shelby County, District 12 | 0.34669177 | 0.50591716 | 35.04728936 | -89.91751135 |
| Sevier County, District 1 | 0.36417429 | 0.50484131 | 35.84307085 | -83.28558334 |
| Macon County, District 4 | 0.48440022 | 0.50376547 | 36.48589322 | -86.01329749 |
| Dickson County, District 9 | 0.35260893 | 0.5034965 | 36.12190786 | -87.38514649 |
| Blount County, District 10 | 0.17576654 | 0.50322754 | 35.84906422 | -83.98871653 |
| Jefferson County, District 1 | 0.41460463 | 0.50215169 | 36.01574221 | -83.42588249 |
| McMinn County, District 2 | 0.19096288 | 0.50134481 | 35.45800093 | -84.58246462 |
| Franklin County, District 8 | 0.24852071 | 0.50026896 | 35.21317489 | -86.09625935 |
| Macon County, District 2 | 0.43625605 | 0.49784831 | 36.4391605 | -85.9912112 |
| DeKalb County, District 1 | 0.4338354 | 0.49757934 | 36.05711267 | -85.98555977 |
| Smith County, District 3 | 0.44176977 | 0.49757934 | 36.17279752 | -86.02659241 |
| Cumberland County, District 4 | 0.2049489 | 0.49704142 | 35.99569509 | -85.03650324 |
| Hamilton County, District 1 | 0.23614847 | 0.49596557 | 35.22429976 | -85.20313277 |
| Madison County, District 3 | 0.29115116 | 0.49408284 | 35.65989986 | -88.86319146 |
| Macon County, District 8 | 0.45185584 | 0.49273803 | 36.57700876 | -86.02950874 |
| Coffee County, District 15 | 0.49193115 | 0.49193115 | 35.38025135 | -86.21407683 |
| Sevier County, District 4 | 0.39187735 | 0.49058634 | 35.78106908 | -83.54935107 |
| Macon County, District 10 | 0.49004841 | 0.49004841 | 36.53495302 | -86.06343853 |
| Montgomery County, District 10 | 0.13421194 | 0.48977945 | 36.57453312 | -87.44641948 |
| Hamblen County, District 3 | 0.46100054 | 0.48897257 | 36.22216573 | -83.24975571 |
| Campbell County, District 3 | 0.29747176 | 0.48682087 | 36.33931141 | -84.16527144 |
| Hamblen County, District 10 | 0.29666487 | 0.48655191 | 36.13367907 | -83.26475475 |
| Blount County, District 5 | 0.42872512 | 0.48493814 | 35.7576486 | -83.97510962 |
| Coffee County, District 19 | 0.48359333 | 0.48359333 | 35.35573467 | -86.18932854 |
| Anderson County, District 3 | 0.17509414 | 0.48332437 | 36.17315371 | -84.07833761 |
| Knox County, District 4 | 0.30849919 | 0.48305541 | 35.94047182 | -84.00235622 |
| Humphreys County, District 5 | 0.22135557 | 0.48278644 | 35.88266275 | -87.80505311 |
| Smith County, District 8 | 0.4260355 | 0.48197956 | 36.21014705 | -85.93699359 |
| Warren County, District 2 | 0.36498117 | 0.4817106 | 35.68055232 | -85.78305485 |
| Madison County, District 6 | 0.05392684 | 0.48117267 | 35.58973329 | -88.81394775 |
| Jefferson County, District 4 | 0.33916084 | 0.48090371 | 36.11307258 | -83.49443361 |
| Coffee County, District 17 | 0.4760624 | 0.4760624 | 35.37795215 | -86.18646831 |
| Bedford County, District 7 | 0.46019365 | 0.47579344 | 35.48729937 | -86.42383286 |
| Loudon County, District 5 | 0.19876278 | 0.4739107 | 35.8741325 | -84.2381427 |
| Putnam County, District 7 | 0.26681011 | 0.47337278 | 36.13210432 | -85.55266744 |
| Washington County, District 8 | 0.24878967 | 0.4725659 | 36.28123599 | -82.55916741 |
| Bradley County, District 7 | 0.45629371 | 0.47229693 | 35.15167588 | -84.906959 |
| Sullivan County, District 7 | 0.30876815 | 0.47175901 | 36.47693923 | -82.49556231 |
| Blount County, District 3 | 0.39537386 | 0.47122109 | 35.79912594 | -83.94037939 |
| Sevier County, District 11 | 0.31374395 | 0.47014524 | 35.77965735 | -83.32440175 |
| Carter County, District 7 | 0.23614847 | 0.46906939 | 36.27474634 | -82.30496713 |
| Roane County, District 1 | 0.16648736 | 0.46880043 | 35.88991906 | -84.62979789 |
| Sevier County, District 5 | 0.44607316 | 0.46853147 | 35.84455586 | -83.51071977 |
| Johnson County, District 5 | 0.13152232 | 0.46853147 | 36.49412269 | -81.86617746 |
| Shelby County, District 7 | 0.30796127 | 0.46584185 | 35.16161924 | -89.96390131 |
| Dyer County, District 8 | 0.35153308 | 0.46557289 | 36.02465287 | -89.36204567 |
| Bedford County, District 9 | 0.464766 | 0.464766 | 35.49177986 | -86.47914468 |
| Rhea County, District 7 | 0.22538999 | 0.46422808 | 35.51961315 | -84.99323217 |
| Jefferson County, District 5 | 0.32571275 | 0.46422808 | 36.03787143 | -83.4221889 |
| Marshall County, District 6 | 0.30204411 | 0.46395912 | 35.62925858 | -86.79770845 |
| Sevier County, District 8 | 0.32167832 | 0.46342119 | 35.9668921 | -83.60331205 |
| DeKalb County, District 3 | 0.42899408 | 0.46180742 | 35.95650009 | -85.85826635 |
| Montgomery County, District 14 | 0.43074233 | 0.46100054 | 36.56645691 | -87.33410038 |
| Hamblen County, District 7 | 0.38138784 | 0.46046261 | 36.24726966 | -83.27382279 |
| Sullivan County, District 6 | 0.2288865 | 0.45992469 | 36.514786 | -82.43180079 |
| Knox County, District 3 | 0.39833244 | 0.45965573 | 35.90736099 | -84.12030097 |
| Giles County, District 7 | 0.36363636 | 0.45911781 | 35.19949903 | -87.044181 |
| Putnam County, District 4 | 0.23023131 | 0.45831092 | 36.15164952 | -85.2678142 |
| Cocke County, District 3 | 0.28160301 | 0.45696611 | 36.01709077 | -83.22797568 |
| Carroll County, District 9 | 0.26035503 | 0.45642819 | 36.13075092 | -88.50944367 |
| Montgomery County, District 17 | 0.43760086 | 0.45589026 | 36.58868062 | -87.33153223 |
| Hawkins County, District 2 | 0.25255514 | 0.45535234 | 36.52100423 | -82.68682191 |
| Hickman County, District 5 | 0.19150081 | 0.45400753 | 35.88736641 | -87.24347937 |
| Hardin County, District 4 | 0.20831092 | 0.45293168 | 35.22256355 | -88.3205512 |
| Montgomery County, District 11 | 0.45077999 | 0.45077999 | 36.57252392 | -87.41833745 |
| Anderson County, District 2 | 0.43208714 | 0.45077999 | 36.109499 | -84.11665993 |
| Bedford County, District 6 | 0.4499731 | 0.4499731 | 35.46732125 | -86.45427528 |
| Coffee County, District 20 | 0.44916622 | 0.44916622 | 35.35803325 | -86.21692913 |
| Dyer County, District 6 | 0.44607316 | 0.44916622 | 36.02620766 | -89.3899703 |
| White County, District 2 | 0.39564282 | 0.44889726 | 35.94968642 | -85.52162027 |
| Knox County, District 5 | 0.23668639 | 0.44862829 | 35.88798825 | -84.15150867 |
| Warren County, District 6 | 0.41191501 | 0.44782141 | 35.70275354 | -85.78006391 |
| Jefferson County, District 8 | 0.29962345 | 0.44755245 | 36.02603556 | -83.31119687 |
| Maury County, District 9 | 0.14254976 | 0.44701452 | 35.58382467 | -87.0827818 |
| Warren County, District 5 | 0.3337816 | 0.44674556 | 35.67815299 | -85.7553508 |
| Knox County, District 7 | 0.2619688 | 0.4464766 | 36.03187159 | -84.0160288 |
| Rhea County, District 5 | 0.44620764 | 0.44620764 | 35.49742911 | -84.99644466 |
| Blount County, District 4 | 0.21301775 | 0.44620764 | 35.75482754 | -83.9474396 |
| Maury County, District 10 | 0.15438408 | 0.44593868 | 35.56783684 | -87.16849553 |
| Obion County, District 7 | 0.39443249 | 0.44459387 | 36.41109272 | -89.07409488 |
| Hamblen County, District 14 | 0.18531469 | 0.44459387 | 36.18980804 | -83.36846616 |
| Coffee County, District 18 | 0.44432491 | 0.44432491 | 35.38254409 | -86.241687 |
| Smith County, District 6 | 0.44432491 | 0.44432491 | 36.25453929 | -85.93105848 |
| Hickman County, District 2 | 0.23910705 | 0.44405594 | 35.91369494 | -87.29657316 |
| Smith County, District 7 | 0.43356643 | 0.44351802 | 36.25924727 | -85.98690202 |
| Montgomery County, District 9 | 0.43275955 | 0.44351802 | 36.61898477 | -87.44134754 |
| Bedford County, District 5 | 0.21086606 | 0.44324906 | 35.46507838 | -86.42662815 |
| Haywood County, District 7 | 0.09601937 | 0.4429801 | 35.58400552 | -89.09356061 |
| Hamblen County, District 9 | 0.16863905 | 0.44190425 | 36.2604185 | -83.18658897 |
| Clay County, District 2 | 0.30688542 | 0.43786982 | 36.51372514 | -85.81125574 |
| Hamblen County, District 8 | 0.33472297 | 0.43679398 | 36.24129641 | -83.21818034 |
| Jackson County, District 5 | 0.33068854 | 0.43679398 | 36.31162243 | -85.81037925 |
| Coffee County, District 21 | 0.43652501 | 0.43652501 | 35.36032538 | -86.24453136 |
| Putnam County, District 9 | 0.37278107 | 0.43652501 | 36.17892663 | -85.5743928 |
| Marshall County, District 8 | 0.2105971 | 0.43544917 | 35.4471079 | -86.76389433 |
| Marshall County, District 3 | 0.14470145 | 0.43410436 | 35.42487822 | -86.76658925 |
| Madison County, District 4 | 0.42926304 | 0.43410436 | 35.70443461 | -88.85900875 |
| Montgomery County, District 12 | 0.40572889 | 0.43087682 | 36.59474934 | -87.41579399 |
| Sullivan County, District 4 | 0.29343733 | 0.42980097 | 36.51448232 | -82.23257743 |
| Jefferson County, District 9 | 0.31145777 | 0.42953201 | 36.06000018 | -83.41849315 |
| Blount County, District 9 | 0.27864443 | 0.42899408 | 35.8212744 | -83.9368462 |
| Sevier County, District 9 | 0.32894029 | 0.42657343 | 35.8706269 | -83.75997186 |
| Warren County, District 8 | 0.11928456 | 0.42522862 | 35.65835063 | -85.78604404 |
| Macon County, District 1 | 0.37305003 | 0.42442173 | 36.51848827 | -85.86728404 |
| Hawkins County, District 3 | 0.13569123 | 0.42388381 | 36.58417595 | -82.64708455 |
| Shelby County, District 11 | 0.22431415 | 0.42334588 | 35.10173344 | -90.10716703 |
| Shelby County, District 1 | 0.15519096 | 0.42119419 | 35.33711346 | -89.89444544 |
| DeKalb County, District 7 | 0.39080151 | 0.41931146 | 35.95411883 | -85.83045917 |
| Dyer County, District 9 | 0.26734804 | 0.41931146 | 36.07074775 | -89.38607706 |
| Dickson County, District 4 | 0.32477138 | 0.4190425 | 36.30763145 | -87.1663813 |
| Coffee County, District 13 | 0.267617 | 0.41850457 | 35.4047623 | -86.23884098 |
| Marshall County, District 7 | 0.2894029 | 0.41527703 | 35.47150169 | -86.78885616 |
| Warren County, District 9 | 0.24690694 | 0.41527703 | 35.60126482 | -85.90570715 |
| Decatur County, District 9 | 0.2267348 | 0.41500807 | 35.83481677 | -88.09045427 |
| Humphreys County, District 2 | 0.32813341 | 0.4133943 | 36.08284038 | -87.78333874 |
| Carter County, District 4 | 0.26062399 | 0.4133943 | 36.35344342 | -82.20534859 |
| Hamblen County, District 12 | 0.37506724 | 0.4112426 | 36.17063702 | -83.39998198 |
| McMinn County, District 3 | 0.14523938 | 0.41097364 | 35.43582706 | -84.58579374 |
| Cumberland County, District 1 | 0.32261969 | 0.41097364 | 35.95134117 | -85.04294136 |
| Overton County, District 4 | 0.26089295 | 0.41070468 | 36.2528774 | -85.39464826 |
| Bradley County, District 6 | 0.2204142 | 0.41043572 | 35.10208312 | -84.85846557 |
| White County, District 1 | 0.35018827 | 0.40989779 | 35.90038176 | -85.472217 |
| Jefferson County, District 7 | 0.1993007 | 0.40693921 | 36.00533842 | -83.5405368 |
| Gibson County, District 10 | 0.30016138 | 0.40693921 | 35.9782822 | -88.94539213 |
| Shelby County, District 4 | 0.29935449 | 0.40640129 | 35.06235834 | -89.77781888 |
| DeKalb County, District 4 | 0.32235073 | 0.40505648 | 35.97392807 | -85.79965784 |
| Fayette County, District 5 | 0.09884346 | 0.40478752 | 35.03267502 | -89.64176858 |
| Marion County, District 2 | 0.13340506 | 0.4042496 | 35.2364234 | -85.84250946 |
| Franklin County, District 7 | 0.20602474 | 0.4042496 | 35.33581464 | -86.21977977 |
| White County, District 3 | 0.20091447 | 0.4042496 | 35.94476014 | -85.46603421 |
| DeKalb County, District 6 | 0.21490048 | 0.40344271 | 35.91210211 | -85.86422073 |
| Sevier County, District 2 | 0.23910705 | 0.40317375 | 35.75893162 | -83.55299519 |
| Franklin County, District 2 | 0.36511565 | 0.40155998 | 35.21549555 | -86.1238058 |
| Rhea County, District 8 | 0.32732652 | 0.39967725 | 35.47524456 | -84.99965529 |
| Blount County, District 8 | 0.19526627 | 0.39940828 | 35.74916605 | -83.89210573 |
| Weakley County, District 5 | 0.37049489 | 0.39887036 | 36.35336936 | -88.85387189 |
| Roane County, District 2 | 0.27218935 | 0.3986014 | 35.90941702 | -84.59871182 |
| Grundy County, District 3 | 0.19970414 | 0.39779451 | 35.23880332 | -85.87005649 |
| Coffee County, District 6 | 0.27380312 | 0.39725659 | 35.45521146 | -86.0368344 |
| Weakley County, District 2 | 0.16541151 | 0.39725659 | 36.13250531 | -88.53738884 |
| Hamilton County, District 3 | 0.31710597 | 0.39564282 | 35.13804053 | -85.24317007 |
| Weakley County, District 8 | 0.10731576 | 0.39564282 | 36.35168933 | -88.8258377 |
| Anderson County, District 1 | 0.2119419 | 0.39564282 | 36.06520051 | -84.12364598 |
| Coffee County, District 16 | 0.39537386 | 0.39537386 | 35.40246893 | -86.21122287 |
| White County, District 7 | 0.26788596 | 0.3951049 | 35.96941572 | -85.49074027 |
| Rhea County, District 4 | 0.24341044 | 0.39402905 | 35.5221908 | -85.02085386 |
| Coffee County, District 4 | 0.36982249 | 0.39268424 | 35.45754548 | -86.06446631 |
| Sevier County, District 7 | 0.29128564 | 0.39080151 | 35.8754414 | -83.59014801 |
| Lawrence County, District 16 | 0.39026358 | 0.39026358 | 35.24447885 | -87.34500095 |
| Coffee County, District 2 | 0.38864981 | 0.38864981 | 35.45987304 | -86.0920999 |
| Lincoln County, District 5 | 0.302851 | 0.38838085 | 35.14059843 | -86.57848557 |
| Sevier County, District 3 | 0.30782679 | 0.38703604 | 35.88295692 | -83.44799882 |
| Anderson County, District 7 | 0.35664336 | 0.38676708 | 36.03197247 | -84.24173167 |
| Giles County, District 1 | 0.15599785 | 0.38515331 | 35.0045682 | -86.84498837 |
| Montgomery County, District 8 | 0.37210866 | 0.38192577 | 36.63919919 | -87.41070257 |
| Carter County, District 8 | 0.38138784 | 0.38138784 | 36.33458277 | -82.23722588 |
| Shelby County, District 10 | 0.32423346 | 0.38004303 | 35.11417941 | -89.91220391 |
| Carroll County, District 8 | 0.23897257 | 0.37950511 | 36.11025118 | -88.5395872 |
| Lincoln County, District 3 | 0.25080689 | 0.37923615 | 35.16282963 | -86.57574612 |
| Blount County, District 6 | 0.20642819 | 0.37923615 | 35.74111909 | -84.03395905 |
| Montgomery County, District 7 | 0.14281872 | 0.37923615 | 36.52807207 | -87.42341985 |
| Fentress County, District 1 | 0.32867133 | 0.37923615 | 36.4365524 | -84.94374939 |
| Humphreys County, District 3 | 0.27797203 | 0.37896719 | 36.08476482 | -87.81124863 |
| Shelby County, District 5 | 0.32840237 | 0.37869822 | 35.15733598 | -89.88104159 |
| Fentress County, District 2 | 0.2436794 | 0.37789134 | 36.45871964 | -84.94048137 |
| Benton County, District 1 | 0.18423884 | 0.37520172 | 35.85520695 | -88.06029918 |
| Anderson County, District 5 | 0.14093599 | 0.3746638 | 36.11227985 | -84.14446583 |
| Bedford County, District 4 | 0.1936525 | 0.3746638 | 35.54067981 | -86.52893131 |
| Dyer County, District 7 | 0.37412587 | 0.37412587 | 36.0484779 | -89.38802425 |
| Marshall County, District 5 | 0.26250672 | 0.37358795 | 35.45142948 | -86.81919659 |
| Coffee County, District 14 | 0.27218935 | 0.37305003 | 35.37564648 | -86.15886146 |
| Moore County, District 5 | 0.30433029 | 0.37278107 | 35.38483038 | -86.26929882 |
| Jefferson County, District 6 | 0.22969338 | 0.37278107 | 36.01116257 | -83.59603108 |
| Scott County, District 6 | 0.24731038 | 0.37197418 | 36.50773525 | -84.50767517 |
| DeKalb County, District 5 | 0.24475524 | 0.37170522 | 35.9493368 | -85.77485005 |
| Anderson County, District 6 | 0.1366326 | 0.36820871 | 36.01256969 | -84.27295962 |
| Carroll County, District 5 | 0.15599785 | 0.36820871 | 35.99015194 | -88.41109131 |
| Benton County, District 5 | 0.29410974 | 0.36767079 | 36.05915715 | -88.09508088 |
| Lawrence County, District 15 | 0.32302313 | 0.36740183 | 35.2465073 | -87.37259404 |
| Blount County, District 7 | 0.15599785 | 0.36713287 | 35.63872227 | -84.13433675 |
| Lawrence County, District 17 | 0.36659494 | 0.36659494 | 35.26468951 | -87.31488074 |
| McNairy County, District 3 | 0.095078 | 0.36632598 | 35.1717158 | -88.57556986 |
| Shelby County, District 6 | 0.29800968 | 0.36578806 | 35.18106486 | -89.90688721 |
| Bradley County, District 1 | 0.19943518 | 0.36525013 | 35.19605255 | -84.90050934 |
| Madison County, District 8 | 0.0739645 | 0.36498117 | 35.65314609 | -88.75208234 |
| Madison County, District 9 | 0.19795589 | 0.36498117 | 35.72670132 | -88.85691555 |
| Coffee County, District 10 | 0.25954814 | 0.36363636 | 35.6736361 | -86.23227466 |
| Coffee County, District 1 | 0.34050565 | 0.36309844 | 35.48208608 | -86.08920748 |
| Henry County, District 4 | 0.2027972 | 0.36282948 | 36.29807879 | -88.32370837 |
| Claiborne County, District 6 | 0.21490048 | 0.36202259 | 36.57894117 | -83.64407201 |
| Hamilton County, District 7 | 0.22861754 | 0.36175363 | 35.06130631 | -85.14267921 |
| Shelby County, District 2 | 0.2056213 | 0.36148467 | 35.03416557 | -89.66933811 |
| Giles County, District 5 | 0.15034965 | 0.36121571 | 35.17515316 | -87.01922656 |
| Decatur County, District 7 | 0.23749328 | 0.36067778 | 35.63082978 | -88.05578738 |
| White County, District 6 | 0.2119419 | 0.36067778 | 35.9274961 | -85.52469652 |
| Gibson County, District 12 | 0.23184508 | 0.36067778 | 35.87748672 | -88.75865226 |
| Gibson County, District 15 | 0.26654115 | 0.3598709 | 35.92030627 | -88.72652361 |
| Warren County, District 1 | 0.33270576 | 0.35852609 | 35.70514705 | -85.8077777 |
| Bradley County, District 2 | 0.22377622 | 0.35798817 | 35.25738974 | -84.83577539 |
| Franklin County, District 6 | 0.14093599 | 0.3577192 | 35.33351668 | -86.19218712 |
| Coffee County, District 3 | 0.33633674 | 0.35718128 | 35.43765949 | -86.09499063 |
| Loudon County, District 2 | 0.31818182 | 0.35718128 | 35.80766186 | -84.24848006 |
| Jackson County, District 4 | 0.29236148 | 0.35637439 | 36.22330146 | -85.56824722 |
| Hamilton County, District 2 | 0.22162453 | 0.35610543 | 35.14056073 | -85.27066267 |
| Perry County, District 2 | 0.19674556 | 0.35529855 | 35.62896194 | -88.02803767 |
| McNairy County, District 4 | 0.07006455 | 0.35529855 | 35.17345613 | -88.60316696 |
| Lawrence County, District 18 | 0.35529855 | 0.35529855 | 35.24244395 | -87.31740932 |
| Lawrence County, District 14 | 0.29182356 | 0.35502959 | 35.26672497 | -87.34248031 |
| Gibson County, District 14 | 0.35502959 | 0.35502959 | 35.92201009 | -88.75439623 |
| McMinn County, District 4 | 0.1366326 | 0.3542227 | 35.31690308 | -84.51980787 |
| Campbell County, District 4 | 0.11820871 | 0.35368478 | 36.27563588 | -84.20360477 |
| Hardin County, District 8 | 0.23614847 | 0.35126412 | 35.23938949 | -88.23546999 |
| Polk County, District 1 | 0.16272189 | 0.35099516 | 35.17236271 | -84.65307617 |
| Madison County, District 10 | 0.07880581 | 0.3507262 | 35.69767713 | -88.74783501 |
| Cumberland County, District 7 | 0.20925229 | 0.3485745 | 36.0426152 | -85.0578702 |
| Monroe County, District 1 | 0.15465304 | 0.34749866 | 35.6024239 | -84.44856342 |
| Gibson County, District 9 | 0.19311458 | 0.34722969 | 35.97993468 | -88.97329089 |
| Grainger County, District 5 | 0.19499731 | 0.34696073 | 36.33873099 | -83.28666837 |
| Hardin County, District 9 | 0.31750941 | 0.34615385 | 35.2171247 | -88.23773044 |
| Weakley County, District 7 | 0.18531469 | 0.34534696 | 36.33278581 | -88.88401057 |
| Carter County, District 6 | 0.20952125 | 0.34373319 | 36.33135846 | -82.20941404 |
| Lawrence County, District 1 | 0.24986552 | 0.34265734 | 35.07254394 | -87.4477068 |
| Gibson County, District 4 | 0.34211942 | 0.34211942 | 35.82077039 | -88.93204515 |
| Franklin County, District 4 | 0.14873588 | 0.3415815 | 35.16641114 | -86.07448914 |
| Henderson County, District 1 | 0.16325982 | 0.34131253 | 35.78807072 | -88.40330914 |
| Dyer County, District 10 | 0.34104357 | 0.34104357 | 36.05002658 | -89.41595819 |
| Obion County, District 2 | 0.0654922 | 0.34077461 | 36.43335196 | -89.07204559 |
| Anderson County, District 4 | 0.07100592 | 0.34077461 | 36.22580039 | -84.15485301 |
| Anderson County, District 8 | 0.24206563 | 0.3394298 | 36.02921413 | -84.21395168 |
| Rhea County, District 9 | 0.21436256 | 0.33781603 | 35.43857914 | -85.08884663 |
| Campbell County, District 2 | 0.26506186 | 0.33754707 | 36.35868164 | -84.13388059 |
| Hamilton County, District 9 | 0.14335664 | 0.33674018 | 35.0758311 | -85.05712921 |
| Lawrence County, District 9 | 0.21274879 | 0.33674018 | 35.28693456 | -87.31235069 |
| Shelby County, District 13 | 0.27622378 | 0.33647122 | 35.06381693 | -89.80540167 |
| Polk County, District 2 | 0.14416353 | 0.33566434 | 35.16969839 | -84.62559307 |
| Weakley County, District 9 | 0.21812803 | 0.33405056 | 36.27642845 | -88.69216414 |
| McNairy County, District 6 | 0.20871436 | 0.33324368 | 35.15118305 | -88.60531873 |
| Clay County, District 5 | 0.27662722 | 0.33162991 | 36.55113797 | -85.46589033 |
| Lawrence County, District 5 | 0.25739645 | 0.33136095 | 35.22223222 | -87.34752012 |
| Cumberland County, District 3 | 0.11820871 | 0.33136095 | 35.94619143 | -84.98738595 |
| Gibson County, District 6 | 0.1457773 | 0.33055406 | 35.84137353 | -88.90212261 |
| Chester County, District 6 | 0.32947821 | 0.32947821 | 35.44589275 | -88.66033447 |
| Grundy County, District 1 | 0.13542227 | 0.32840237 | 35.25863477 | -85.83955311 |
| Coffee County, District 7 | 0.15949435 | 0.32786444 | 35.5019698 | -86.05866389 |
| Obion County, District 4 | 0.21705218 | 0.32759548 | 36.43172309 | -89.04397685 |
| Lincoln County, District 6 | 0.17132867 | 0.32732652 | 35.16503825 | -86.60328862 |
| Giles County, District 6 | 0.2859064 | 0.32705756 | 35.20159798 | -87.07174981 |
| Scott County, District 5 | 0.09870898 | 0.32625067 | 36.59094604 | -84.43807184 |
| Lewis County, District 9 | 0.32598171 | 0.32598171 | 35.55172576 | -87.56144168 |
| Fentress County, District 3 | 0.17832168 | 0.32544379 | 36.38962427 | -84.92234183 |
| Clay County, District 3 | 0.30809575 | 0.32517483 | 36.55360338 | -85.49390641 |
| Loudon County, District 3 | 0.16030124 | 0.3246369 | 35.64150003 | -84.16197147 |
| Warren County, District 12 | 0.21449704 | 0.32356105 | 35.8256716 | -85.90386936 |
| Decatur County, District 3 | 0.26815492 | 0.32329209 | 35.6067089 | -88.03037022 |
| Sequatchie County, District 4 | 0.32302313 | 0.32302313 | 35.39725736 | -85.37418627 |
| Henry County, District 2 | 0.10408822 | 0.32221625 | 36.15300418 | -88.50723585 |
| Chester County, District 5 | 0.16137708 | 0.32140936 | 35.44416624 | -88.63263769 |
| Claiborne County, District 1 | 0.25174825 | 0.32087144 | 36.47881328 | -83.54698286 |
| Lawrence County, District 7 | 0.302851 | 0.32060247 | 35.22019787 | -87.31993643 |
| Hickman County, District 1 | 0.17509414 | 0.32006455 | 35.9742947 | -87.51430859 |
| Monroe County, District 2 | 0.20064551 | 0.31979559 | 35.52504653 | -84.34826376 |
| White County, District 5 | 0.24394836 | 0.31979559 | 35.81653769 | -85.54005079 |
| Humphreys County, District 1 | 0.20979021 | 0.31925767 | 36.08668272 | -87.83915992 |
| Cumberland County, District 6 | 0.06401291 | 0.31791286 | 35.8978846 | -84.71303527 |
| Warren County, District 3 | 0.28348575 | 0.31683701 | 35.66074284 | -85.81374186 |
| Clay County, District 1 | 0.25820334 | 0.31656805 | 36.61262701 | -85.65591553 |
| Claiborne County, District 7 | 0.21637977 | 0.31629909 | 36.58183126 | -83.67203365 |
| Henderson County, District 6 | 0.24045186 | 0.31468531 | 35.65273666 | -88.38888845 |
| Franklin County, District 5 | 0.07261969 | 0.31280258 | 35.21896361 | -85.90054409 |
| Fentress County, District 5 | 0.14981173 | 0.31280258 | 36.45352378 | -84.88455894 |
| Dyer County, District 1 | 0.13152232 | 0.31226466 | 36.00393704 | -89.3919152 |
| Campbell County, District 5 | 0.14806347 | 0.31226466 | 36.3836042 | -84.15828444 |
| Lincoln County, District 7 | 0.21221087 | 0.31118881 | 35.01831232 | -86.45470703 |
| Shelby County, District 3 | 0.18275955 | 0.31038193 | 35.20192399 | -89.87747804 |
| Gibson County, District 11 | 0.11592254 | 0.31011296 | 35.92370739 | -88.7822701 |
| Warren County, District 7 | 0.19176977 | 0.309844 | 35.71773288 | -85.69391641 |
| Stewart County, District 4 | 0.16917698 | 0.30876815 | 36.51111844 | -87.82131847 |
| Claiborne County, District 4 | 0.12399139 | 0.30849919 | 36.45668668 | -83.55066141 |
| Weakley County, District 4 | 0.1161915 | 0.30823023 | 36.26266283 | -88.83430897 |
| Haywood County, District 3 | 0.27703066 | 0.30796127 | 35.59361761 | -89.2601241 |
| Rhea County, District 3 | 0.22270038 | 0.30769231 | 35.595345 | -84.84212472 |
| Marion County, District 1 | 0.09373319 | 0.30688542 | 34.99990331 | -85.70720492 |
| Chester County, District 1 | 0.10220549 | 0.30473373 | 35.4218982 | -88.6347885 |
| Decatur County, District 5 | 0.28859602 | 0.30446477 | 35.63269112 | -88.08353844 |
| Clay County, District 4 | 0.23883808 | 0.30392684 | 36.54866603 | -85.4378761 |
| Scott County, District 4 | 0.1810113 | 0.30338892 | 36.48826893 | -84.53903174 |
| Henderson County, District 4 | 0.2493276 | 0.30311996 | 35.69725438 | -88.38442875 |
| Lawrence County, District 10 | 0.1788596 | 0.30258203 | 35.37386496 | -87.27457789 |
| Decatur County, District 8 | 0.2732652 | 0.30258203 | 35.65494471 | -88.08122062 |
| Jackson County, District 3 | 0.24300699 | 0.30231307 | 36.29474227 | -85.61486141 |
| Lawrence County, District 2 | 0.24731038 | 0.30204411 | 35.07053276 | -87.42017424 |
| Sevier County, District 10 | 0.24354492 | 0.30150619 | 35.84273781 | -83.70817066 |
| Lincoln County, District 8 | 0.21180742 | 0.30043034 | 34.9960808 | -86.45747634 |
| Lincoln County, District 1 | 0.12345347 | 0.29989242 | 35.00241579 | -86.8174965 |
| Wayne County, District 3 | 0.21328671 | 0.29881657 | 35.32066686 | -87.78187483 |
| Gibson County, District 3 | 0.16702528 | 0.2972028 | 35.81910805 | -88.90420572 |
| Lawrence County, District 8 | 0.23332437 | 0.29693384 | 35.24040258 | -87.28981916 |
| White County, District 4 | 0.18181818 | 0.29666487 | 35.91762028 | -85.41356373 |
| Jackson County, District 1 | 0.26143088 | 0.29639591 | 36.31936093 | -85.63973056 |
| Coffee County, District 11 | 0.24771382 | 0.29612695 | 35.55567364 | -86.16352648 |
| Lake County, District 1 | 0.26788596 | 0.29585799 | 36.41398614 | -89.52514016 |
| Scott County, District 7 | 0.10973642 | 0.29451318 | 36.51042425 | -84.53564233 |
| Bledsoe County, District 4 | 0.27084454 | 0.29343733 | 35.62880693 | -85.20164283 |
| Carter County, District 2 | 0.23076923 | 0.29289941 | 36.13847981 | -82.10314647 |
| Pickett County, District 3 | 0.22754169 | 0.29182356 | 36.56027472 | -85.06739224 |
| Dickson County, District 3 | 0.18692846 | 0.2915546 | 36.21486272 | -87.43087979 |
| Henderson County, District 5 | 0.154922 | 0.29020979 | 35.63225993 | -88.41887528 |
| Loudon County, District 6 | 0.17724583 | 0.29020979 | 35.82981922 | -84.24503628 |
| Union County, District 3 | 0.15976331 | 0.28994083 | 36.25844747 | -83.81004675 |
| Roane County, District 4 | 0.1225121 | 0.2880581 | 36.04294102 | -84.35287179 |
| Johnson County, District 1 | 0.2049489 | 0.2880581 | 36.60655008 | -81.67371262 |
| Bledsoe County, District 5 | 0.15545992 | 0.28752017 | 35.65099401 | -85.19848455 |
| Pickett County, District 1 | 0.24731038 | 0.28617536 | 36.56538494 | -85.12340652 |
| Chester County, District 4 | 0.09642281 | 0.28483055 | 35.42362425 | -88.66247726 |
| Carroll County, District 7 | 0.12910167 | 0.28483055 | 35.99192955 | -88.43898237 |
| Lawrence County, District 13 | 0.20199032 | 0.28402367 | 35.27279254 | -87.4252878 |
| Stewart County, District 6 | 0.21543841 | 0.28375471 | 36.48888356 | -87.82373939 |
| Claiborne County, District 3 | 0.19392146 | 0.28321678 | 36.44037564 | -83.6101492 |
| Lake County, District 2 | 0.2584723 | 0.28321678 | 36.38713248 | -89.44287026 |
| Sequatchie County, District 6 | 0.15465304 | 0.28267886 | 35.35535126 | -85.40795237 |
| Giles County, District 2 | 0.18988704 | 0.28106509 | 35.17935637 | -87.07434683 |
| Obion County, District 3 | 0.08996772 | 0.28106509 | 36.40782934 | -89.01797511 |
| Johnson County, District 6 | 0.13340506 | 0.28079613 | 36.4974264 | -81.89403455 |
| Overton County, District 3 | 0.18047337 | 0.27972028 | 36.20296533 | -85.09105076 |
| Haywood County, District 10 | 0.08741259 | 0.27918236 | 35.59519684 | -89.28788874 |
| Humphreys County, District 4 | 0.23278644 | 0.27918236 | 36.06444134 | -87.84156203 |
| Morgan County, District 2 | 0.11726735 | 0.27918236 | 35.951071 | -84.56425371 |
| Hardin County, District 1 | 0.25161377 | 0.27837547 | 35.21529881 | -88.21012614 |
| Lawrence County, District 4 | 0.20790748 | 0.27783755 | 35.19591099 | -87.2948878 |
| Warren County, District 10 | 0.18504572 | 0.27783755 | 35.73922206 | -85.94342791 |
| Union County, District 4 | 0.12264658 | 0.27783755 | 36.23916102 | -83.8414788 |
| Sequatchie County, District 5 | 0.14147391 | 0.27703066 | 35.35286363 | -85.3803781 |
| Perry County, District 1 | 0.2225659 | 0.27649274 | 35.60483507 | -88.00262992 |
| Campbell County, District 1 | 0.17643895 | 0.27568585 | 36.35589876 | -84.10598427 |
| Henderson County, District 7 | 0.20199032 | 0.27568585 | 35.4470085 | -88.32581661 |
| Wayne County, District 7 | 0.18114578 | 0.27541689 | 35.31873437 | -87.7542444 |
| Carroll County, District 3 | 0.09628833 | 0.27487897 | 36.04609986 | -88.23692442 |
| Haywood County, District 2 | 0.13125336 | 0.27461001 | 35.61589324 | -89.25815115 |
| Wayne County, District 5 | 0.17563206 | 0.27461001 | 35.30797543 | -87.92240263 |
| Lawrence County, District 6 | 0.20010758 | 0.27461001 | 35.11503508 | -87.41518651 |
| Coffee County, District 5 | 0.13206025 | 0.27461001 | 35.40610149 | -85.98741898 |
| Decatur County, District 6 | 0.27003765 | 0.27461001 | 35.61598056 | -88.16909206 |
| Bledsoe County, District 1 | 0.17684239 | 0.27407208 | 35.63133569 | -85.22931106 |
| Tipton County, District 2 | 0.06535772 | 0.27380312 | 35.54828703 | -89.65450482 |
| McNairy County, District 2 | 0.12426036 | 0.27218935 | 35.22615708 | -88.37577154 |
| Henderson County, District 3 | 0.20010758 | 0.27218935 | 35.68641207 | -88.21775753 |
| Bledsoe County, District 6 | 0.19526627 | 0.27218935 | 35.57935714 | -85.1526547 |
| Union County, District 1 | 0.20723507 | 0.27218935 | 36.25558896 | -83.78219955 |
| Pickett County, District 4 | 0.22767617 | 0.27192039 | 36.61734602 | -85.20104785 |
| Monroe County, District 4 | 0.16137708 | 0.27138246 | 35.6082234 | -84.27938395 |
| Tipton County, District 9 | 0.16352878 | 0.2711135 | 35.50668547 | -89.71369563 |
| Fayette County, District 1 | 0.14416353 | 0.27057558 | 35.24028234 | -89.34680854 |
| Overton County, District 2 | 0.20387305 | 0.26896181 | 36.42782653 | -85.34160903 |
| Lawrence County, District 12 | 0.16030124 | 0.26869285 | 35.33751501 | -87.3901609 |
| Decatur County, District 4 | 0.22001076 | 0.26842388 | 35.56592757 | -88.09048377 |
| Sequatchie County, District 3 | 0.25605164 | 0.26815492 | 35.39476193 | -85.34659798 |
| Bledsoe County, District 2 | 0.15088757 | 0.26815492 | 35.60154383 | -85.14948421 |
| Hawkins County, District 4 | 0.11632598 | 0.26788596 | 36.46035819 | -82.9534525 |
| Bledsoe County, District 3 | 0.17993545 | 0.267617 | 35.55716995 | -85.15582334 |
| Gibson County, District 8 | 0.08848843 | 0.26734804 | 35.84303632 | -88.92997015 |
| Sevier County, District 6 | 0.23830016 | 0.26600323 | 35.889897 | -83.72867169 |
| Gibson County, District 2 | 0.22243141 | 0.26465842 | 35.83970421 | -88.87427629 |
| Coffee County, District 9 | 0.14658419 | 0.26438946 | 35.64220931 | -86.12431307 |
| Gibson County, District 7 | 0.13313609 | 0.26412049 | 35.95435973 | -88.9195769 |
| Henry County, District 5 | 0.22942442 | 0.26412049 | 36.31852134 | -88.29343131 |
| Sequatchie County, District 9 | 0.14470145 | 0.26385153 | 35.37754906 | -85.40486526 |
| Sequatchie County, District 1 | 0.1936525 | 0.26358257 | 35.372566 | -85.3497027 |
| Stewart County, District 2 | 0.18544917 | 0.26250672 | 36.53335299 | -87.81889613 |
| Giles County, District 3 | 0.11686391 | 0.26223776 | 35.1814483 | -87.10190923 |
| Tipton County, District 1 | 0.14833244 | 0.2619688 | 35.57056989 | -89.65264824 |
| Roane County, District 6 | 0.1788596 | 0.26169984 | 35.88189523 | -84.54657804 |
| Lewis County, District 3 | 0.15640129 | 0.26143088 | 35.57397033 | -87.55897402 |
| Overton County, District 5 | 0.19634212 | 0.26116192 | 36.33661229 | -85.32626094 |
| Hardin County, District 6 | 0.1845078 | 0.26089295 | 35.23573026 | -88.18024678 |
| Sequatchie County, District 8 | 0.14550834 | 0.26089295 | 35.21468309 | -85.34386199 |
| Obion County, District 1 | 0.09817106 | 0.26089295 | 36.4902475 | -88.89734784 |
| Overton County, District 1 | 0.24583109 | 0.26089295 | 36.35879116 | -85.32310967 |
| Grundy County, District 2 | 0.19069392 | 0.26062399 | 35.3132565 | -85.69280643 |
| Dyer County, District 2 | 0.18410436 | 0.26035503 | 36.10902089 | -89.27035688 |
| Lincoln County, District 2 | 0.12547068 | 0.26008607 | 35.36510044 | -86.57863323 |
| Meigs County, District 2 | 0.18316299 | 0.26008607 | 35.45047748 | -84.97526796 |
| Benton County, District 3 | 0.17859064 | 0.25981711 | 36.03691025 | -88.09740699 |
| Hawkins County, District 5 | 0.18975256 | 0.25954814 | 36.41919068 | -82.98902834 |
| Haywood County, District 6 | 0.10207101 | 0.25793437 | 35.61430706 | -89.23037959 |
| Humphreys County, District 7 | 0.19849381 | 0.25685853 | 36.11562948 | -87.61097081 |
| Van Buren County, District 2 | 0.19741797 | 0.2563206 | 35.77704063 | -85.60164501 |
| Pickett County, District 2 | 0.2528241 | 0.25605164 | 36.54829679 | -85.18263087 |
| Monroe County, District 3 | 0.11013986 | 0.25605164 | 35.48344085 | -84.38264738 |
| Cumberland County, District 8 | 0.19176977 | 0.25578268 | 35.93428591 | -85.1017045 |
| Carter County, District 1 | 0.1366326 | 0.25524476 | 36.37229535 | -82.17345536 |
| Hardin County, District 3 | 0.18934911 | 0.25497579 | 35.03898765 | -88.25576683 |
| Jackson County, District 2 | 0.22619688 | 0.25497579 | 36.47222703 | -85.59029319 |
| Sequatchie County, District 2 | 0.16272189 | 0.25255514 | 35.41945344 | -85.37108766 |
| Stewart County, District 1 | 0.10866057 | 0.25255514 | 36.54395267 | -87.64798612 |
| Lauderdale County, District 1 | 0.13797741 | 0.25201721 | 35.65669956 | -89.58962143 |
| Moore County, District 2 | 0.18585261 | 0.25174825 | 35.31817082 | -86.27780318 |
| Van Buren County, District 5 | 0.19311458 | 0.25174825 | 35.72038375 | -85.46916573 |
| Lake County, District 3 | 0.20952125 | 0.25174825 | 36.25506386 | -89.48249577 |
| Lauderdale County, District 2 | 0.11592254 | 0.25147929 | 35.74278951 | -89.52645197 |
| Hardeman County, District 3 | 0.13340506 | 0.2506724 | 35.31830621 | -89.14538643 |
| Wayne County, District 4 | 0.20790748 | 0.25040344 | 35.1832691 | -87.74105833 |
| Stewart County, District 5 | 0.20441097 | 0.24852071 | 36.40565781 | -87.91749859 |
| Humphreys County, District 6 | 0.20817644 | 0.24825175 | 36.09535739 | -87.64135027 |
| Johnson County, District 3 | 0.18625605 | 0.24825175 | 36.4654263 | -81.81465904 |
| Fentress County, District 4 | 0.16514255 | 0.24744486 | 36.20040537 | -85.06317803 |
| Wayne County, District 6 | 0.19849381 | 0.24663798 | 35.02162929 | -87.67527597 |
| Hardin County, District 7 | 0.16178053 | 0.24636902 | 35.18751899 | -88.12961214 |
| Hawkins County, District 7 | 0.13394298 | 0.24610005 | 36.24836496 | -83.07530745 |
| Van Buren County, District 1 | 0.214766 | 0.24556213 | 35.74010295 | -85.43836519 |
| Johnson County, District 4 | 0.17859064 | 0.24556213 | 36.32740888 | -81.98287622 |
| Van Buren County, District 4 | 0.14658419 | 0.24475524 | 35.6759974 | -85.47532847 |
| Hardeman County, District 5 | 0.17724583 | 0.24448628 | 35.04607518 | -89.0865942 |
| Lauderdale County, District 5 | 0.1936525 | 0.24448628 | 35.6980008 | -89.94925815 |
| Hawkins County, District 6 | 0.11578806 | 0.24421732 | 36.28852604 | -83.23848337 |
| Wayne County, District 2 | 0.23063475 | 0.2436794 | 35.37852698 | -87.97064472 |
| Cumberland County, District 5 | 0.18881119 | 0.2436794 | 35.88992643 | -85.10809932 |
| Union County, District 5 | 0.14120495 | 0.2436794 | 36.3248579 | -83.79926178 |
| Crockett County, District 4 | 0.09386767 | 0.24287251 | 35.83543772 | -89.18265415 |
| Roane County, District 5 | 0.11753631 | 0.24260355 | 35.75758096 | -84.42488787 |
| Perry County, District 3 | 0.15667025 | 0.24233459 | 35.57311806 | -87.86632839 |
| Crockett County, District 11 | 0.12116729 | 0.24206563 | 35.71600934 | -89.05360973 |
| Meigs County, District 1 | 0.19983862 | 0.2415277 | 35.40868778 | -85.00927598 |
| Decatur County, District 2 | 0.20252824 | 0.24098978 | 35.50835392 | -88.23595139 |
| Obion County, District 5 | 0.15868747 | 0.24045186 | 36.49067078 | -89.29266854 |
| Lewis County, District 6 | 0.24018289 | 0.24018289 | 35.55370297 | -87.58915155 |
| Wayne County, District 1 | 0.23332437 | 0.23910705 | 35.3409875 | -87.75184032 |
| Lawrence County, District 3 | 0.18719742 | 0.23830016 | 35.08874138 | -87.36260448 |
| Meigs County, District 3 | 0.17482517 | 0.2380312 | 35.53141871 | -84.87951813 |
| McNairy County, District 5 | 0.12076385 | 0.23776224 | 35.08087731 | -88.55663726 |
| Lewis County, District 8 | 0.20763852 | 0.23695535 | 35.54974206 | -87.53373325 |
| Warren County, District 11 | 0.16231845 | 0.23587951 | 35.74715453 | -85.77407681 |
| Hardin County, District 2 | 0.16594944 | 0.23507262 | 35.17259357 | -88.24224739 |
| Houston County, District 3 | 0.19123185 | 0.23507262 | 36.3216278 | -87.67265223 |
| Carroll County, District 4 | 0.10516407 | 0.2345347 | 36.10496983 | -88.45578008 |
| Van Buren County, District 3 | 0.20293168 | 0.23426573 | 35.64637244 | -85.39535882 |
| Houston County, District 5 | 0.20589026 | 0.23265196 | 36.22483093 | -87.57064195 |
| Crockett County, District 8 | 0.11955352 | 0.23211404 | 35.73990714 | -89.0793862 |
| Perry County, District 4 | 0.19311458 | 0.23130715 | 35.79560148 | -87.84247176 |
| Benton County, District 2 | 0.09628833 | 0.23130715 | 36.08140365 | -88.0927534 |
| Moore County, District 4 | 0.190156 | 0.23103819 | 35.34494192 | -86.33016689 |
| Johnson County, District 2 | 0.2105971 | 0.23103819 | 36.46210409 | -81.78681727 |
| Hickman County, District 7 | 0.16299086 | 0.23050027 | 35.70344641 | -87.48860989 |
| Henry County, District 3 | 0.12587413 | 0.23023131 | 36.33895596 | -88.26313784 |
| Marion County, District 4 | 0.11995697 | 0.22996235 | 35.2688429 | -85.69880674 |
| Loudon County, District 4 | 0.12600861 | 0.22969338 | 35.69108807 | -84.43504635 |
| Hardin County, District 5 | 0.16783217 | 0.22942442 | 35.26709555 | -88.31607682 |
| Scott County, District 3 | 0.17549758 | 0.22807961 | 36.41911466 | -84.52125326 |
| Houston County, District 1 | 0.20898332 | 0.22781065 | 36.31670272 | -87.92704681 |
| Hancock County, District 4 | 0.10247445 | 0.22781065 | 36.54490448 | -83.10940437 |
| Roane County, District 3 | 0.1570737 | 0.22592792 | 35.96786041 | -84.50535776 |
| Haywood County, District 9 | 0.07100592 | 0.22592792 | 35.57134153 | -89.26209589 |
| Tipton County, District 4 | 0.19620764 | 0.22565896 | 35.42193723 | -89.80417079 |
| Gibson County, District 19 | 0.15115654 | 0.22565896 | 36.02446222 | -88.96915519 |
| Benton County, District 4 | 0.15371167 | 0.22538999 | 36.05729911 | -88.06717333 |
| Union County, District 6 | 0.12103281 | 0.22538999 | 36.21702212 | -83.84505946 |
| Decatur County, District 1 | 0.18934911 | 0.22243141 | 35.40454118 | -88.02363393 |
| Hamblen County, District 13 | 0.21463152 | 0.22216245 | 36.17358471 | -83.42778324 |
| Stewart County, District 7 | 0.14685315 | 0.21597633 | 36.40185288 | -87.86143728 |
| Lawrence County, District 11 | 0.15303927 | 0.21570737 | 35.42039415 | -87.29713948 |
| Lauderdale County, District 8 | 0.11484669 | 0.21516945 | 35.89257791 | -89.4016226 |
| Polk County, District 3 | 0.13905325 | 0.21463152 | 35.03182749 | -84.36776026 |
| Hickman County, District 6 | 0.19338354 | 0.2140936 | 35.76816361 | -87.45333802 |
| Lauderdale County, District 3 | 0.18477676 | 0.21247983 | 35.74430611 | -89.55427644 |
| Dyer County, District 4 | 0.11242604 | 0.21247983 | 36.11059707 | -89.29831073 |
| Lincoln County, District 4 | 0.17173211 | 0.21113502 | 35.07390159 | -86.58669441 |
| Hardeman County, District 1 | 0.12479828 | 0.21113502 | 35.30853083 | -88.97943675 |
| Lewis County, District 2 | 0.12856374 | 0.21086606 | 35.56999535 | -87.50354256 |
| Perry County, District 6 | 0.18881119 | 0.2105971 | 35.74919191 | -87.81946587 |
| Stewart County, District 3 | 0.17670791 | 0.21032813 | 36.57589787 | -87.78595426 |
| Benton County, District 6 | 0.1520979 | 0.20925229 | 36.32047716 | -87.98304816 |
| McNairy County, District 1 | 0.08364712 | 0.2084454 | 35.35509119 | -88.6412334 |
| Gibson County, District 18 | 0.1422808 | 0.20656267 | 36.08291124 | -88.82325711 |
| Lauderdale County, District 6 | 0.10193653 | 0.20548682 | 35.76658378 | -89.55238573 |
| Marshall County, District 4 | 0.13259817 | 0.20521786 | 35.32018445 | -86.86285288 |
| Hancock County, District 7 | 0.11027434 | 0.20521786 | 36.51269254 | -83.22868401 |
| Rhea County, District 1 | 0.08835395 | 0.20441097 | 35.74795606 | -84.79155645 |
| Franklin County, District 3 | 0.11874664 | 0.20414201 | 35.00702948 | -86.31729508 |
| Dyer County, District 5 | 0.10529855 | 0.20333513 | 36.10523008 | -89.60776273 |
| Gibson County, District 1 | 0.10099516 | 0.20306616 | 35.85522441 | -88.7607784 |
| Crockett County, District 3 | 0.08606778 | 0.20252824 | 35.81476395 | -89.21249817 |
| Grainger County, District 1 | 0.13327058 | 0.20252824 | 36.27382834 | -83.52432196 |
| Hancock County, District 3 | 0.18827327 | 0.20252824 | 36.55092718 | -83.16526268 |
| Tipton County, District 5 | 0.14604626 | 0.20225928 | 35.42338913 | -89.83188383 |
| Moore County, District 1 | 0.19540075 | 0.20225928 | 35.28279679 | -86.39376967 |
| Scott County, District 2 | 0.09561592 | 0.20225928 | 36.37748647 | -84.55594893 |
| Claiborne County, District 2 | 0.15182894 | 0.20225928 | 36.39901373 | -83.64535615 |
| Claiborne County, District 8 | 0.1048951 | 0.20091447 | 36.44976043 | -83.92083398 |
| Rhea County, District 2 | 0.1246638 | 0.20037655 | 35.64492231 | -84.89092138 |
| Crockett County, District 5 | 0.13340506 | 0.20010758 | 35.78929379 | -89.15882054 |
| Roane County, District 7 | 0.10866057 | 0.19795589 | 35.84024418 | -84.58101447 |
| Lewis County, District 7 | 0.14766003 | 0.19768693 | 35.52948072 | -87.56390791 |
| Crockett County, District 12 | 0.10677784 | 0.197149 | 35.77954052 | -88.99184157 |
| Tipton County, District 3 | 0.14443249 | 0.19688004 | 35.52897006 | -89.71185735 |
| Fayette County, District 2 | 0.12049489 | 0.19688004 | 35.2164323 | -89.32111681 |
| Weakley County, District 1 | 0.10623991 | 0.19634212 | 36.2969644 | -88.66199572 |
| Coffee County, District 8 | 0.10502959 | 0.19499731 | 35.61999845 | -86.12720133 |
| Morgan County, District 5 | 0.13878429 | 0.19499731 | 36.2082099 | -84.66648736 |
| Hancock County, District 2 | 0.12331899 | 0.19499731 | 36.52580615 | -83.14113473 |
| Hardeman County, District 2 | 0.09306079 | 0.19445939 | 35.26397646 | -88.9835278 |
| Gibson County, District 17 | 0.12143626 | 0.19445939 | 36.01444374 | -88.80168325 |
| Houston County, District 4 | 0.15895643 | 0.19445939 | 36.23463556 | -87.71044067 |
| Hancock County, District 6 | 0.19445939 | 0.19445939 | 36.5348084 | -83.22490807 |
| Marion County, District 3 | 0.14739107 | 0.19284562 | 35.0370501 | -85.6188196 |
| Fayette County, District 8 | 0.09440559 | 0.19284562 | 35.22877463 | -89.542182 |
| Lewis County, District 4 | 0.11430877 | 0.19150081 | 35.53145739 | -87.59160977 |
| Fayette County, District 4 | 0.09628833 | 0.19096288 | 35.22725453 | -89.51454497 |
| Giles County, District 4 | 0.10005379 | 0.19096288 | 35.30009053 | -86.89311392 |
| Carroll County, District 6 | 0.10112964 | 0.18961807 | 35.83258414 | -88.39884995 |
| Crockett County, District 7 | 0.18881119 | 0.18881119 | 35.78768456 | -89.13098774 |
| Carroll County, District 2 | 0.09844002 | 0.18692846 | 36.06748814 | -88.57190202 |
| Henry County, District 1 | 0.07920925 | 0.18692846 | 36.30166758 | -88.37971518 |
| Grainger County, District 2 | 0.09951587 | 0.18639053 | 36.36085102 | -83.28291814 |
| Meigs County, District 4 | 0.13555675 | 0.18612157 | 35.54574554 | -84.7933915 |
| Hardeman County, District 8 | 0.08781603 | 0.18558365 | 35.33080728 | -88.97738944 |
| Hardeman County, District 4 | 0.11780527 | 0.18396988 | 35.06835905 | -89.08458498 |
| Hardin County, District 10 | 0.18316299 | 0.18316299 | 35.21346645 | -88.18252316 |
| Gibson County, District 16 | 0.11377084 | 0.18289403 | 36.13742335 | -88.98675177 |
| Moore County, District 3 | 0.15330823 | 0.18128026 | 35.25605239 | -86.34142621 |
| Fayette County, District 7 | 0.14416353 | 0.1810113 | 35.03023322 | -89.19882988 |
| Morgan County, District 4 | 0.14819796 | 0.1810113 | 36.19661462 | -84.78125213 |
| Marion County, District 5 | 0.14456697 | 0.18074233 | 35.20736192 | -85.51208001 |
| Hardeman County, District 7 | 0.07920925 | 0.17751479 | 35.26232542 | -88.95588966 |
| Perry County, District 5 | 0.16863905 | 0.17697687 | 35.61761839 | -87.86156821 |
| Haywood County, District 8 | 0.08203335 | 0.17455621 | 35.50451055 | -89.26800437 |
| Fayette County, District 3 | 0.07100592 | 0.17455621 | 35.28641024 | -89.37058452 |
| Hancock County, District 5 | 0.11363636 | 0.17455621 | 36.58202835 | -83.24529456 |
| Haywood County, District 5 | 0.07988166 | 0.17374933 | 35.6174729 | -89.28592387 |
| Claiborne County, District 9 | 0.08257127 | 0.1732114 | 36.56546119 | -83.73159401 |
| Tipton County, District 8 | 0.14712211 | 0.17267348 | 35.40110064 | -89.83368458 |
| Lewis County, District 1 | 0.12694997 | 0.17159763 | 35.49322069 | -87.37255621 |
| Crockett County, District 6 | 0.10691232 | 0.16917698 | 35.8224262 | -88.95988579 |
| McNairy County, District 7 | 0.09642281 | 0.16890802 | 35.07736807 | -88.50151298 |
| Tipton County, District 6 | 0.0887574 | 0.16675632 | 35.54481949 | -90.0171669 |
| Houston County, District 2 | 0.16325982 | 0.16325982 | 36.3274582 | -87.75664257 |
| Grainger County, District 3 | 0.09709521 | 0.16272189 | 36.18531066 | -83.53900991 |
| Weakley County, District 3 | 0.12385691 | 0.16272189 | 36.14800024 | -88.78895298 |
| Hardeman County, District 6 | 0.13192577 | 0.16137708 | 35.05179163 | -88.80893913 |
| Union County, District 2 | 0.12963959 | 0.16137708 | 36.21131478 | -83.78939504 |
| Grainger County, District 4 | 0.09709521 | 0.16110812 | 36.29887601 | -83.54849669 |
| Carroll County, District 1 | 0.08472297 | 0.15868747 | 35.83966779 | -88.51018951 |
| Houston County, District 7 | 0.1070468 | 0.15384615 | 36.29350562 | -87.59115808 |
| Houston County, District 6 | 0.14362561 | 0.1514255 | 36.34581246 | -87.69819568 |
| Chester County, District 2 | 0.07799892 | 0.1514255 | 35.48522649 | -88.57291042 |
| Henderson County, District 2 | 0.06334051 | 0.15034965 | 35.58951385 | -88.45105519 |
| Claiborne County, District 5 | 0.07988166 | 0.1492738 | 36.548106 | -83.56387914 |
| Morgan County, District 3 | 0.11390533 | 0.14550834 | 36.08672928 | -84.57188799 |
| Cumberland County, District 9 | 0.0922539 | 0.14470145 | 36.00493767 | -84.89428682 |
| Lauderdale County, District 4 | 0.10946746 | 0.14389457 | 35.72051183 | -89.52834968 |
| Dyer County, District 3 | 0.08418505 | 0.14389457 | 36.08517125 | -89.24439458 |
| Lewis County, District 5 | 0.11646046 | 0.13905325 | 35.41428088 | -87.52089809 |
| Obion County, District 6 | 0.07100592 | 0.13878429 | 36.24103165 | -89.23043727 |
| Crockett County, District 2 | 0.04397526 | 0.13824637 | 35.90702197 | -89.26027559 |
| Crockett County, District 9 | 0.11565358 | 0.13824637 | 35.74152853 | -89.10720043 |
| Loudon County, District 7 | 0.09521248 | 0.13824637 | 35.74118685 | -84.25879936 |
| Scott County, District 1 | 0.07826789 | 0.1366326 | 36.34184508 | -84.41979379 |
| Union County, District 7 | 0.12506724 | 0.13555675 | 36.18631443 | -83.76516858 |
| Chester County, District 3 | 0.05876815 | 0.13286713 | 35.387614 | -88.80514134 |
| Lauderdale County, District 7 | 0.06764389 | 0.12963959 | 35.83341423 | -89.54670694 |
| Crockett County, District 1 | 0.08579882 | 0.12829478 | 35.85930521 | -89.20851171 |
| Meigs County, District 5 | 0.09413663 | 0.12479828 | 35.62917912 | -84.72497942 |
| Hancock County, District 1 | 0.07261969 | 0.12452932 | 36.49356234 | -83.26037051 |
| Morgan County, District 1 | 0.08687466 | 0.12318451 | 36.03703788 | -84.52301585 |
| Morgan County, District 6 | 0.08216783 | 0.11861216 | 36.10620928 | -84.5407062 |
| Tipton County, District 7 | 0.05930608 | 0.11807423 | 35.41901393 | -89.74874789 |
| Fayette County, District 6 | 0.08176439 | 0.11780527 | 35.200369 | -89.4335529 |
