## Supplementary material for "Mapping long-distance jump dispersal risk by leveraging transport infrastructure data and sampling bias correction: a case study with the invasive spotted lanternfly (*Lycorma delicatula*)": S16 Virginia risk

*Supplemental material S16.*  Mapping long-distance jump dispersal risk by leveraging transport infrastructure data and sampling bias correction: a case study with the invasive spotted lanternfly (*Lycorma delicatula*)
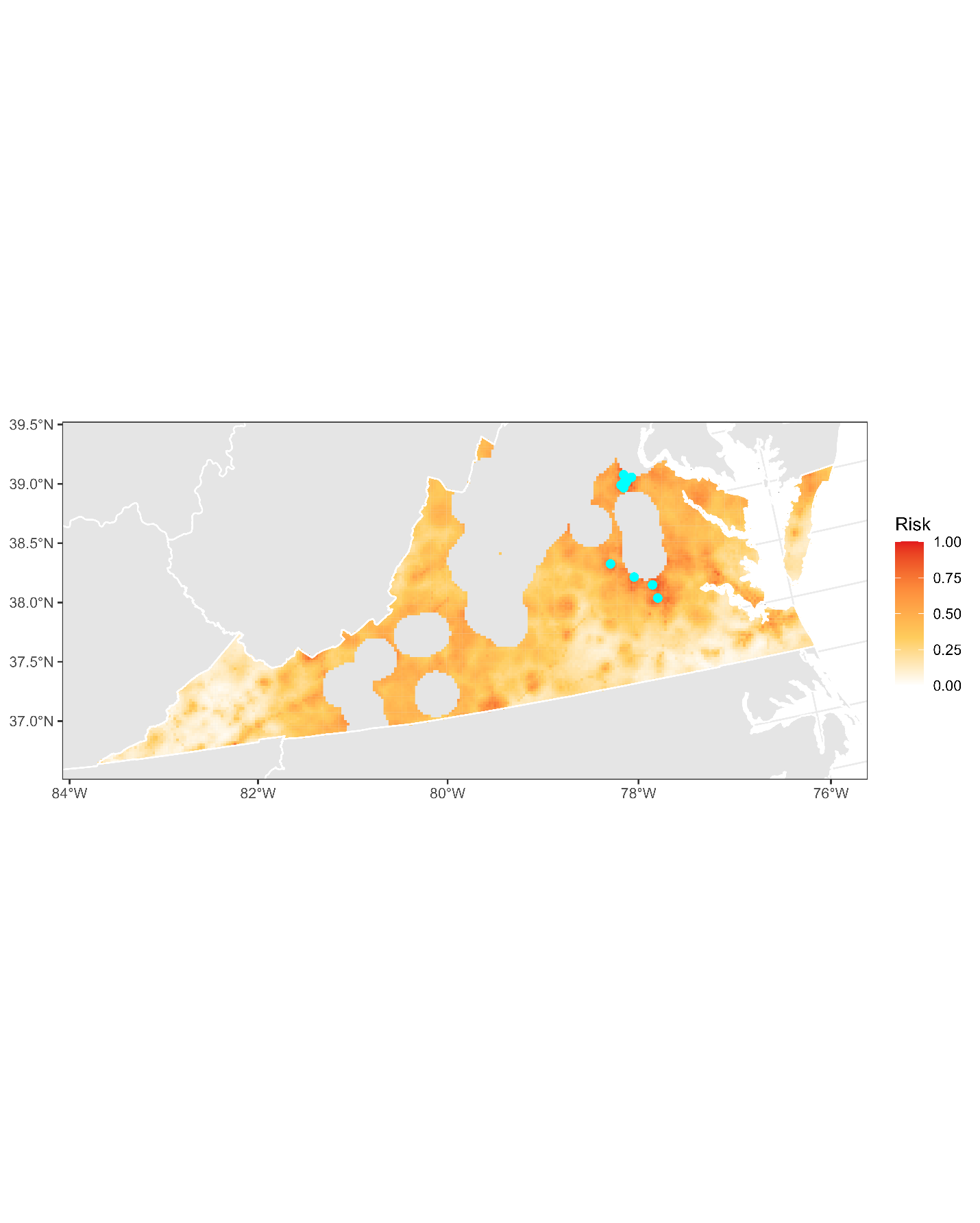

**Figure S16.1:** Projected risk of colonization by *L. delicatula* jump dispersal for Virginia. Risk projections are based on an ensemble random forest model including distances from existing *L. delicatula* populations. Note that areas falling less than 20 km from existing *L. delicatula* populations are not included in the map because any colonization there would be categorized as diffusive spread based on our methodology (regardless of the actual source of the resulting population). Darker, redder colors indicate increased risk of colonization by jump dispersal. Cyan points show the top ten highest risk locations in the state.

**Table S16.1:** Ranked list of Virginia county subdivisions sorted by the maximum projected risk of colonization by jump dispersal occurring there. For each subdivision, we identified the 2.5 x 2.5 km grid cell centroid included there with the highest projected risk of colonization by jump dispersal and report that location and risk value.

| **County, subdivision** | **Median risk value** | **Maximum risk value** | **Latitude of maximum risk** | **Longitude of maximum risk** |
| --- | --- | --- | --- | --- |
| Spotsylvania County, Courtland district | 0.69634212 | 0.88192577 | 38.30673434 | -77.52946483 |
| Spotsylvania County, Berkeley district | 0.63232921 | 0.88138784 | 38.19730922 | -77.55729488 |
| Chesterfield County, Bermuda district | 0.78214094 | 0.86498117 | 37.3401112 | -77.453088 |
| Chesterfield County, Clover Hill district | 0.72485207 | 0.85718128 | 37.43591639 | -77.63236884 |
| Spotsylvania County, Lee Hill district | 0.77003765 | 0.85126412 | 38.27190102 | -77.45012147 |
| Spotsylvania County, Battlefield district | 0.81629909 | 0.83862292 | 38.23677292 | -77.51788035 |
| Powhatan County, District 3 | 0.70817644 | 0.82759548 | 37.58373116 | -77.8573561 |
| Spotsylvania County, Livingston district | 0.53106509 | 0.81871974 | 38.22349425 | -77.58002317 |
| Spotsylvania County, Salem district | 0.76519634 | 0.81871974 | 38.24108013 | -77.5461728 |
| Petersburg city, Petersburg city | 0.72027972 | 0.8154922 | 37.22202724 | -77.42486227 |
| Fredericksburg city, Fredericksburg city | 0.71718666 | 0.81387843 | 38.28054012 | -77.50672881 |
| Stafford County, Hartwood district | 0.64483593 | 0.81253362 | 38.35480949 | -77.5466459 |
| Chesterfield County, Matoaca district | 0.58700914 | 0.80661646 | 37.39640956 | -77.67129806 |
| Dinwiddie County, District 3 | 0.52716514 | 0.80607854 | 37.20447068 | -77.45827852 |
| Stafford County, George Washington district | 0.68746638 | 0.79666487 | 38.2675717 | -77.42182279 |
| Bristol city, Bristol city | 0.4704142 | 0.79558903 | 36.59958424 | -82.18837874 |
| Chesterfield County, Dale district | 0.66581495 | 0.79343733 | 37.4011439 | -77.55390657 |
| York County, District 1 | 0.29532006 | 0.79182356 | 37.31735631 | -76.73255039 |
| Powhatan County, District 1 | 0.59803658 | 0.78778913 | 37.57949516 | -77.82929644 |
| Dinwiddie County, District 2 | 0.66581495 | 0.7802582 | 37.20879271 | -77.48617575 |
| Tazewell County, Eastern district | 0.45024207 | 0.77972028 | 37.23413523 | -81.26431073 |
| Prince George County, Rives district | 0.654922 | 0.77272727 | 37.18276409 | -77.31884104 |
| Spotsylvania County, Chancellor district | 0.54101668 | 0.76869285 | 38.29344987 | -77.59166475 |
| Dinwiddie County, District 1 | 0.37385691 | 0.76600323 | 37.18690479 | -77.49167904 |
| Colonial Heights city, Colonial Heights city | 0.73695535 | 0.75981711 | 37.28333986 | -77.38035375 |
| Prince George County, Bland district | 0.6253362 | 0.75658956 | 37.30087439 | -77.34688375 |
| James City County, Stonehouse district | 0.49905863 | 0.75605164 | 37.34369812 | -76.75471883 |
| Danville city, Danville city | 0.68746638 | 0.75551372 | 36.54531051 | -79.45761039 |
| Henrico County, Varina district | 0.5922539 | 0.74072082 | 37.39274591 | -77.35260542 |
| James City County, Berkeley district | 0.54074771 | 0.7105971 | 37.25628366 | -76.77761133 |
| Bland County, District 4 | 0.46745562 | 0.70763852 | 37.24580828 | -81.1755882 |
| Chesterfield County, Midlothian district | 0.67469069 | 0.70118343 | 37.49253344 | -77.70544482 |
| Hopewell city, Hopewell city | 0.62802582 | 0.69472835 | 37.29216004 | -77.29103982 |
| Fluvanna County, Columbia district | 0.52447552 | 0.69230769 | 37.9355714 | -78.23763981 |
| Louisa County, Green Spring district | 0.68961807 | 0.68961807 | 37.97940424 | -78.22696116 |
| Richmond County, District 5 | 0.54760624 | 0.68423884 | 37.98691954 | -76.78993045 |
| Goochland County, District 5 | 0.49193115 | 0.68289403 | 37.65433243 | -77.72329072 |
| Accomack County, District 3 | 0.54115116 | 0.68074233 | 37.9845101 | -75.52749121 |
| Goochland County, District 4 | 0.57961269 | 0.67993545 | 37.67622754 | -77.71782115 |
| New Kent County, District 3 | 0.4887036 | 0.67859064 | 37.50199402 | -77.0336308 |
| Richmond County, District 3 | 0.57383002 | 0.67724583 | 37.9650685 | -76.79569812 |
| King George County, James Monroe district | 0.5274341 | 0.67616998 | 38.28082433 | -77.21249904 |
| Powhatan County, District 5 | 0.46167294 | 0.6753631 | 37.61408769 | -77.90807618 |
| Richmond County, District 2 | 0.58068854 | 0.67348037 | 37.98243819 | -76.76178056 |
| New Kent County, District 5 | 0.53429263 | 0.66756321 | 37.49756825 | -77.00565011 |
| Pittsylvania County, Westover district | 0.47928994 | 0.66568047 | 36.5491724 | -79.48537184 |
| Norfolk city, Norfolk city | 0.45535234 | 0.66487359 | 36.85512191 | -76.27497447 |
| Chesapeake city, Chesapeake city | 0.15088757 | 0.6605702 | 36.81145235 | -76.2866324 |
| Montgomery County, District C-01 | 0.49825175 | 0.65949435 | 37.15723964 | -80.30092689 |
| Bedford County, District 5 | 0.4338354 | 0.65868747 | 37.39314343 | -79.52762118 |
| Powhatan County, District 4 | 0.44728349 | 0.654922 | 37.51802441 | -77.87361122 |
| Prince George County, Blackwater district | 0.51075847 | 0.65384615 | 37.15215843 | -77.26865498 |
| New Kent County, District 4 | 0.56374395 | 0.65196342 | 37.49336395 | -77.12322743 |
| New Kent County, District 1 | 0.58821947 | 0.65115654 | 37.5282791 | -77.0559611 |
| King George County, James Madison district | 0.57705756 | 0.64981173 | 38.25457161 | -77.18988415 |
| Powhatan County, District 2 | 0.51855836 | 0.64712211 | 37.50105628 | -77.76148745 |
| Bedford County, District 7 | 0.50820334 | 0.64685315 | 37.37116928 | -79.53253627 |
| Westmoreland County, Cople district | 0.51802044 | 0.64658419 | 38.00428758 | -76.75600148 |
| Caroline County, Port Royal district | 0.45642819 | 0.64281872 | 38.15817067 | -77.44979808 |
| Halifax County, District 6 | 0.2084454 | 0.63959118 | 36.67630503 | -78.9408535 |
| Dinwiddie County, District 4 | 0.29935449 | 0.63824637 | 37.10365816 | -77.5415211 |
| New Kent County, District 2 | 0.59131253 | 0.63770845 | 37.47589229 | -77.15683688 |
| Charlotte County, Walton district | 0.3542227 | 0.63474987 | 37.06271318 | -78.47647899 |
| King George County, Shiloh district | 0.56213018 | 0.63474987 | 38.25018239 | -77.16160759 |
| Pittsylvania County, Dan River district | 0.32571275 | 0.63313609 | 36.62158794 | -79.35469964 |
| Prince Edward County, District 301 | 0.26681011 | 0.62937063 | 37.08055066 | -78.44336736 |
| Cumberland County, District 5 | 0.49488973 | 0.6253362 | 37.32587692 | -78.41381384 |
| James City County, Jamestown district | 0.59897795 | 0.6253362 | 37.27365165 | -76.74401232 |
| Prince Edward County, District 201 | 0.25242066 | 0.62479828 | 37.29572871 | -78.36308452 |
| Tazewell County, Northern district | 0.26923077 | 0.62452932 | 37.26305669 | -81.31614794 |
| Accomack County, District 2 | 0.47458311 | 0.62318451 | 37.87555796 | -75.55815906 |
| Prince Edward County, District 101 | 0.60462614 | 0.62291555 | 37.2738026 | -78.36833635 |
| Bland County, District 2 | 0.34722969 | 0.61861216 | 37.08112361 | -81.12233051 |
| Goochland County, District 3 | 0.56280258 | 0.61807423 | 37.69747064 | -77.85830292 |
| Grayson County, Old Town district | 0.53792361 | 0.61646046 | 36.63093024 | -80.95506558 |
| Accomack County, District 5 | 0.4373319 | 0.61538462 | 37.85376636 | -75.56428177 |
| Grayson County, Providence district | 0.4408284 | 0.61511565 | 36.68203008 | -81.00193059 |
| Richmond County, District 1 | 0.37345347 | 0.61403981 | 37.93425319 | -76.74519794 |
| Pittsylvania County, Banister district | 0.31710597 | 0.61323292 | 36.64356217 | -79.34978158 |
| Hanover County, South Anna district | 0.50941366 | 0.61323292 | 37.72001665 | -77.70687228 |
| Newport News city, Newport News city | 0.46637977 | 0.61188811 | 37.07099749 | -76.47815665 |
| Virginia Beach city, Virginia Beach city | 0.20615923 | 0.61000538 | 36.83995668 | -76.04762752 |
| Fluvanna County, Fork Union district | 0.50322754 | 0.60973642 | 37.75608892 | -78.25207738 |
| Hampton city, Hampton city | 0.46584185 | 0.6070468 | 37.03547751 | -76.40062075 |
| Prince Edward County, District 701 | 0.60381926 | 0.60381926 | 37.30394885 | -78.41905285 |
| Henry County, Iriswood district | 0.44513179 | 0.60274341 | 36.67539934 | -79.74393984 |
| Prince Edward County, District 501 | 0.34238838 | 0.60274341 | 37.28202039 | -78.42428878 |
| James City County, Powhatan district | 0.49704142 | 0.59897795 | 37.40084228 | -76.82703309 |
| King William County, Acquinton district | 0.51398601 | 0.59870898 | 37.70337916 | -77.15657874 |
| Orange County, District 4 | 0.47552448 | 0.5979021 | 38.34524014 | -77.7844647 |
| Fluvanna County, Palmyra district | 0.51721356 | 0.59628833 | 37.91779481 | -78.27118447 |
| Botetourt County, Buchanan district | 0.37520172 | 0.59467456 | 37.4570576 | -79.8318475 |
| Franklin County, Rocky Mount district | 0.49448628 | 0.59440559 | 37.00974695 | -79.87198169 |
| Hanover County, Henry district | 0.51398601 | 0.59332975 | 37.65091138 | -77.25762163 |
| Portsmouth city, Portsmouth city | 0.40613233 | 0.5887574 | 36.8033709 | -76.3754873 |
| Prince George County, Templeton district | 0.43249059 | 0.58337816 | 37.01205167 | -77.39097203 |
| Westmoreland County, Montross district | 0.49193115 | 0.58230231 | 38.0524706 | -76.77260693 |
| Wythe County, Black Lick district | 0.42092523 | 0.58041958 | 36.89659282 | -81.2739548 |
| Louisa County, Mountain Road district | 0.46503497 | 0.57988166 | 37.76740517 | -77.87012481 |
| Charles City County, District 1 | 0.46342119 | 0.57934373 | 37.45402151 | -77.16245693 |
| Alleghany County, Jackson River district | 0.44055944 | 0.57934373 | 37.80192804 | -79.87205189 |
| Charles City County, District 2 | 0.42415277 | 0.57638515 | 37.41453197 | -77.05624401 |
| Bedford County, District 6 | 0.46221087 | 0.57504034 | 37.34919478 | -79.53744845 |
| Orange County, District 5 | 0.5478752 | 0.57019903 | 38.34098783 | -77.75611819 |
| Suffolk city, Suffolk city | 0.13017751 | 0.56885422 | 36.76062644 | -76.53120619 |
| Bedford County, District 2 | 0.50457235 | 0.56885422 | 37.2393167 | -79.56196591 |
| Goochland County, District 2 | 0.46866595 | 0.56804734 | 37.72782589 | -77.90910133 |
| Hanover County, Cold Harbor district | 0.52555137 | 0.56508876 | 37.65084414 | -77.11175709 |
| King William County, West Point district | 0.4436525 | 0.5648198 | 37.53196903 | -76.79269455 |
| Culpeper County, Stevensburg district | 0.5147929 | 0.56347499 | 38.37624699 | -77.68837729 |
| Montgomery County, District A-01 | 0.48345885 | 0.56293706 | 37.28191148 | -80.38983114 |
| Covington city, Covington city | 0.53429263 | 0.5626681 | 37.77296503 | -79.99469869 |
| Prince Edward County, District 601 | 0.45131791 | 0.56239914 | 37.31214313 | -78.47503352 |
| Williamsburg city, Williamsburg city | 0.54599247 | 0.56024744 | 37.28650979 | -76.68251559 |
| Louisa County, Mineral district | 0.51210328 | 0.55997848 | 38.10437005 | -77.84477977 |
| Franklin County, Union Hall district | 0.4675901 | 0.5591716 | 37.05297234 | -79.68978439 |
| King and Queen County, Buena Vista district | 0.31804734 | 0.55513717 | 37.50563575 | -76.77045643 |
| Northampton County, District 5 | 0.22135557 | 0.55513717 | 37.53335536 | -75.82921366 |
| James City County, Roberts district | 0.41608392 | 0.55433029 | 37.24730722 | -76.7218685 |
| Montgomery County, District B-01 | 0.52165143 | 0.55379236 | 37.18291251 | -80.32429211 |
| Prince Edward County, District 801 | 0.55298548 | 0.55298548 | 37.29984201 | -78.39106714 |
| Halifax County, District 7 | 0.18208714 | 0.55056482 | 36.69027783 | -78.88023811 |
| Essex County, Tappahannock district | 0.51909629 | 0.55029586 | 37.90398291 | -76.84110204 |
| Giles County, Western district | 0.46369016 | 0.54922001 | 37.33589941 | -80.811361 |
| Tazewell County, Northwestern district | 0.18692846 | 0.54895105 | 37.09654732 | -81.80874009 |
| Tazewell County, Western district | 0.2915546 | 0.54868209 | 37.09322478 | -81.78066332 |
| Halifax County, District 5 | 0.29209252 | 0.54706832 | 36.76412794 | -78.92066496 |
| Westmoreland County, Washington district | 0.4873588 | 0.54626143 | 38.2544226 | -77.04282174 |
| Patrick County, Dan River district | 0.45884884 | 0.54572351 | 36.56655692 | -80.4539188 |
| Halifax County, District 4 | 0.38999462 | 0.54464766 | 36.76811347 | -78.94847933 |
| Giles County, Central district | 0.34454008 | 0.54276493 | 37.32521912 | -80.72700625 |
| Smyth County, Atkins district | 0.43033889 | 0.54195804 | 36.88484615 | -81.36224817 |
| Halifax County, District 3 | 0.29343733 | 0.54168908 | 36.69826146 | -78.93581079 |
| Lunenburg County, Rehoboth district | 0.17697687 | 0.54088219 | 37.05861921 | -78.44858049 |
| Charles City County, District 3 | 0.36040882 | 0.54034427 | 37.41011129 | -77.02829489 |
| Washington County, Harrison district | 0.45387305 | 0.54034427 | 36.72473271 | -81.90813673 |
| Giles County, Eastern district | 0.42227004 | 0.53953739 | 37.31120529 | -80.44135985 |
| King George County, Dahlgren district | 0.45373857 | 0.53926842 | 38.28070236 | -77.06538401 |
| Campbell County, Altavista district | 0.37587413 | 0.53899946 | 37.13469751 | -79.2967877 |
| Mecklenburg County, District 4 | 0.29276493 | 0.5387305 | 36.7156907 | -78.12724799 |
| Floyd County, Indian Valley district | 0.46611081 | 0.53846154 | 36.95536673 | -80.48730459 |
| Pittsylvania County, Chatham-Blairs district | 0.37520172 | 0.53630984 | 36.69139802 | -79.36774478 |
| Grayson County, Elk Creek district | 0.43114578 | 0.53442711 | 36.62291011 | -81.0709627 |
| Craig County, New Castle district | 0.47875202 | 0.53335126 | 37.49450172 | -80.11318812 |
| Franklin County, Gills Creek district | 0.45427649 | 0.53254438 | 37.09693825 | -79.68008579 |
| Halifax County, District 8 | 0.53254438 | 0.53254438 | 36.7162277 | -78.90296946 |
| King William County, Mangohick district | 0.46180742 | 0.53254438 | 37.79963832 | -77.1901978 |
| Bedford County, District 3 | 0.46530393 | 0.53146853 | 37.30524466 | -79.54726412 |
| Amelia County, District 2 | 0.39725659 | 0.53146853 | 37.36047131 | -77.88344807 |
| Craig County, Craig City district | 0.49959656 | 0.53093061 | 37.53477691 | -80.07551969 |
| Prince George County, Brandon district | 0.37318451 | 0.53066165 | 37.27026158 | -77.15148733 |
| Rockingham County, Plains district | 0.44042496 | 0.53066165 | 38.69349691 | -78.93657171 |
| Franklin County, Blackwater district | 0.44109736 | 0.52931684 | 36.98077704 | -79.99332926 |
| Bedford County, District 1 | 0.48063475 | 0.52877891 | 37.21066091 | -79.68380718 |
| Floyd County, Locust Grove district | 0.46180742 | 0.52528241 | 37.02880373 | -80.18434163 |
| Appomattox County, Courthouse district | 0.42630446 | 0.5239376 | 37.36477518 | -78.83920623 |
| Craig County, Craig Creek district | 0.48251748 | 0.52366864 | 37.49078663 | -80.08504132 |
| Montgomery County, District D-01 | 0.49556213 | 0.52259279 | 37.06921082 | -80.31959711 |
| Henry County, Ridgeway district | 0.43464228 | 0.52097902 | 36.54345371 | -79.77272665 |
| Franklin County, Blue Ridge district | 0.47283486 | 0.51936525 | 36.92968353 | -80.11929631 |
| Essex County, South district | 0.44096288 | 0.51936525 | 37.89951328 | -76.81298065 |
| Floyd County, Court House district | 0.48574502 | 0.51694459 | 36.93715593 | -80.34752007 |
| Amelia County, District 4 | 0.47095212 | 0.51613771 | 37.38660049 | -77.90603531 |
| Essex County, Central district | 0.47807961 | 0.51425498 | 37.8691897 | -76.90881182 |
| Craig County, Simmonsville district | 0.48372781 | 0.51344809 | 37.3295753 | -80.40861546 |
| Floyd County, Little River district | 0.47444863 | 0.51210328 | 37.0068434 | -80.36154638 |
| Hanover County, Beaverdam district | 0.50618612 | 0.51210328 | 37.8294832 | -77.67944338 |
| Louisa County, Jackson district | 0.49919311 | 0.5112964 | 38.02169405 | -77.74824827 |
| Augusta County, Pastures district | 0.38300161 | 0.51022055 | 38.14007008 | -79.35876268 |
| Louisa County, Cuckoo district | 0.50268962 | 0.50887574 | 38.0478442 | -77.7709902 |
| Alleghany County, Clifton Forge East district | 0.48050027 | 0.50753093 | 37.81636327 | -79.81069251 |
| Alleghany County, Sharon district | 0.42522862 | 0.50672405 | 37.8594836 | -79.6264709 |
| Nottoway County, District 4 | 0.38327058 | 0.50645508 | 37.10179847 | -77.97589586 |
| York County, District 5 | 0.45736955 | 0.50591716 | 37.11012354 | -76.43876885 |
| Accomack County, District 4 | 0.46570737 | 0.50511027 | 37.86801636 | -75.62033753 |
| Wythe County, West Wytheville district | 0.46987628 | 0.50484131 | 36.97439917 | -81.17243041 |
| Wythe County, Speedwell district | 0.38649812 | 0.50242066 | 36.85250127 | -81.28269077 |
| Smyth County, Rye Valley district | 0.33862292 | 0.50107585 | 36.85593976 | -81.31065488 |
| Pittsylvania County, Tunstall district | 0.41258741 | 0.50026896 | 36.55403706 | -79.68459872 |
| Northumberland County, District 5 | 0.47283486 | 0.49973104 | 37.88343842 | -76.28967045 |
| Tazewell County, Southern district | 0.31038193 | 0.49865519 | 37.11051662 | -81.54754009 |
| Fluvanna County, Cunningham district | 0.39456697 | 0.49784831 | 37.88221389 | -78.33822534 |
| Appomattox County, Appomattox River district | 0.36847768 | 0.49677246 | 37.3607654 | -78.81117471 |
| Bland County, District 1 | 0.31468531 | 0.49569661 | 37.06602753 | -81.18282635 |
| Amelia County, District 5 | 0.40693921 | 0.49515869 | 37.28265846 | -77.67064067 |
| Franklin County, Snow Creek district | 0.41608392 | 0.49488973 | 36.9172577 | -79.69090052 |
| Northumberland County, District 2 | 0.41715976 | 0.49488973 | 37.92815734 | -76.42425002 |
| Alleghany County, Covington district | 0.30930608 | 0.49488973 | 37.79121185 | -79.96163935 |
| Botetourt County, Fincastle district | 0.36390533 | 0.49354492 | 37.46083132 | -79.85996877 |
| Northumberland County, District 3 | 0.42388381 | 0.49139322 | 37.88448924 | -76.43598904 |
| Rockingham County, Linville district | 0.39214632 | 0.4908553 | 38.67552991 | -78.97029332 |
| Nottoway County, District 2 | 0.32517483 | 0.49004841 | 37.1060014 | -78.00378425 |
| Accomack County, District 7 | 0.25309306 | 0.49004841 | 37.69836208 | -75.72500823 |
| King and Queen County, Shanghai district | 0.42052179 | 0.48708983 | 37.5493396 | -76.75897425 |
| Cumberland County, District 1 | 0.42657343 | 0.48682087 | 37.6215638 | -78.11003655 |
| Buckingham County, District 1 | 0.40102205 | 0.48628295 | 37.6944677 | -78.29614645 |
| Prince Edward County, District 401 | 0.35691232 | 0.48601399 | 37.23816225 | -78.43475139 |
| Caroline County, Bowling Green district | 0.41204949 | 0.4852071 | 38.03586776 | -77.24643017 |
| Essex County, North district | 0.43195266 | 0.4852071 | 38.14951862 | -77.09958594 |
| Craig County, Potts Mountain district | 0.38972566 | 0.48466918 | 37.39924477 | -80.42277302 |
| Highland County, Stonewall district | 0.43006993 | 0.48466918 | 38.25883531 | -79.56587876 |
| Pittsylvania County, Staunton River district | 0.37789134 | 0.48359333 | 37.09076339 | -79.30671679 |
| Smyth County, Royal Oak district | 0.43854223 | 0.48305541 | 36.83571871 | -81.51507235 |
| Northampton County, District 6 | 0.27003765 | 0.48278644 | 37.51154846 | -75.83522857 |
| Patrick County, Peters Creek district | 0.43948359 | 0.48224852 | 36.64732177 | -80.20815647 |
| Patrick County, Mayo River district | 0.4168908 | 0.48197956 | 36.66932974 | -80.20348658 |
| Carroll County, Pine Creek district | 0.47821409 | 0.48197956 | 36.85611742 | -80.59408216 |
| Patrick County, Blue Ridge district | 0.44486283 | 0.48197956 | 36.68037734 | -80.28702902 |
| York County, District 2 | 0.43679398 | 0.48117267 | 37.136521 | -76.46077194 |
| Franklin County, Boone district | 0.43598709 | 0.48036579 | 37.07948656 | -79.88557779 |
| Bath County, Millboro district | 0.37089833 | 0.48009683 | 37.88528066 | -79.64982329 |
| Pulaski County, Ingles district | 0.47848306 | 0.47848306 | 36.94779885 | -80.60375325 |
| Sussex County, Waverly district | 0.38004303 | 0.47794513 | 37.02510574 | -77.04051633 |
| Alleghany County, Boiling Springs district | 0.29653039 | 0.47740721 | 37.75097458 | -79.99950026 |
| Amelia County, District 3 | 0.38528779 | 0.47552448 | 37.33350864 | -78.00611621 |
| King and Queen County, St. Stephens Church district | 0.41151157 | 0.47525551 | 37.81692921 | -77.01025693 |
| Richmond County, District 4 | 0.37520172 | 0.47498655 | 37.94260361 | -76.65502282 |
| Accomack County, District 6 | 0.36847768 | 0.47444863 | 37.71543491 | -75.69096387 |
| Highland County, Monterey district | 0.38838085 | 0.47283486 | 38.32858727 | -79.57941541 |
| Floyd County, Burks Fork district | 0.4197149 | 0.47283486 | 36.72794681 | -80.47746329 |
| Carroll County, Laurel Fork district | 0.44862829 | 0.47014524 | 36.70592899 | -80.4820565 |
| Patrick County, Smith River district | 0.43464228 | 0.46853147 | 36.62900028 | -80.24065252 |
| Buckingham County, District 3 | 0.34198494 | 0.46745562 | 37.33816217 | -78.49781334 |
| Sussex County, Wakefield district | 0.37224314 | 0.46718666 | 37.00323663 | -77.04613652 |
| Goochland County, District 1 | 0.42240452 | 0.46718666 | 37.6873001 | -78.09396848 |
| Pittsylvania County, Callands-Gretna district | 0.40236686 | 0.46637977 | 36.84647152 | -79.50538116 |
| Smyth County, Park district | 0.36350188 | 0.46584185 | 36.83910331 | -81.54304076 |
| Appomattox County, Piney Mountain district | 0.32436794 | 0.46369016 | 37.38672072 | -78.83408686 |
| Rockbridge County, Walkers Creek district | 0.39913932 | 0.46288327 | 37.92157736 | -79.58341779 |
| Halifax County, District 1 | 0.34117805 | 0.4612695 | 36.79006916 | -78.94343268 |
| Dinwiddie County, District 5 | 0.31737493 | 0.45535234 | 37.13415116 | -77.5917862 |
| Halifax County, District 2 | 0.34010221 | 0.45508338 | 36.82383305 | -78.7055912 |
| Grayson County, Wilson Creek district | 0.3844809 | 0.44674556 | 36.62169405 | -81.24258001 |
| Carroll County, Sulphur Springs district | 0.44620764 | 0.44620764 | 36.77368593 | -81.01207769 |
| Accomack County, District 8 | 0.18719742 | 0.44566971 | 37.67656131 | -75.73106721 |
| Botetourt County, Amsterdam district | 0.4316837 | 0.44324906 | 37.46134341 | -80.03352896 |
| King and Queen County, Stevensville district | 0.404922 | 0.4429801 | 37.78175922 | -76.93168484 |
| Campbell County, Brookneal district | 0.39214632 | 0.4408284 | 37.03953107 | -78.94355113 |
| Charlotte County, Roanoke district | 0.33122647 | 0.4373319 | 37.03155228 | -78.88772389 |
| Mecklenburg County, District 5 | 0.41729424 | 0.43625605 | 36.73761175 | -78.12196479 |
| Accomack County, District 1 | 0.3746638 | 0.43598709 | 37.917024 | -75.39953567 |
| Smyth County, Chilhowie district | 0.3119957 | 0.4351802 | 36.80845698 | -81.6634463 |
| Mecklenburg County, District 3 | 0.19553523 | 0.43464228 | 36.71151549 | -78.09949772 |
| Nottoway County, District 5 | 0.33109198 | 0.43464228 | 37.1194983 | -77.94264777 |
| Northumberland County, District 1 | 0.39591178 | 0.4338354 | 38.01186983 | -76.51916594 |
| Bath County, Williamsville district | 0.32571275 | 0.43329747 | 38.17477683 | -79.61408851 |
| Northampton County, District 3 | 0.13797741 | 0.43329747 | 37.52865291 | -75.80129526 |
| Buckingham County, District 6 | 0.35032275 | 0.4316837 | 37.74924143 | -78.5160317 |
| Washington County, Wilson district | 0.32167832 | 0.43087682 | 36.63726478 | -82.12435887 |
| Cumberland County, District 4 | 0.30486821 | 0.43033889 | 37.36150465 | -78.34731047 |
| Albemarle County, Scottsville district | 0.42872512 | 0.42899408 | 37.79309994 | -78.50554499 |
| Middlesex County, Saluda district | 0.32423346 | 0.42845616 | 37.59236407 | -76.60173097 |
| Northumberland County, District 4 | 0.3816568 | 0.4260355 | 37.87078158 | -76.35173192 |
| Appomattox County, Wreck Island district | 0.42173211 | 0.42522862 | 37.39072516 | -78.8621294 |
| Buckingham County, District 5 | 0.29827864 | 0.42442173 | 37.73138793 | -78.5494249 |
| Smyth County, North Fork district | 0.27918236 | 0.42307692 | 36.84248141 | -81.57101169 |
| Middlesex County, Jamaica district | 0.36027434 | 0.42307692 | 37.72863173 | -76.74095867 |
| Cumberland County, District 2 | 0.35180204 | 0.42280796 | 37.5777378 | -78.12073278 |
| Sussex County, Stony Creek district | 0.22027972 | 0.42092523 | 36.99016629 | -77.39648872 |
| Cumberland County, District 3 | 0.3816568 | 0.42092523 | 37.48477867 | -78.25957923 |
| Charlotte County, Midway district | 0.36538462 | 0.41662184 | 37.05350269 | -78.8826416 |
| Bath County, Valley Springs district | 0.36403981 | 0.41662184 | 37.97402205 | -79.80491183 |
| Highland County, Blue Grass district | 0.36229155 | 0.41500807 | 38.53783315 | -79.62018012 |
| Lancaster County, District 4 | 0.36094675 | 0.41366326 | 37.71795929 | -76.39287577 |
| Rockbridge County, Buffalo district | 0.29343733 | 0.41204949 | 37.6463857 | -79.55788255 |
| Rockingham County, Ashby district | 0.41151157 | 0.41151157 | 38.48363291 | -79.22113896 |
| Mecklenburg County, District 9 | 0.3041958 | 0.41097364 | 36.63208946 | -78.63543694 |
| Washington County, Monroe district | 0.27366864 | 0.41070468 | 36.72143268 | -81.88019466 |
| King and Queen County, Newtown district | 0.38246369 | 0.41070468 | 37.9132263 | -77.04373691 |
| Wise County, District 2 | 0.13986014 | 0.41016676 | 36.9818489 | -82.57561788 |
| Brunswick County, Sturgeon district | 0.20871436 | 0.40559441 | 36.8397091 | -77.89563077 |
| Surry County, Dendron district | 0.28187197 | 0.4042496 | 36.98994634 | -76.96276208 |
| Brunswick County, Red Oak district | 0.19876278 | 0.40398063 | 36.86161853 | -77.89027107 |
| Accomack County, District 9 | 0.25201721 | 0.40155998 | 37.55516182 | -75.82319522 |
| Norton city, Norton city | 0.29101668 | 0.40075309 | 36.94080523 | -82.61164914 |
| Bath County, Cedar Creek district | 0.31091985 | 0.40021517 | 37.99978446 | -79.82836286 |
| Washington County, Madison district | 0.27541689 | 0.39940828 | 36.67164562 | -82.03234461 |
| Lancaster County, District 2 | 0.36874664 | 0.39806347 | 37.74435991 | -76.41504318 |
| Rockbridge County, Kerrs Creek district | 0.32907477 | 0.39618074 | 37.81553249 | -79.63630139 |
| Pulaski County, Massie district | 0.36659494 | 0.3951049 | 37.15259609 | -80.79143189 |
| Washington County, Tyler district | 0.11780527 | 0.39456697 | 36.60603507 | -82.24420144 |
| Brunswick County, Powellton district | 0.16003228 | 0.39187735 | 36.76128434 | -77.82842121 |
| Surry County, Claremont district | 0.34480904 | 0.39187735 | 37.15147774 | -76.83398886 |
| Washington County, Jefferson district | 0.15384615 | 0.39160839 | 36.74113543 | -82.0478838 |
| Buckingham County, District 2 | 0.33539537 | 0.39133943 | 37.5355292 | -78.45073845 |
| Greensville County, District 3 | 0.20979021 | 0.38999462 | 36.73614208 | -77.51788138 |
| Scott County, District 1 | 0.24623453 | 0.38838085 | 36.61895753 | -82.55537551 |
| York County, District 3 | 0.28846154 | 0.38838085 | 37.18475521 | -76.47699331 |
| Surry County, Carsley district | 0.33136095 | 0.38757396 | 37.07326358 | -77.05708756 |
| Brunswick County, Totaro district | 0.30970952 | 0.38703604 | 36.74361298 | -77.86153749 |
| Nottoway County, District 3 | 0.28536848 | 0.38676708 | 37.2197575 | -78.00494845 |
| Lee County, District 1 | 0.11215707 | 0.38488435 | 36.60106906 | -83.64041496 |
| Charlotte County, Bacon district | 0.27124798 | 0.3816568 | 36.83768612 | -78.64481286 |
| Lancaster County, District 1 | 0.27972028 | 0.37789134 | 37.79347249 | -76.57750856 |
| Gloucester County, Petsworth district | 0.26976869 | 0.37762238 | 37.57052006 | -76.60752339 |
| Charlotte County, Madison district | 0.30016138 | 0.37547068 | 37.08714157 | -78.64393398 |
| Lee County, District 5 | 0.1542496 | 0.37278107 | 36.75993779 | -83.01516621 |
| Buckingham County, District 4 | 0.31791286 | 0.37197418 | 37.54778799 | -78.53498135 |
| York County, District 4 | 0.33041958 | 0.37170522 | 37.15836122 | -76.45497025 |
| Wise County, District 1 | 0.15384615 | 0.3690156 | 36.98498694 | -82.60368713 |
| Washington County, Taylor district | 0.20656267 | 0.36793975 | 36.70266274 | -81.91230968 |
| Alleghany County, Falling Spring district | 0.27434104 | 0.36740183 | 37.86092394 | -79.97546381 |
| Russell County, Lebanon district | 0.14779451 | 0.3655191 | 36.81712883 | -82.11942085 |
| Franklin city, Franklin city | 0.2802582 | 0.36309844 | 36.66603983 | -76.93074155 |
| Nottoway County, District 1 | 0.20037655 | 0.36229155 | 37.08828467 | -78.03701183 |
| Appomattox County, Falling River district | 0.29532006 | 0.36094675 | 37.26897238 | -78.8036574 |
| Bath County, Warm Springs district | 0.28644432 | 0.35556751 | 38.18696786 | -79.8745209 |
| Charlotte County, Central district | 0.29599247 | 0.35529855 | 36.95144686 | -78.64694752 |
| Scott County, District 2 | 0.10570199 | 0.35476062 | 36.59686435 | -82.5593525 |
| Nelson County, West district | 0.35260893 | 0.35260893 | 37.84359016 | -79.0183323 |
| Mecklenburg County, District 6 | 0.21490048 | 0.35233997 | 36.60607813 | -78.15361779 |
| Amelia County, District 1 | 0.29115116 | 0.35207101 | 37.37216681 | -78.11271178 |
| Wise County, District 4 | 0.13179129 | 0.34803658 | 36.95975912 | -82.57960818 |
| Lee County, District 3 | 0.24878967 | 0.34615385 | 36.74086663 | -83.04701146 |
| Sussex County, Newville district | 0.22175901 | 0.34561592 | 37.04276152 | -77.29643689 |
| Buckingham County, District 7 | 0.29128564 | 0.34534696 | 37.6245739 | -78.28393439 |
| Southampton County, Berlin and Ivor district | 0.18706294 | 0.34185046 | 36.95919081 | -76.91285145 |
| Emporia city, Emporia city | 0.30231307 | 0.33996772 | 36.6966602 | -77.55650797 |
| Rockbridge County, Natural Bridge district | 0.31172673 | 0.33969876 | 37.62056971 | -79.53463742 |
| Buchanan County, South Grundy district | 0.23493814 | 0.33566434 | 37.23976889 | -82.06889797 |
| Poquoson city, Poquoson city | 0.23117267 | 0.33405056 | 37.12739849 | -76.40515702 |
| Bland County, District 3 | 0.29908553 | 0.33351264 | 37.14473537 | -80.90815665 |
| Lancaster County, District 5 | 0.3006993 | 0.33136095 | 37.70524965 | -76.45478084 |
| Surry County, Surry district | 0.28160301 | 0.33055406 | 37.10776073 | -76.84536593 |
| Mecklenburg County, District 8 | 0.22081764 | 0.32920925 | 36.66784985 | -78.56965317 |
| Lunenburg County, Hounds Creek district | 0.16433566 | 0.32732652 | 36.96515313 | -78.12464471 |
| Mecklenburg County, District 1 | 0.24636902 | 0.32705756 | 36.59795475 | -78.55735359 |
| Henry County, Horsepasture district | 0.32665412 | 0.3267886 | 36.54784144 | -79.97195542 |
| Gloucester County, Abingdon district | 0.24515869 | 0.32544379 | 37.39575309 | -76.65374023 |
| Middlesex County, Pine Top district | 0.29209252 | 0.32033351 | 37.51688401 | -76.41757366 |
| Lunenburg County, Meherrin River district | 0.14389457 | 0.31979559 | 37.07234106 | -78.38756374 |
| Mathews County, Piankatank district | 0.28294782 | 0.31979559 | 37.49504834 | -76.42341435 |
| Southampton County, Franklin district | 0.20199032 | 0.31791286 | 36.67935449 | -77.01375074 |
| Lancaster County, District 3 | 0.28765465 | 0.31387843 | 37.68697654 | -76.34271218 |
| Isle of Wight County, Carrsville district | 0.1401291 | 0.31226466 | 36.66158885 | -76.90307837 |
| Brunswick County, Meherrin district | 0.19042496 | 0.3119957 | 36.72170352 | -77.86689716 |
| Smyth County, Saltville district | 0.20441097 | 0.30957504 | 36.86261353 | -81.73889005 |
| Lee County, District 2 | 0.14416353 | 0.30930608 | 36.70269848 | -83.11065342 |
| Greensville County, District 2 | 0.11511565 | 0.30365788 | 36.67476389 | -77.56195323 |
| Mecklenburg County, District 2 | 0.21301775 | 0.302851 | 36.66524817 | -78.39795942 |
| Isle of Wight County, Smithfield district | 0.23063475 | 0.30150619 | 36.97990912 | -76.61808527 |
| Isle of Wight County, Hardy district | 0.16352878 | 0.29935449 | 36.98442492 | -76.64584911 |
| Scott County, District 6 | 0.08808499 | 0.29209252 | 36.71331435 | -82.79512769 |
| Gloucester County, Ware district | 0.24556213 | 0.29101668 | 37.45593274 | -76.46301426 |
| Southampton County, Jerusalem district | 0.11202259 | 0.29074771 | 36.70565225 | -77.03582936 |
| Isle of Wight County, Newport district | 0.19674556 | 0.2894029 | 36.94447751 | -76.54057737 |
| Mathews County, Westville district | 0.25228618 | 0.28778913 | 37.45137571 | -76.43508543 |
| Gloucester County, York district | 0.20656267 | 0.28671329 | 37.35128116 | -76.51998588 |
| Sussex County, Courthouse district | 0.2105971 | 0.28187197 | 36.89837487 | -77.24634566 |
| Wise County, District 3 | 0.15599785 | 0.27891339 | 36.86800308 | -82.76768301 |
| Mathews County, Chesapeake district | 0.20790748 | 0.27729962 | 37.38941951 | -76.33512452 |
| Greensville County, District 4 | 0.15330823 | 0.26842388 | 36.81939304 | -77.46825745 |
| Gloucester County, Gloucester Point district | 0.2563206 | 0.26815492 | 37.25482728 | -76.48743598 |
| Buchanan County, Garden district | 0.1344809 | 0.2654653 | 37.2520306 | -81.98029759 |
| Russell County, Castlewood district | 0.10247445 | 0.26358257 | 36.93075301 | -82.12683353 |
| Isle of Wight County, Windsor district | 0.14981173 | 0.26331361 | 36.90531546 | -76.57982478 |
| Buchanan County, Hurricane district | 0.13071544 | 0.25443787 | 37.21116997 | -82.01678646 |
| Lee County, District 4 | 0.12937063 | 0.25147929 | 36.77595027 | -82.95529692 |
| Dickenson County, Willis district | 0.09951587 | 0.2493276 | 37.14907978 | -82.45922151 |
| Sussex County, Henry district | 0.19096288 | 0.24663798 | 36.79769461 | -77.32953324 |
| Southampton County, Capron district | 0.12614309 | 0.24556213 | 36.68819903 | -77.06910652 |
| Russell County, New Garden district | 0.11202259 | 0.24475524 | 37.02479072 | -81.96587654 |
| Northampton County, District 4 | 0.18047337 | 0.24018289 | 37.34643871 | -75.93892783 |
| Greensville County, District 1 | 0.11565358 | 0.23991393 | 36.6047699 | -77.55059005 |
| Lunenburg County, Plymouth district | 0.18316299 | 0.23023131 | 37.00371303 | -78.23077623 |
| Lunenburg County, Browns Store district | 0.13972566 | 0.22485207 | 36.94323443 | -78.12994319 |
| Northampton County, District 1 | 0.16594944 | 0.22377622 | 37.26384727 | -75.99060506 |
| Surry County, Bacons Castle district | 0.14604626 | 0.2197418 | 37.12067114 | -76.78402335 |
| Buchanan County, North Grundy district | 0.16998386 | 0.21839699 | 37.28064477 | -82.03242368 |
| Mecklenburg County, District 7 | 0.16823561 | 0.21436256 | 36.7186788 | -78.29904747 |
| Scott County, District 4 | 0.12883271 | 0.21382464 | 36.64419453 | -82.57933711 |
| Buchanan County, Rock Lick district | 0.13649812 | 0.20925229 | 37.31897163 | -82.16911208 |
| Southampton County, Drewryville district | 0.13743948 | 0.2084454 | 36.78459878 | -77.24633735 |
| Buchanan County, Prater district | 0.13206025 | 0.20414201 | 37.22420097 | -82.12932767 |
| Dickenson County, Clintwood district | 0.13824637 | 0.19983862 | 37.15540295 | -82.5154784 |
| Scott County, District 5 | 0.09306079 | 0.19419042 | 36.81462054 | -82.49147403 |
| Buchanan County, Knox district | 0.1218397 | 0.190156 | 37.33700984 | -81.93537281 |
| Northampton County, District 2 | 0.18961807 | 0.18961807 | 37.24203047 | -75.99655098 |
| Southampton County, Boykins and Branchville district | 0.09803658 | 0.18612157 | 36.55702637 | -77.2463208 |
| Dickenson County, Kenady district | 0.07154384 | 0.18423884 | 37.1301594 | -82.49137429 |
| Lunenburg County, Loves Mill district | 0.14201183 | 0.17993545 | 36.95986665 | -78.24131336 |
| Southampton County, Newsoms district | 0.0661646 | 0.16810113 | 36.63118142 | -76.99729415 |
| Dickenson County, Ervinton district | 0.06535772 | 0.16783217 | 37.18051532 | -82.33859666 |
| Lunenburg County, Beaver Creek district | 0.10919849 | 0.16648736 | 36.79919209 | -78.0783151 |
| Dickenson County, Sand Lick district | 0.10664336 | 0.14792899 | 37.21505106 | -82.24601089 |
| Scott County, District 3 | 0.07853685 | 0.13851533 | 36.6946737 | -82.42760729 |
