## Supplementary material for "Mapping long-distance jump dispersal risk by leveraging transport infrastructure data and sampling bias correction: a case study with the invasive spotted lanternfly (*Lycorma delicatula*)": S17 West Virginia risk

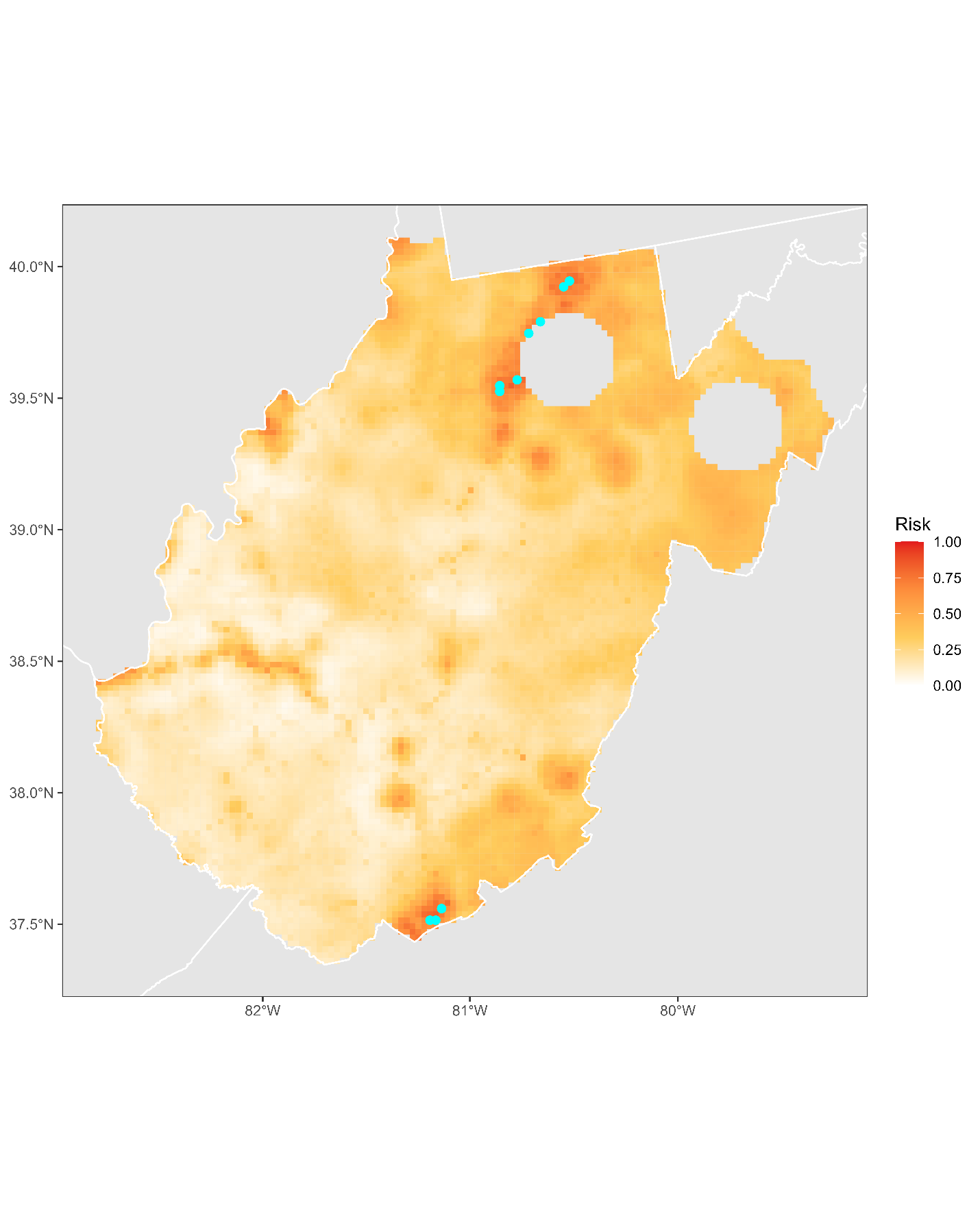

**Figure S17.1:** Projected risk of colonization by *L. delicatula* jump dispersal for West Virginia. Risk projections are based on an ensemble random forest model including distances from existing *L. delicatula* populations. Note that areas falling less than 20 km from existing *L. delicatula* populations are not included in the map because any colonization there would be categorized as diffusive spread based on our methodology (regardless of the actual source of the resulting population). Darker, redder colors indicate increased risk of colonization by jump dispersal. Cyan points show the top ten highest risk locations in the state.

**Table S17.1:** Ranked list of West Virginia county subdivisions sorted by the maximum projected risk of colonization by jump dispersal occurring there. For each subdivision, we identified the 2.5 x 2.5 km grid cell centroid included there with the highest projected risk of colonization by jump dispersal and report that location and risk value.

| **County, subdivision** | **Median risk value** | **Maximum risk value** | **Latitude of maximum risk** | **Longitude of maximum risk** |
| --- | --- | --- | --- | --- |
| Mercer County, District II | 0.70629371 | 0.85233997 | 37.34555433 | -81.06902859 |
| Mercer County, District I | 0.66218397 | 0.84965035 | 37.3049713 | -81.10606173 |
| Monongalia County, Western district | 0.45535234 | 0.84023669 | 39.62472212 | -79.96739561 |
| Mercer County, District III | 0.35691232 | 0.83754707 | 37.30845008 | -81.13418757 |
| Harrison County, Eastern district | 0.7049489 | 0.83512641 | 39.3024504 | -80.2767443 |
| Monongalia County, Eastern district | 0.64604626 | 0.83001614 | 39.64296375 | -79.93348039 |
| Harrison County, North Urban district | 0.61565358 | 0.79639591 | 39.29143645 | -80.36812614 |
| Marion County, Middletown district | 0.59077461 | 0.77461001 | 39.47108402 | -80.18025156 |
| Harrison County, South Urban district | 0.67348037 | 0.77245831 | 39.26943114 | -80.37291498 |
| Marion County, Palatine district | 0.6007262 | 0.74098978 | 39.5076716 | -80.1126558 |
| Monongalia County, Central district | 0.72377622 | 0.73587951 | 39.62097301 | -79.93842511 |
| Wood County, Parkersburg district | 0.55567509 | 0.72942442 | 39.29989317 | -81.55140516 |
| Lewis County, Hackers Creek-Skin Creek district | 0.34938139 | 0.69983862 | 39.11539192 | -80.4063556 |
| Harrison County, Southern district | 0.46664874 | 0.69768693 | 39.13374968 | -80.37278882 |
| Harrison County, Northern district | 0.45535234 | 0.69688004 | 39.35379268 | -80.32485882 |
| Wood County, Tygart district | 0.37412587 | 0.6810113 | 39.25241527 | -81.53136053 |
| Harrison County, Southwest district | 0.38999462 | 0.67724583 | 39.22542031 | -80.38248393 |
| Wood County, Lubeck district | 0.27810651 | 0.67401829 | 39.25579489 | -81.560262 |
| Upshur County, First district | 0.51048951 | 0.67025282 | 39.00173051 | -80.22417063 |
| Wayne County, Ceredo district | 0.37869822 | 0.65761162 | 38.40145029 | -82.57678965 |
| Marshall County, District 2 | 0.60408822 | 0.65196342 | 39.90316591 | -80.7421335 |
| Greenbrier County, Eastern district | 0.24098978 | 0.65115654 | 37.77335452 | -80.34328553 |
| Wood County, Williams district | 0.43410436 | 0.64389457 | 39.39995728 | -81.44233895 |
| Marshall County, District 3 | 0.30217859 | 0.63905325 | 39.90672929 | -80.77125863 |
| Upshur County, Second district | 0.31226466 | 0.62587413 | 38.97604411 | -80.20025679 |
| Cabell County, District 3 | 0.10570199 | 0.6218397 | 38.42664338 | -82.39692312 |
| Marion County, West Augusta district | 0.37278107 | 0.61861216 | 39.55166502 | -80.10288501 |
| Raleigh County, District 2 | 0.12883271 | 0.612695 | 37.78854924 | -81.18197328 |
| Kanawha County, District 1 | 0.12506724 | 0.60247445 | 38.33495241 | -81.5967393 |
| Preston County, First district | 0.46032813 | 0.59978483 | 39.66808148 | -79.7786461 |
| Kanawha County, District 3 | 0.16783217 | 0.59440559 | 38.38373695 | -81.82086489 |
| Kanawha County, District 2 | 0.36175363 | 0.58768155 | 38.35170715 | -81.73948174 |
| Upshur County, Third district | 0.27555137 | 0.58741259 | 38.95773345 | -80.23379431 |
| Randolph County, Leadsville district | 0.44271114 | 0.58203335 | 38.9312708 | -79.85575986 |
| Cabell County, District 1 | 0.21167294 | 0.57315761 | 38.41091551 | -82.45826069 |
| Fayette County, Plateau district | 0.17146315 | 0.57154384 | 37.98685588 | -81.14180965 |
| Putnam County, District 1 | 0.10462614 | 0.55621302 | 38.43116356 | -81.84085939 |
| Wetzel County, District 2 | 0.31925767 | 0.55513717 | 39.66799681 | -80.85190307 |
| Wetzel County, District 3 | 0.55433029 | 0.55433029 | 39.64597064 | -80.85656922 |
| Nicholas County, Summersville district | 0.29693384 | 0.54410974 | 38.26718305 | -80.85096978 |
| Raleigh County, District 1 | 0.12009145 | 0.53738569 | 37.76997354 | -81.21472767 |
| Hardy County, Old Fields district | 0.36040882 | 0.5274341 | 39.08251227 | -78.96281538 |
| Preston County, Third district | 0.49058634 | 0.52339968 | 39.48083212 | -79.73179046 |
| Monroe County, Eastern district | 0.3577192 | 0.52286175 | 37.67751793 | -80.6537378 |
| Summers County, New River district | 0.19311458 | 0.52286175 | 37.72513504 | -80.67279232 |
| Raleigh County, District 3 | 0.1788596 | 0.51398601 | 37.76651388 | -81.18642262 |
| Greenbrier County, Western district | 0.20131791 | 0.5126412 | 37.86844719 | -80.55558656 |
| Preston County, Fourth district | 0.3725121 | 0.51075847 | 39.45123943 | -79.67900449 |
| Randolph County, Beverly district | 0.35852609 | 0.50968263 | 38.88730307 | -79.86559662 |
| Pendleton County, Central district | 0.44419042 | 0.50806885 | 38.65687629 | -79.32822991 |
| Wayne County, Westmoreland district | 0.31522324 | 0.50618612 | 38.3951571 | -82.51957192 |
| Preston County, Second district | 0.39618074 | 0.48924153 | 39.64905032 | -79.63378645 |
| Greenbrier County, Central district | 0.24825175 | 0.48466918 | 37.80266388 | -80.39513822 |
| Monroe County, Central district | 0.38972566 | 0.48332437 | 37.67392818 | -80.6254948 |
| Lewis County, Courthouse-Collins Settlement district | 0.20306616 | 0.47902098 | 38.91340746 | -80.59745129 |
| Barbour County, North district | 0.37547068 | 0.4760624 | 39.15542595 | -80.01278365 |
| Putnam County, District 2 | 0.28429263 | 0.47579344 | 38.44106624 | -81.9266426 |
| Hardy County, South Fork district | 0.37600861 | 0.47417967 | 39.04258797 | -79.0018994 |
| Barbour County, West district | 0.34211942 | 0.47229693 | 39.15915738 | -80.0415722 |
| Pendleton County, Western district | 0.40667025 | 0.47014524 | 38.71245551 | -79.40382948 |
| Kanawha County, District 4 | 0.12130178 | 0.46987628 | 38.36372465 | -81.64948464 |
| Pendleton County, Eastern district | 0.39725659 | 0.46960732 | 38.74861667 | -79.3365495 |
| Lewis County, Freemans Creek district | 0.31334051 | 0.46906939 | 39.05664107 | -80.47818119 |
| Hardy County, Moorefield district | 0.39268424 | 0.46745562 | 39.06056434 | -78.96801532 |
| Preston County, Fifth district | 0.36215707 | 0.46584185 | 39.41488738 | -79.74676624 |
| Barbour County, South district | 0.37062937 | 0.45938677 | 39.0342538 | -79.95097057 |
| Monroe County, Western district | 0.37439484 | 0.45804196 | 37.40197098 | -80.79774903 |
| Doddridge County, Beech district | 0.35180204 | 0.4556213 | 39.2984293 | -80.60381314 |
| Wood County, Union district | 0.24179666 | 0.45535234 | 39.33041582 | -81.42681042 |
| Hardy County, Lost River district | 0.36672942 | 0.4499731 | 38.8391158 | -78.84301692 |
| Tucker County, Fairfax district | 0.43813878 | 0.44943518 | 39.13876239 | -79.54251859 |
| Wetzel County, District 1 | 0.30849919 | 0.44809037 | 39.59837729 | -80.83688619 |
| Tucker County, St. George district | 0.40344271 | 0.44728349 | 39.1684104 | -79.59502165 |
| Pocahontas County, Greenbank district | 0.3148198 | 0.44701452 | 38.55308163 | -79.73429974 |
| Tucker County, Black Fork district | 0.41151157 | 0.4464766 | 39.09196291 | -79.70128663 |
| Wood County, Clay district | 0.27931684 | 0.43975256 | 39.22698185 | -81.50690329 |
| Randolph County, Roaring Creek district | 0.2915546 | 0.43760086 | 38.91681332 | -79.91804957 |
| Tucker County, Davis district | 0.38259817 | 0.43760086 | 39.13106206 | -79.48501284 |
| Tucker County, Clover district | 0.38488435 | 0.42926304 | 39.12154359 | -79.75382058 |
| Cabell County, District 2 | 0.12479828 | 0.42872512 | 38.41065165 | -82.2538873 |
| Jackson County, Western district | 0.16594944 | 0.42711135 | 38.93844031 | -81.74146603 |
| Mingo County, Williamson district | 0.20871436 | 0.42415277 | 37.6694109 | -82.27673355 |
| Tucker County, Dry Fork district | 0.33620226 | 0.42388381 | 39.09097377 | -79.52383542 |
| Wood County, Slate district | 0.1514255 | 0.42307692 | 39.18288749 | -81.51577827 |
| Tyler County, North district | 0.3422539 | 0.422539 | 39.57989212 | -80.87055057 |
| Mason County, Lewis district | 0.14819796 | 0.41984938 | 38.84881236 | -82.14129193 |
| Grant County, Grant district | 0.35731576 | 0.41204949 | 39.14227418 | -79.06730818 |
| Putnam County, District 3 | 0.1013986 | 0.40263583 | 38.44435386 | -81.95524214 |
| Grant County, Union district | 0.29128564 | 0.40021517 | 39.19697037 | -79.46985782 |
| Mason County, Clendenin district | 0.09736417 | 0.39967725 | 38.82674448 | -82.14551586 |
| Randolph County, New Interest district | 0.33122647 | 0.39725659 | 39.02297698 | -79.86477616 |
| Summers County, Greenbrier River district | 0.34817106 | 0.39349112 | 37.61436657 | -80.86966875 |
| Doddridge County, Oak district | 0.31038193 | 0.38838085 | 39.25440056 | -80.61324573 |
| Nicholas County, Hamilton district | 0.16783217 | 0.38649812 | 38.35526758 | -80.83263229 |
| Mason County, Robinson district | 0.14147391 | 0.37923615 | 38.89294785 | -82.13283639 |
| Summers County, Bluestone River district | 0.302851 | 0.3781603 | 37.59587397 | -80.90242427 |
| Randolph County, Dry Fork district | 0.29424422 | 0.3781603 | 38.89979525 | -79.44934785 |
| Tyler County, South district | 0.24448628 | 0.37600861 | 39.44487968 | -80.69116993 |
| Wayne County, Butler district | 0.16984938 | 0.37278107 | 38.2247889 | -82.60925842 |
| Tucker County, Licking district | 0.35314685 | 0.37170522 | 39.22382997 | -79.67135851 |
| Doddridge County, Maple district | 0.32651963 | 0.36847768 | 39.38239528 | -80.73417573 |
| Doddridge County, Pine district | 0.28698225 | 0.36471221 | 39.27585515 | -80.78643743 |
| Logan County, Central district | 0.17926304 | 0.36282948 | 37.84211047 | -82.01253314 |
| Ritchie County, Clay district | 0.27797203 | 0.35960194 | 39.26352371 | -81.05558074 |
| Pocahontas County, Huntersville district | 0.29585799 | 0.35718128 | 38.19073112 | -79.90292338 |
| Pleasants County, District C | 0.26143088 | 0.35260893 | 39.39454977 | -81.20624684 |
| Randolph County, Mingo district | 0.29424422 | 0.35018827 | 38.57620137 | -80.0817728 |
| Randolph County, Huttonsville district | 0.28738569 | 0.3485745 | 38.59819329 | -80.07694179 |
| Webster County, Northern district | 0.23292093 | 0.34480904 | 38.68271717 | -80.38191134 |
| Pocahontas County, Little Levels district | 0.25739645 | 0.34104357 | 38.18400894 | -80.1966518 |
| Pocahontas County, Edray district | 0.24690694 | 0.33566434 | 38.17661178 | -80.13982507 |
| Tyler County, Central district | 0.24825175 | 0.32974718 | 39.51027697 | -80.85553404 |
| Braxton County, Northern district | 0.14254976 | 0.32947821 | 38.87297607 | -80.63554327 |
| Mineral County, District 1 | 0.29464766 | 0.32759548 | 39.38546713 | -79.1883624 |
| Braxton County, Eastern district | 0.17348037 | 0.32329209 | 38.6968618 | -80.67290918 |
| Tyler County, West district | 0.28859602 | 0.32275417 | 39.568442 | -80.96219095 |
| Hardy County, Capon district | 0.31603012 | 0.32221625 | 39.08849247 | -78.84279408 |
| Roane County, District III | 0.25618612 | 0.31952663 | 38.61291141 | -81.27762317 |
| Randolph County, Valley Bend district | 0.25874126 | 0.31710597 | 38.82511235 | -79.9089794 |
| Gilmer County, City district | 0.21382464 | 0.31468531 | 38.94198853 | -80.82730853 |
| Nicholas County, Wilderness district | 0.14335664 | 0.31387843 | 38.20111752 | -80.86469407 |
| Braxton County, Western district | 0.14806347 | 0.31253362 | 38.65997763 | -80.73947338 |
| Marshall County, District 1 | 0.29747176 | 0.31172673 | 39.88164405 | -80.56744364 |
| Gilmer County, Glenville district | 0.21516945 | 0.30957504 | 38.96400964 | -80.82267866 |
| Nicholas County, Kentucky district | 0.19661108 | 0.3063475 | 38.24161873 | -80.82707639 |
| Jackson County, Northern district | 0.12318451 | 0.3041958 | 38.93510295 | -81.71268216 |
| McDowell County, Browns Creek district | 0.19136633 | 0.3041958 | 37.41922662 | -81.487058 |
| Roane County, District II | 0.24502421 | 0.30392684 | 38.69274744 | -81.37886002 |
| Ritchie County, Grant district | 0.20548682 | 0.30311996 | 39.26701368 | -81.08446412 |
| Jackson County, Eastern district | 0.19338354 | 0.30043034 | 38.77404759 | -81.68569726 |
| Ritchie County, Murphy district | 0.21893491 | 0.29989242 | 39.07912362 | -81.21192781 |
| Nicholas County, Beaver district | 0.20575578 | 0.29827864 | 38.23144875 | -80.56630195 |
| Ritchie County, Union district | 0.20037655 | 0.29774072 | 39.21596715 | -81.03587096 |
| Logan County, Eastern district | 0.17563206 | 0.29128564 | 37.97776644 | -82.01567259 |
| Mingo County, Tug Hardee district | 0.14631522 | 0.29101668 | 37.66619763 | -82.24841904 |
| Logan County, Western district | 0.16366326 | 0.28994083 | 37.95570233 | -82.01988423 |
| Randolph County, Middle Fork district | 0.22538999 | 0.28859602 | 38.84754452 | -80.08094046 |
| Clay County, District A | 0.22485207 | 0.28617536 | 38.50401832 | -81.12392646 |
| Braxton County, Southern district | 0.13932222 | 0.28160301 | 38.61594276 | -80.74874195 |
| Wayne County, Stonewall district | 0.14443249 | 0.27918236 | 37.86518345 | -82.4140501 |
| Gilmer County, De Kalb-Troy district | 0.23292093 | 0.27810651 | 38.96755341 | -80.85143193 |
| Roane County, District I | 0.23440022 | 0.27729962 | 38.65548422 | -81.44506223 |
| McDowell County, North Elkin district | 0.21221087 | 0.27649274 | 37.40732626 | -81.57593796 |
| Clay County, District B | 0.16137708 | 0.2732652 | 38.47502134 | -81.07128739 |
| Fayette County, New Haven district | 0.15008069 | 0.27299623 | 37.96672487 | -80.79679435 |
| Wyoming County, District 1 | 0.12990855 | 0.27218935 | 37.48347514 | -81.27222227 |
| Webster County, Central district | 0.21422808 | 0.27084454 | 38.37464831 | -80.44807962 |
| Calhoun County, District 5 | 0.22713825 | 0.2654653 | 38.68026487 | -81.08774663 |
| Gilmer County, Center district | 0.16863905 | 0.2654653 | 38.9235089 | -80.86067194 |
| Wood County, Walker district | 0.14739107 | 0.26223776 | 39.23882979 | -81.41578166 |
| Pleasants County, District B | 0.19795589 | 0.25524476 | 39.37251251 | -81.21078751 |
| Nicholas County, Grant district | 0.17213556 | 0.25416891 | 38.29979926 | -80.93186858 |
| Wirt County, Northeast district | 0.19889726 | 0.25147929 | 39.0350518 | -81.22096356 |
| Pleasants County, District D | 0.22377622 | 0.24986552 | 39.42821205 | -81.11029351 |
| Calhoun County, District 1 | 0.17980097 | 0.24610005 | 38.97014344 | -81.05734496 |
| McDowell County, Big Creek district | 0.16178053 | 0.24206563 | 37.24282894 | -81.52168056 |
| Clay County, District C | 0.12399139 | 0.24179666 | 38.51441994 | -81.20969591 |
| Pleasants County, District A | 0.13999462 | 0.23964497 | 39.3422054 | -81.3355456 |
| Fayette County, Valley district | 0.12345347 | 0.23641743 | 38.1398066 | -81.28557712 |
| Boone County, District 3 | 0.10597095 | 0.23426573 | 38.0495667 | -81.85655298 |
| Boone County, District 1 | 0.10570199 | 0.23184508 | 37.99073956 | -81.54818796 |
| Nicholas County, Jefferson district | 0.16299086 | 0.22861754 | 38.20717402 | -81.09692937 |
| Wyoming County, District 2 | 0.17495966 | 0.22754169 | 37.59515042 | -81.82824732 |
| Calhoun County, District 2 | 0.1964766 | 0.22592792 | 38.9666474 | -81.02858069 |
| Wood County, Harris district | 0.08781603 | 0.22216245 | 39.20308612 | -81.68905685 |
| Boone County, District 2 | 0.15572889 | 0.22135557 | 37.97676903 | -81.81250398 |
| Mingo County, Stafford district | 0.16231845 | 0.22081764 | 37.66464091 | -81.84378878 |
| Lincoln County, District 3 | 0.14066703 | 0.21947283 | 38.27765998 | -82.0748359 |
| Wayne County, Union district | 0.13999462 | 0.21947283 | 38.36359009 | -82.4378731 |
| Mason County, Waggener district | 0.15976331 | 0.21866595 | 38.99351072 | -82.02518144 |
| Wyoming County, District 3 | 0.14846692 | 0.2175901 | 37.62260583 | -81.67835968 |
| Wirt County, Southwest district | 0.15572889 | 0.21732114 | 38.90971791 | -81.30550699 |
| Webster County, Southern district | 0.15814954 | 0.21005917 | 38.33788547 | -80.51446449 |
| McDowell County, Sandy River district | 0.1366326 | 0.20736955 | 37.44282102 | -81.68441831 |
| Mingo County, Lee district | 0.12130178 | 0.19956966 | 37.78893675 | -82.13854206 |
| Wirt County, Central district | 0.15976331 | 0.1993007 | 39.05566047 | -81.3938075 |
| Mason County, Arbuckle district | 0.09185046 | 0.197149 | 38.76633403 | -82.01027658 |
| Calhoun County, District 4 | 0.16702528 | 0.19607316 | 38.73475816 | -81.16469619 |
| Lincoln County, District 2 | 0.14080151 | 0.18665949 | 38.17705661 | -82.18137544 |
| Mingo County, Kermit Harvey district | 0.12506724 | 0.18289403 | 37.86200221 | -82.38565463 |
| Calhoun County, District 3 | 0.1577461 | 0.18262507 | 38.95160192 | -81.09066277 |
| Lincoln County, District 1 | 0.08795051 | 0.17697687 | 38.29219521 | -81.80948708 |
| Mingo County, Beech Ben Mate district | 0.1401291 | 0.17428725 | 37.64352059 | -82.05028662 |
| Mingo County, Magnolia district | 0.11430877 | 0.16729424 | 37.55525155 | -82.06700108 |
| Wood County, Steele district | 0.06643357 | 0.16245293 | 39.12012561 | -81.55791792 |
| Mason County, Graham district | 0.08795051 | 0.16137708 | 38.93626379 | -81.91852357 |
| Mason County, Hannan district | 0.07342657 | 0.15976331 | 38.62489432 | -82.15473493 |
| Mason County, Cooper district | 0.08041958 | 0.15034965 | 38.91420333 | -81.92282006 |
| Mason County, Cologne district | 0.11659494 | 0.14523938 | 38.80060948 | -81.91552352 |
| Mason County, Union district | 0.08068854 | 0.13770845 | 38.74326978 | -81.8092336 |
